# GeneticsMakie.jl: A versatile and scalable toolkit for visualizing locus-level genetic and genomic data

**DOI:** 10.1101/2022.04.18.488573

**Authors:** Minsoo Kim, Daniel D. Vo, Michi E. Kumagai, Connor T. Jops, Michael J. Gandal

## Abstract

With the continued deluge of results from genome-wide association and functional genomic studies, it has become increasingly imperative to quickly combine and visualize different layers of genetic and genomic data within a given locus to facilitate exploratory and integrative data analyses. While several tools have been developed to visualize locus-level genetic results, the limited speed, scalability, and flexibility of current approaches remains a significant bottleneck. Here, we present a Julia package GeneticsMakie.jl for high-performance genetics and genomics-related data visualization that enables fast, simultaneous plotting of hundreds of association results along with multiple relevant genomic annotations. Leveraging the powerful plotting and layout utilities from Makie.jl facilitates the customization and extensibility of every component of a plot, enabling generation of publication-ready figures. The GeneticsMakie.jl package is open source and distributed under the MIT license via GitHub (https://github.com/mmkim1210/GeneticsMakie.jl). The GitHub repository contains installation instructions as well as examples and documentation for built-in functions.

## Introduction

The last decade has seen an exponential increase in the volume of large-scale genetic association results such as those from genome-wide association studies (GWAS) and phenome-wide association studies (PheWAS). The rapid advancements in high-throughput sequencing technologies have further led to an increase in the volume and diversity of molecular genomic readouts, such as 3D genome contacts, ChIP-seq, and ATAC-seq. As these efforts continue to scale, it is becoming increasingly critical to develop efficient ways for simultaneous visualization and integration of multiple such datasets to develop an intuitive understanding of potential underlying biological relationships.

Several tools have been developed to visualize genetic association results within a specific locus along with corresponding genomic annotations, exemplified by the original “LocusZoom” style plots (Pruim et al., 2010). Multiple extensions to these LocusZoom style plots have since been built, spanning a wide array of programming languages, including JavaScript, Python, and R (Jorgenson *et al*., 2009; Pruim *et al*., 2010; Boughton *et al*., 2021; Machiela and Chanock, 2018; Dadaev *et al*., 2016; Machiela and Chanock, 2015; Juliusdottir *et al*., 2018; Kramer *et al*., 2022; Schilder *et al*., 2021; Geihs *et al*., 2015; Kwong *et al*., 2021). However, efficient customization and extension is limited with these tools, and in general they are not well-suited for parallel visualization of large numbers of data points. For example, visualizing GWAS loci across ~100 complex traits for the major histocompatibility complex (MHC) region—the single most pleiotropic region in the human genome that harbors long-range linkage disequilibrium (LD)—requires plotting 66,000 SNPs x 120 traits ≈8 million data points with computation of LD, which existing tools struggle with. Along the same lines, existing tools allow only a certain genomic range to be shown or a certain number of genes to be plotted (see **Table 1** for more complete comparisons), which can misdirect exploratory data analysis (EDA). The Julia programming language (Bezanson *et al*., 2017) is an optimal platform to address this critical gap by providing performance of a low-level language while retaining the readability and ease-of-use of a high-level language.

**Table 1.** Comparison of functionalities provided by different platforms for creating LocusZoom plots. Shown in parentheses are the names of packages that are suited for a particular function or some caveats to note for. We note that plotting functions not provided by the current version of GeneticsMakie.jl can be easily added in the future.

|  | GeneticsMakie.jl | LocusZoom | LocusZoom.js | R ecosystem |
| --- | --- | --- | --- | --- |
| Easily extensible | O | - (designed to run as a command-line tool) | - (designed to run on a web server) | O (will require custom Rcpp code for performance) |
| Actively maintained | O | - | O | O |
| Requires web API access (e.g. pre-computed LD) | - | - | O | - |
| Web browser interface | - | O (legacy service) | O | - |
| Interactivity | - (could be done using Pluto notebooks or GLMakie backend) | - | O | - |
| Automate drawing many figures | O (through scripting) | O (batch mode) | - | O |
| Drawing figures with a custom layout | O | O (not as flexible or advanced as Makie.jl) | - (not through web interface) | O (not as flexible or advanced as Makie.jl) |
| Munge GWAS summary statistics | O | - | - | O (MungeSumstats) |
| Plot an arbitrary genomic range | O | - | - | O (scatter plots yes, gene bodies no) |
| Plot an arbitrary number of genes | O | - | - | - (existing tools reach performance bottleneck) |
| Color SNPs by LD | O | O (LD precomputed) | O (LD precomputed) | O (bigsnpr + custom Rcpp code to compute LD) |
| Plotting various functional genomic data modalities (e.g. ATAC-seq peaks, enhancer-promoter loops) | O | - | - | O (plotgardener) |
| Can visualize the entire ~10 Mb extended MHC region | O | - | - | - (not gene bodies) |

Makie.jl (Danisch and Krumbiegel, 2021) is a Julia plotting package that provides powerful plotting utilities and recipes that can be easily extended to visualize most (if not all) genomic data. Makie.jl permits visualization of millions of data points with ease and speed. For example, to the best of our knowledge, Makie.jl is the only plotting package, where it is possible to visualize the entire LD matrix among ~66,000 SNPs in the MHC region on a personal laptop (e.g. 2.3 GHz 8-Core Intel Core i9, 32 GB RAM). This corresponds to visualization of ~2.2 billion unique data points. In addition, Makie.jl is distinguished from other plotting packages (and their extensions) in that it comes with advanced layout tools. By using Makie.jl’s flexible layout tools, it can be also almost effortless to combine and plot various genetic and genomic data with very complex layouts (https://docs.makie.org/stable/tutorials/layout-tutorial/). Reproducible scientific research involves being able to entirely reproduce its scientific figures, however complex they may be, and this can be easily attained by using Makie.jl.

### Plotting phenome-scale LocusZoom plots

Here, we present the Julia package GeneticsMakie.jl, which builds upon and extends Makie.jl’s plotting tools to generate publication-quality figures visualizing multiple genetic and genomic data modalities on different layers, as shown for the *GCKR* locus (**Figure 1**). To further ease this process, we provide functions for munging GWAS (or other association) summary statistics, which can come in various formats (Lyon *et al*., 2021; Murphy *et al*., 2021; Bulik-Sullivan *et al*., 2015) and therefore require harmonization. Briefly, GeneticsMakie.jl harmonizes column names of GWAS or QTL summary statistics, their SNP IDs, and calculates Z-scores if they are missing. We note that GWAS loci oftentimes harbor extremely small *P* values, in which cases the *P* values cannot be represented by a floating-point number and hence shared as zero. GeneticsMakie.jl mitigates this issue by clamping *P* values of such SNPs to the smallest floating-point number, when munging summary statistics. Such cases tend to be more common in phenotypes that are reaching saturation in terms of GWAS discovery such as height and weight. To fundamentally address this issue, the *P* values in GWAS summary statistics need to be shared in a −log_10_ scale or it might be more appropriate to plot alternative measures of strength of association such as Z-scores. Once the summary statistics are munged, we recommend storing and loading them as memory friendly Arrow files using Arrow.jl package, since loading hundreds of genetic association results simultaneously is memory intensive and infeasible, especially on a personal laptop. Then one can iterate through arbitrary genomic regions of interest. For example, GeneticsMakie.jl conveniently provides functions for identifying GWAS loci and their closest (protein-coding) genes so that one can iterate through either GWAS loci or their cognate genes. To color SNPs by LD with a designated SNP of interest (e.g. index or sentinel SNP), any custom reference panel can be loaded using SnpArrays.jl package (Zhou *et al*., 2020) and LD is computed on the fly. Additionally, genes and isoforms with constituent exons and introns can be plotted with any custom transcriptome annotation file in GTF format. These functionalities form the backbone of LocusZoom plots and other functional genomic data can be added as separate layers as needed, using GeneticsMakie.jl’s provided functions or custom Makie.jl code. It is worth nothing that all these functions can be customized or extended easily with minimal loss of performance, which is an inherent strength of the Julia programming language. We share example code for such a workflow in https://github.com/mmkim1210/GeneticsMakieExamples.

**Figure 1.**
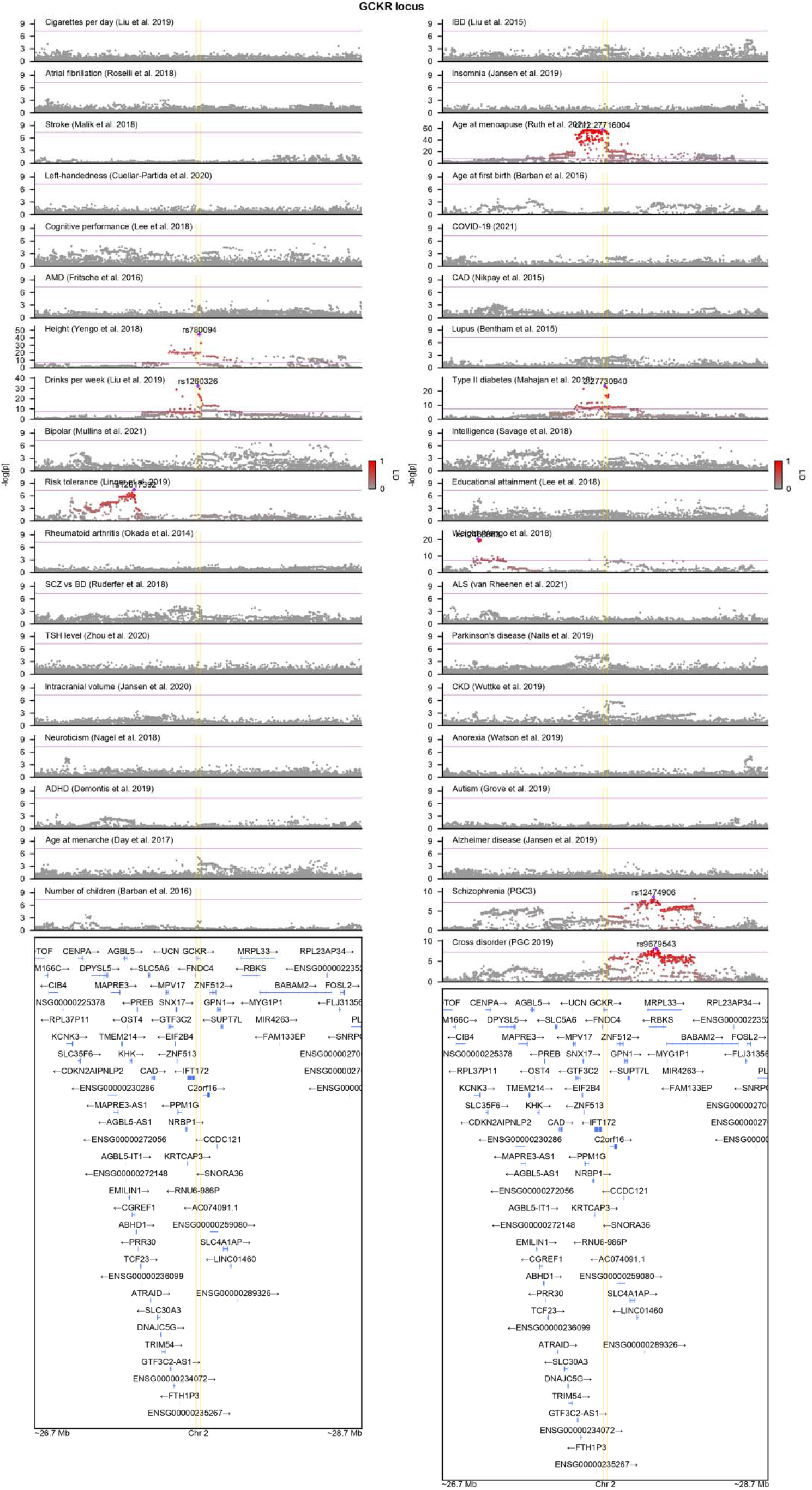
A close look at the known pleiotropic *GCKR* locus using GeneticsMakie.jl. Index SNPs for phenotypes harboring GWAS hits are labeled and corresponding linkage disequilibrium (LD) between other SNPs are displayed with the intensity of red color. Purple line denotes genome-wide significance (*P* = 5e-8), and yellow lines denote gene start and end sites for *GCKR* gene. Figure generated on a personal laptop (e.g. 2.3 GHz 8-Core Intel Core i9, 32 GB RAM) without any further modifications. Each GWAS result contains ~3,000 SNPs.

To further showcase the power of GeneticsMakie.jl, we share phenome-wide LocusZoom plots for 239 GWAS loci for schizophrenia (**Supplementary Data**; Schizophrenia Working Group of the Psychiatric Genomics Consortium *et al*., 2020), which are defined as non-overlapping ±1 Mb windows around the most significantly associated SNPs. We also share such LocusZoom plots for genomic regions known to harbor long-range LD (Anderson *et al*., 2010), which includes the MHC region. Finally, we share LocusZoom plots for high-confidence neuropsychiatric risk genes, which loss-of-function is implicated in increased risk for schizophrenia (Singh *et al*., 2020), autism spectrum disorder (Satterstrom *et al*., 2020), and developmental delay disorder (Kaplanis *et al*., 2020). This permits qualitative assessment of convergence of common variant and rare variant signals for neurodevelopmental and neuropsychiatric disorders.

## Conclusion

In summary, GeneticsMakie.jl allows scalable and flexible visual display of high-dimensional genetic and genomic data within the Julia ecosystem. It produces high-quality, publication-ready figures by default. As the volume and diversity of molecular readouts as well as genetic association results continue to increase, trans-phenotype, trans-tissue, trans-cell-type, trans-ethnic, and trans-omic analyses that consider multiple layers of data will become ever more important. We envision GeneticsMakie.jl facilitating these types of multivariate analyses by enabling fast and seamless data visualization within the larger Julia data science and OpenMendel ecosystems (Zhou *et al*., 2020). GeneticsMakie.jl further supports efficient generation of Manhattan plots and corresponding QQ plots for GWAS summary statistics. It can be also used to visualize gene-level association results. In the future, we foresee other data modalities being plotted on top of what we already have implemented in GeneticsMakie.jl to provide better interpretation of underlying genetic association.

## Acknowledgments

The authors thank members of the Gandal lab and OpenMendel group for testing the package and providing helpful comments.

