## Supplementary figures and images for "GeneticsMakie.jl: A versatile and scalable toolkit for visualizing locus-level genetic and genomic data"

### 1:48000000-52000000-locuszoom.png

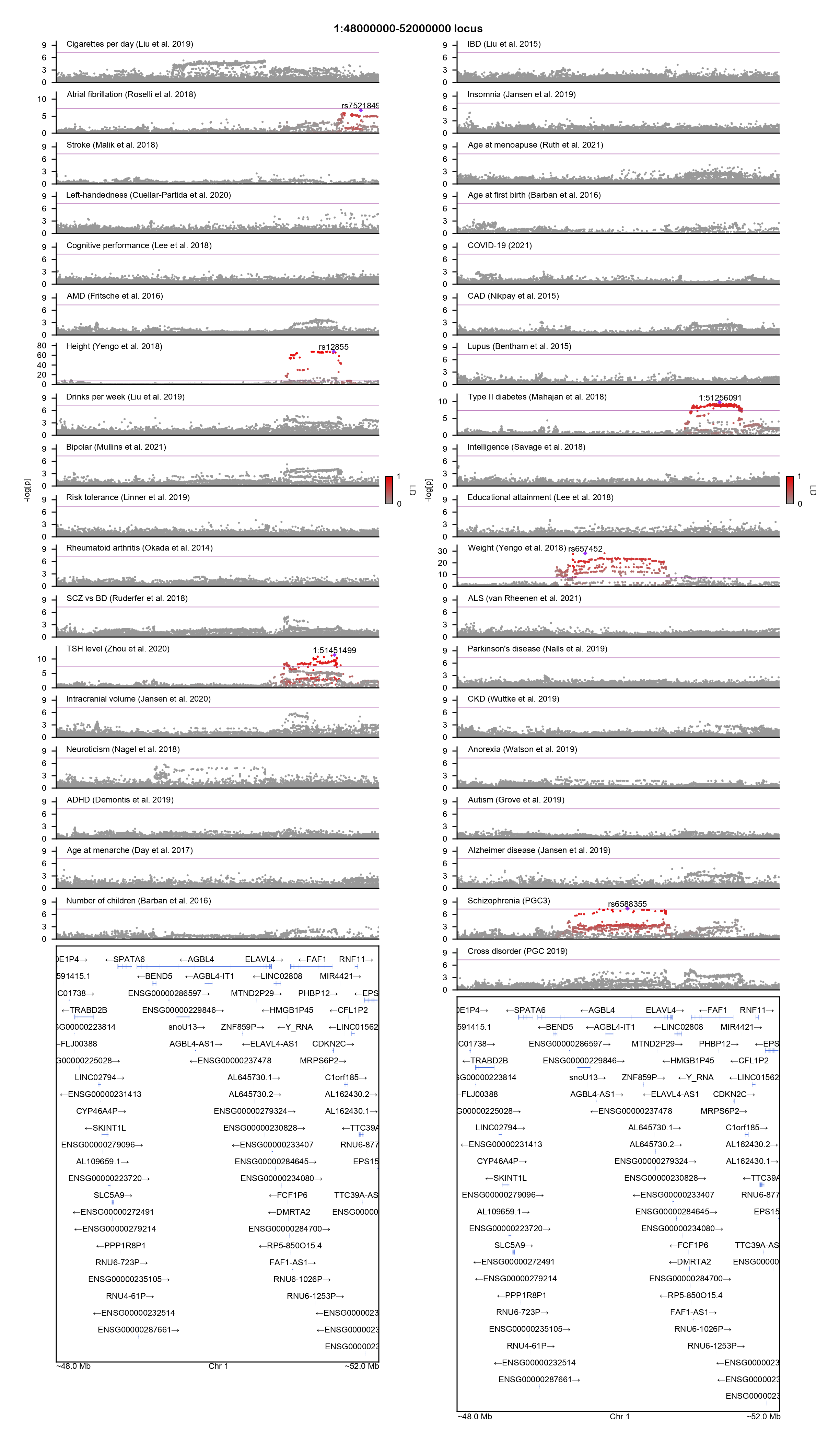

### 2:86000000-100500000-locuszoom.png

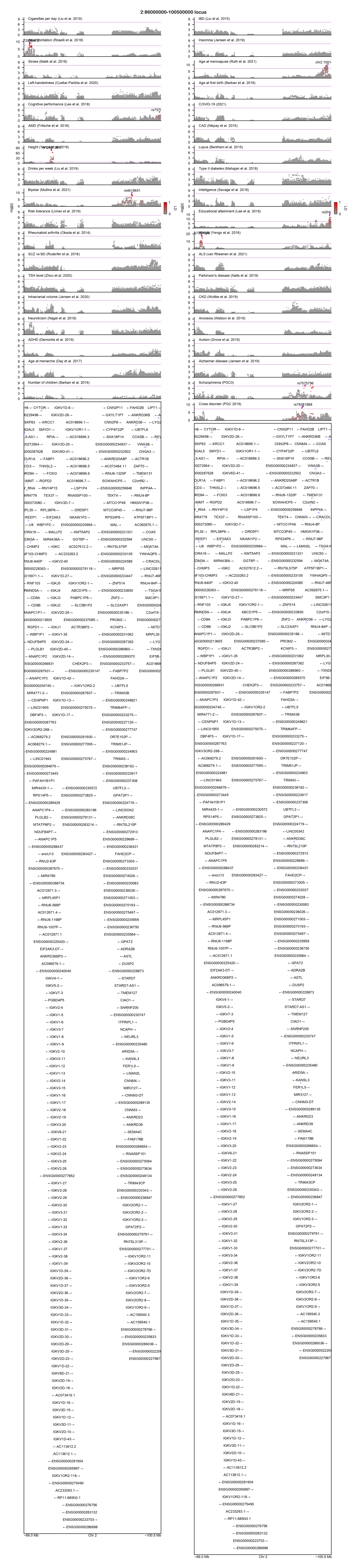

### 2:134500000-138000000-locuszoom.png

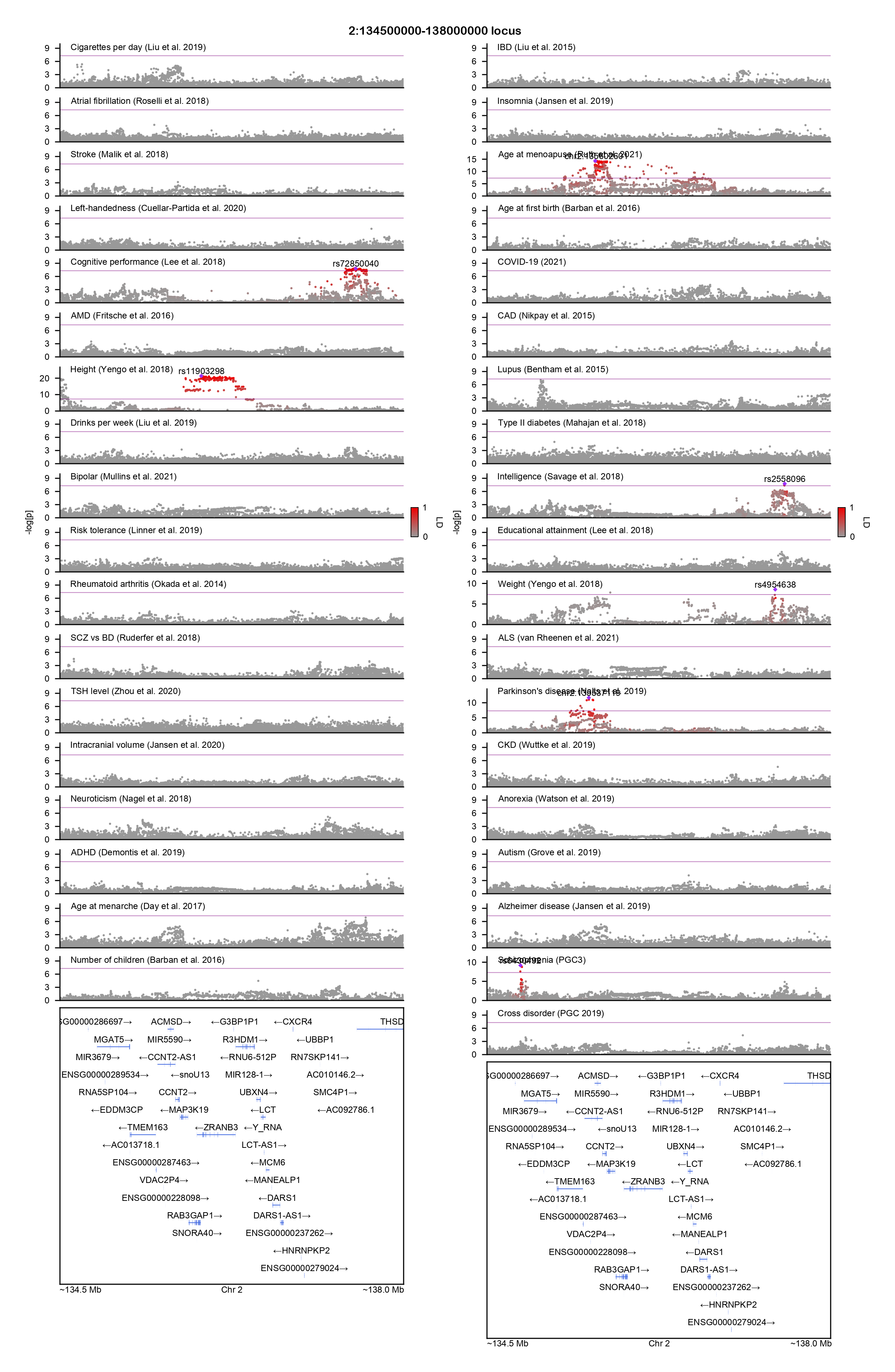

### 2:183000000-190000000-locuszoom.png

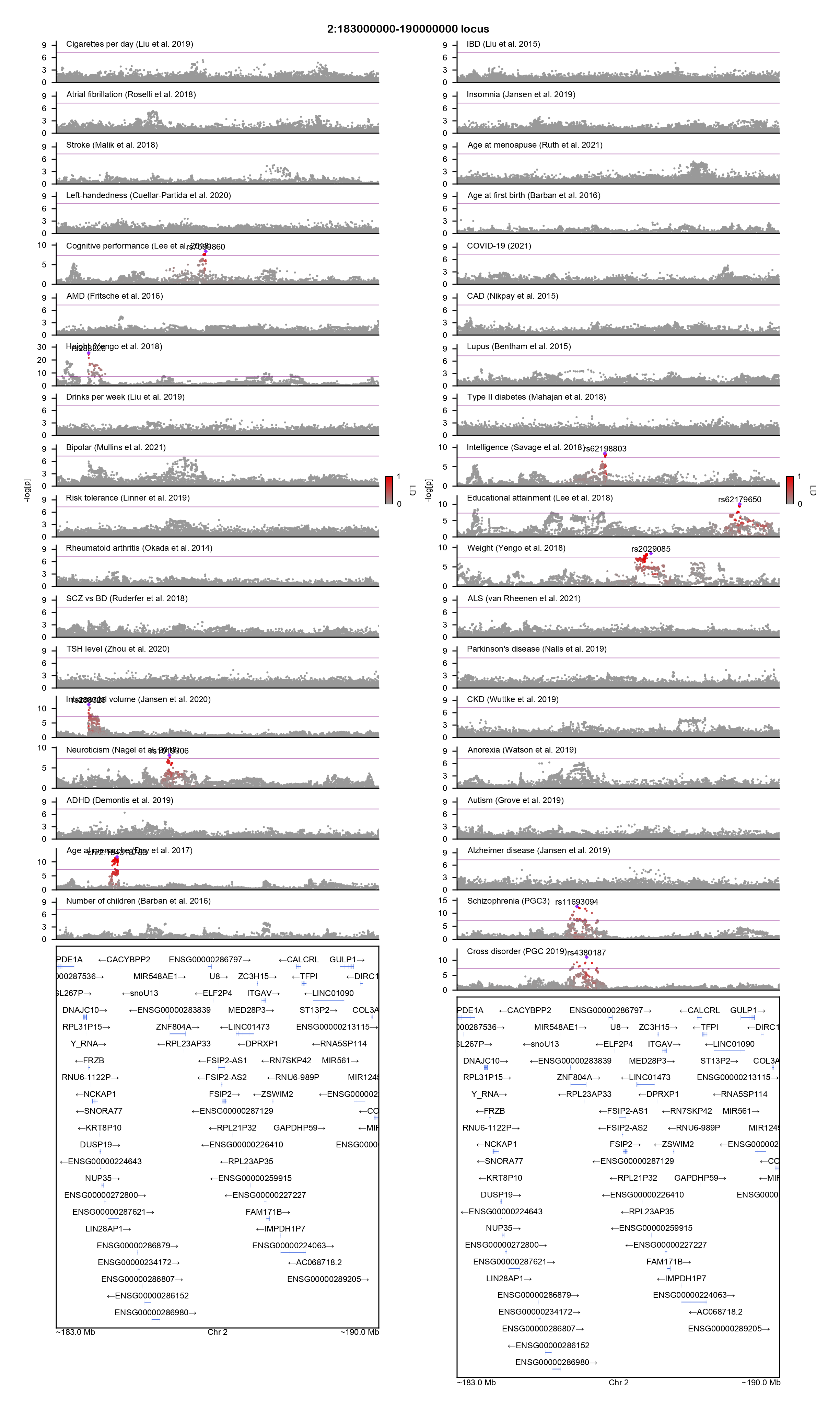

### 3:47500000-50000000-locuszoom.png

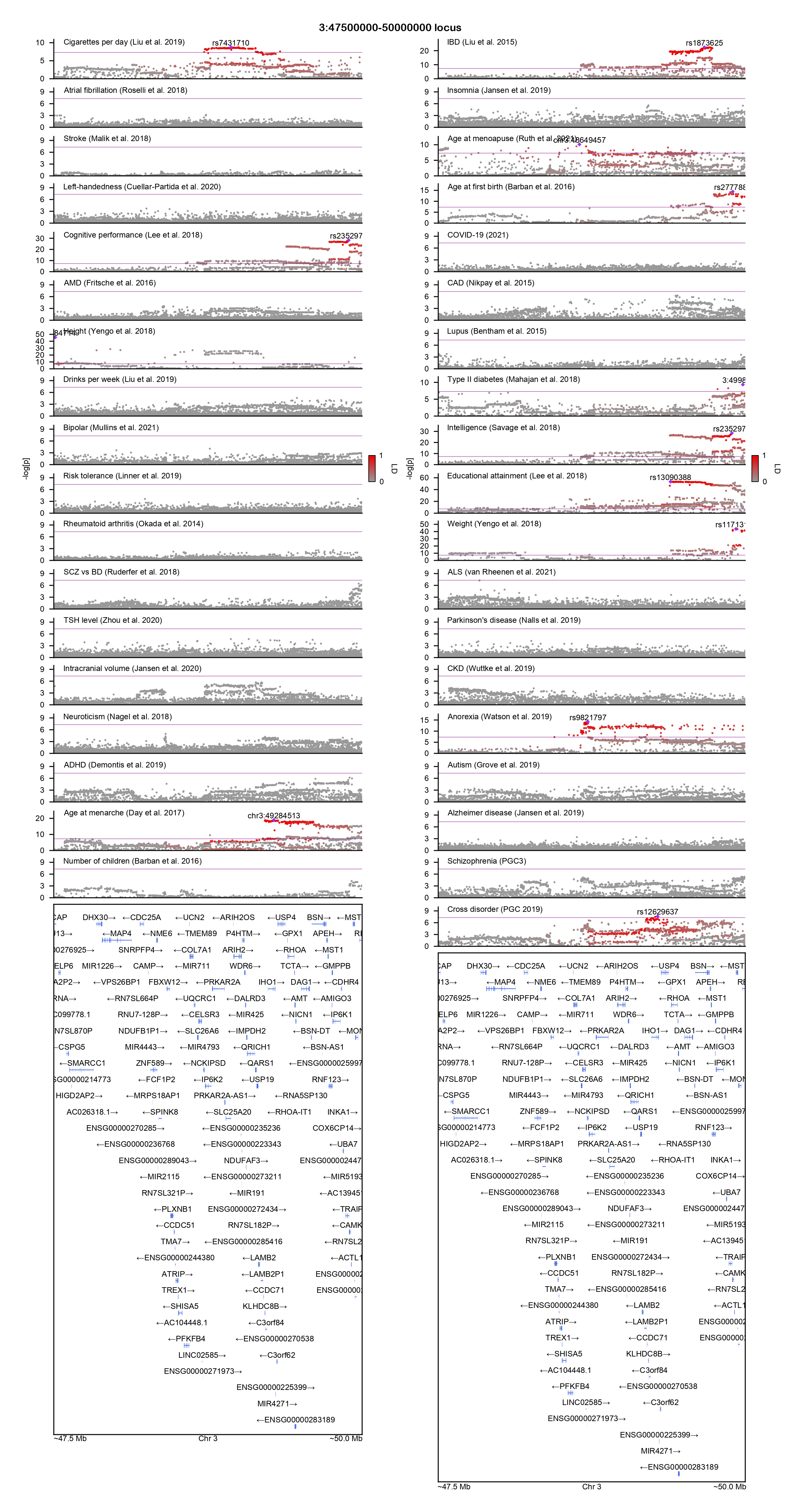

### 3:83500000-87000000-locuszoom.png

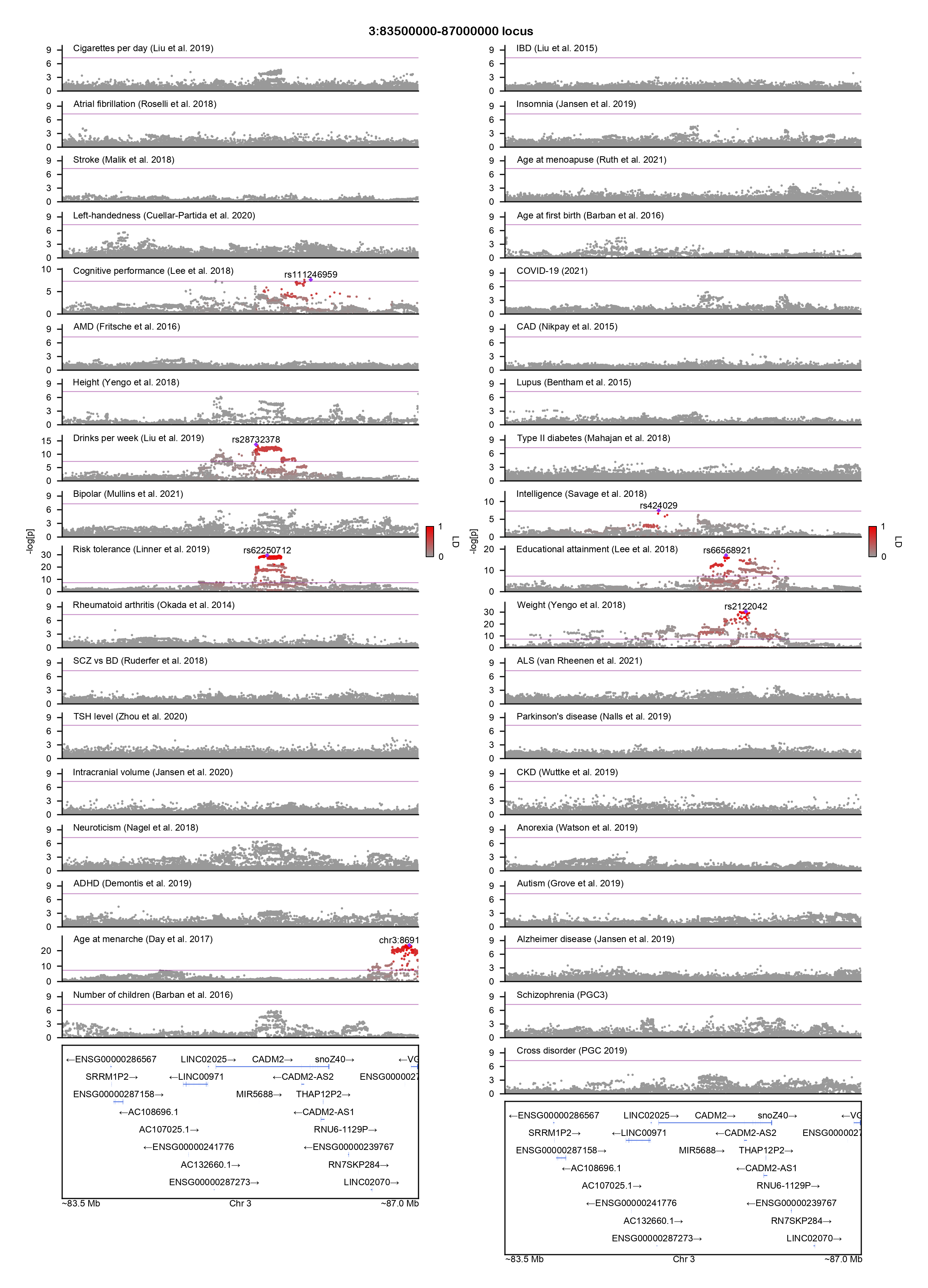

### 3:89000000-97500000-locuszoom.png

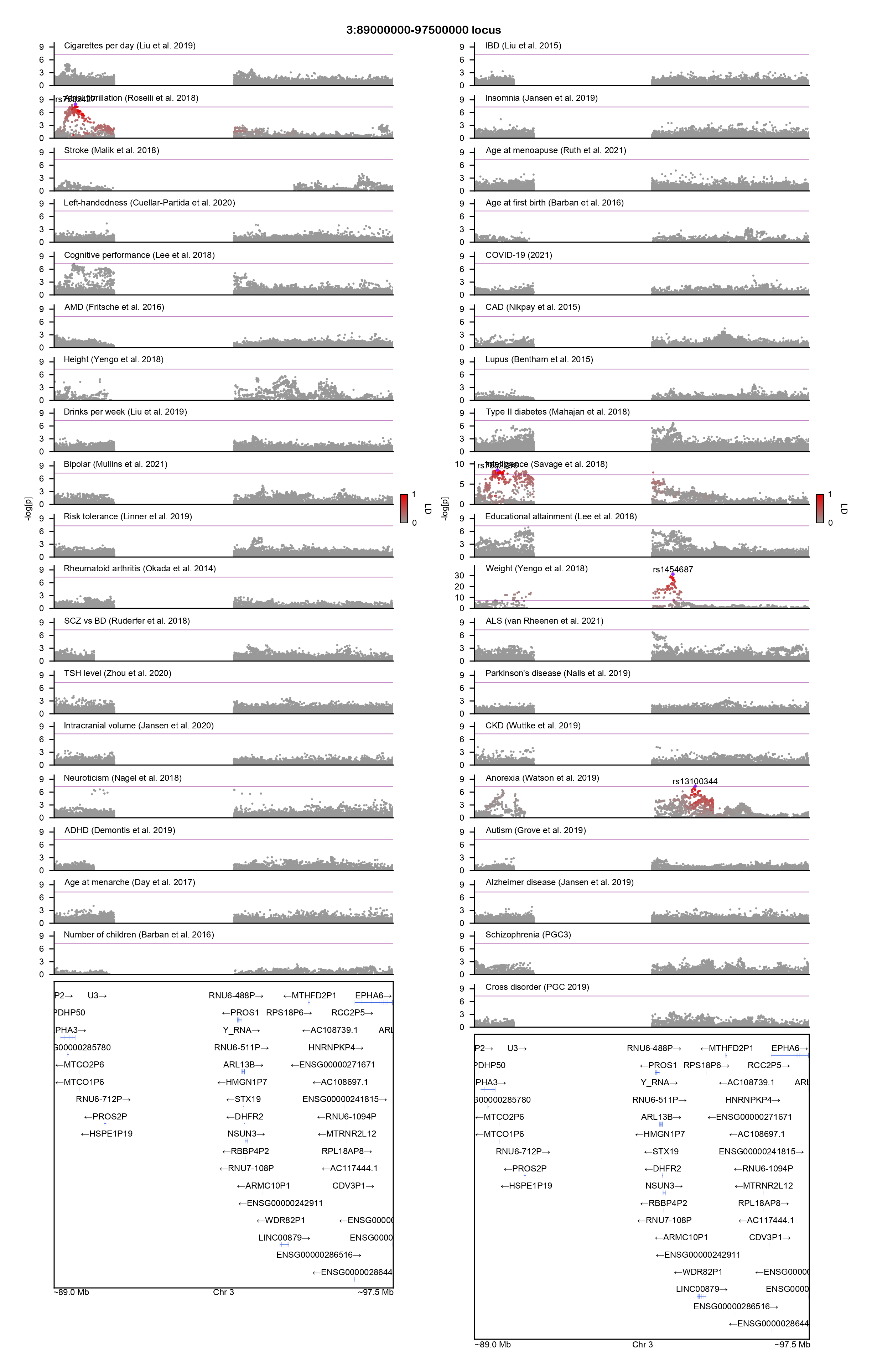

### 5:44500000-50500000-locuszoom.png

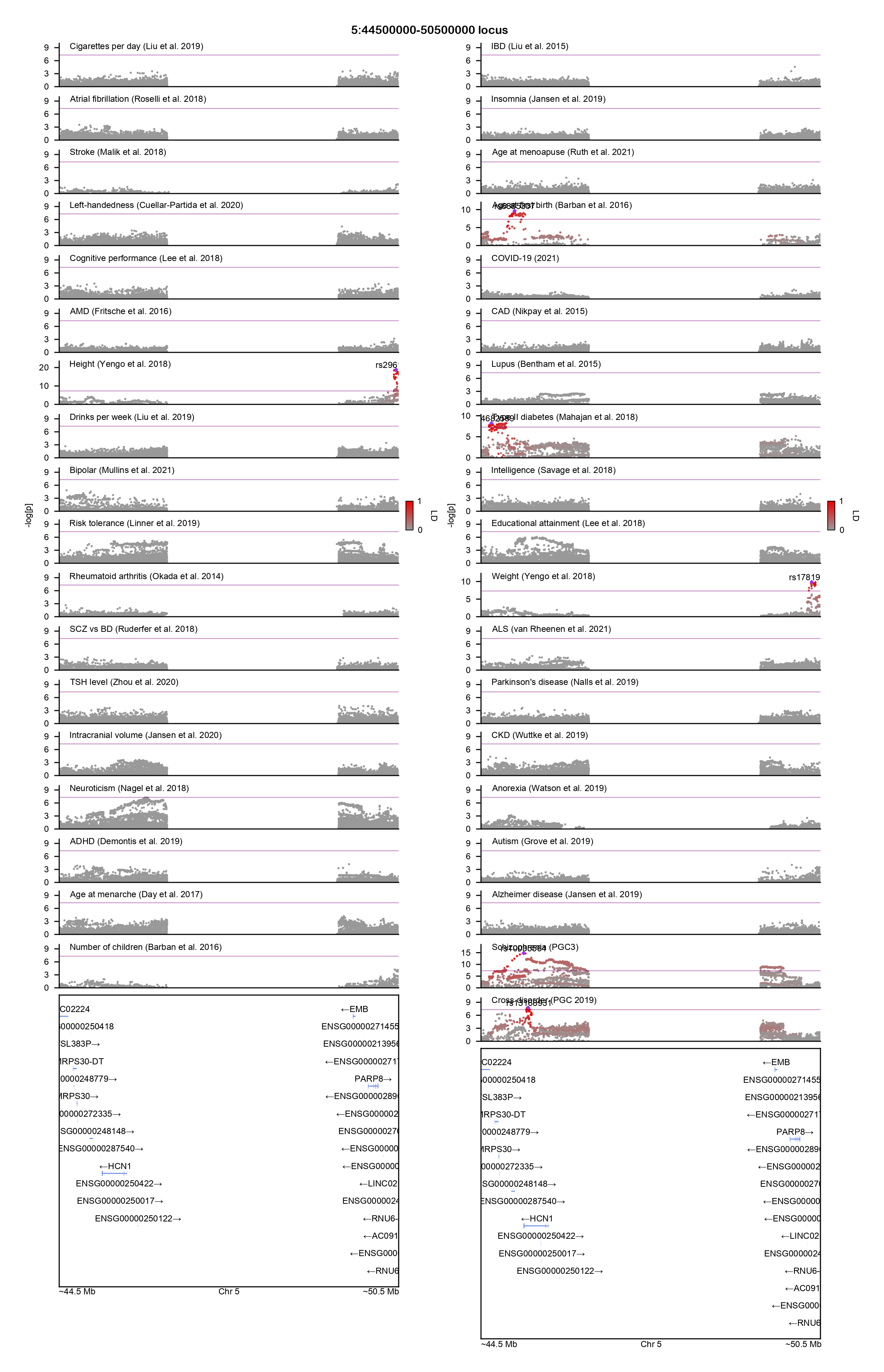

### 5:98000000-100500000-locuszoom.png

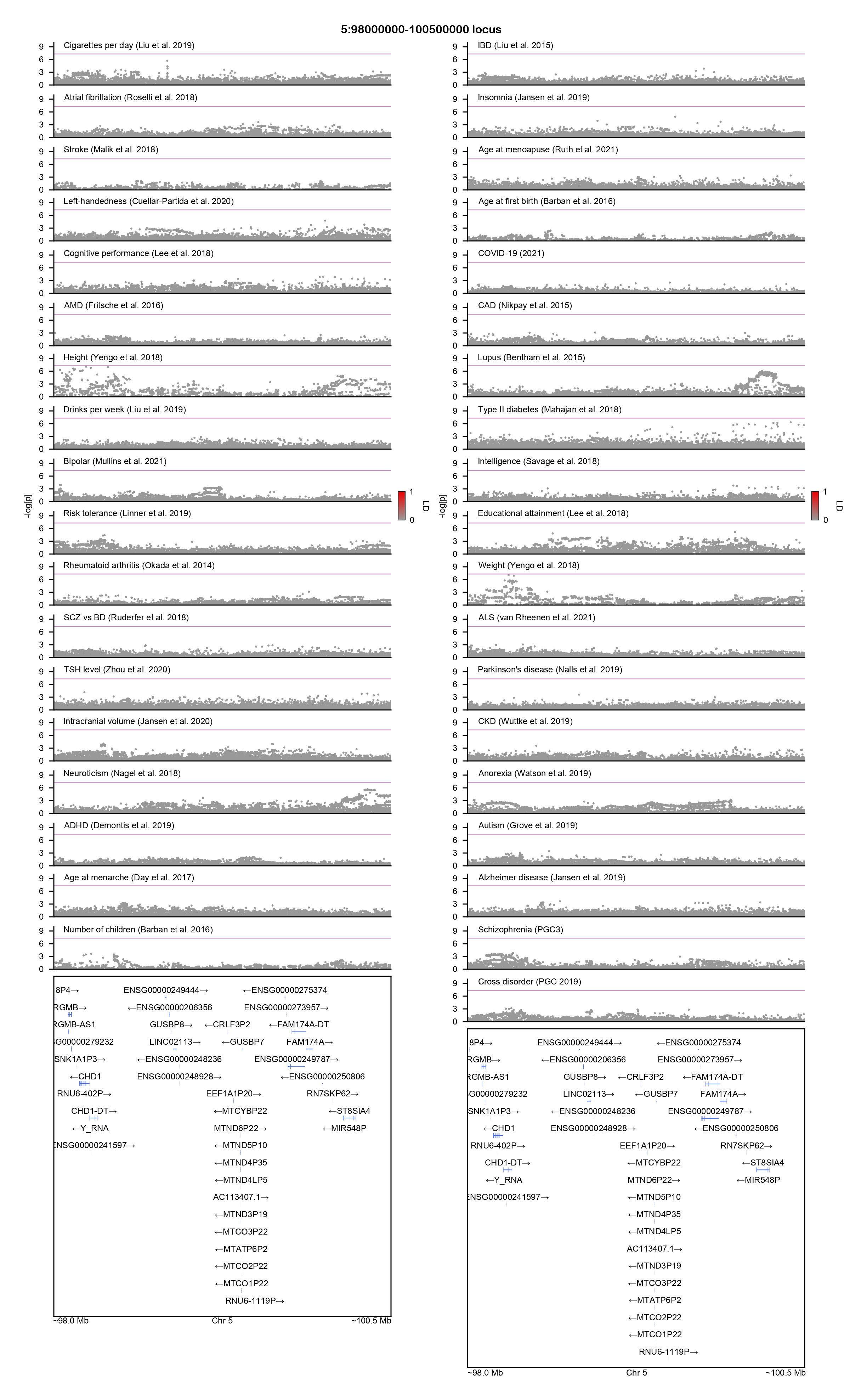

### 5:129000000-132000000-locuszoom.png

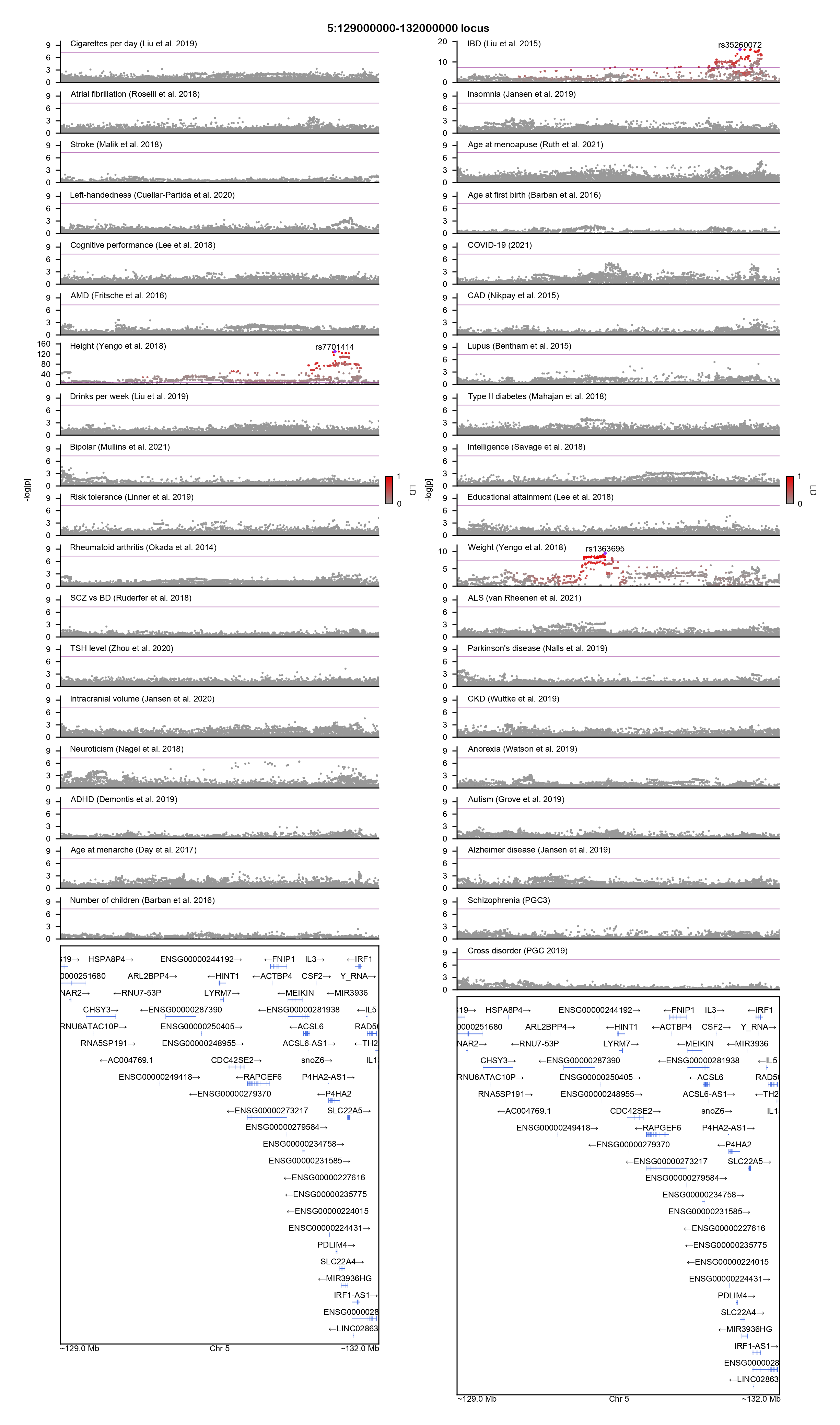

### 5:135500000-138500000-locuszoom.png

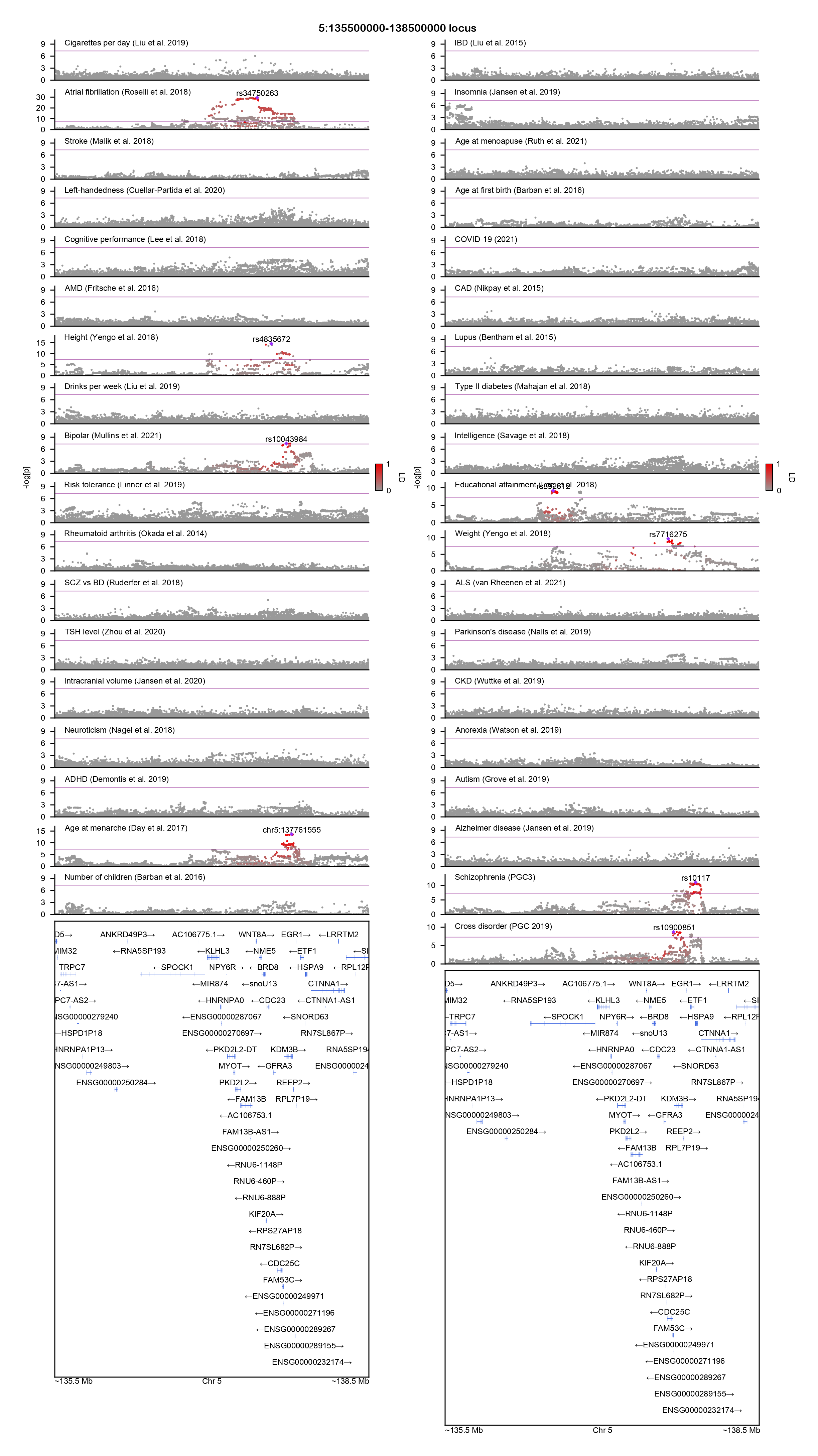

### 6:24000000-36000000-locuszoom.png

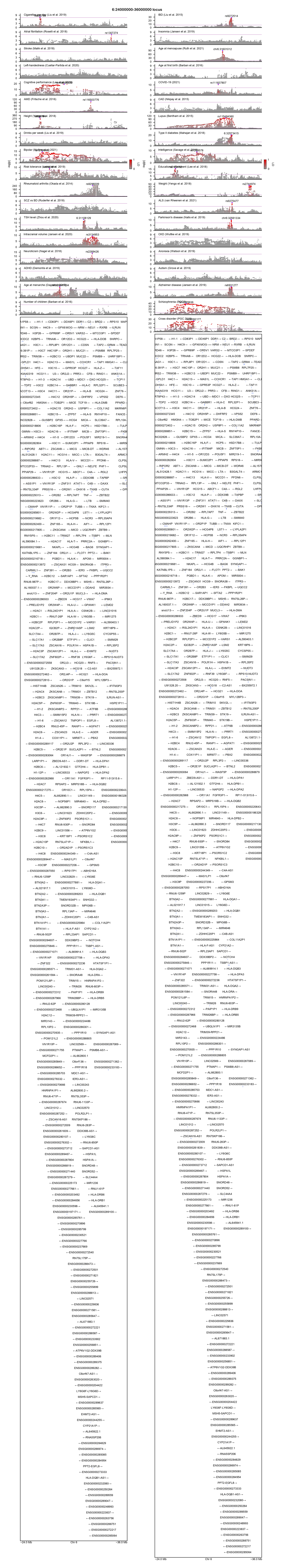

### 6:57000000-64000000-locuszoom.png

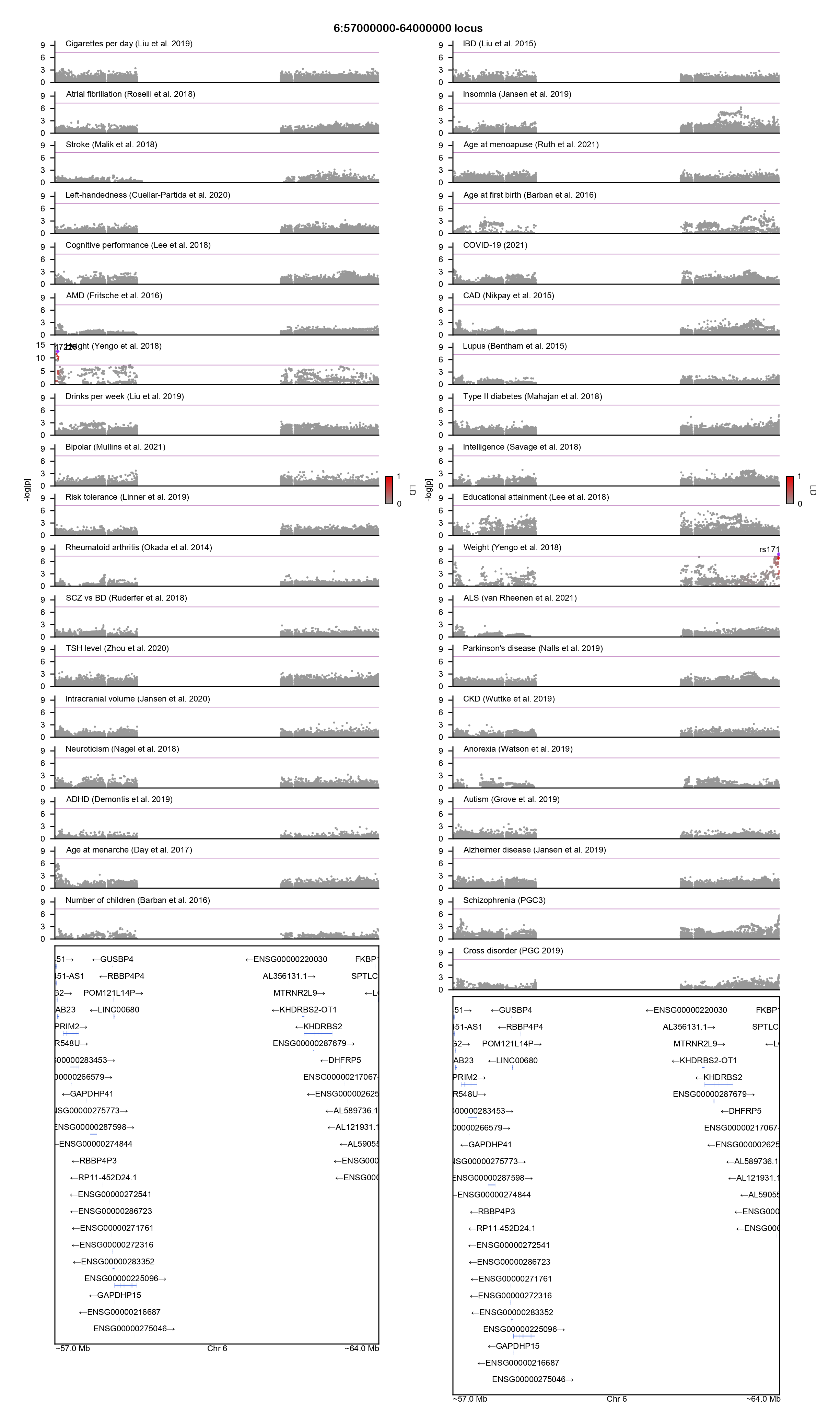

### 6:140000000-142500000-locuszoom.png

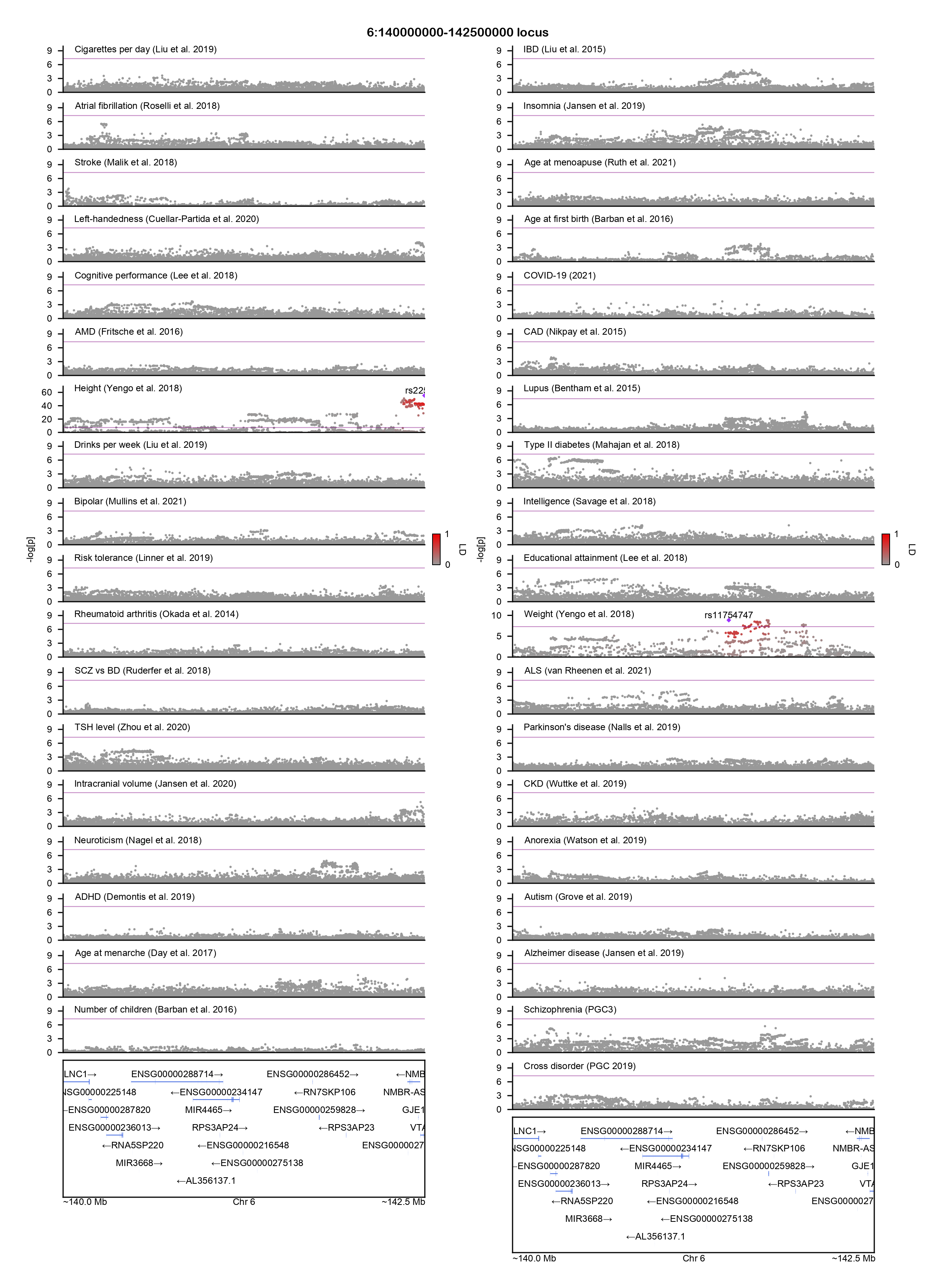

### 7:55000000-66000000-locuszoom.png

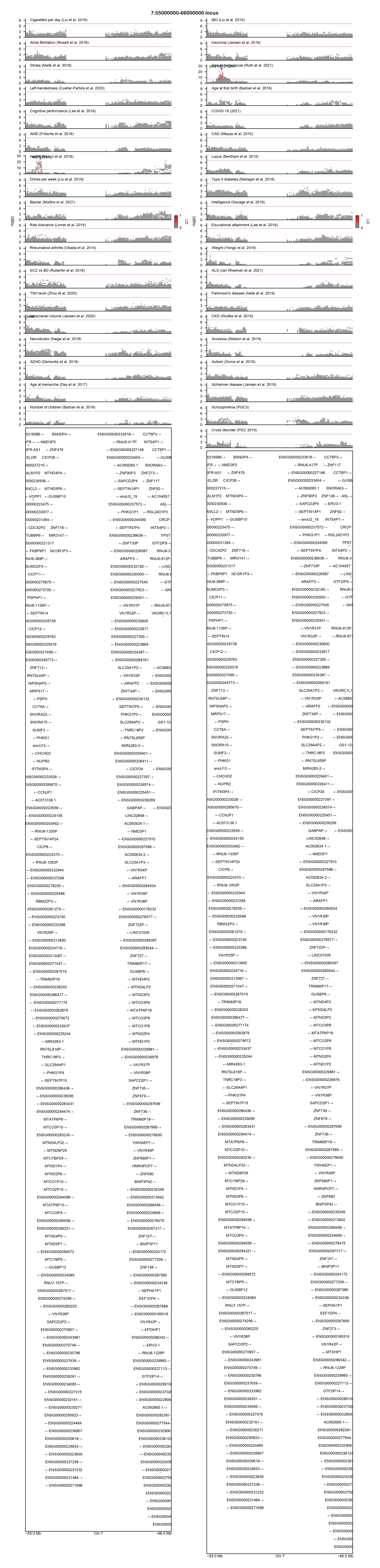

### 8:7000000-13000000-locuszoom.png

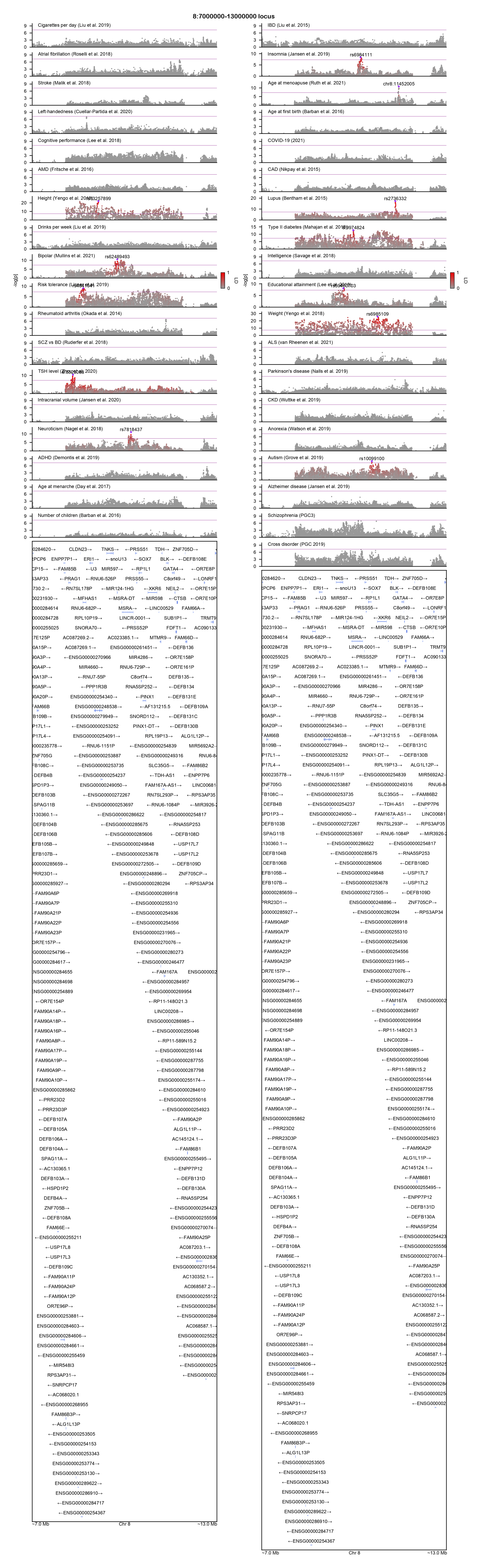

### 8:43000000-50000000-locuszoom.png

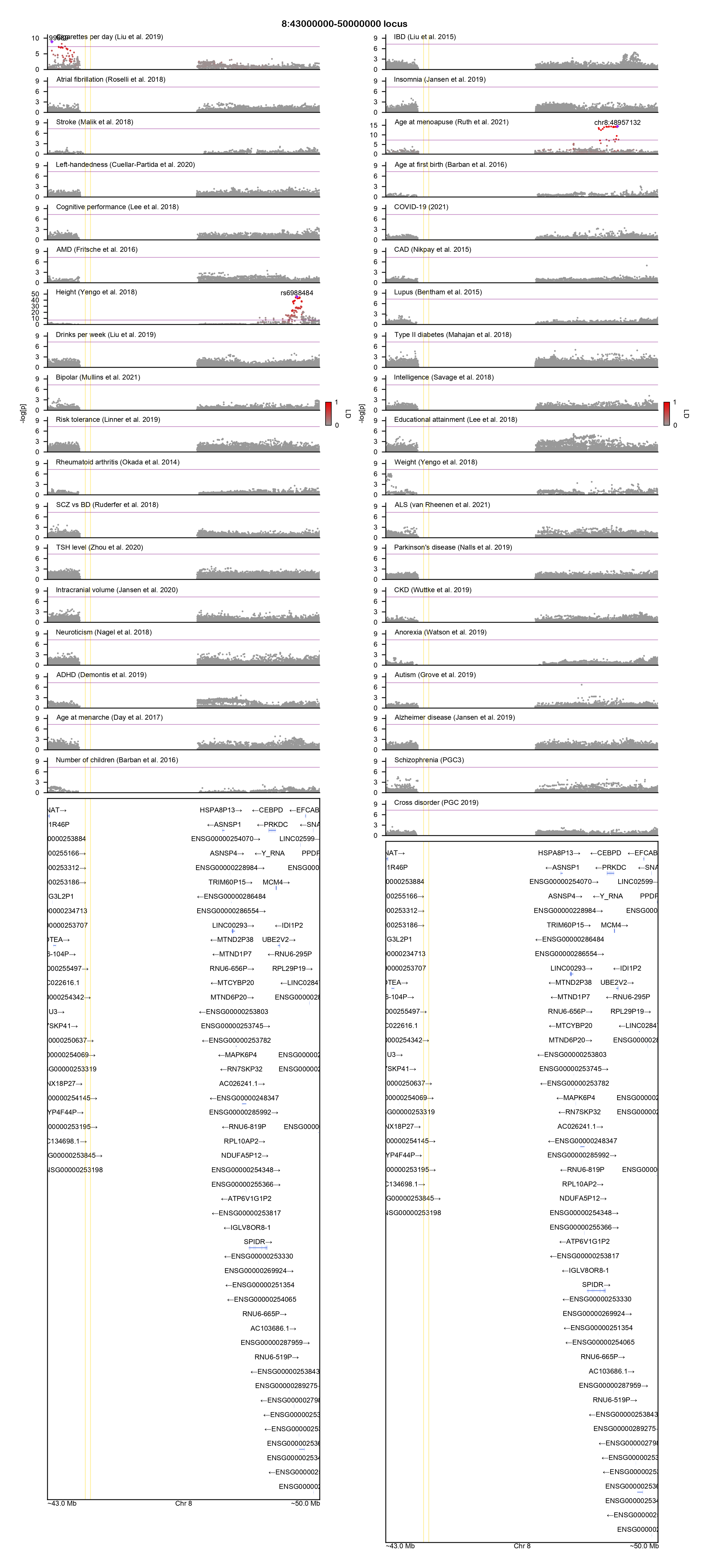

### 8:112000000-115000000-locuszoom.png

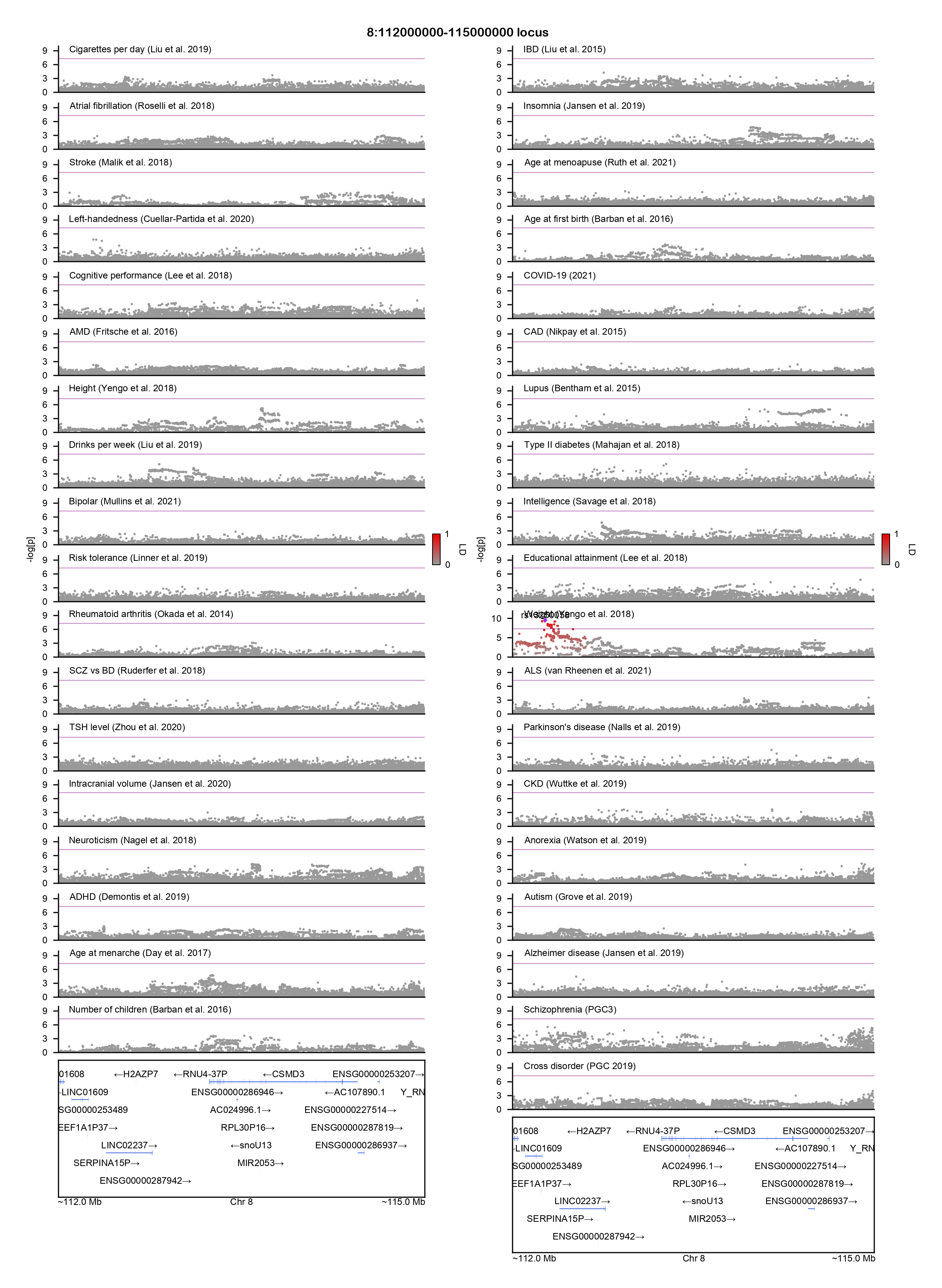

### 10:37000000-43000000-locuszoom.png

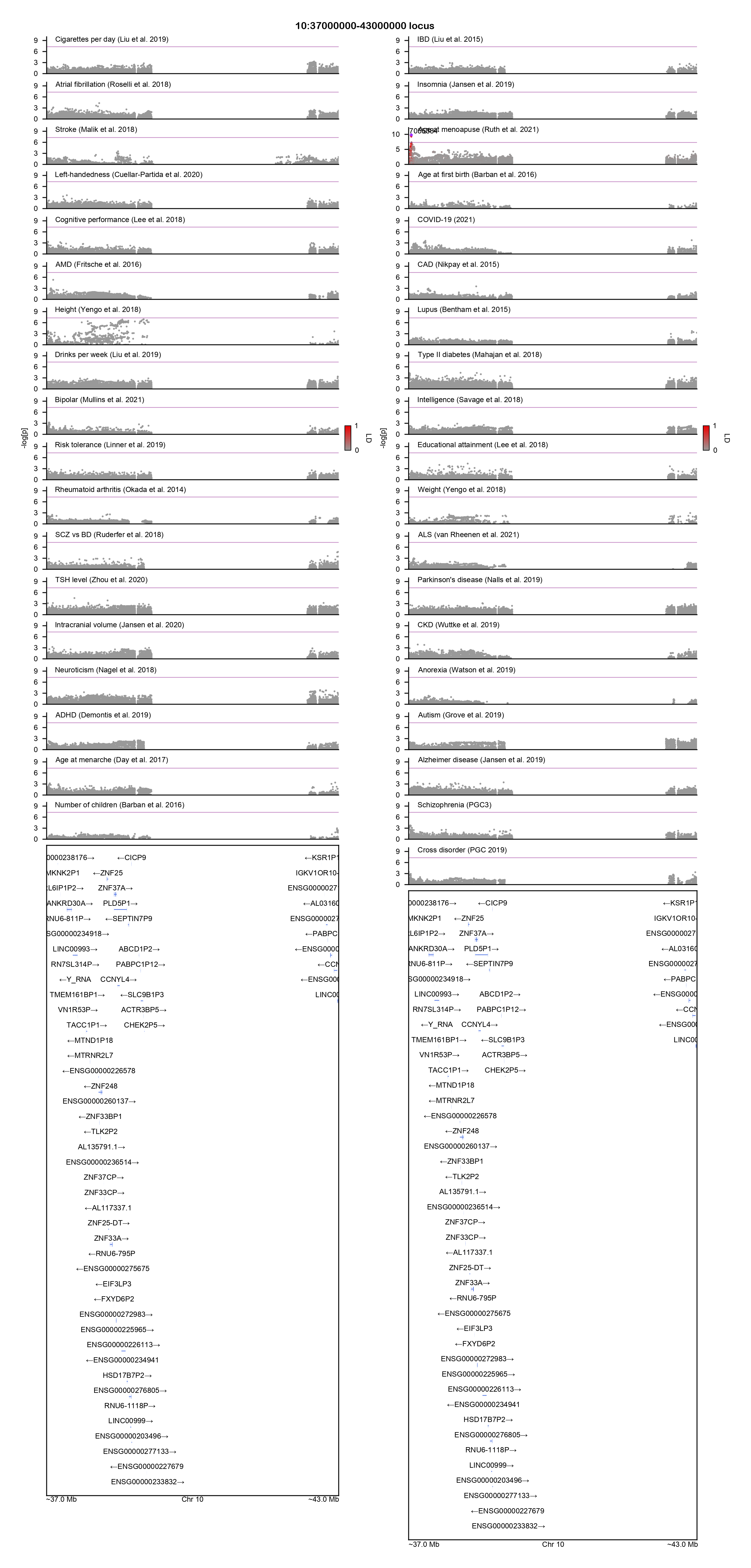

### 11:46000000-57000000-locuszoom.png

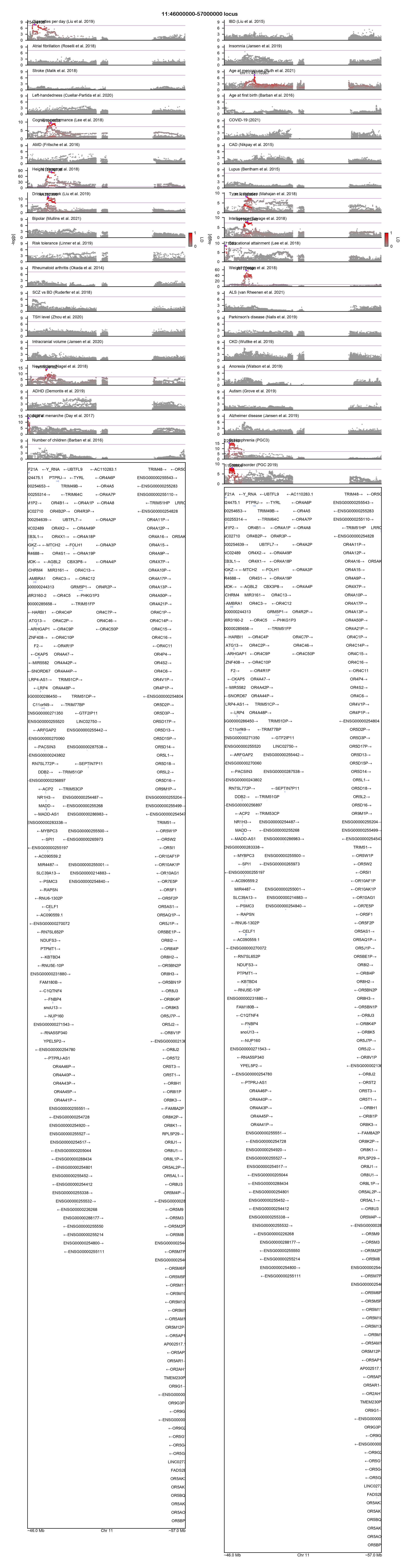

### 11:87500000-90500000-locuszoom.png

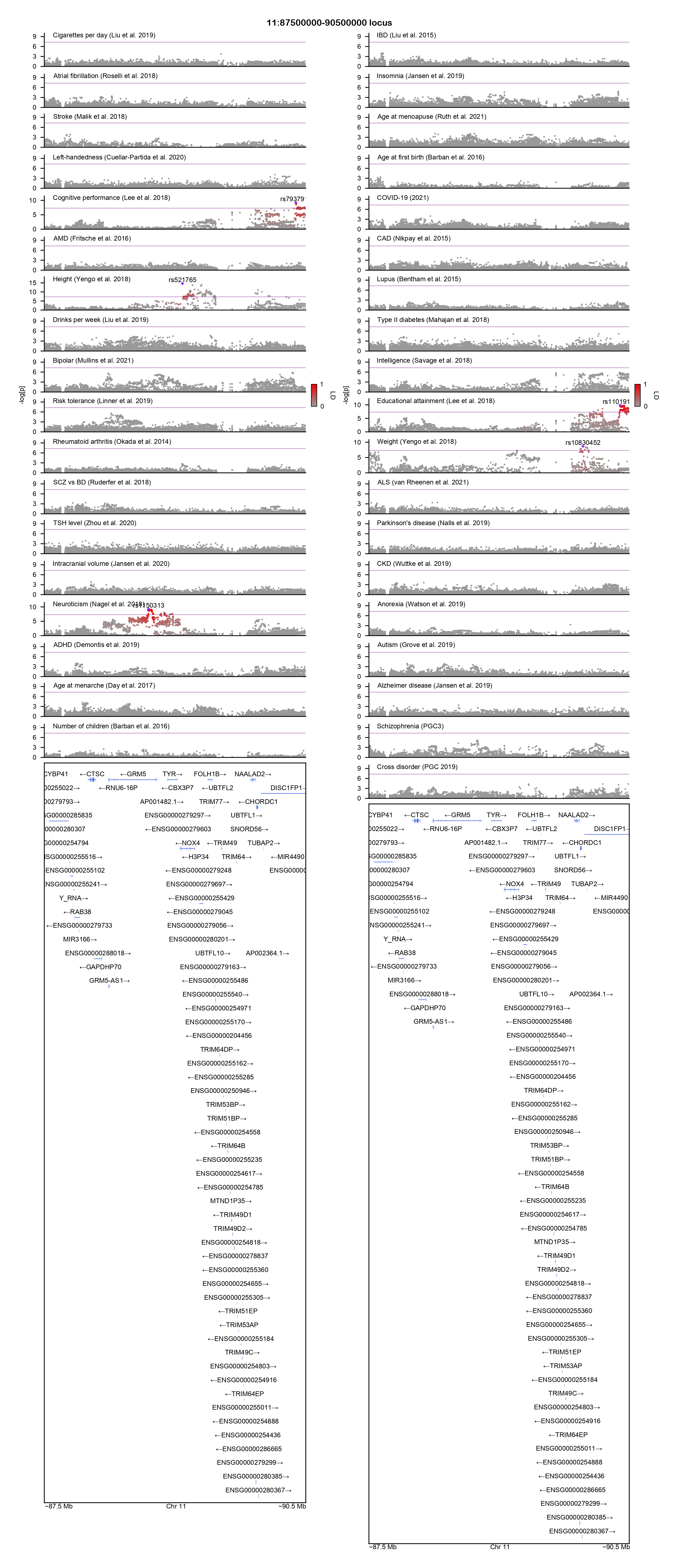

### 12:33000000-40000000-locuszoom.png

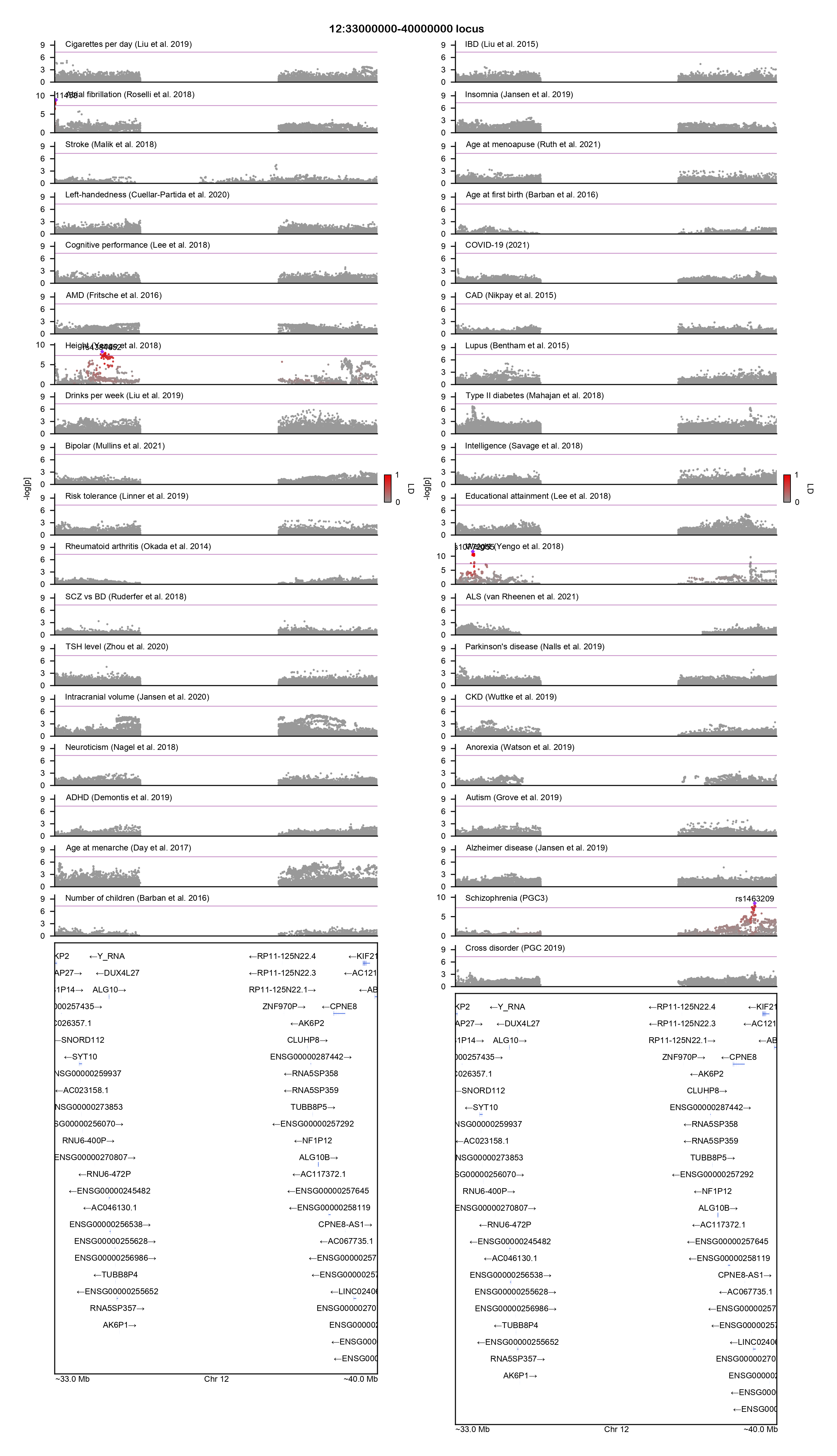

### 12:109500000-112000000-locuszoom.png

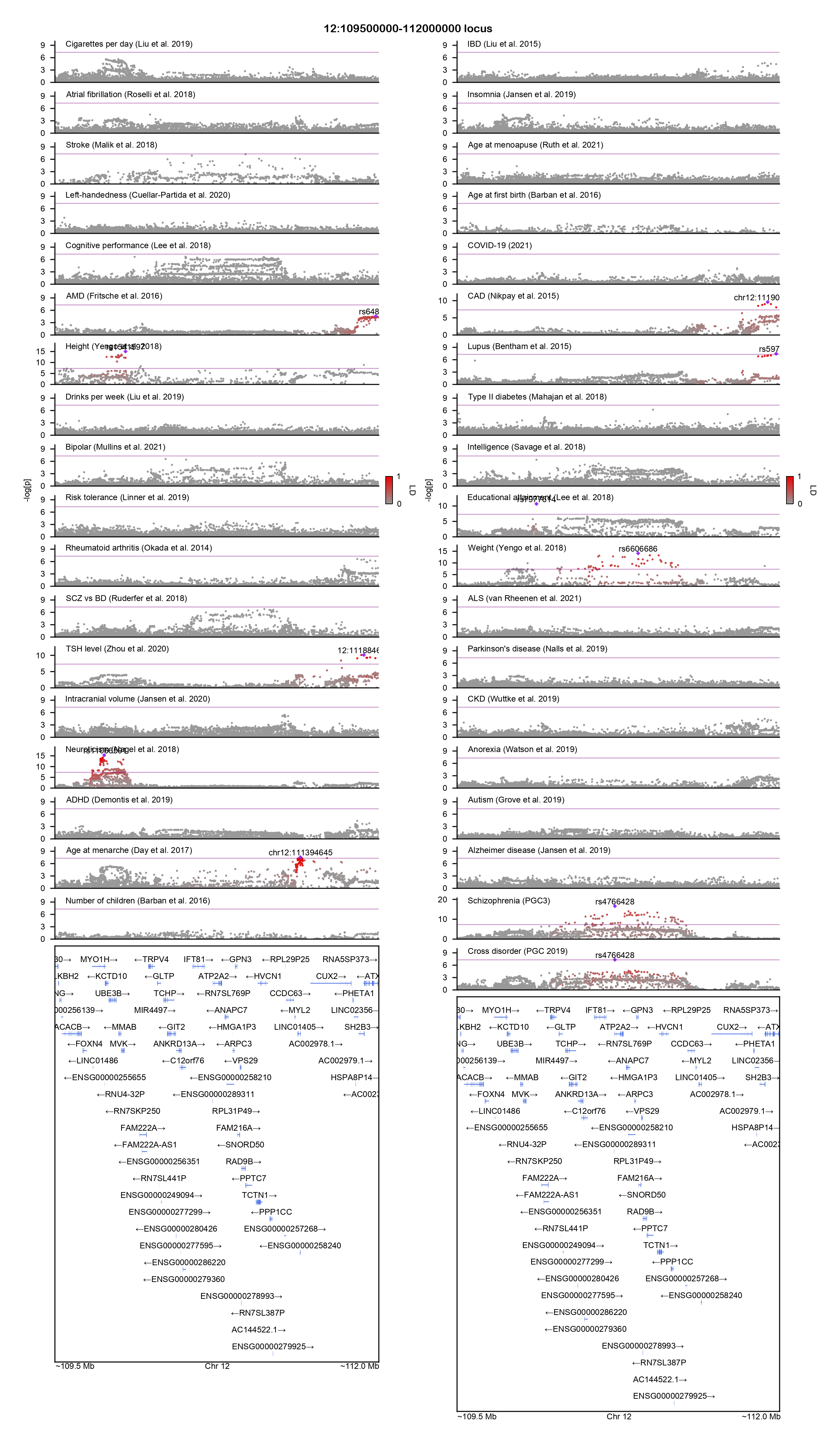

### 17:42900000-45100000-locuszoom.png

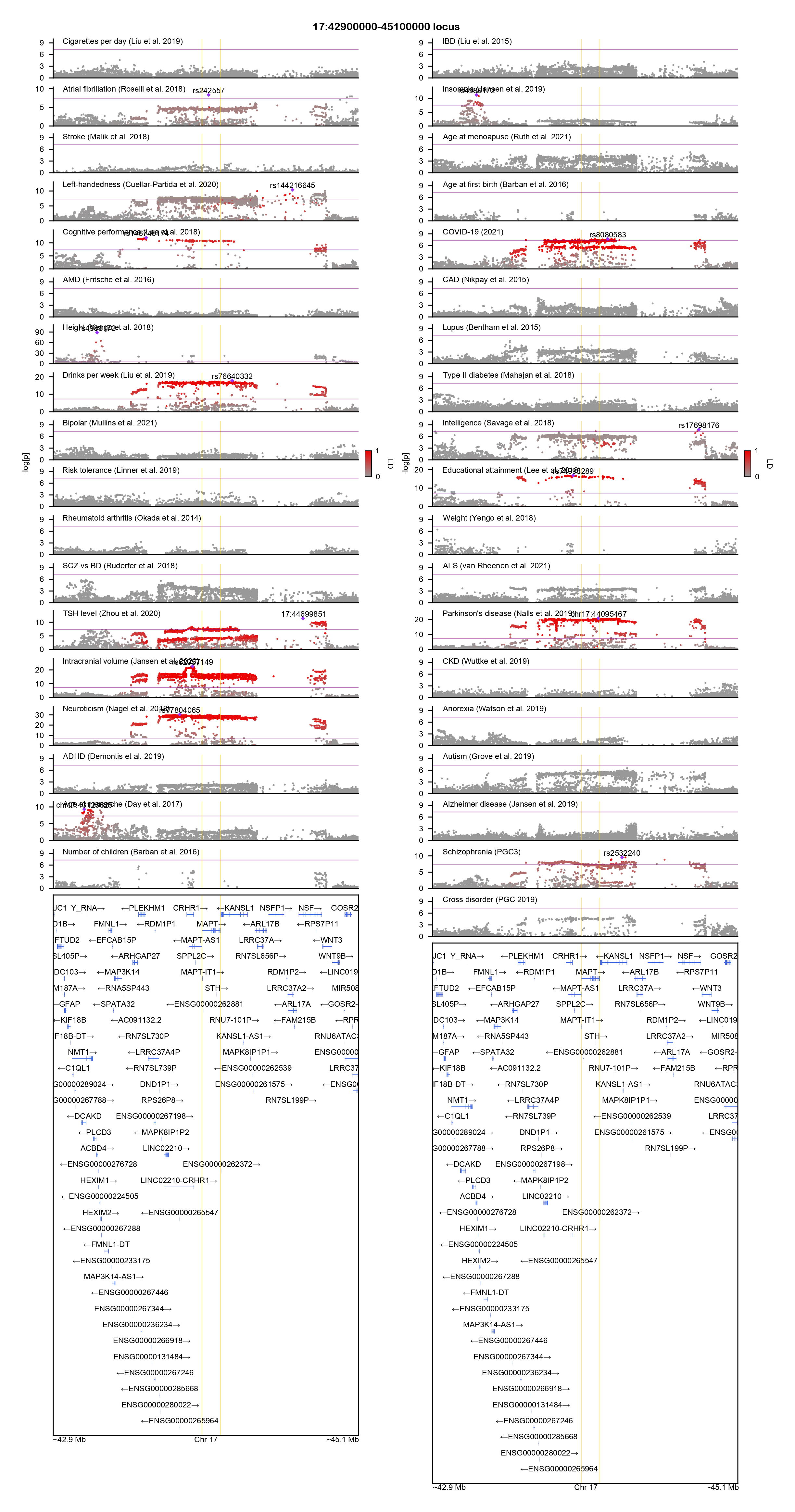

### 20:32000000-34500000-locuszoom.png

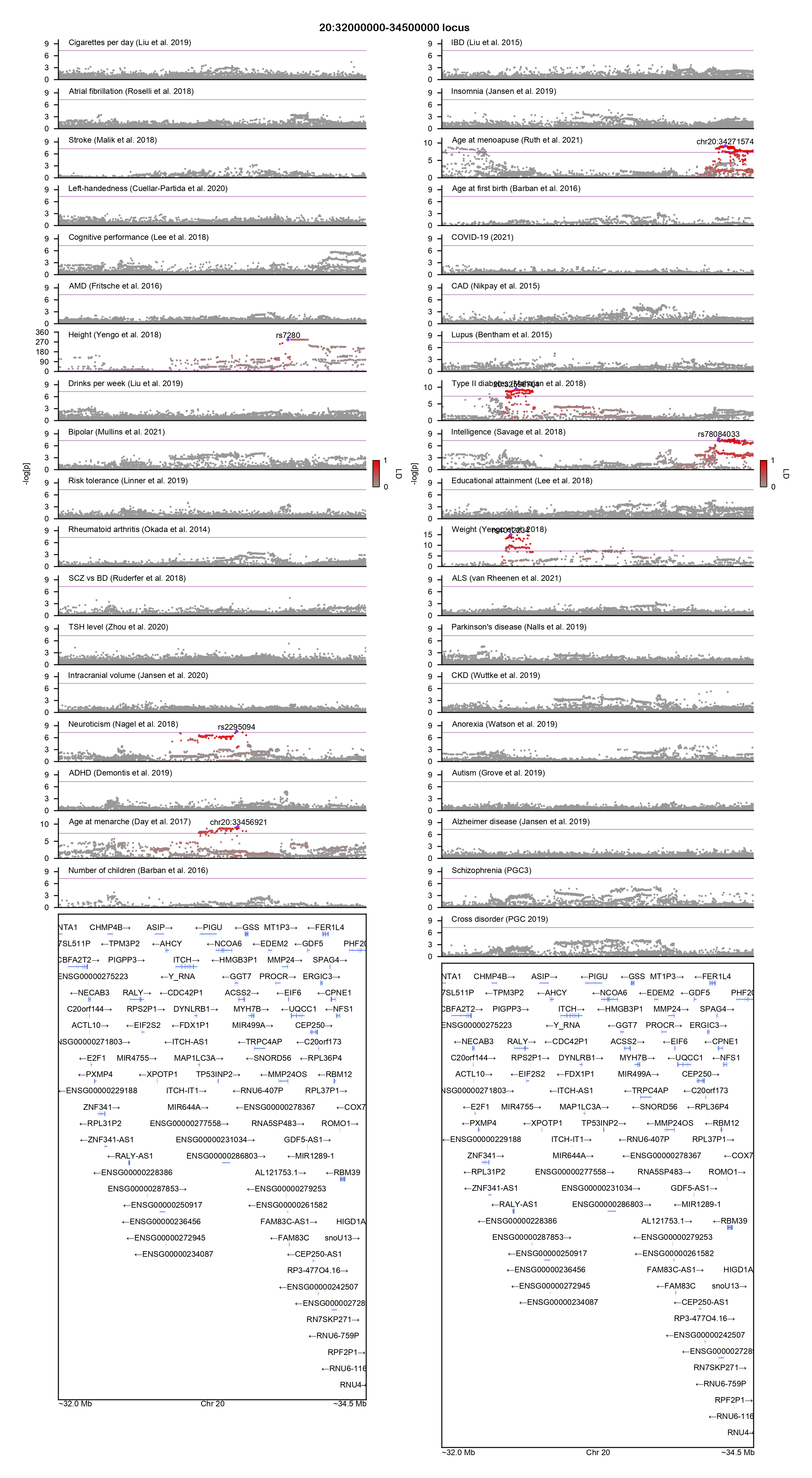

### AGO3-locuszoom.png

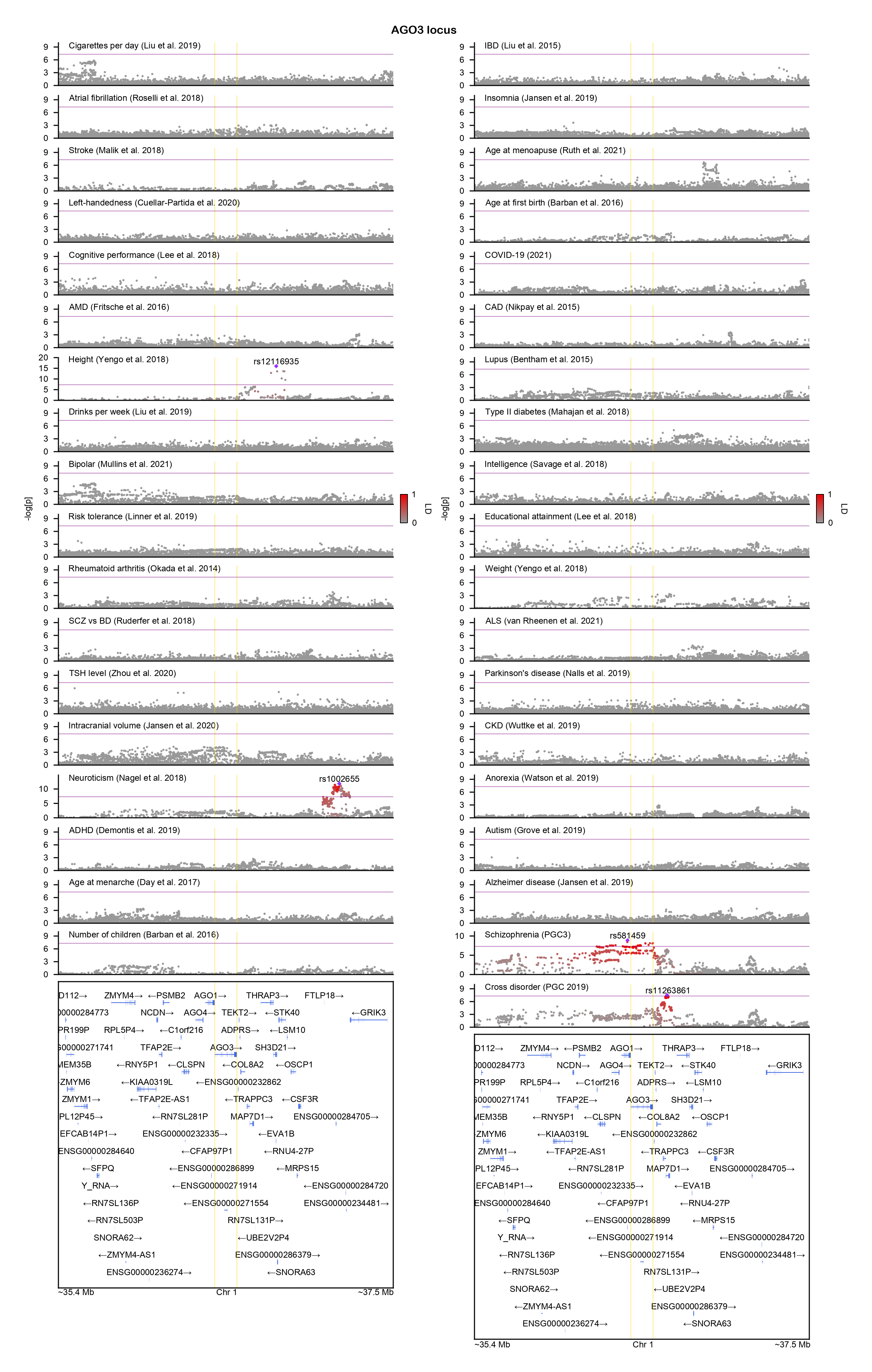

### AL645730.2-locuszoom.png

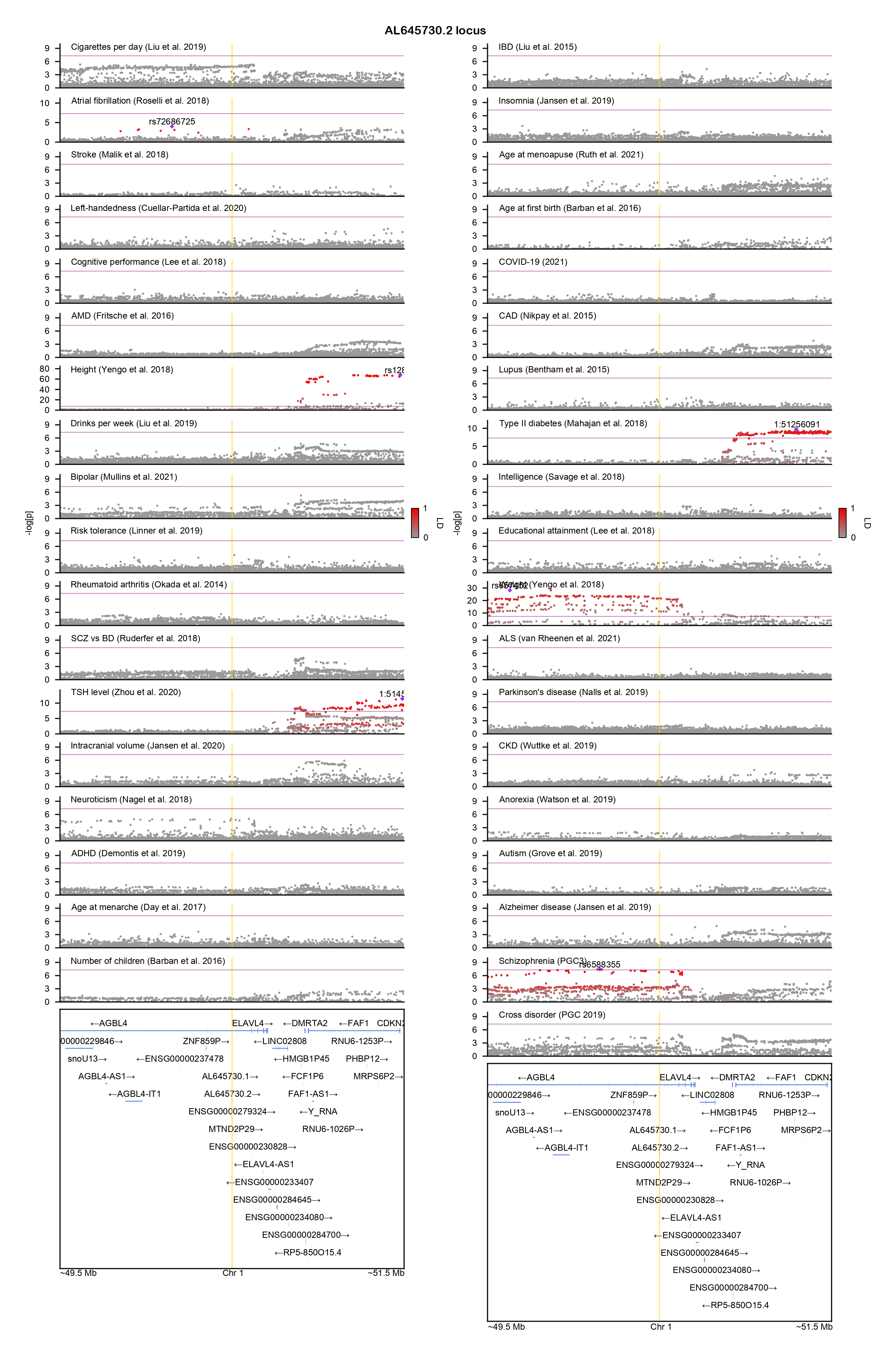

### ARAP2-locuszoom.png

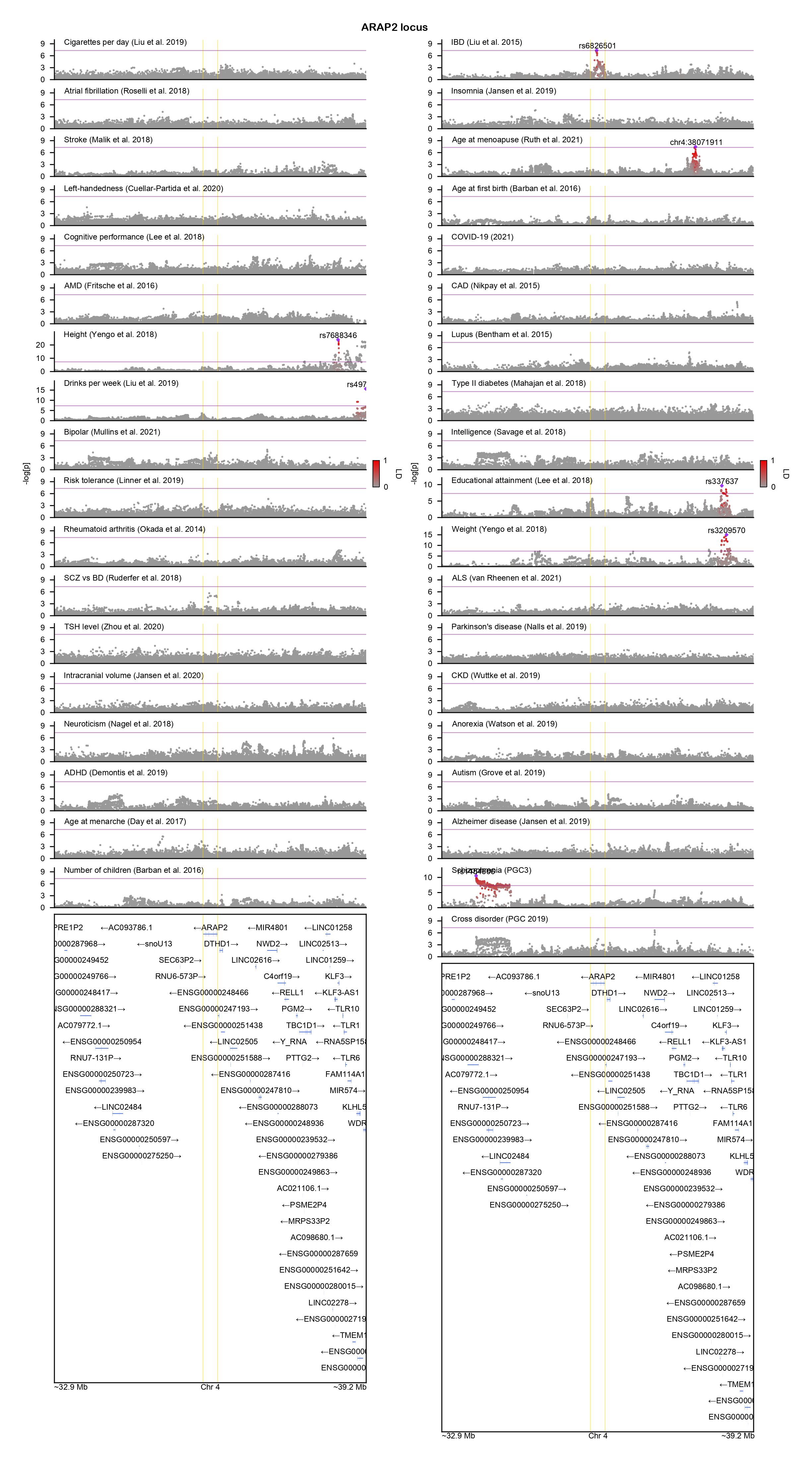

### BCL11A-locuszoom.png

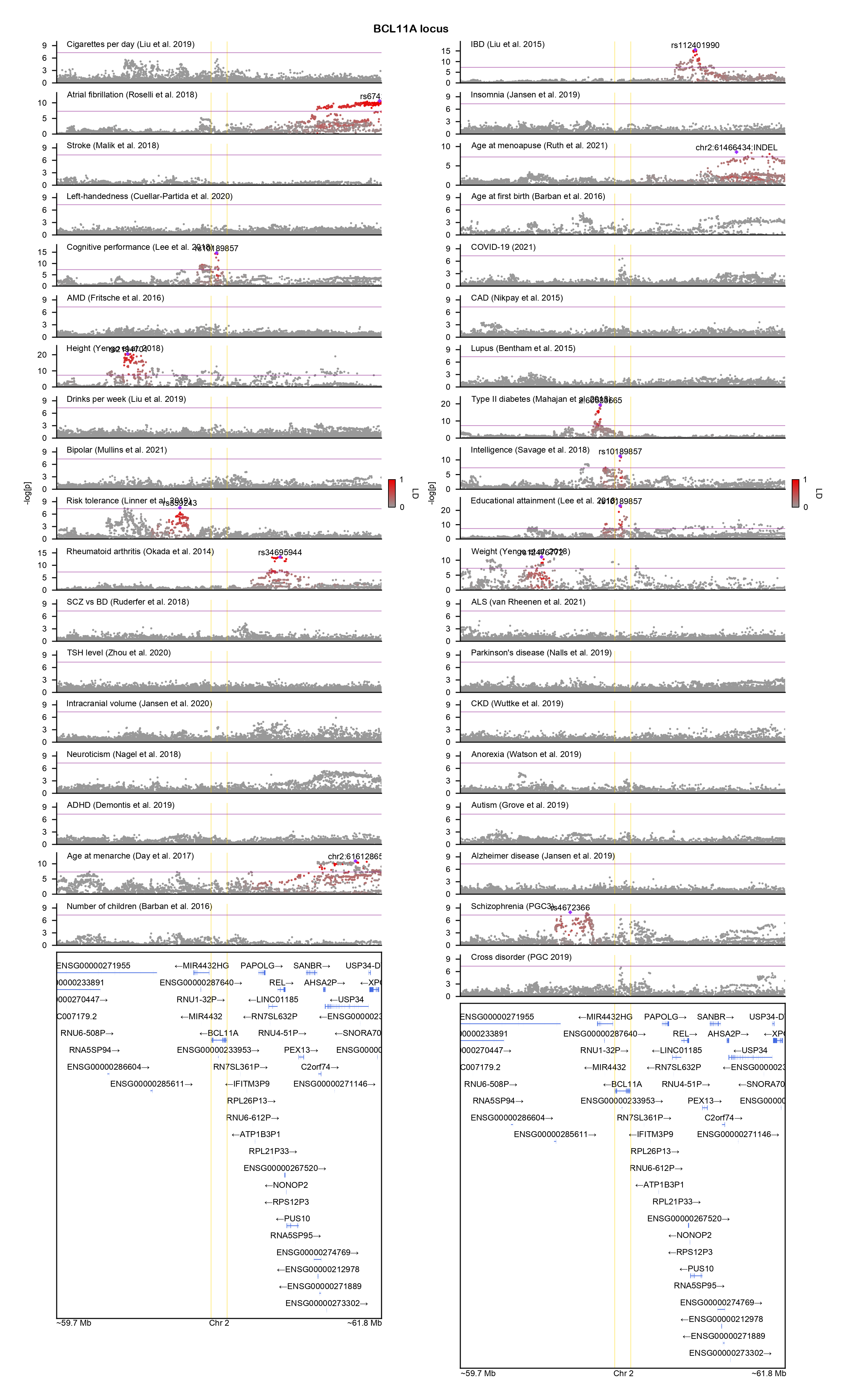

### BCL11B-locuszoom.png

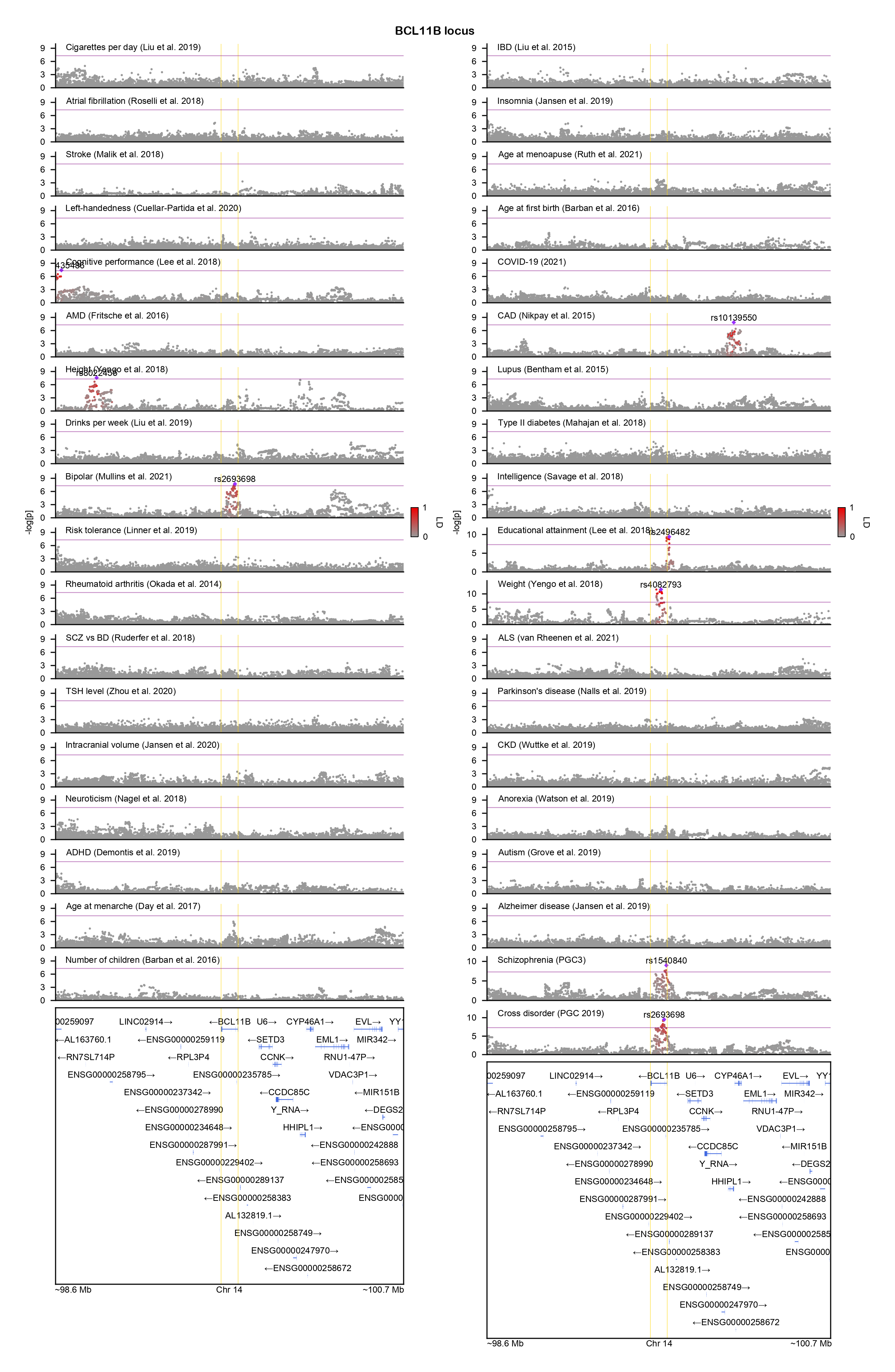

### CACNA1I-locuszoom.png

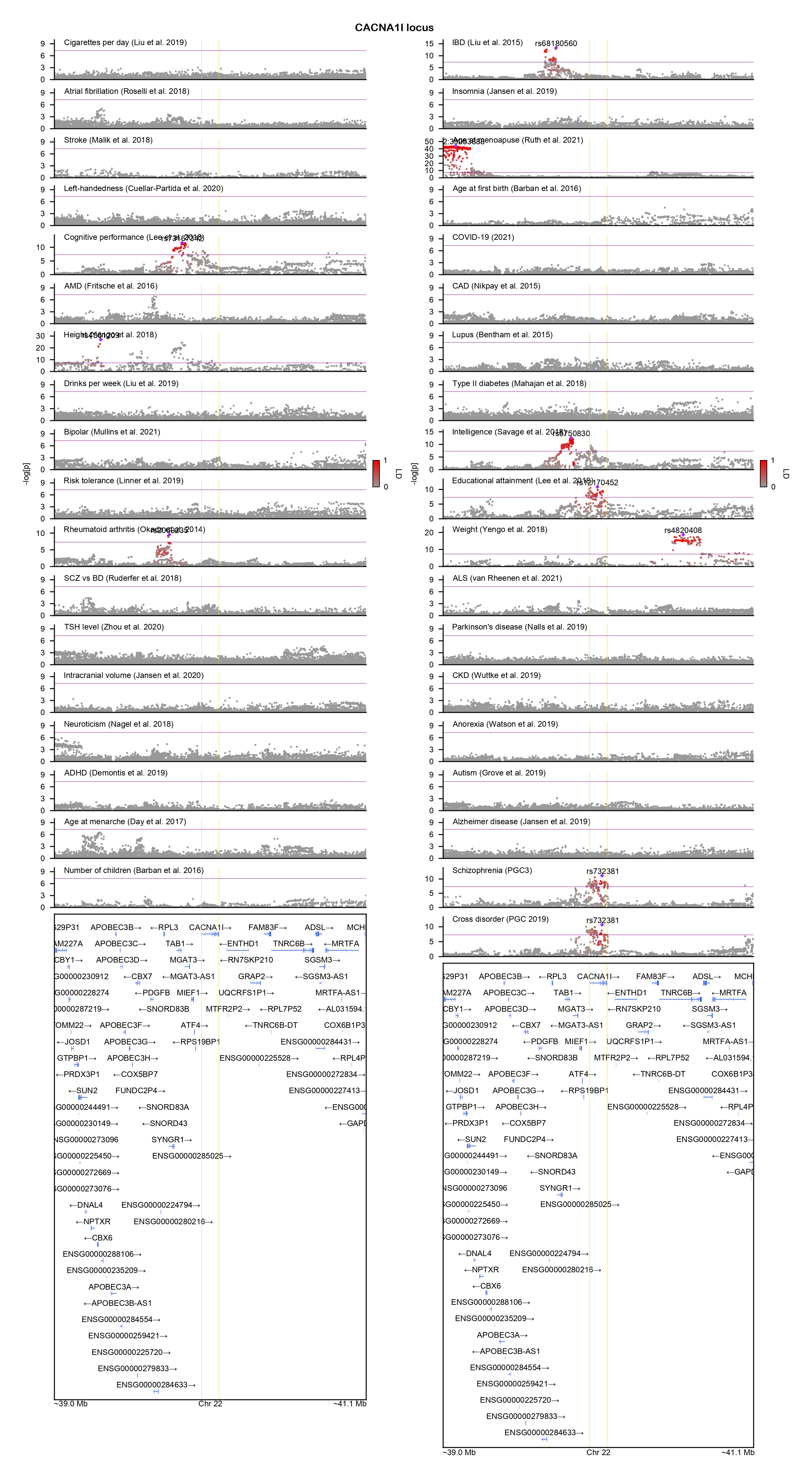

### CACNA2D2-locuszoom.png

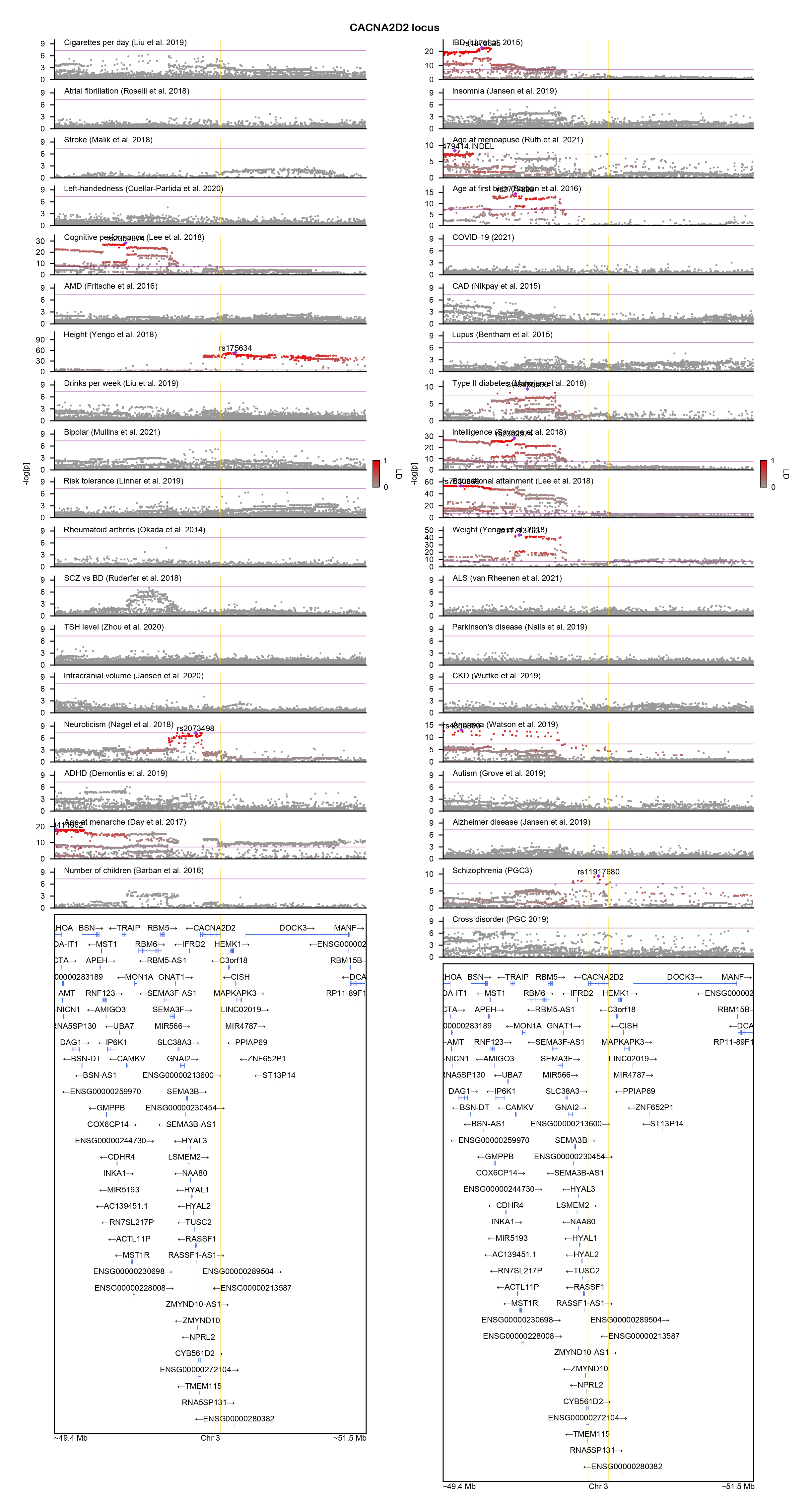

### CBLN2-locuszoom.png

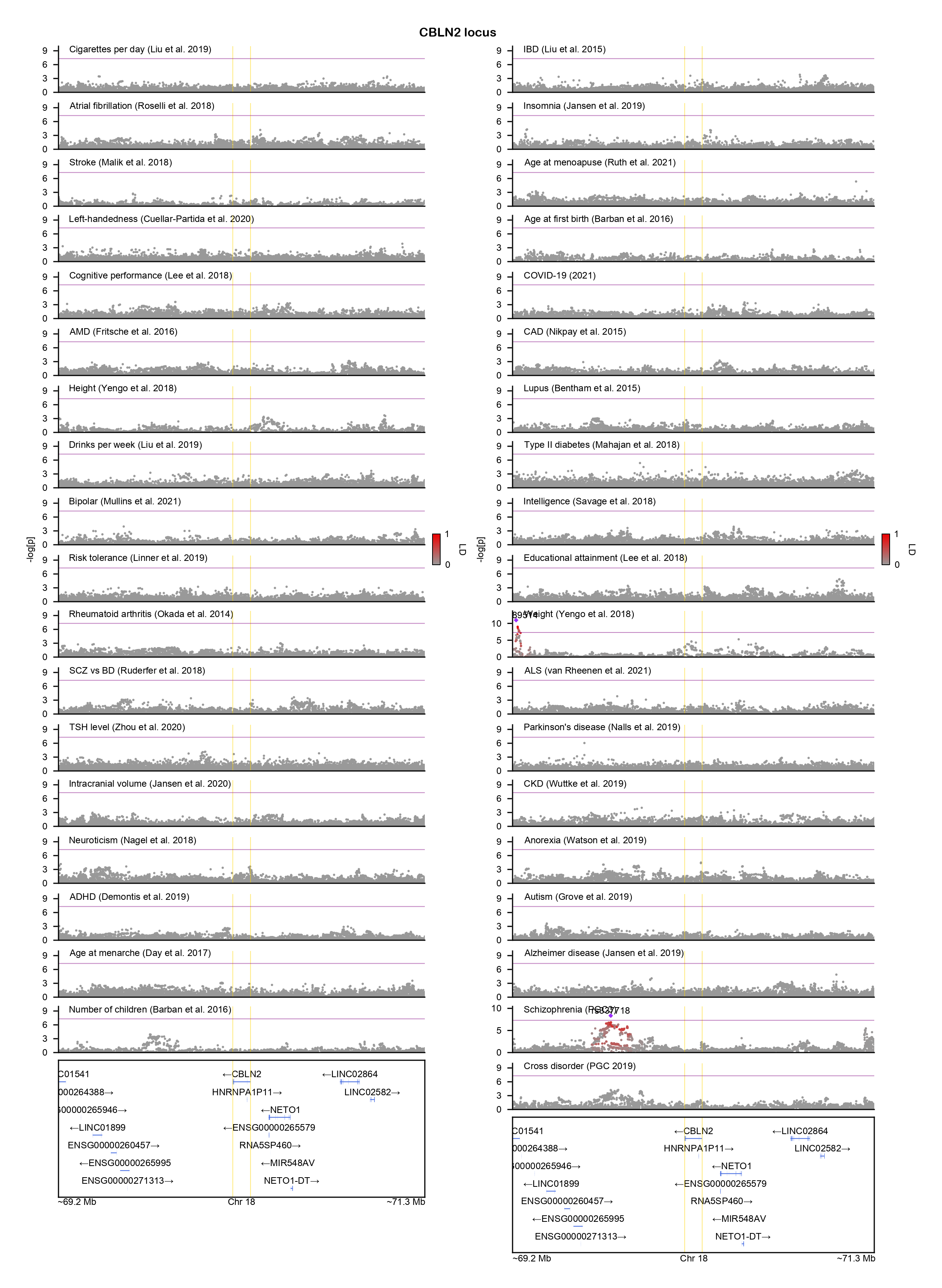

### COL19A1-locuszoom.png

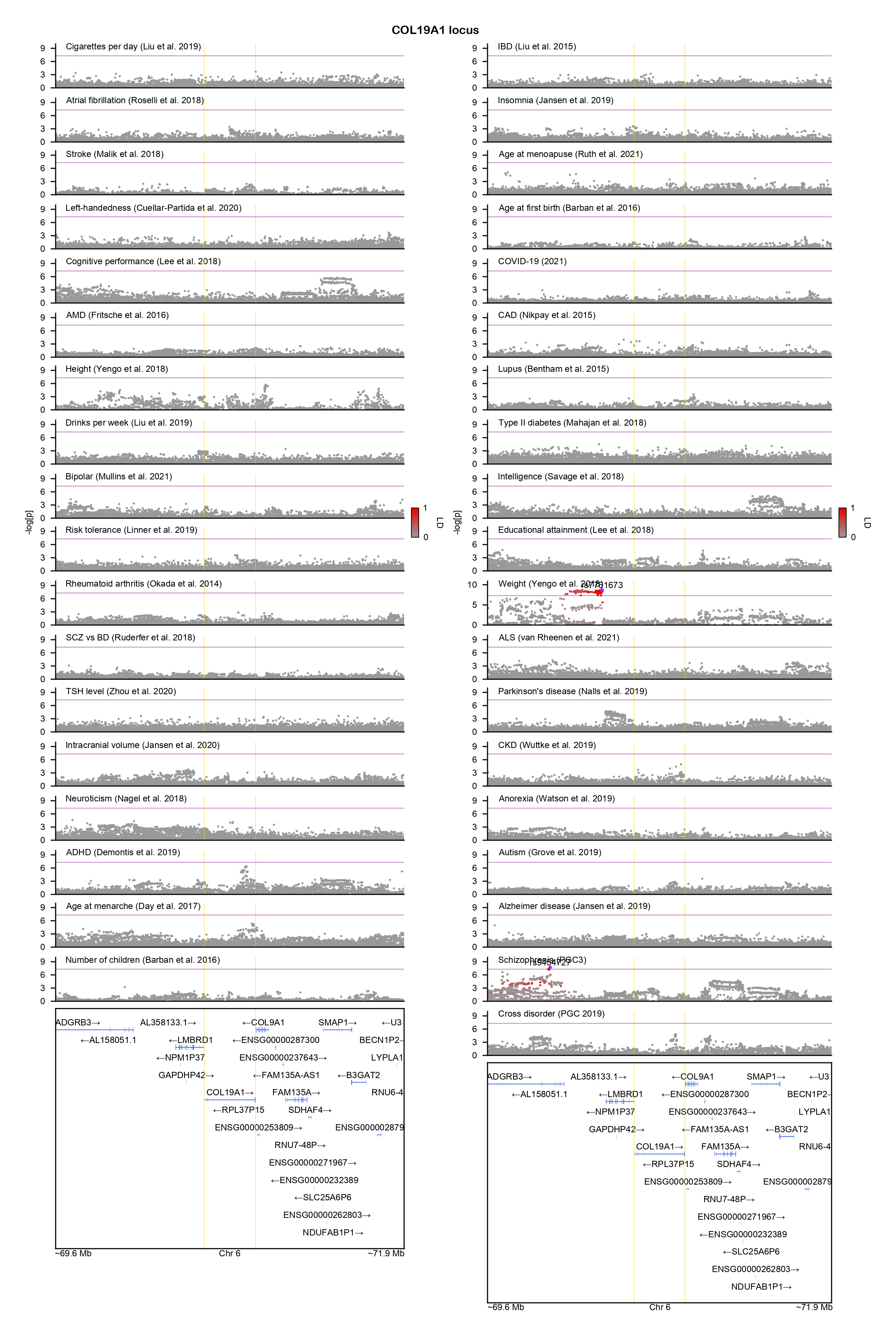

### CPNE8-locuszoom.png

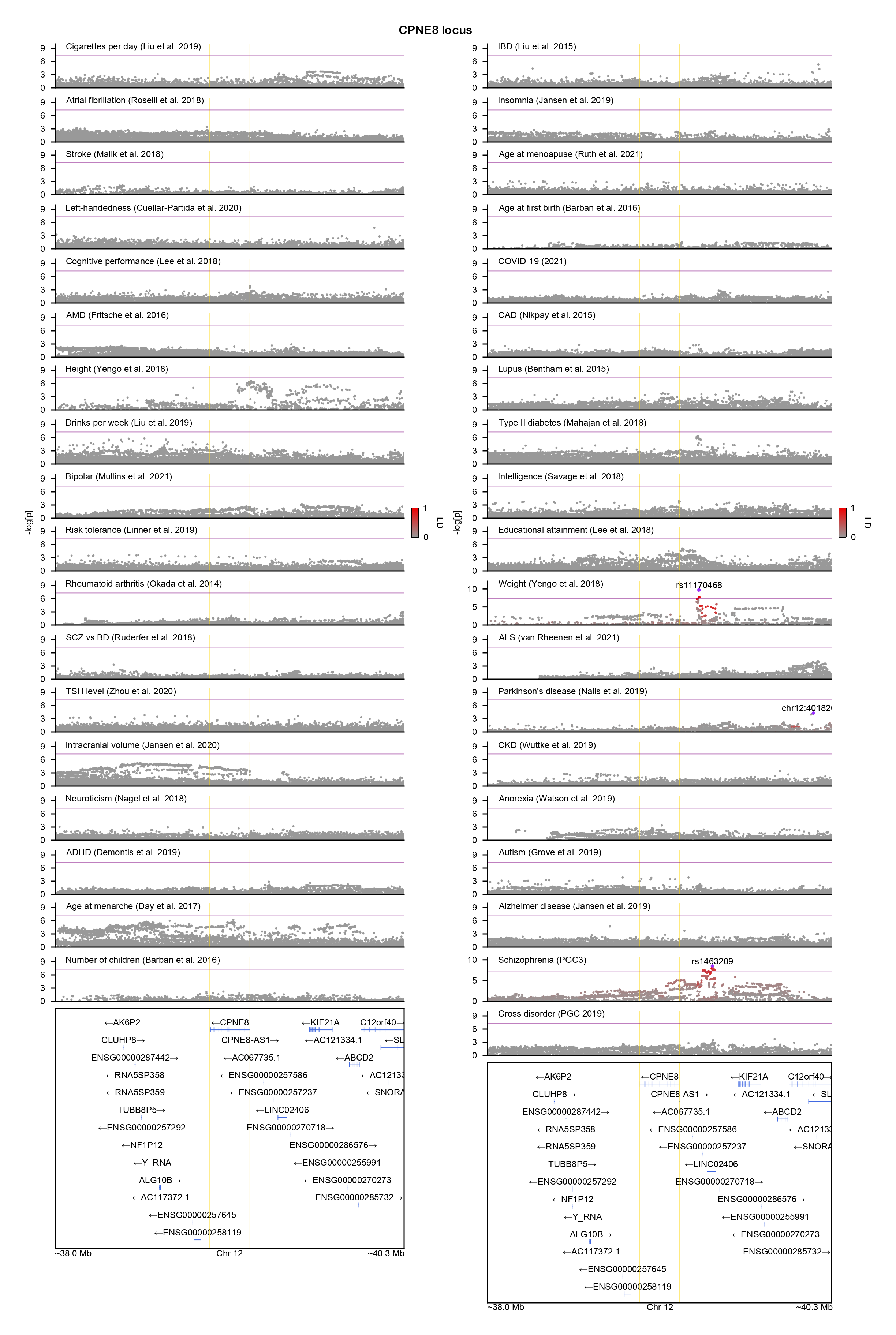

### CSDE1-locuszoom.png

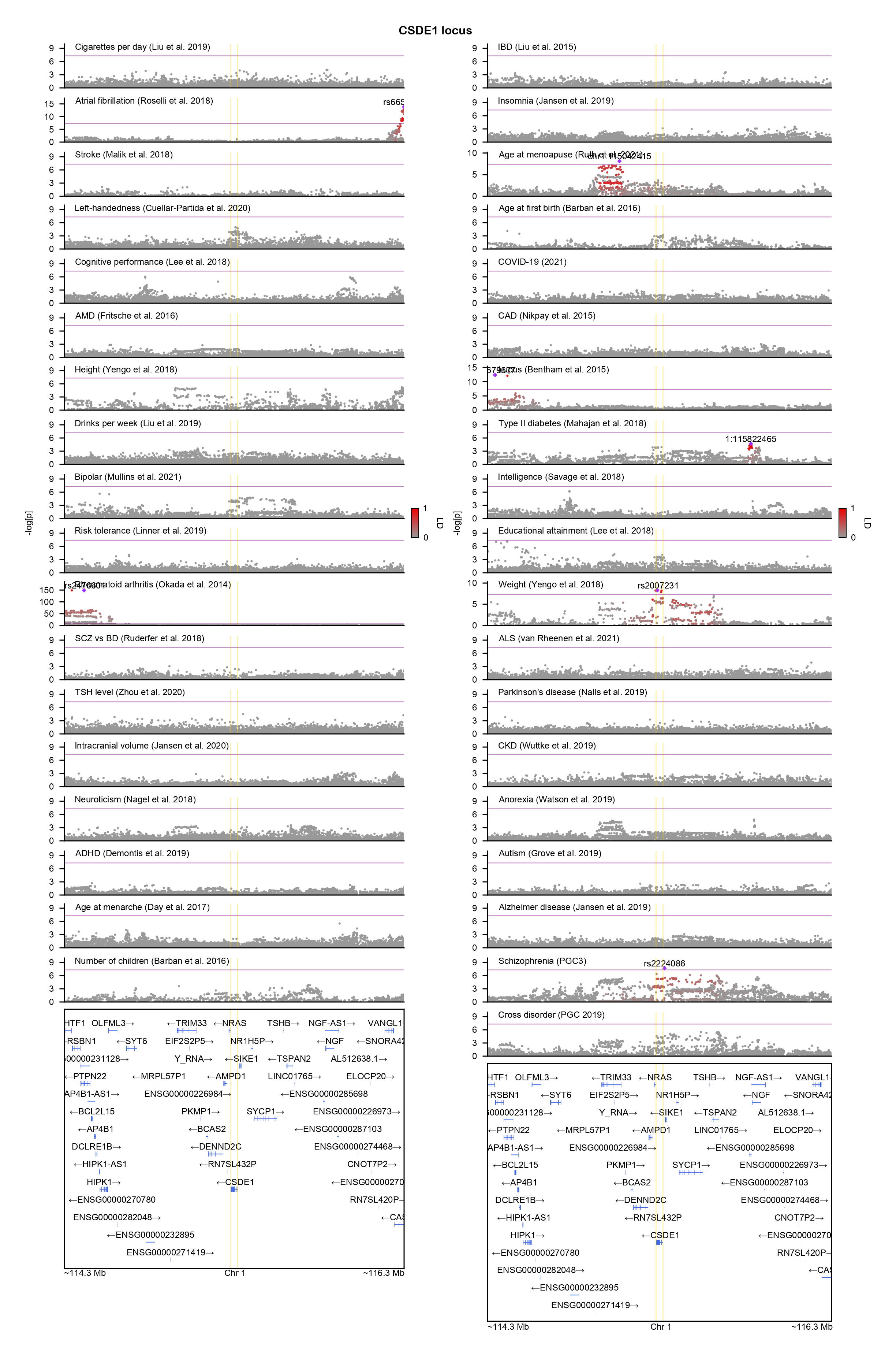

### CSMD1-locuszoom.png

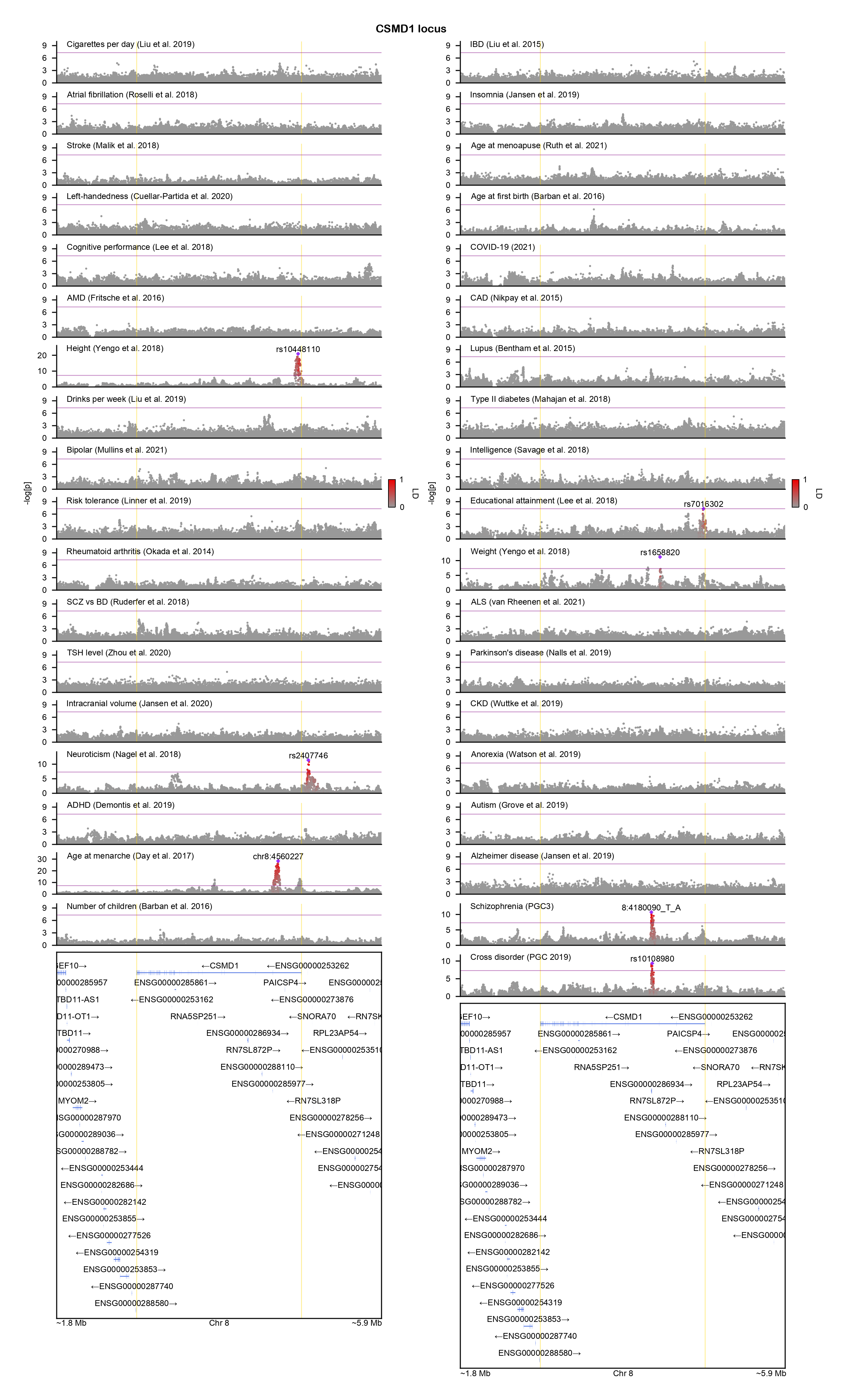

### CYP7B1-locuszoom.png

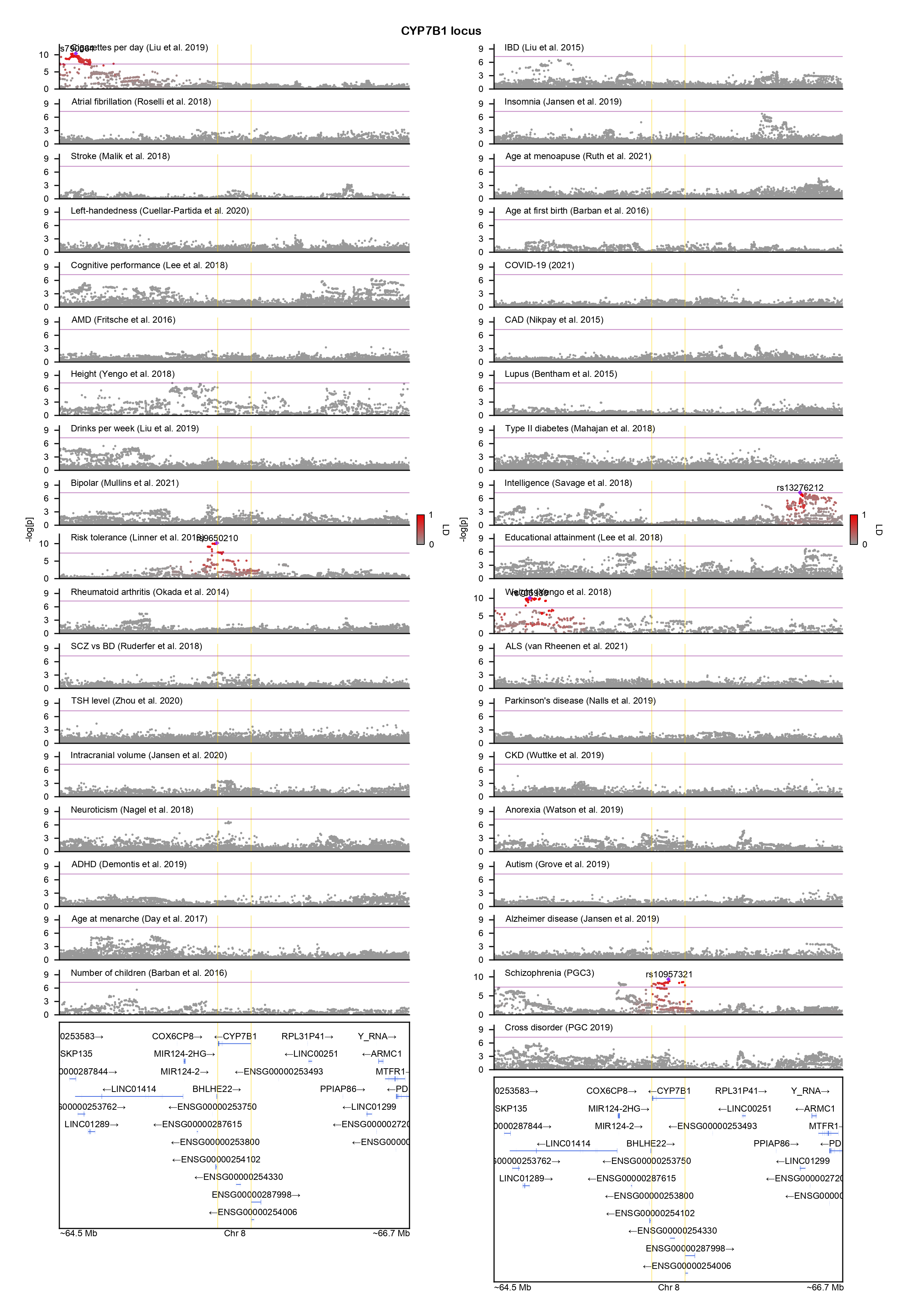

### DCP1B-locuszoom.png

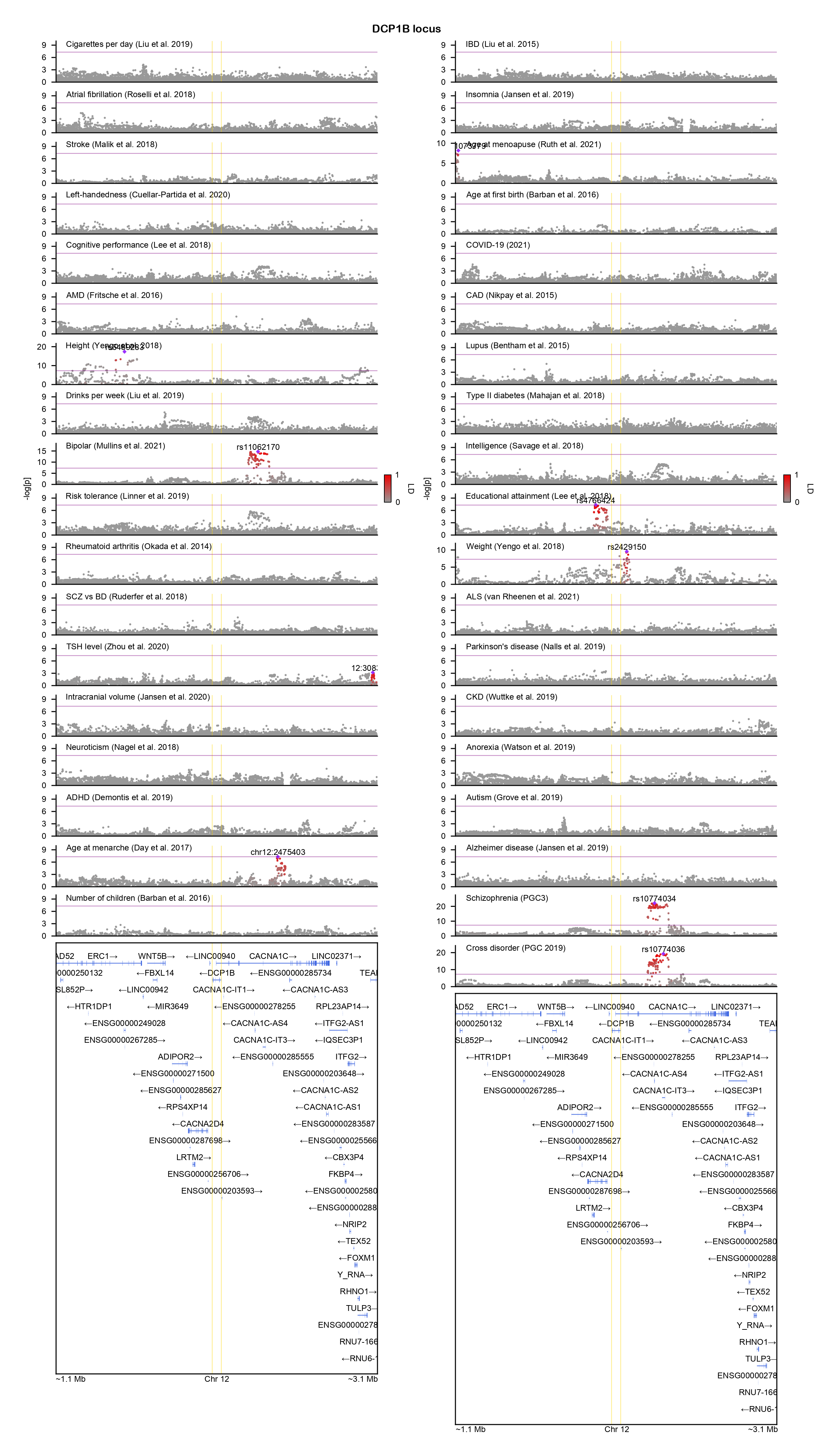

### DGKZ-locuszoom.png

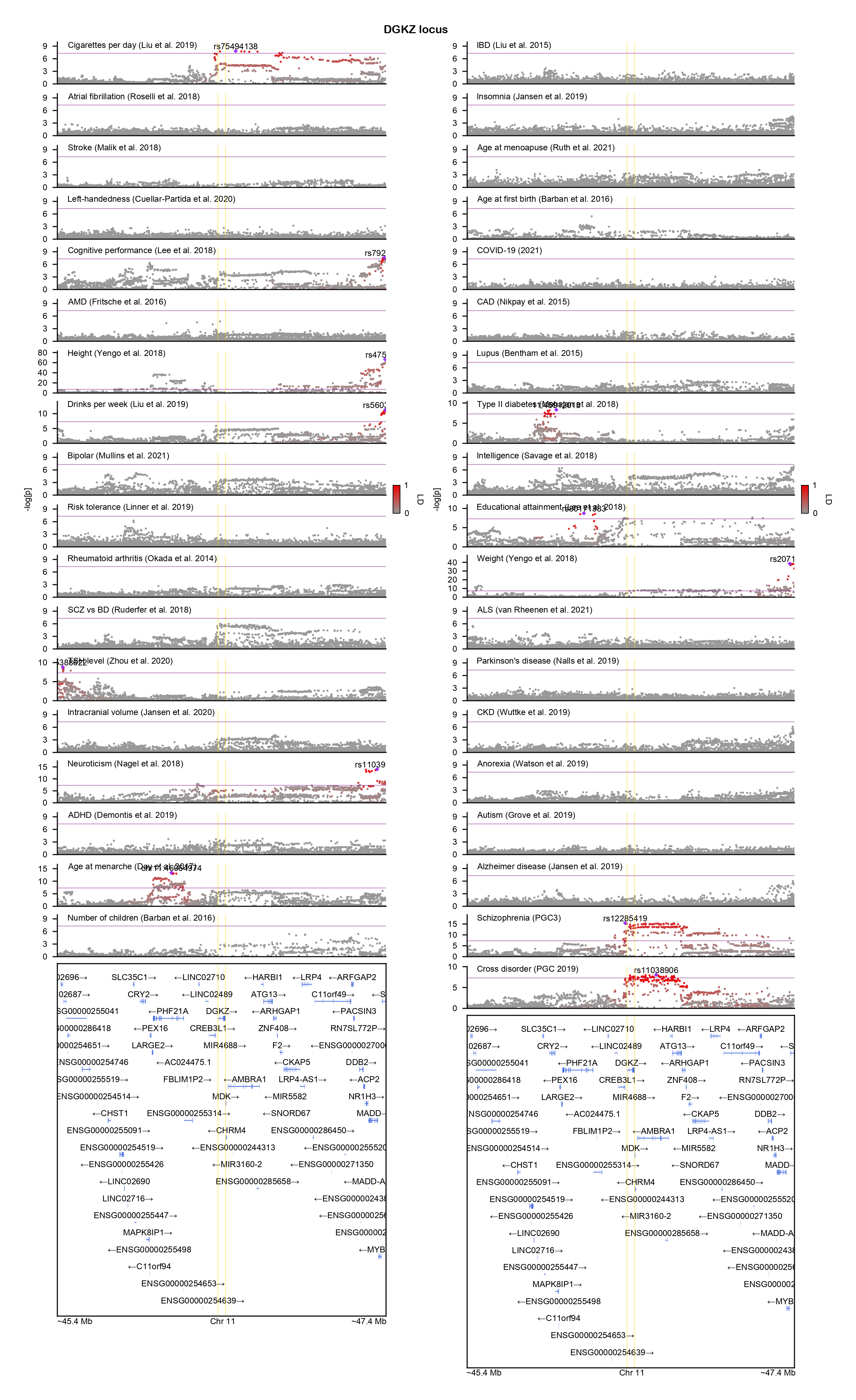

### DNAH11-locuszoom.png

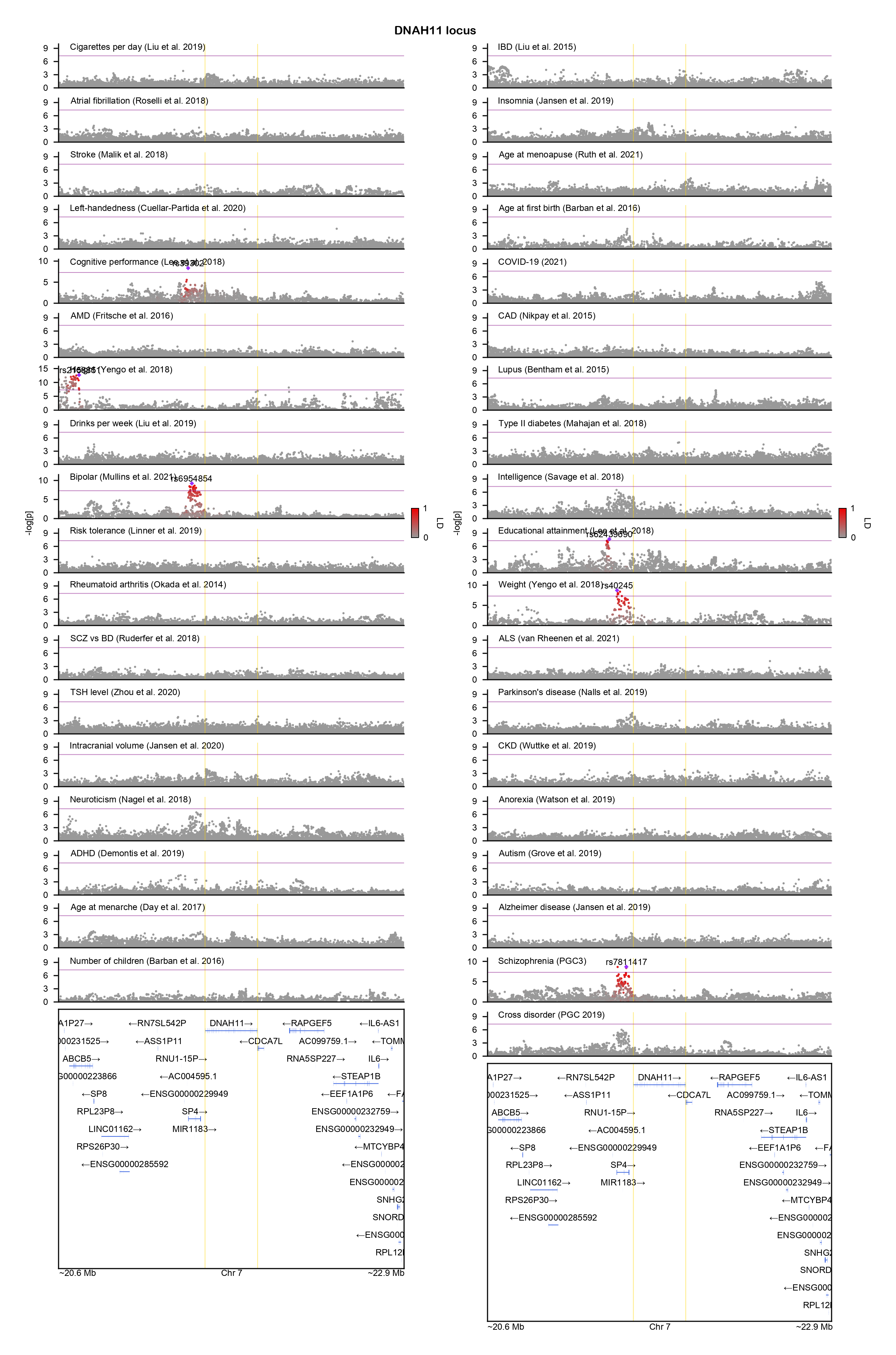

### DNMT3A-locuszoom.png

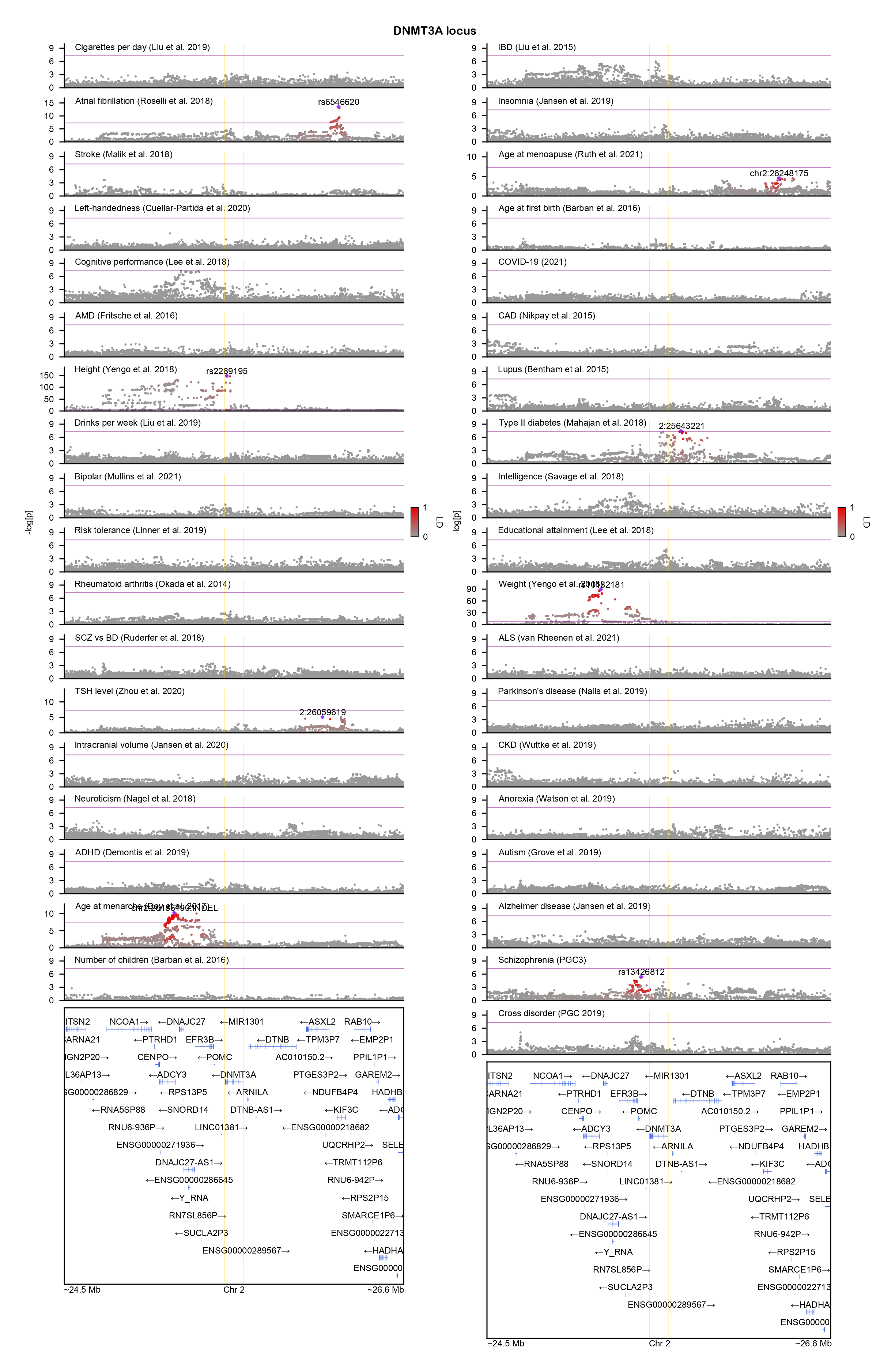

### DPP4-locuszoom.png

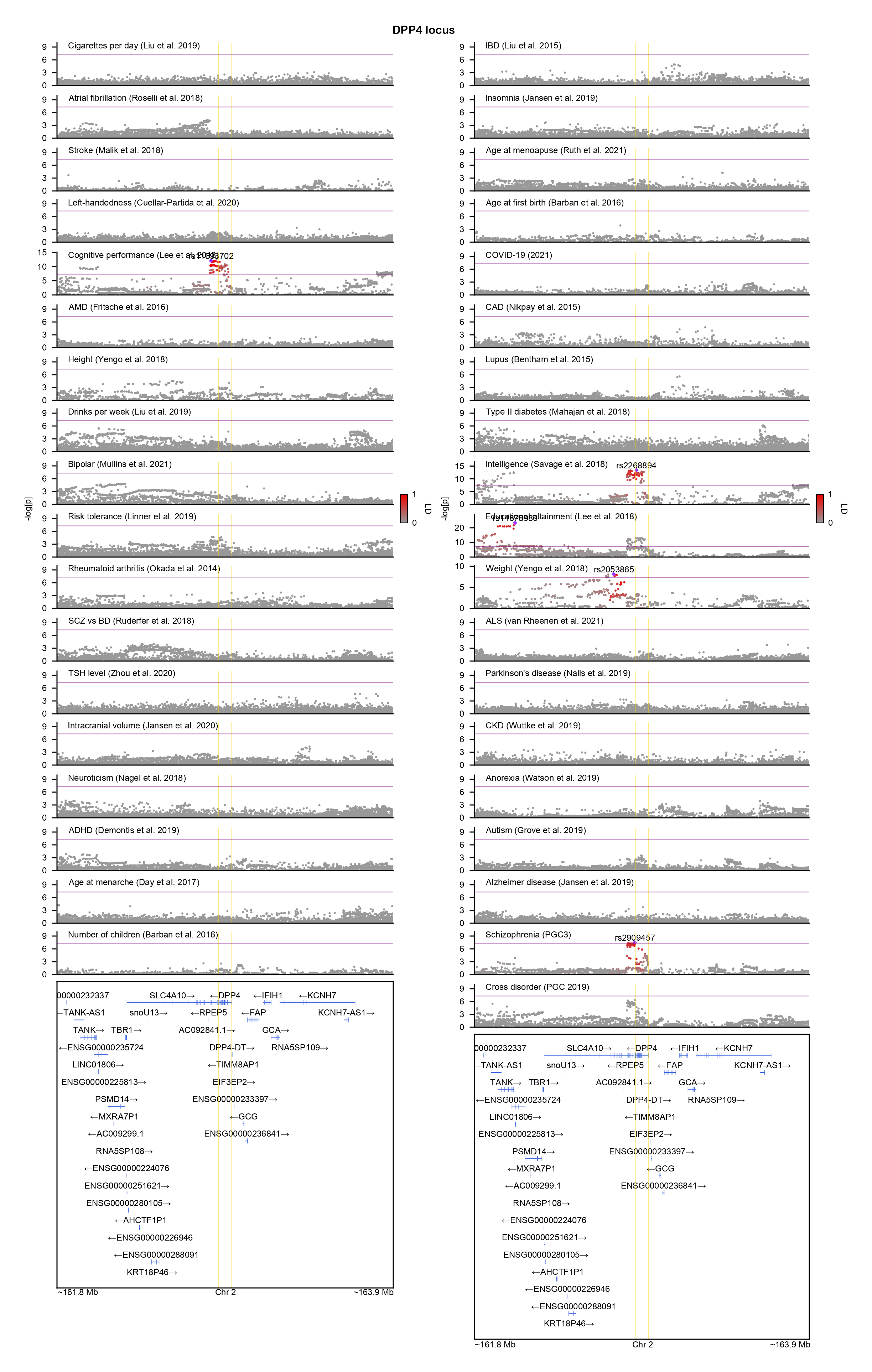

### DUSP26-locuszoom.png

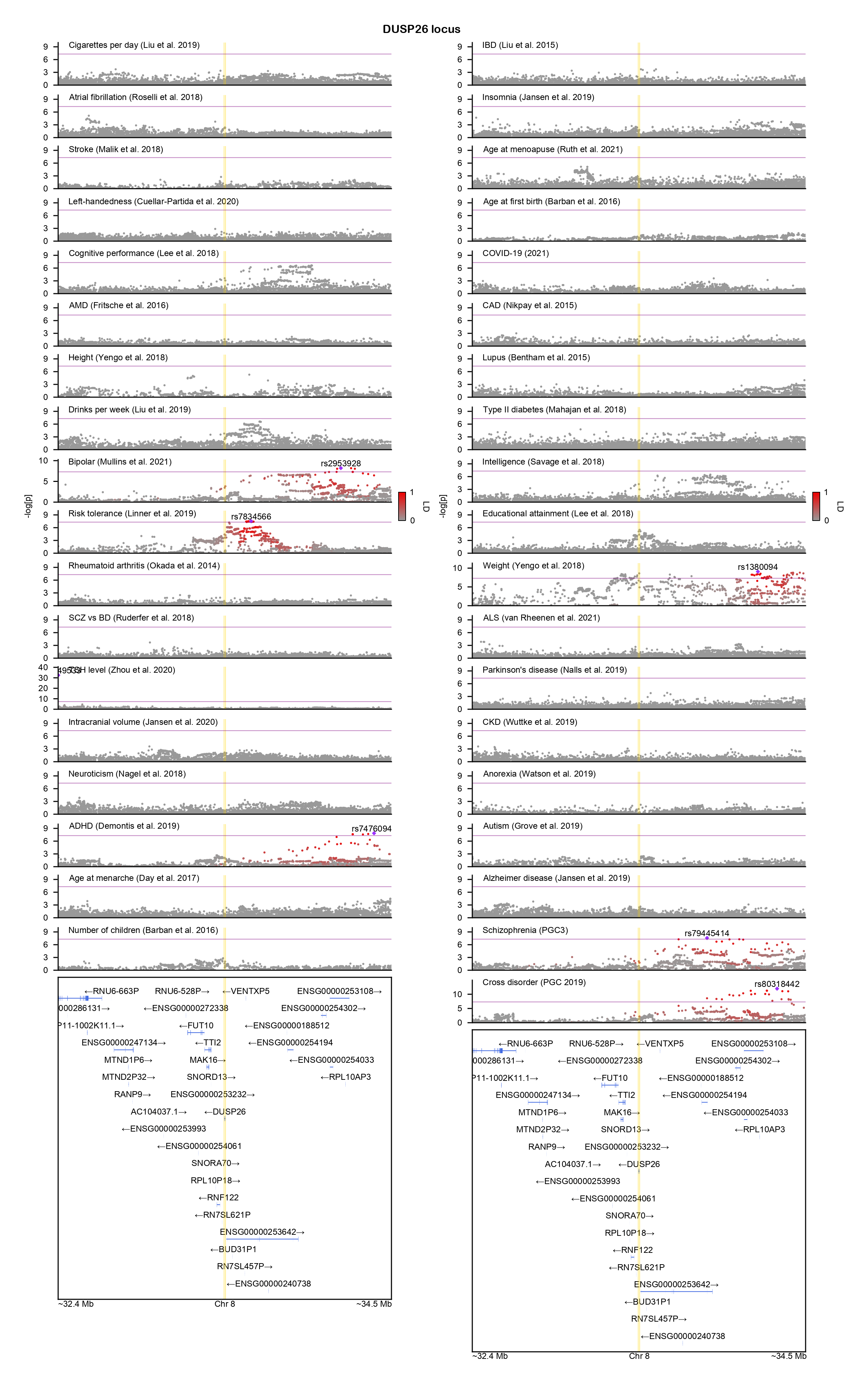

### ERICH1-locuszoom.png

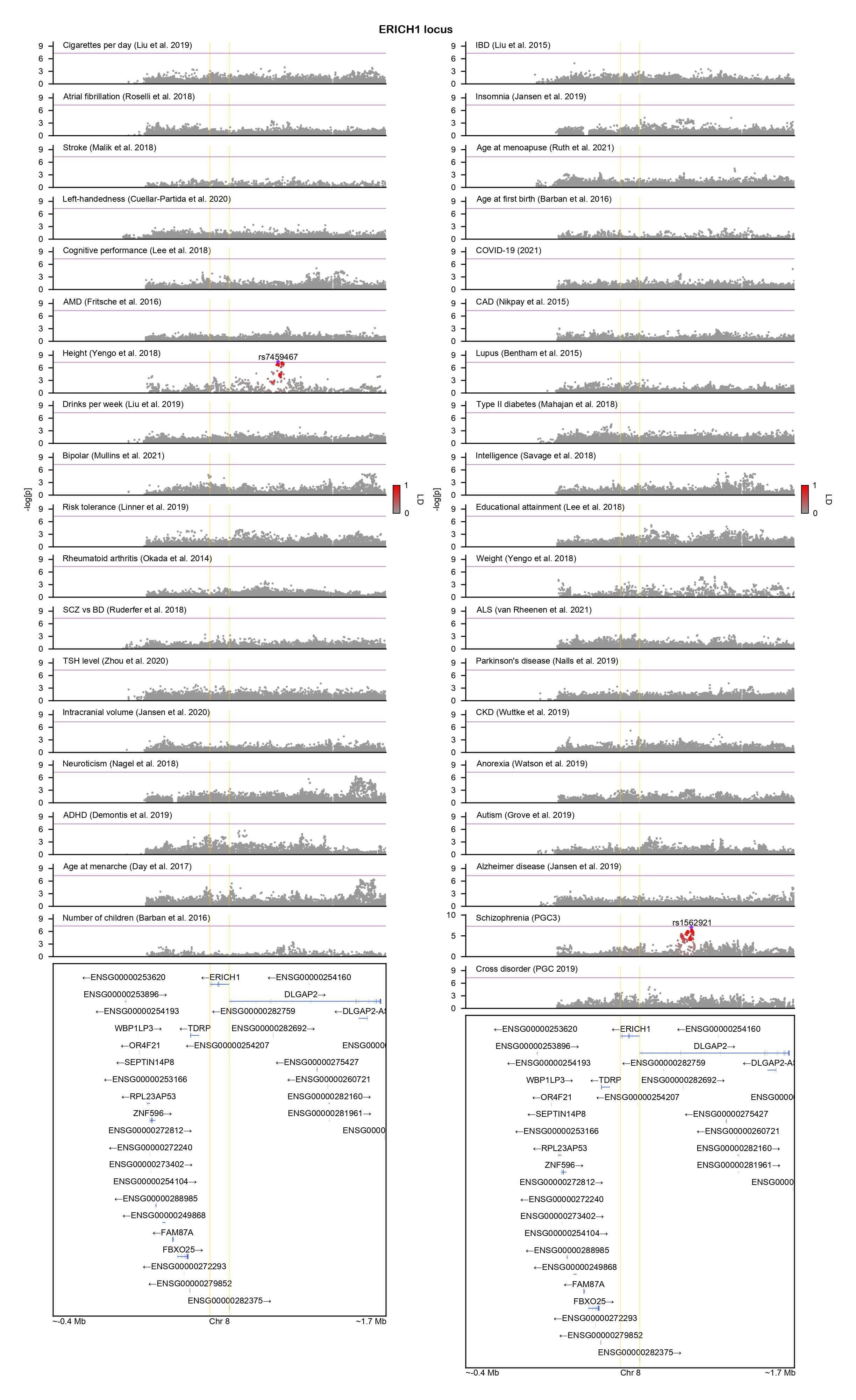

### EXOC4-locuszoom.png

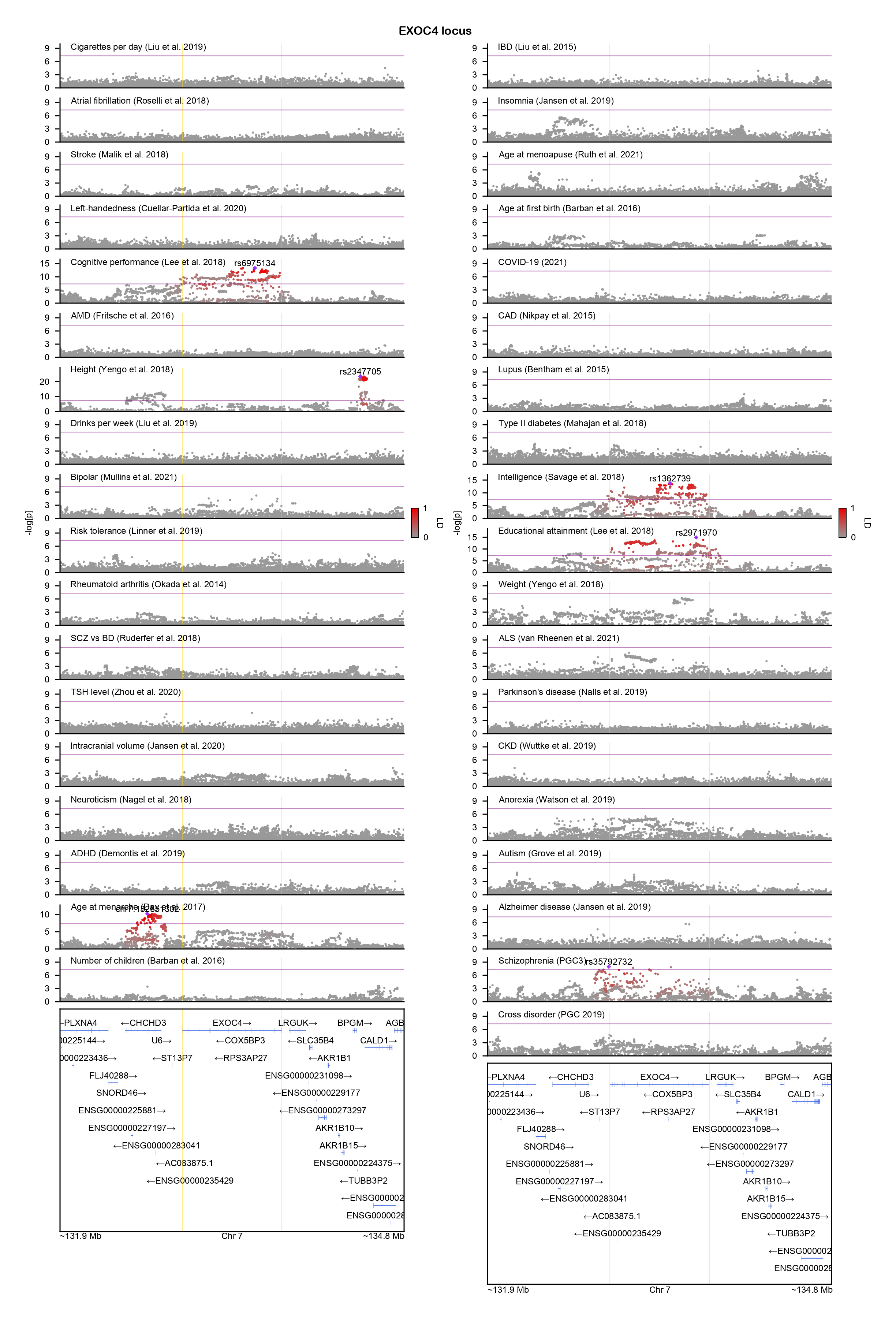

### FAM181B-locuszoom.png

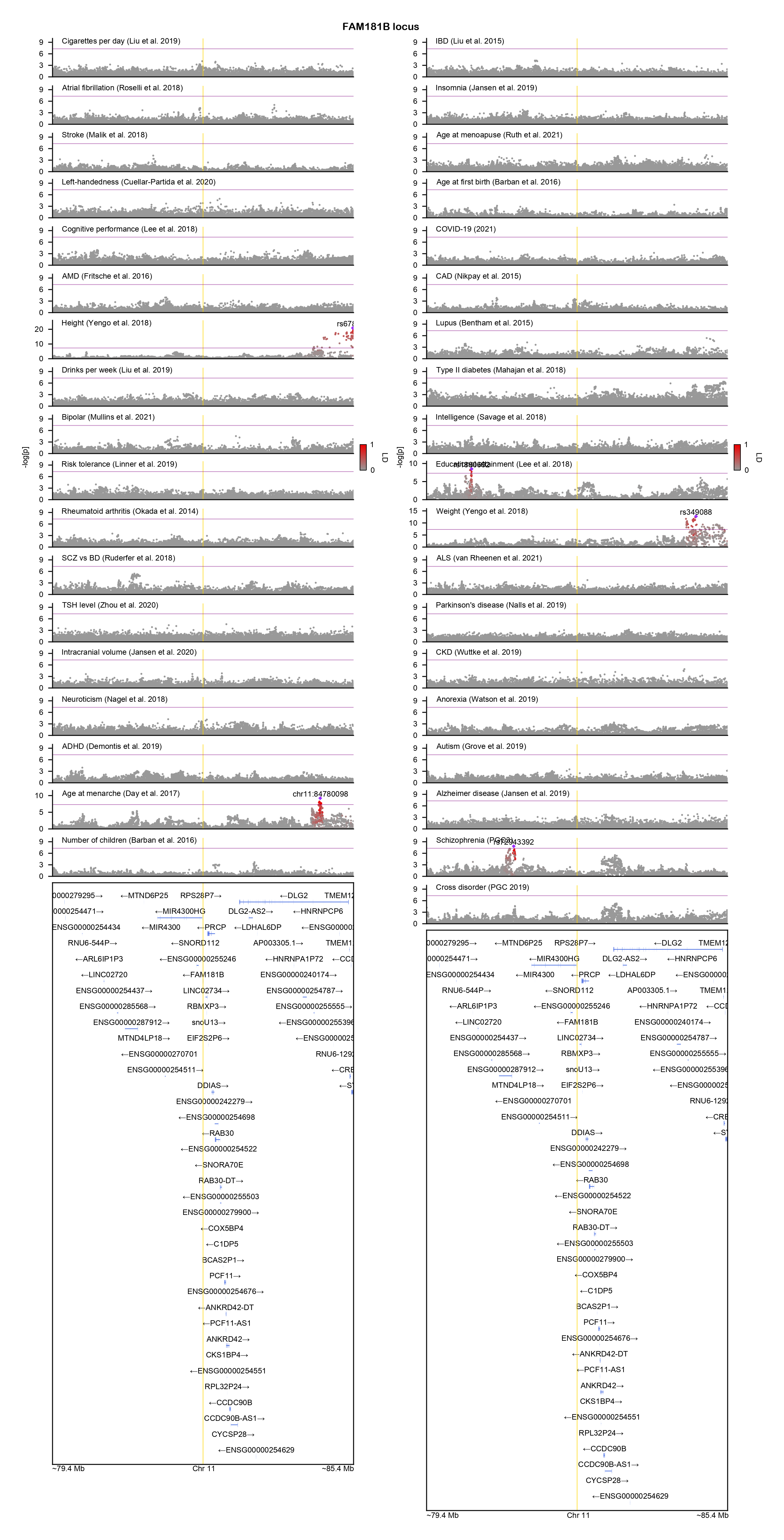

### FLRT2-locuszoom.png

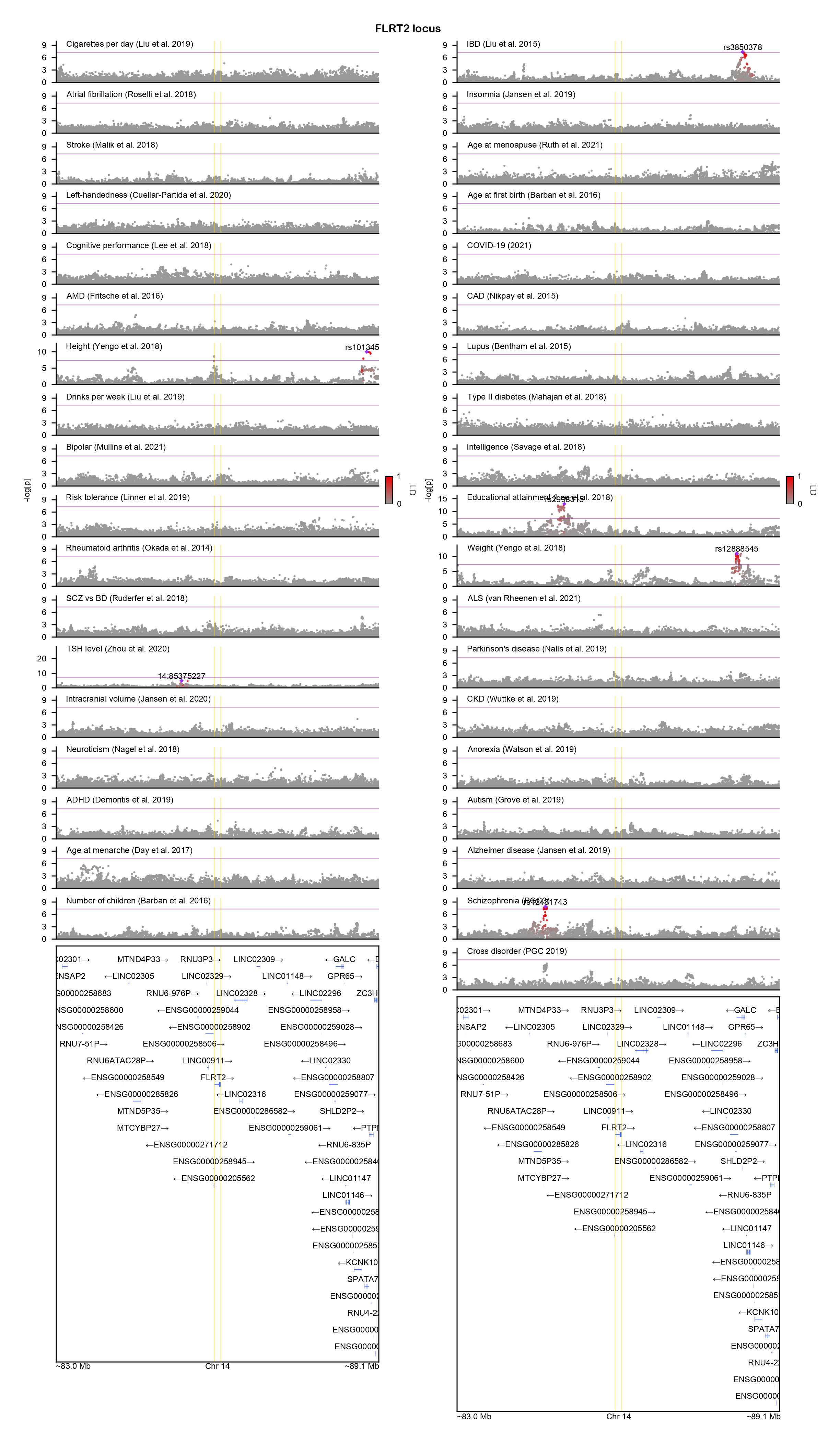

### FOXO3-locuszoom.png

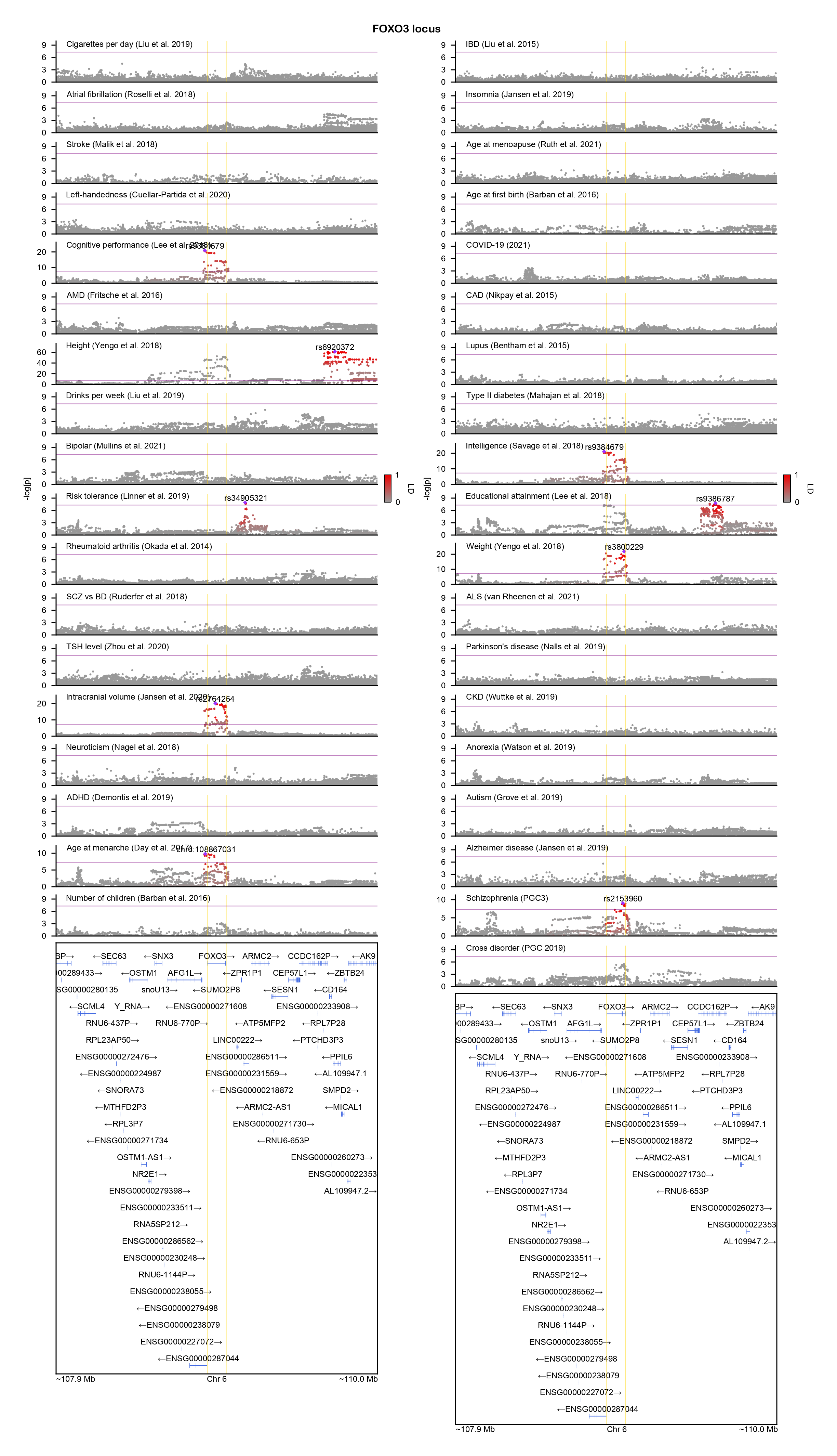

### GALNT2-locuszoom.png

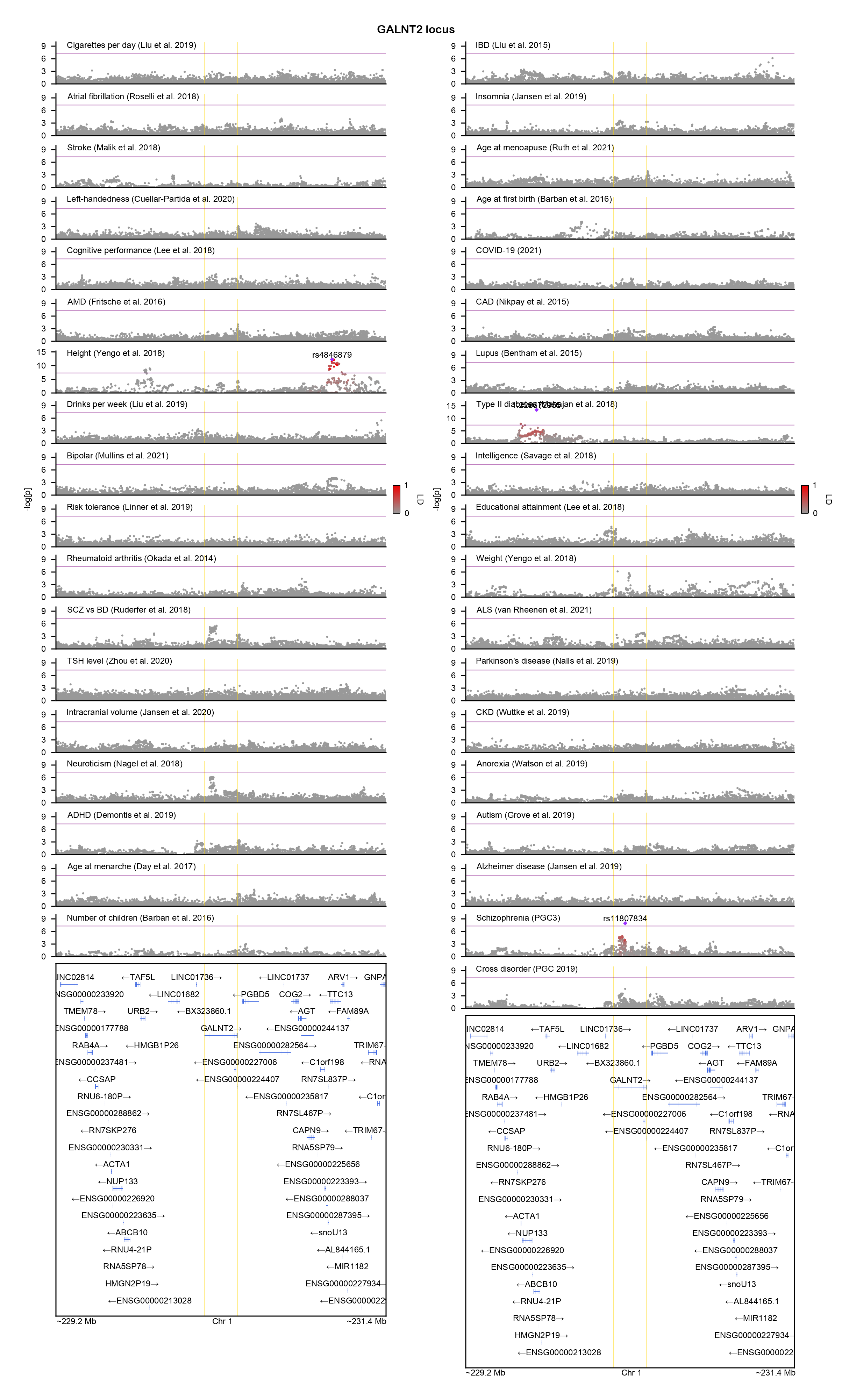

### GARNL3-locuszoom.png

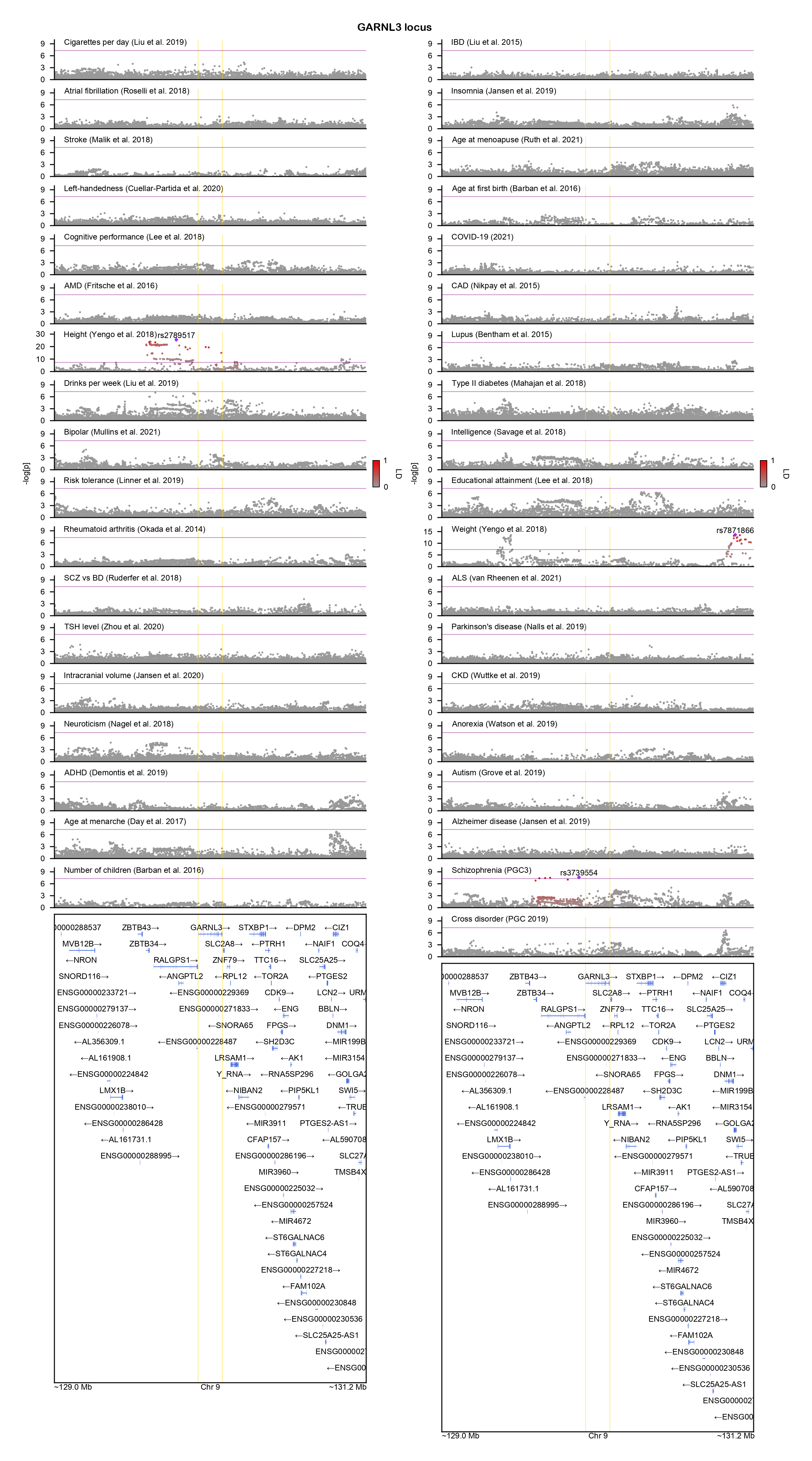

### GNAI1-locuszoom.png

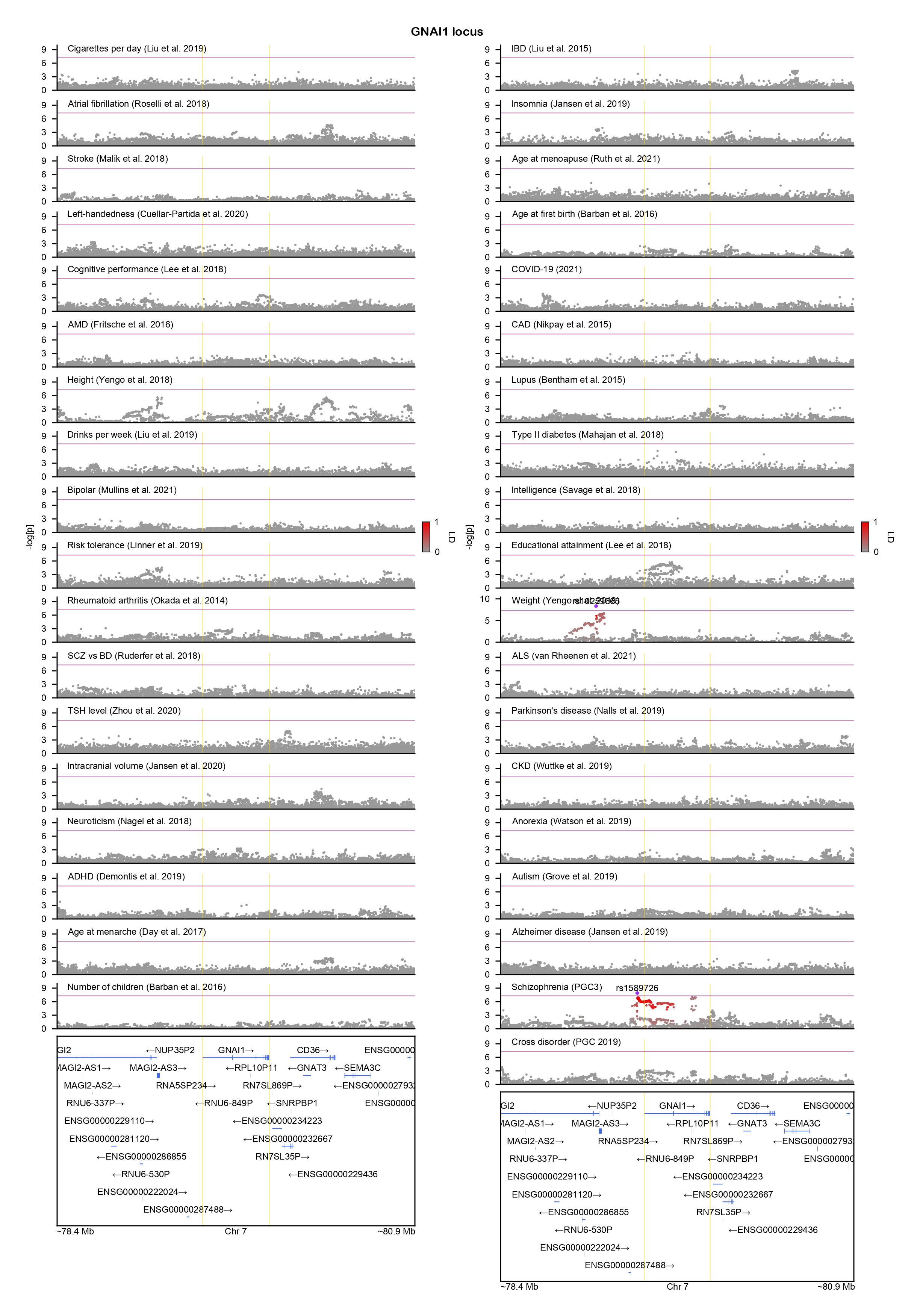

### GRIN2A-locuszoom.png

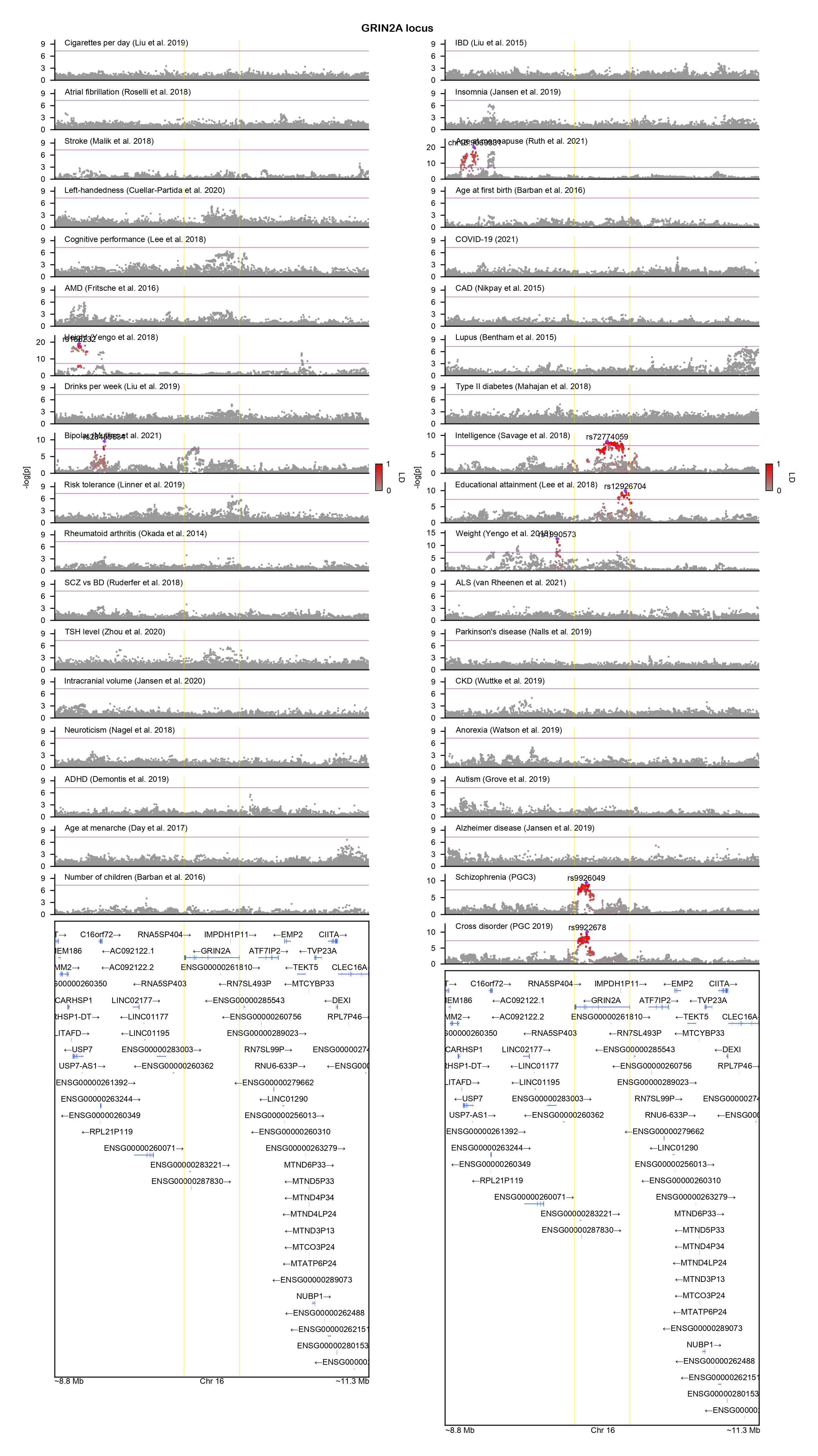

### HLA-G-locuszoom.png

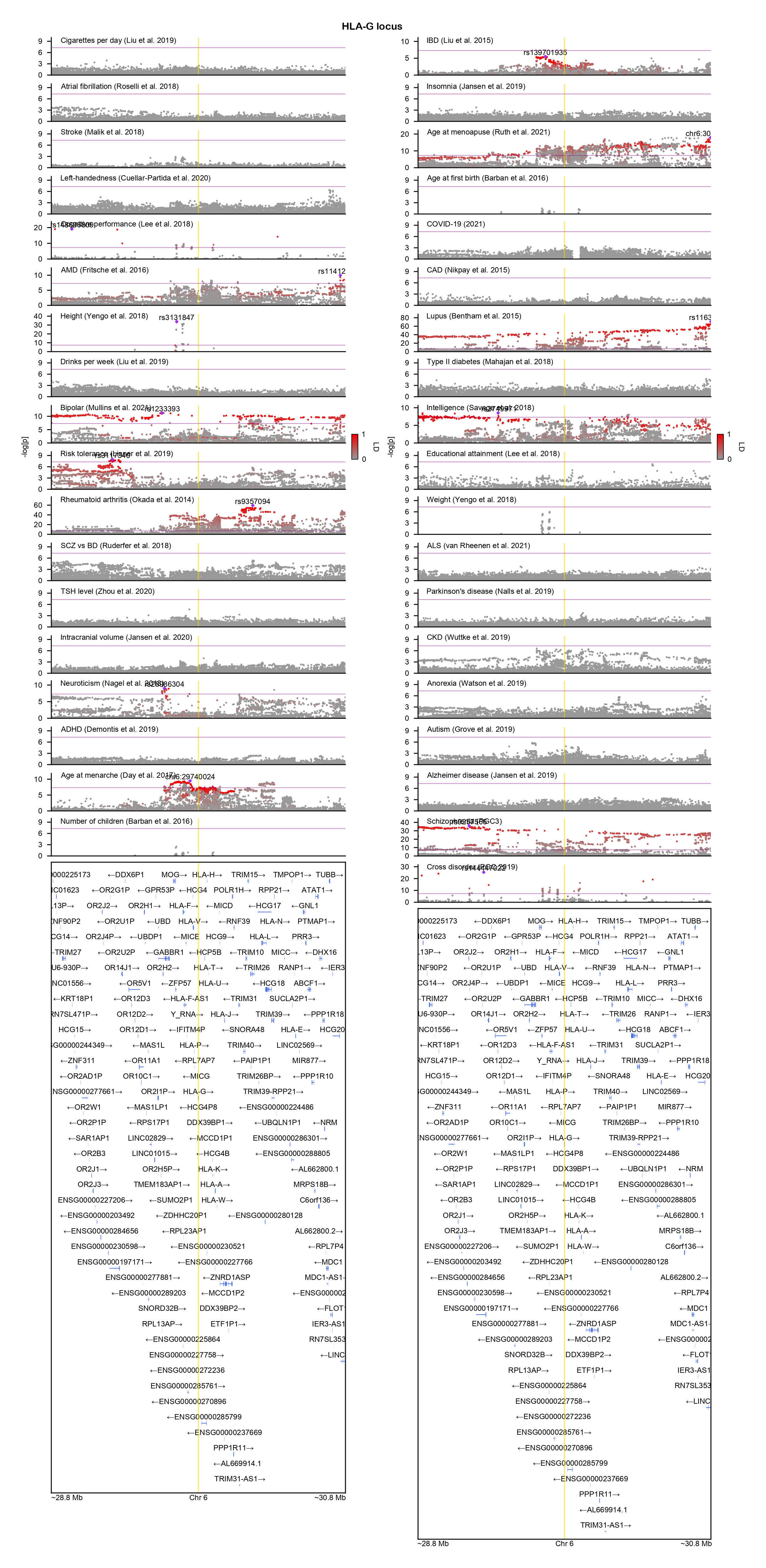

### KCNB1-locuszoom.png

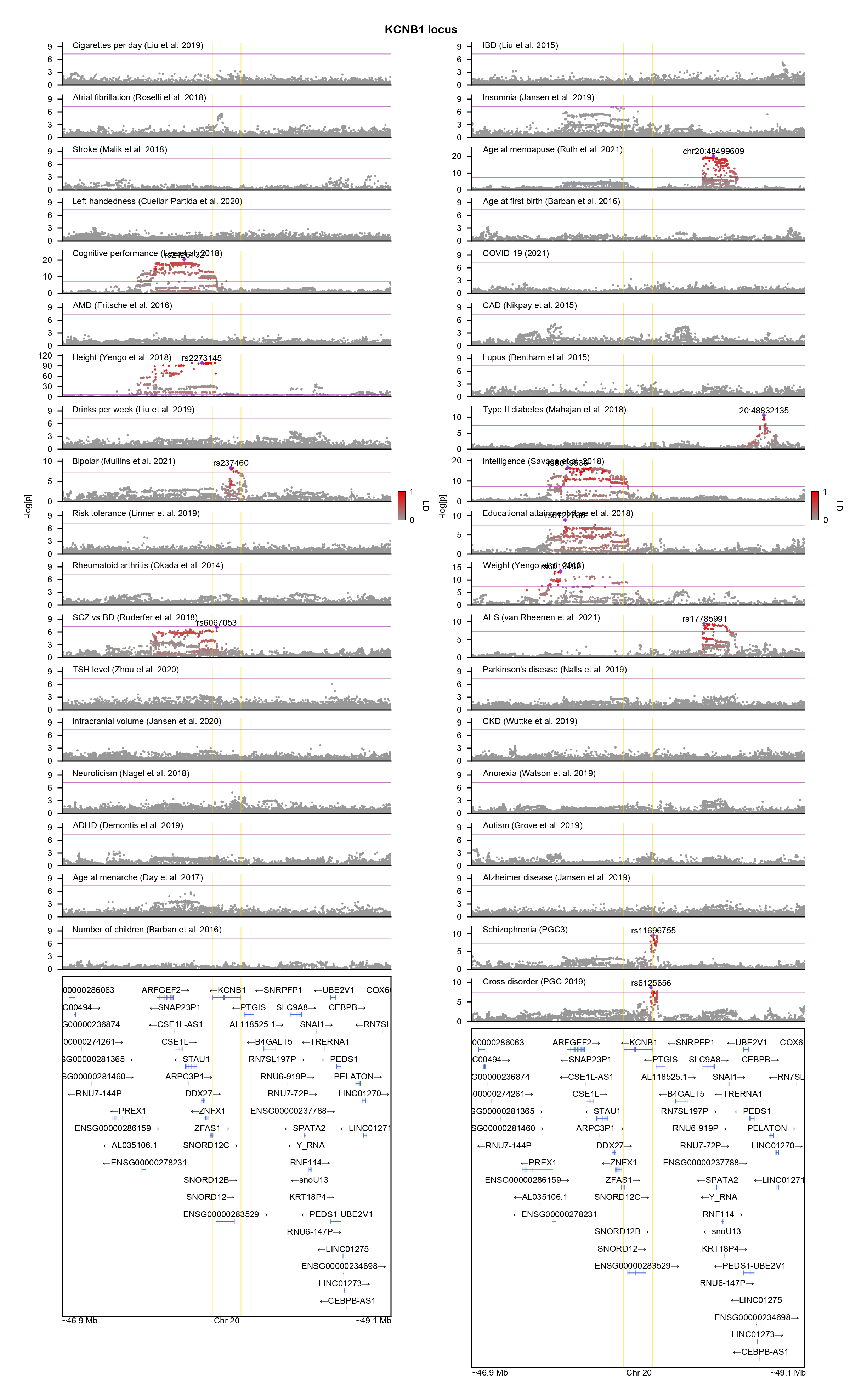

### KIAA1549-locuszoom.png

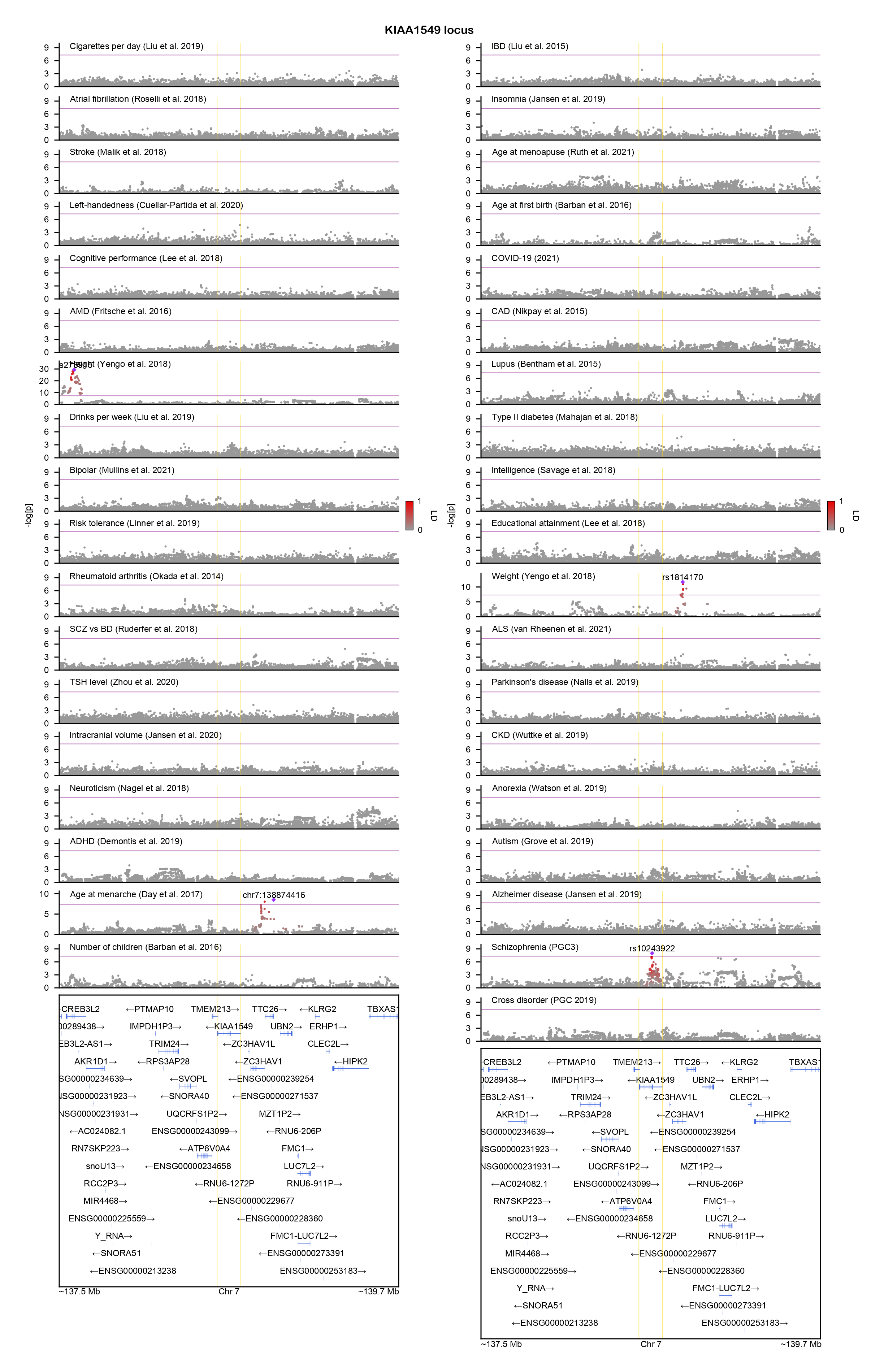

### KLF6-locuszoom.png

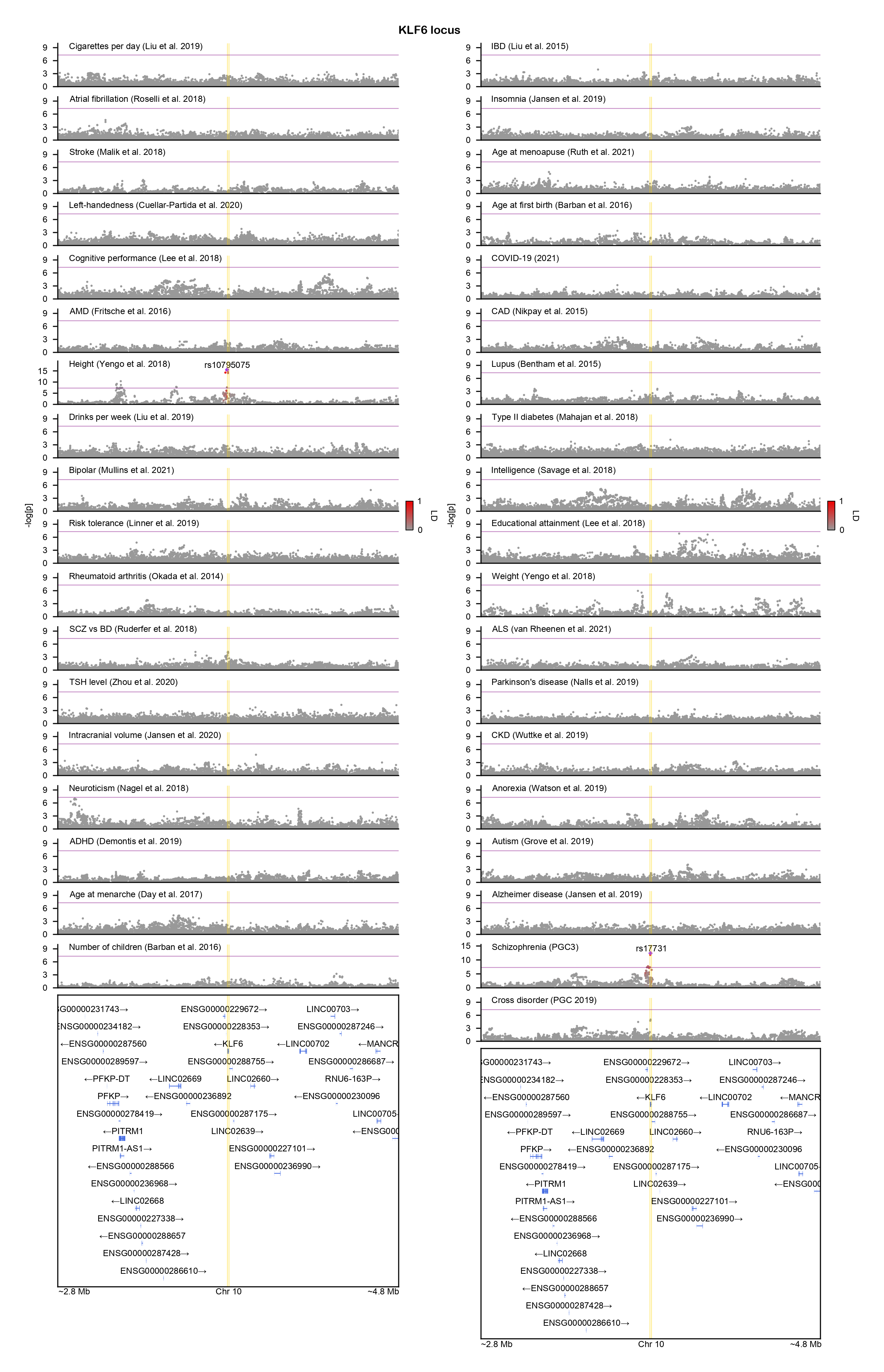

### KLF10-locuszoom.png

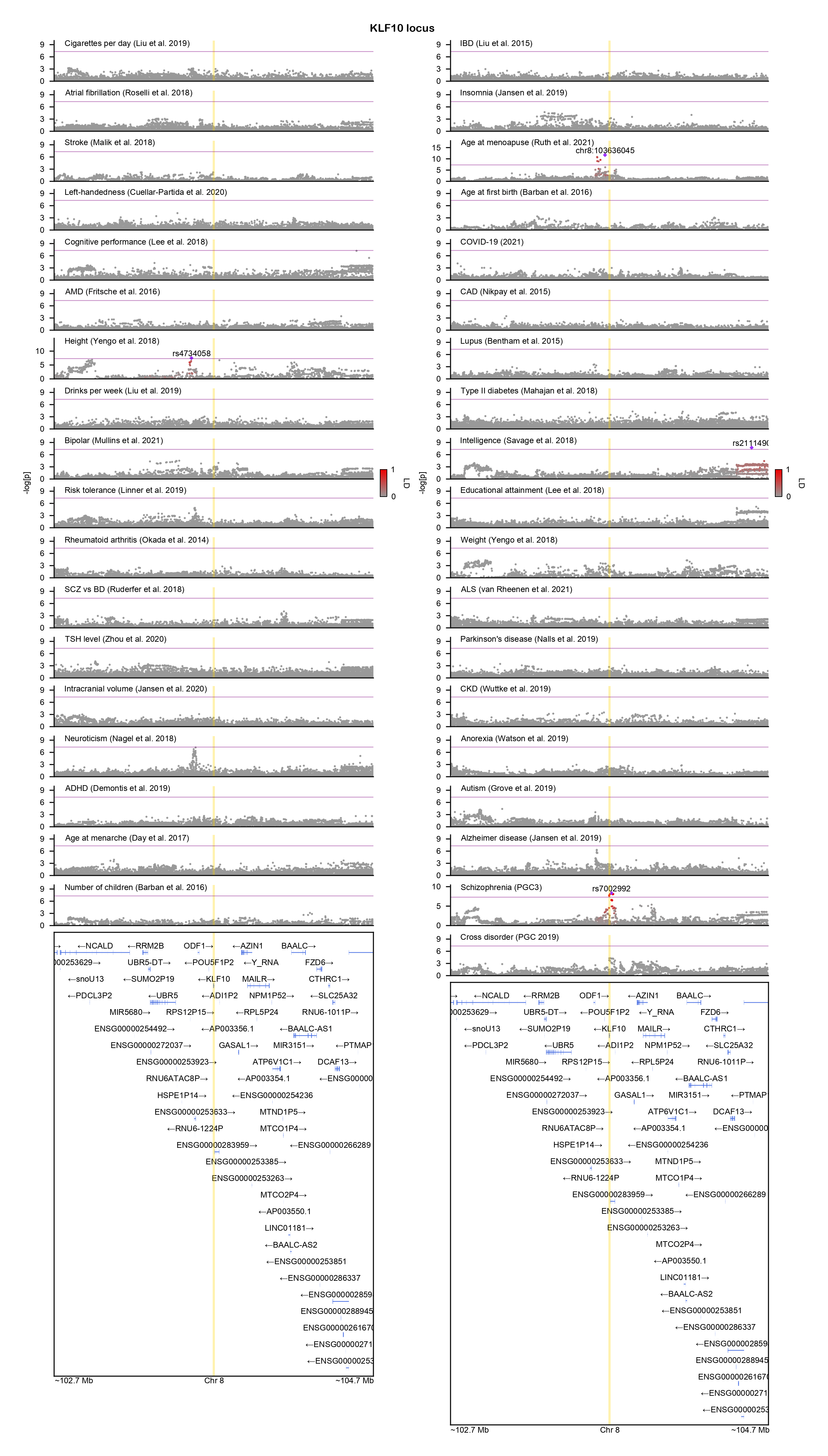

### KMT2E-locuszoom.png

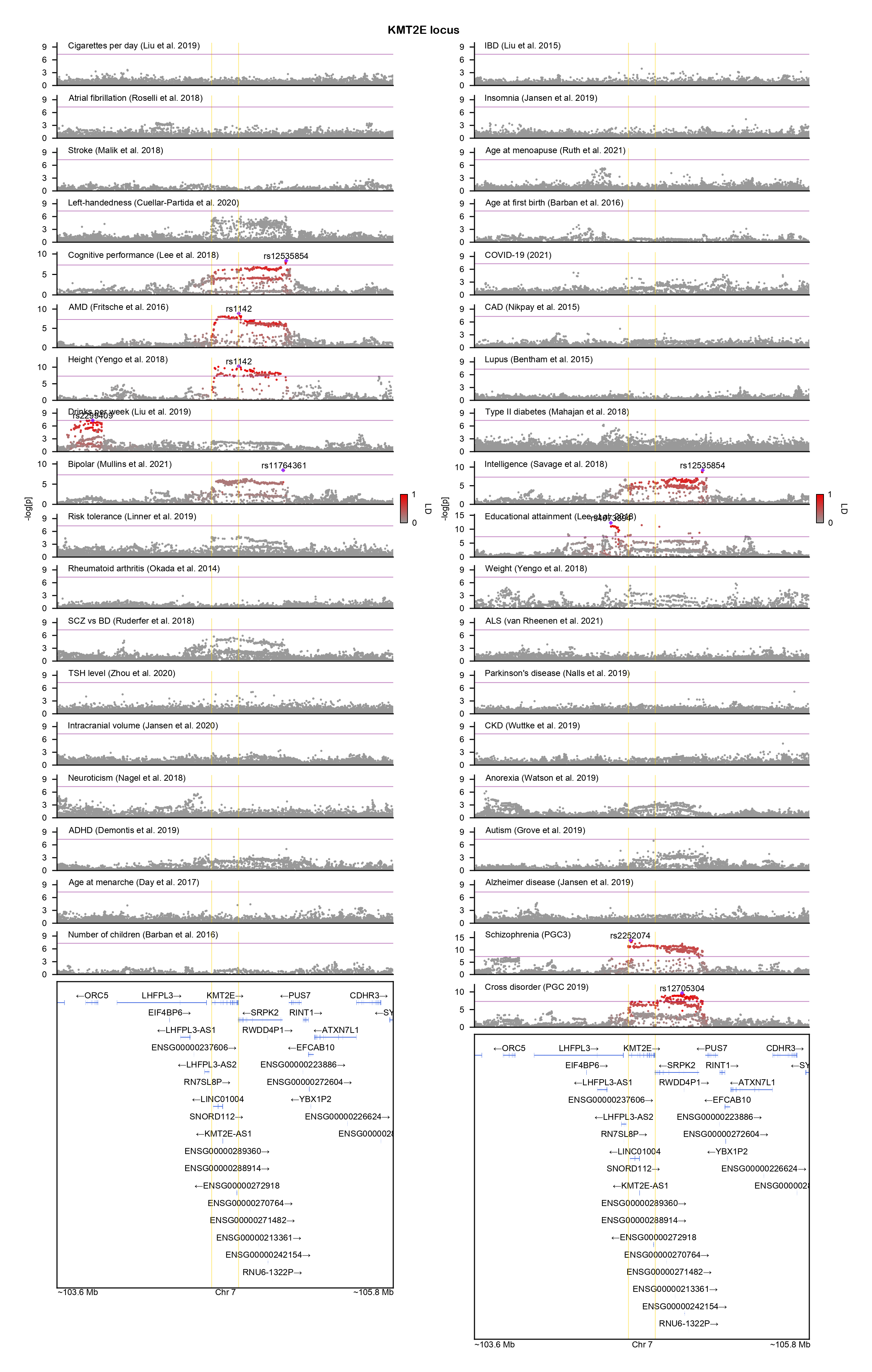

### LPCAT4-locuszoom.png

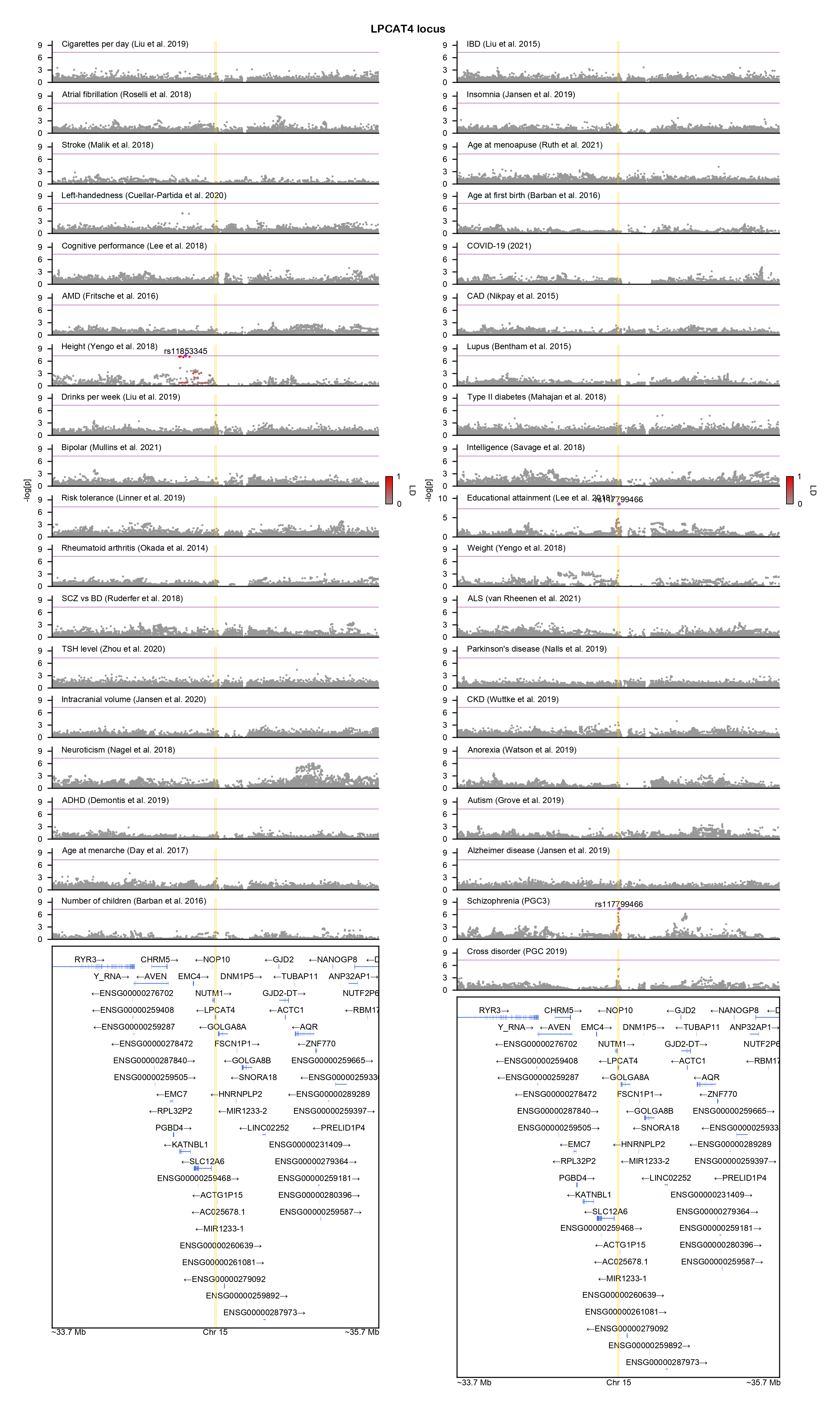

### MAD1L1-locuszoom.png

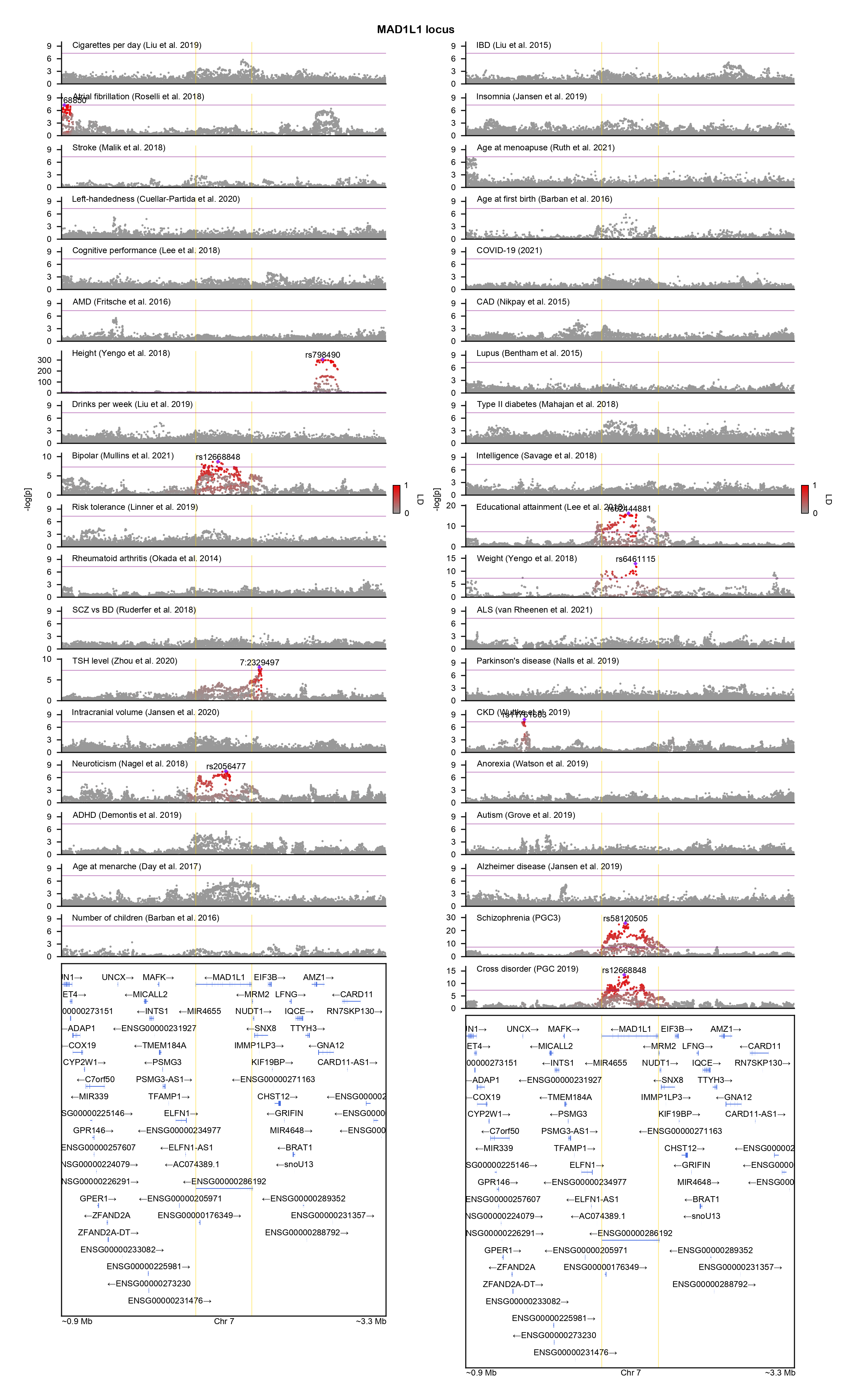

### MAIP1-locuszoom.png

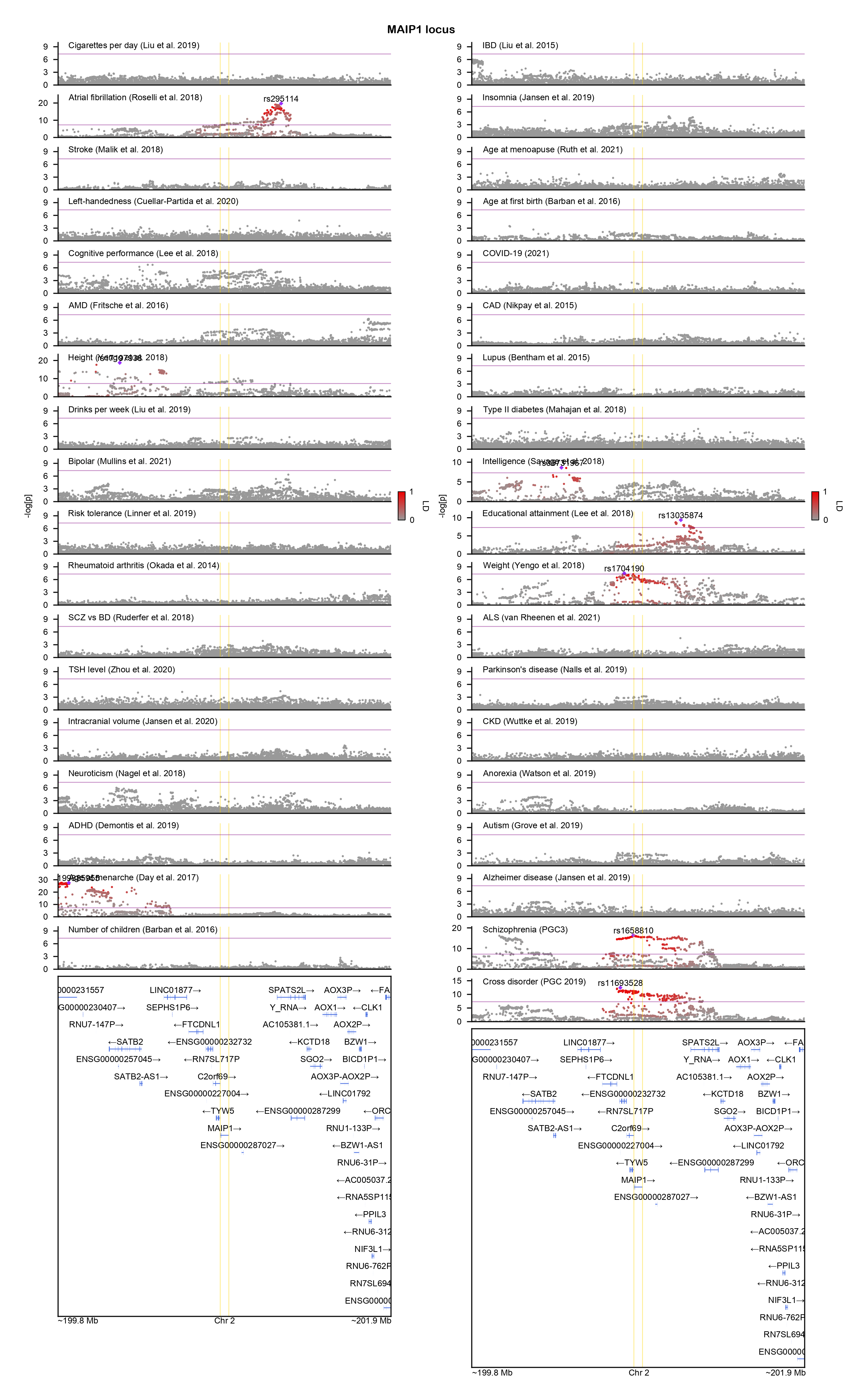

### MEF2C-locuszoom.png

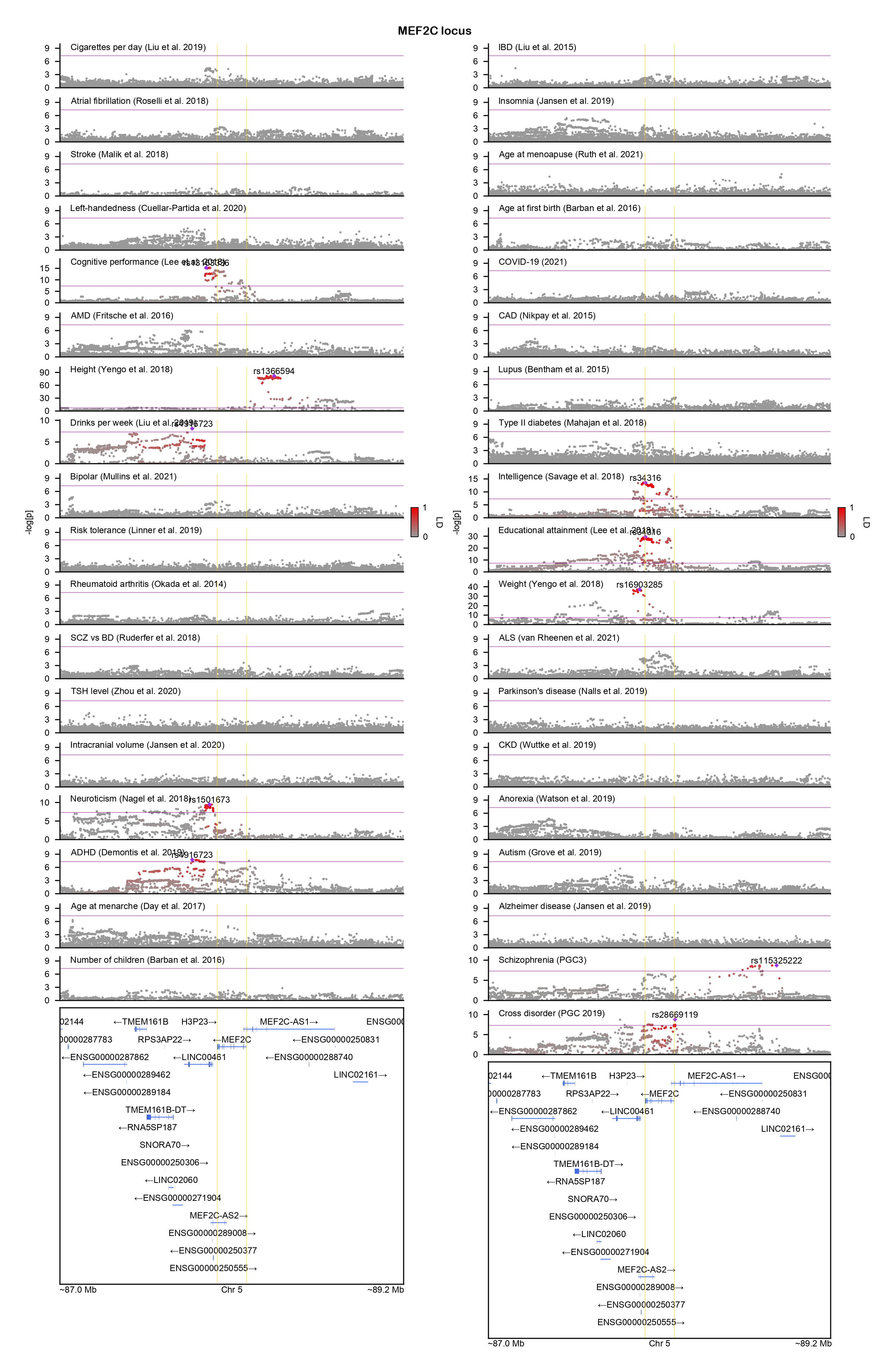

### MLXIP-locuszoom.png

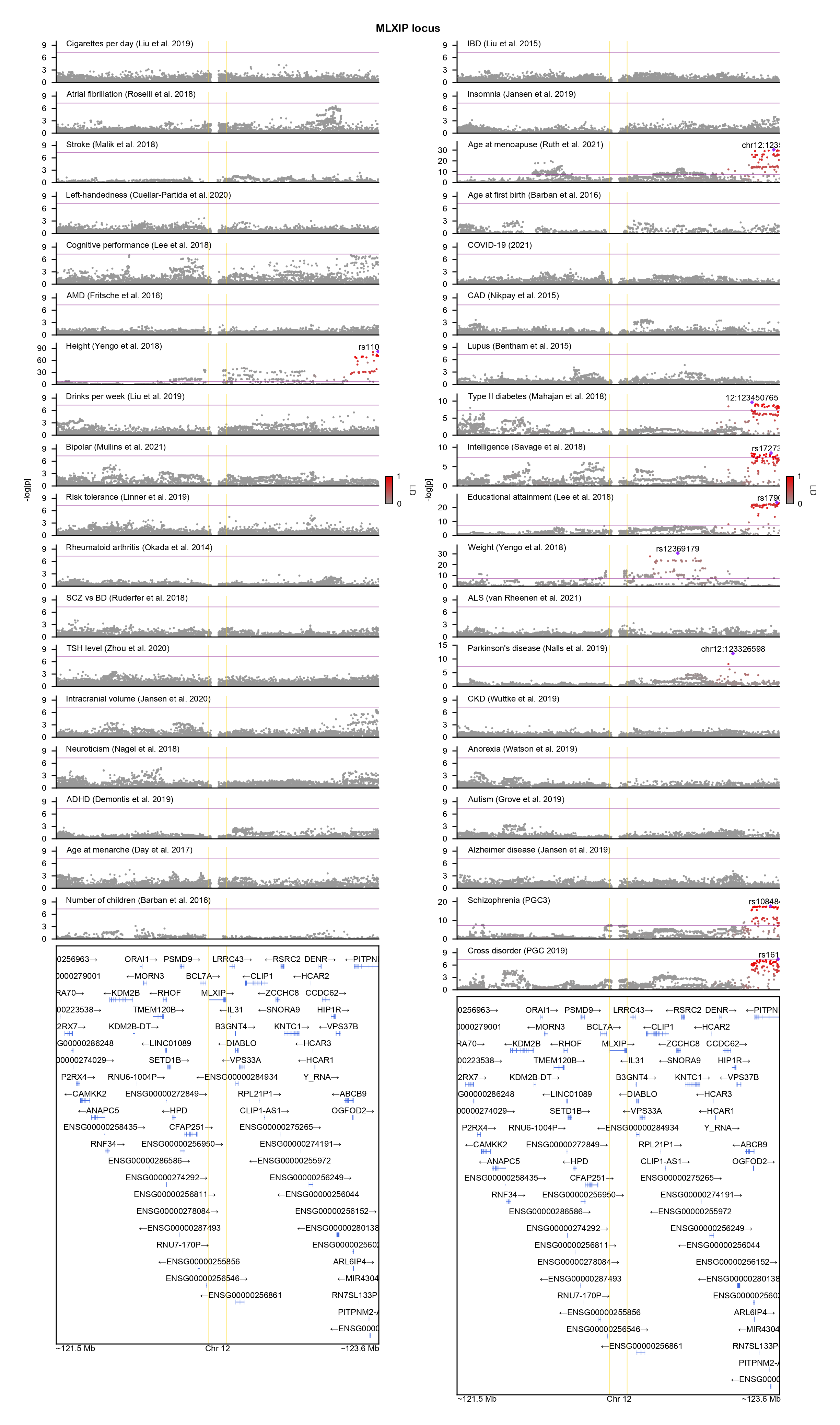

### MYT1L-locuszoom.png

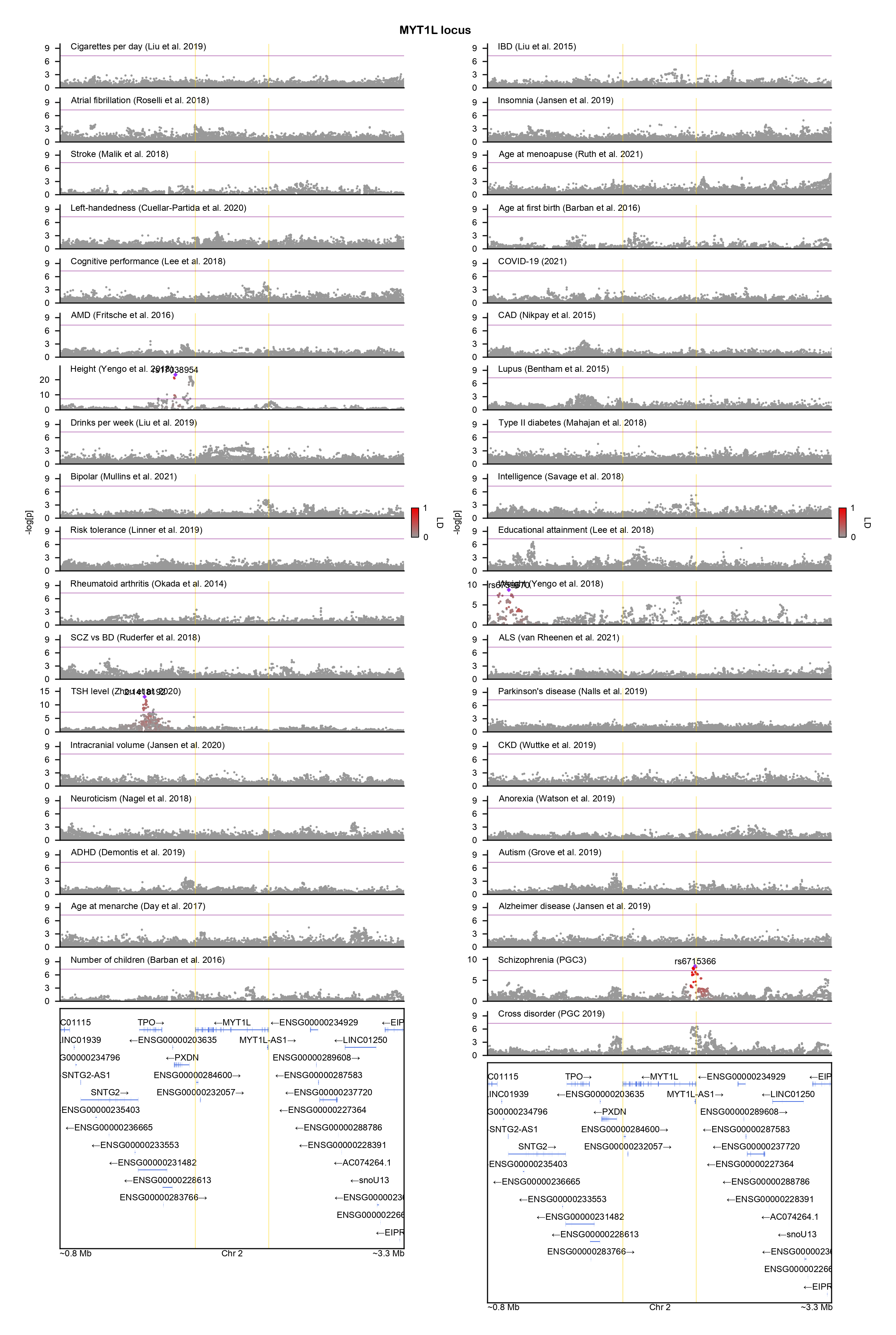

### NAT8-locuszoom.png

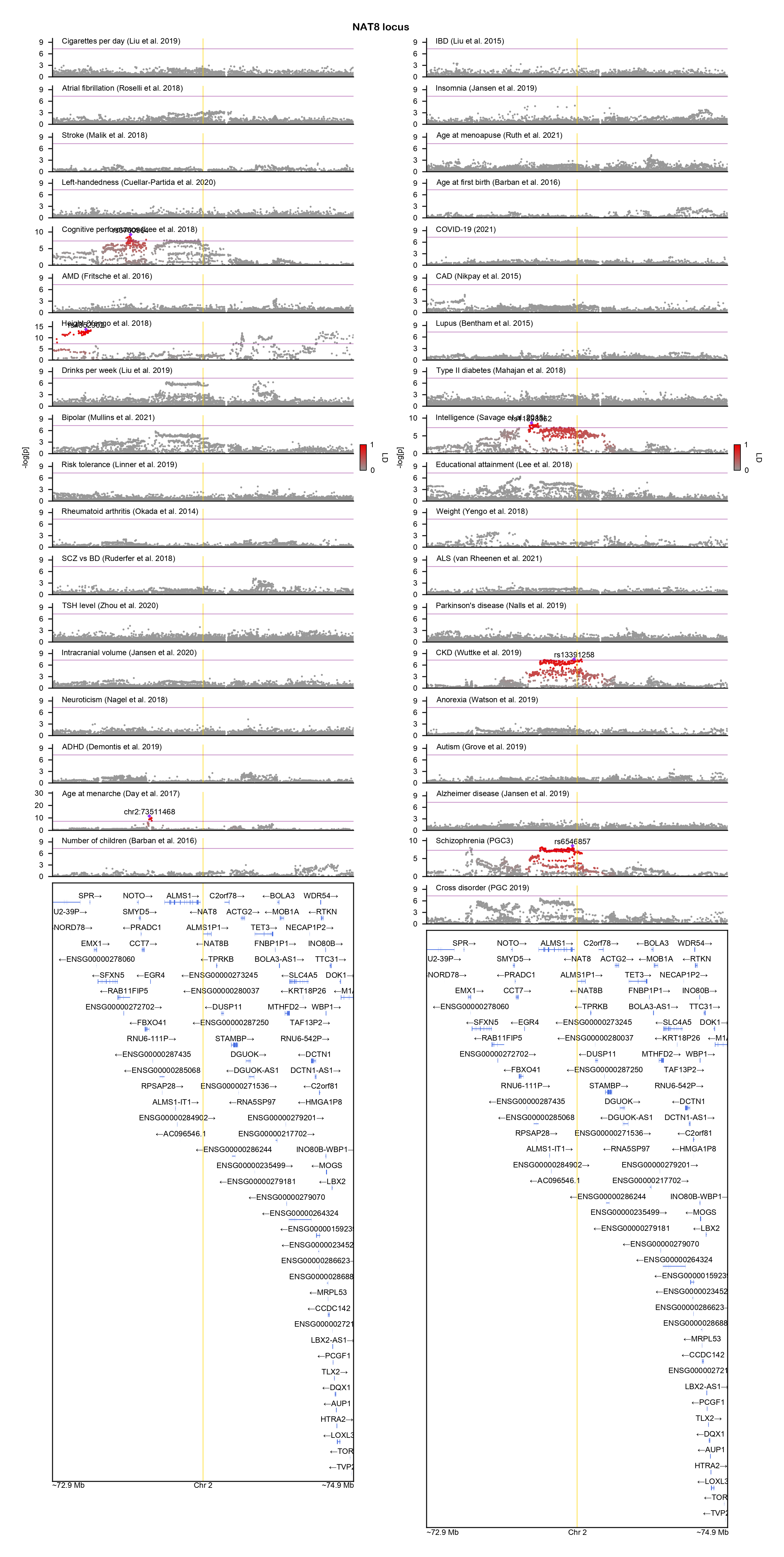

### NCAM2-locuszoom.png

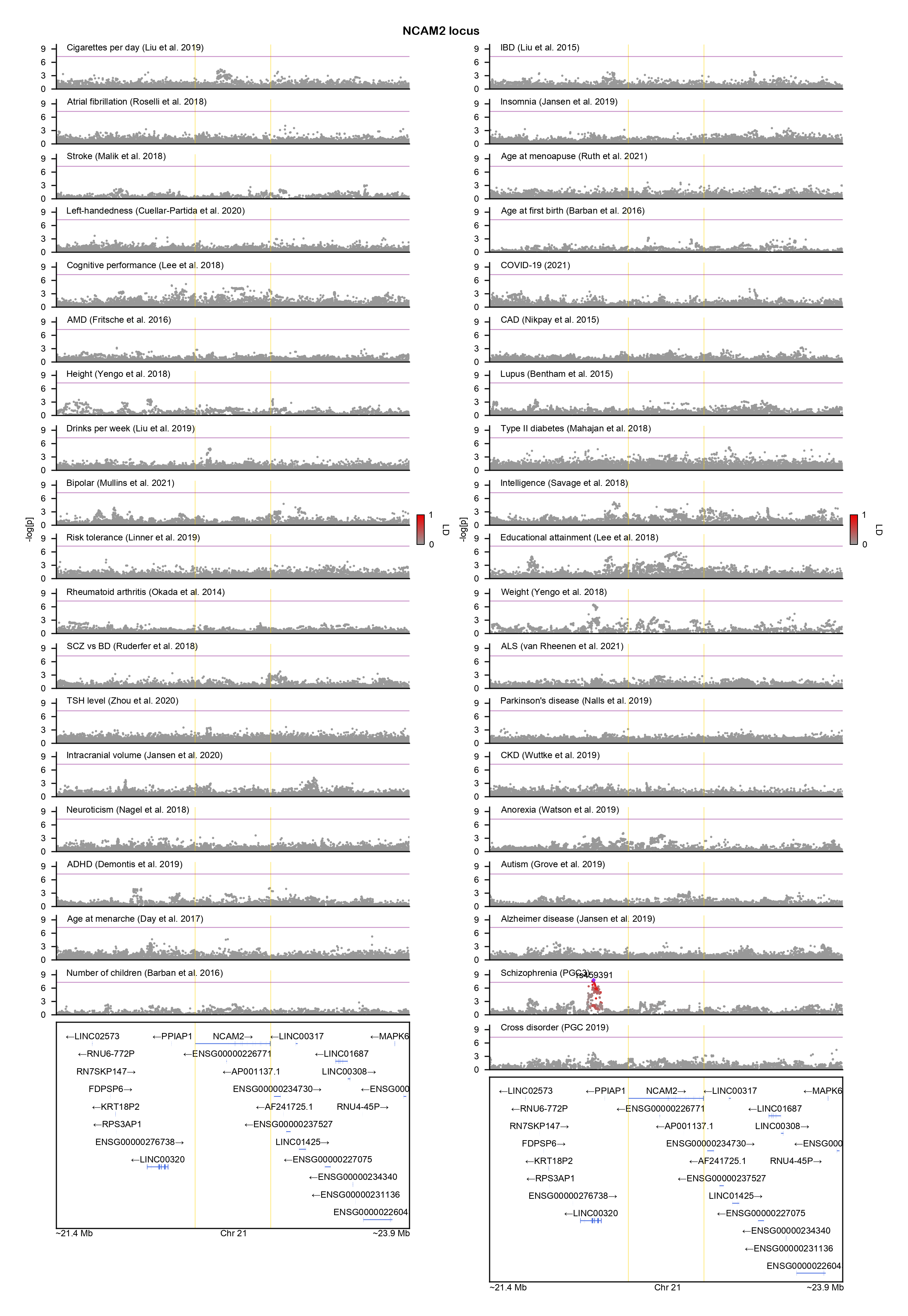

### NFIB-locuszoom.png

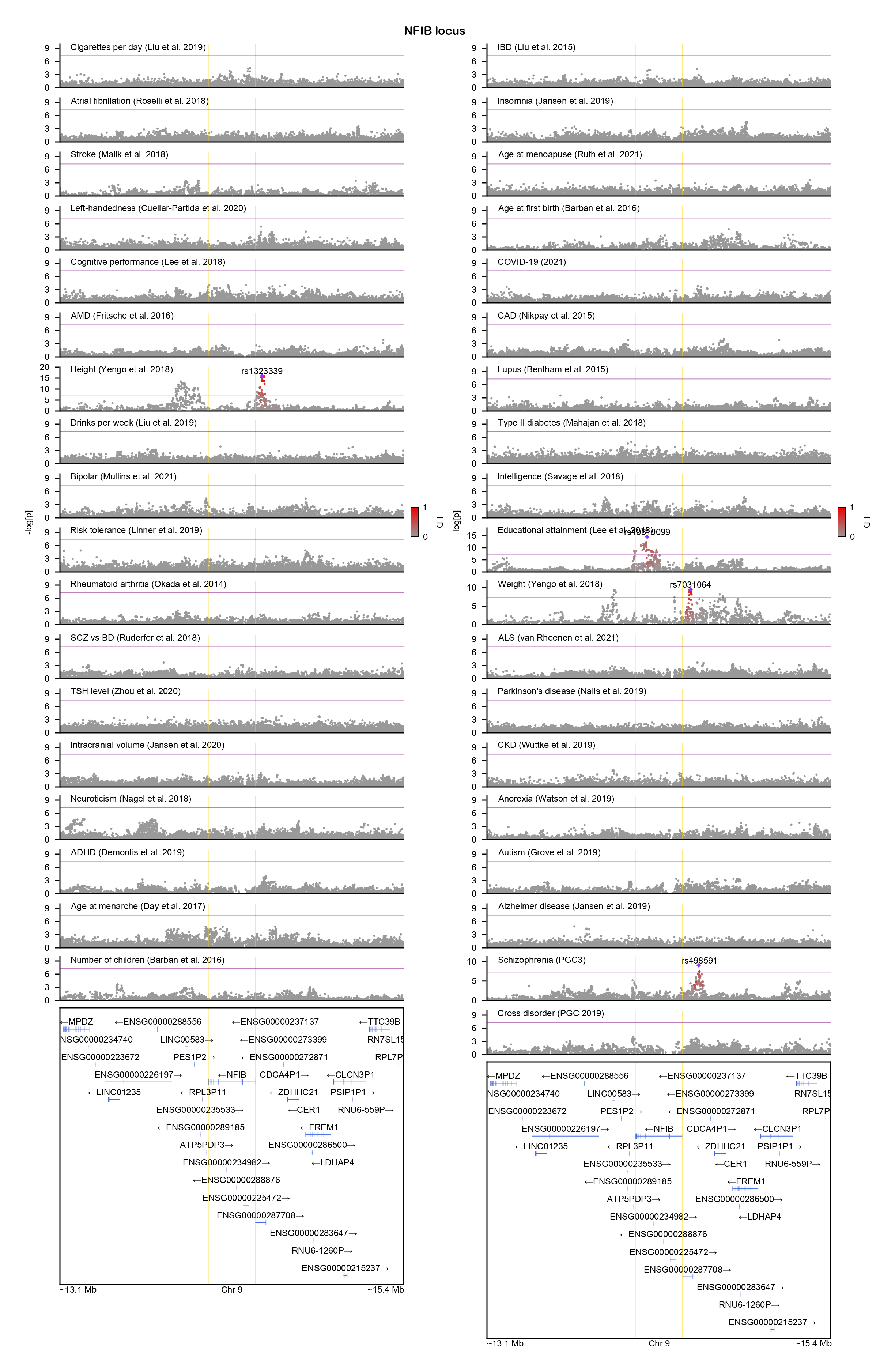

### NOTCH4-locuszoom.png

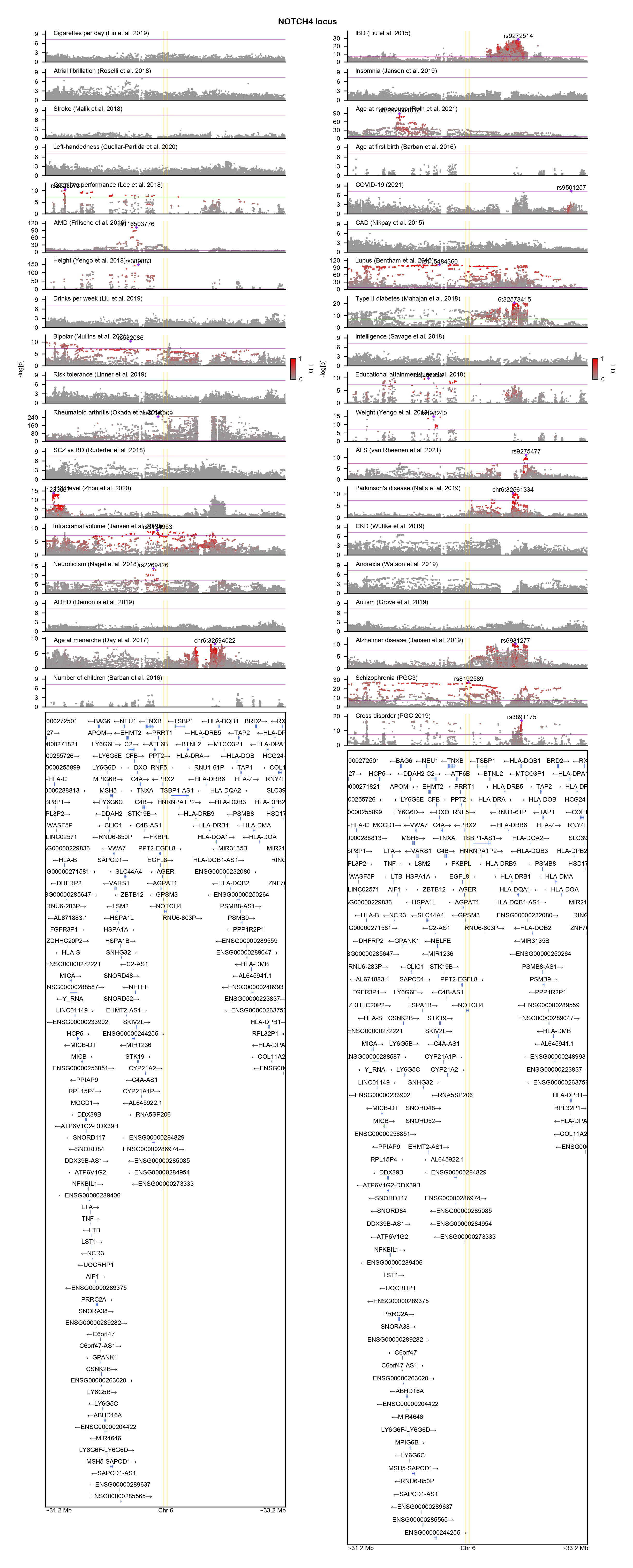

### NSUN6-locuszoom.png

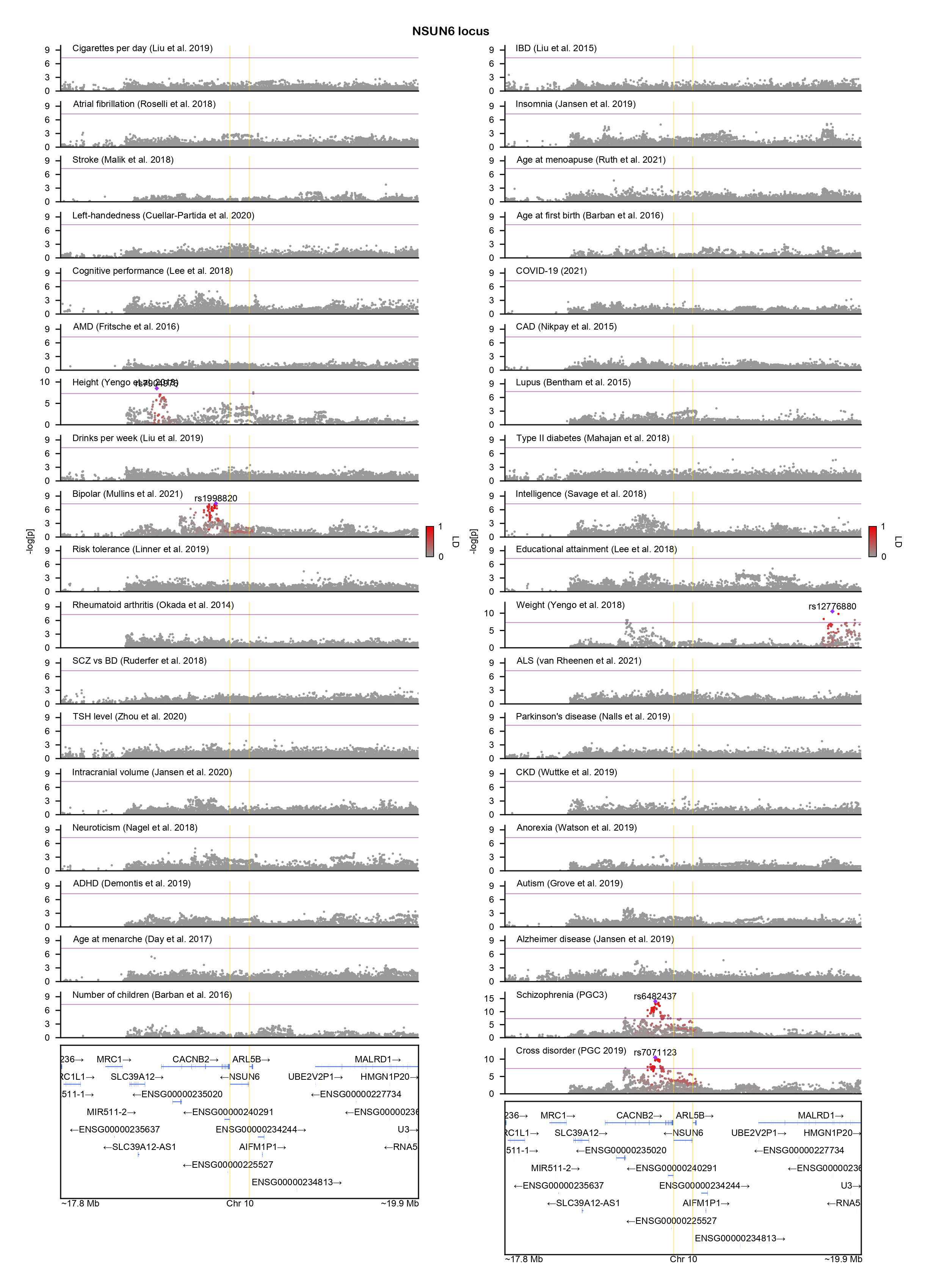

### PCDHA1-locuszoom.png

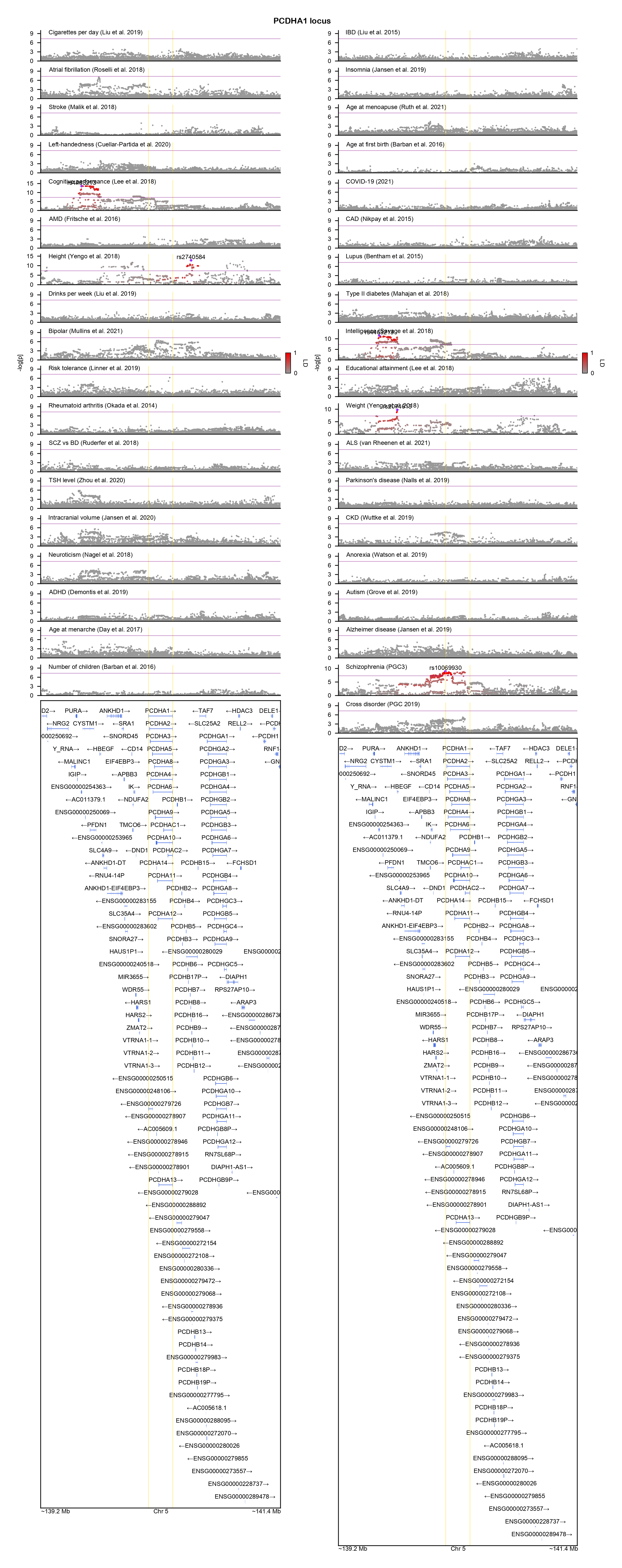

### PCNX3-locuszoom.png

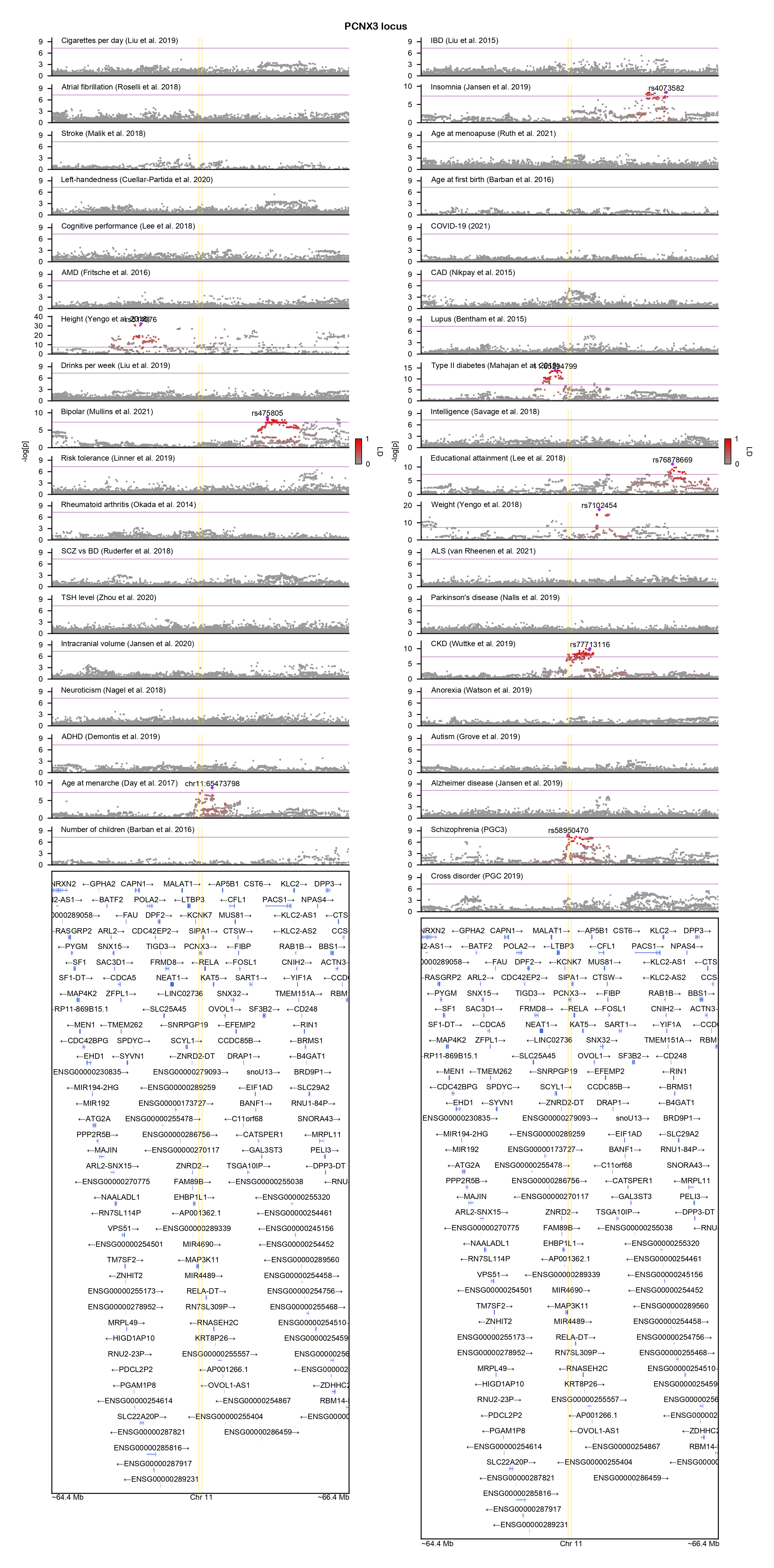

### PITPNM2-locuszoom.png

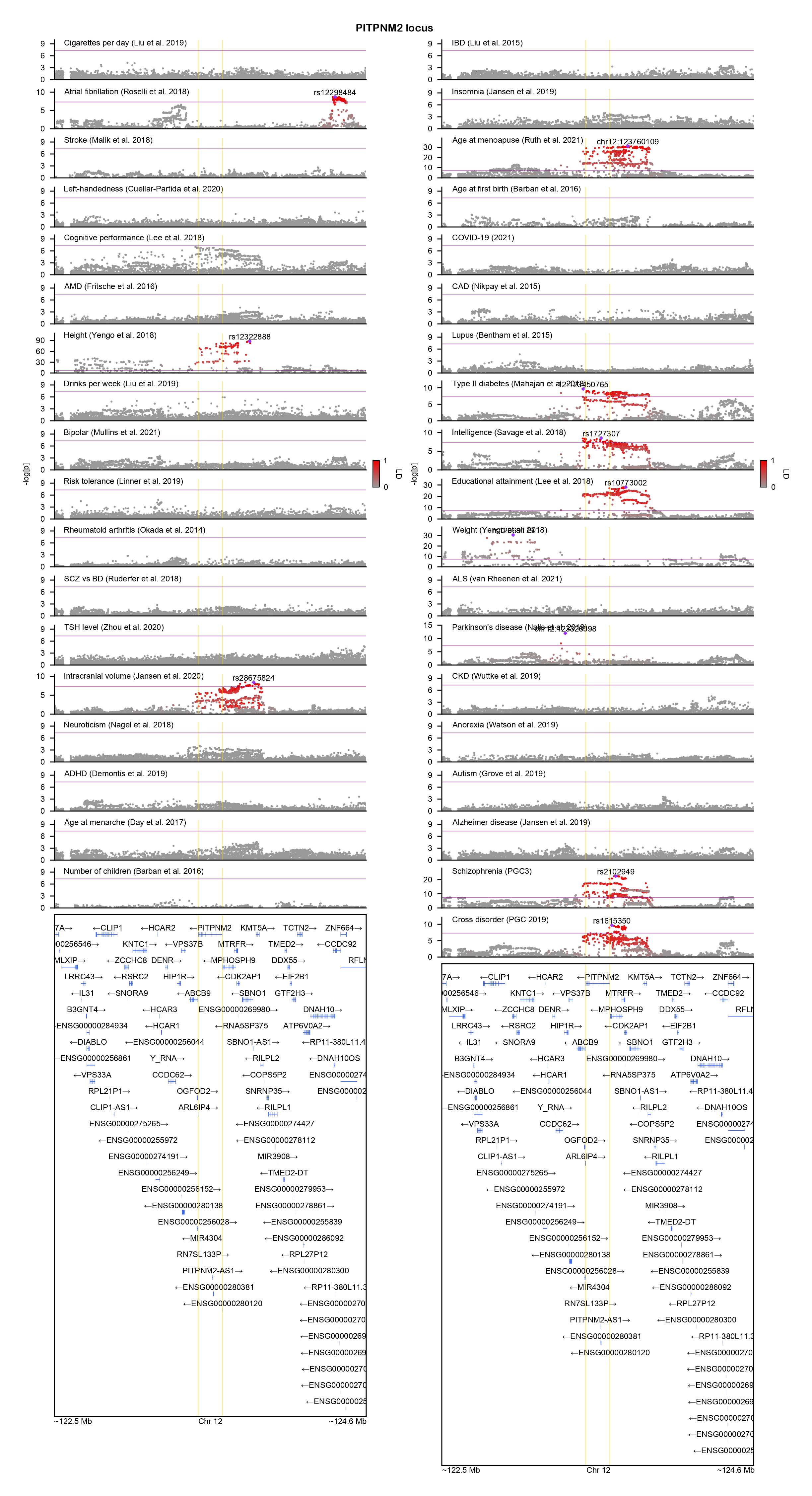

### PODXL-locuszoom.png

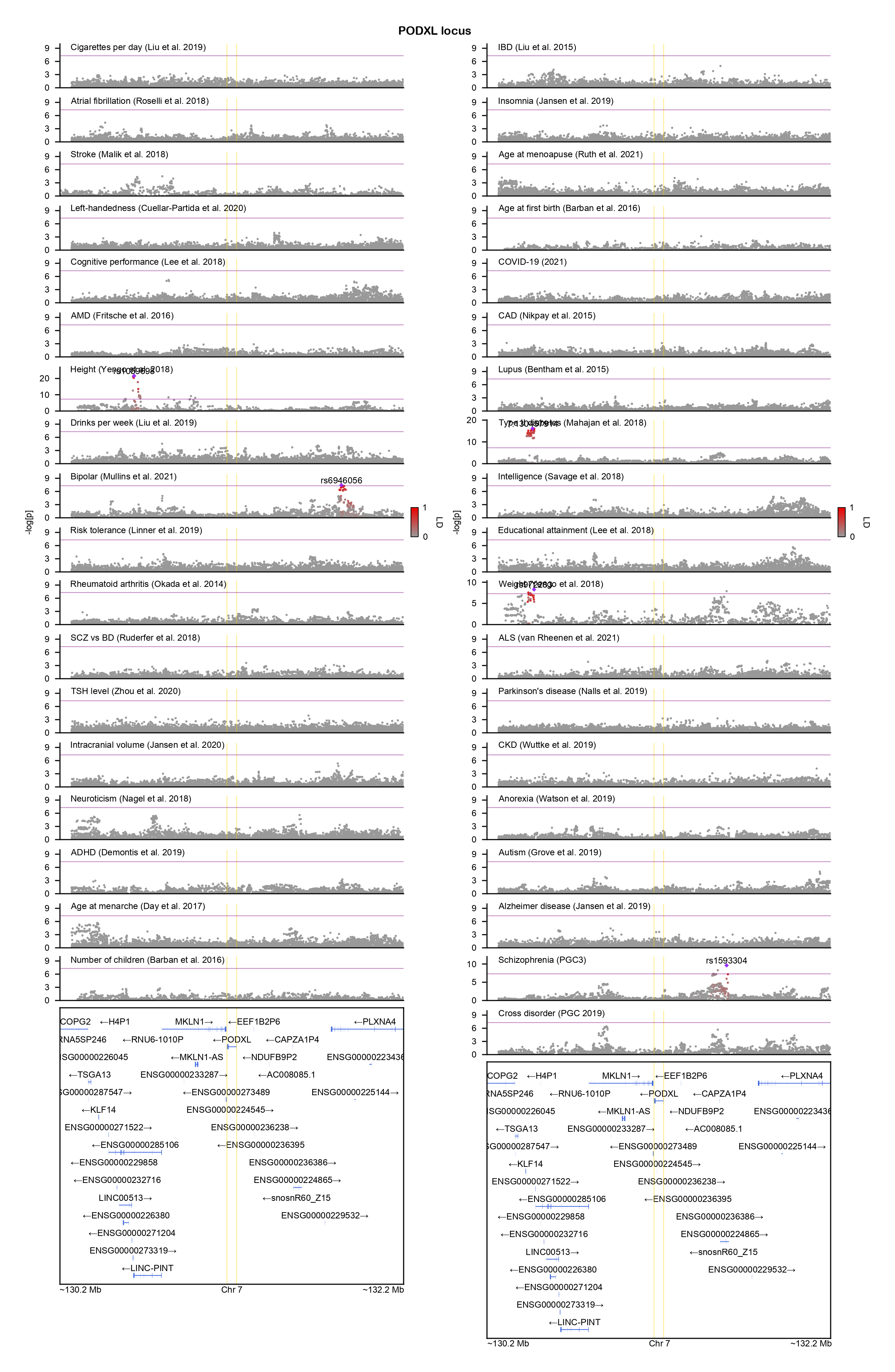

### POU3F3-locuszoom.png

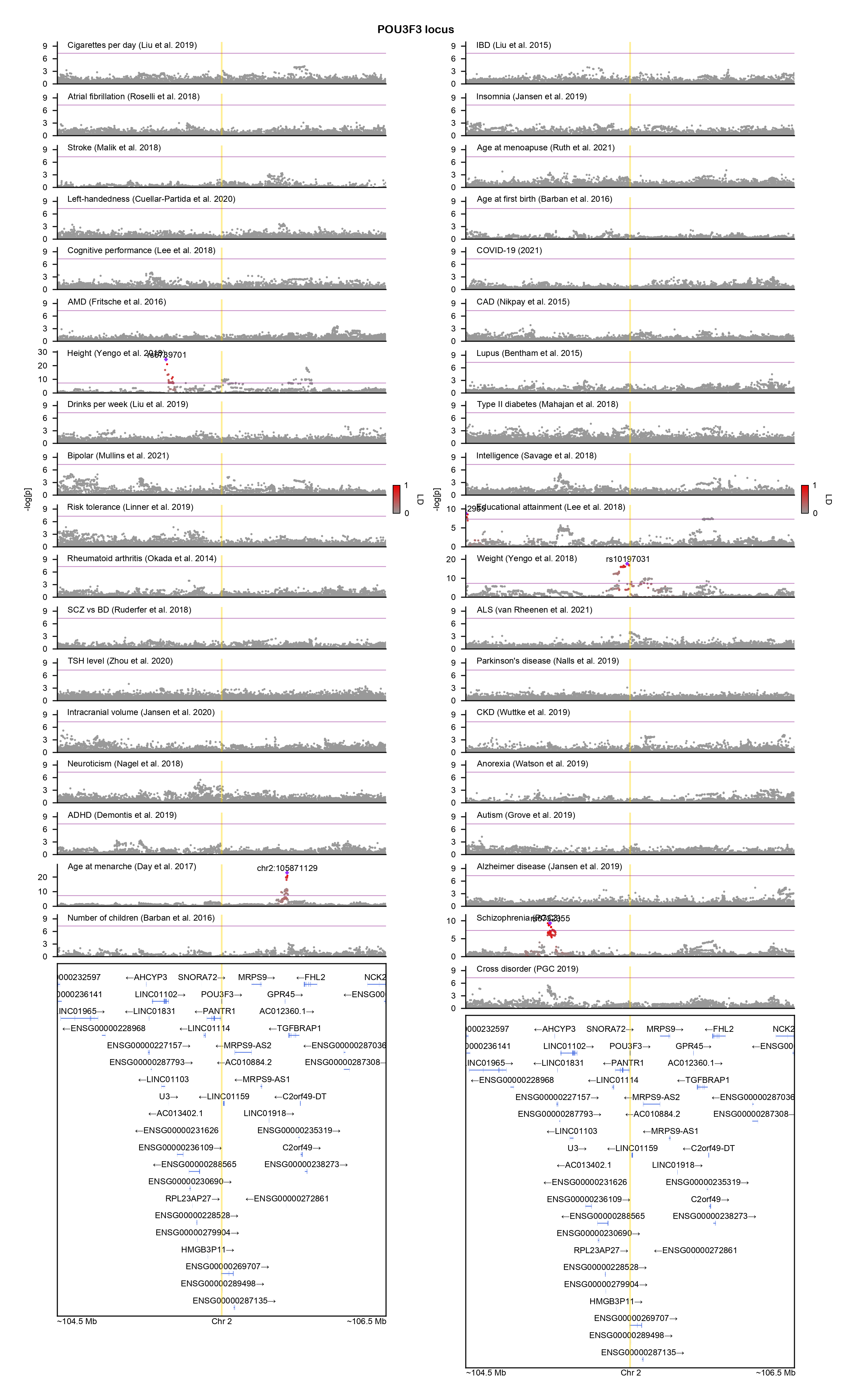

### PPARGC1A-locuszoom.png

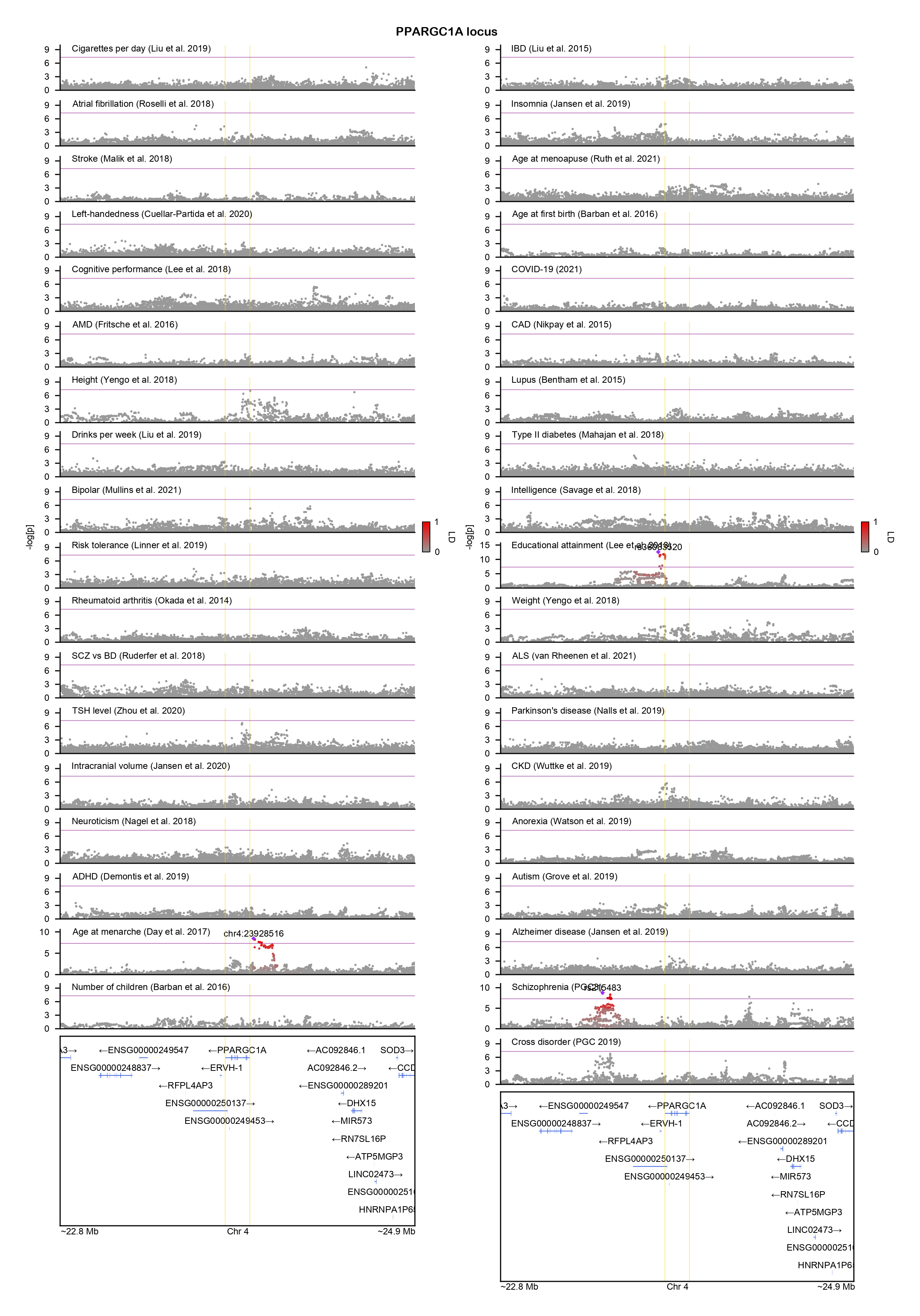

### PPP1R13B-locuszoom.png

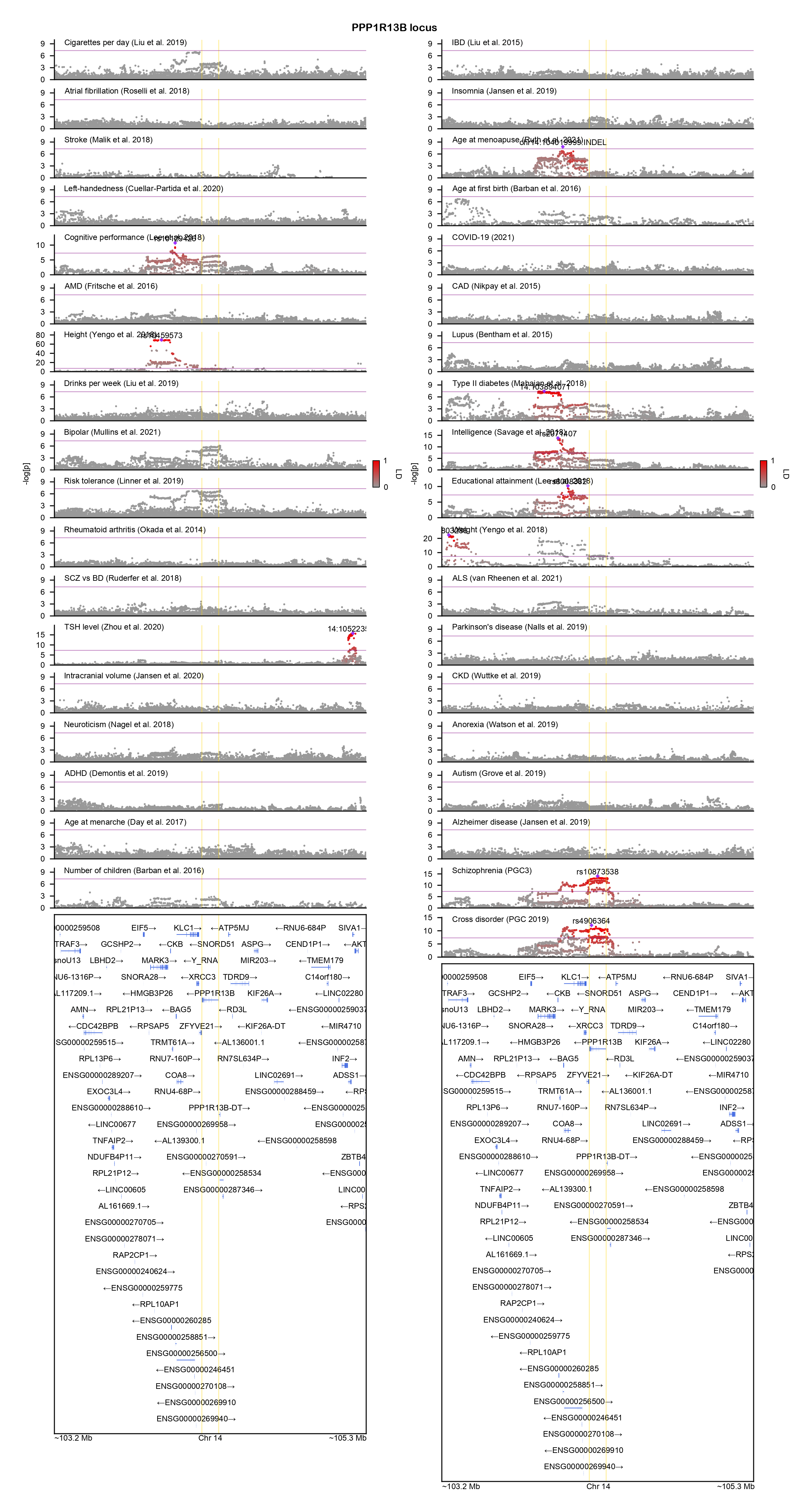

### PPP1R16B-locuszoom.png

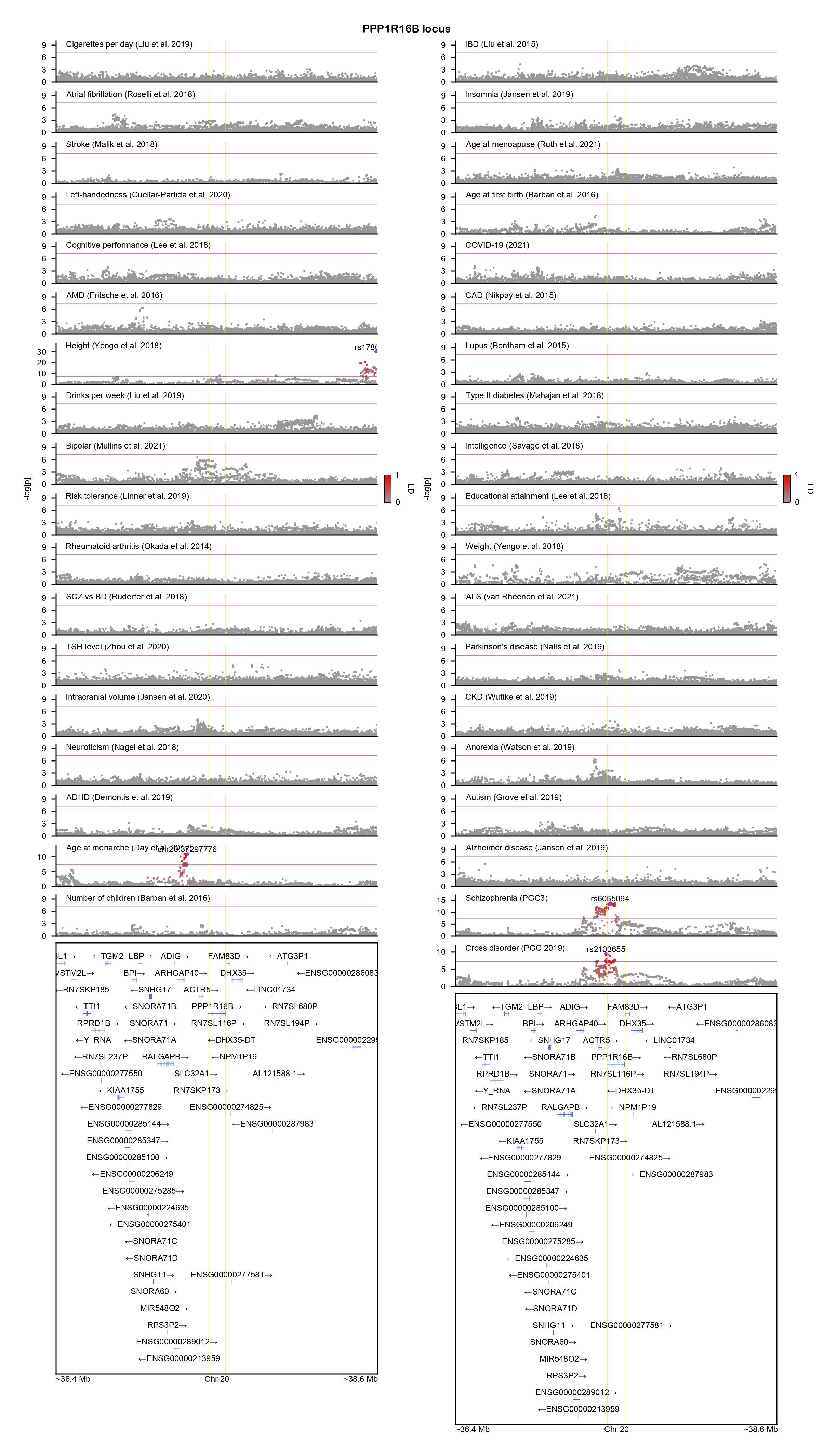

### PPP1R26-locuszoom.png

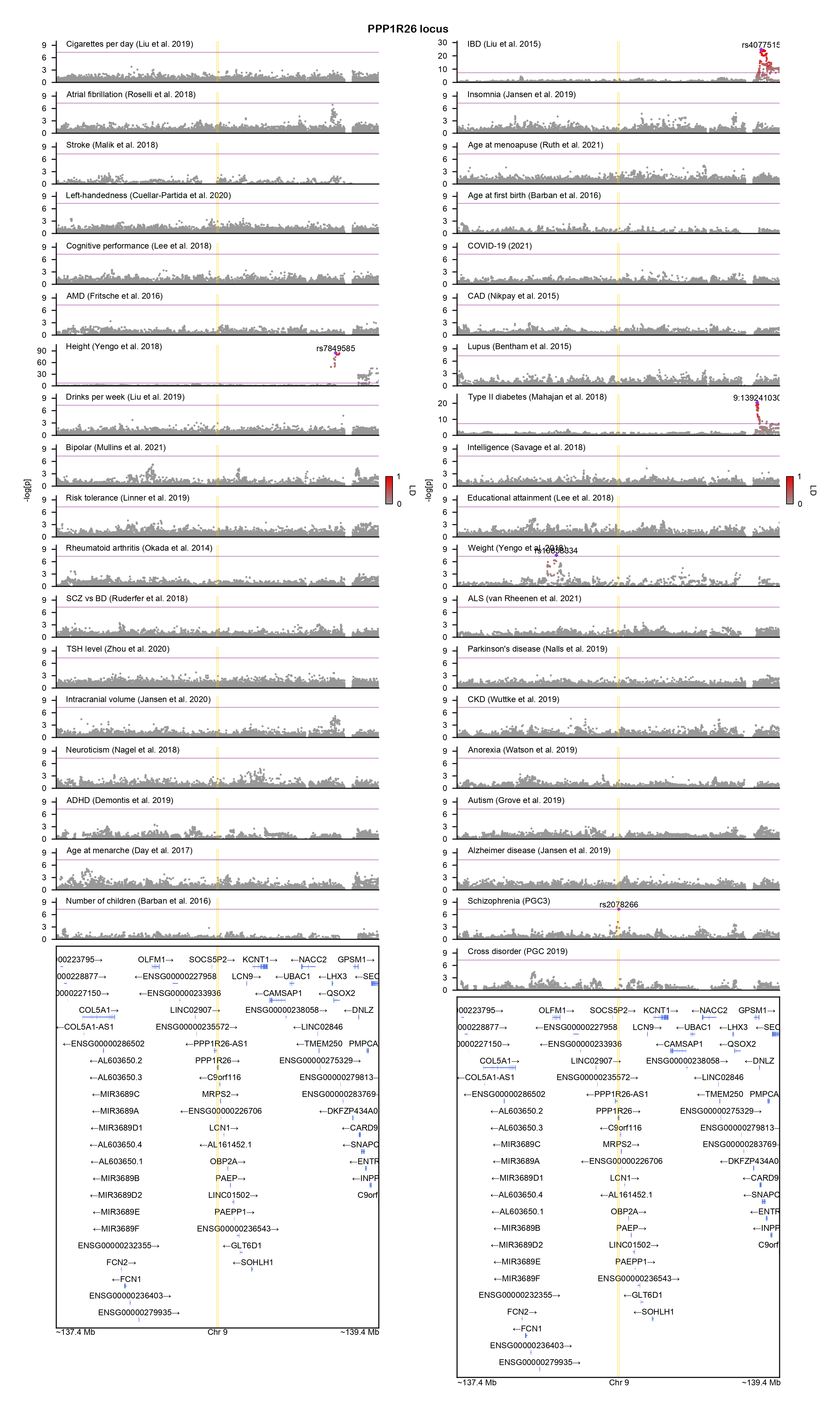

### PRR20A-locuszoom.png

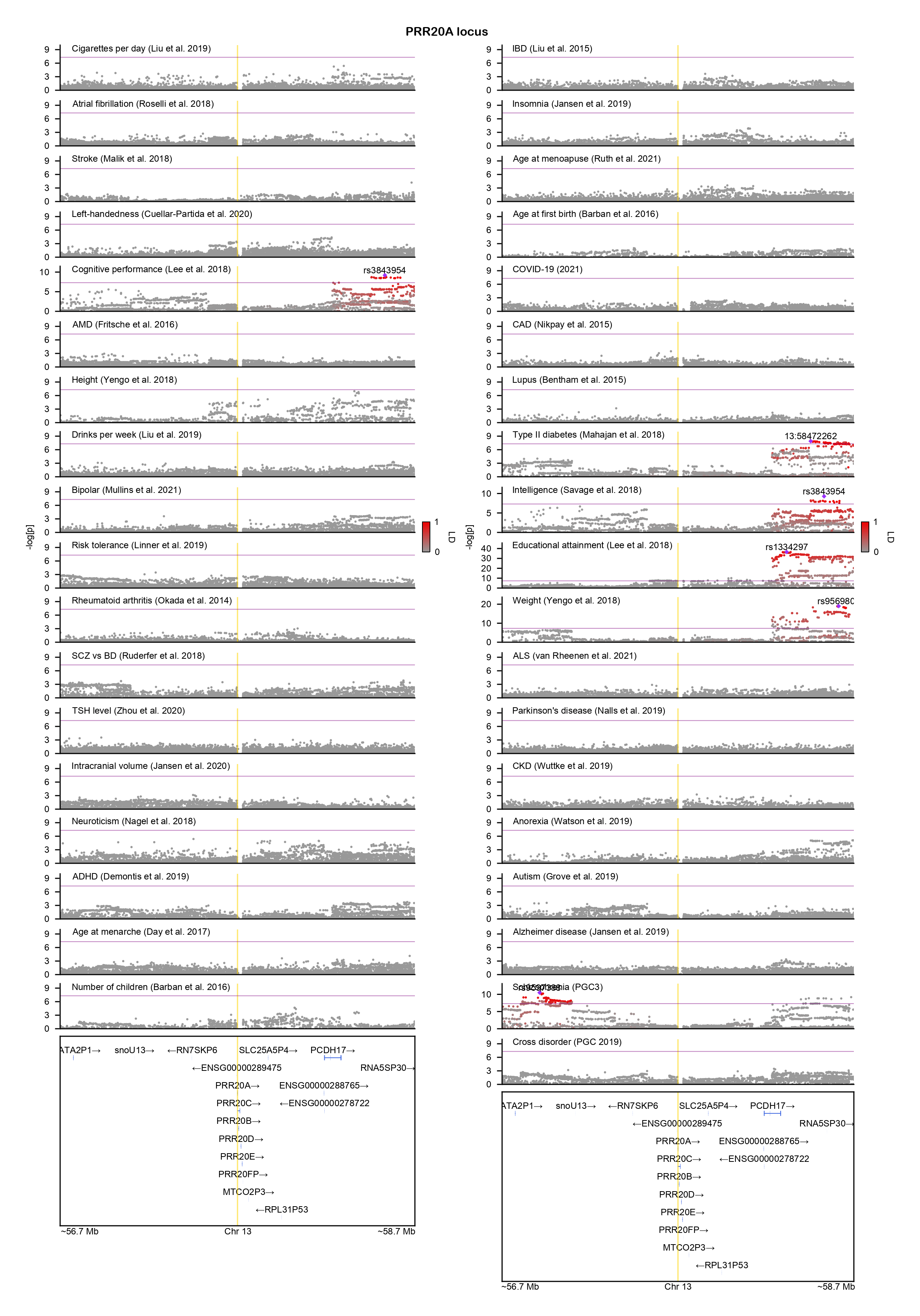

### PTN-locuszoom.png

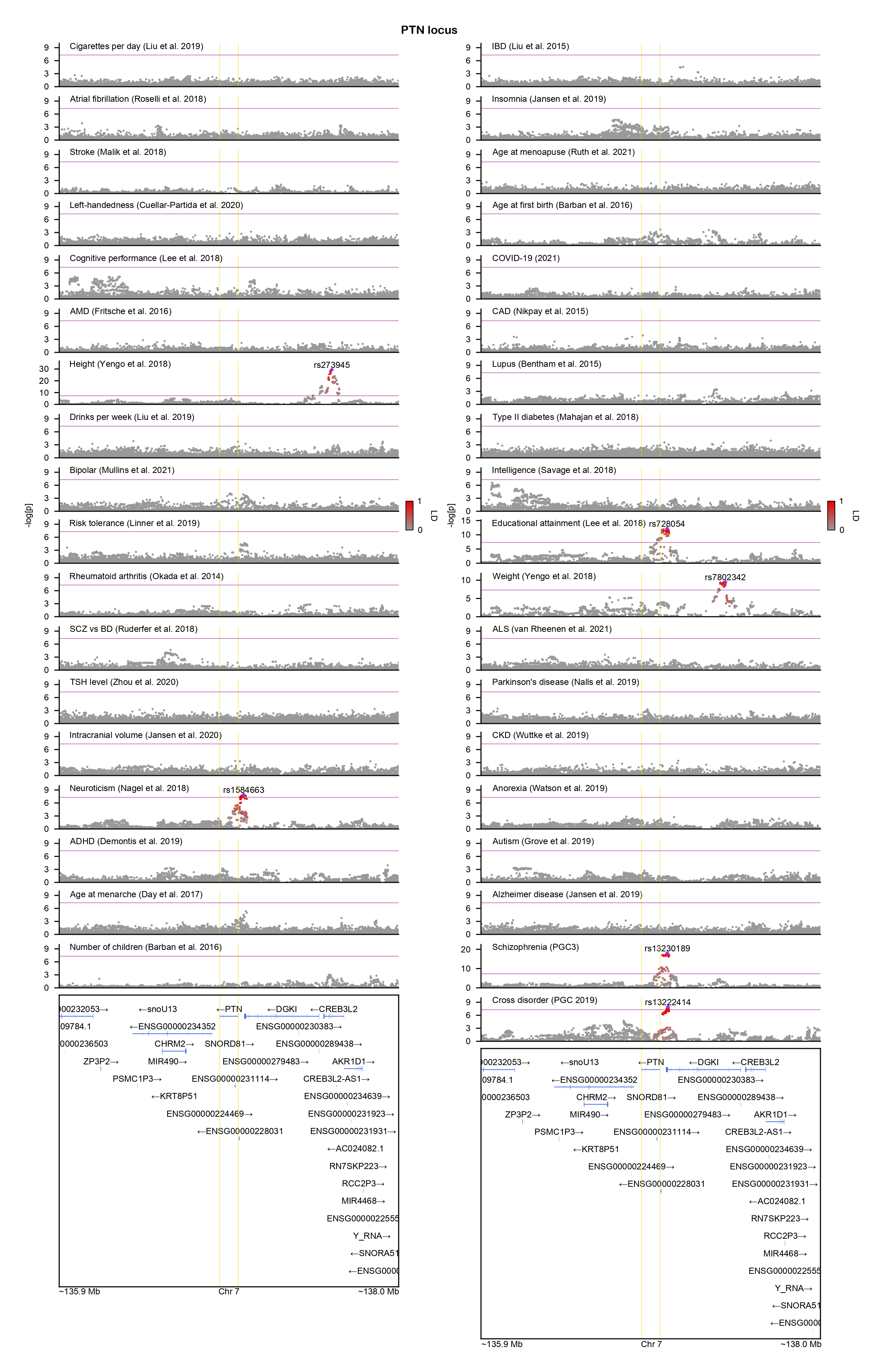

### RBM26-locuszoom.png

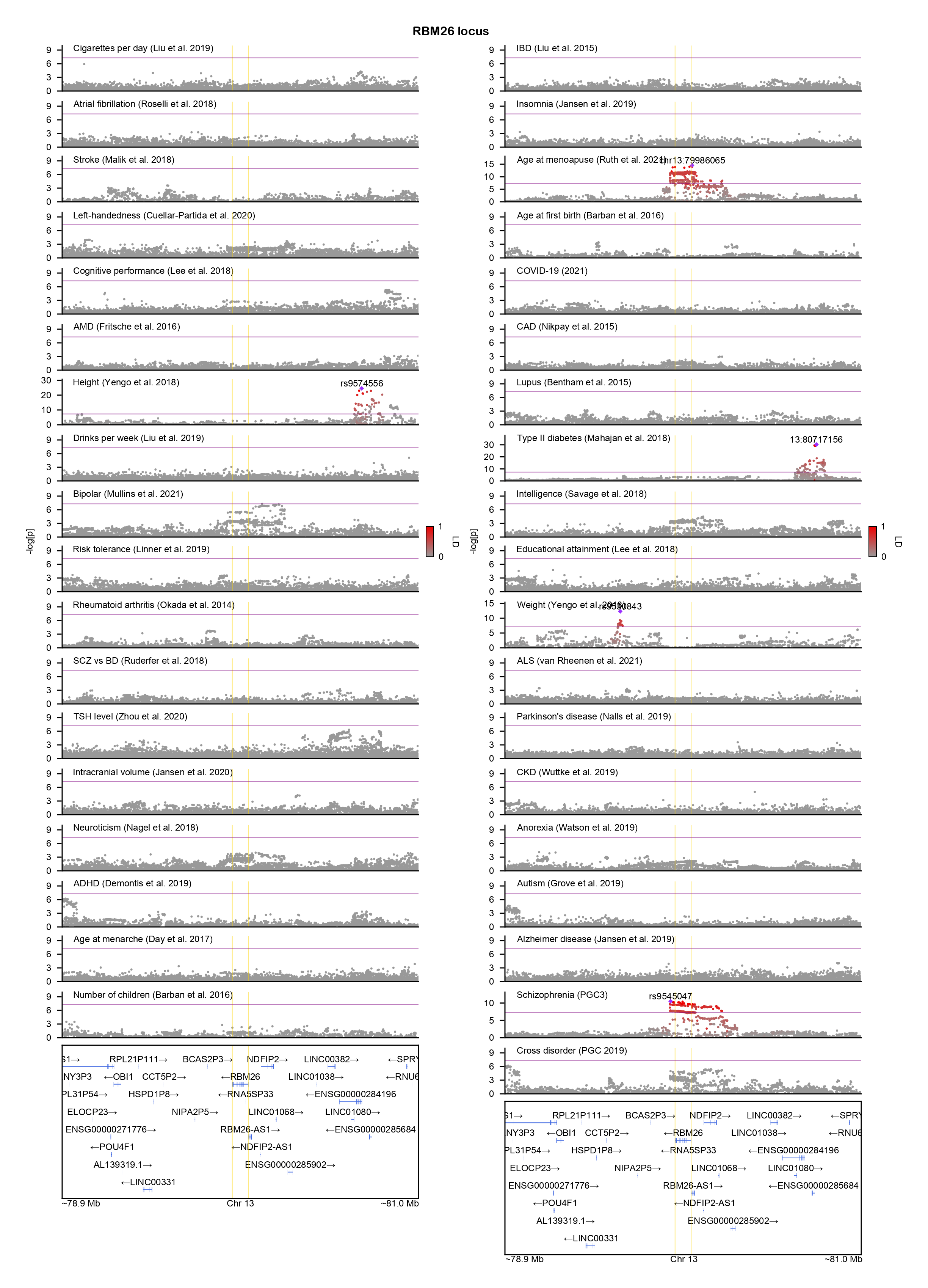

### SCN3B-locuszoom.png

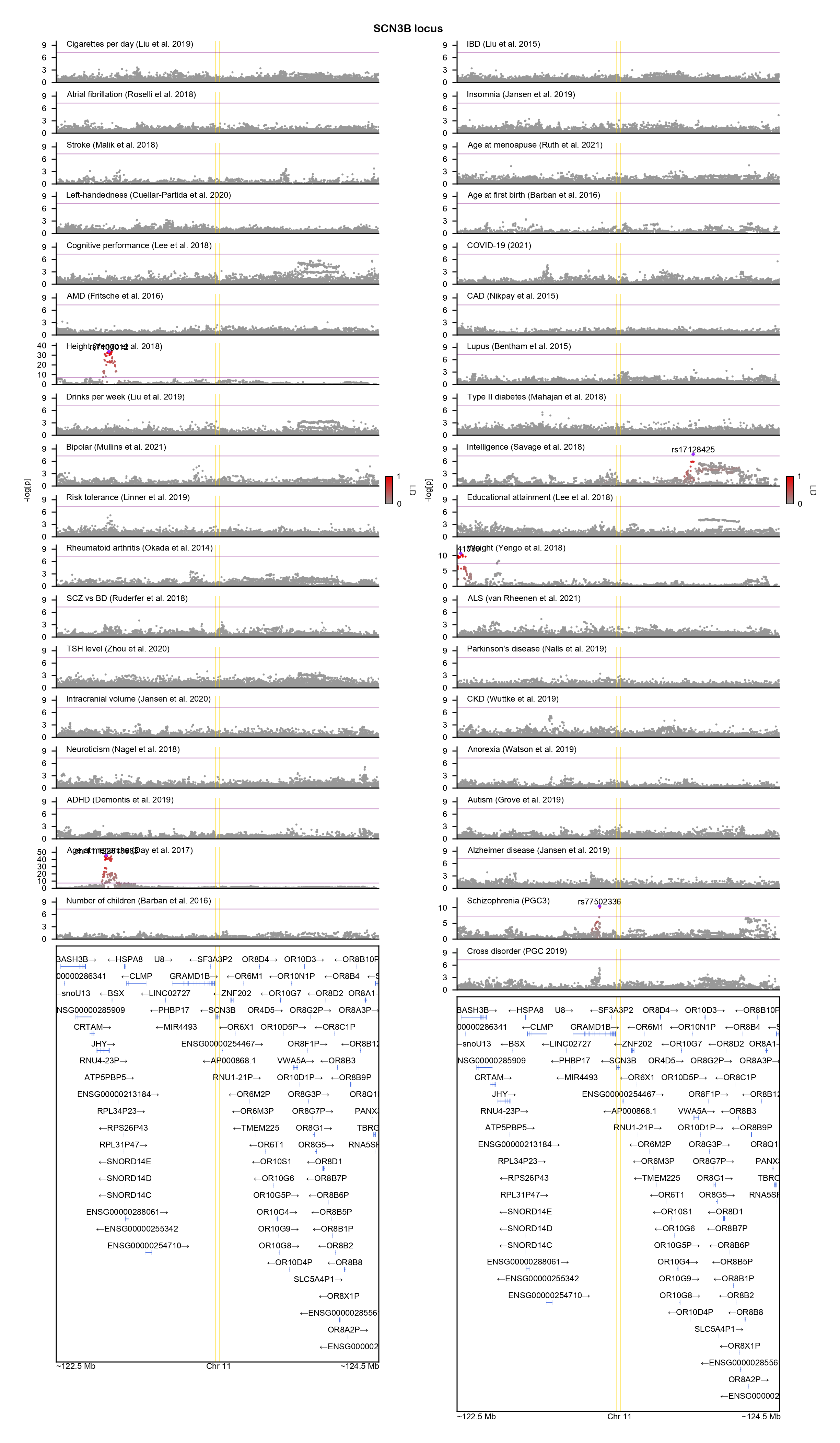

### SGCD-locuszoom.png

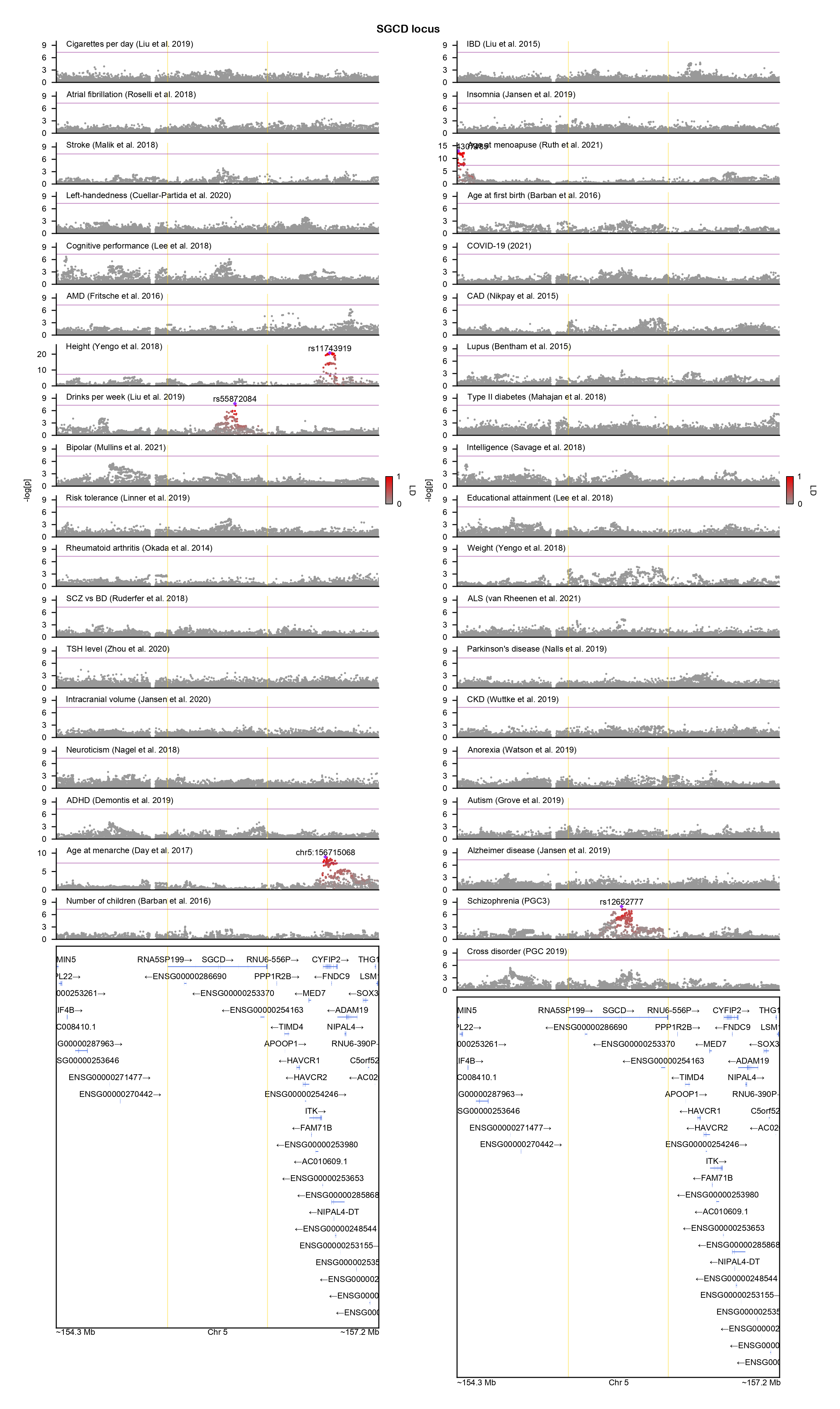

### SHANK3-locuszoom.png

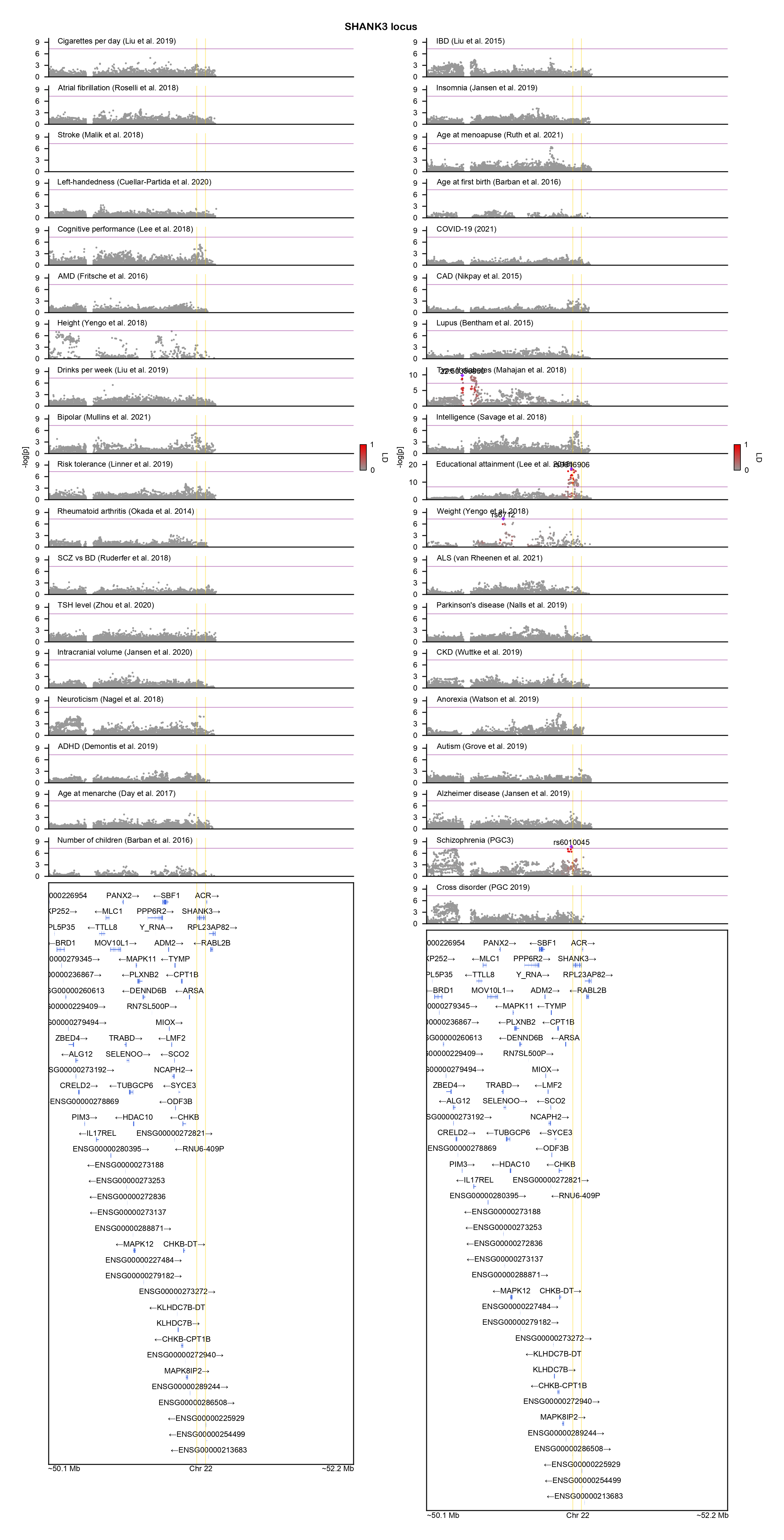

### SLC39A8-locuszoom.png

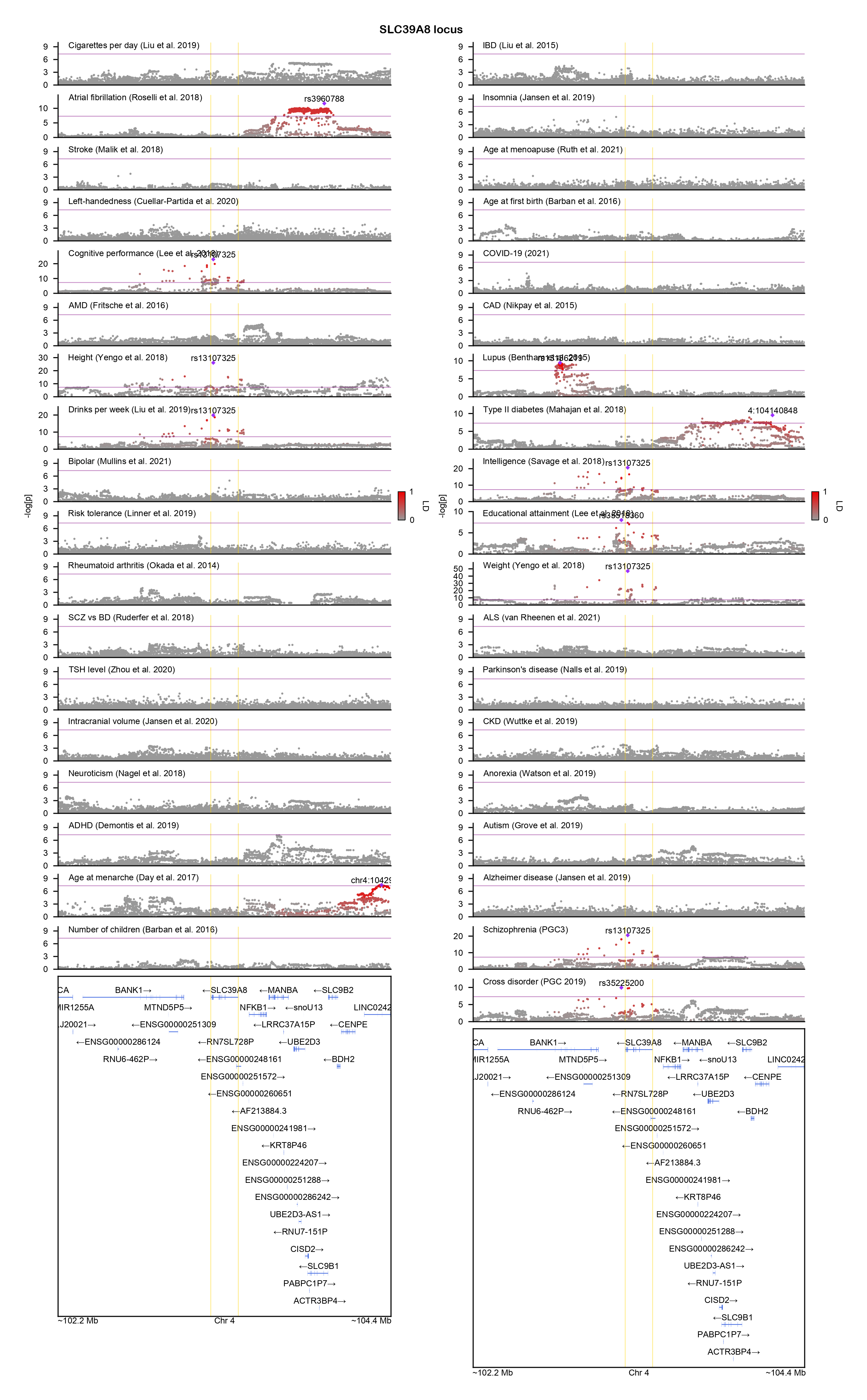

### SLC45A1-locuszoom.png

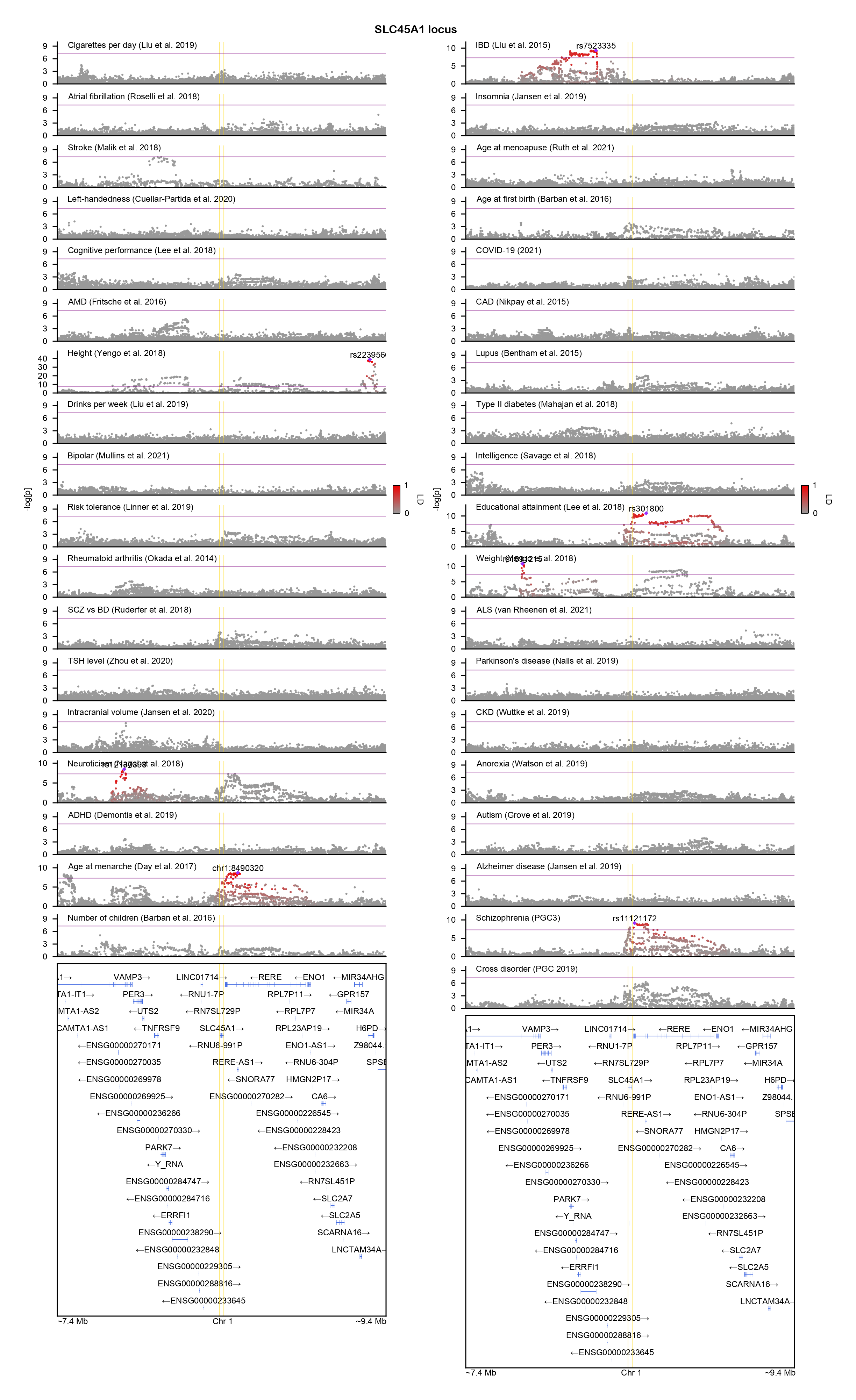

### SNORC-locuszoom.png

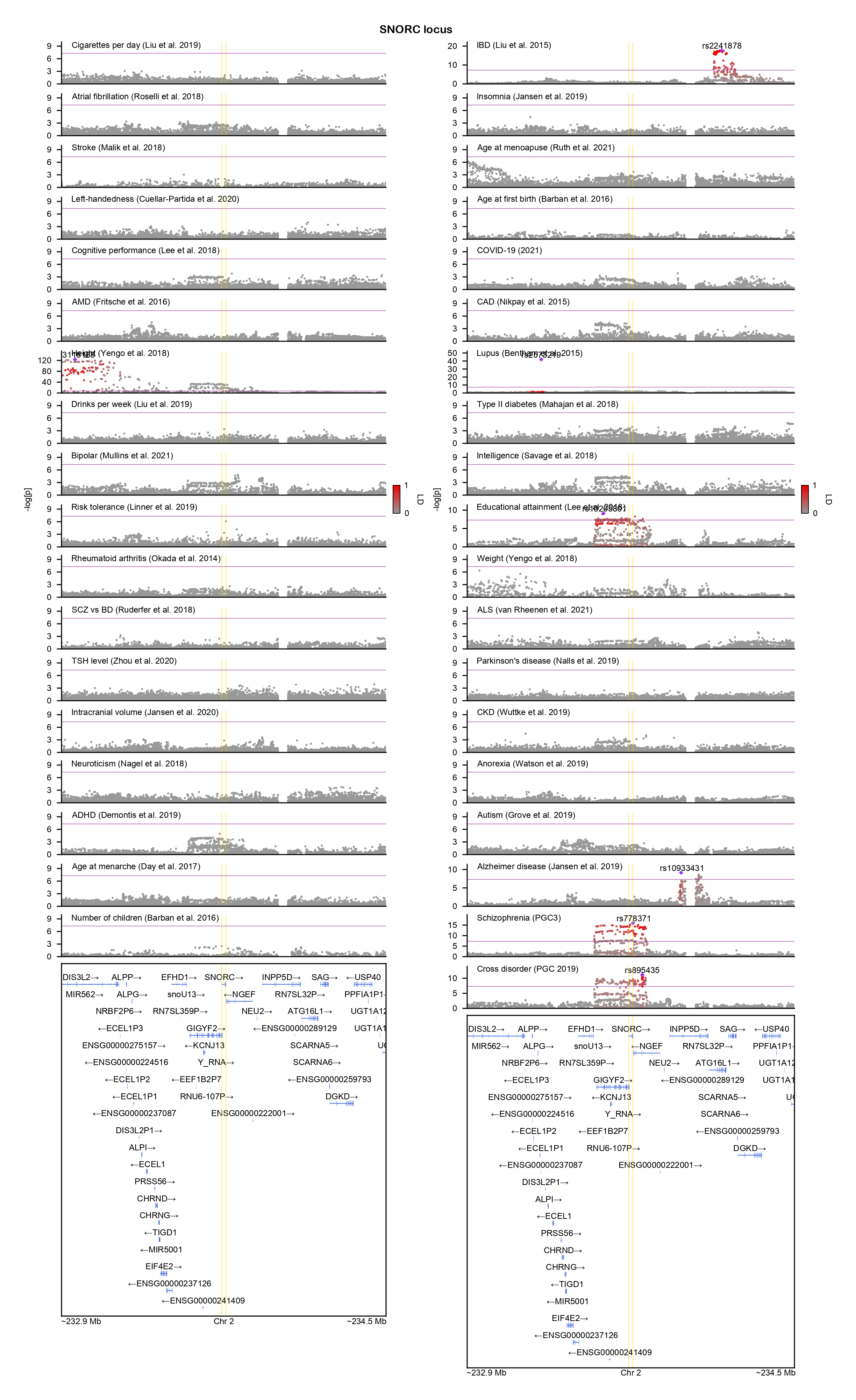

### SNX19-locuszoom.png

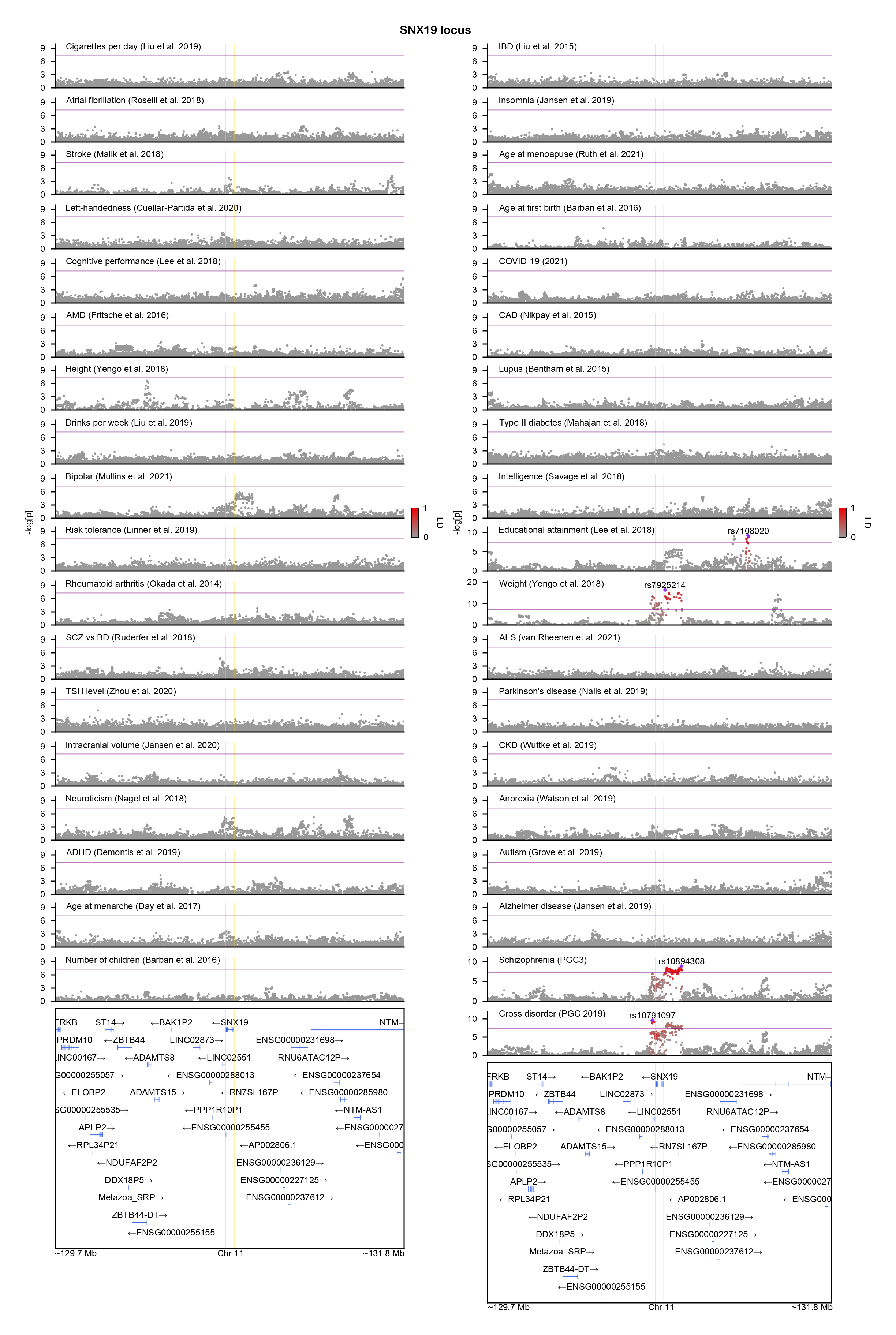

### TBC1D15-locuszoom.png

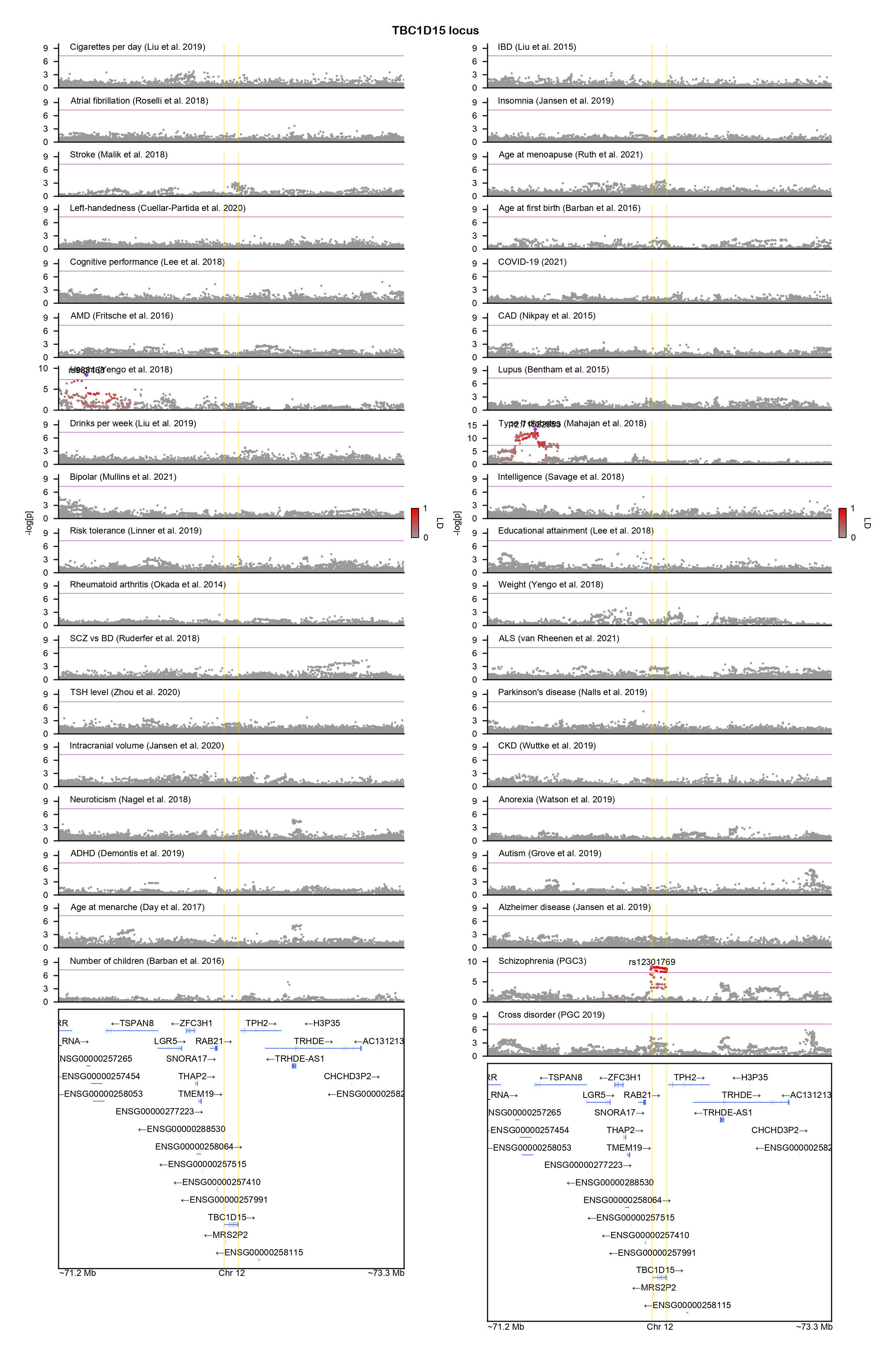

### TGFBR2-locuszoom.png

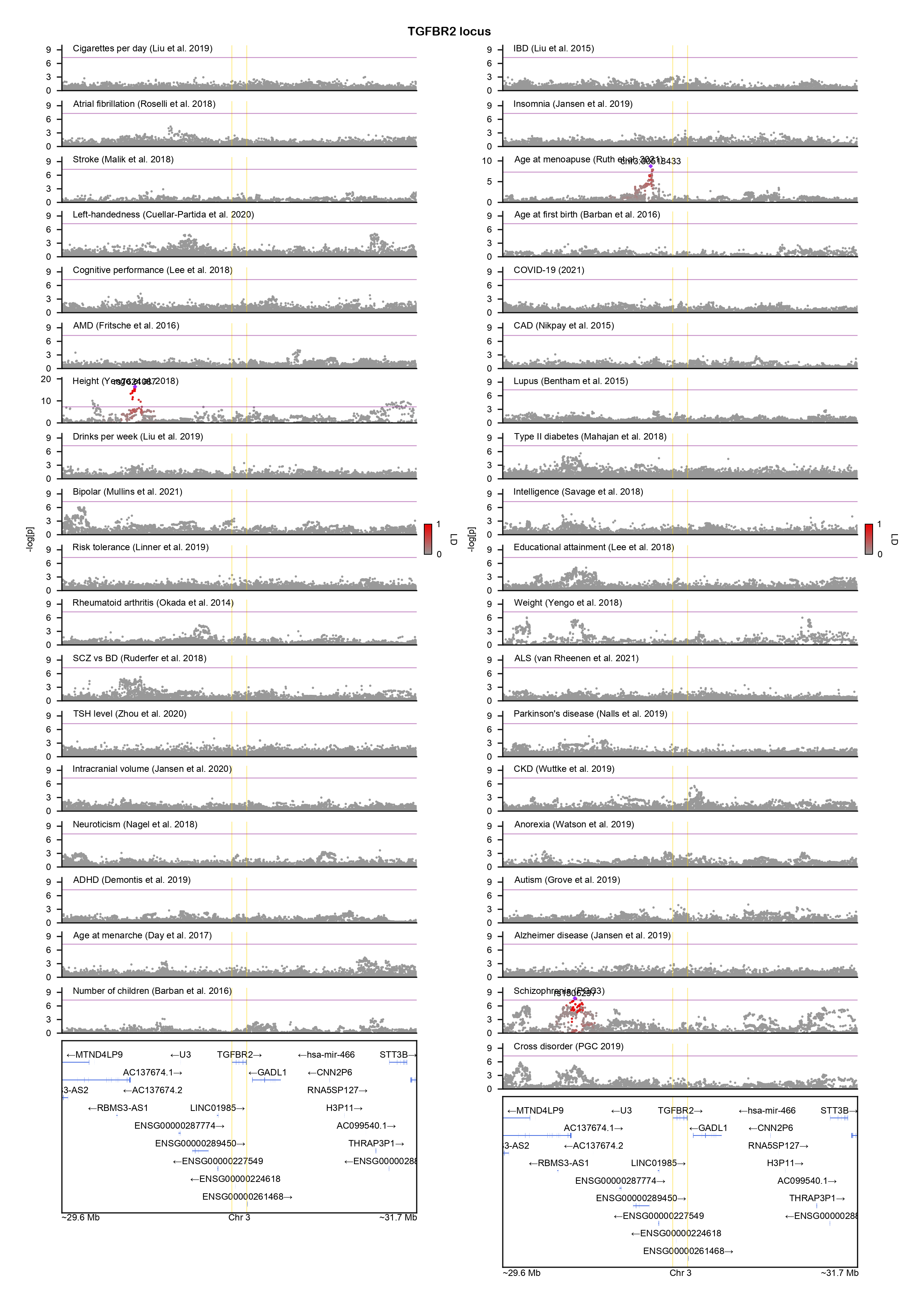

### THEMIS-locuszoom.png

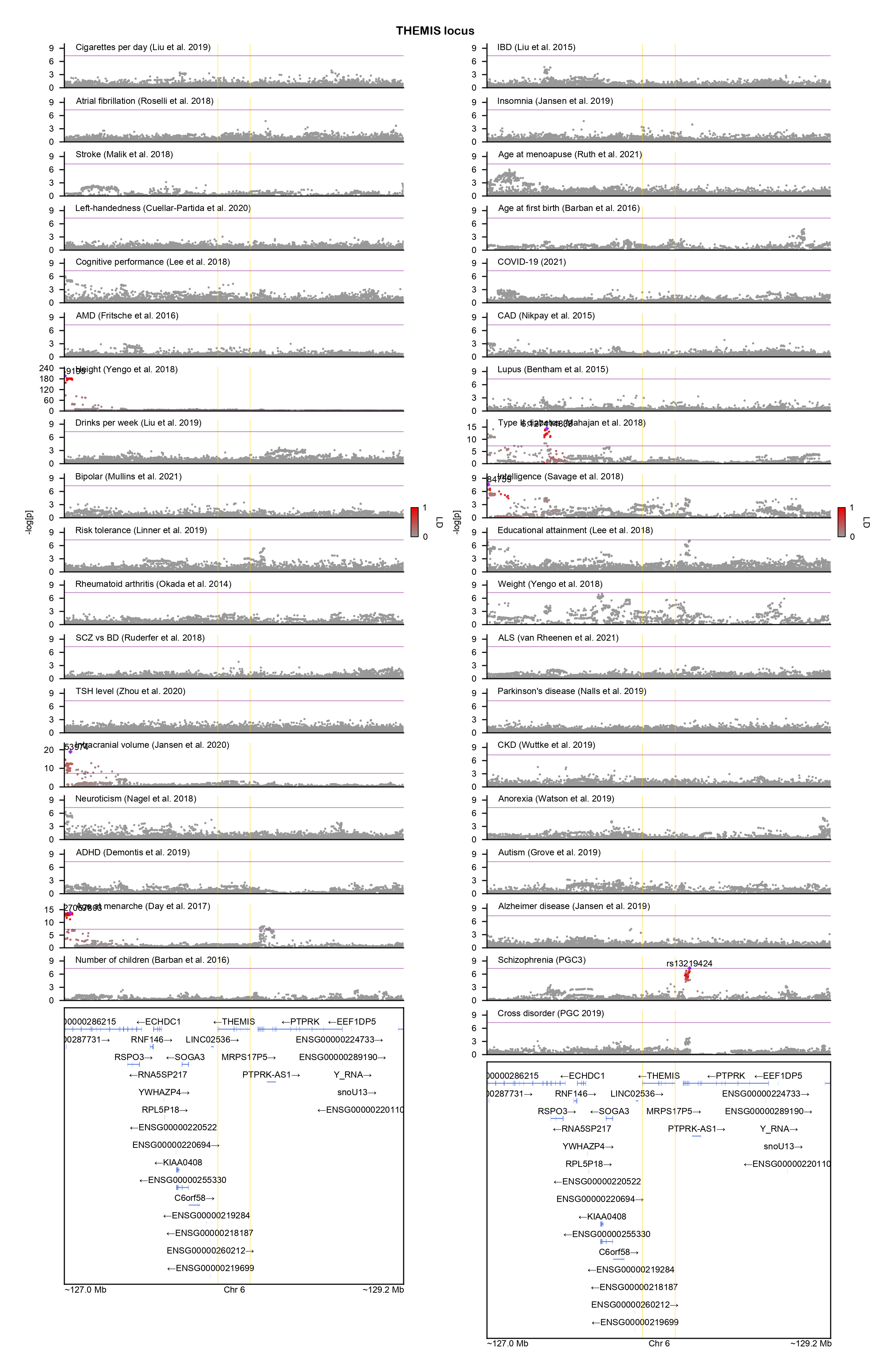

### TMEFF2-locuszoom.png

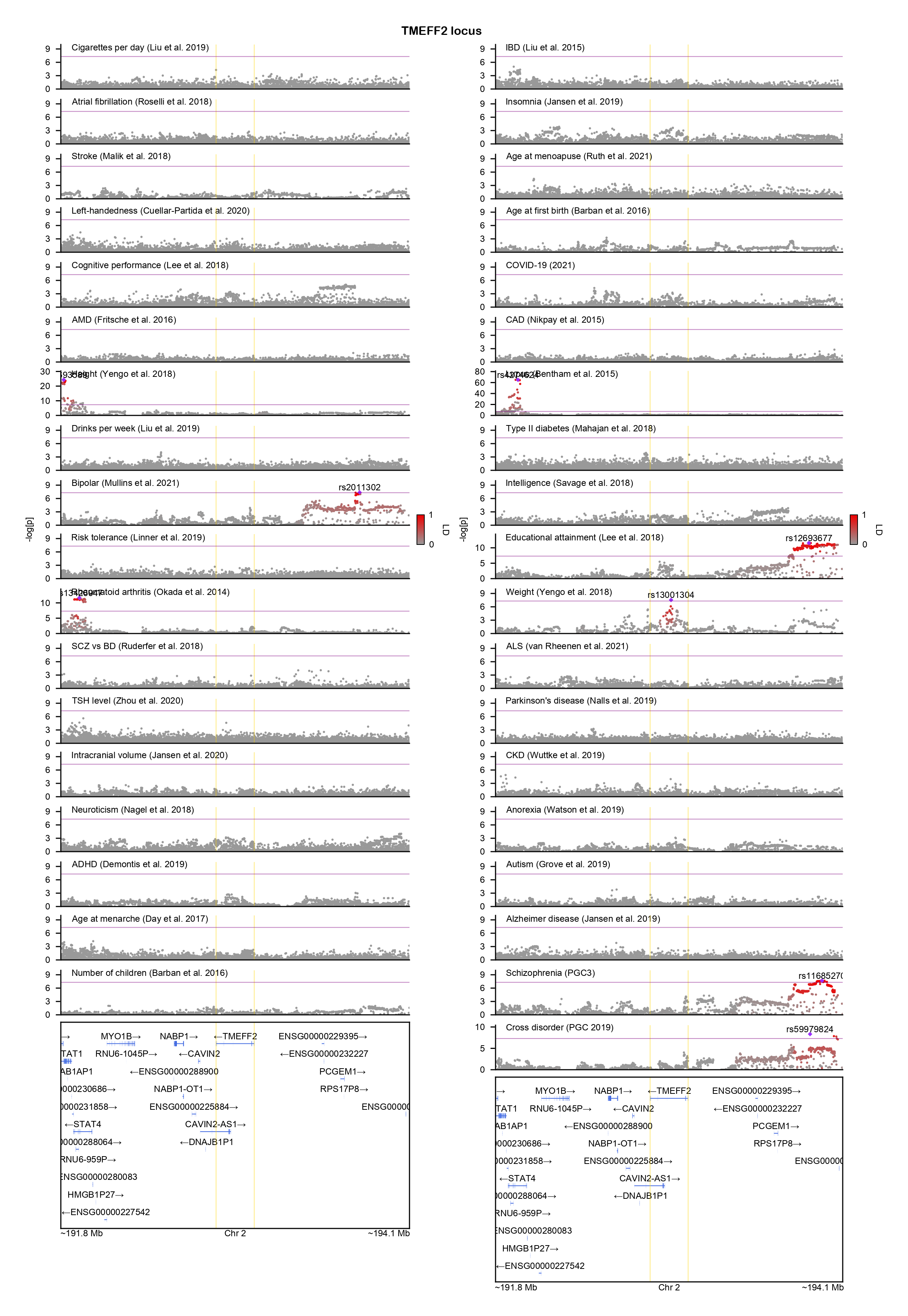

### TMTC1-locuszoom.png

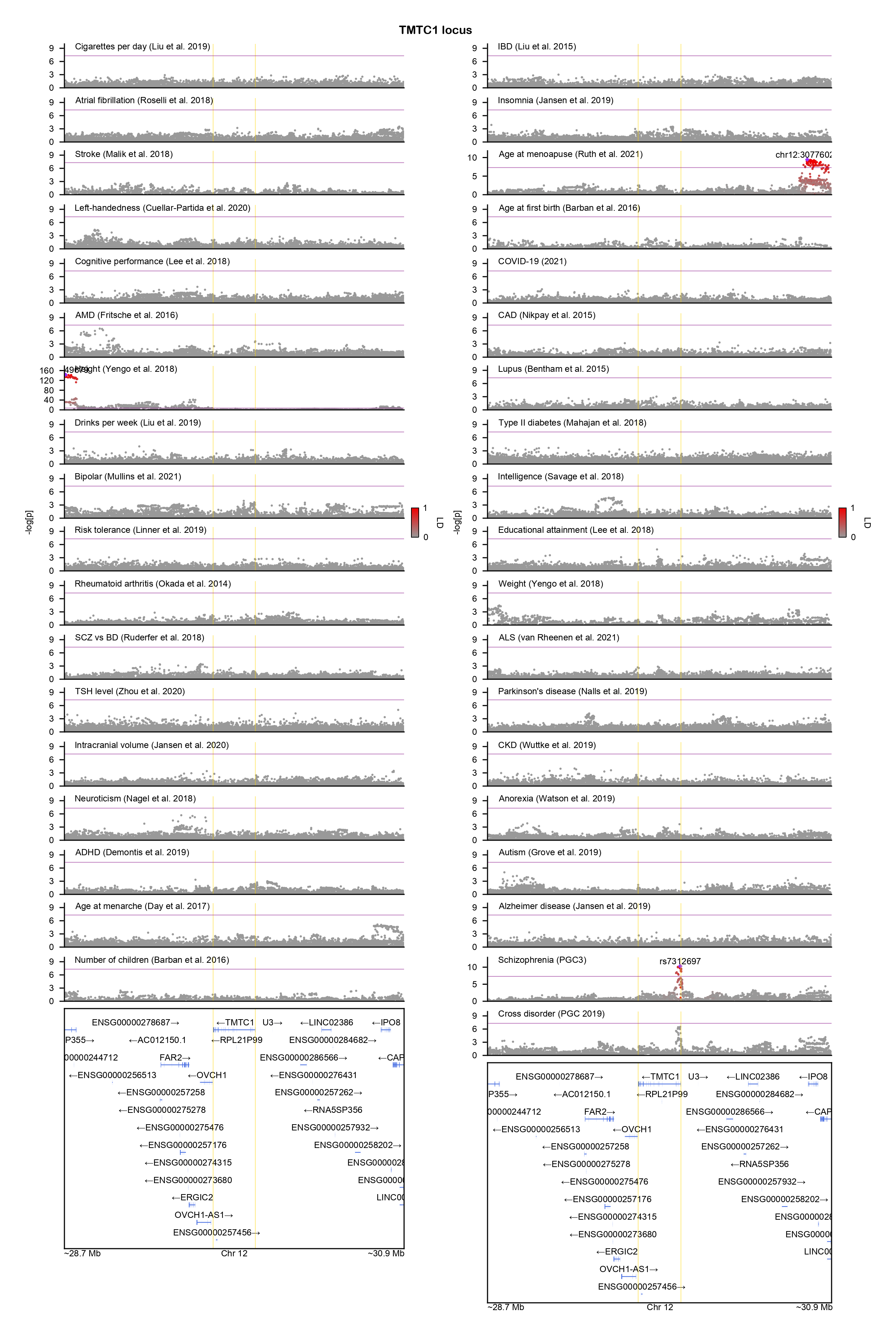

### TSNARE1-locuszoom.png

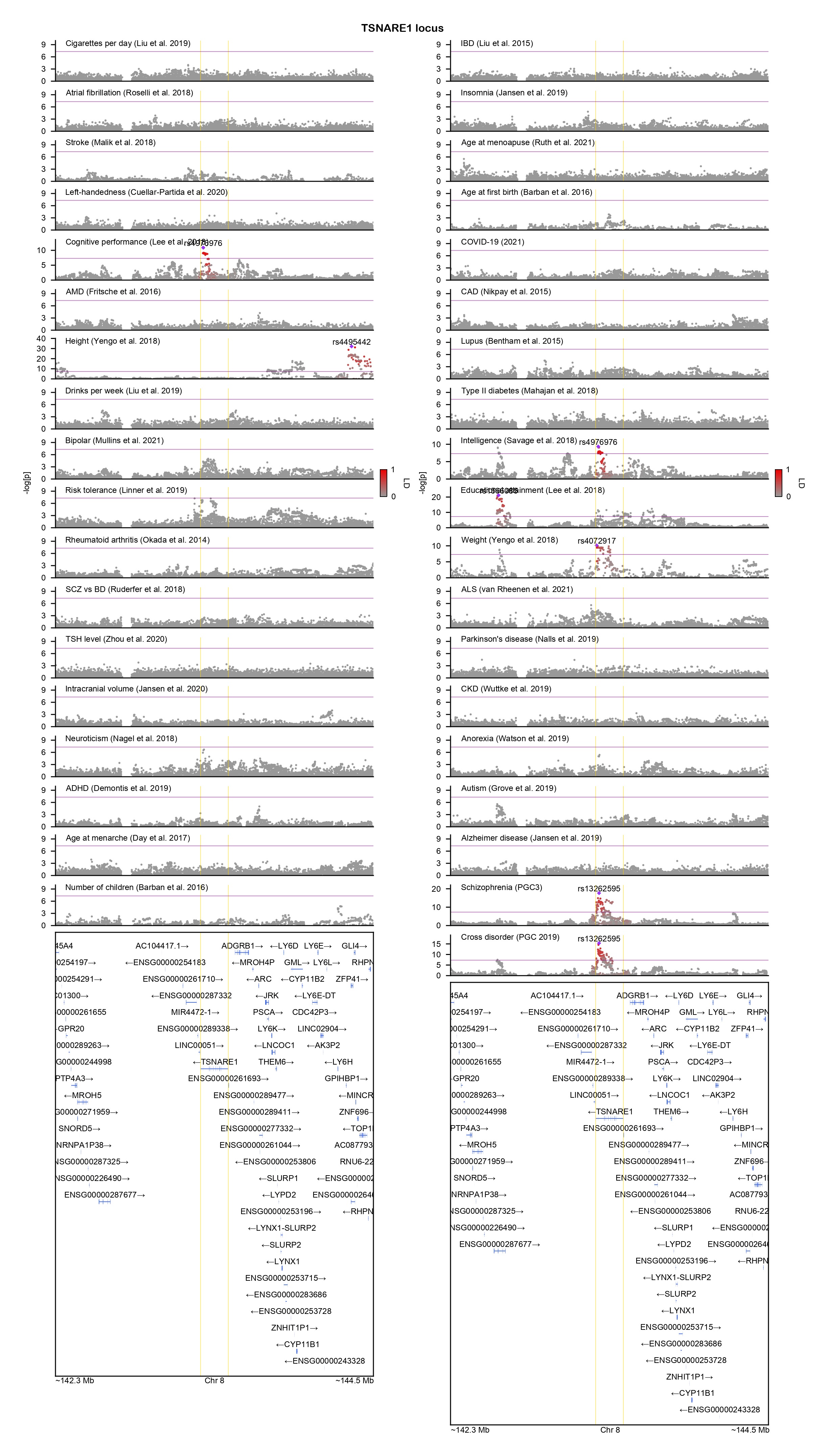

### WDR76-locuszoom.png

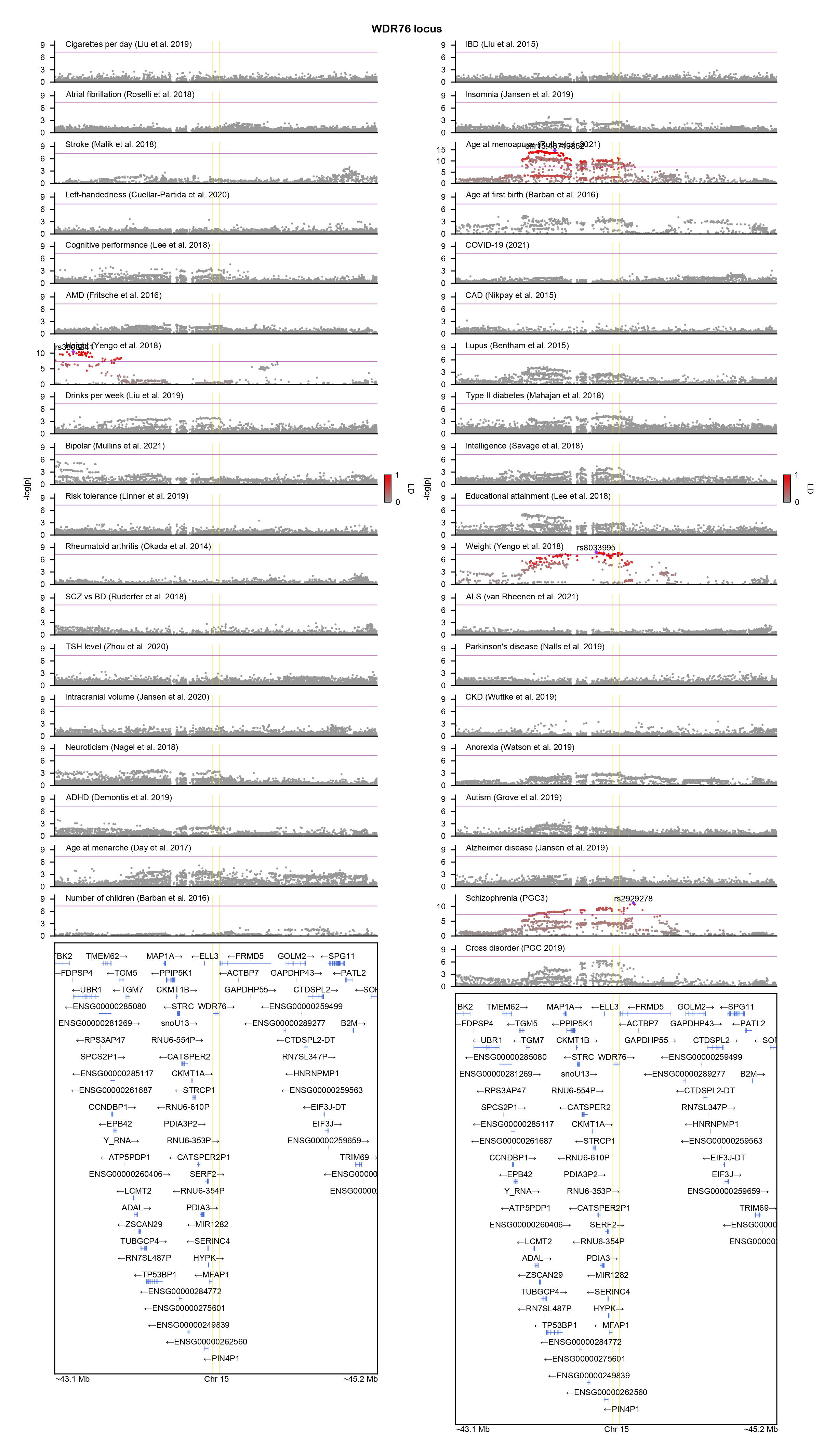

### YTHDF2-locuszoom.png

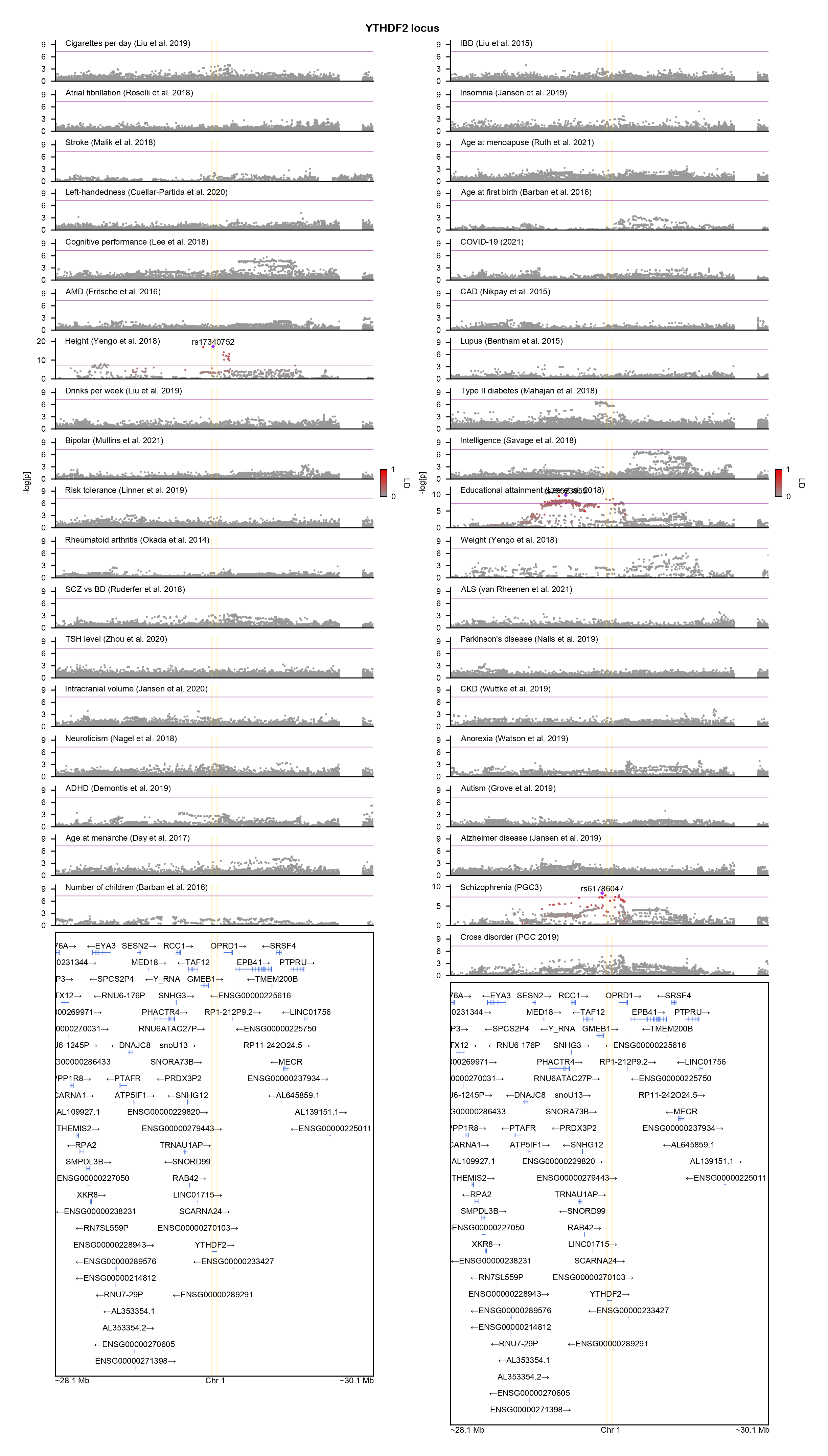

### ZNF184-locuszoom.png

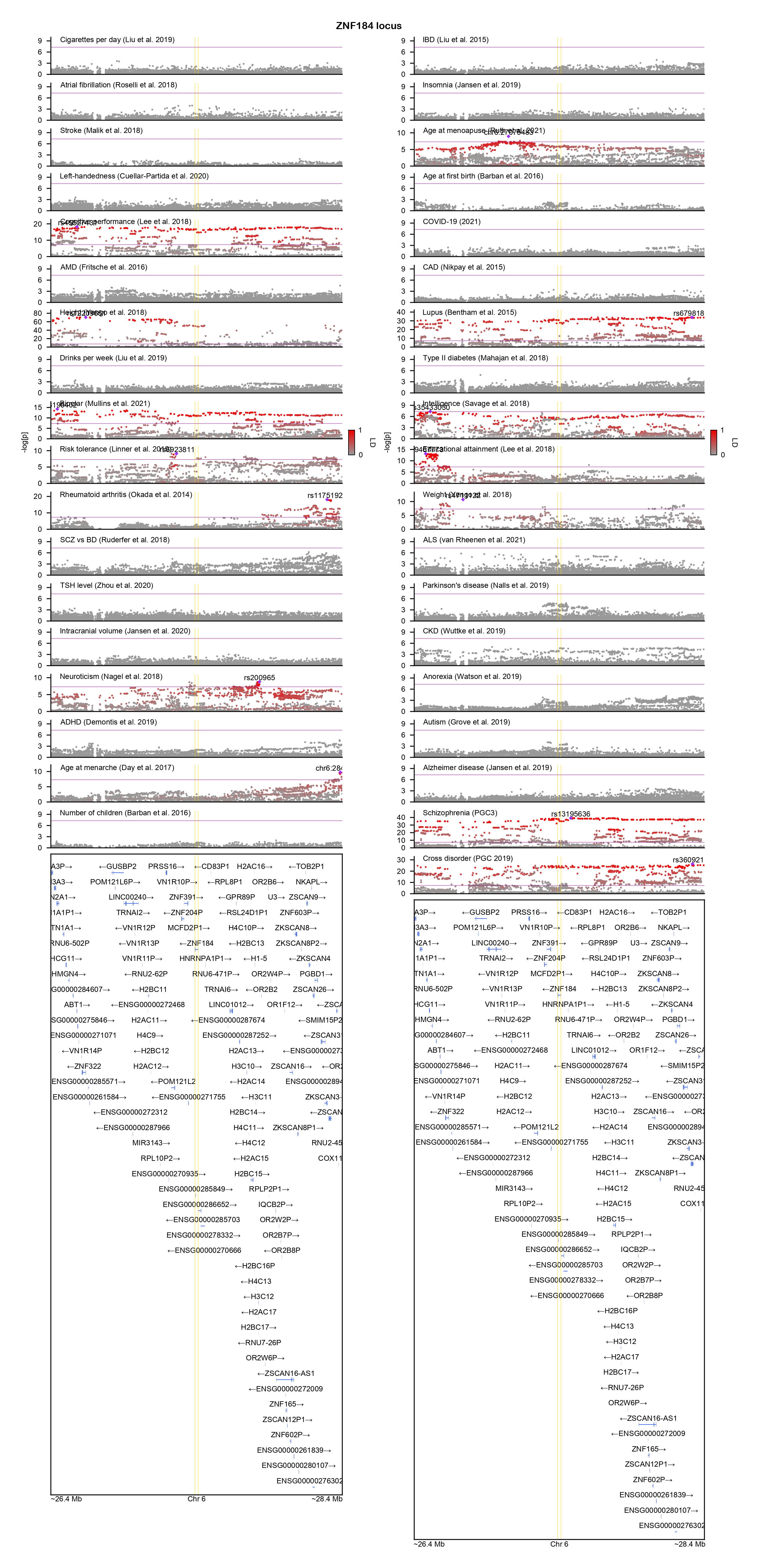

### ZNF536-locuszoom.png

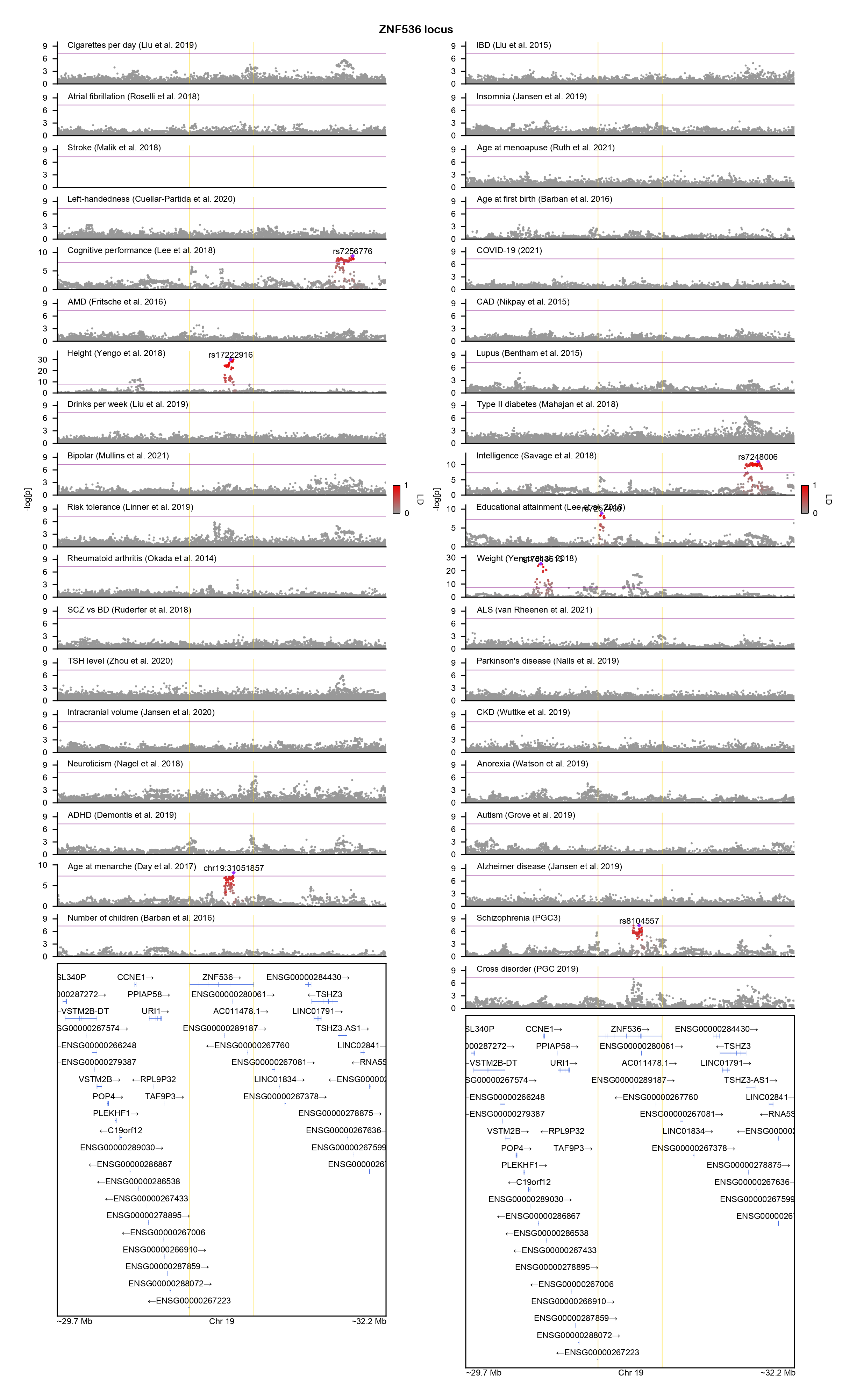

### ZNF823-locuszoom.png

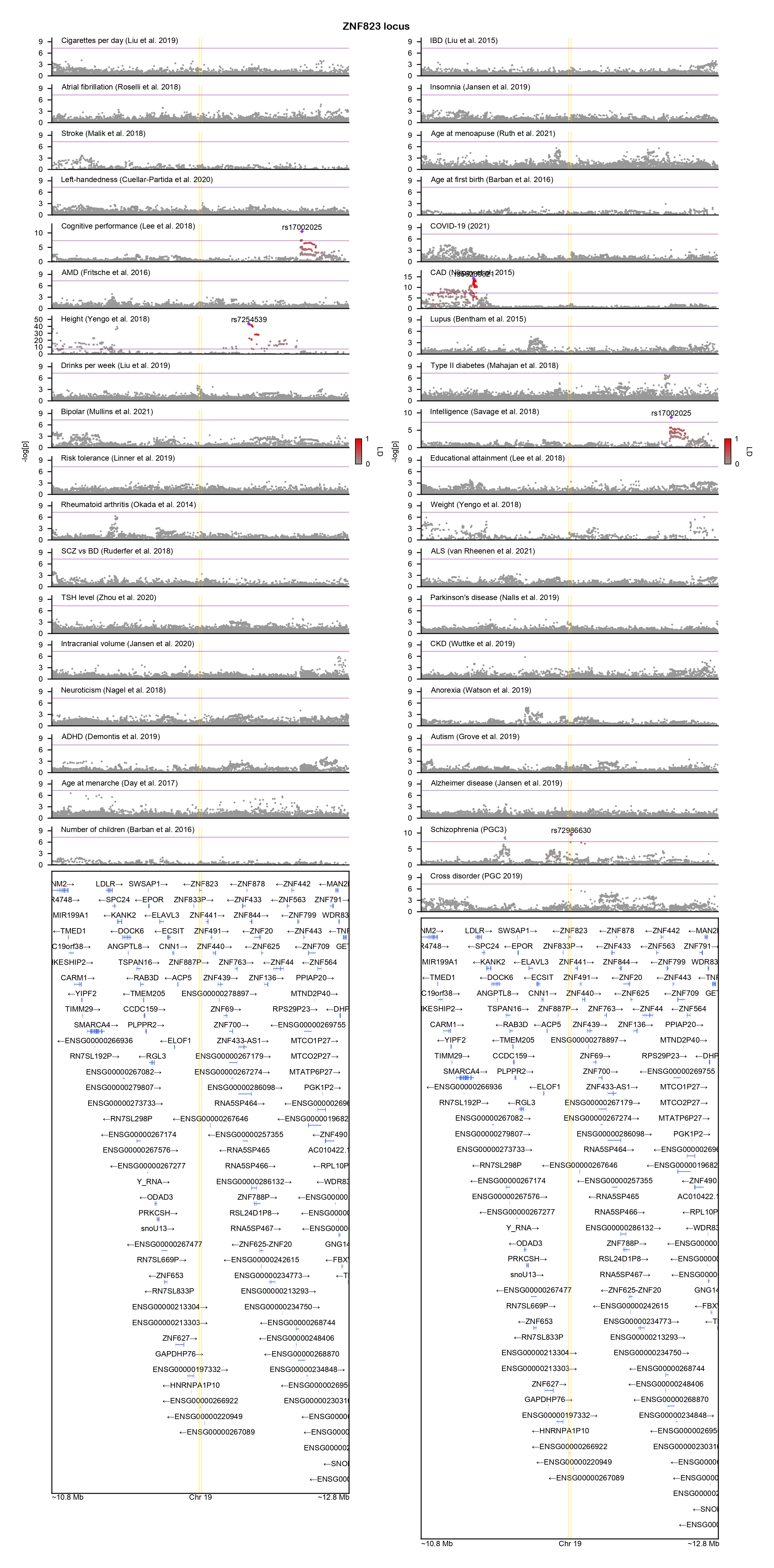
