## Supplementary figures and images for "GeneticsMakie.jl: A versatile and scalable toolkit for visualizing locus-level genetic and genomic data"

### ANKRD11-locuszoom.png

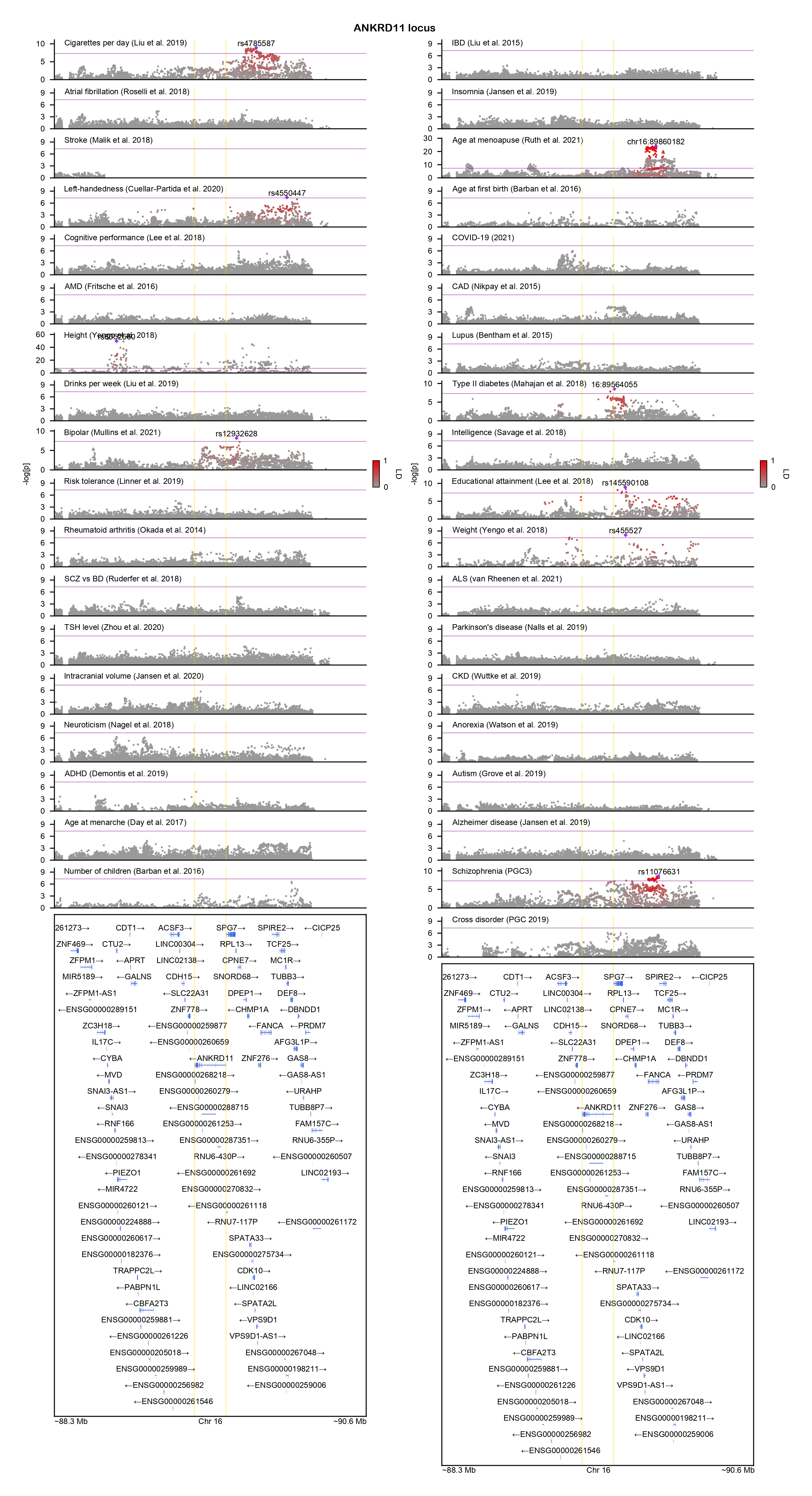

### ANKRD63-locuszoom.png

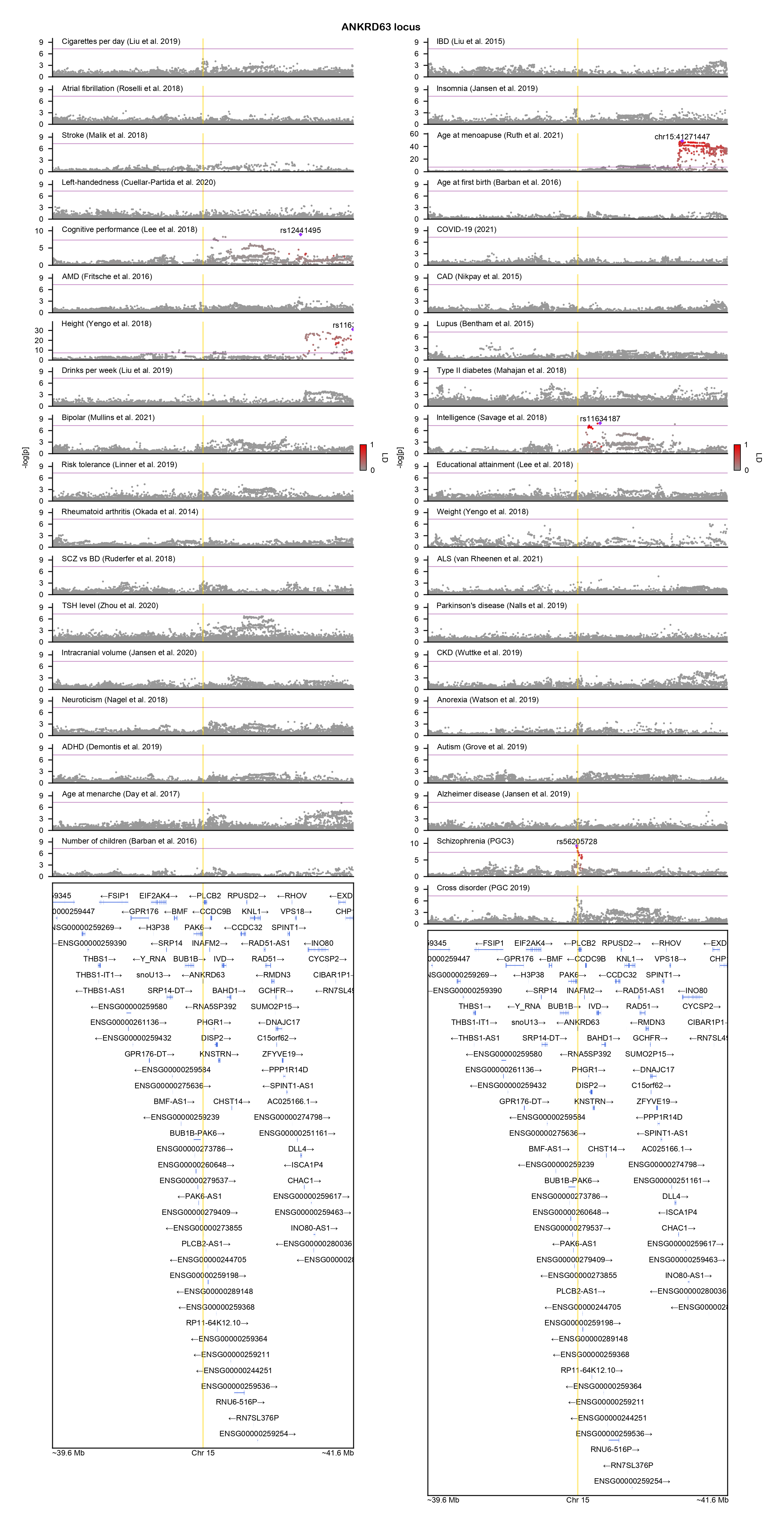

### ASXL1-locuszoom.png

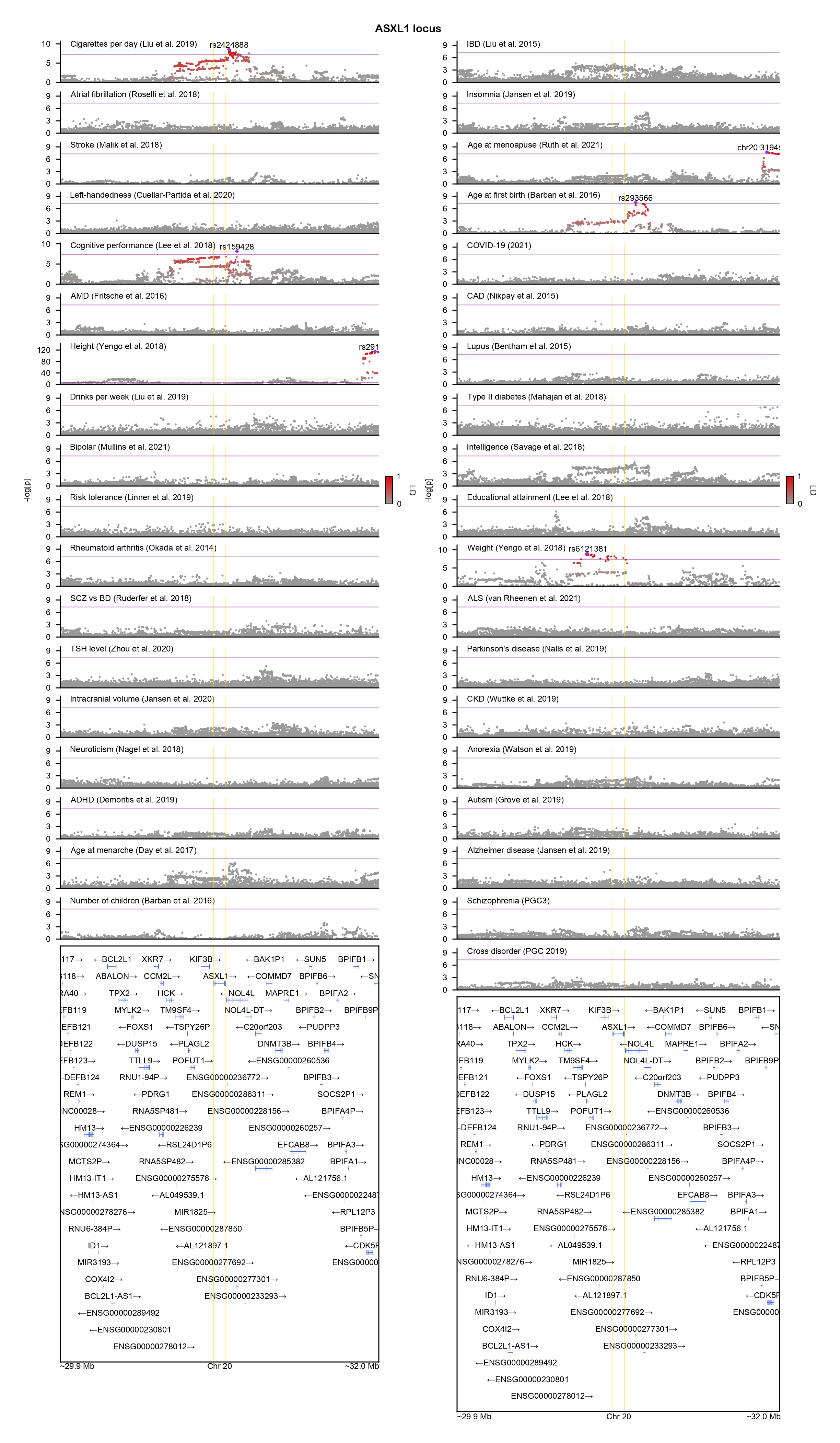

### ATP6V1A-locuszoom.png

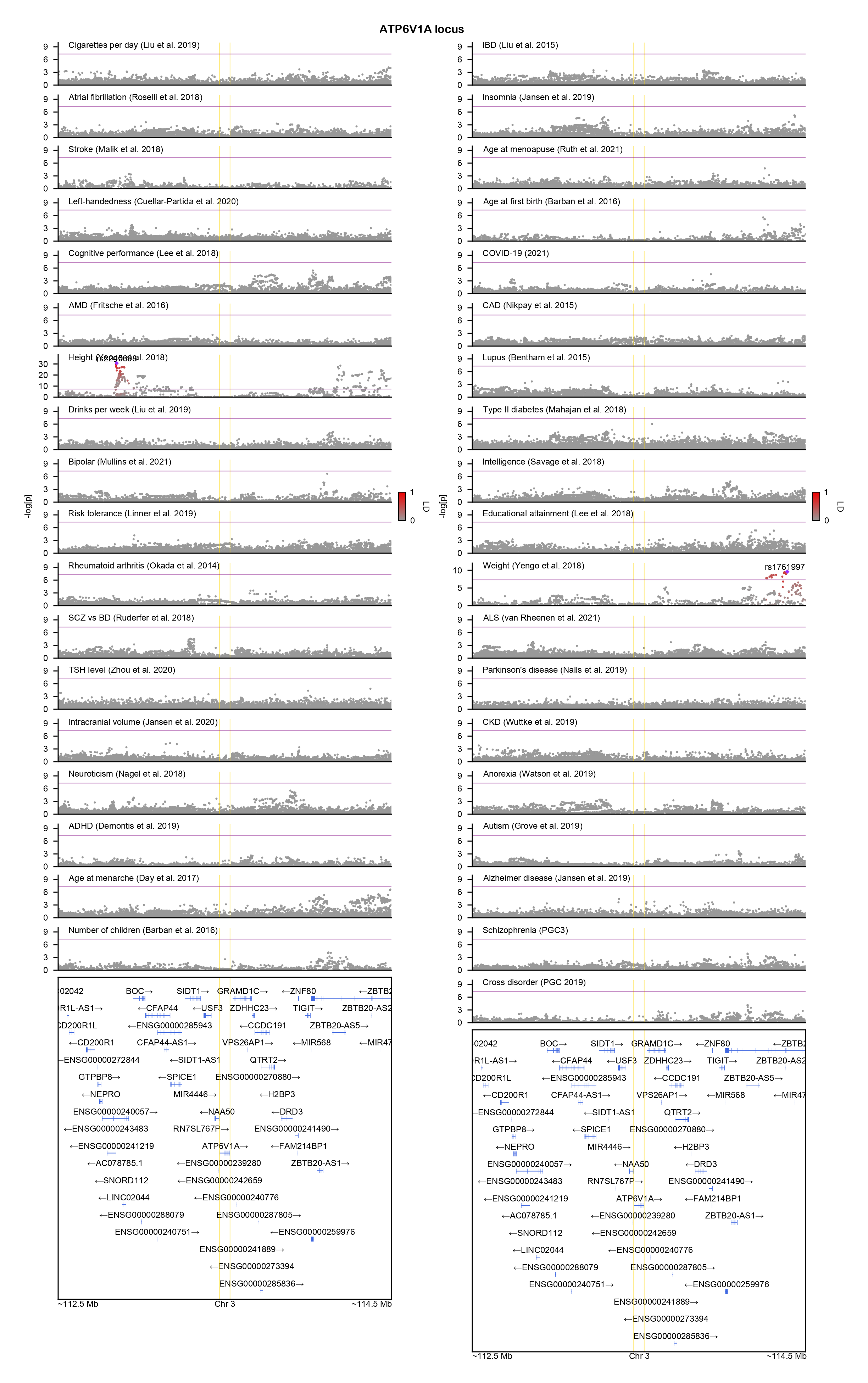

### CAMTA1-locuszoom.png

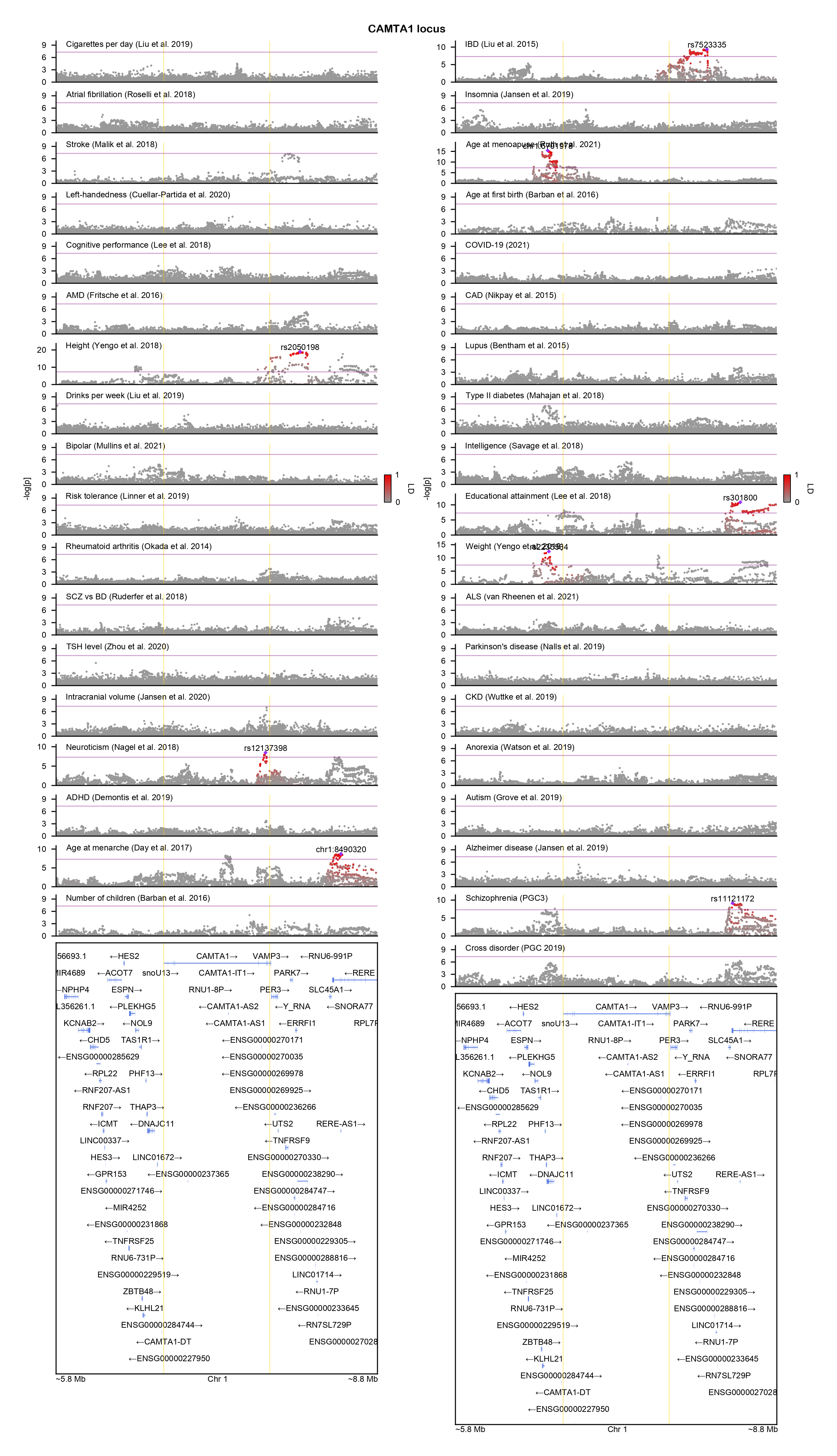

### CASK-locuszoom.png

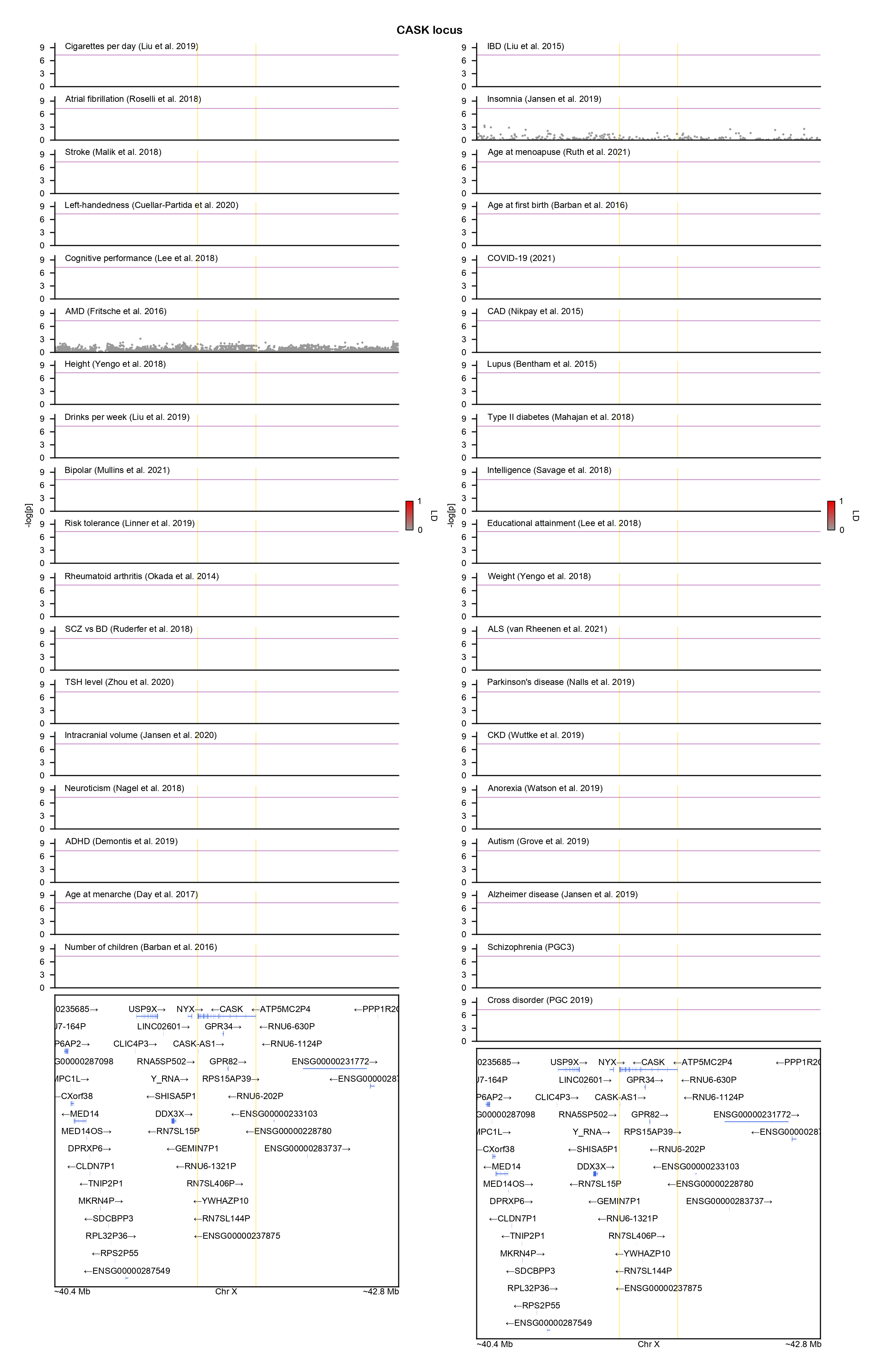

### CCDC182-locuszoom.png

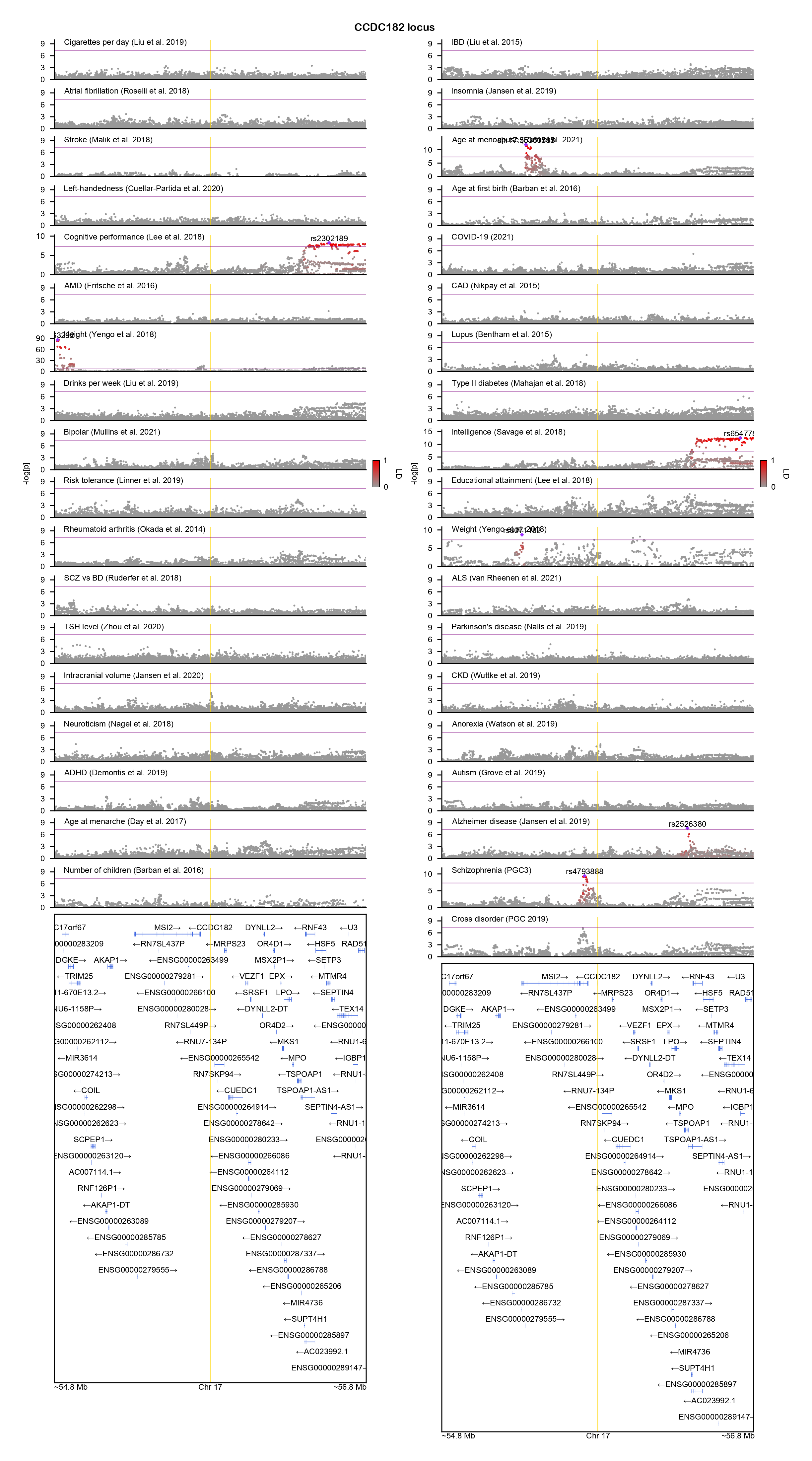

### CELF4-locuszoom.png

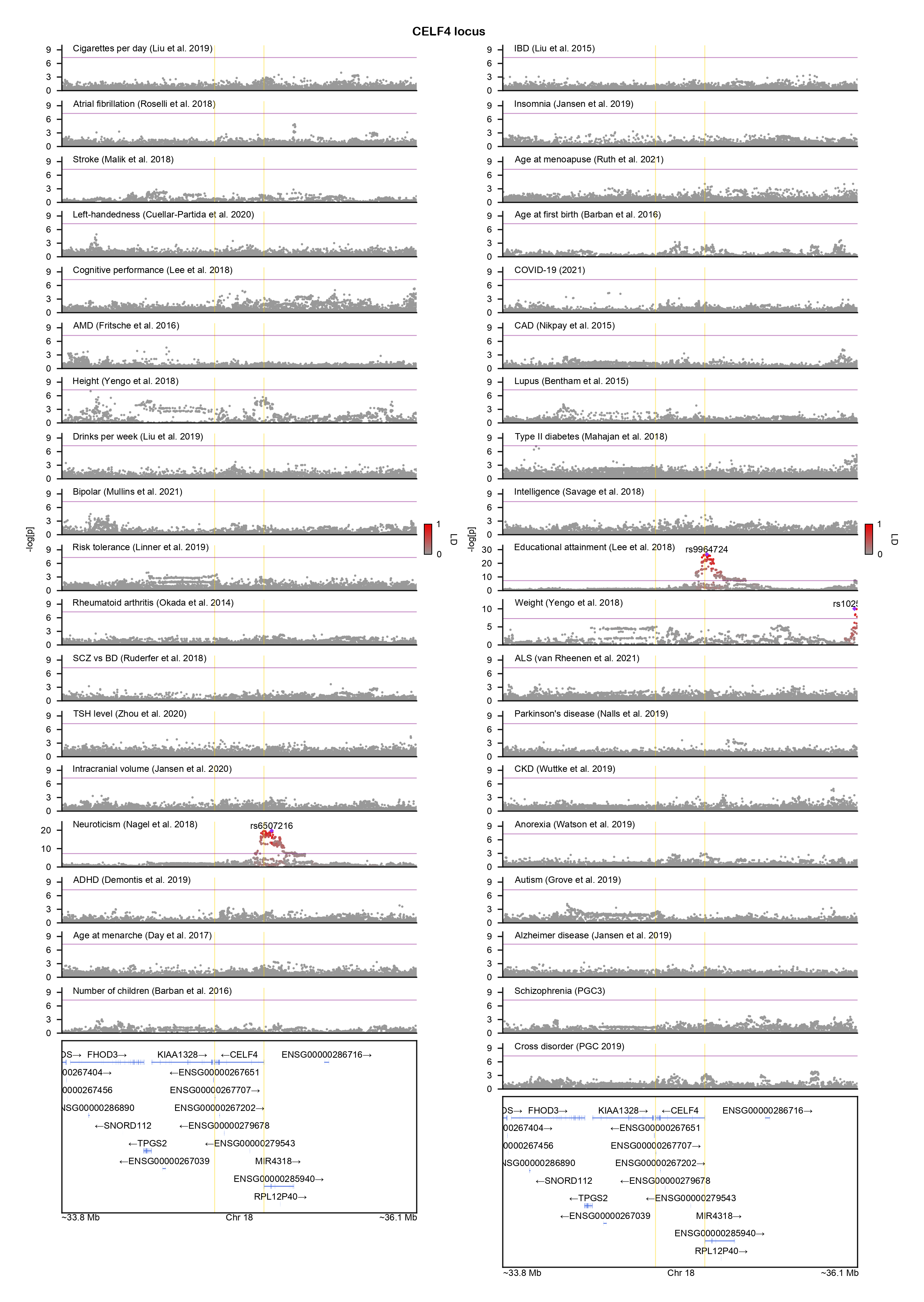

### CHAMP1-locuszoom.png

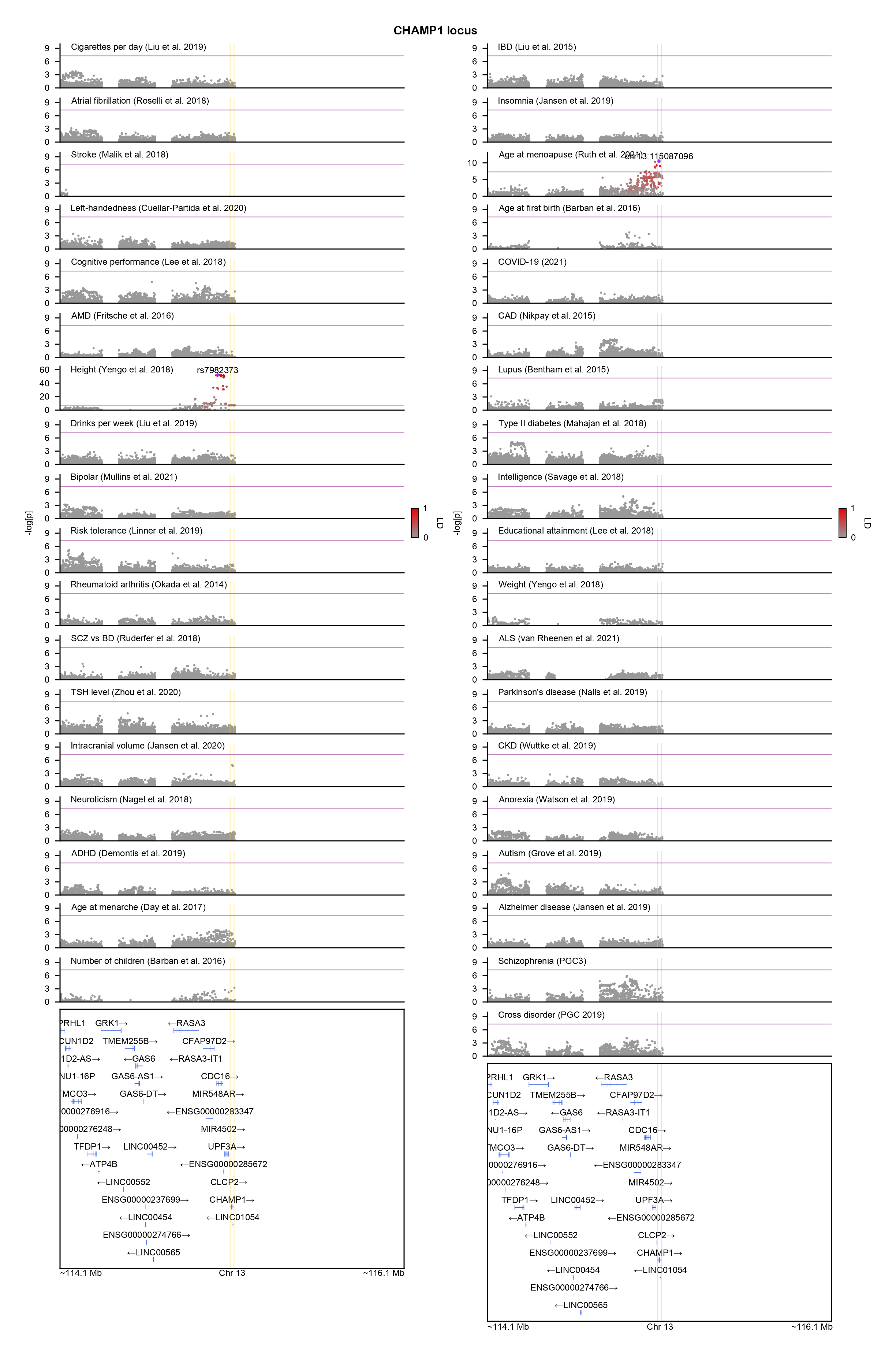

### CHD2-locuszoom.png

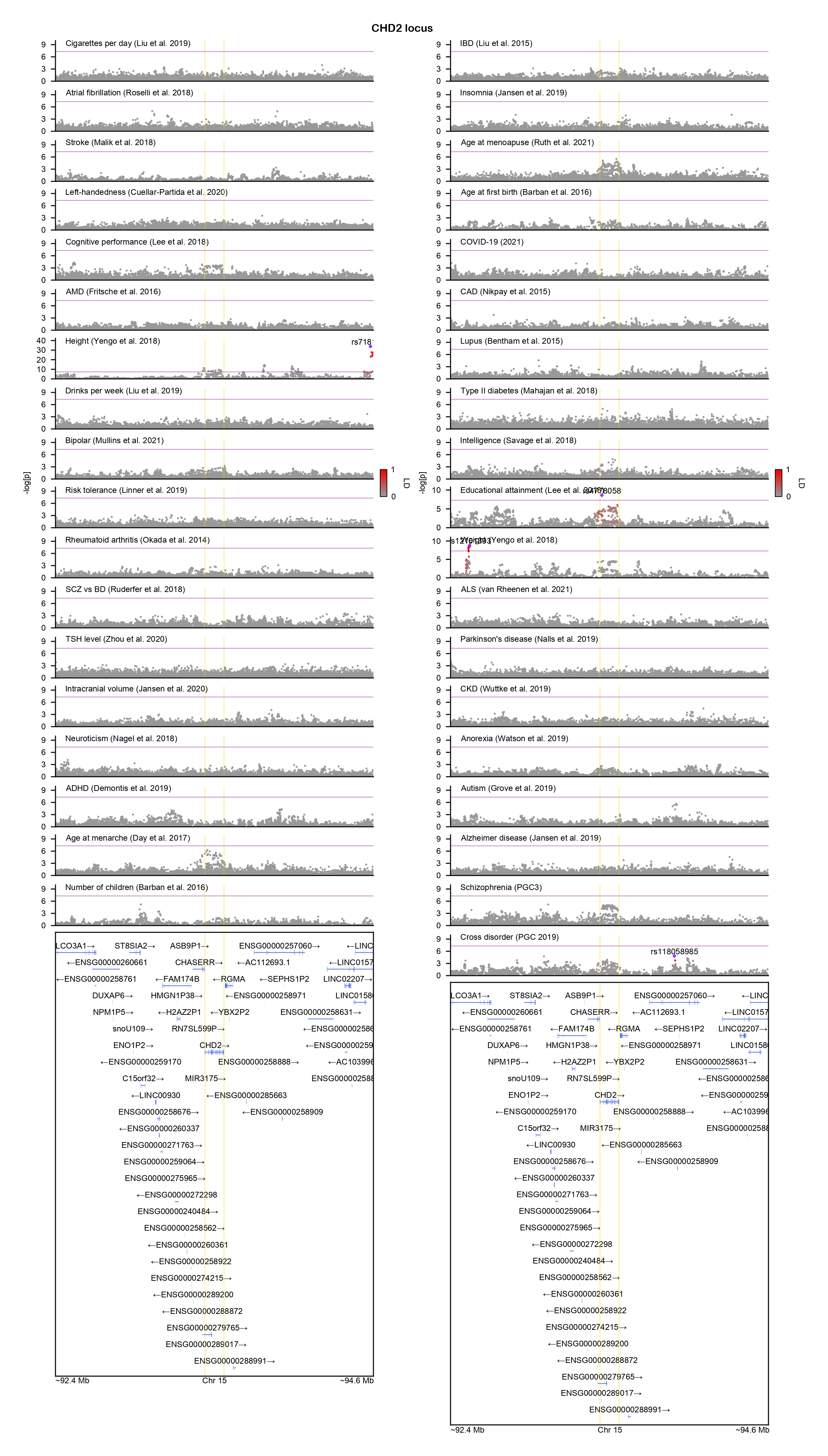

### CHRNA5-locuszoom.png

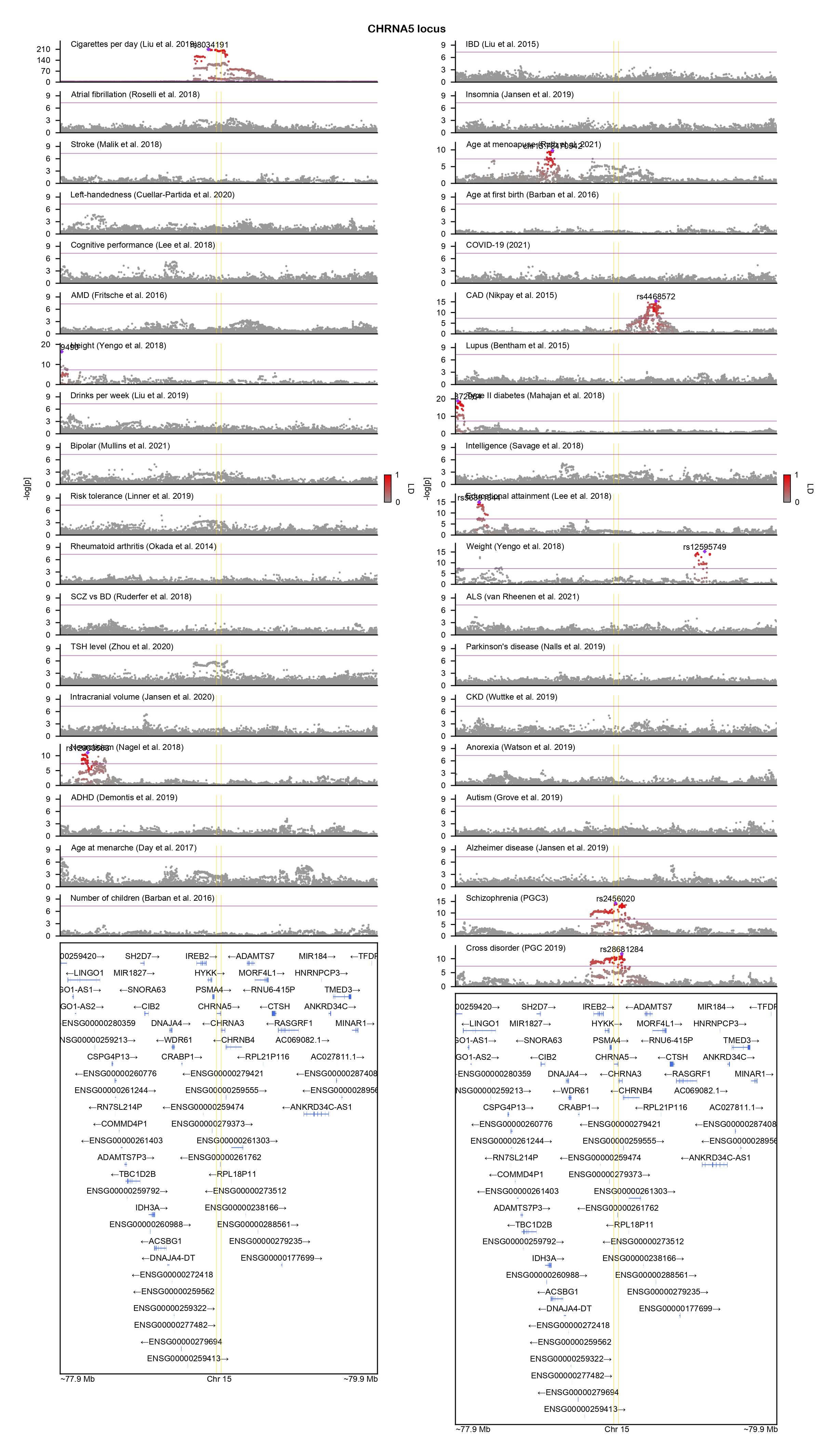

### COL6A1-locuszoom.png

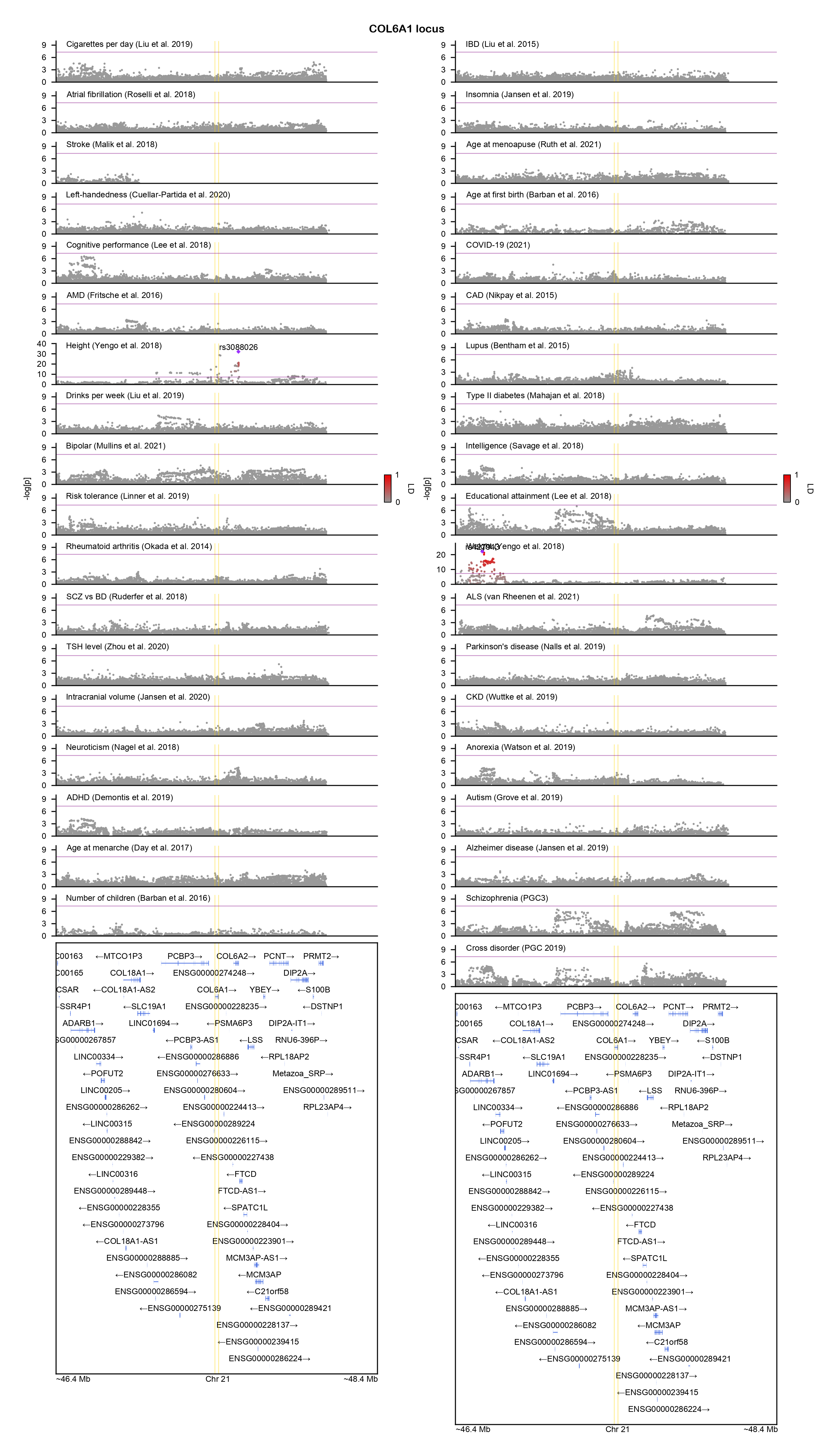

### CSNK2A1-locuszoom.png

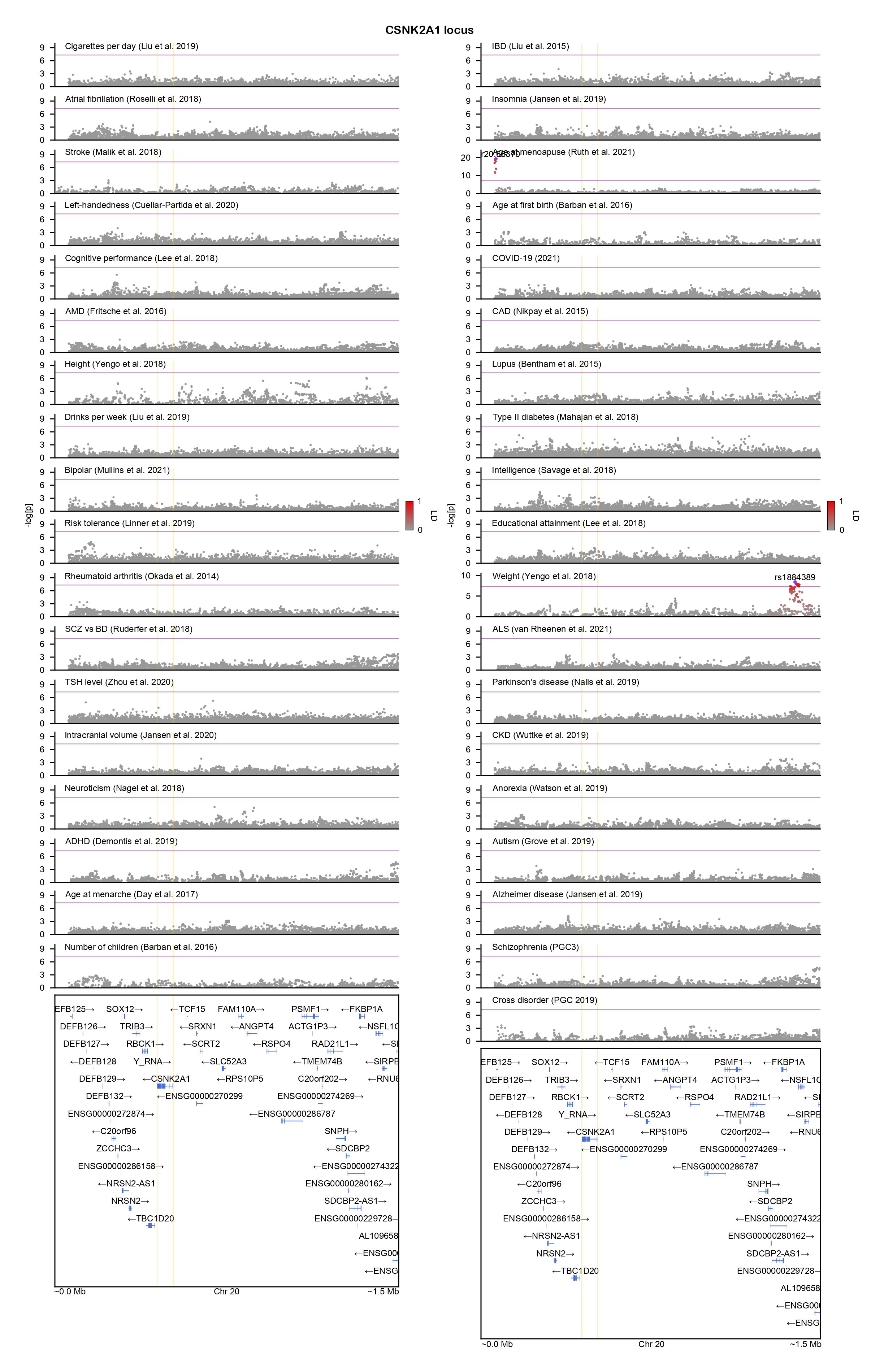

### CTNNB1-locuszoom.png

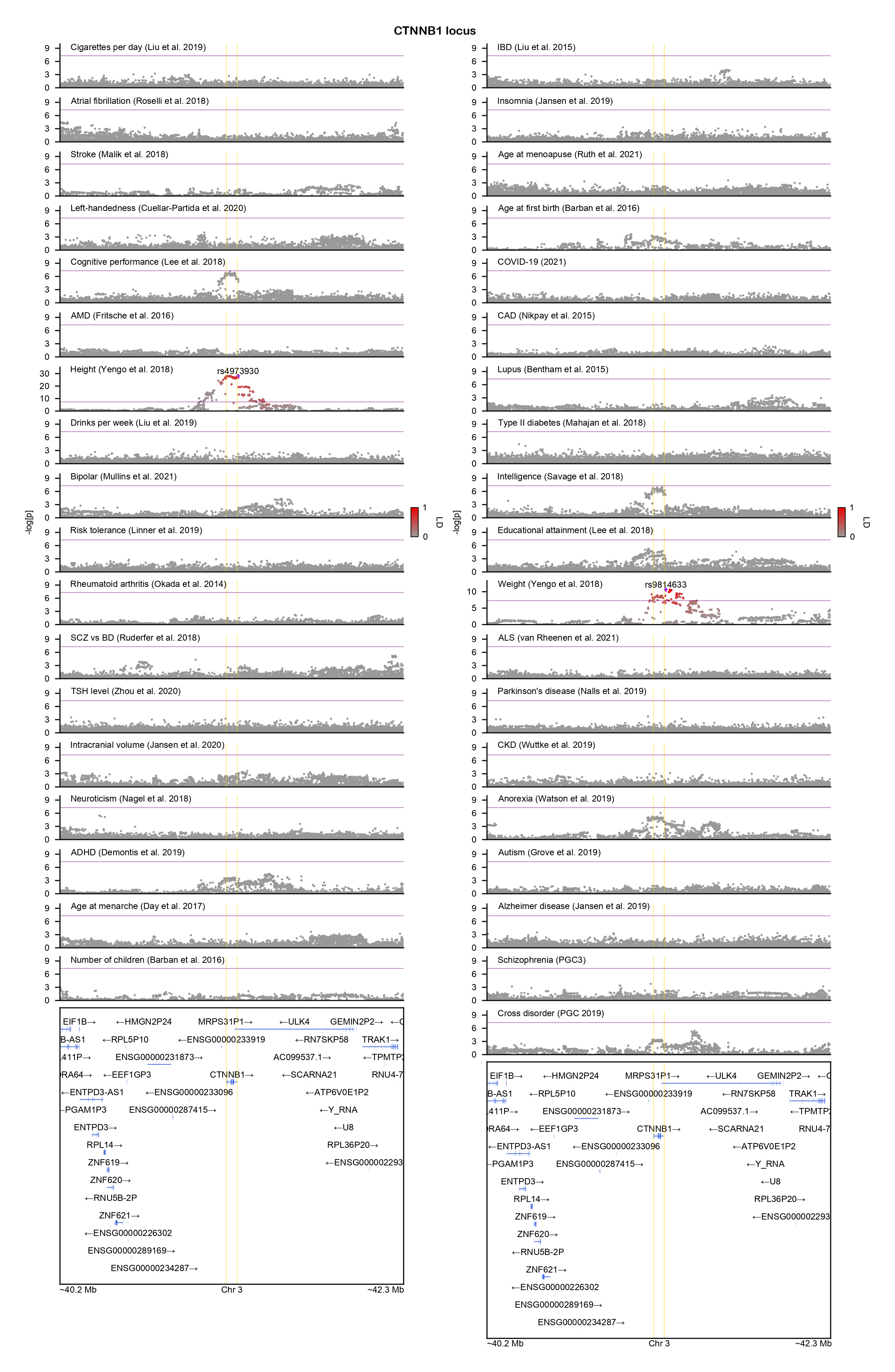

### CUL1-locuszoom.png

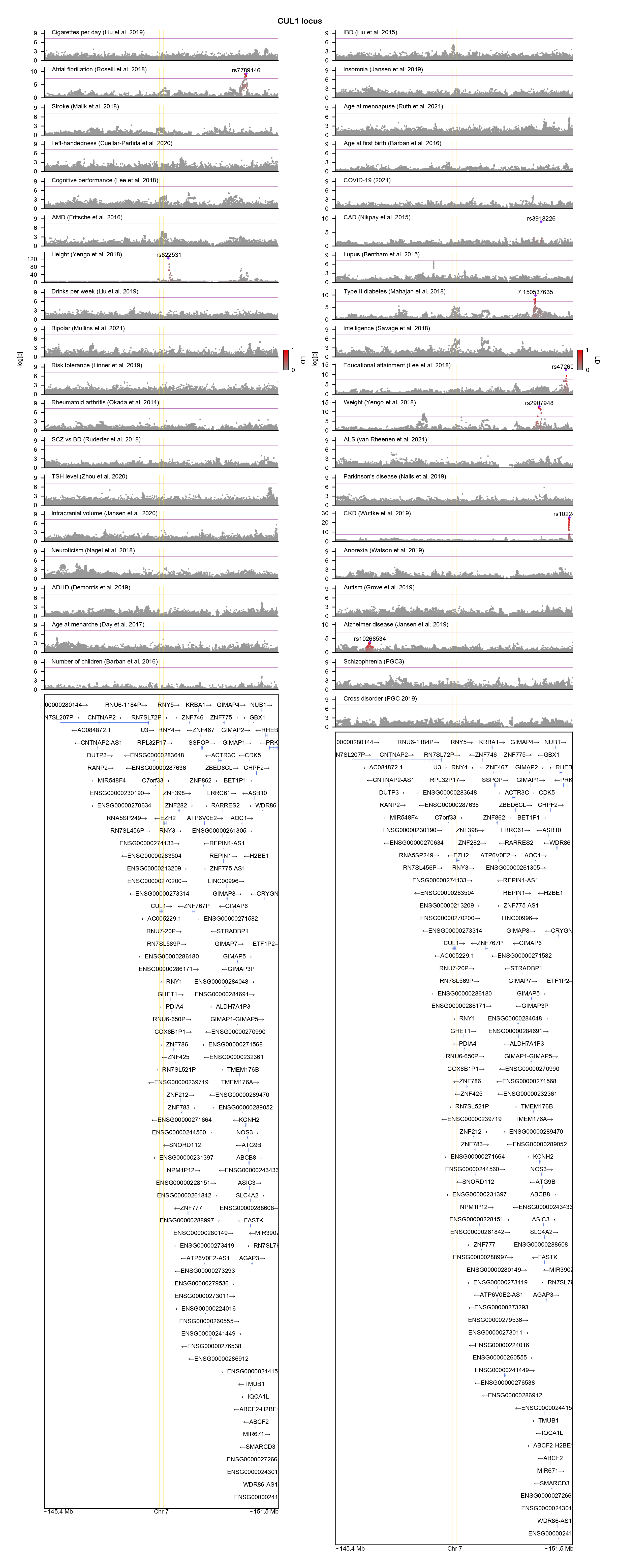

### CUL3-locuszoom.png

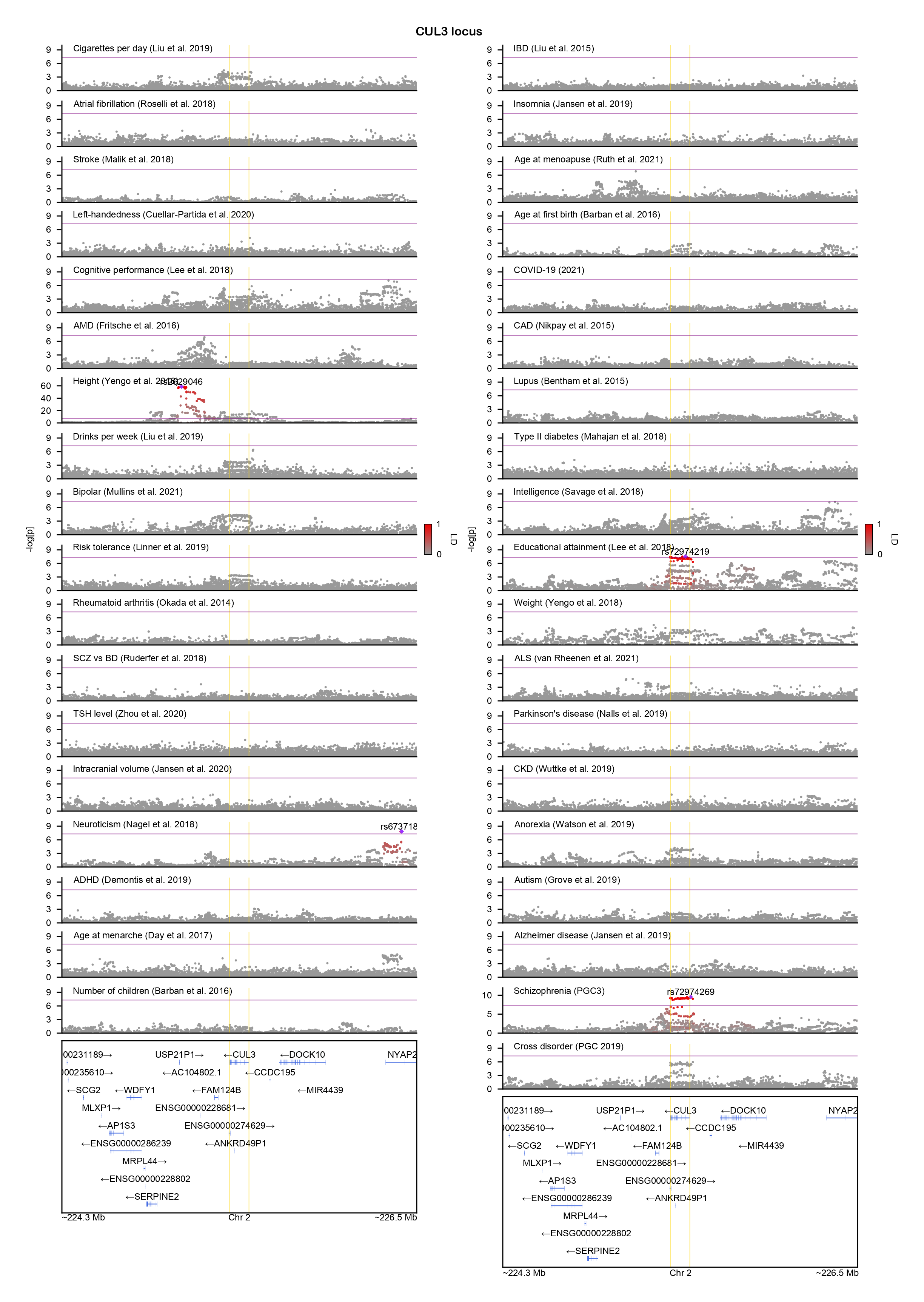

### DCC-locuszoom.png

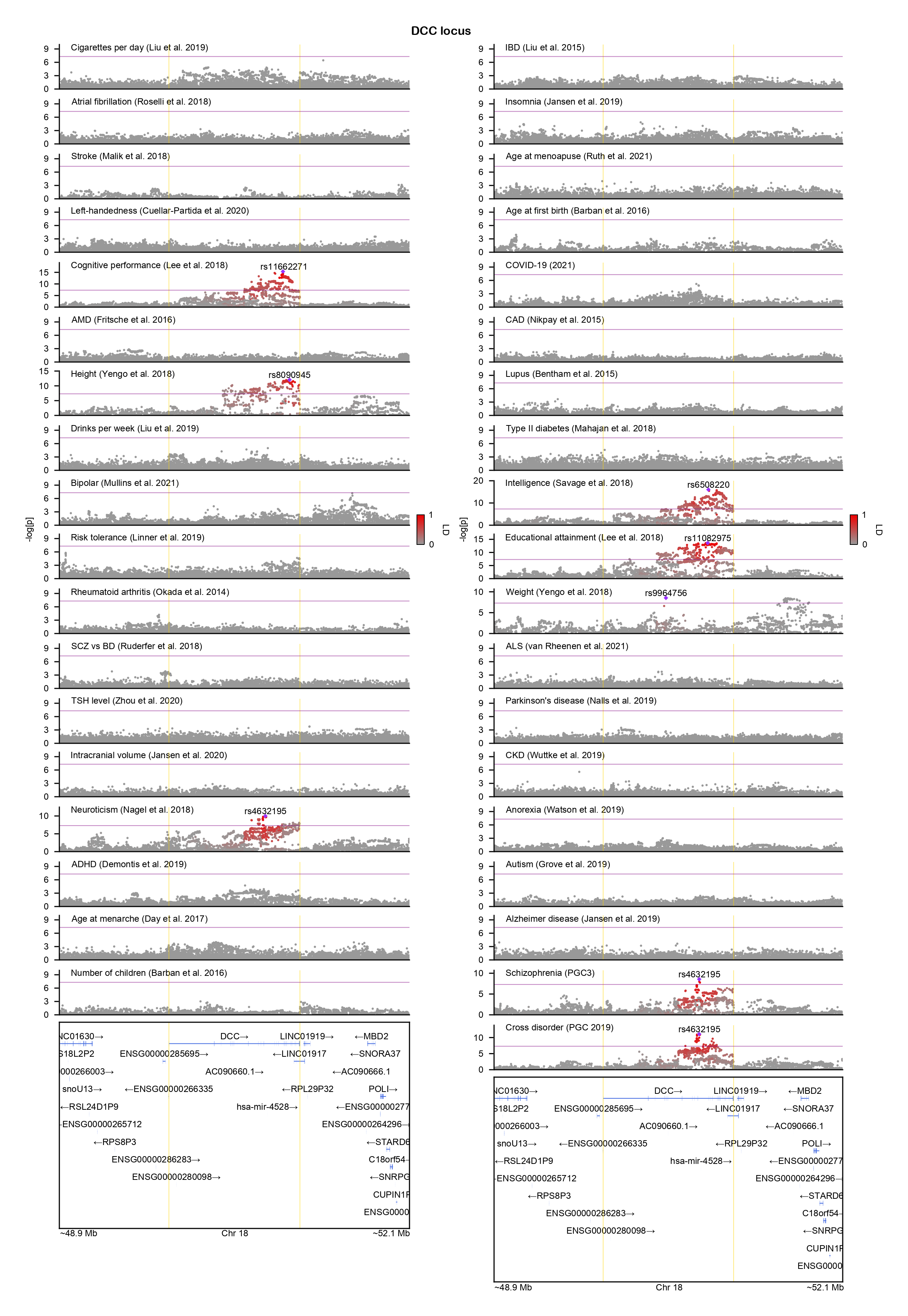

### DEAF1-locuszoom.png

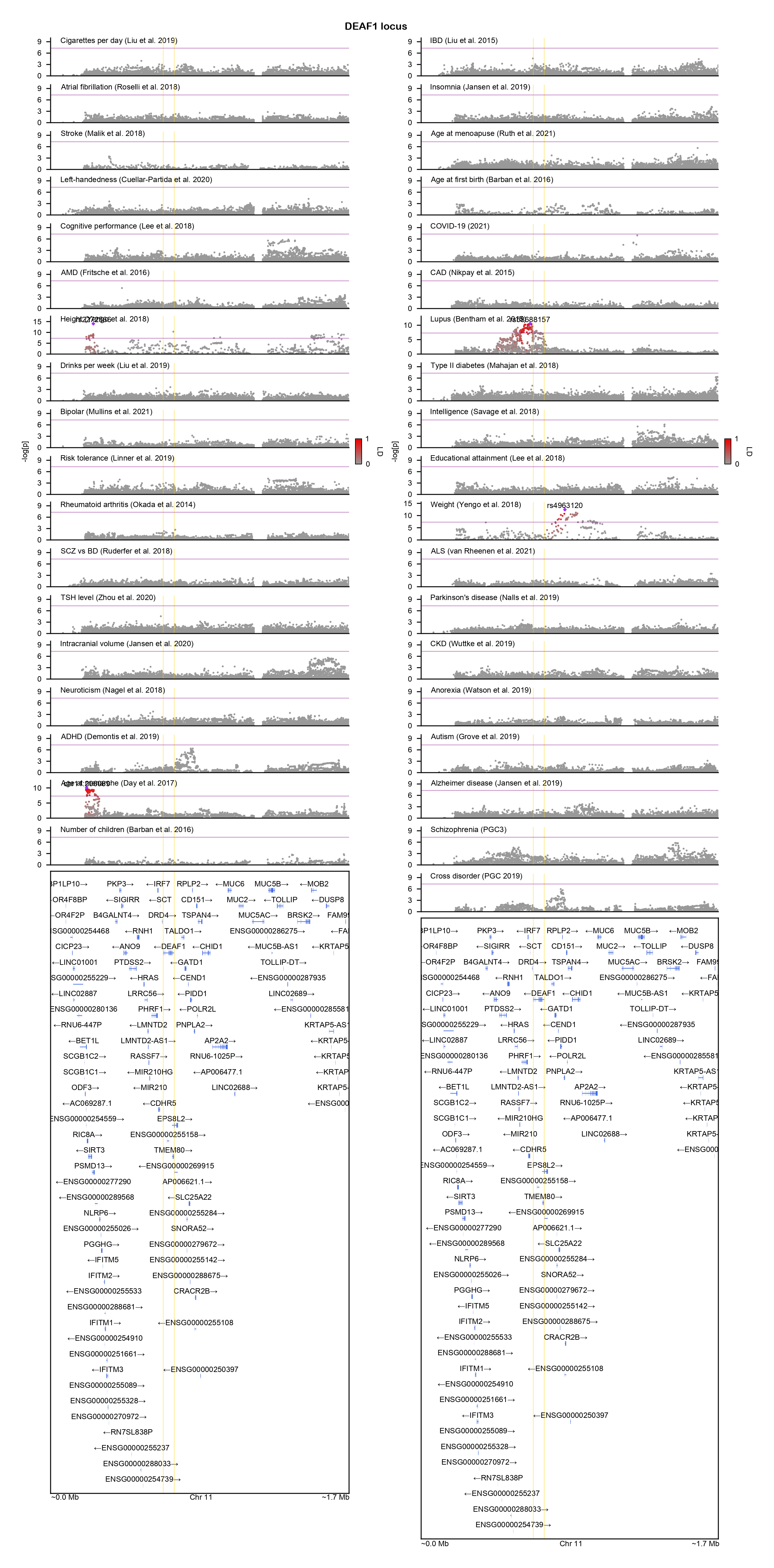

### DNAJC19-locuszoom.png

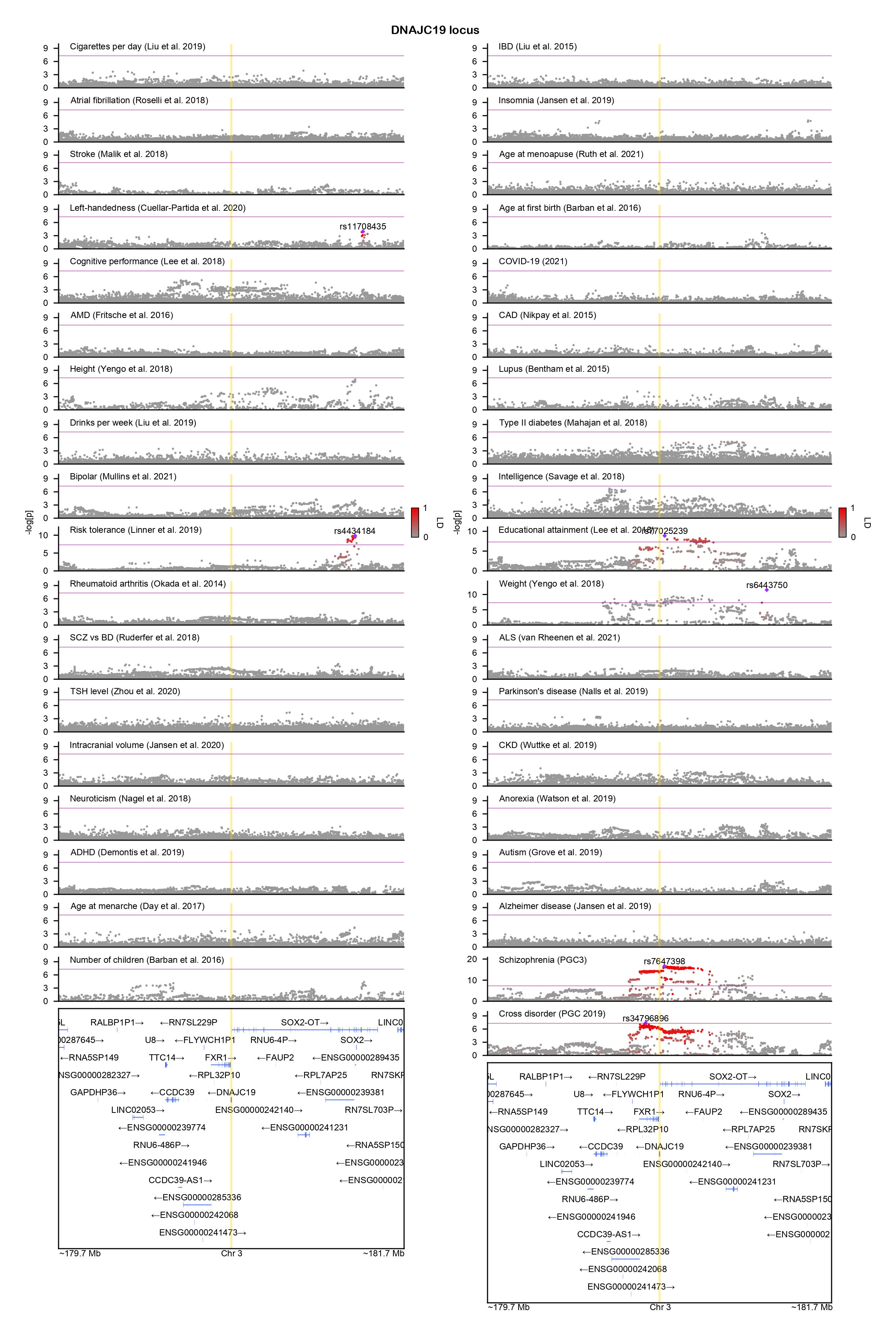

### DPYSL2-locuszoom.png

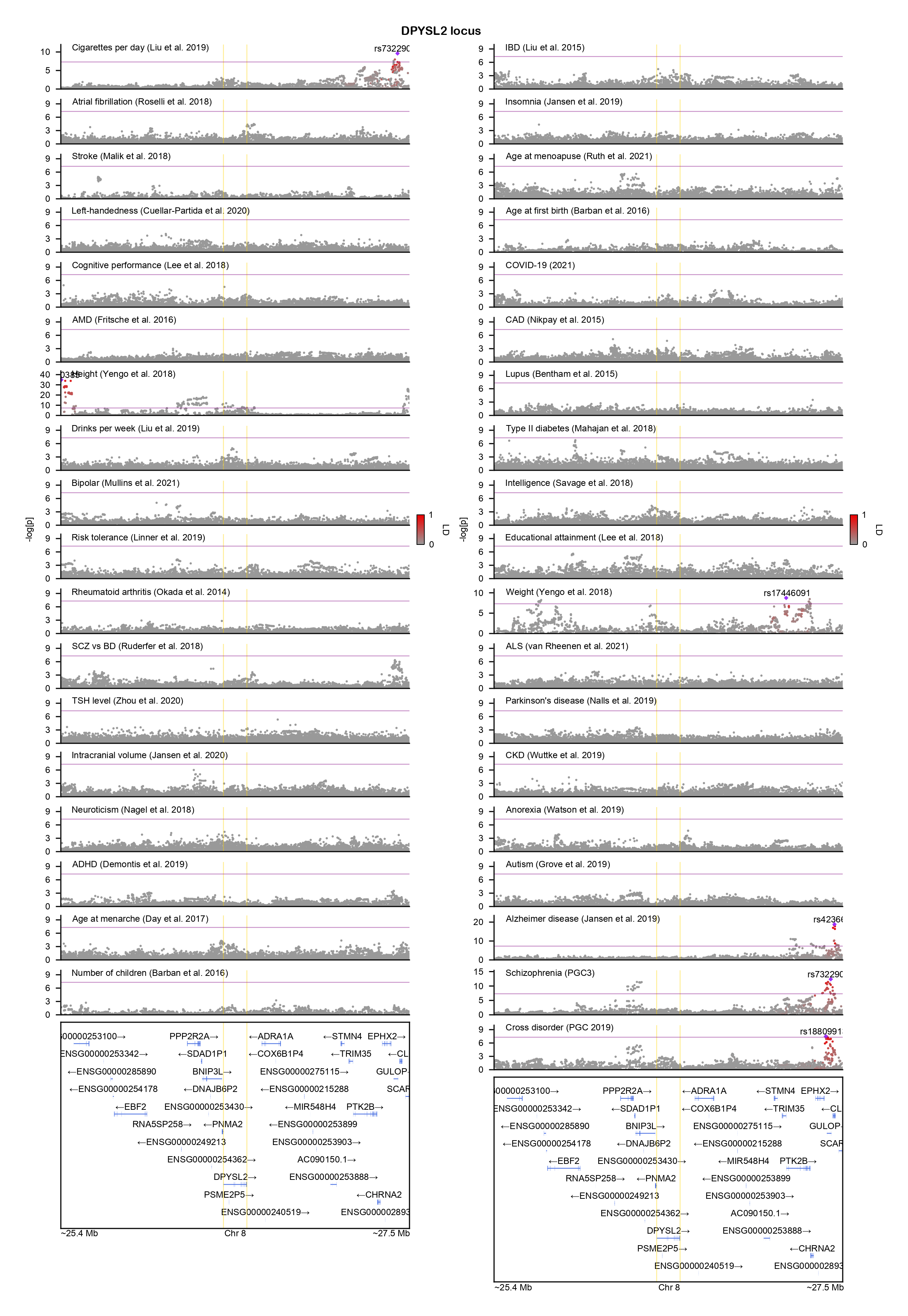

### DSCAM-locuszoom.png

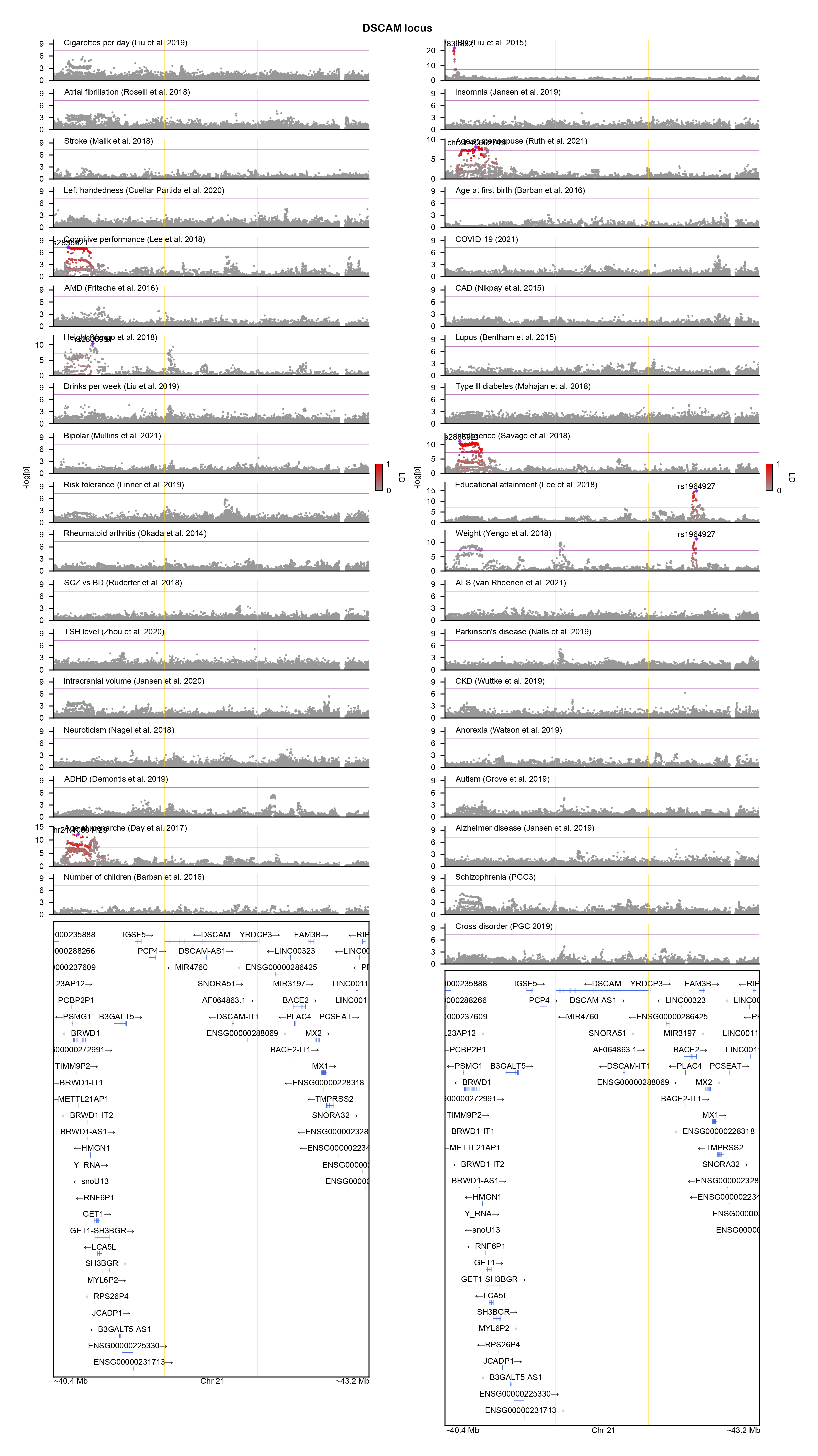

### DYRK1A-locuszoom.png

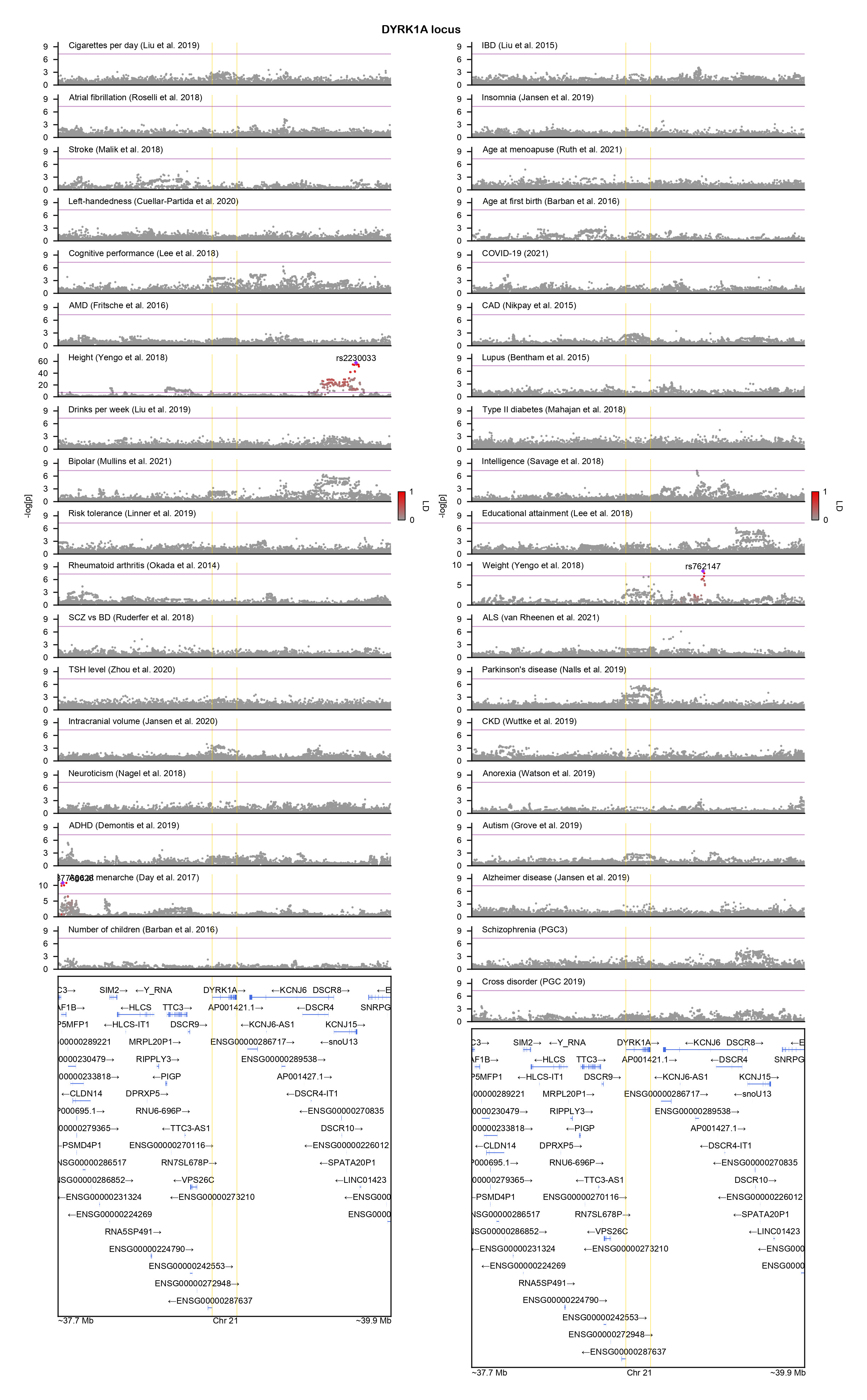

### EIF3G-locuszoom.png

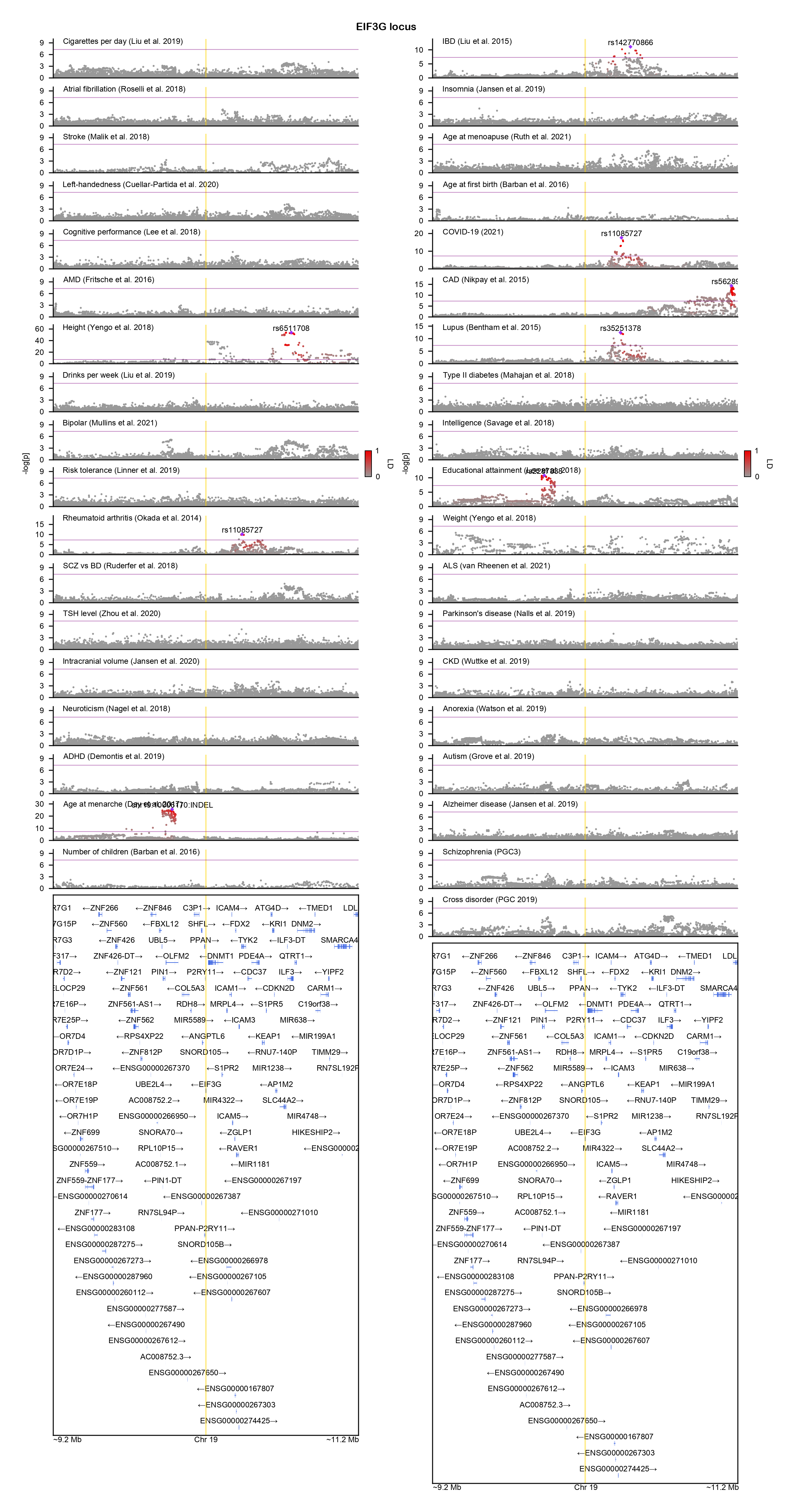

### ELAVL3-locuszoom.png

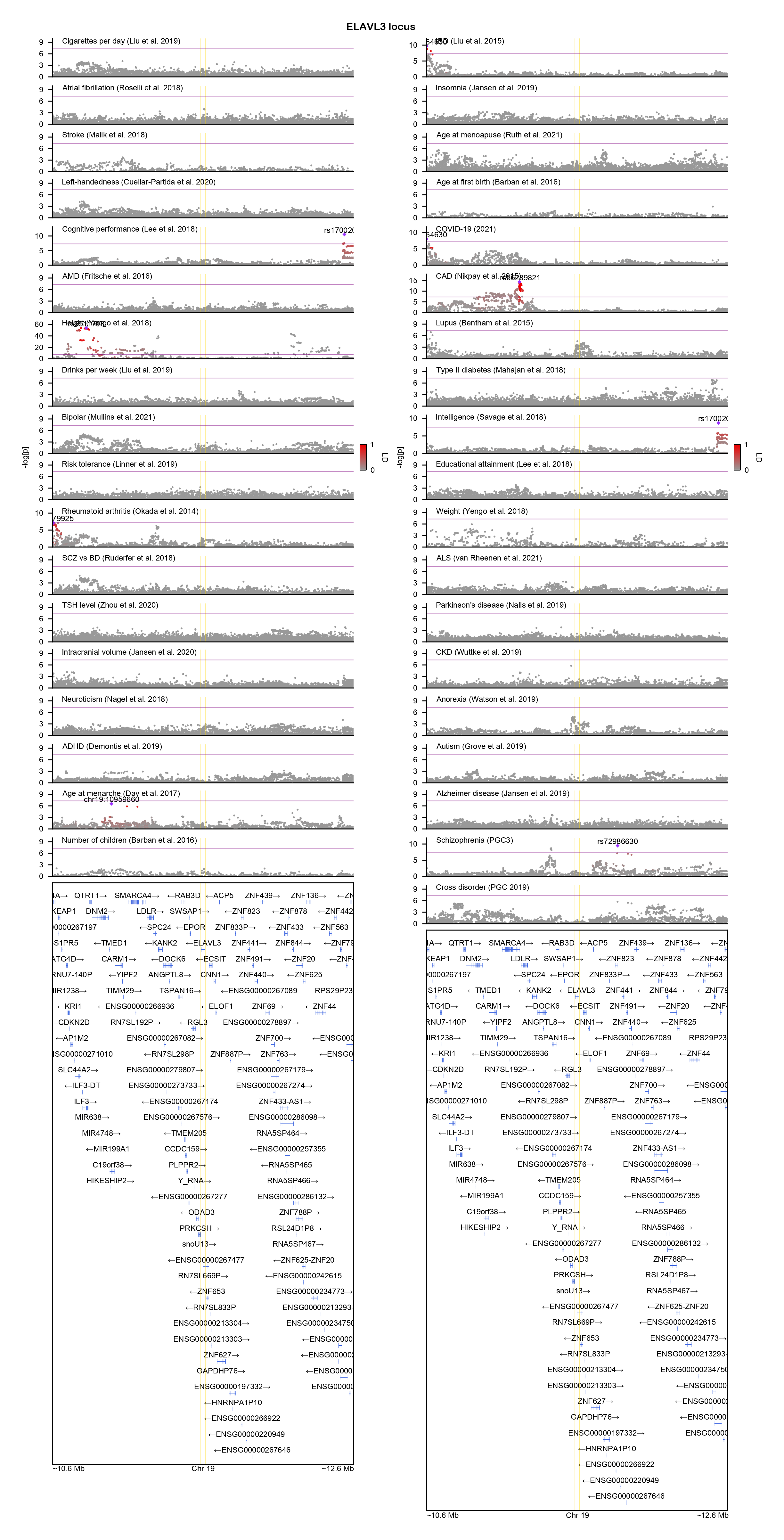

### FBXW7-locuszoom.png

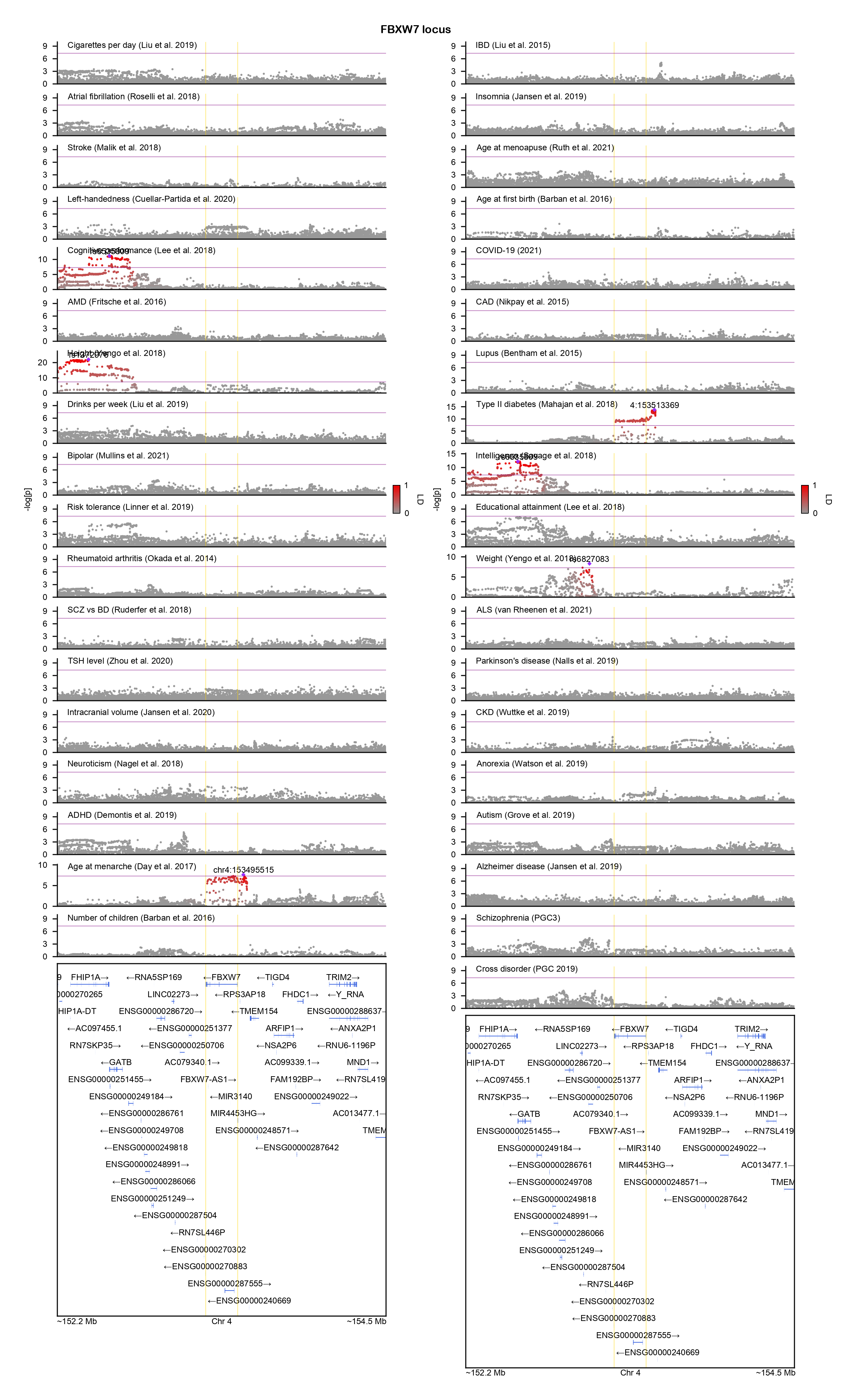

### FES-locuszoom.png

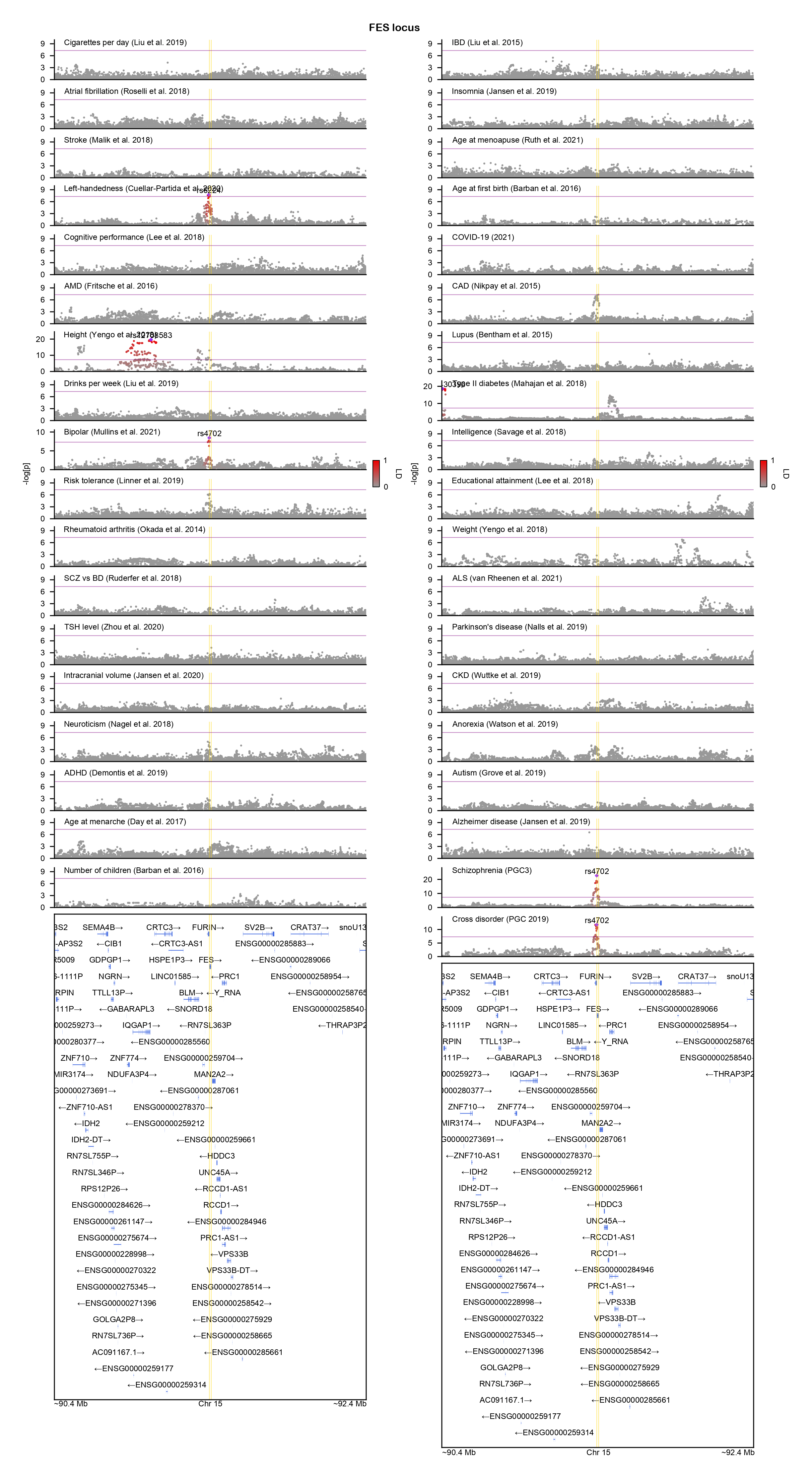

### FGF12-locuszoom.png

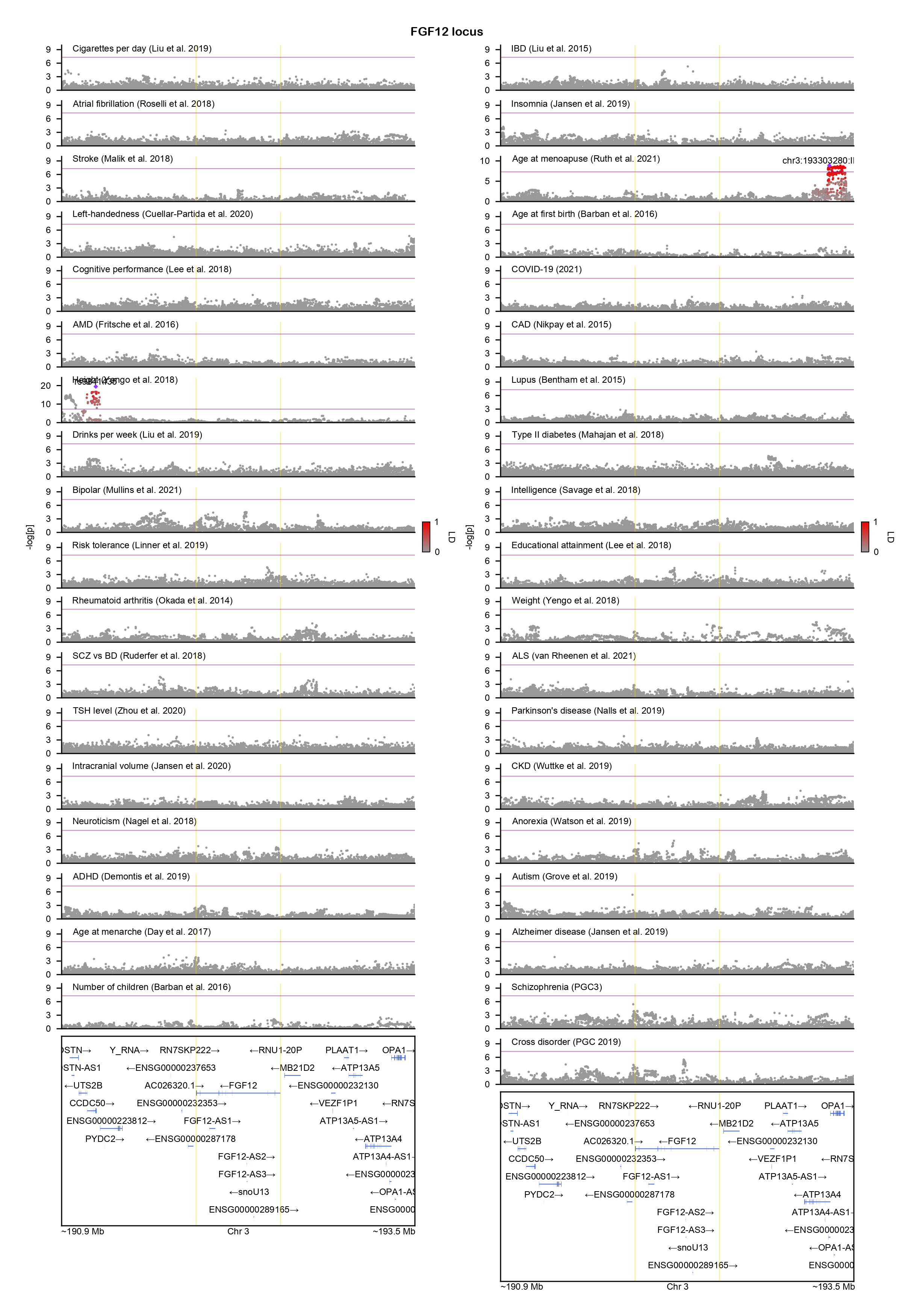

### GATAD2A-locuszoom.png

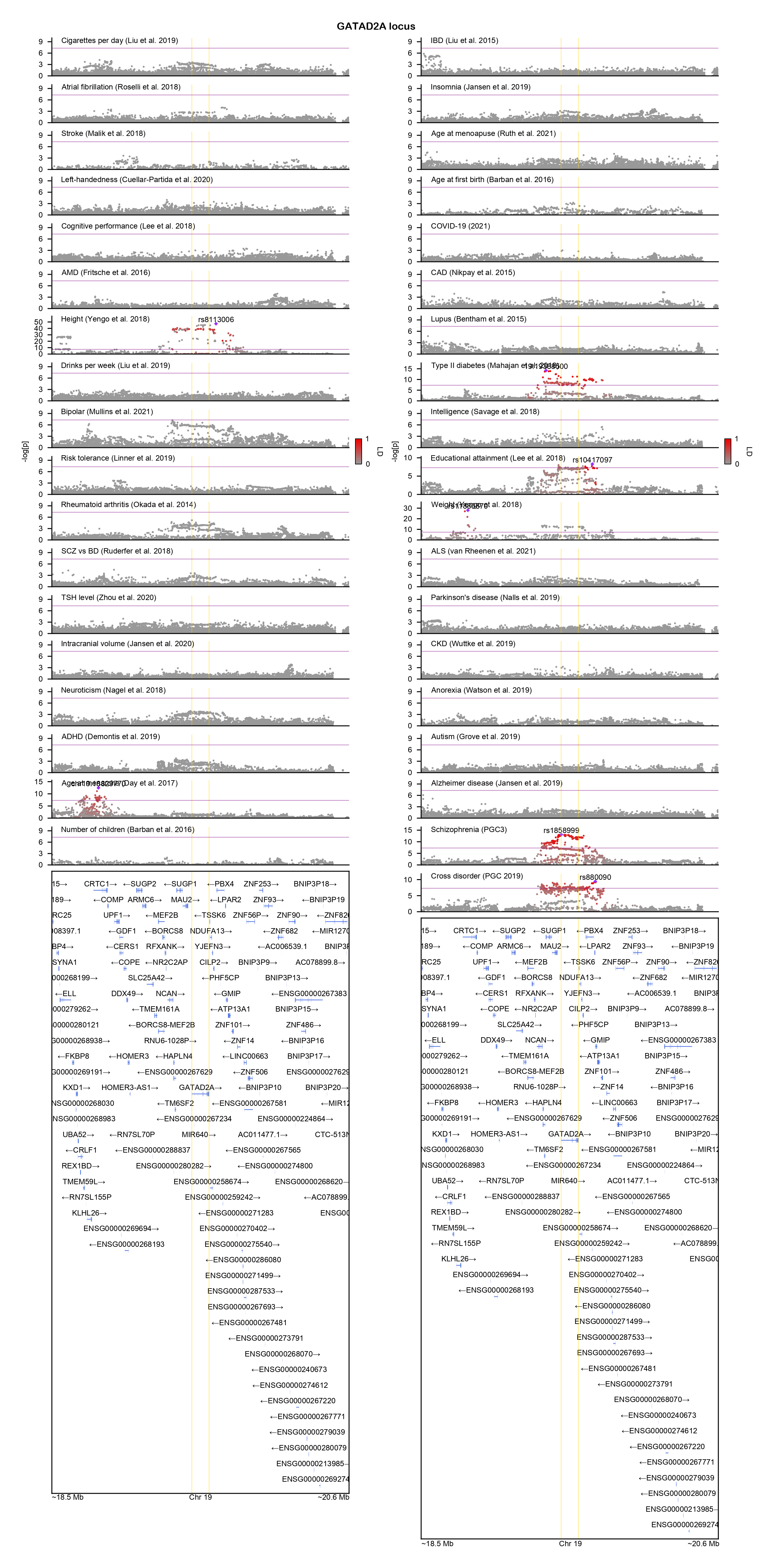

### GBE1-locuszoom.png

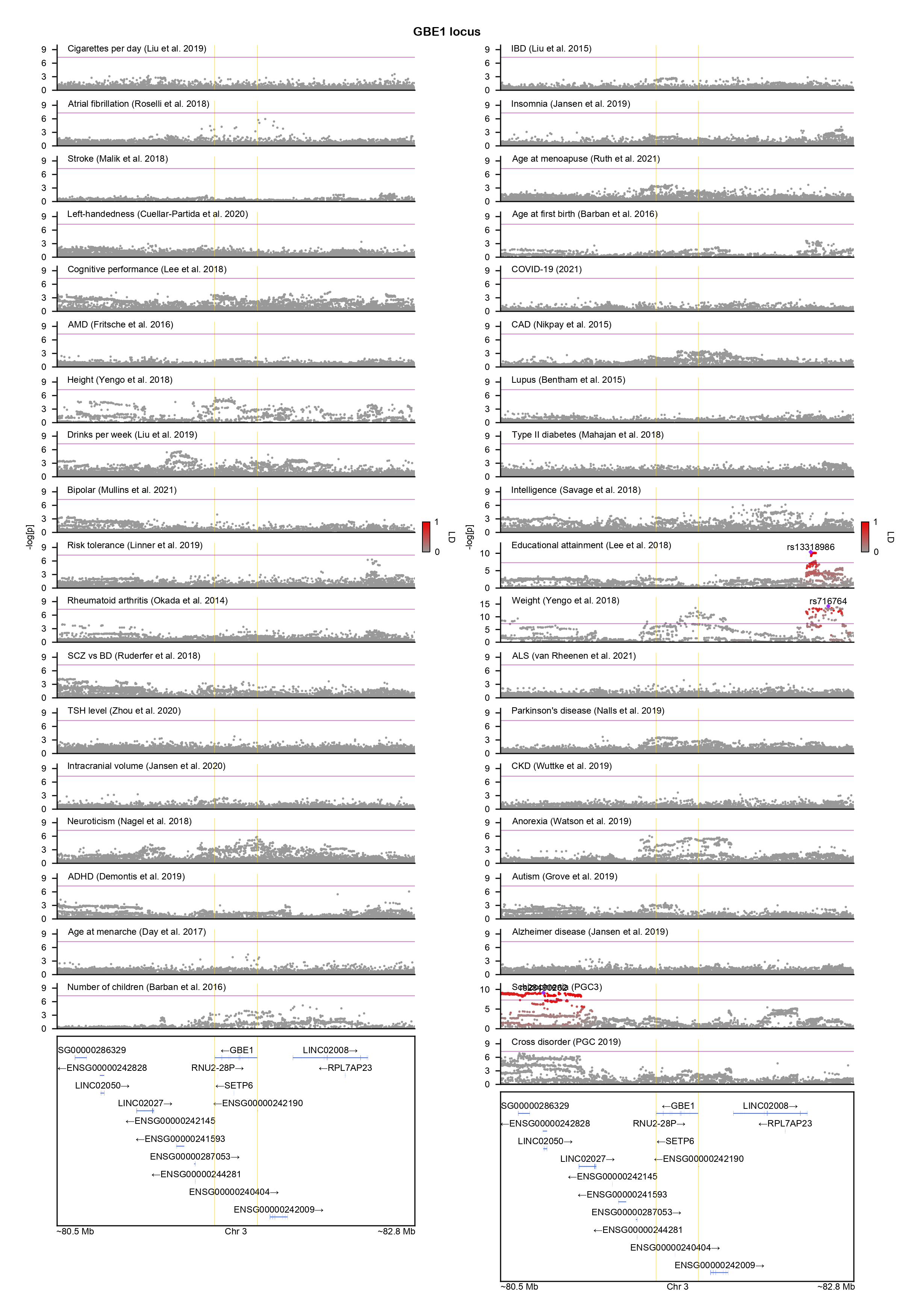

### GNAI1-locuszoom.png

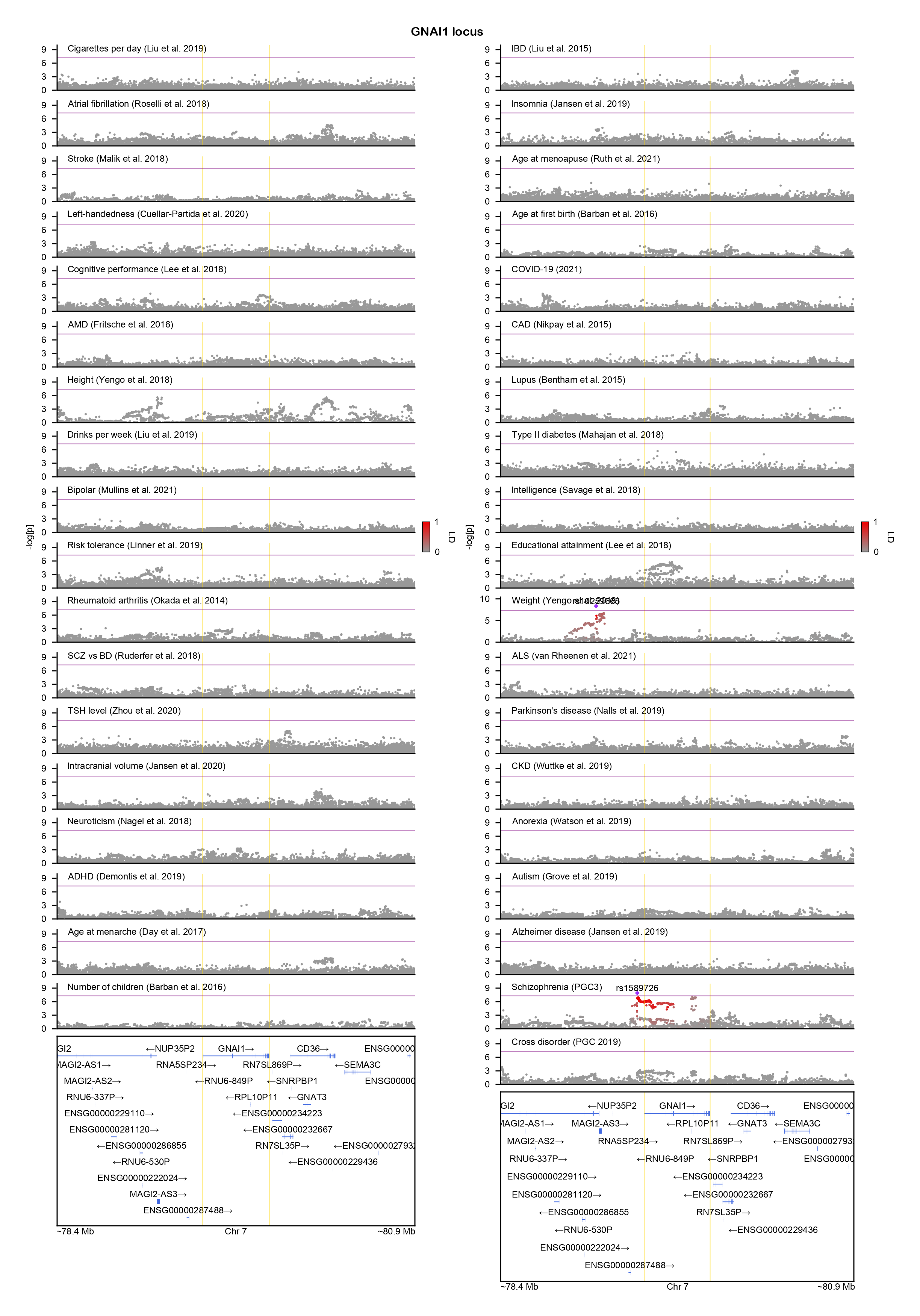

### GNAO1-locuszoom.png

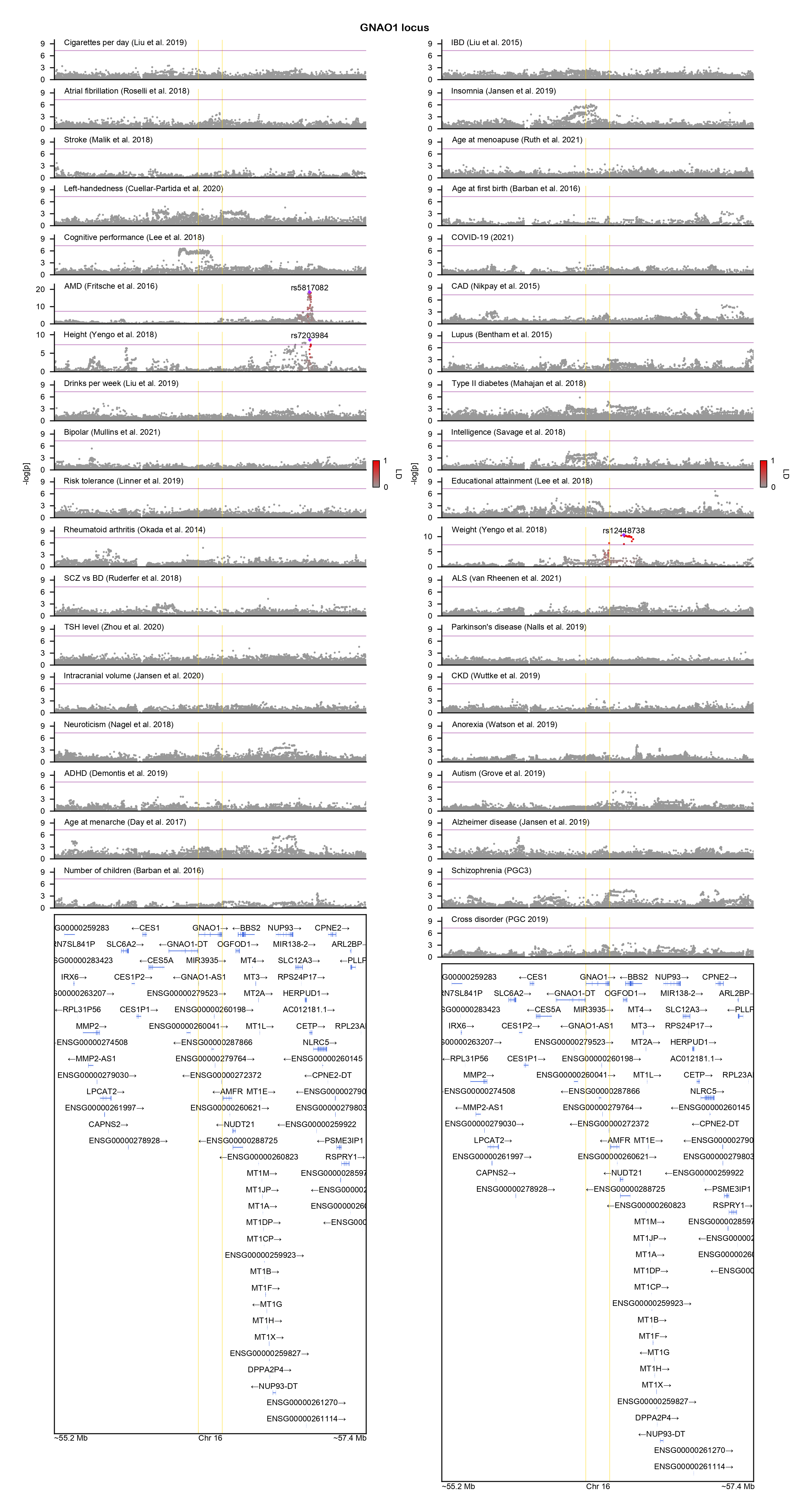

### GNE-locuszoom.png

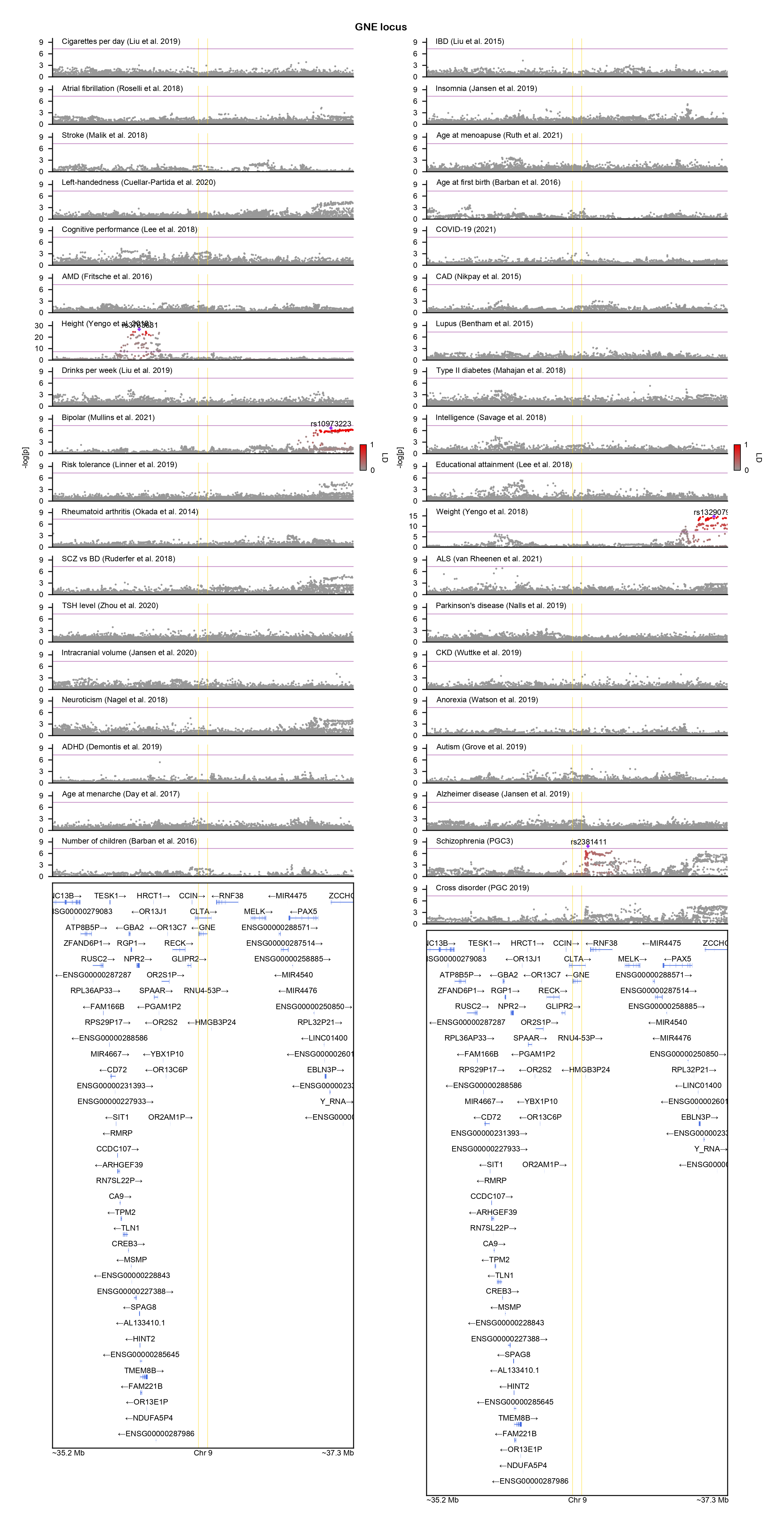

### GPM6A-locuszoom.png

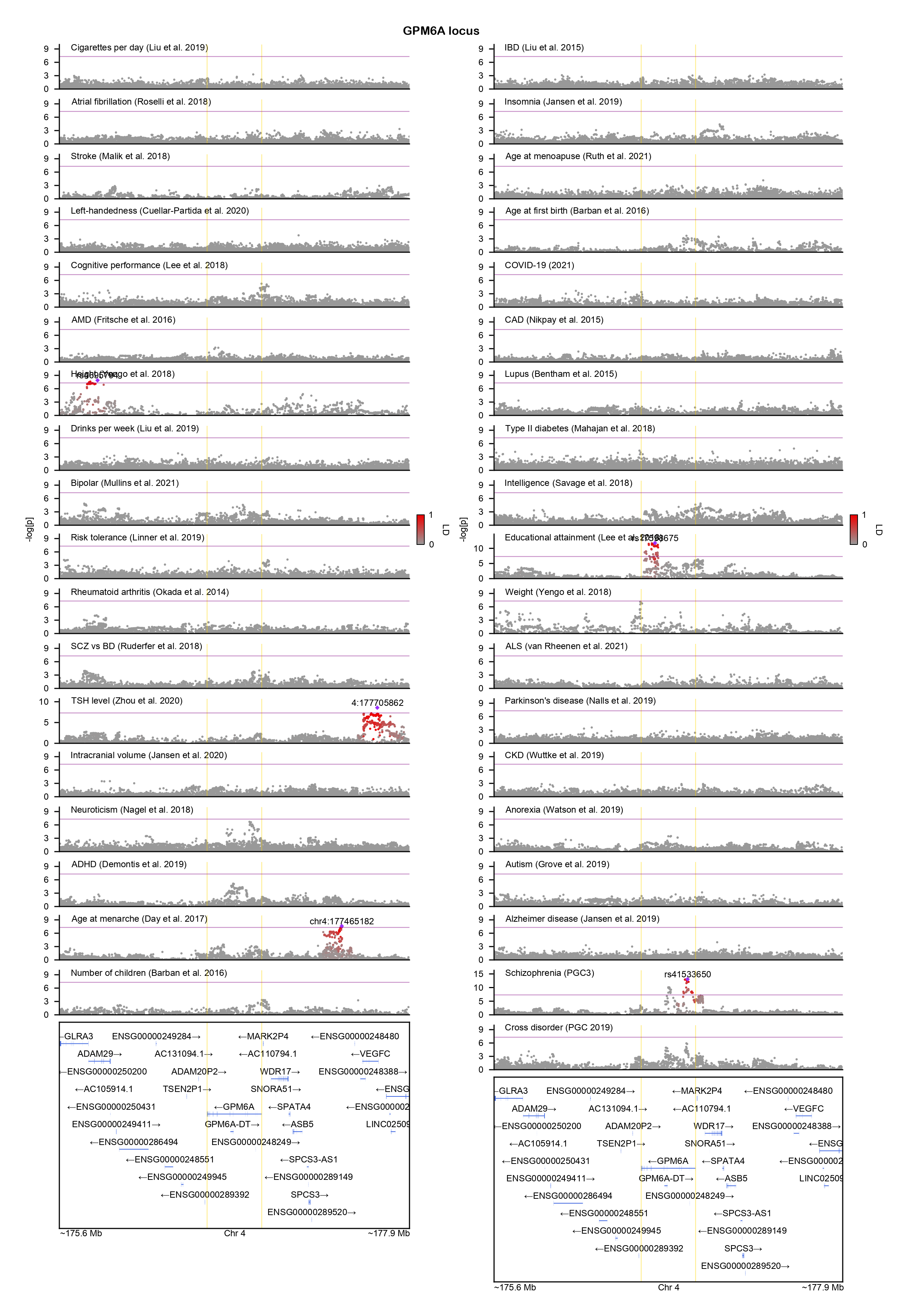

### GRIA3-locuszoom.png

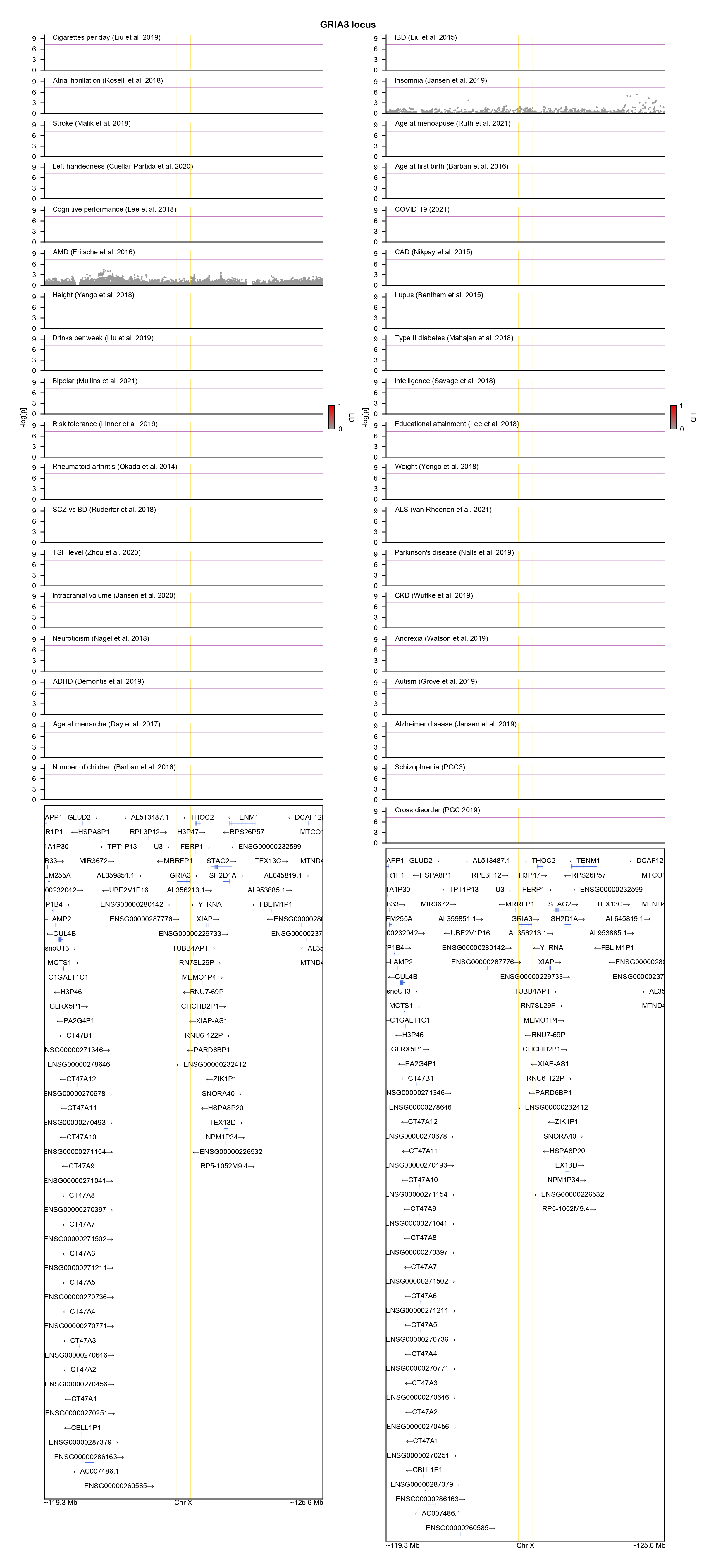

### GRIN2A-locuszoom.png

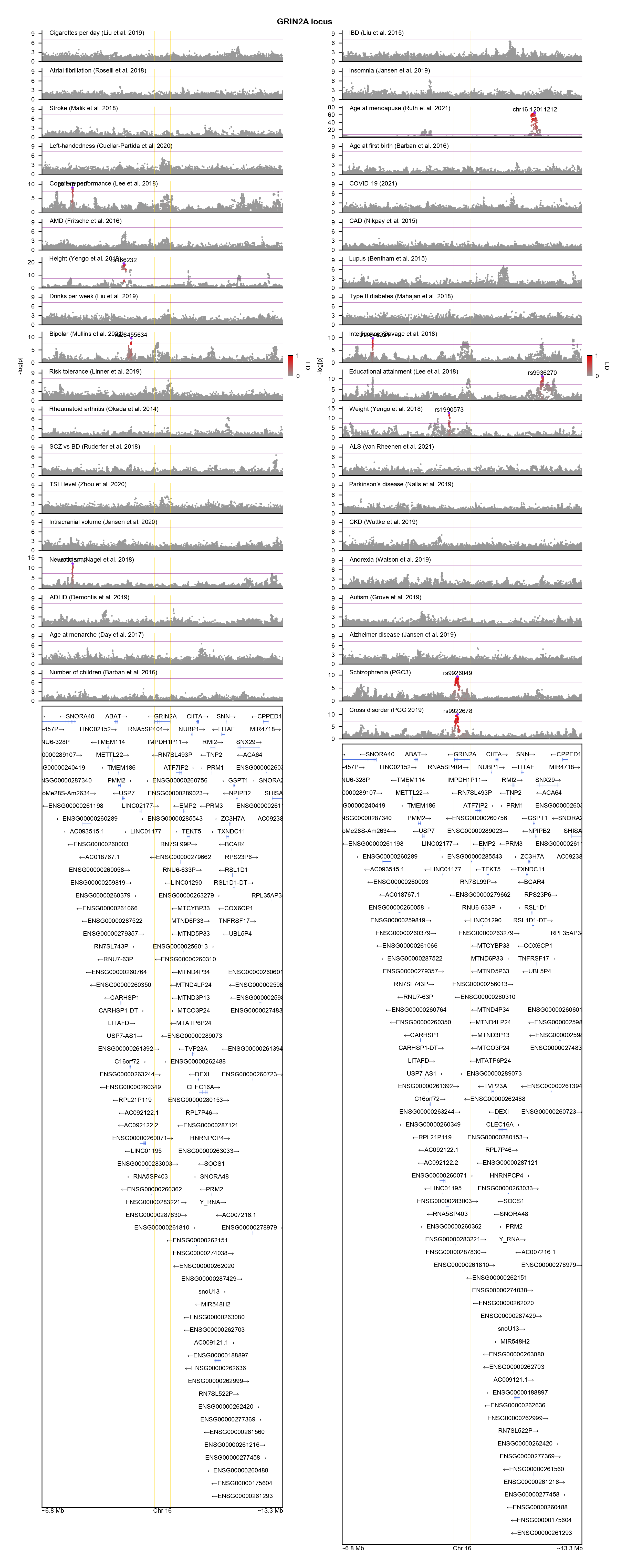

### GRIN2B-locuszoom.png

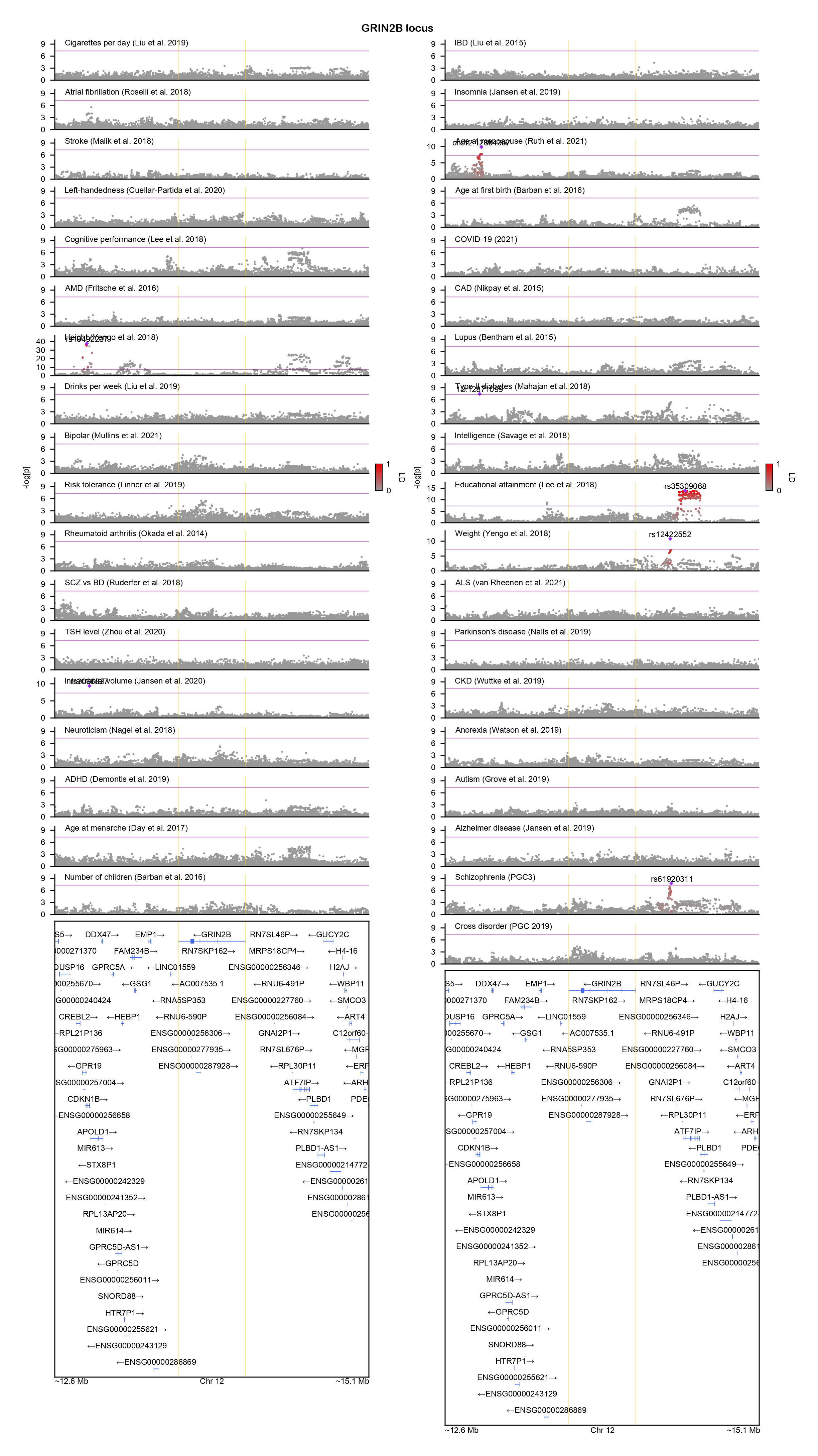

### HCN1-locuszoom.png

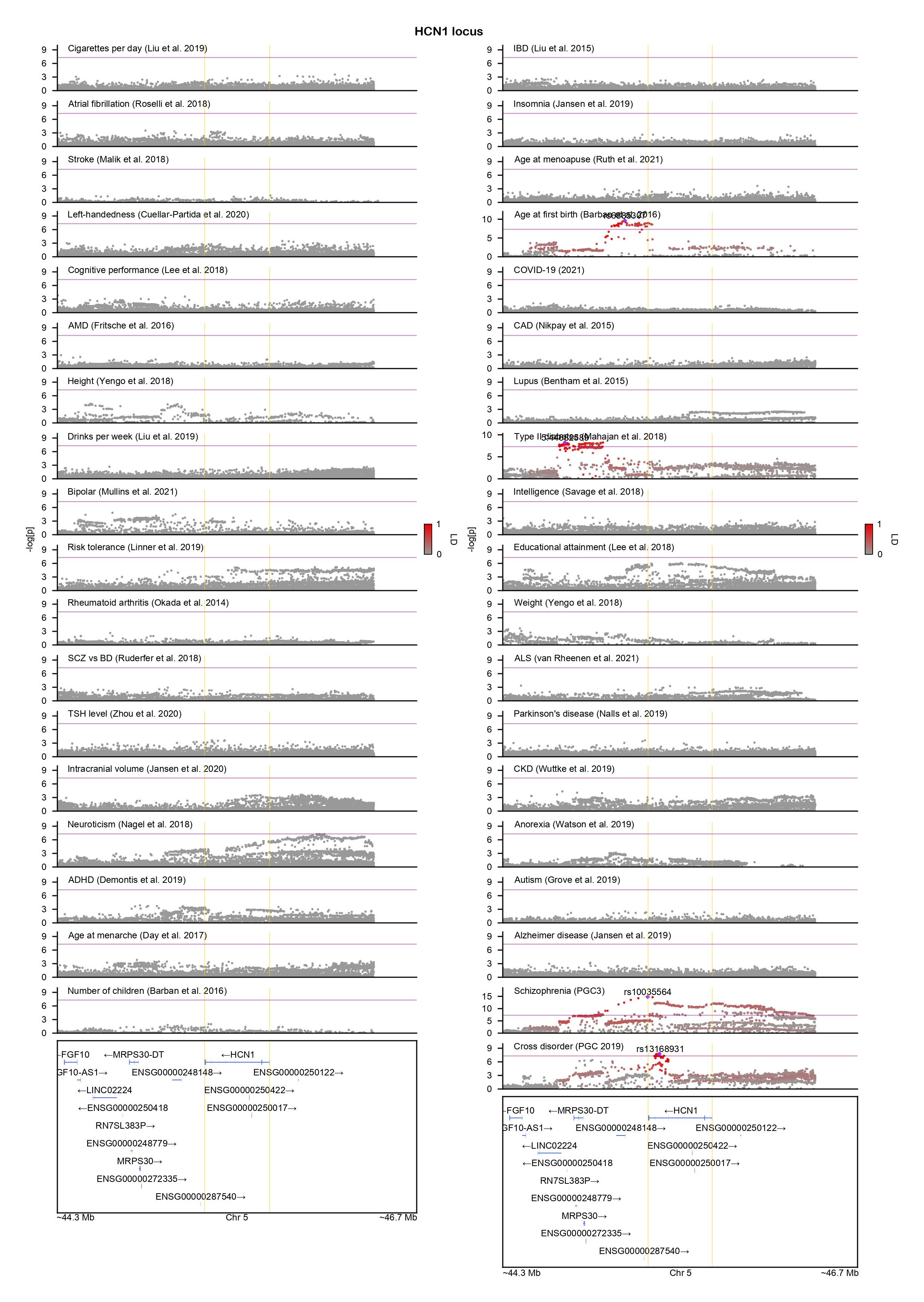

### HK1-locuszoom.png

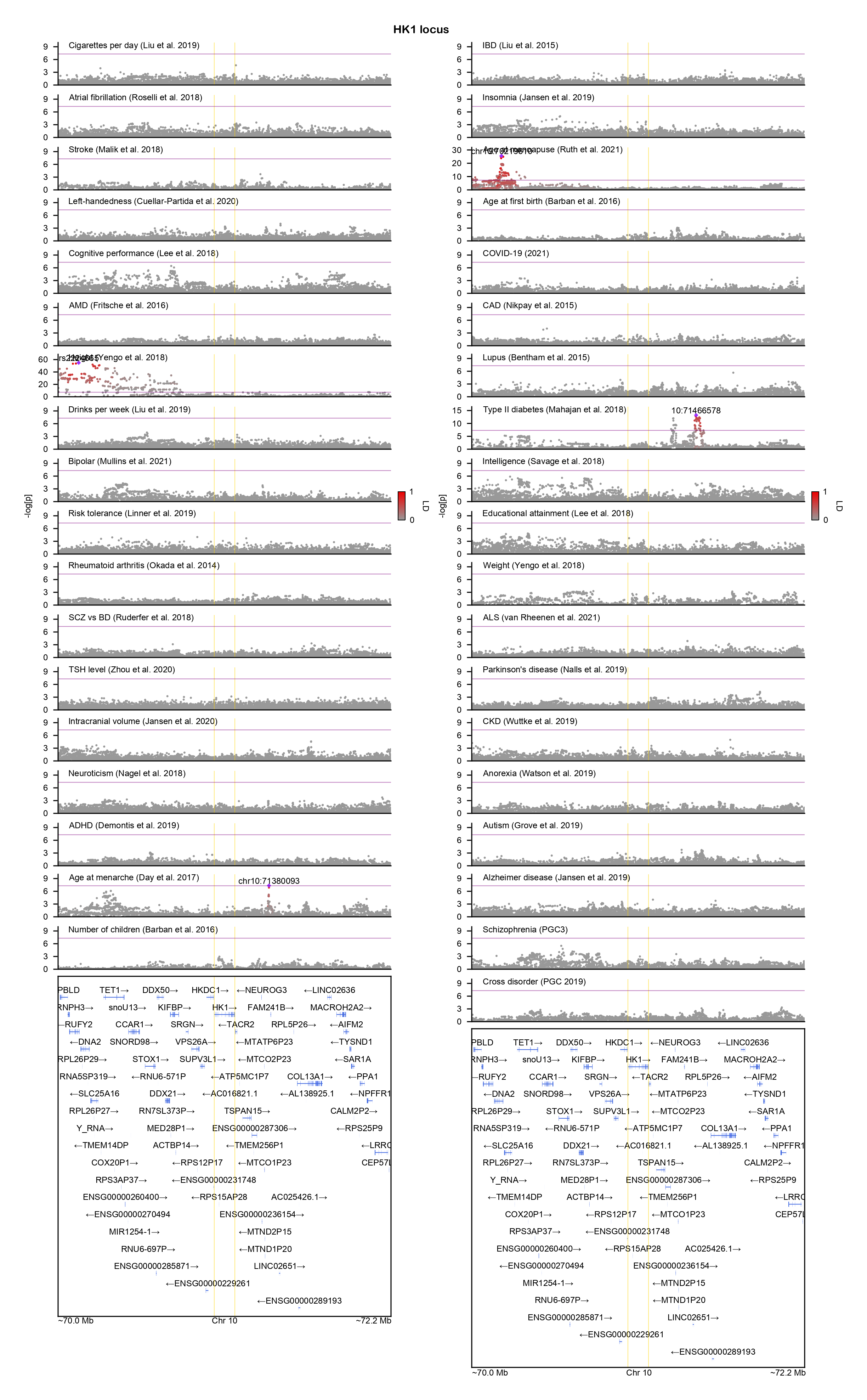

### HNRNPD-locuszoom.png

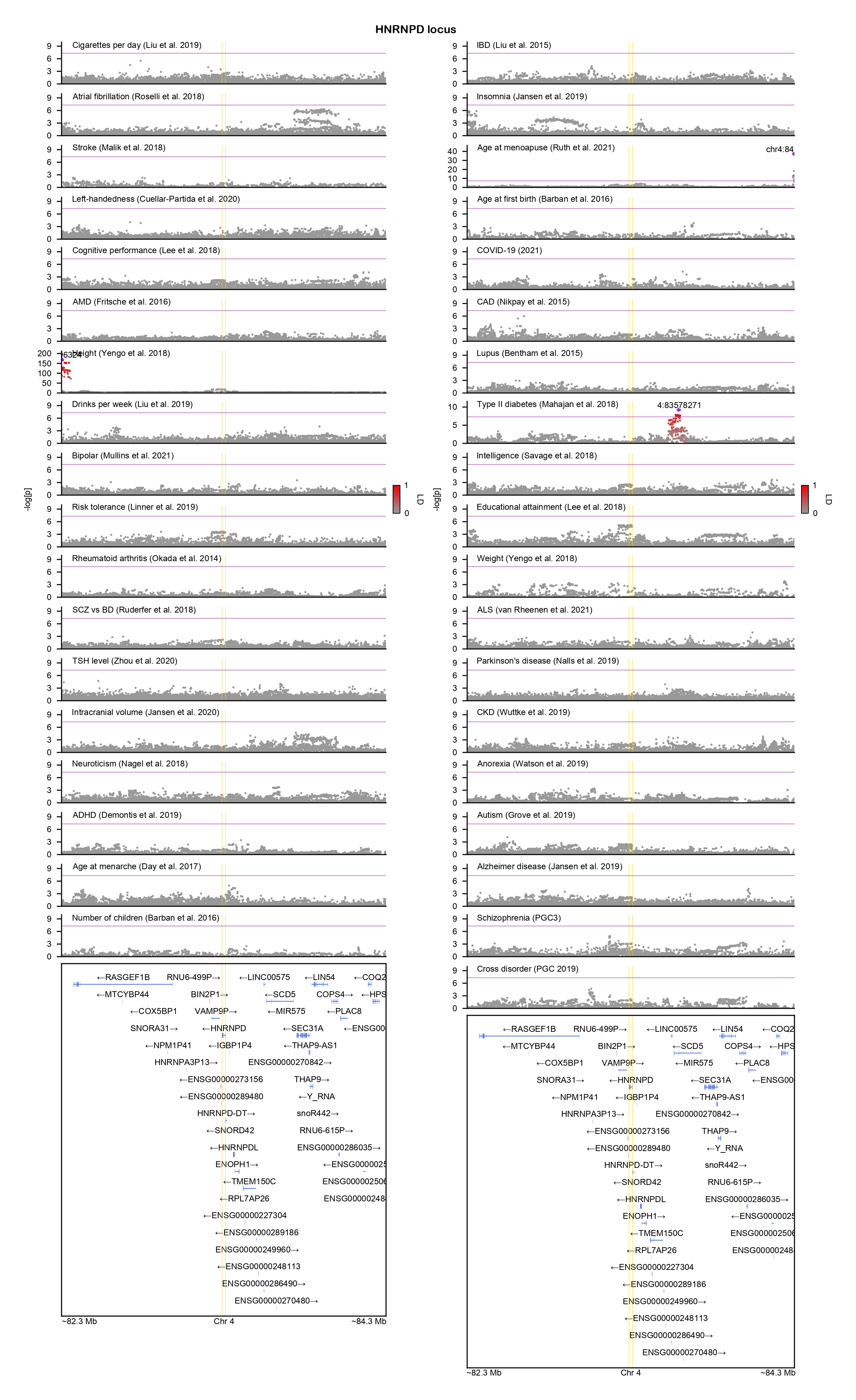

### HNRNPU-locuszoom.png

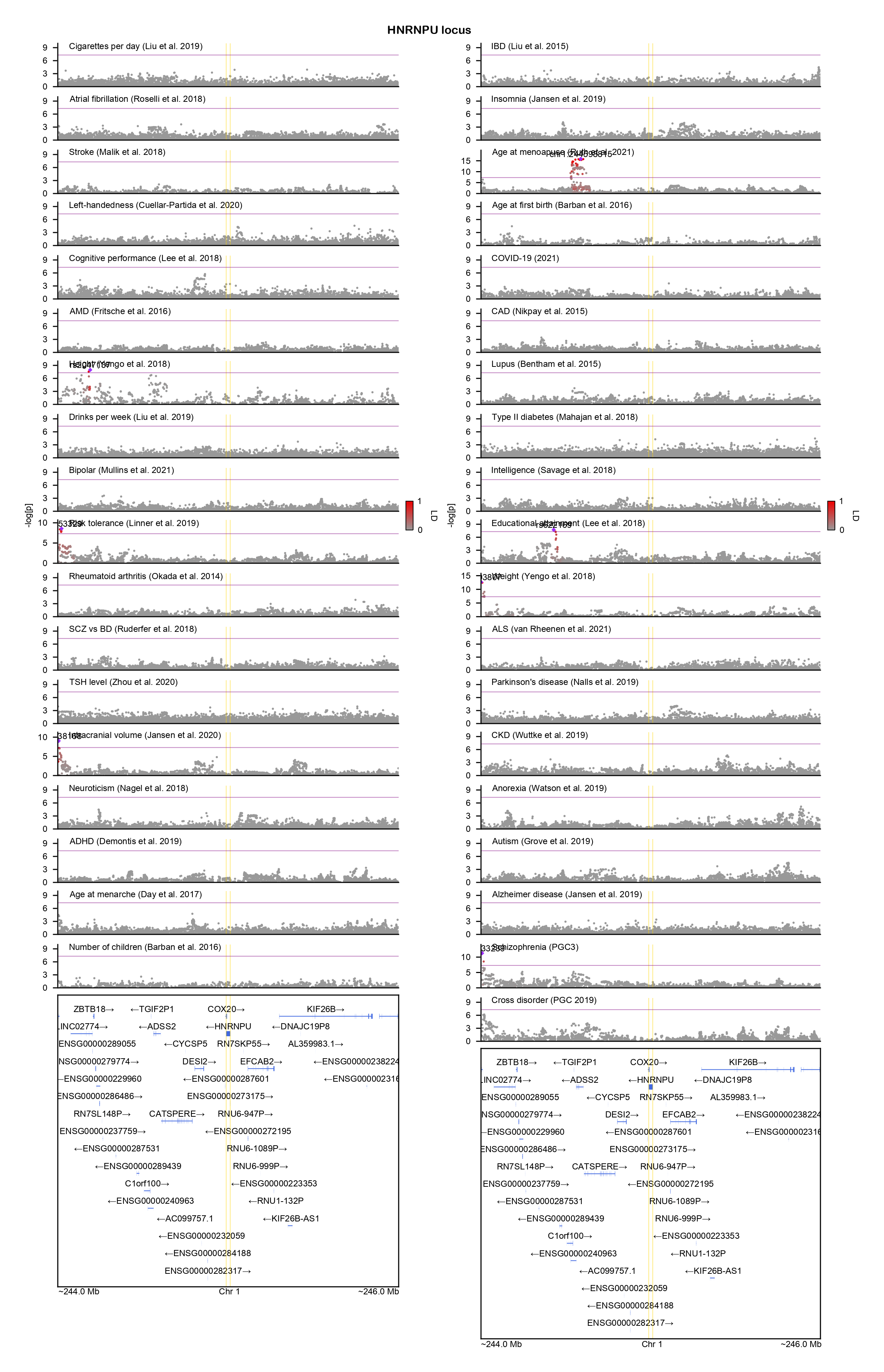

### IMMP2L-locuszoom.png

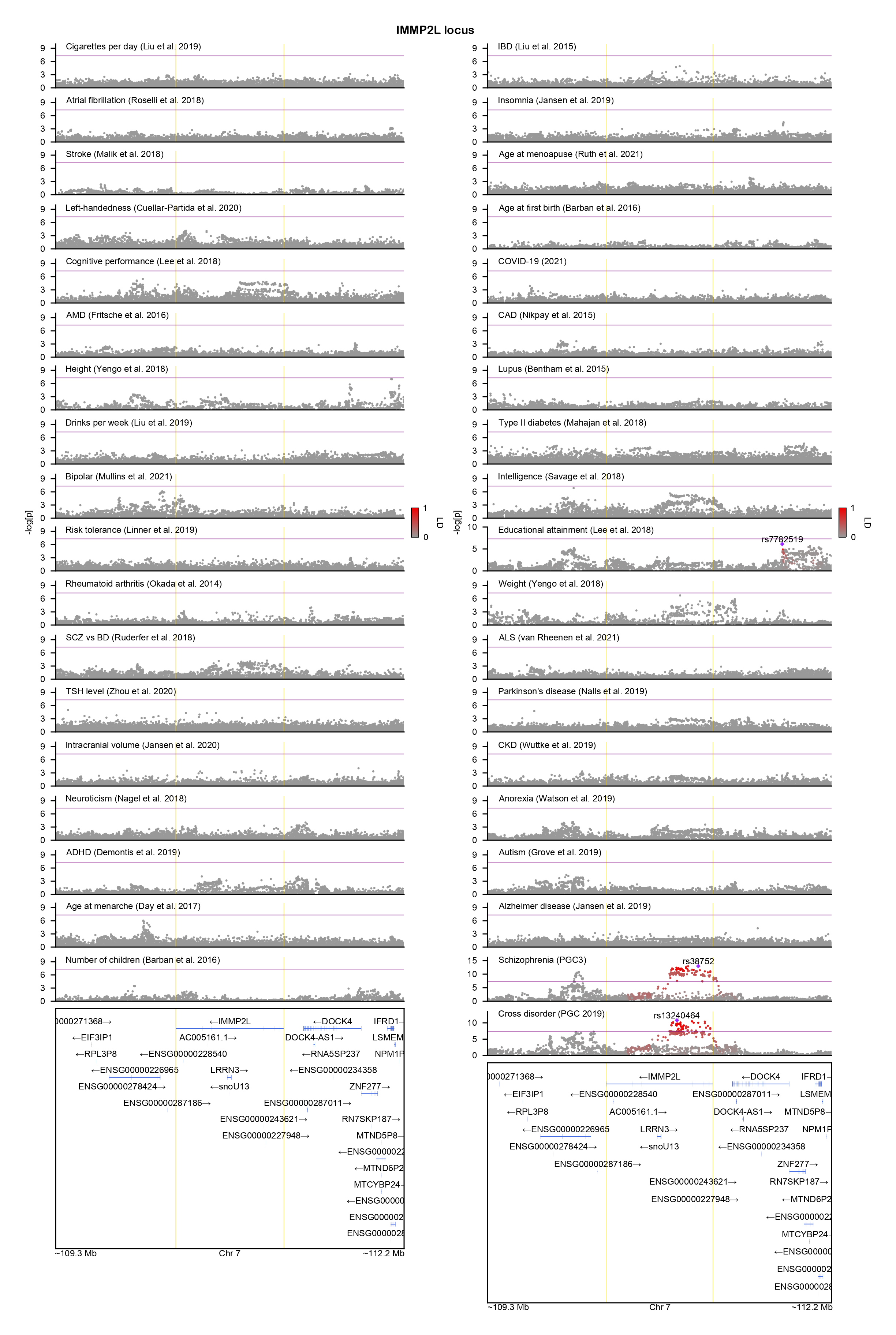

### IRF3-locuszoom.png

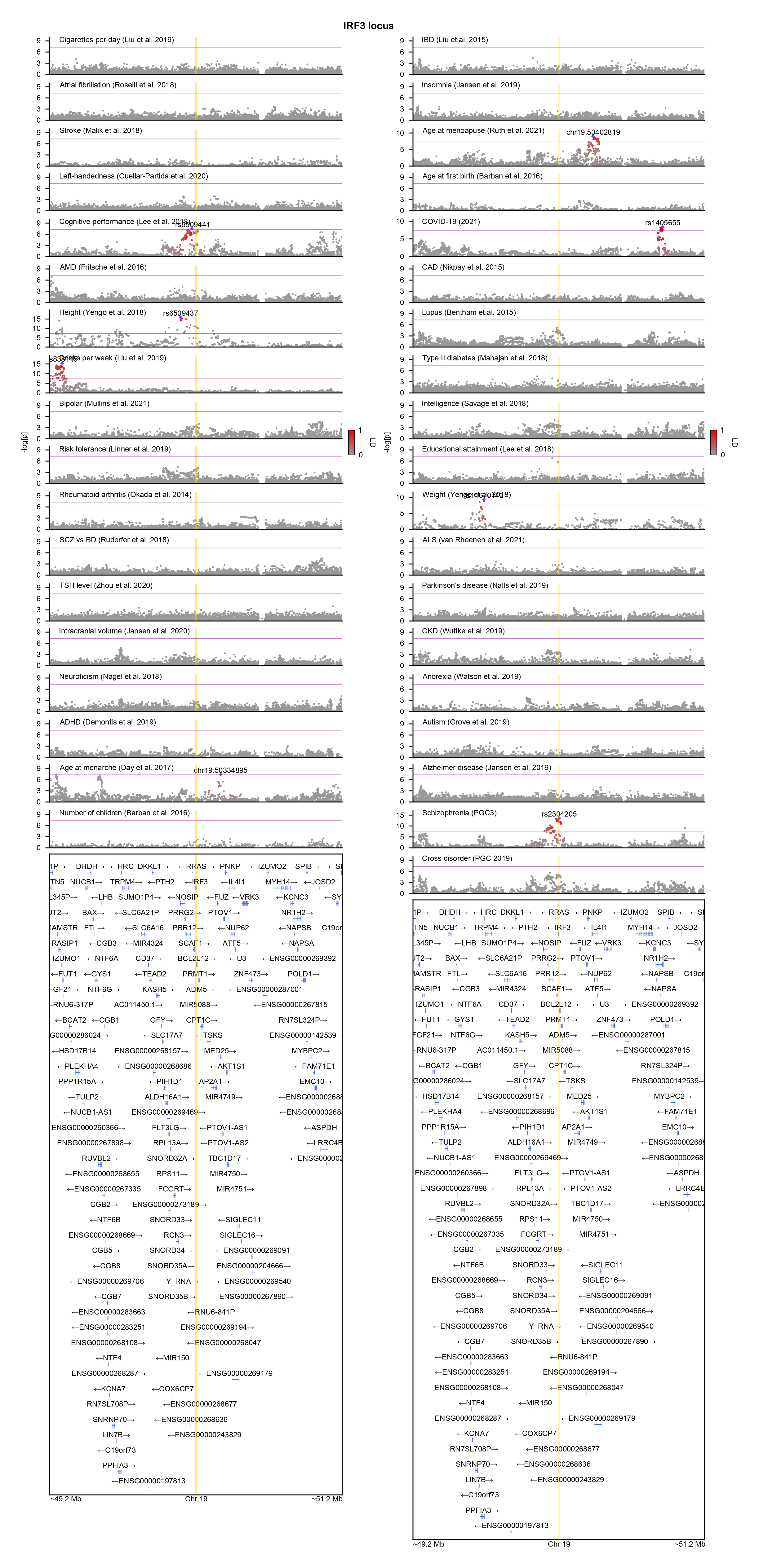

### KCNQ3-locuszoom.png

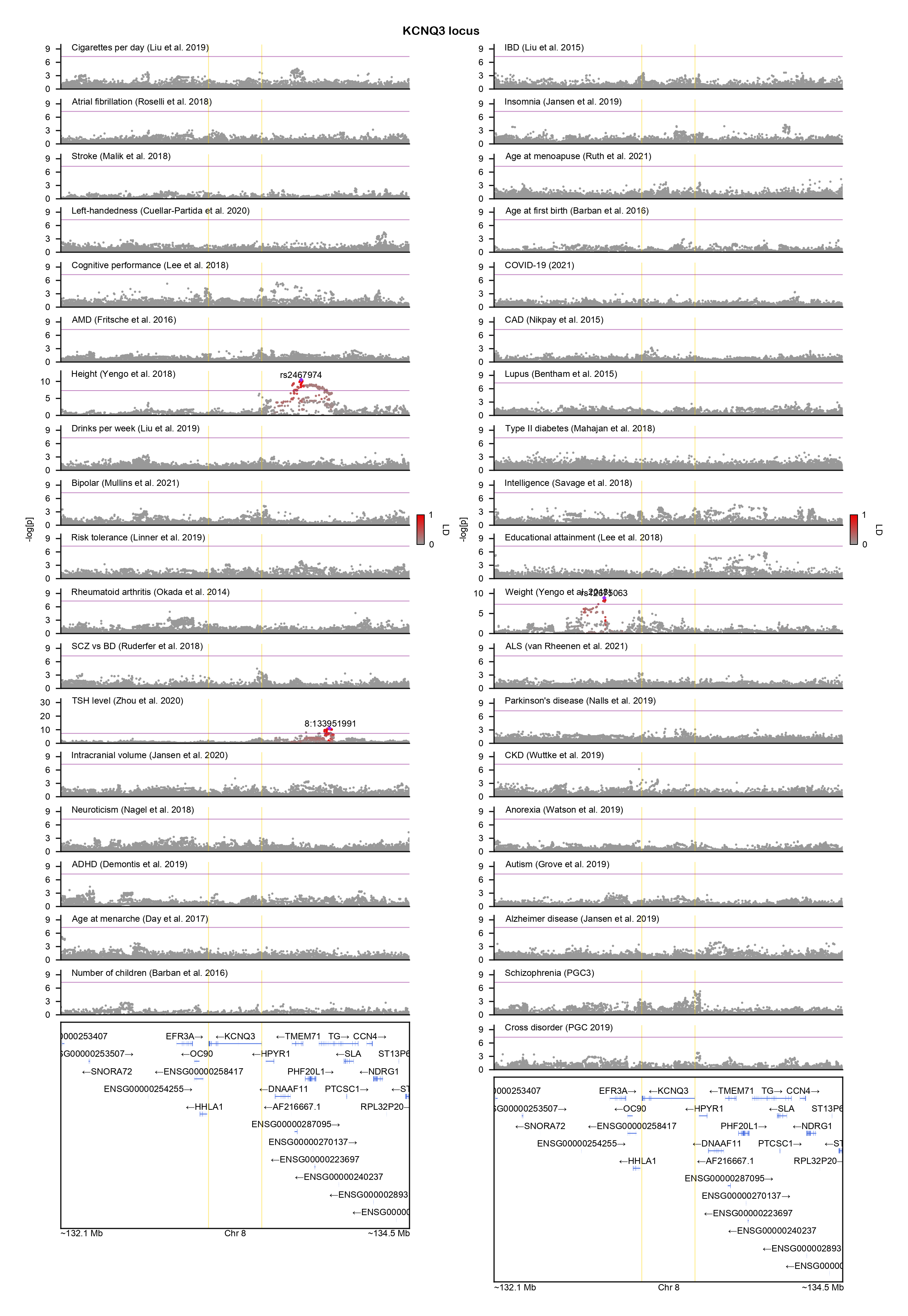

### KCNQ5-locuszoom.png

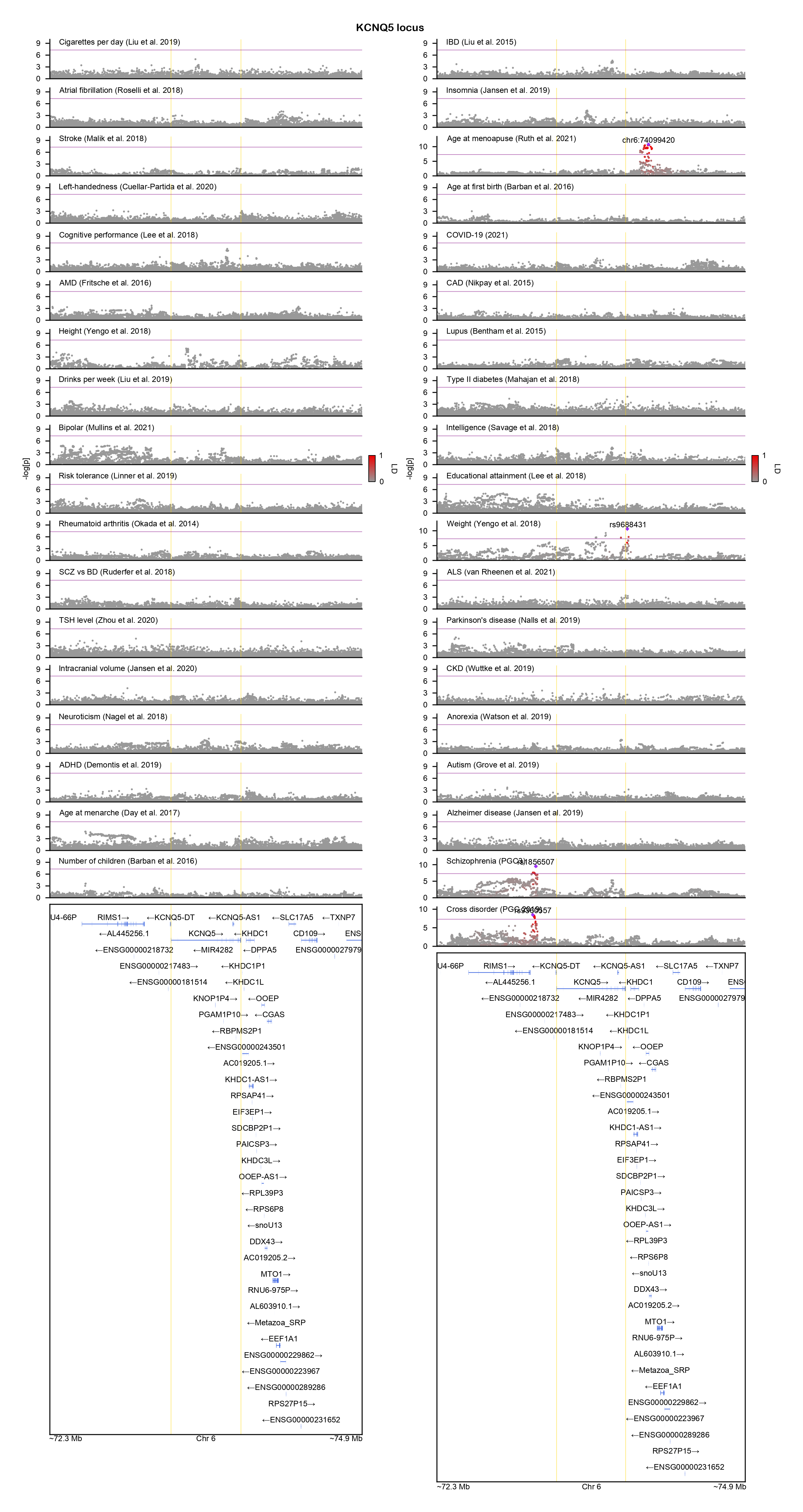

### KCNV1-locuszoom.png

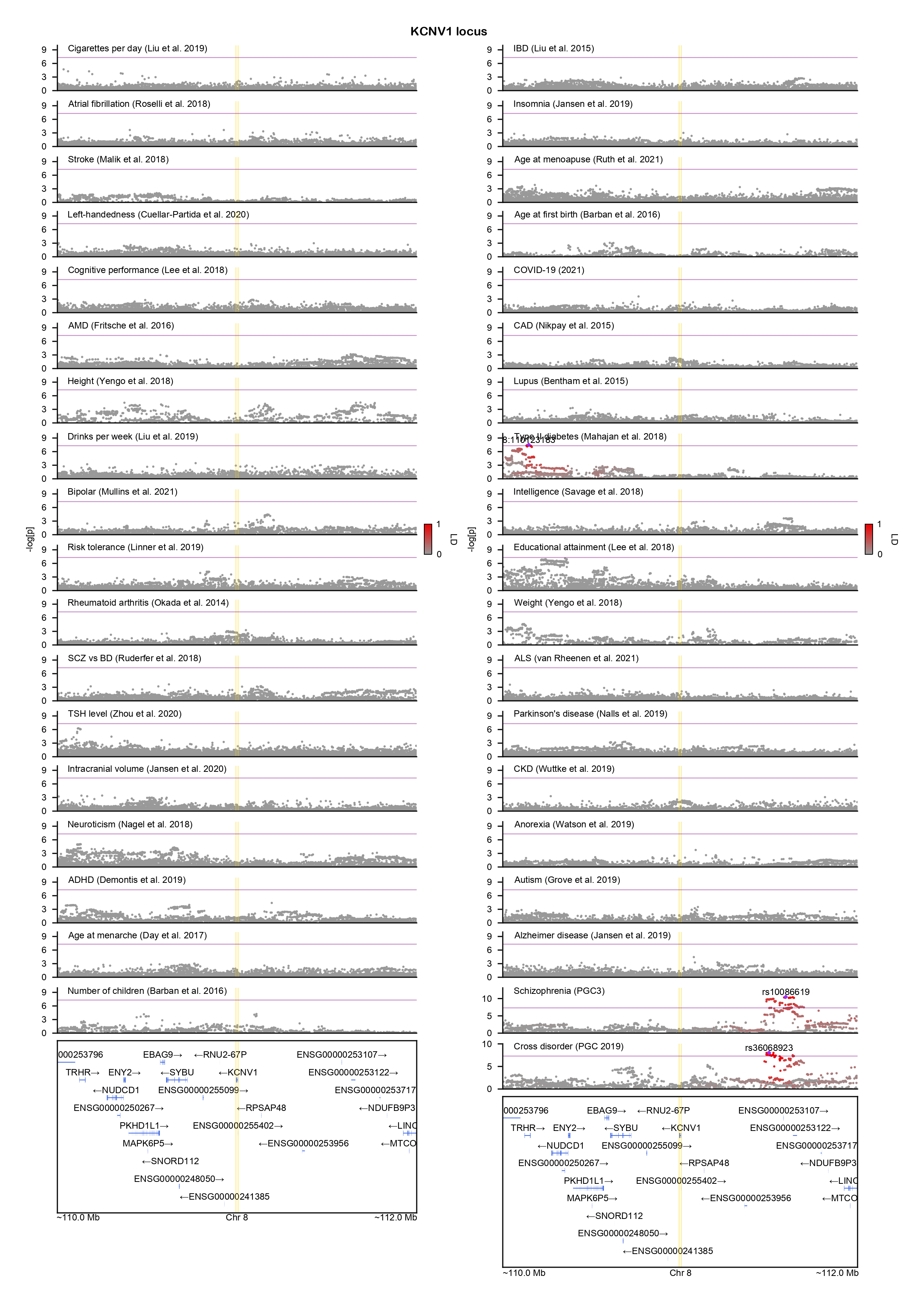

### KDM6A-locuszoom.png

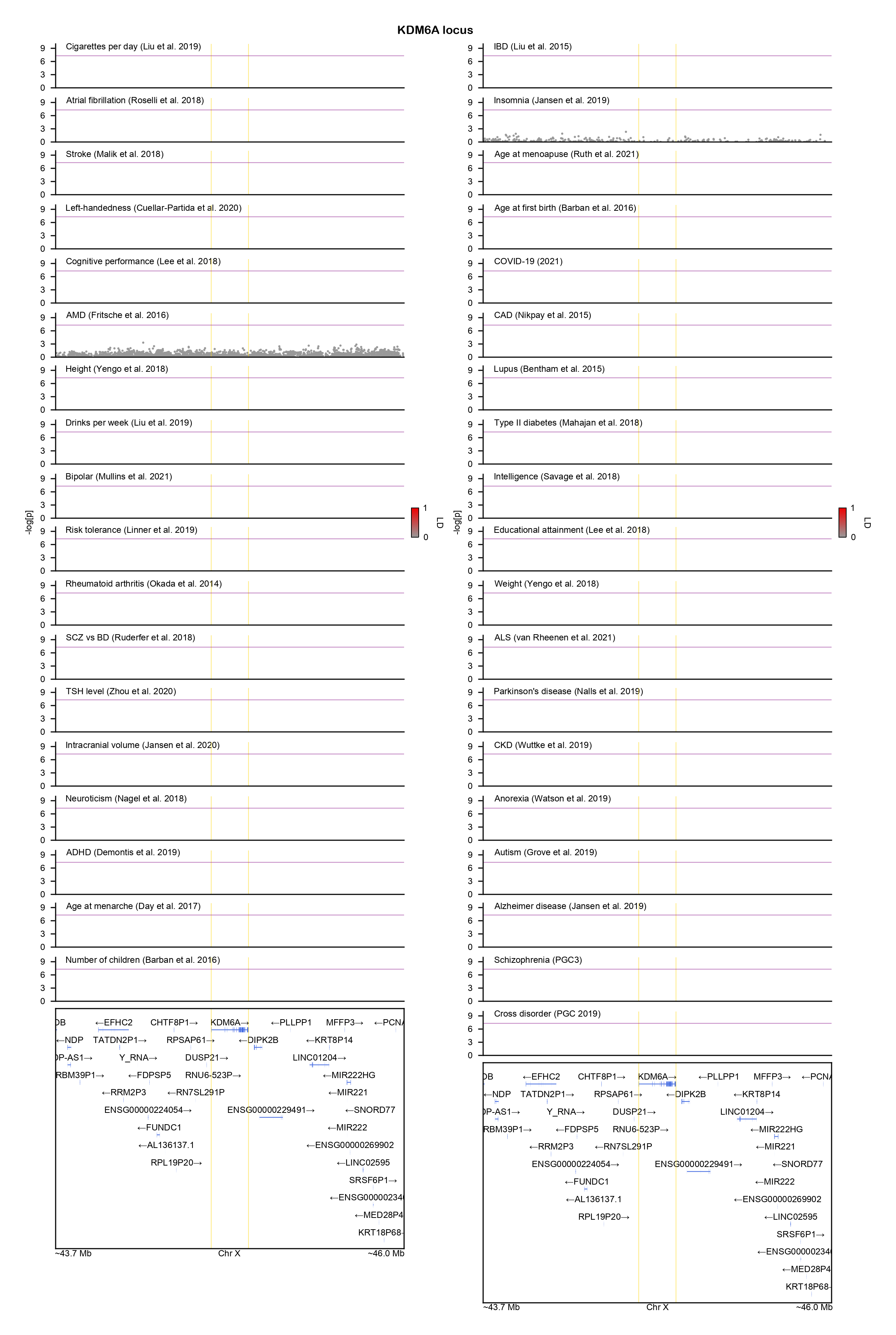

### KDM6B-locuszoom.png

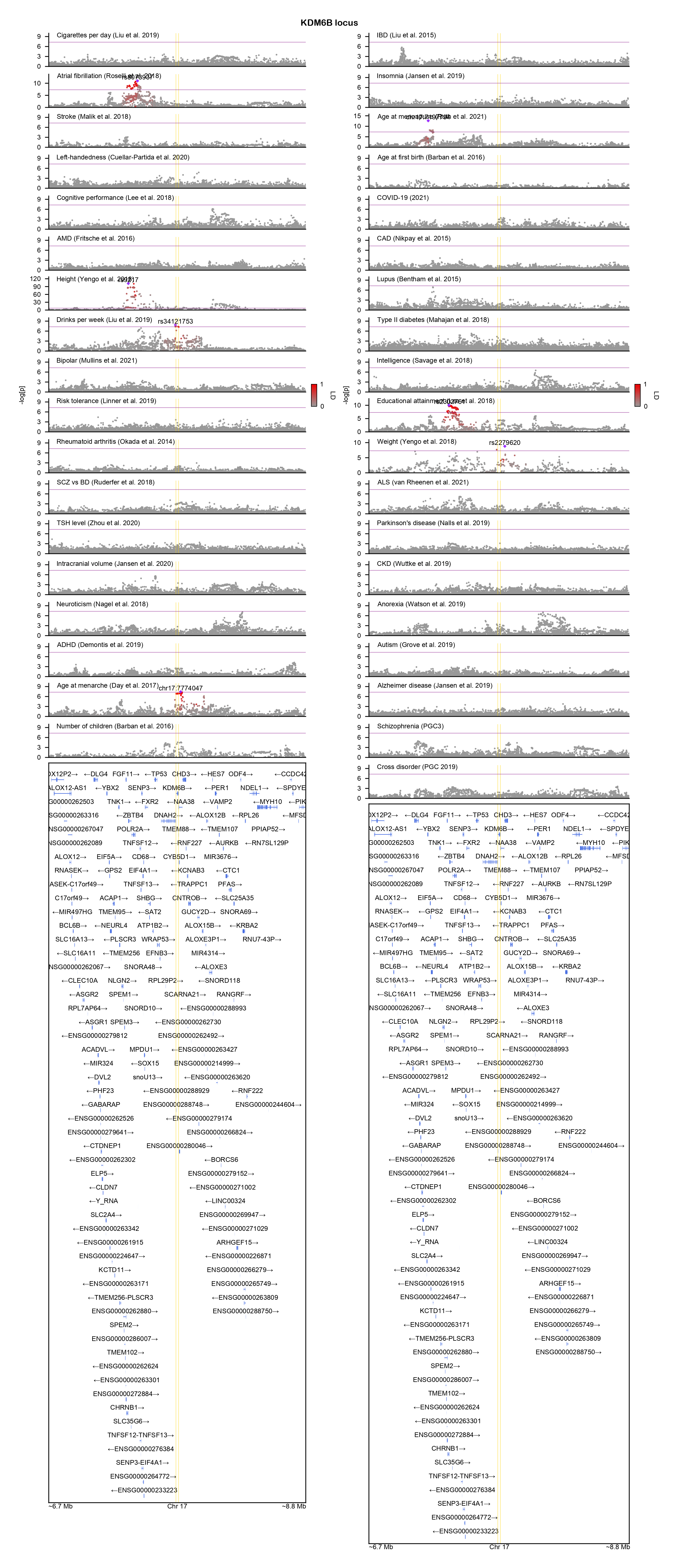

### KIF1A-locuszoom.png

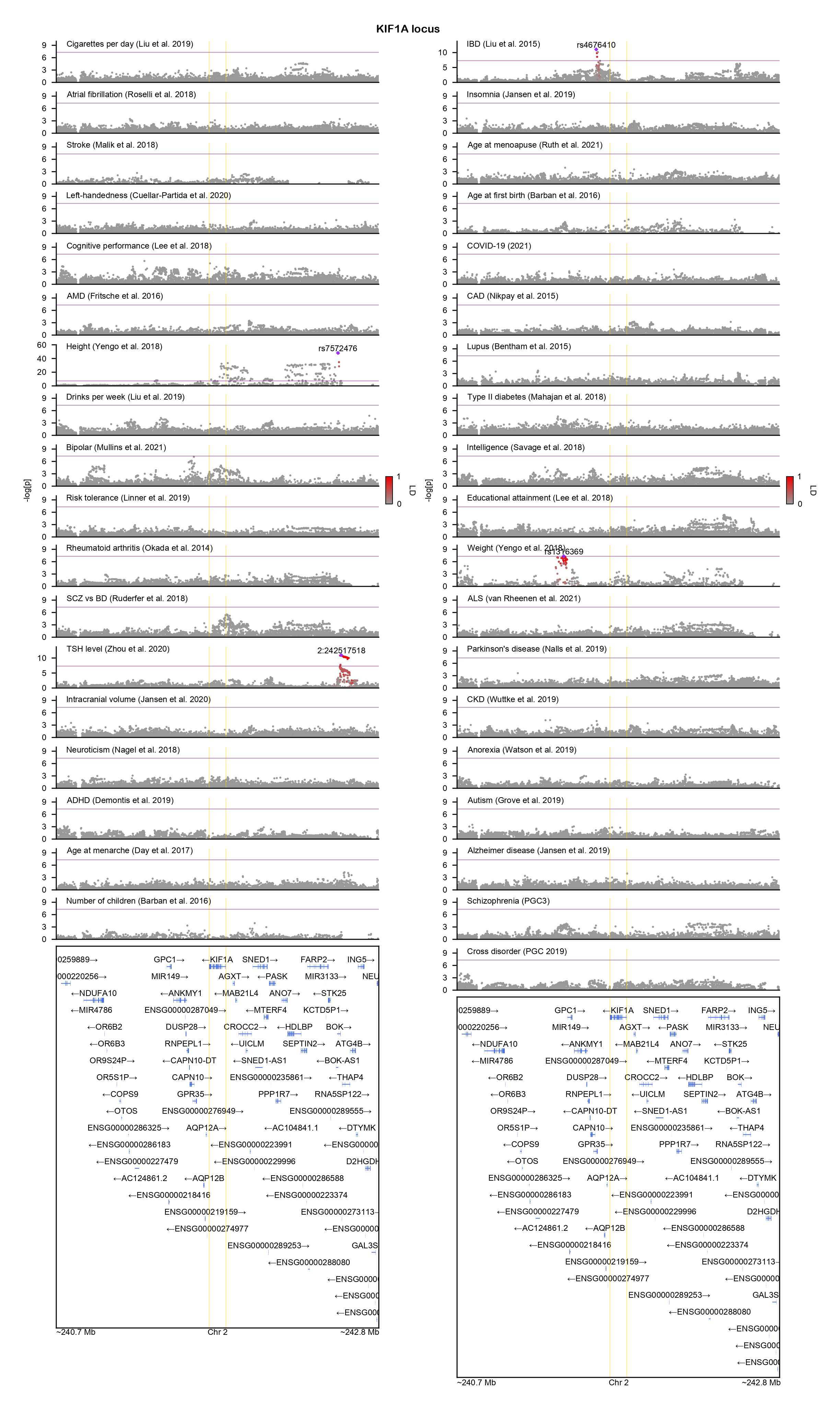

### KRAS-locuszoom.png

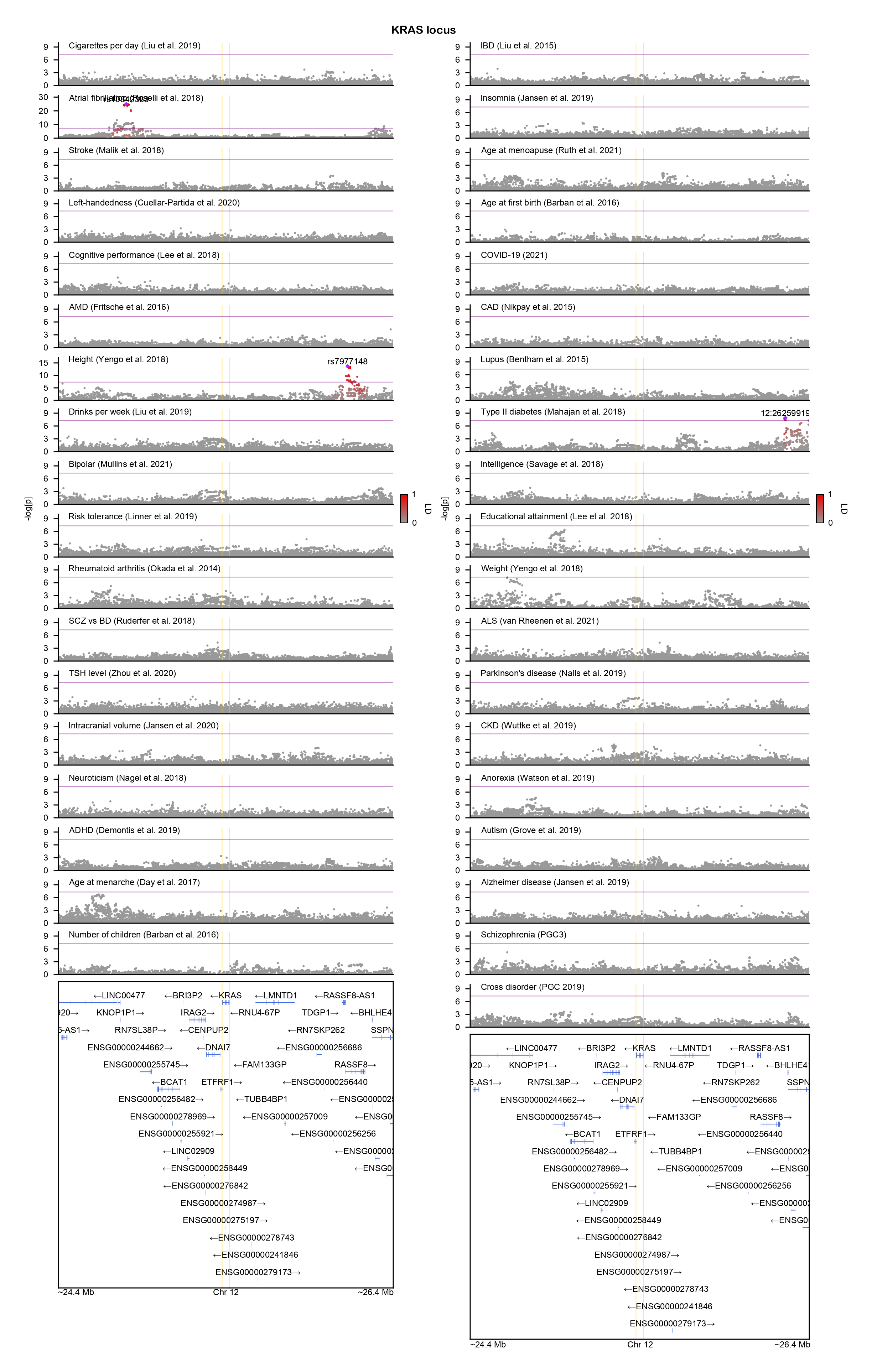

### LRRN3-locuszoom.png

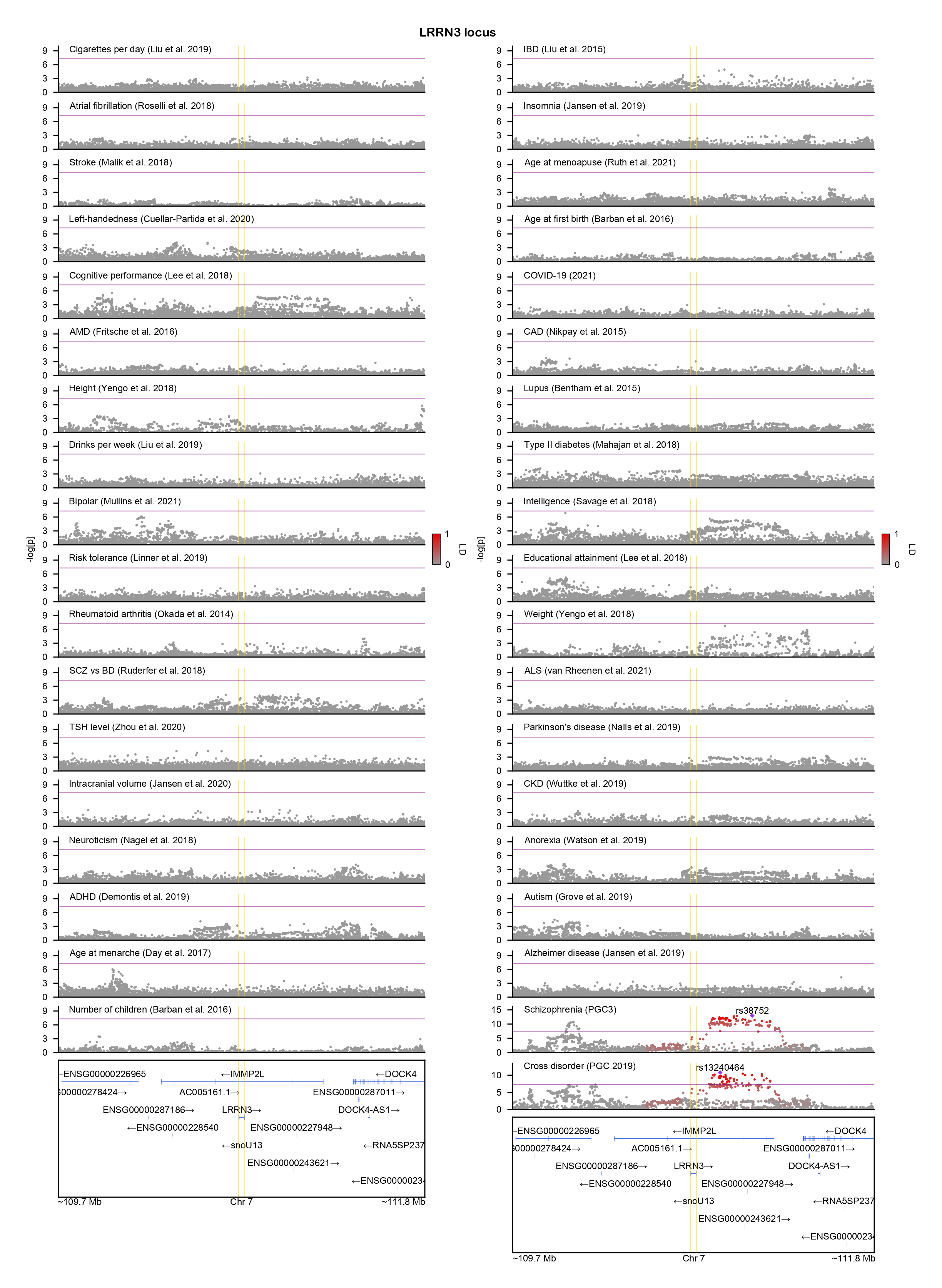

### MAP1A-locuszoom.png

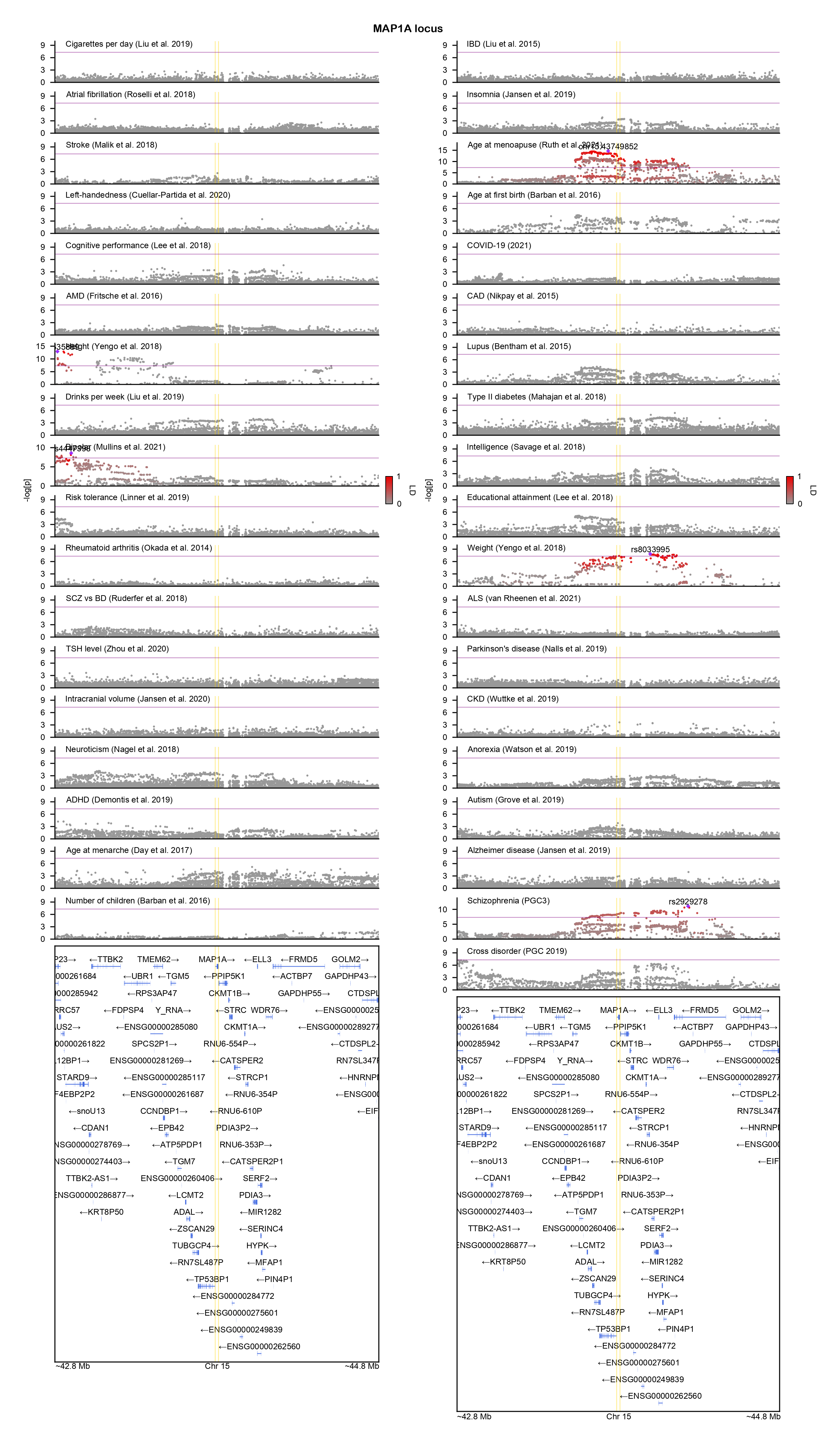

### MAP3K7-locuszoom.png

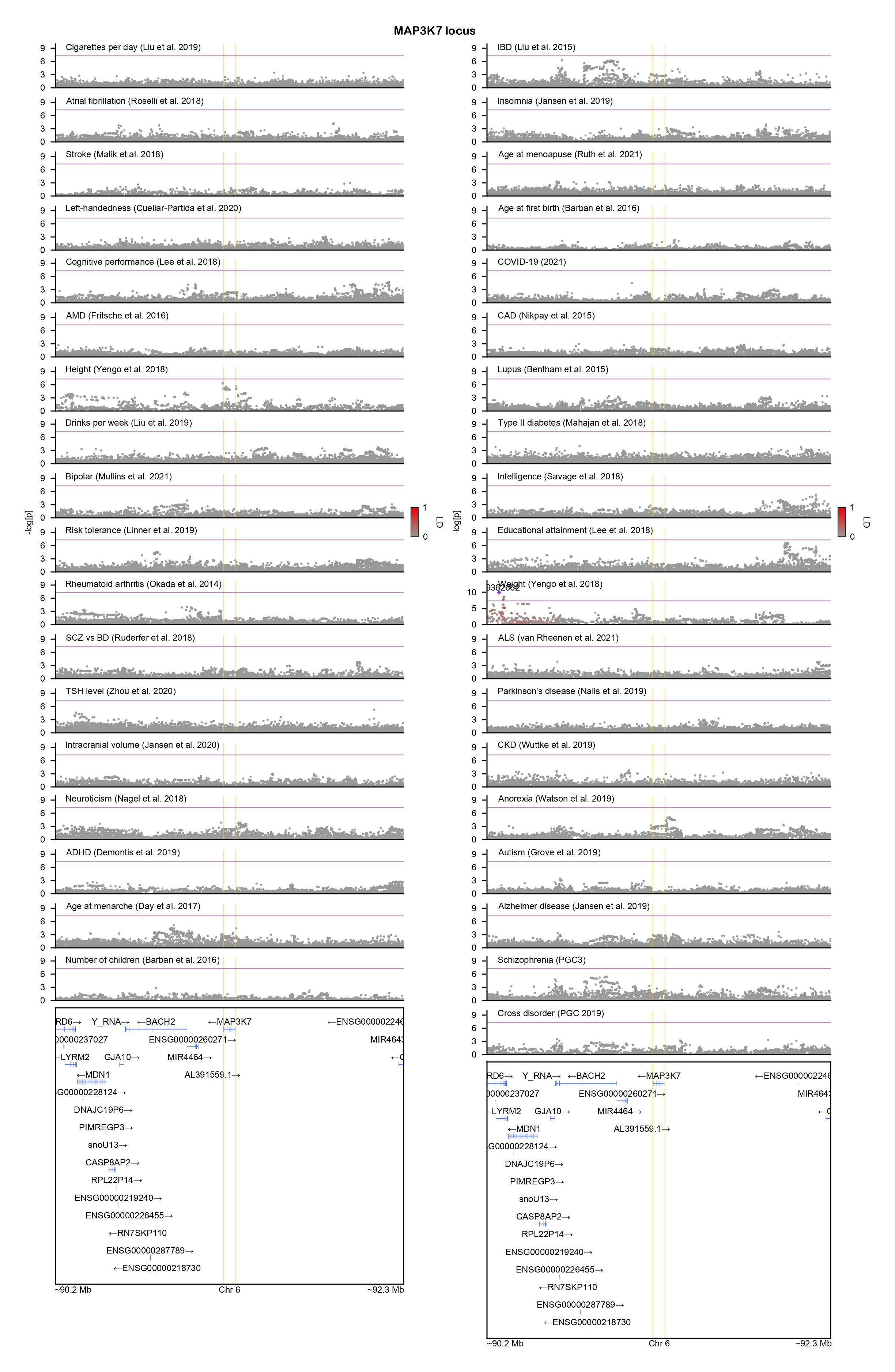

### MIB1-locuszoom.png

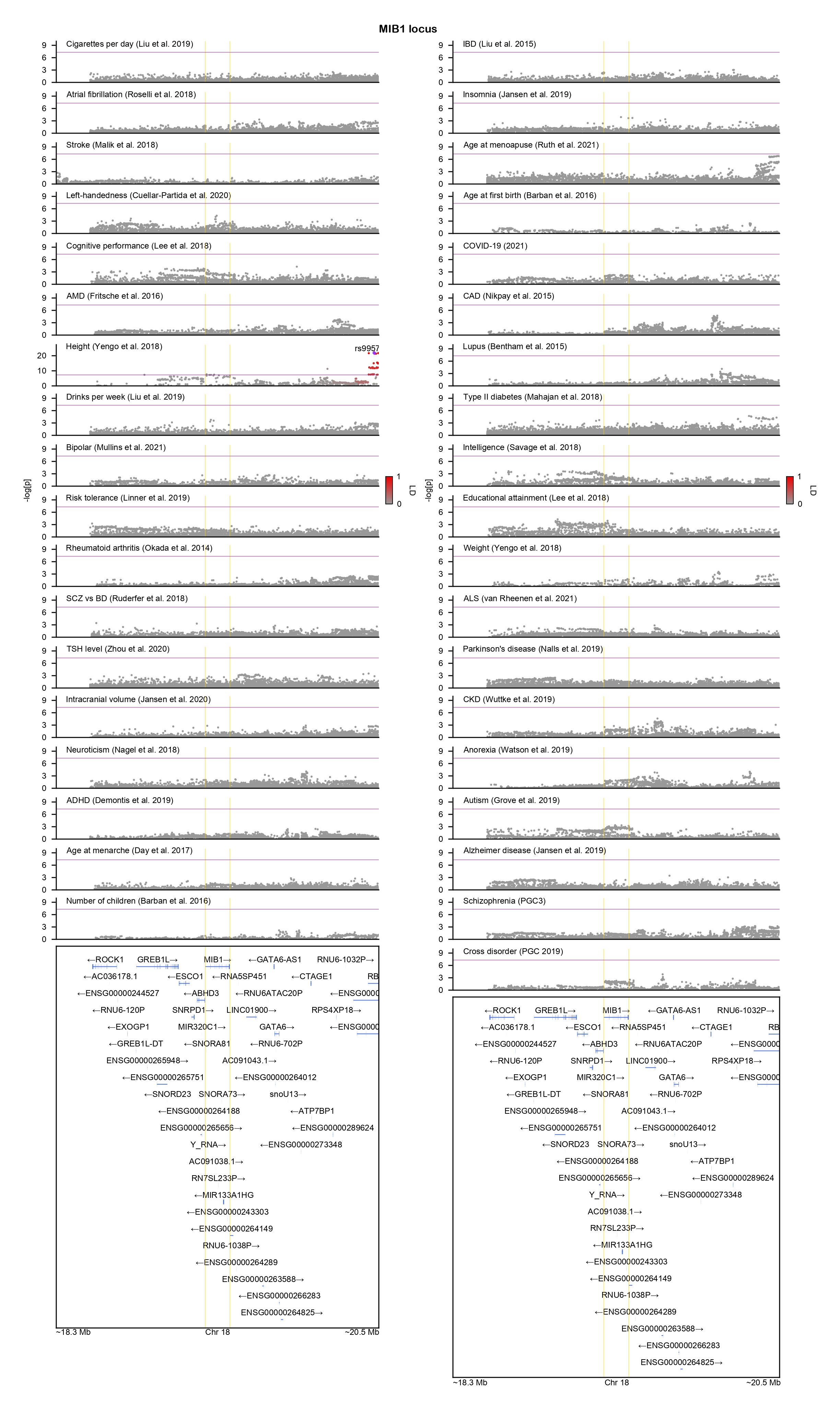

### MSL2-locuszoom.png

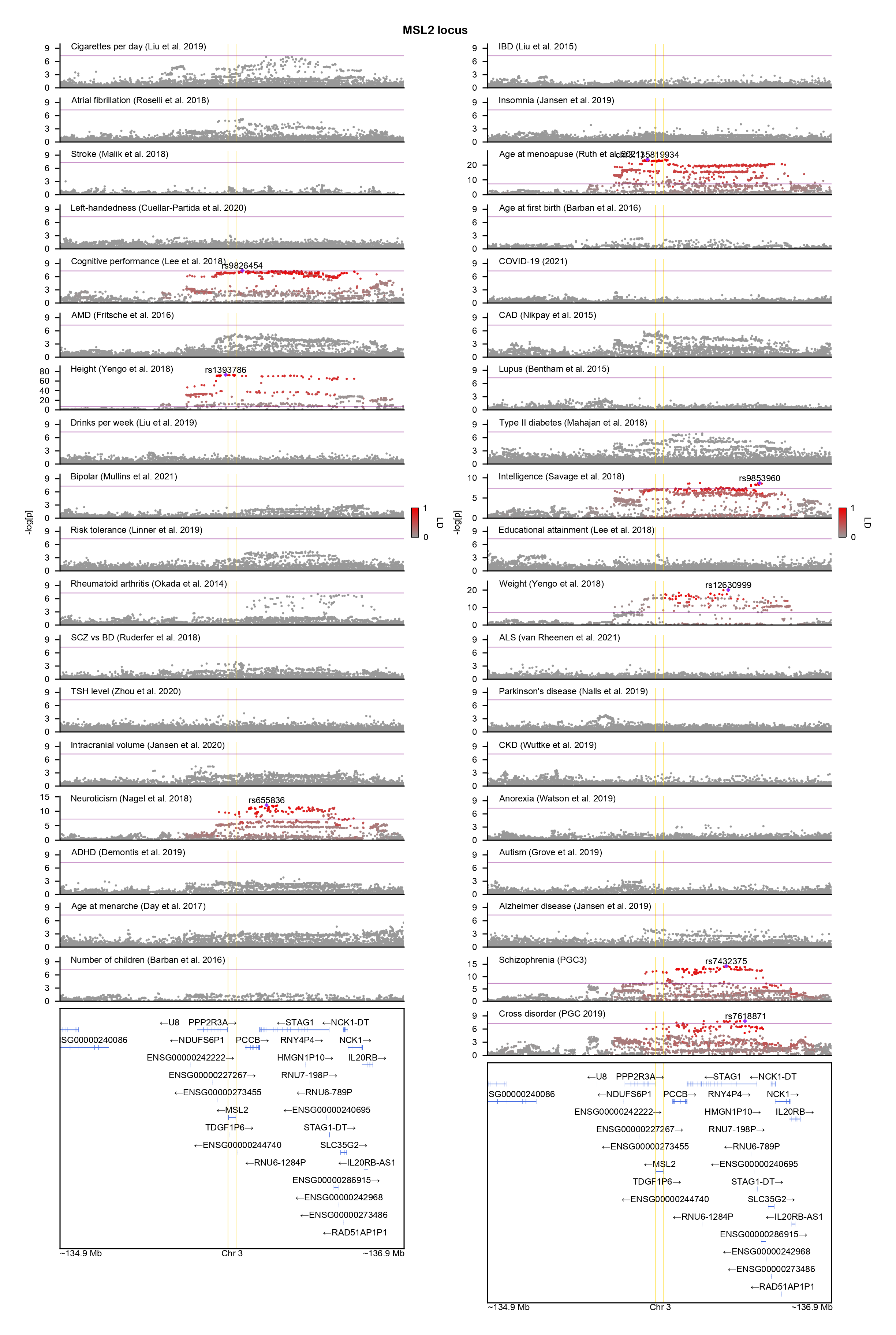

### MTOR-locuszoom.png

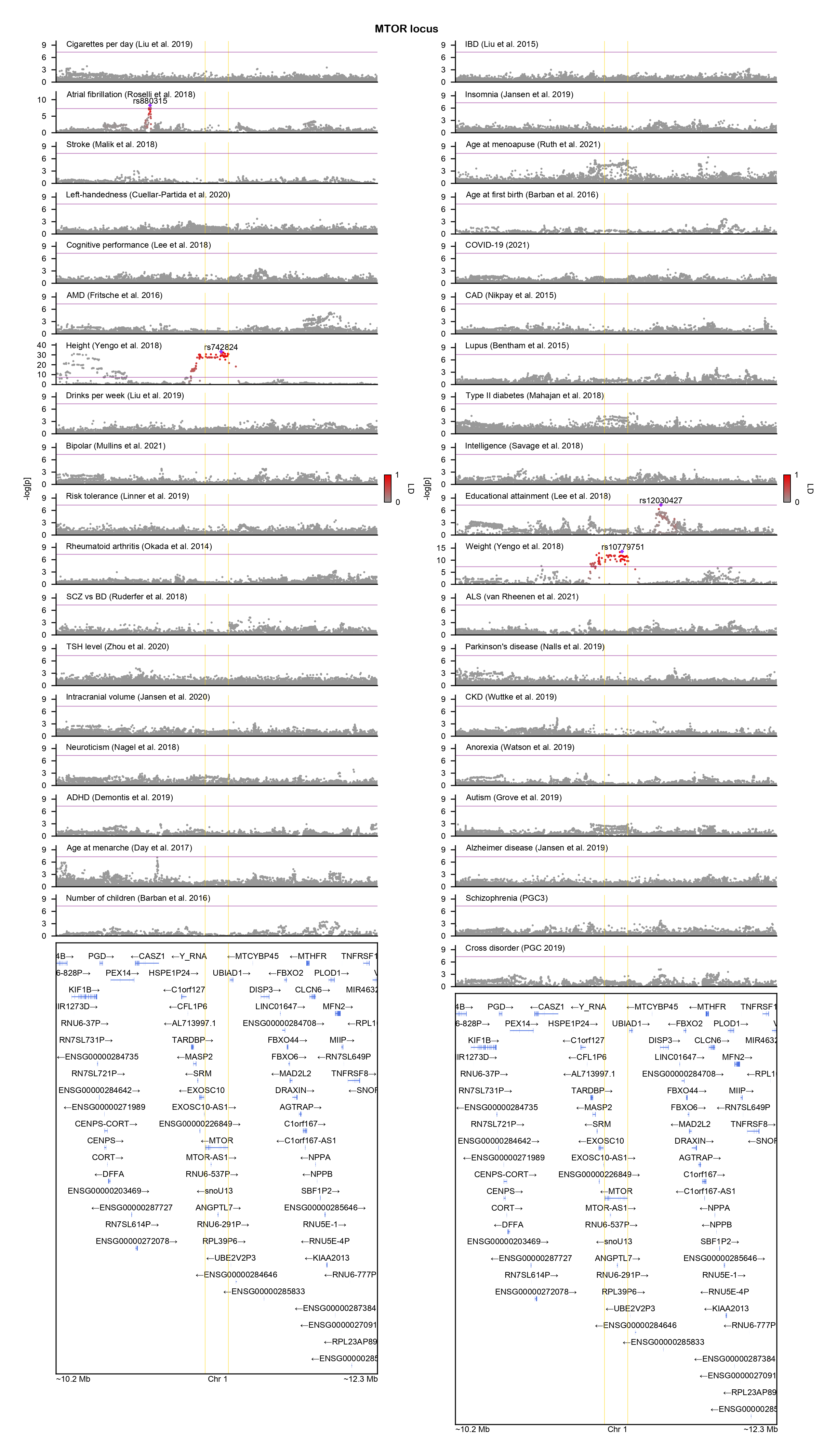

### NACC1-locuszoom.png

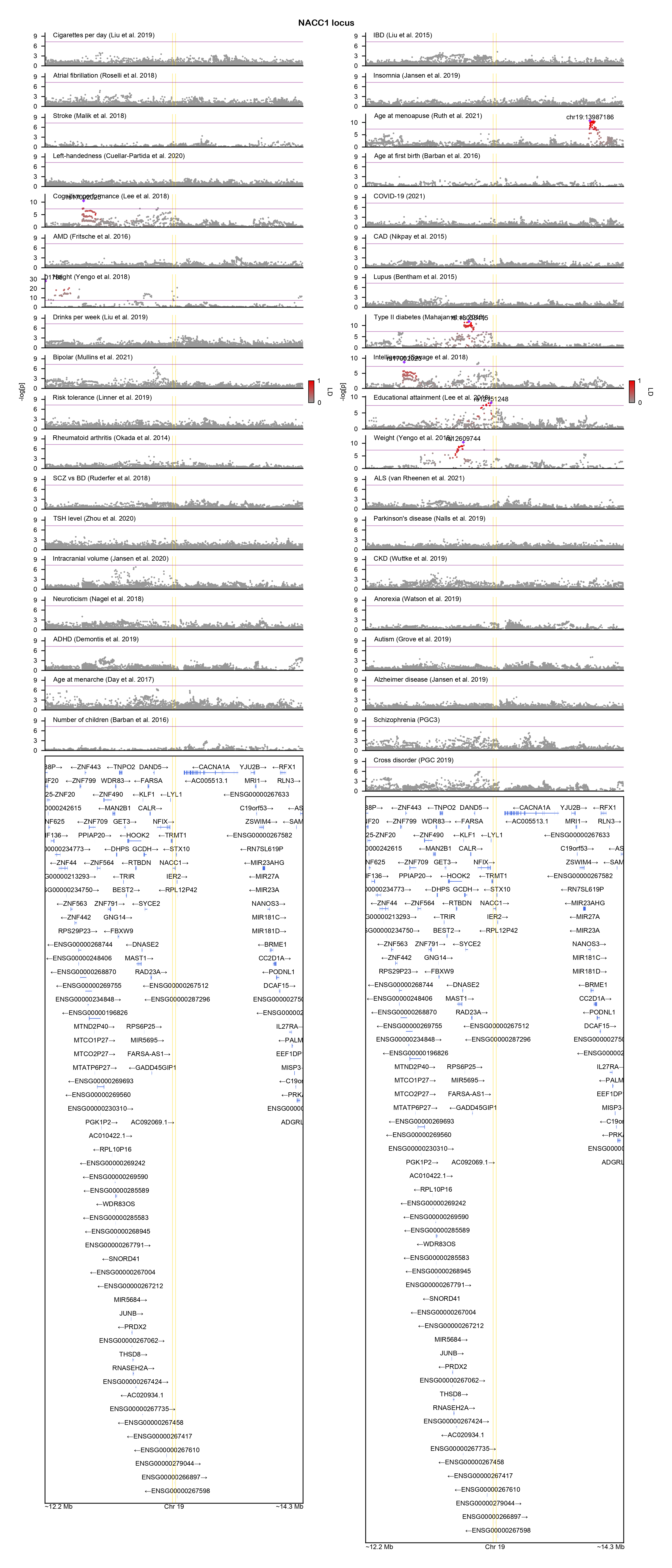

### NCOA1-locuszoom.png

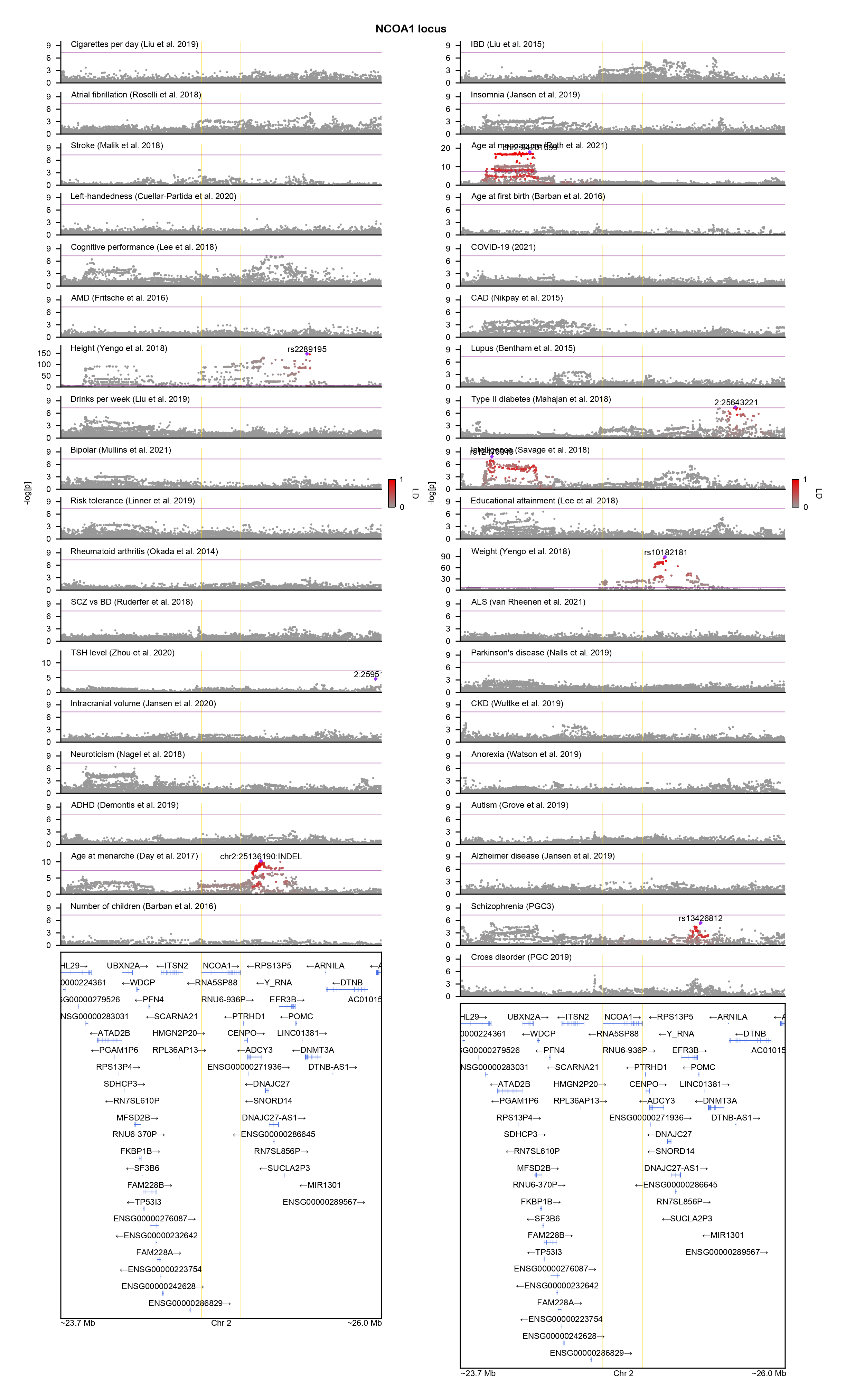

### NFIA-locuszoom.png

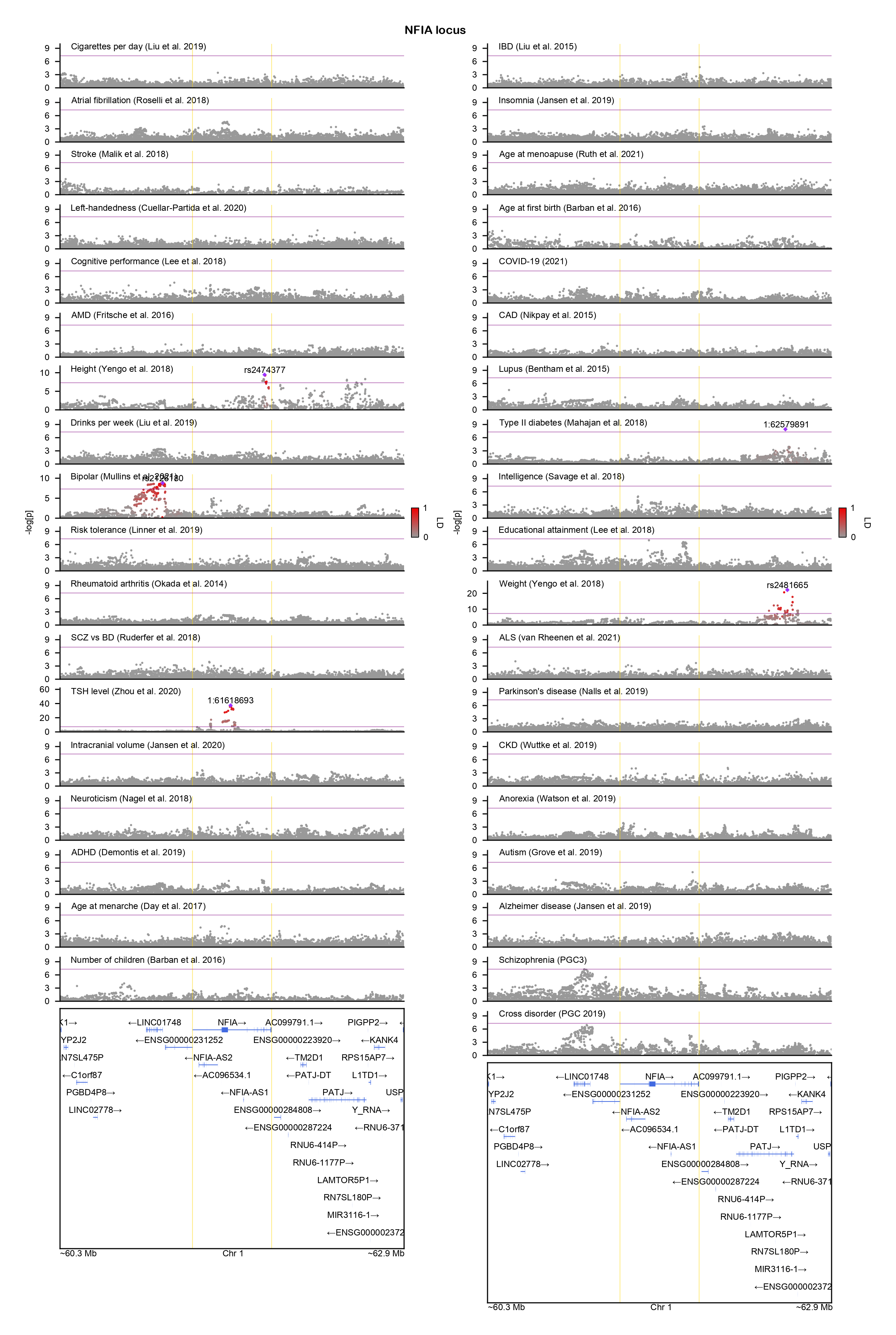

### NR4A2-locuszoom.png

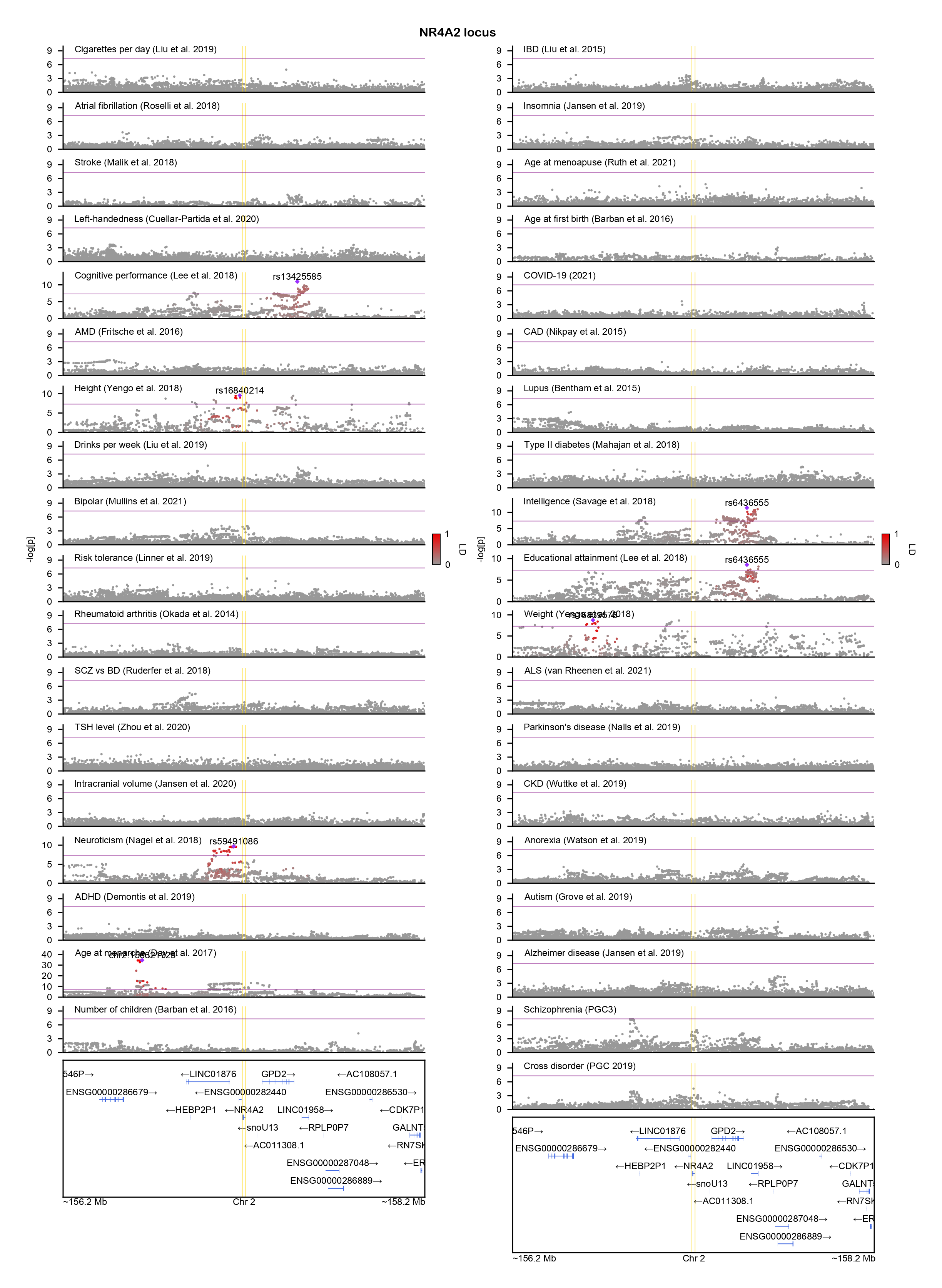

### NUP155-locuszoom.png

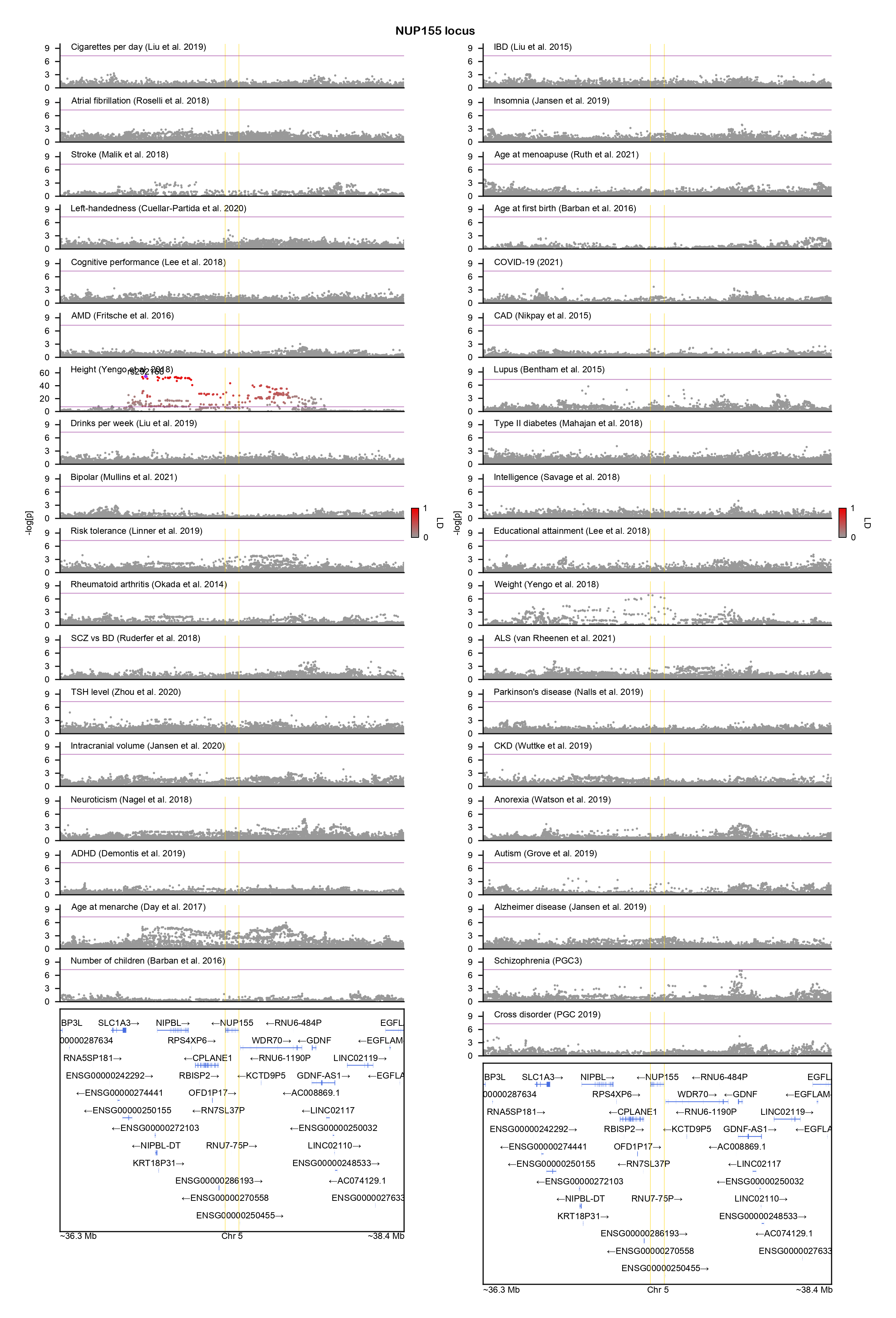

### OPCML-locuszoom.png

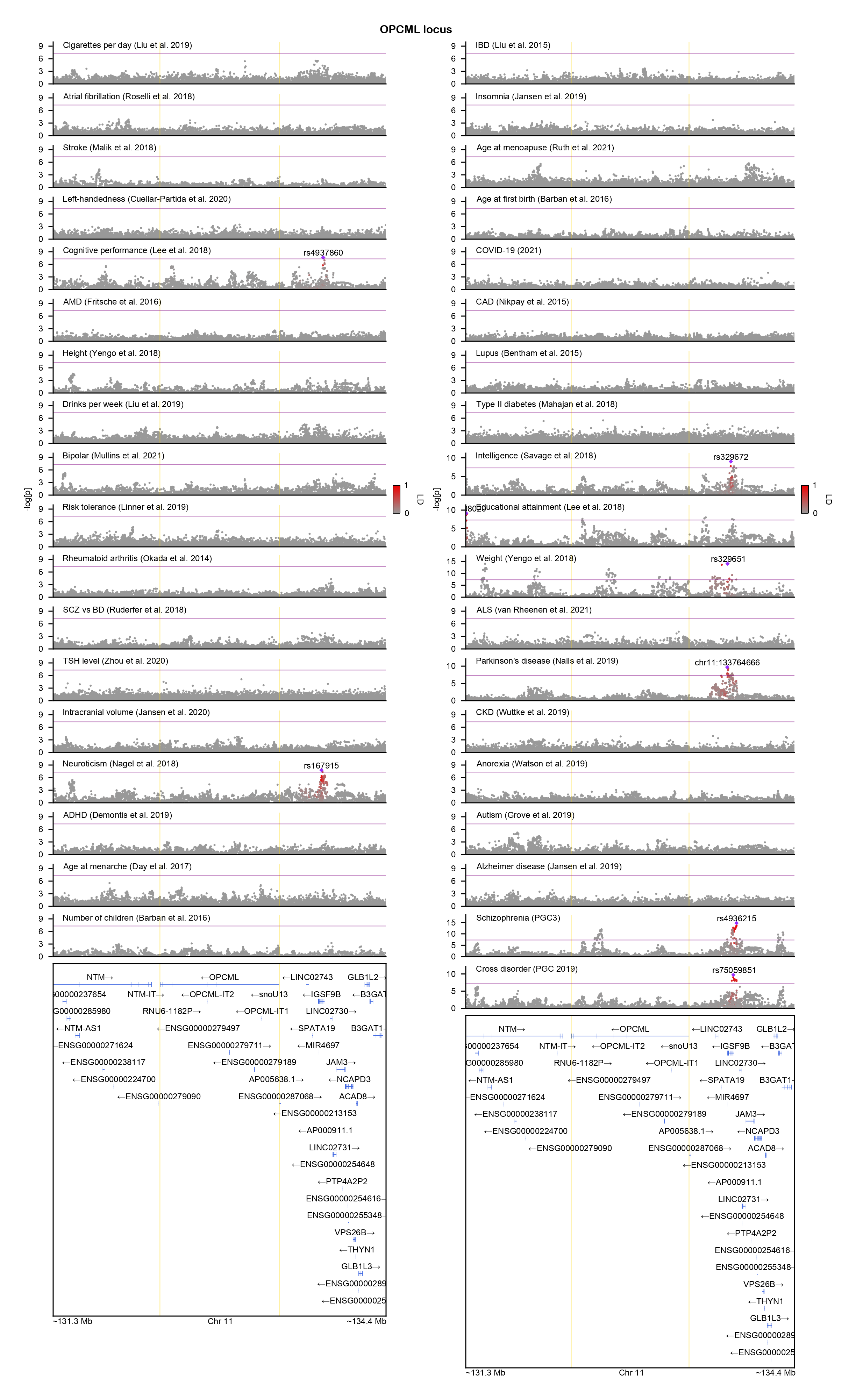

### PCDH19-locuszoom.png

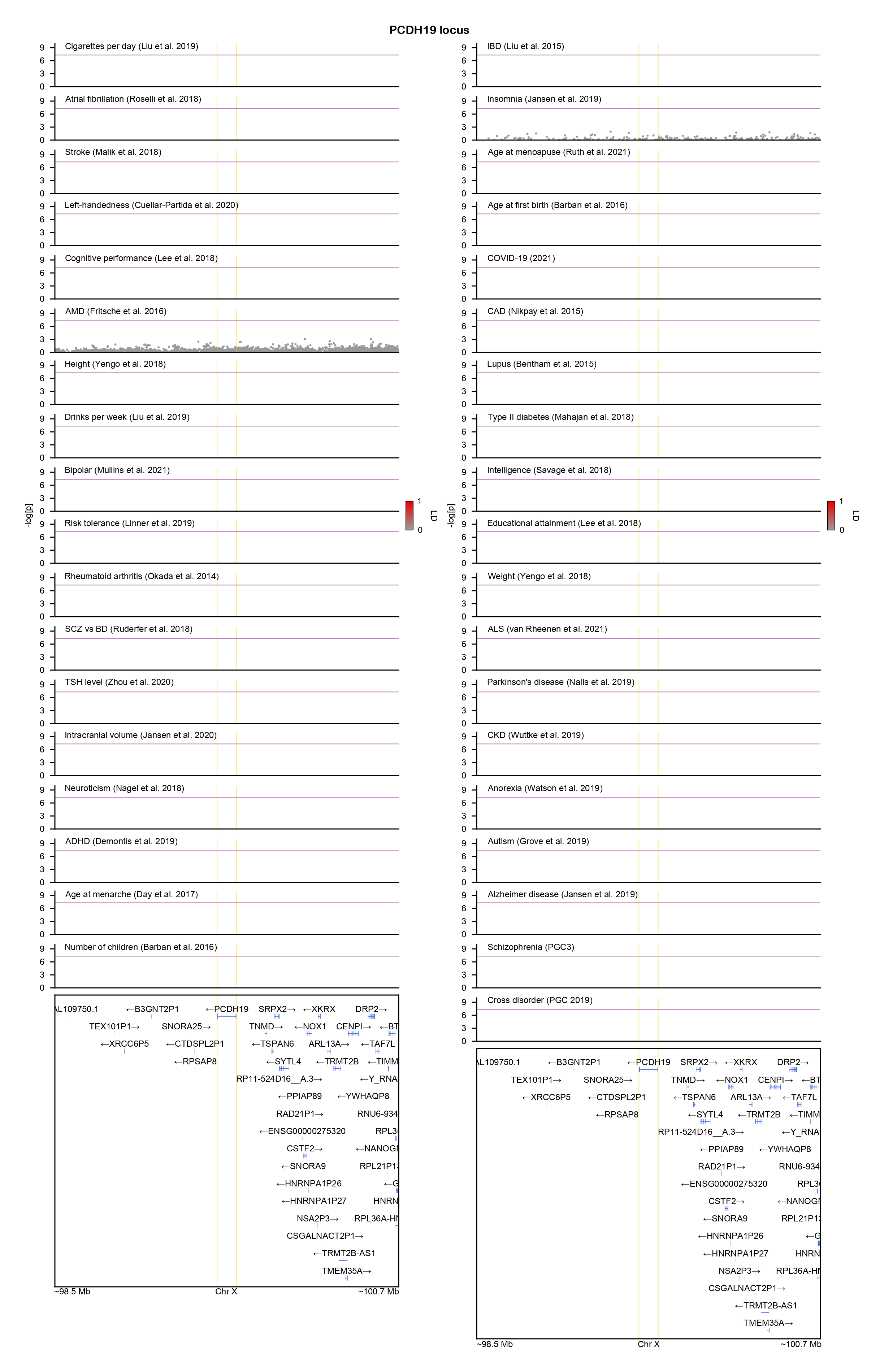

### PHF21A-locuszoom.png

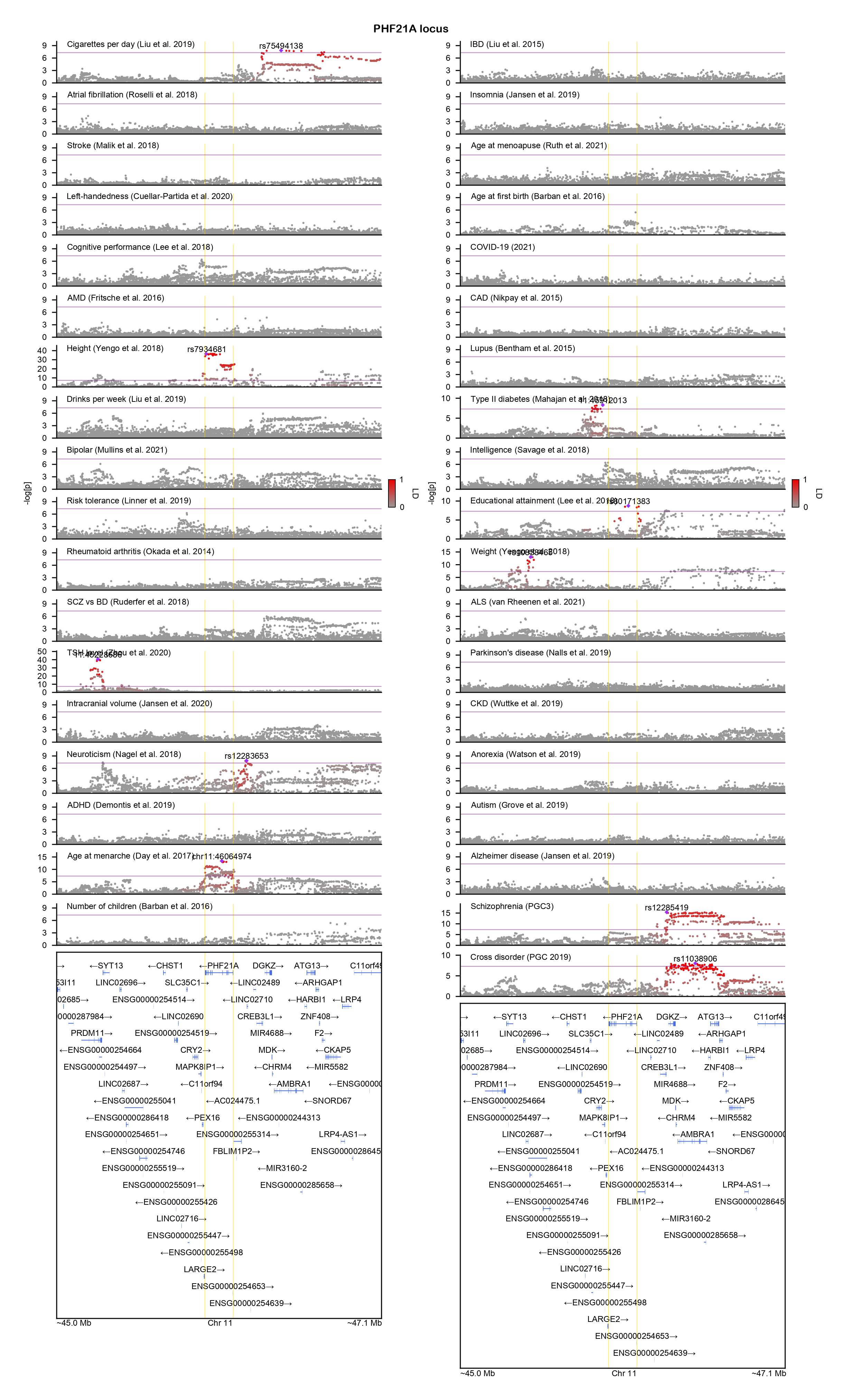

### PHIP-locuszoom.png

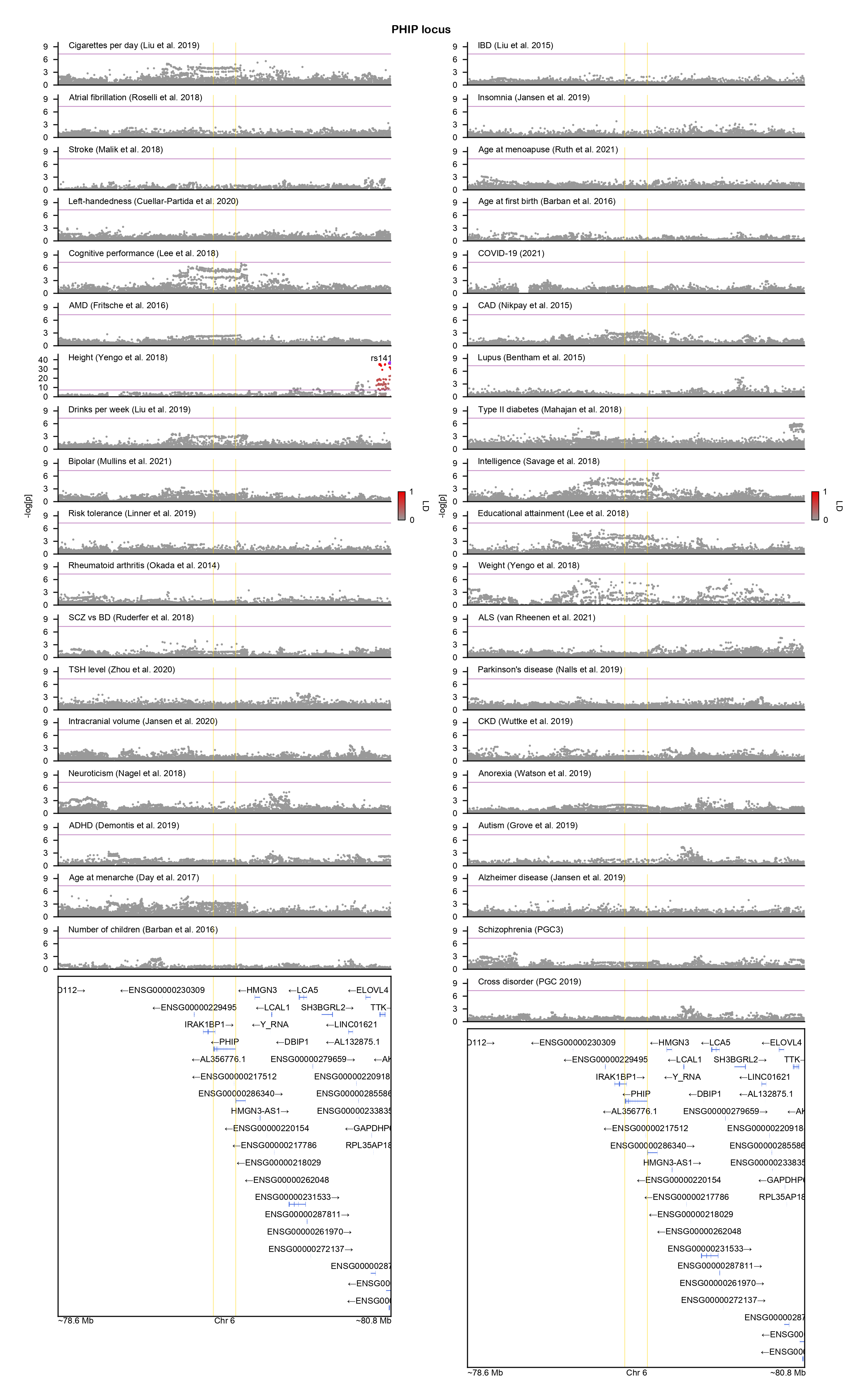

### PIK3CA-locuszoom.png

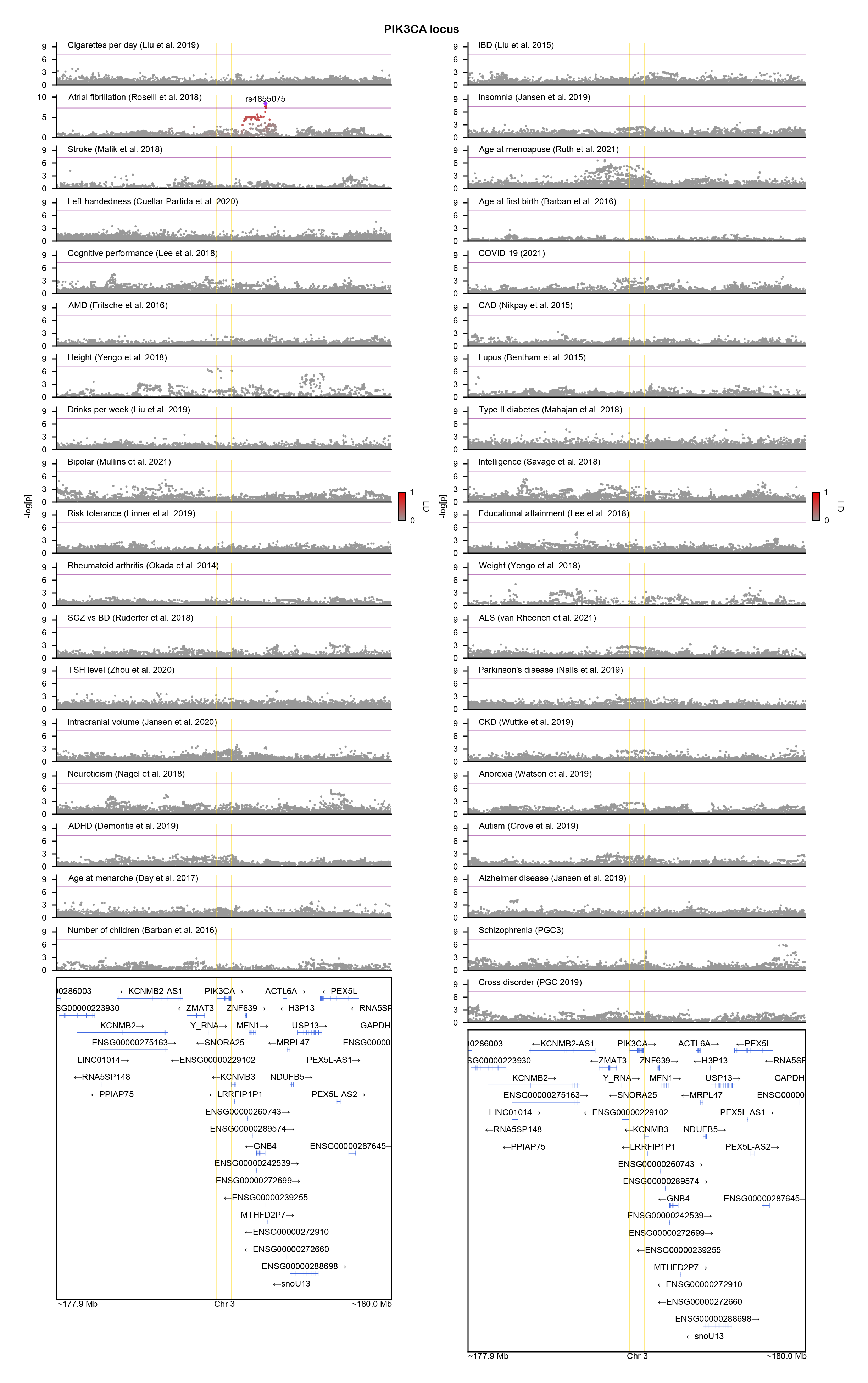

### PIK3R1-locuszoom.png

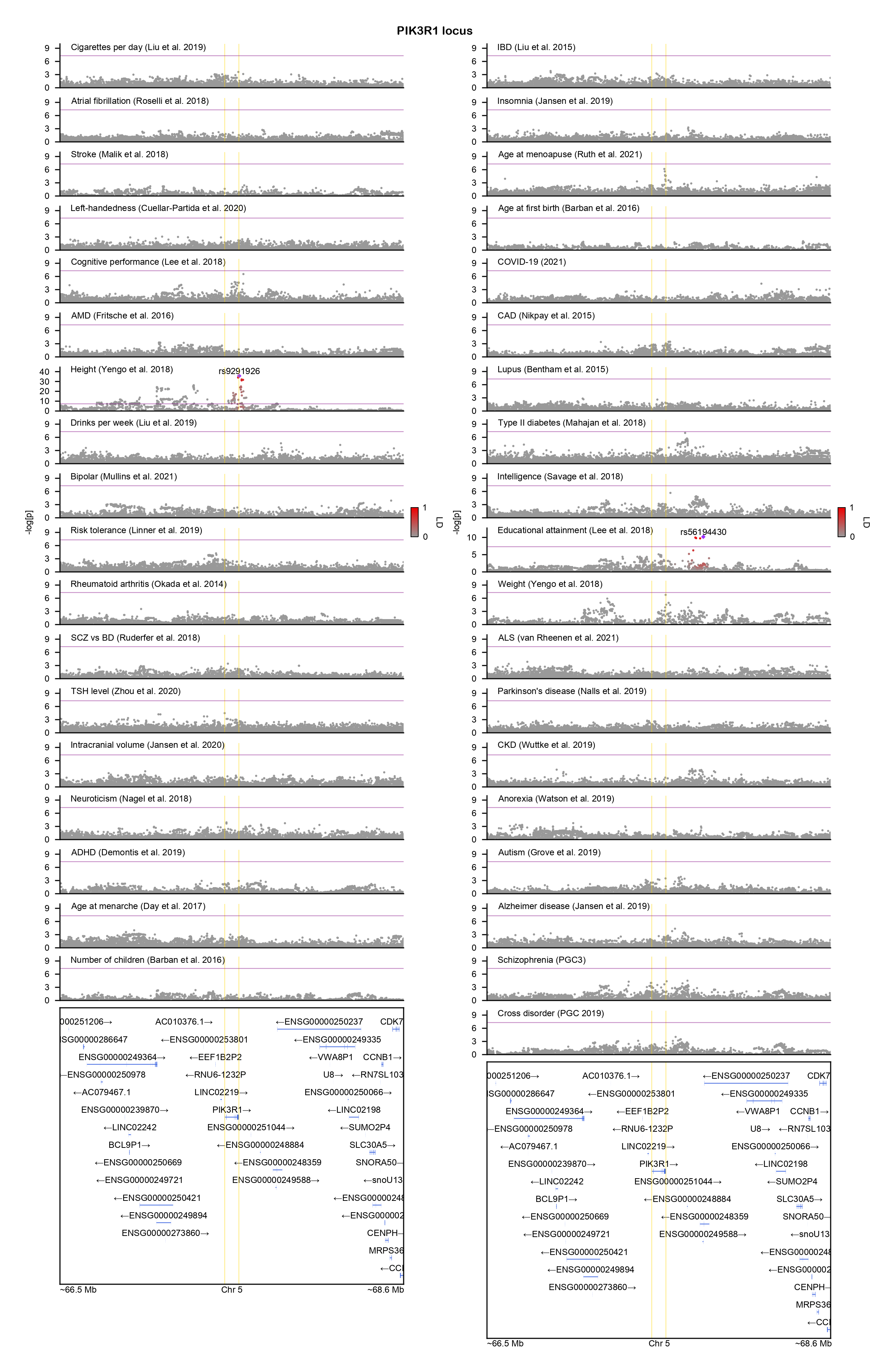

### PIK3R2-locuszoom.png

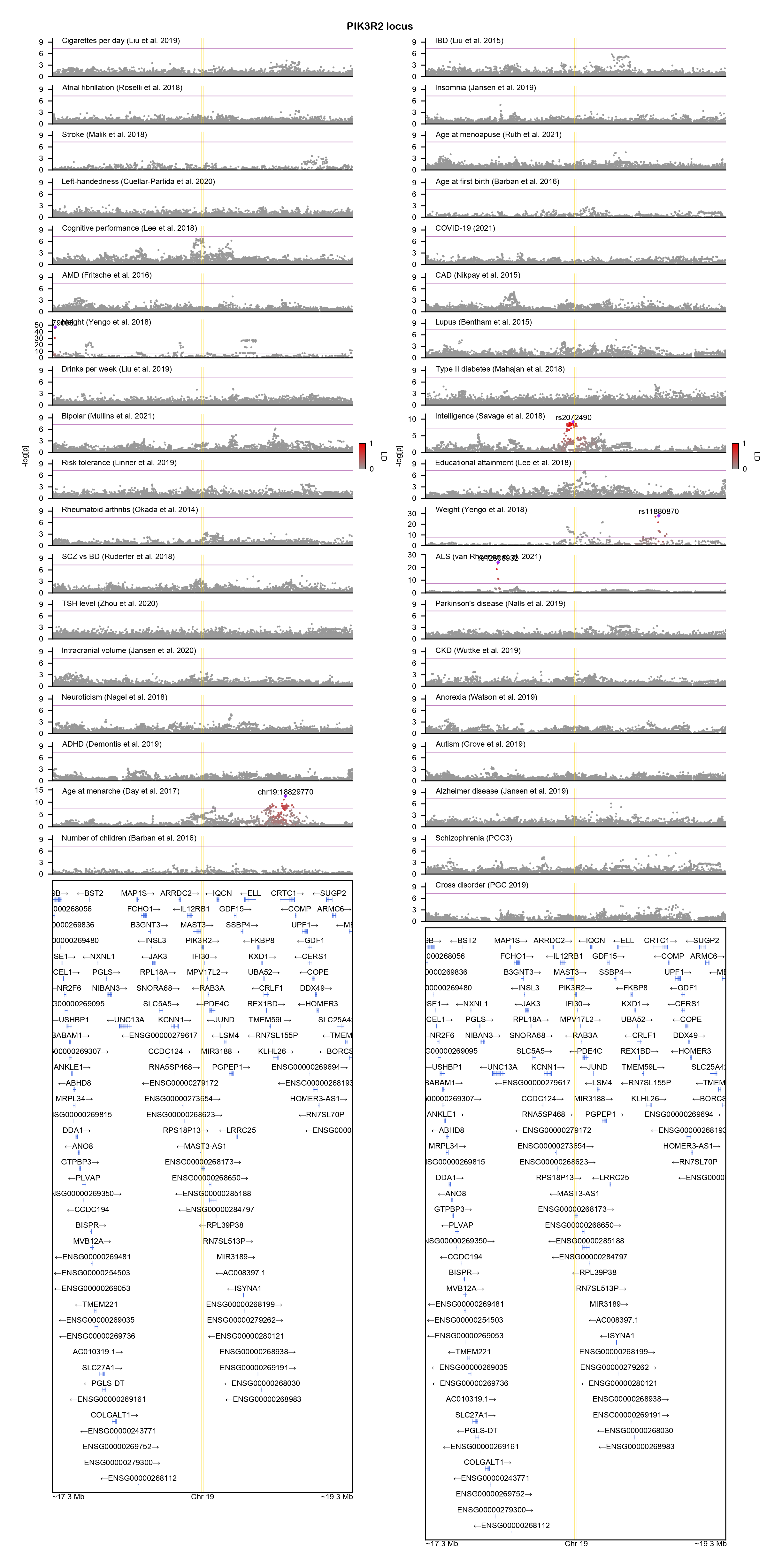

### PLCL2-locuszoom.png

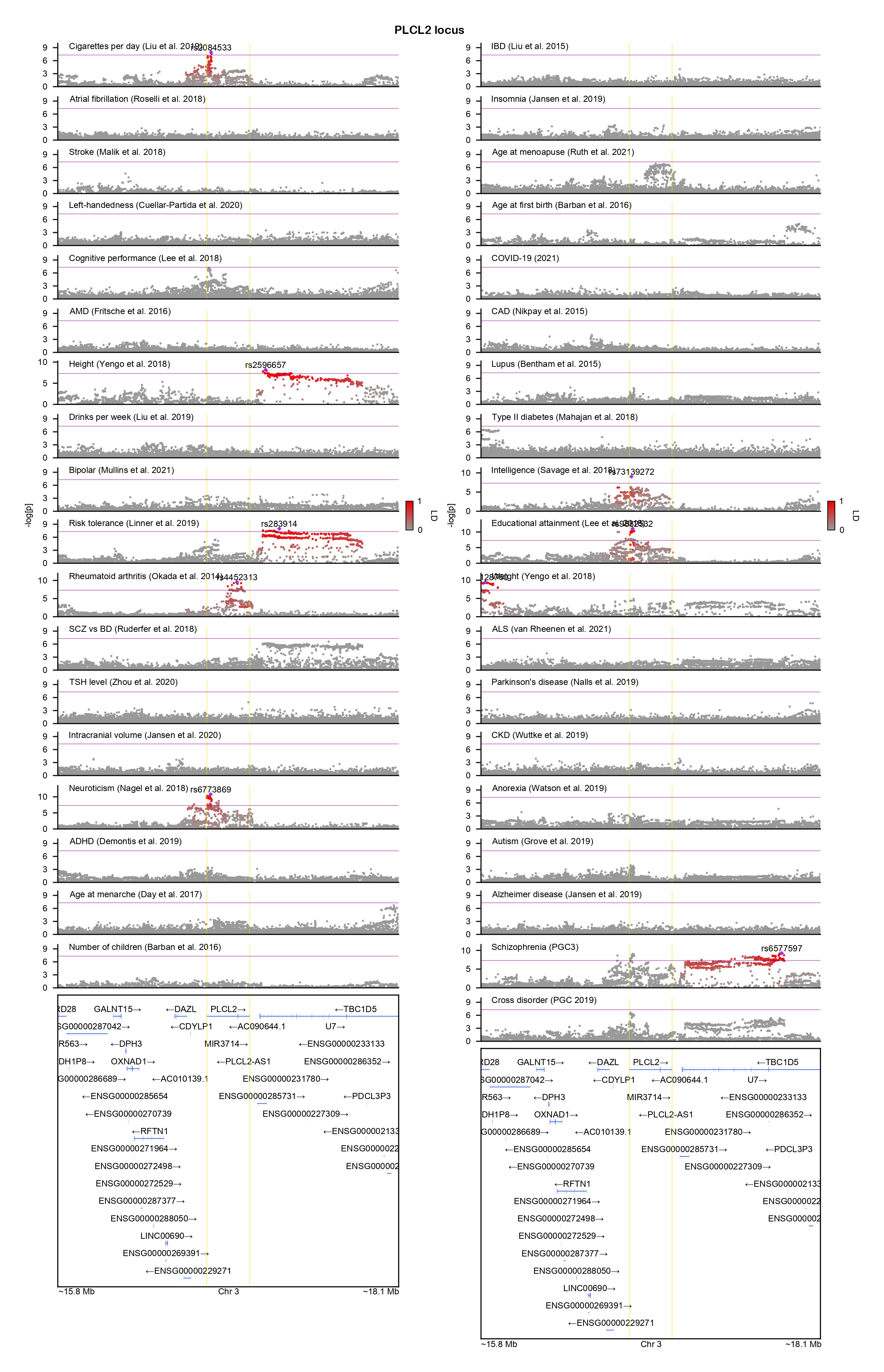

### PPP2CA-locuszoom.png

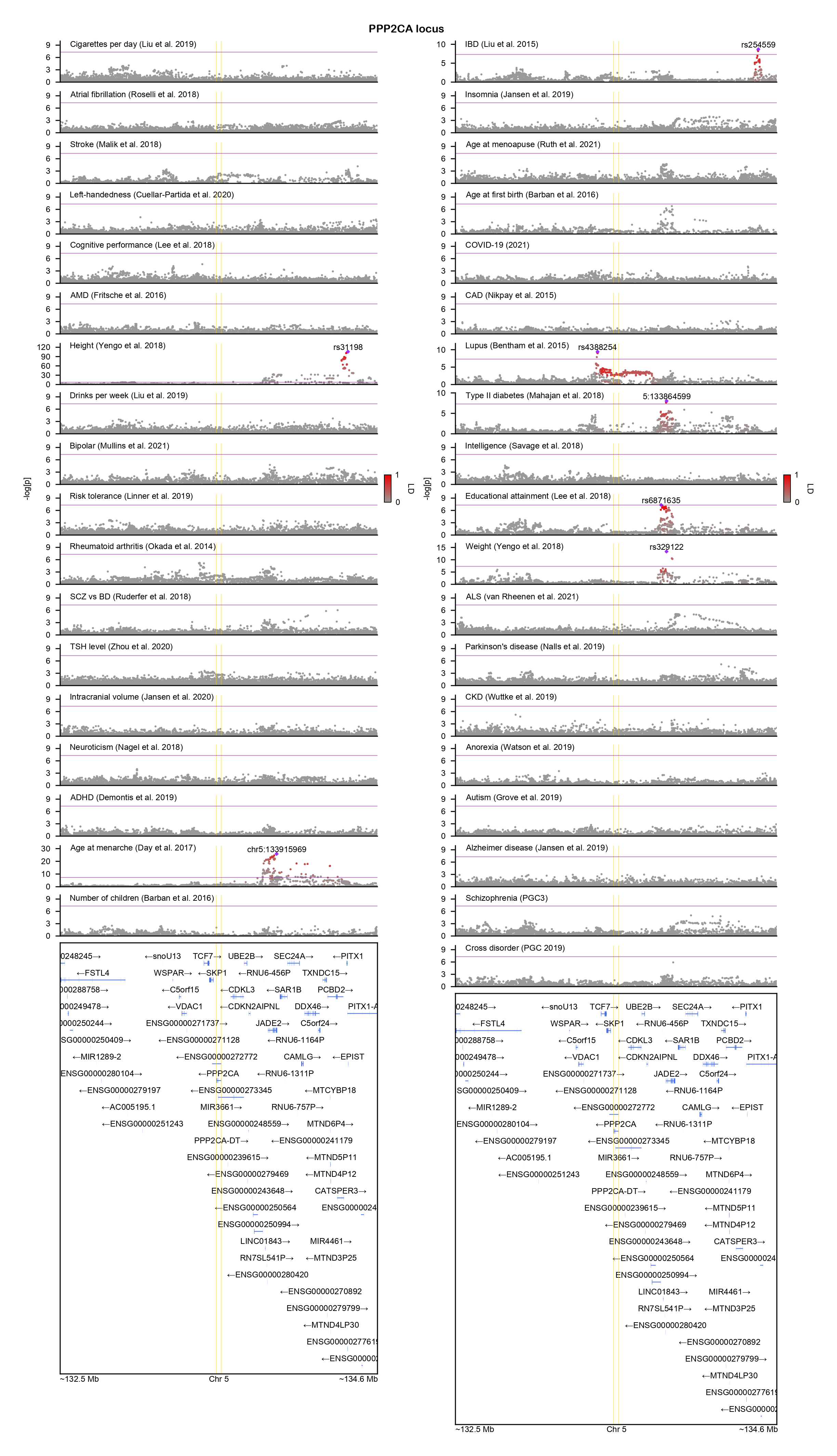

### PPP2R1A-locuszoom.png

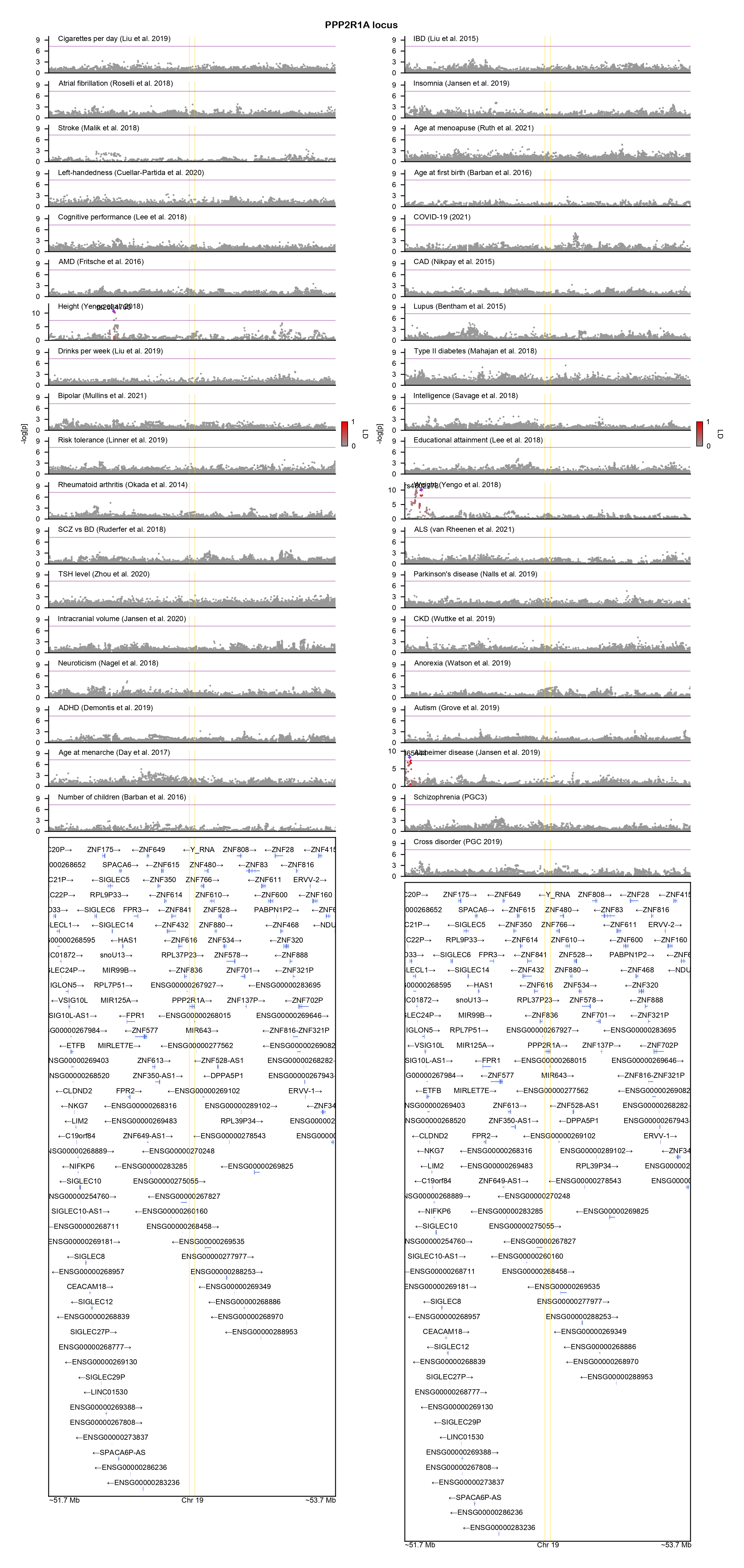

### PTCH1-locuszoom.png

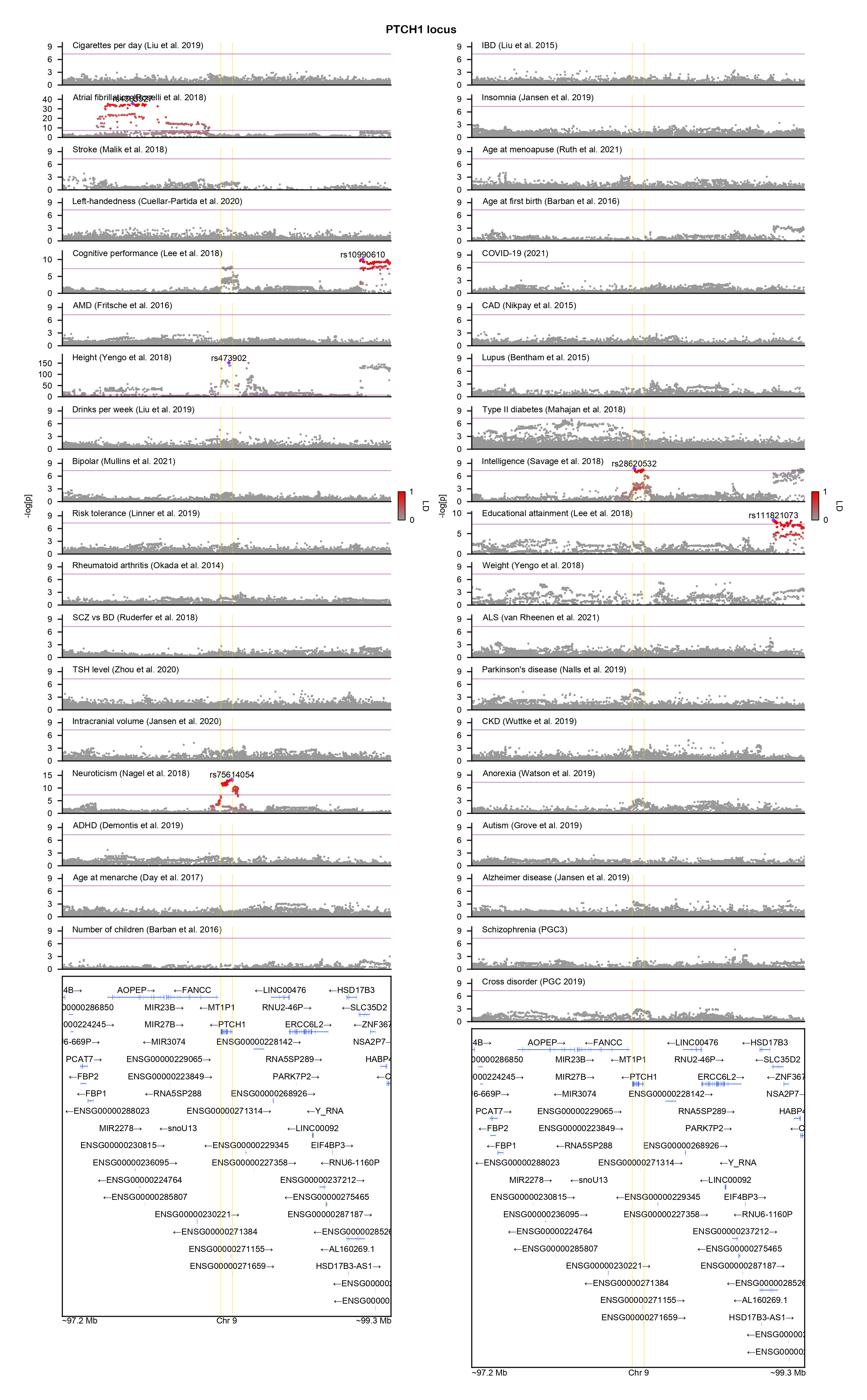

### PTK7-locuszoom.png

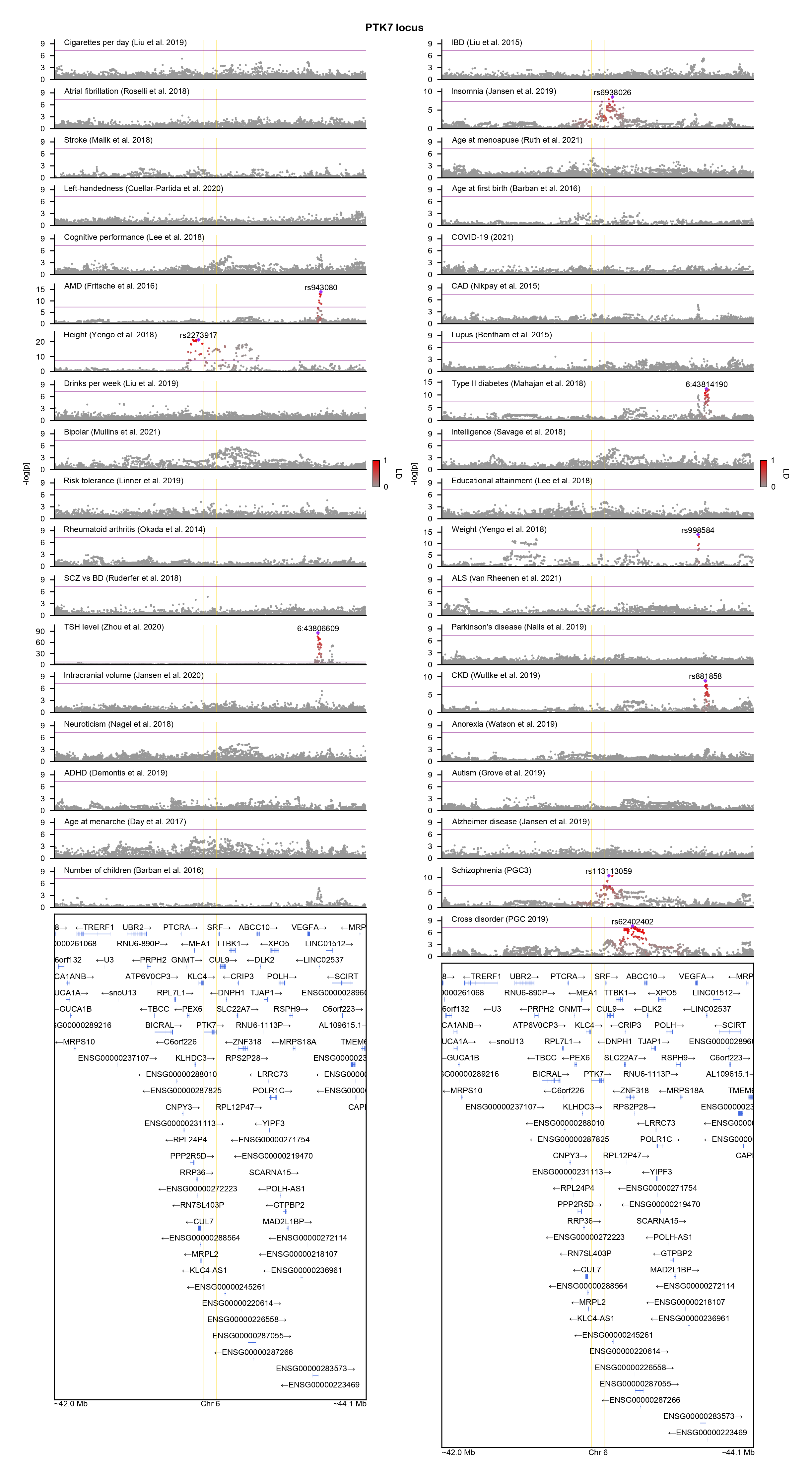

### PTPN11-locuszoom.png

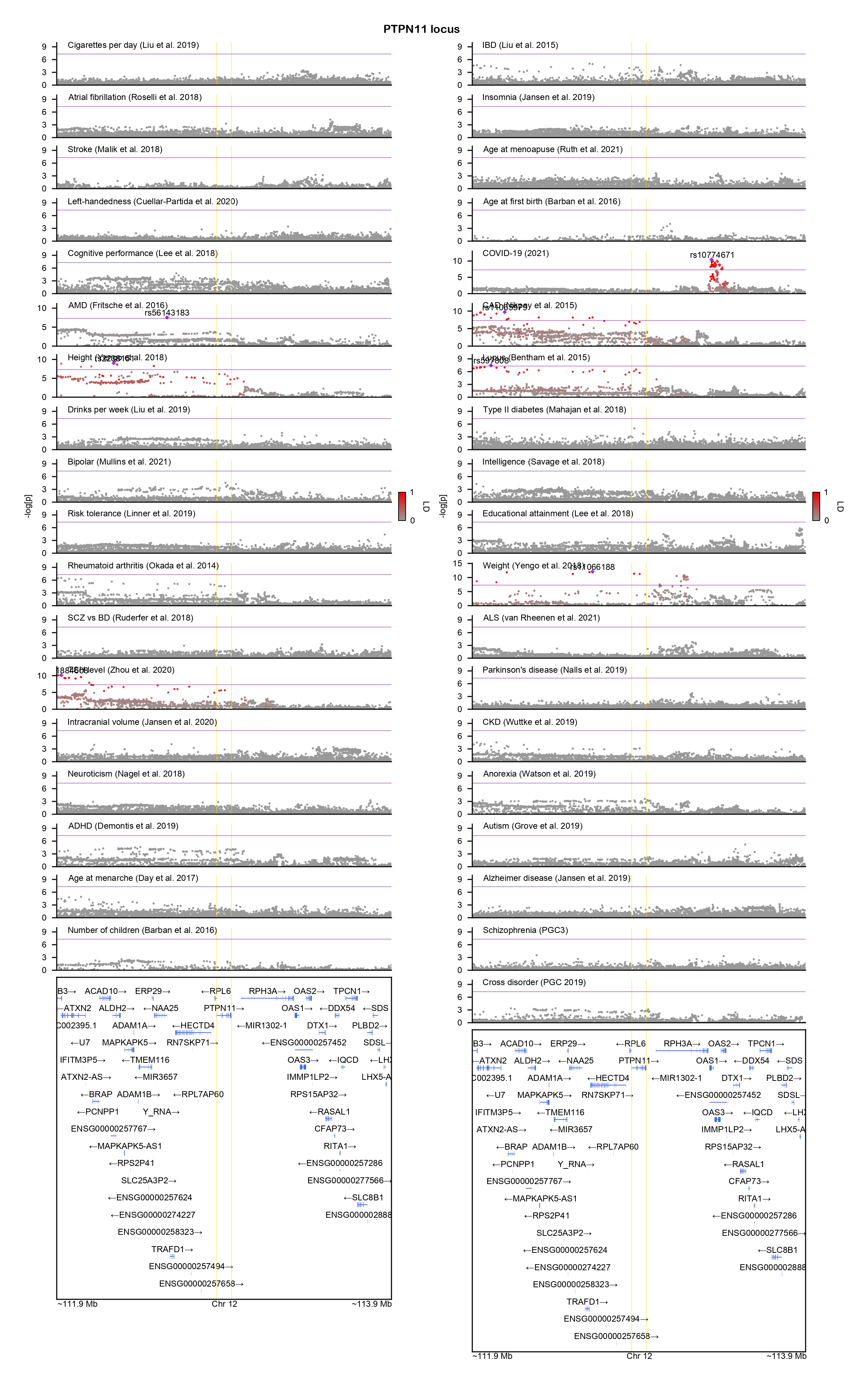

### RAC1-locuszoom.png

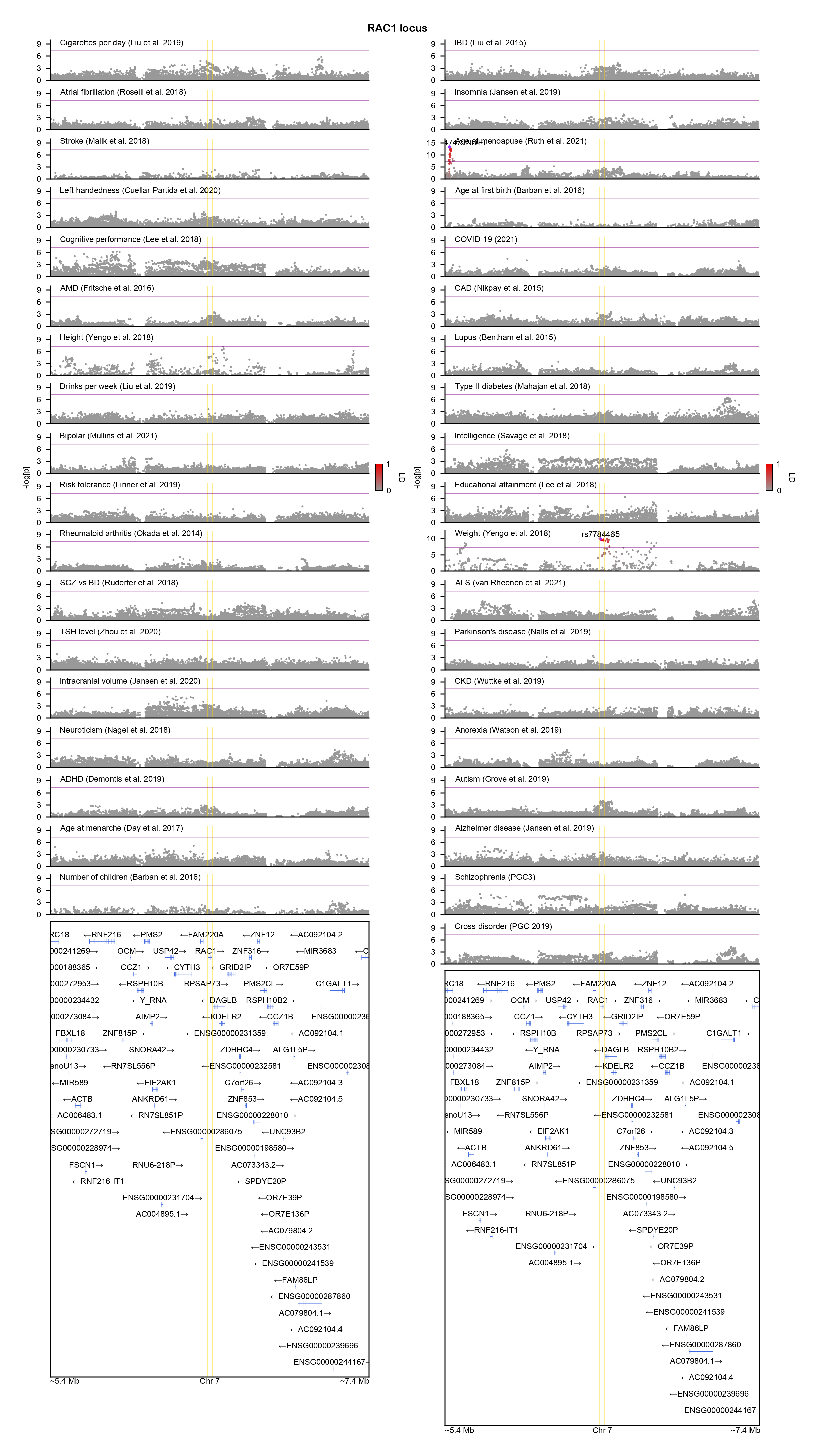

### RFX3-locuszoom.png

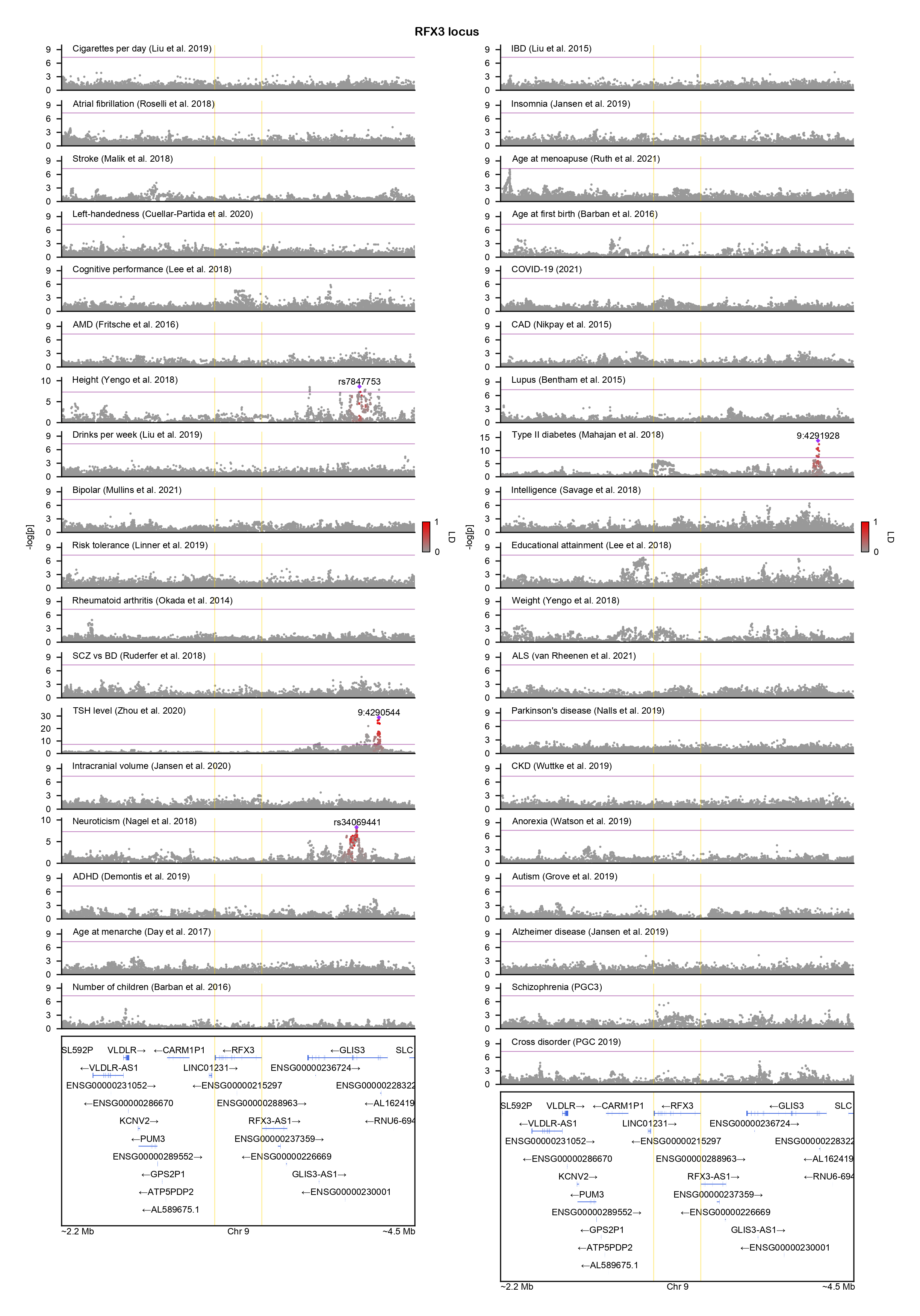

### RGS6-locuszoom.png

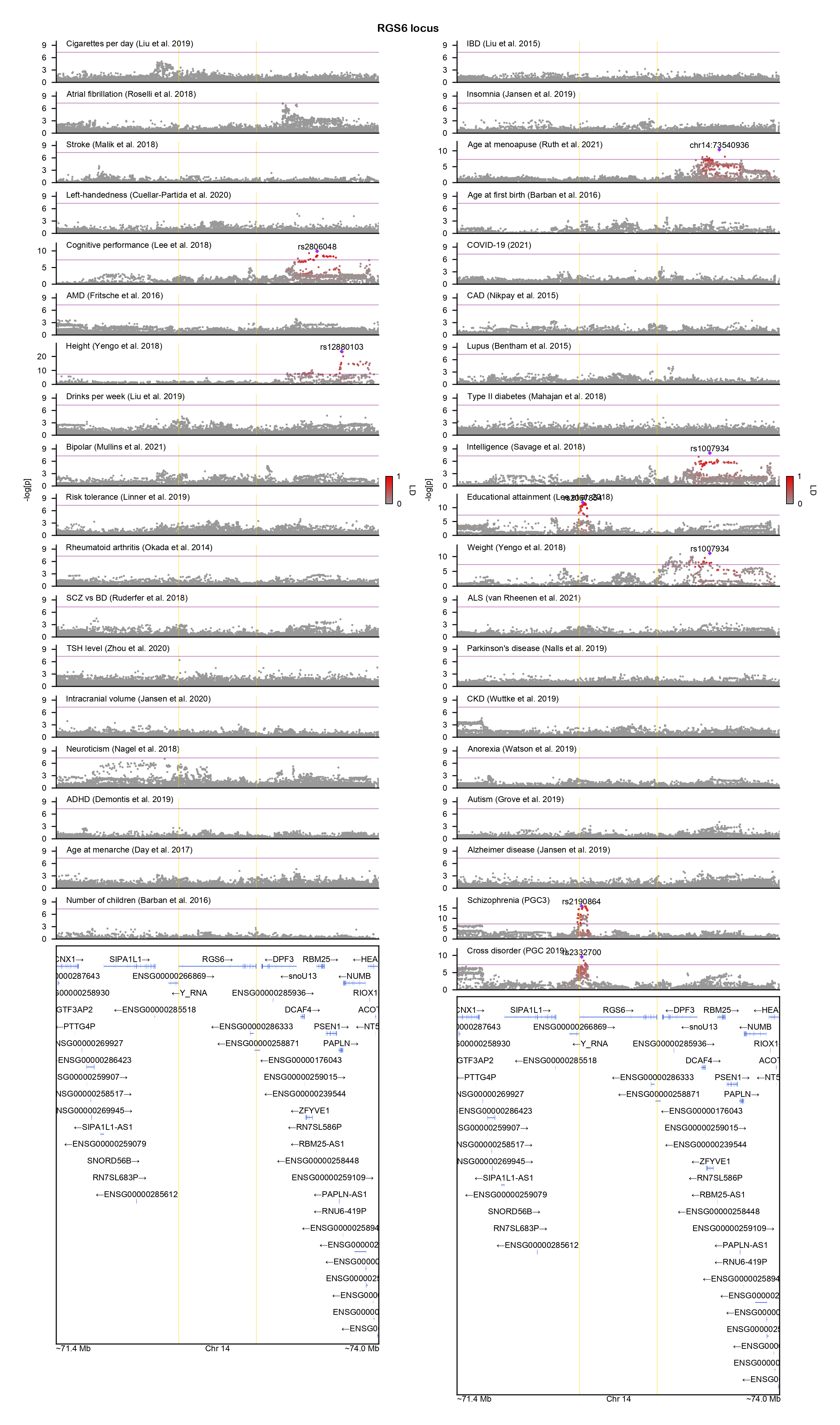

### RPS6KA3-locuszoom.png

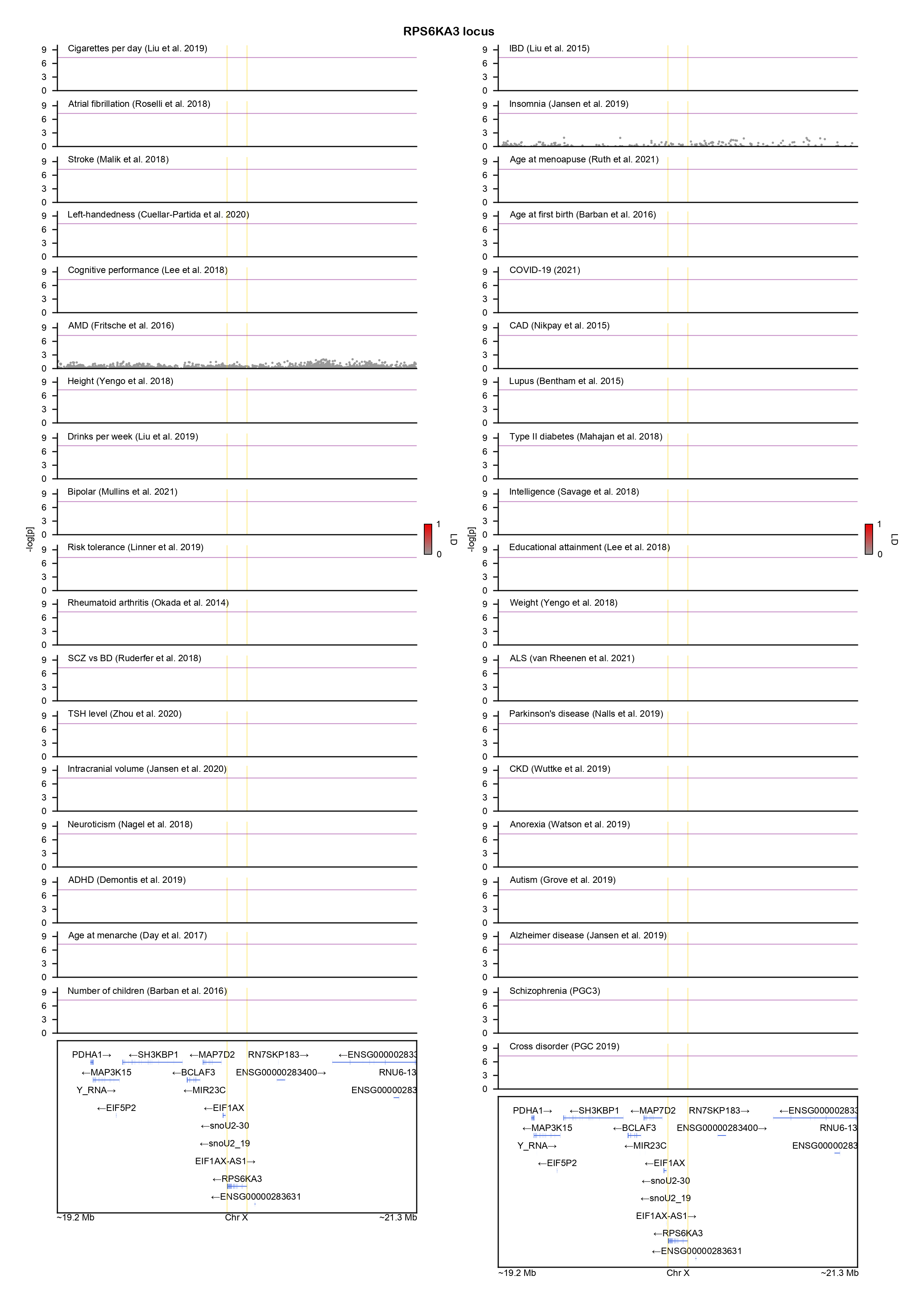

### SCN1A-locuszoom.png

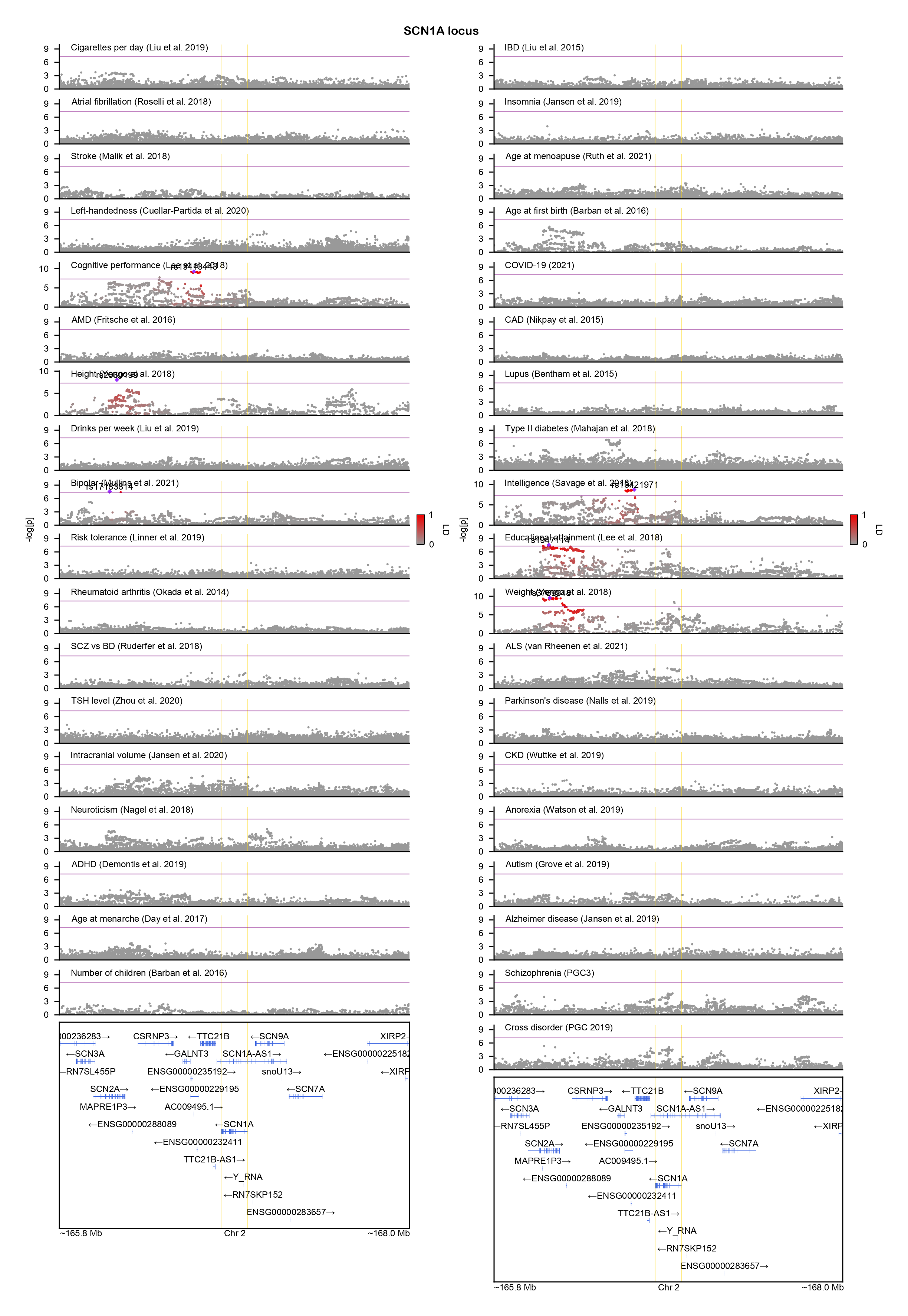

### SCN2A-locuszoom.png

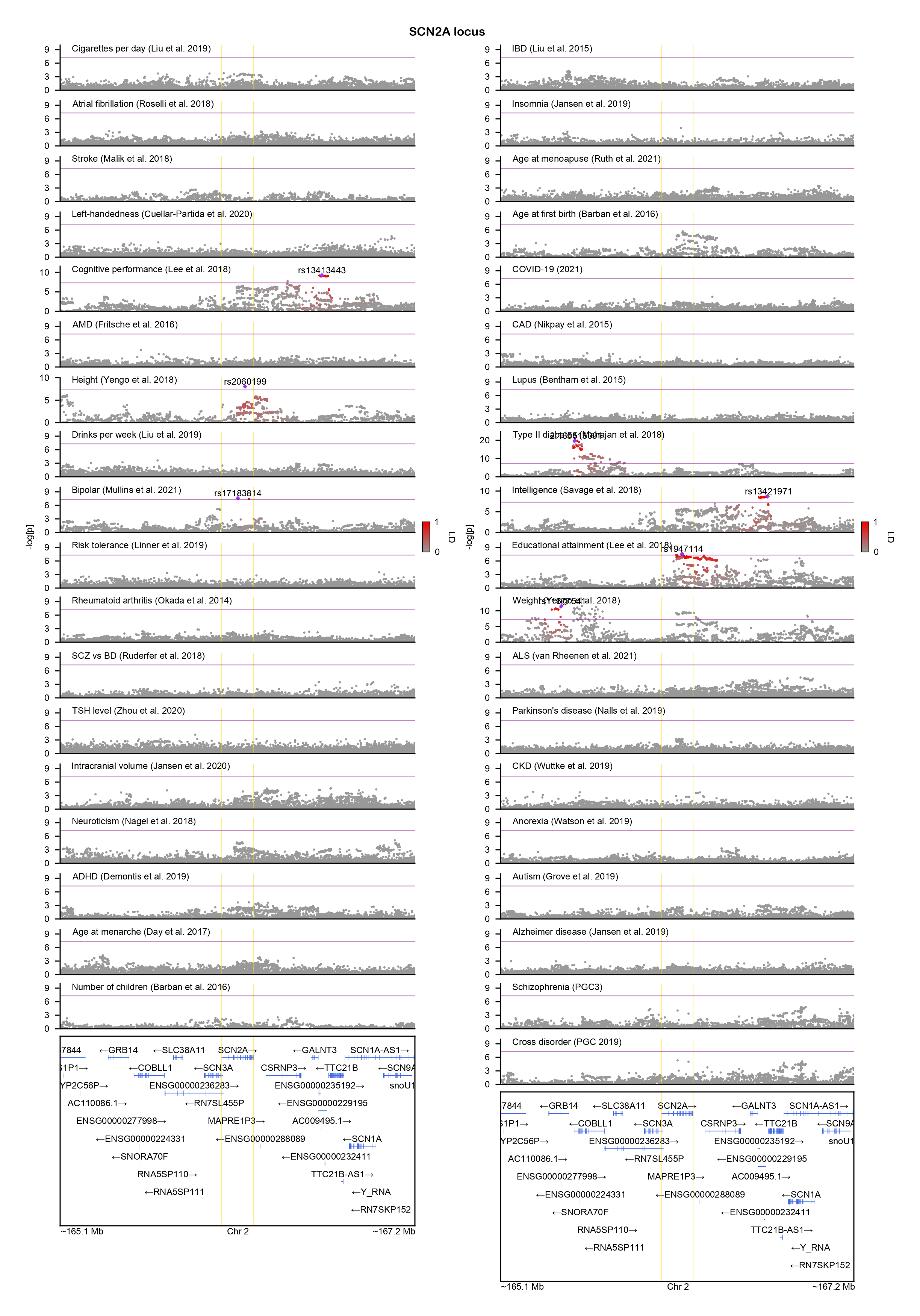

### SCN8A-locuszoom.png

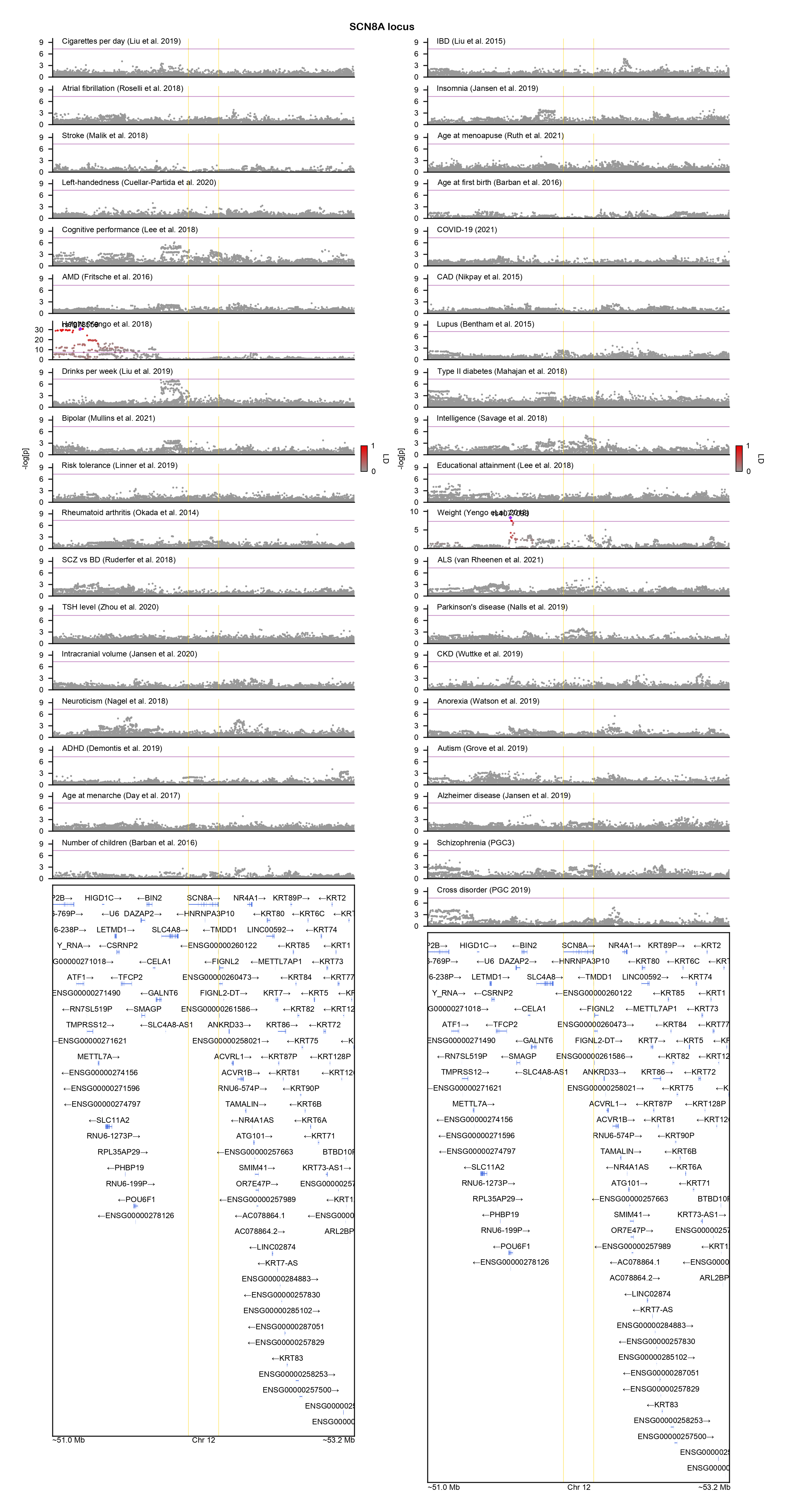

### SETBP1-locuszoom.png

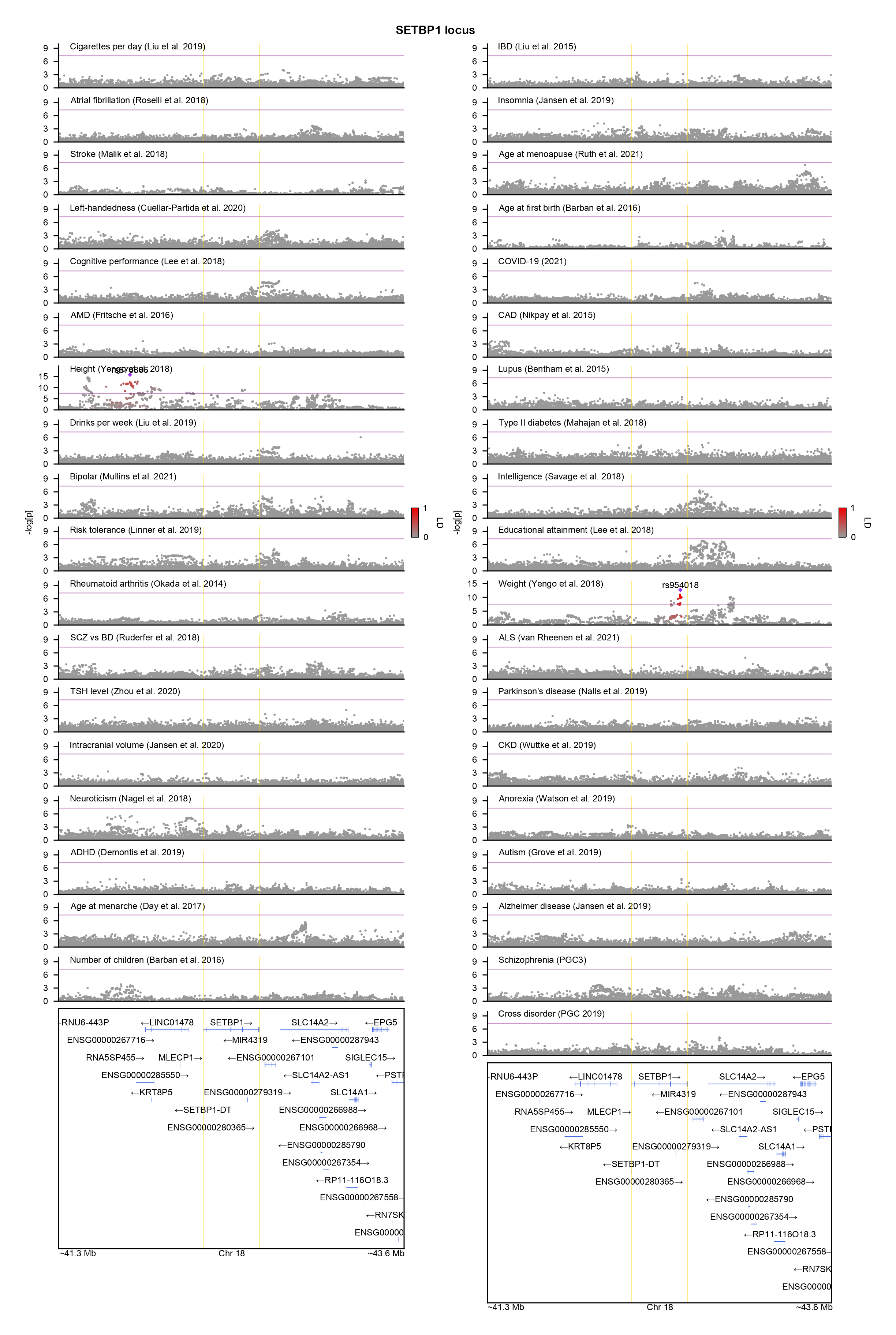

### SH3RF1-locuszoom.png

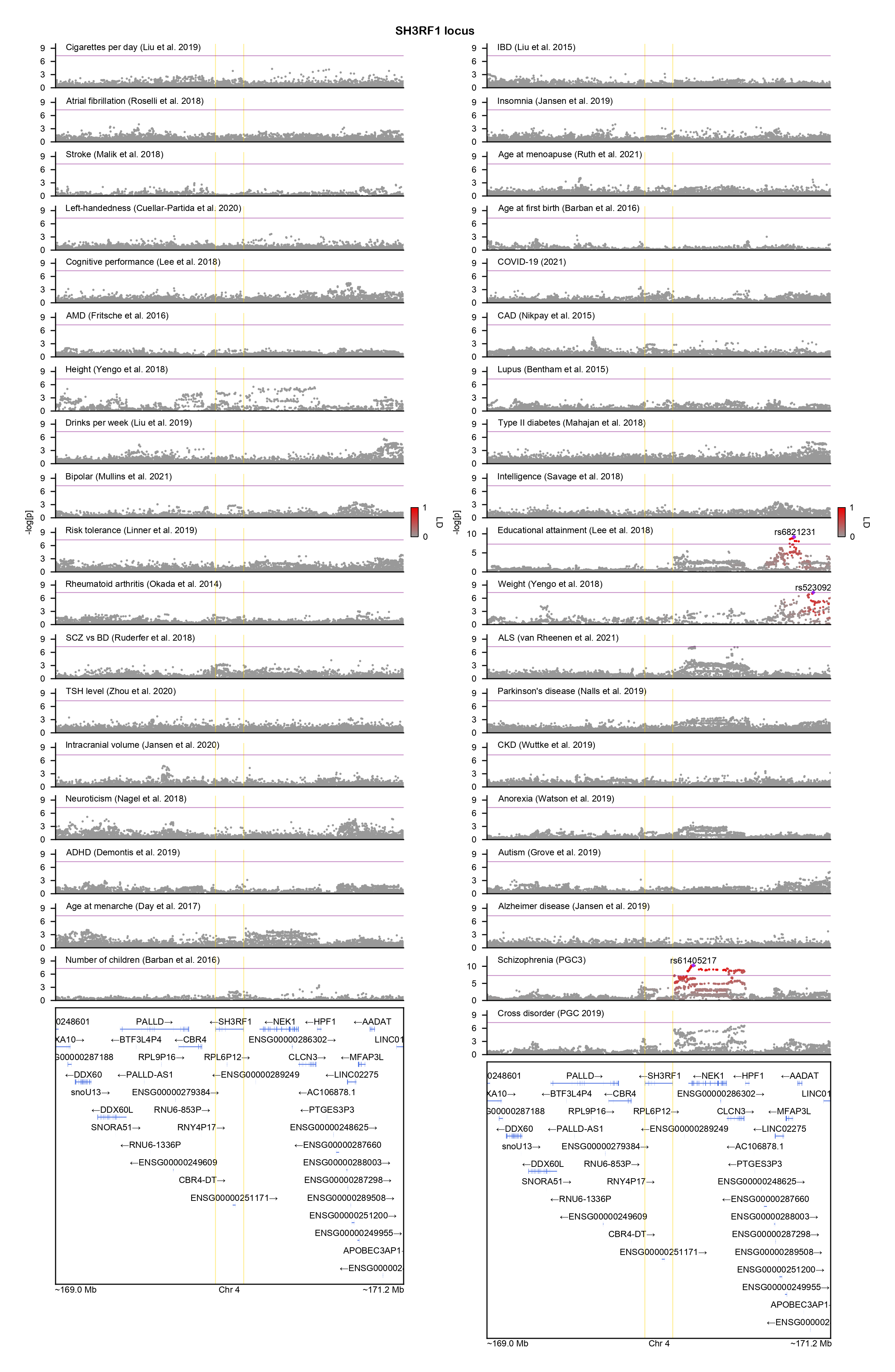

### SHOC2-locuszoom.png

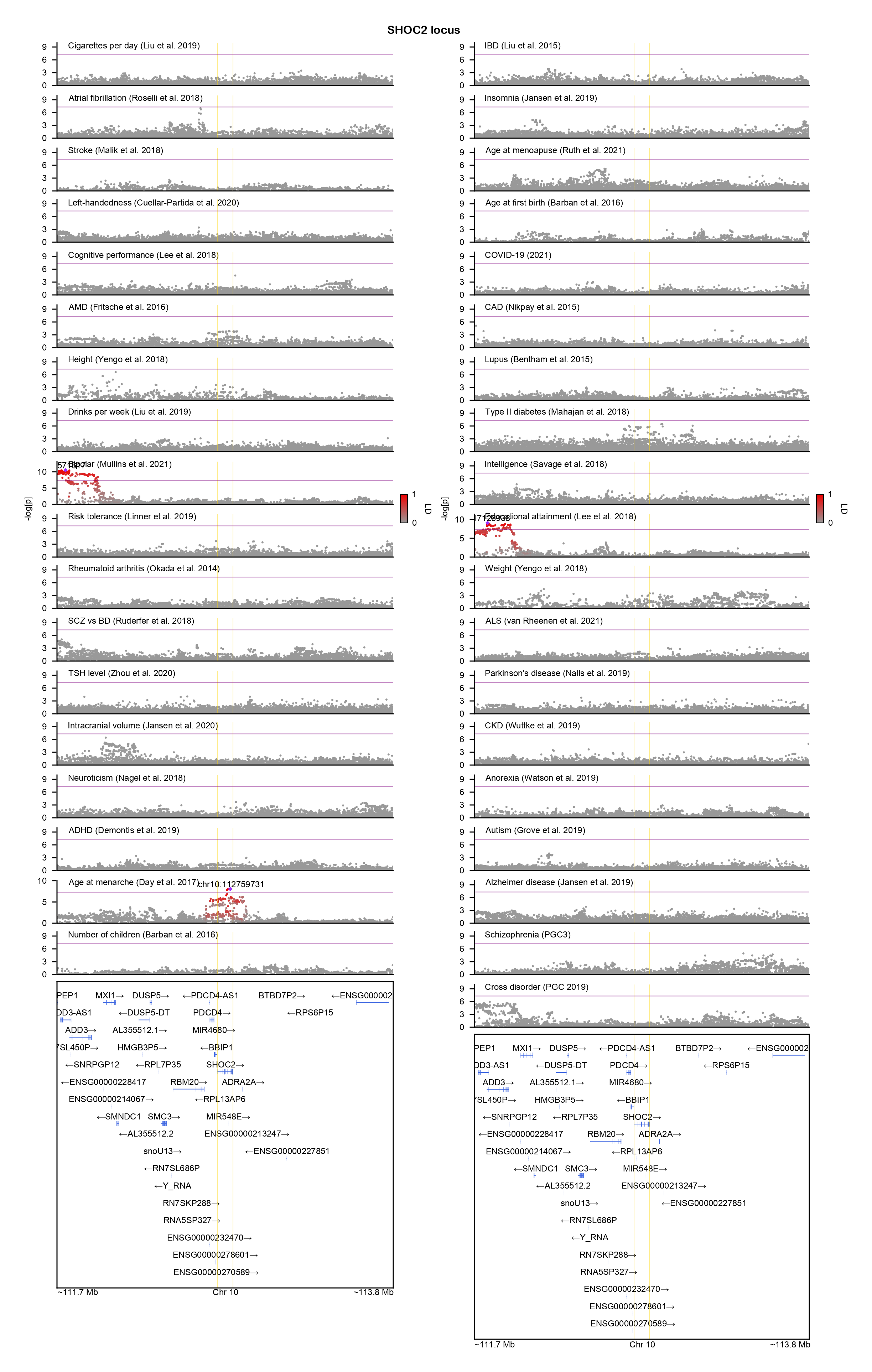

### SIN3A-locuszoom.png

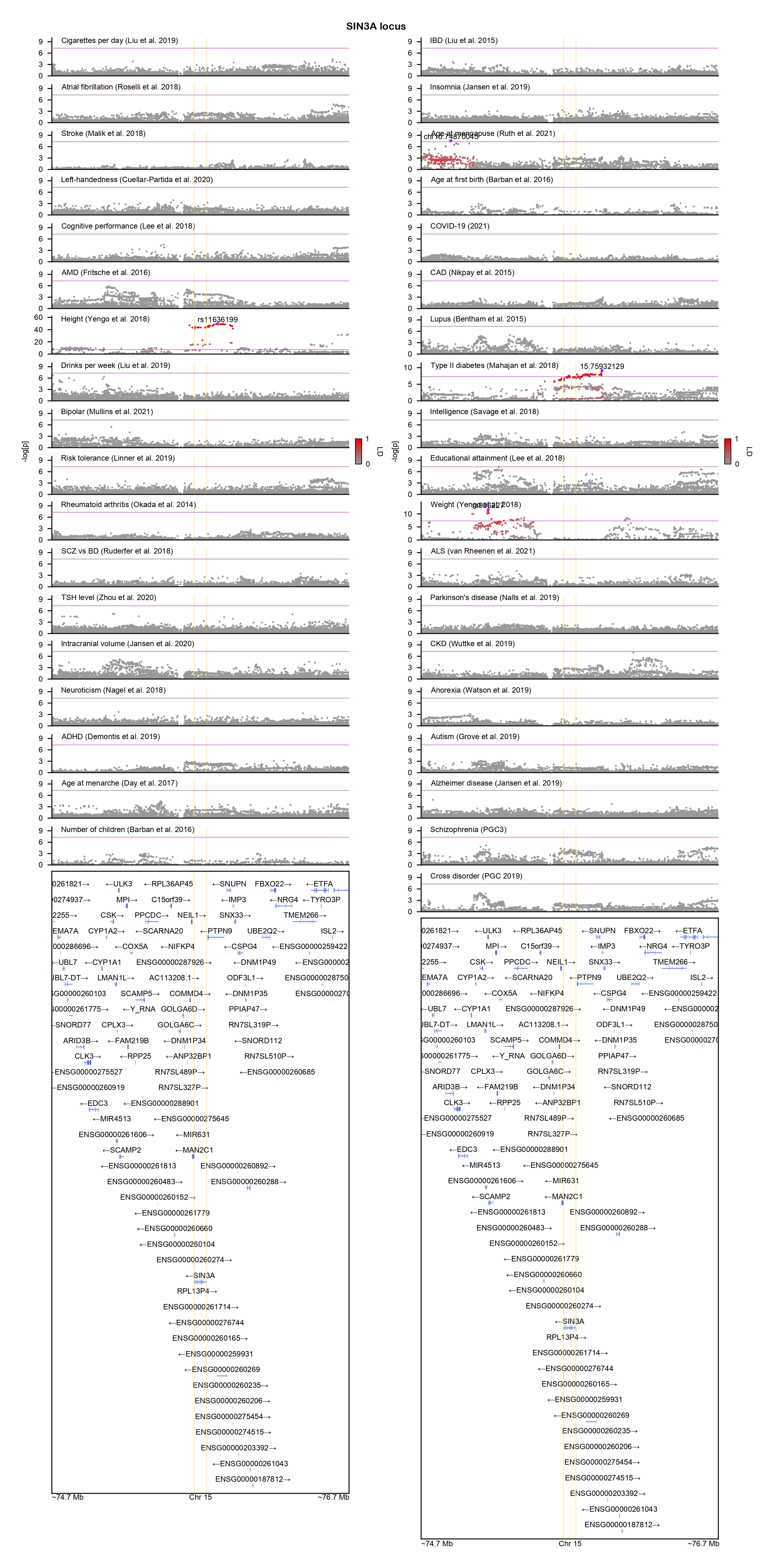

### SMARCA2-locuszoom.png

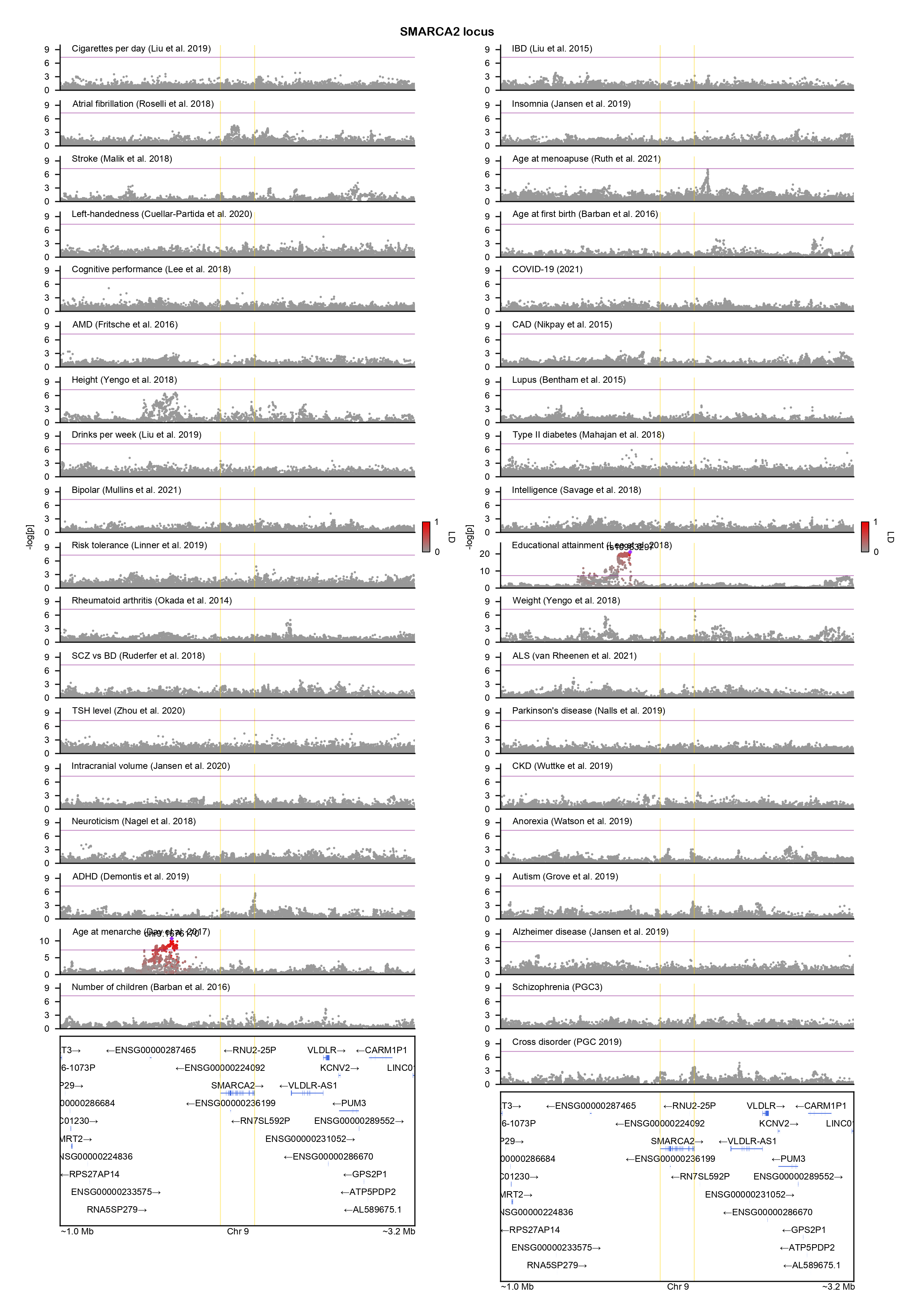

### SON-locuszoom.png

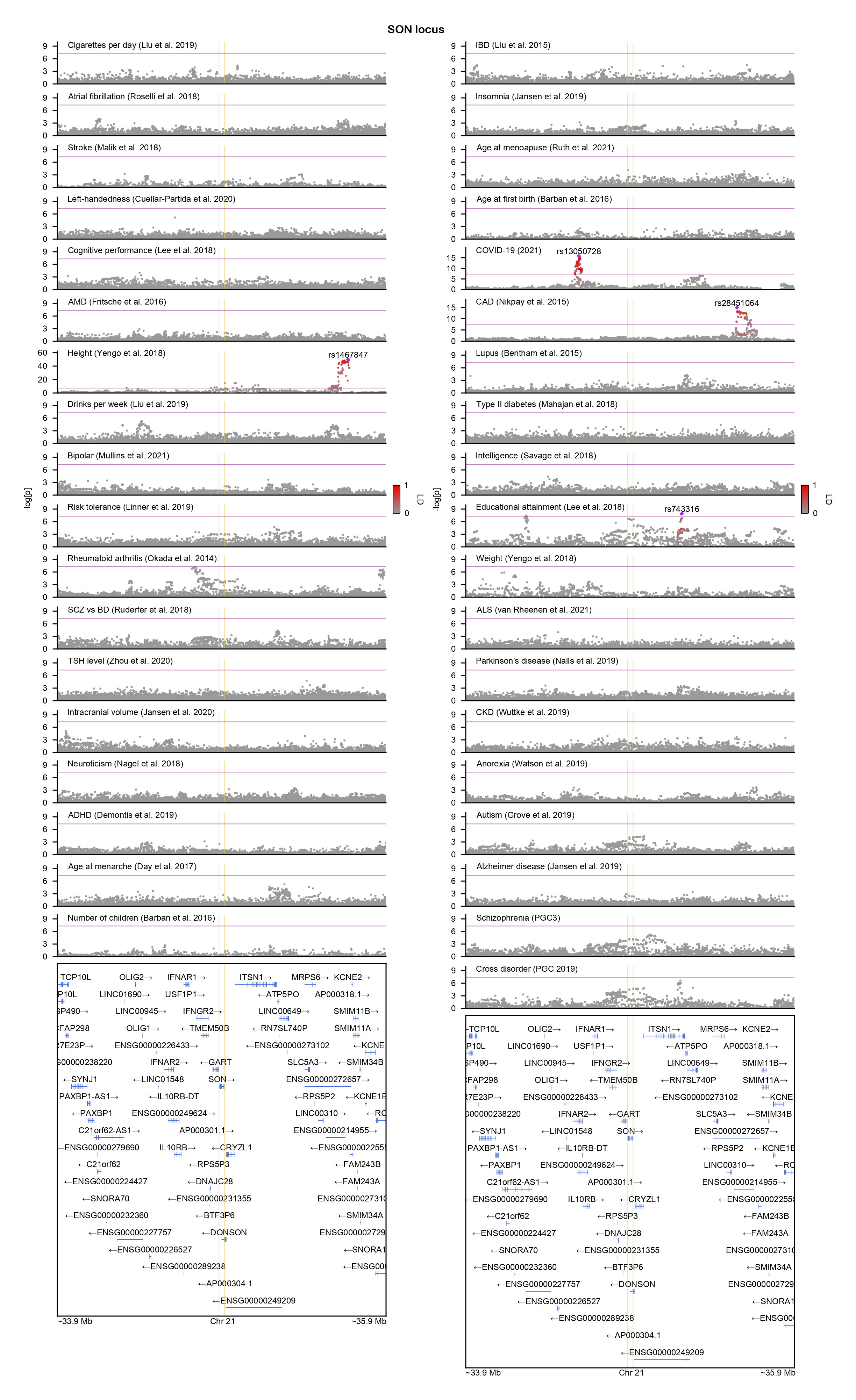

### SOX10-locuszoom.png

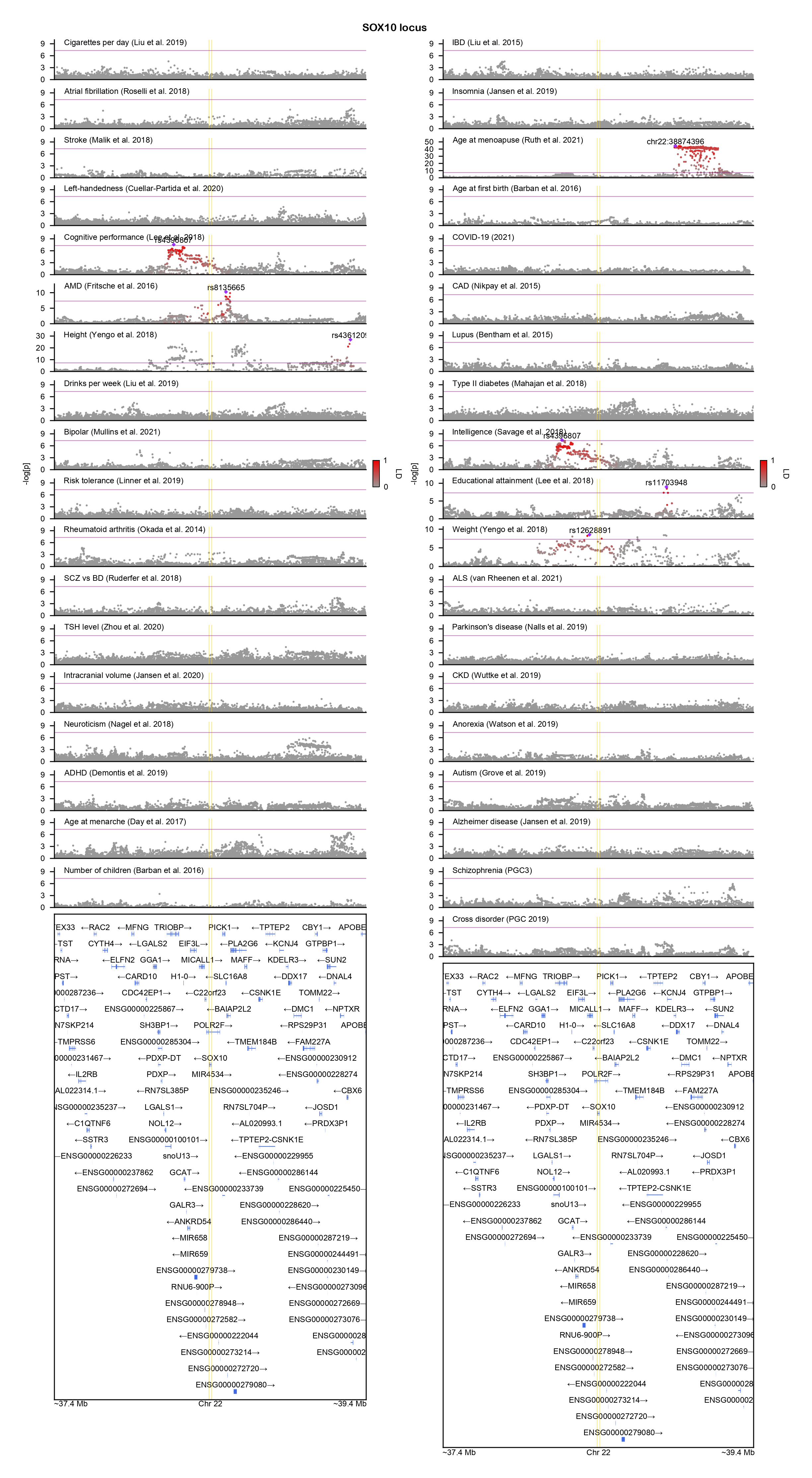

### SRCAP-locuszoom.png

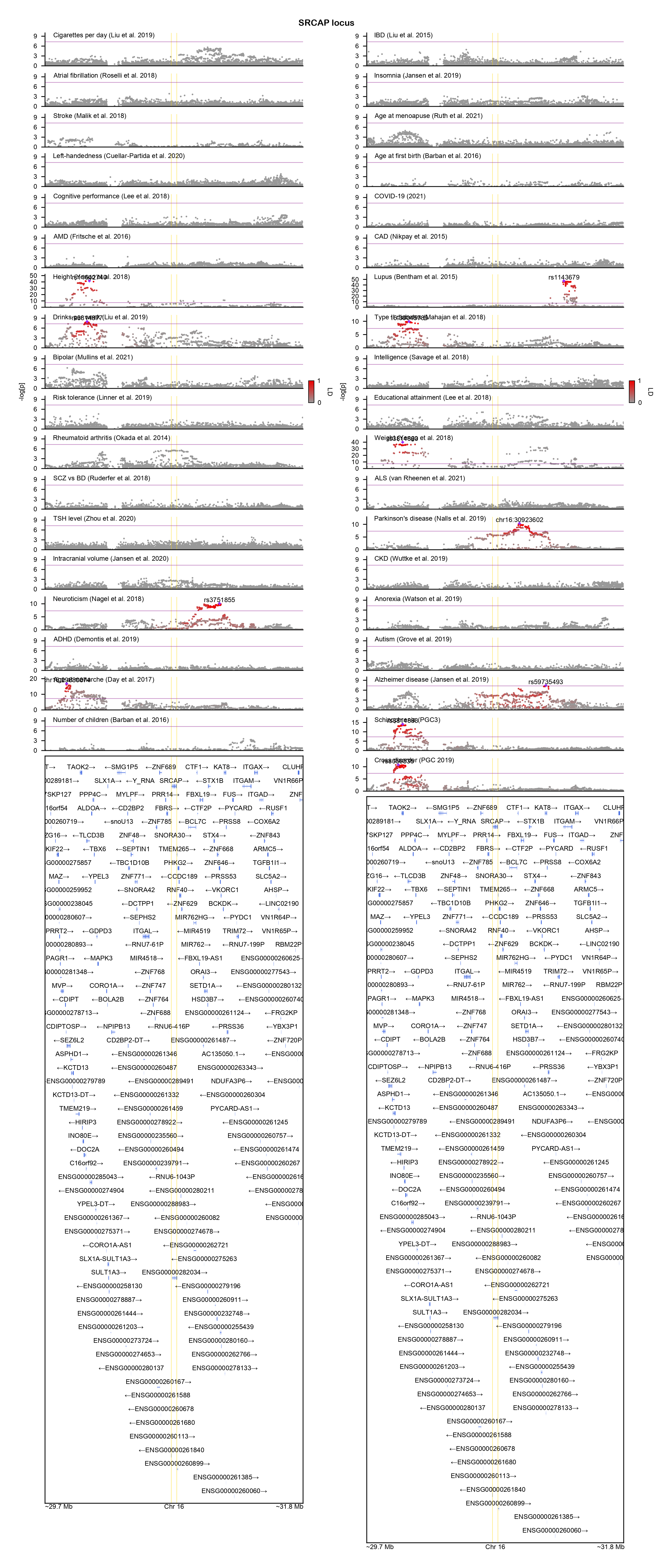

### SRRM2-locuszoom.png

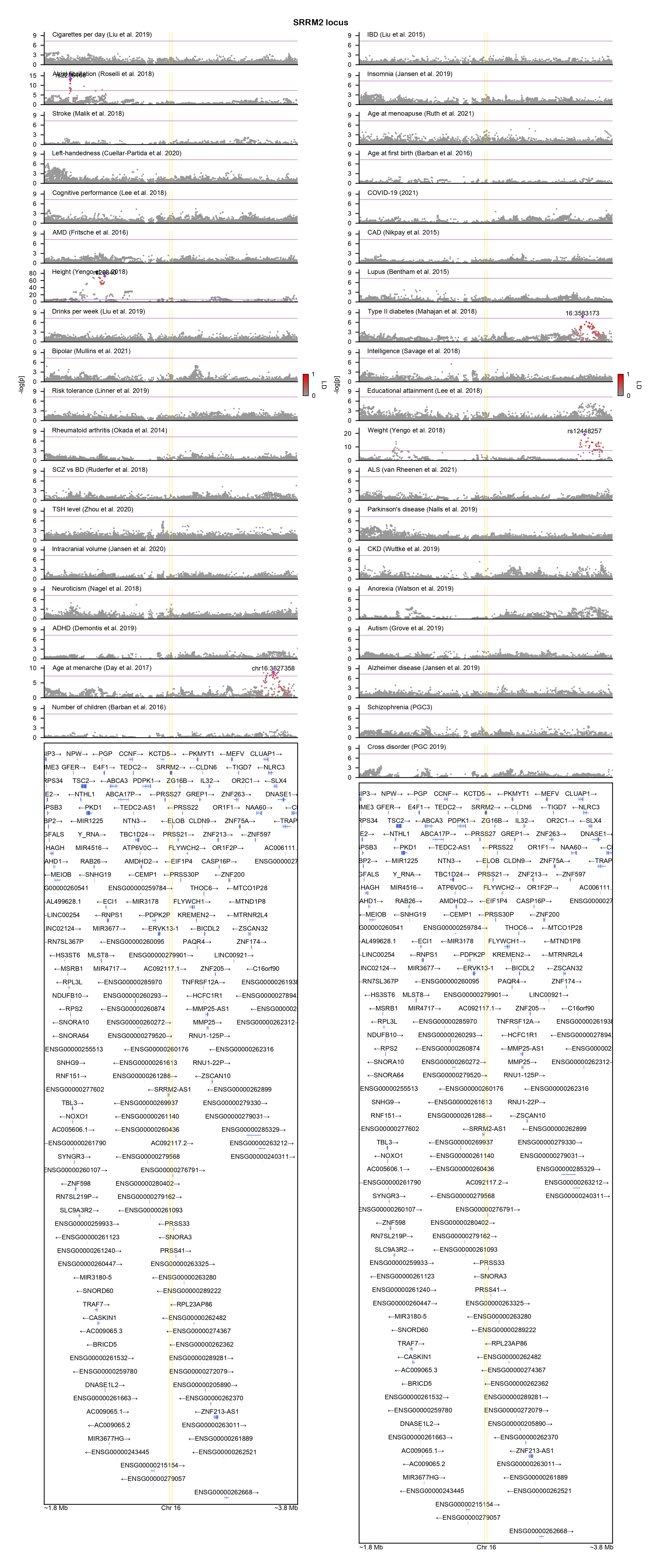

### SYT1-locuszoom.png

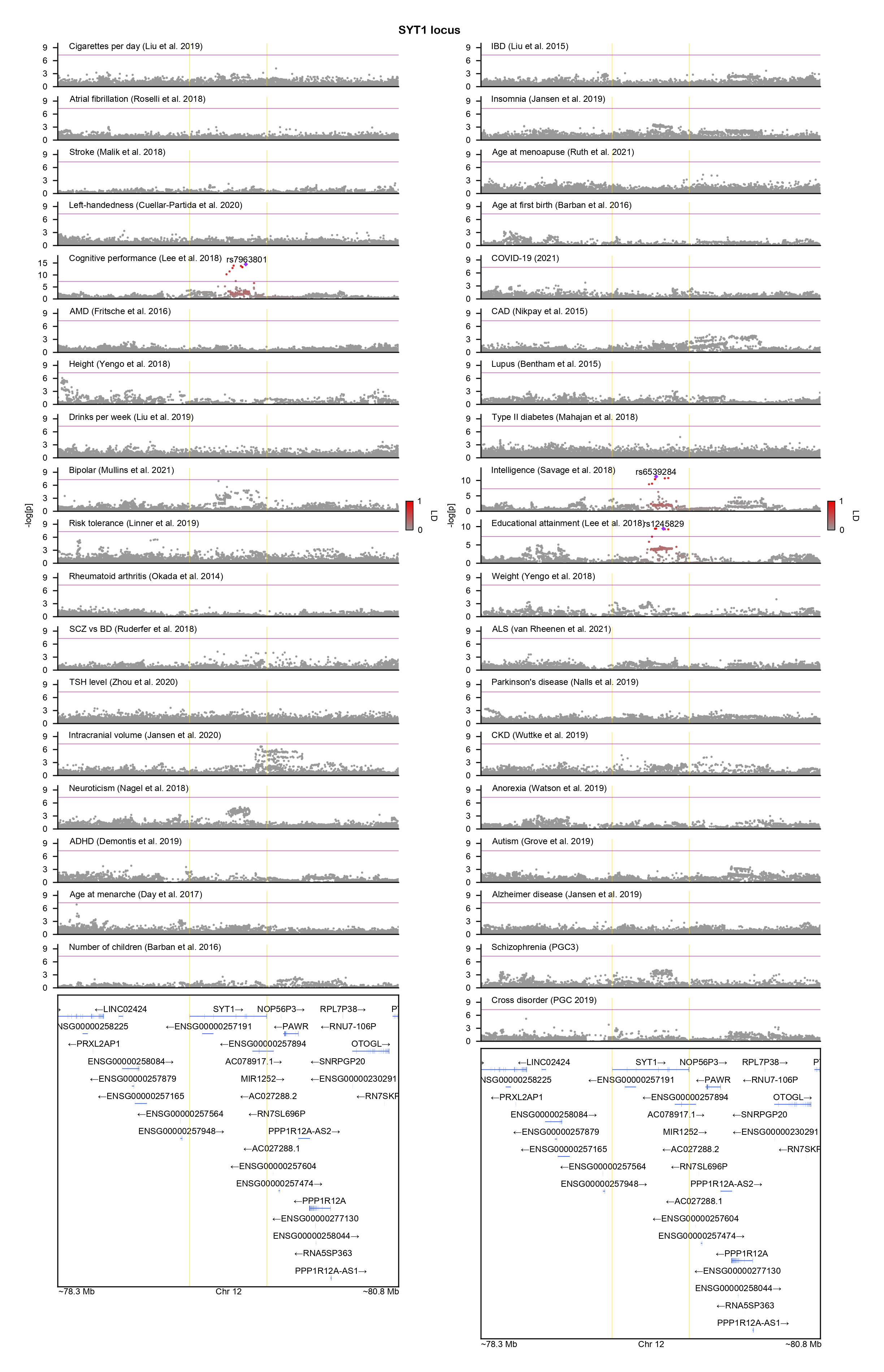

### TLK2-locuszoom.png

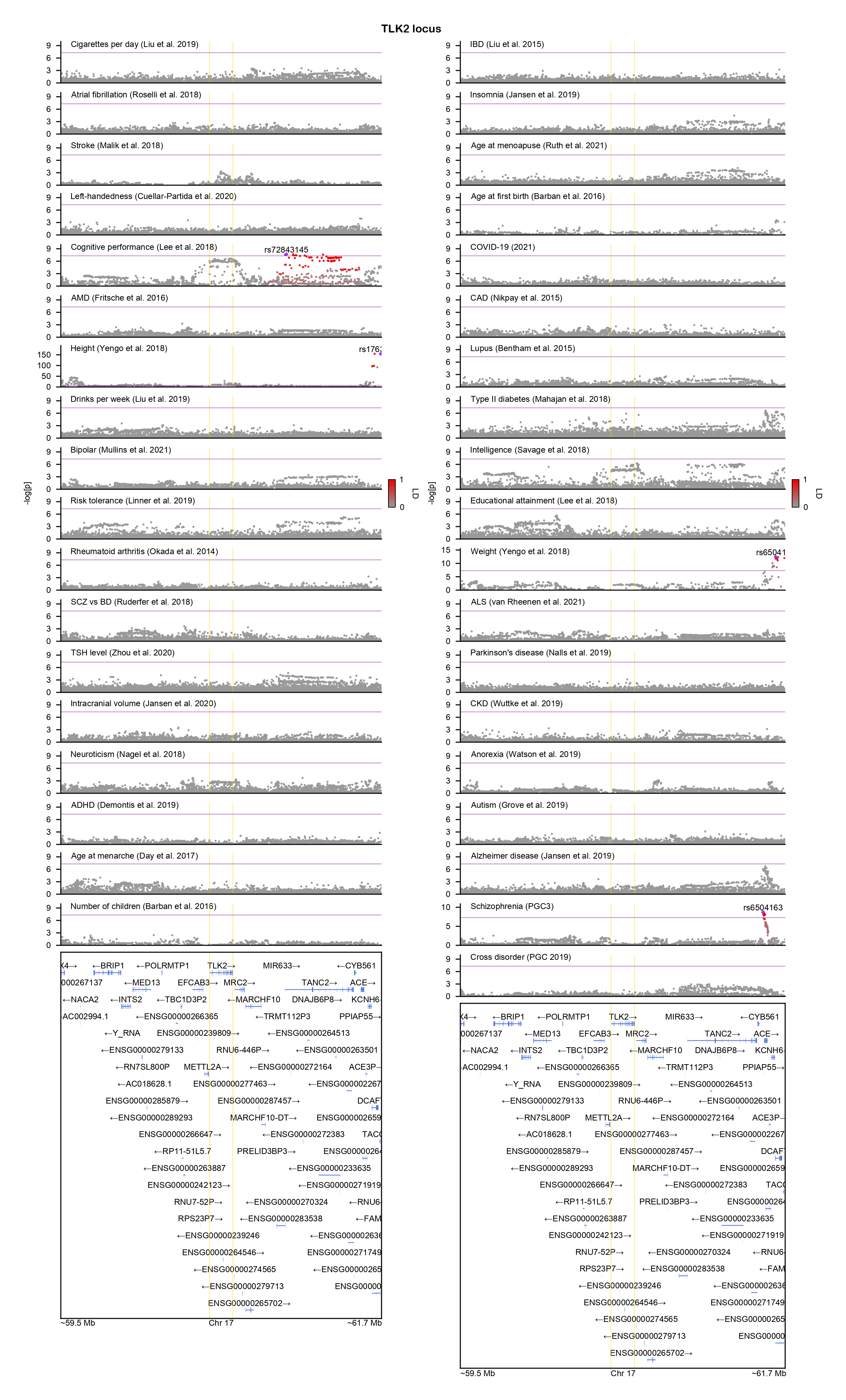

### TMX2-locuszoom.png

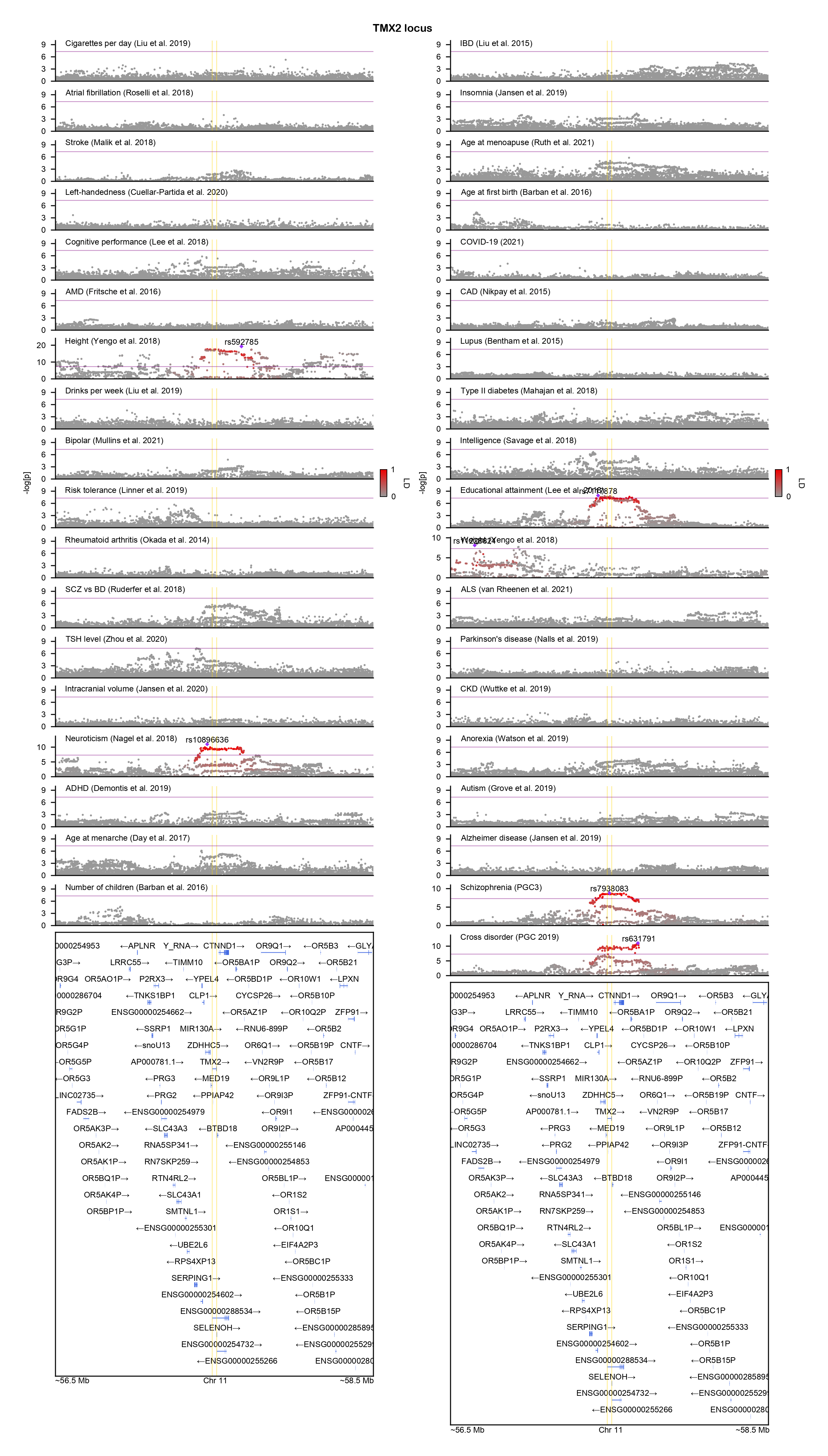

### TRPM3-locuszoom.png

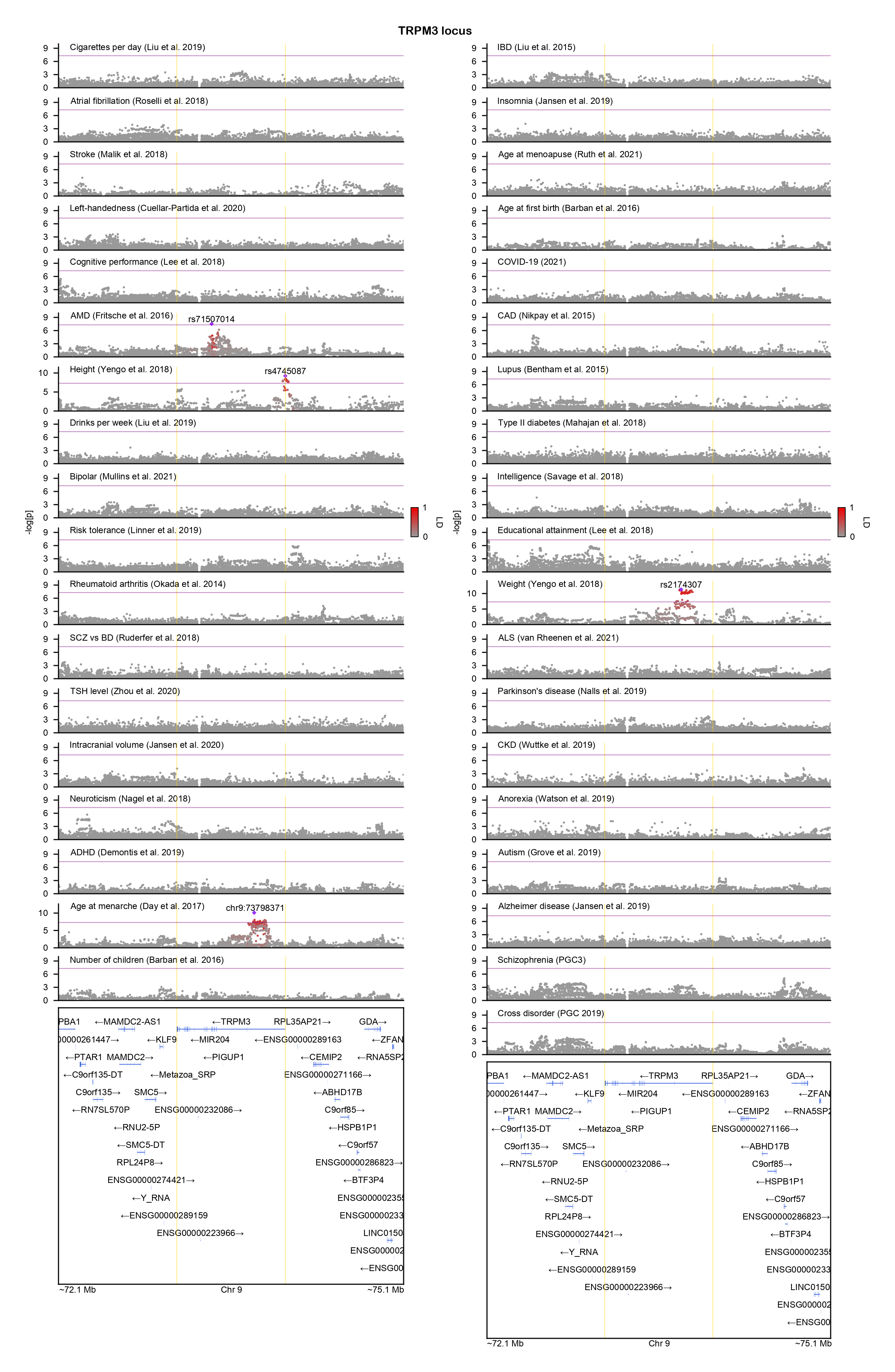

### TRPM6-locuszoom.png

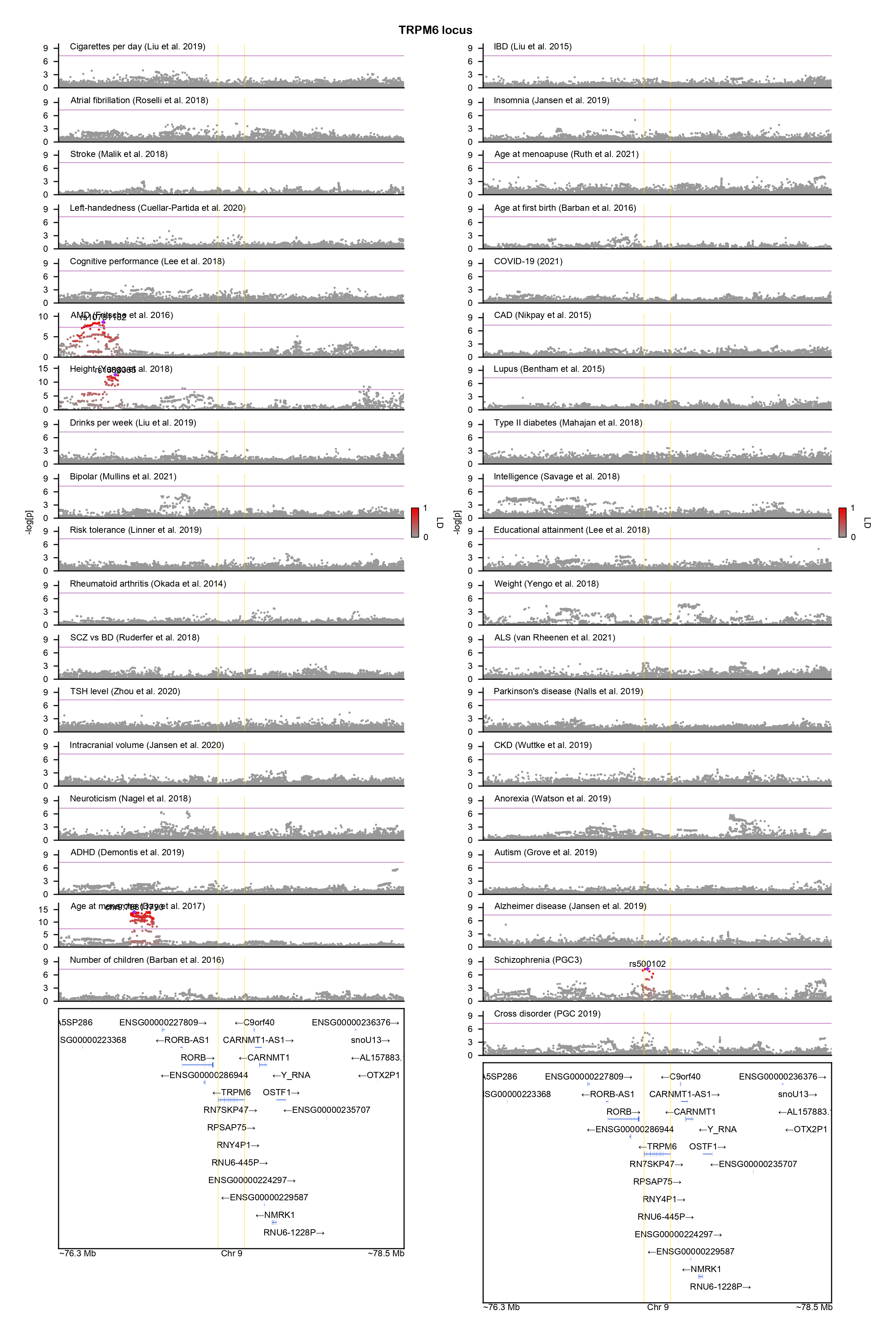

### UPF1-locuszoom.png

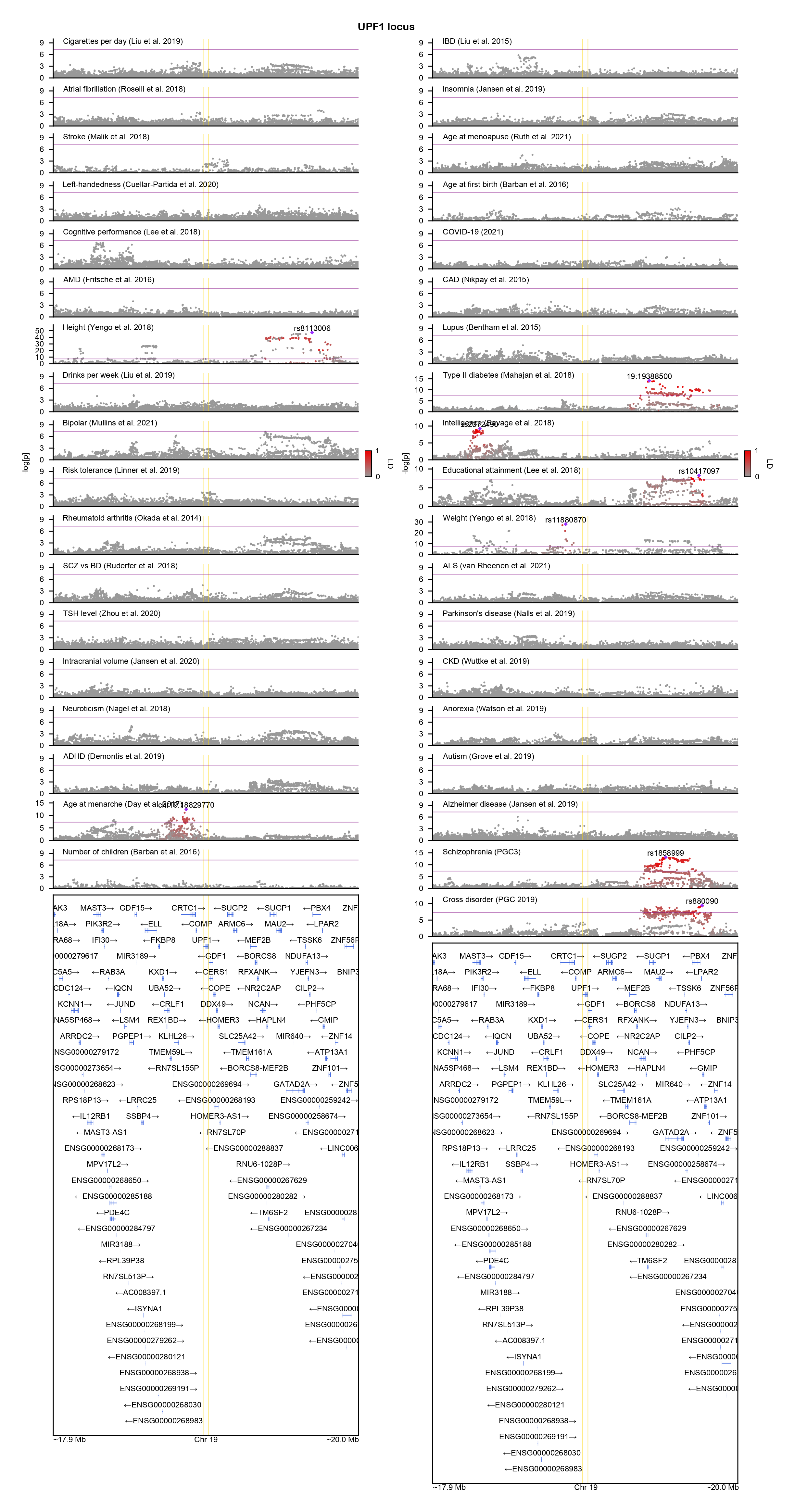

### WDR45-locuszoom.png

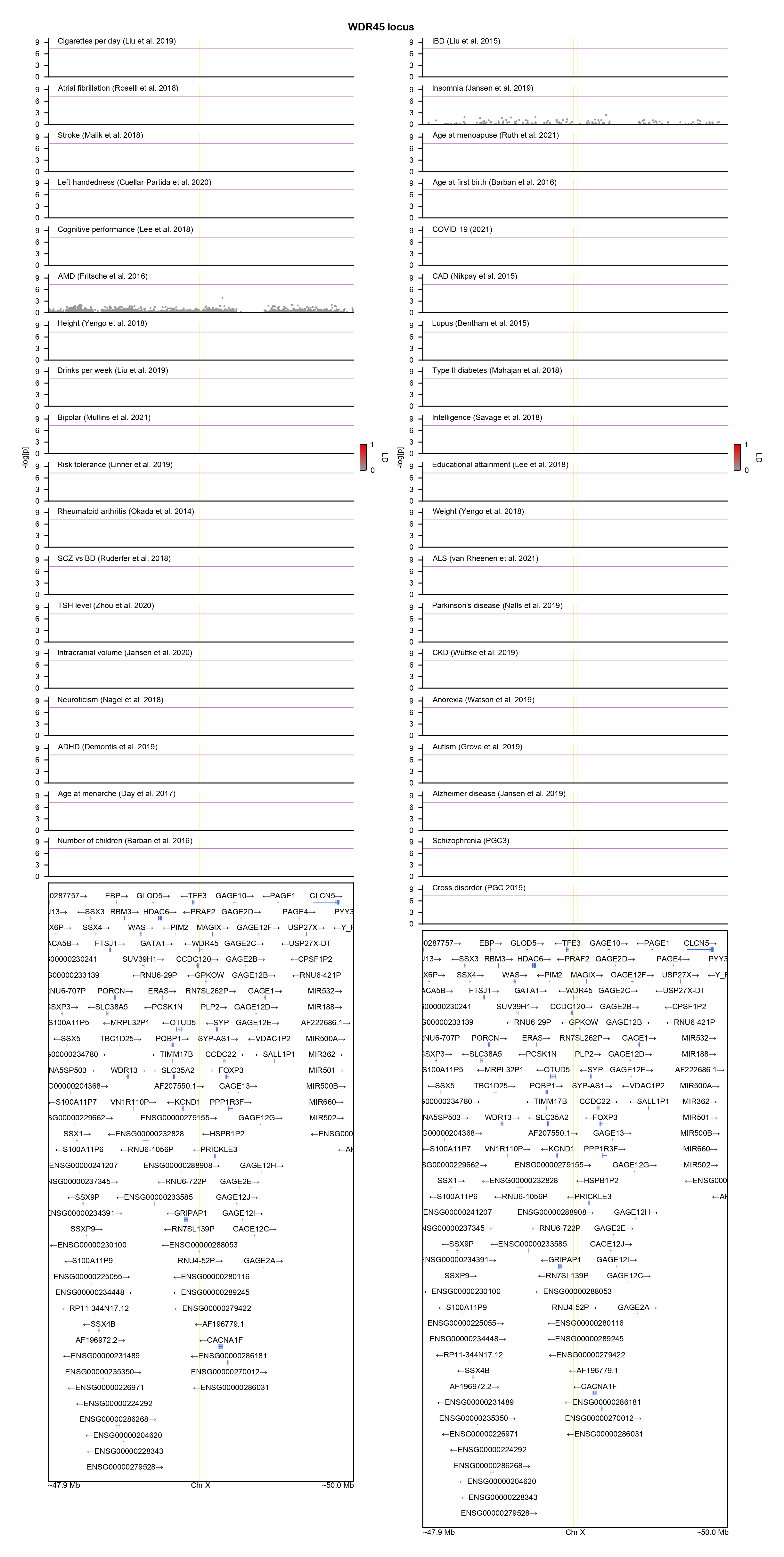

### XPO7-locuszoom.png

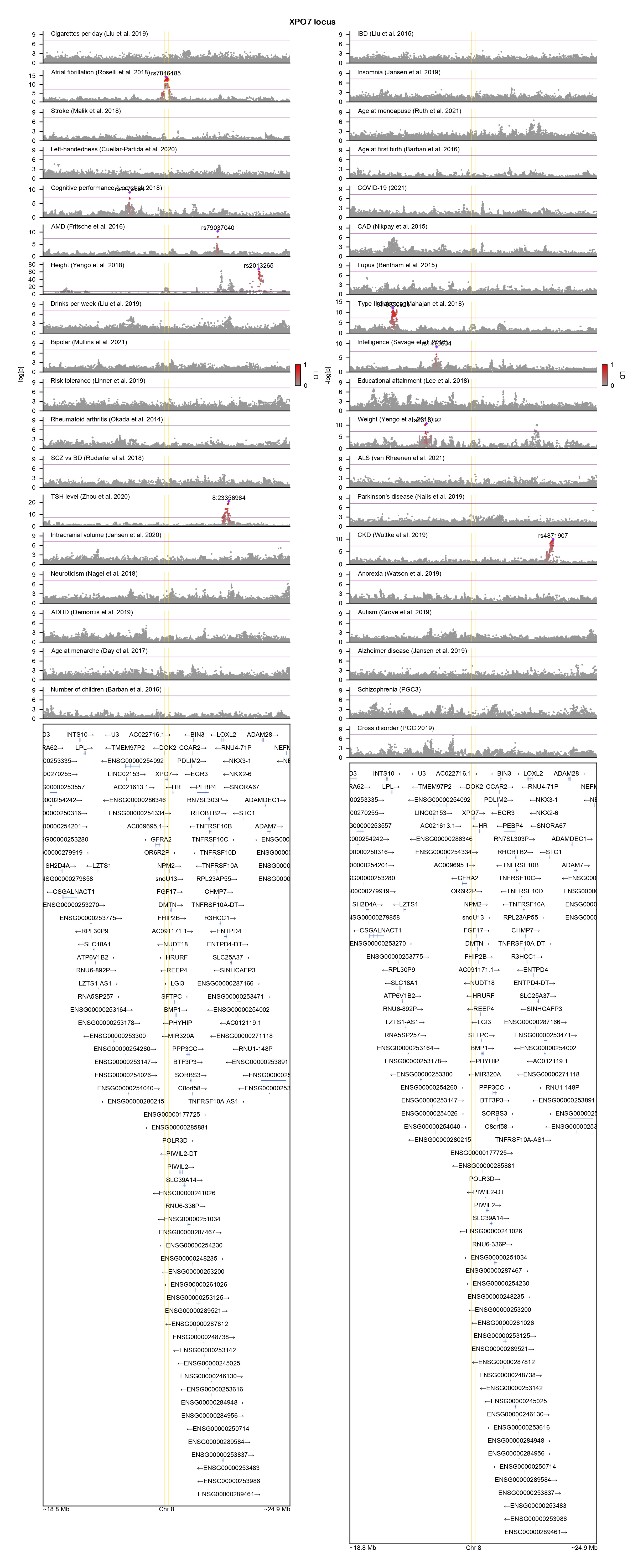

### YY1-locuszoom.png

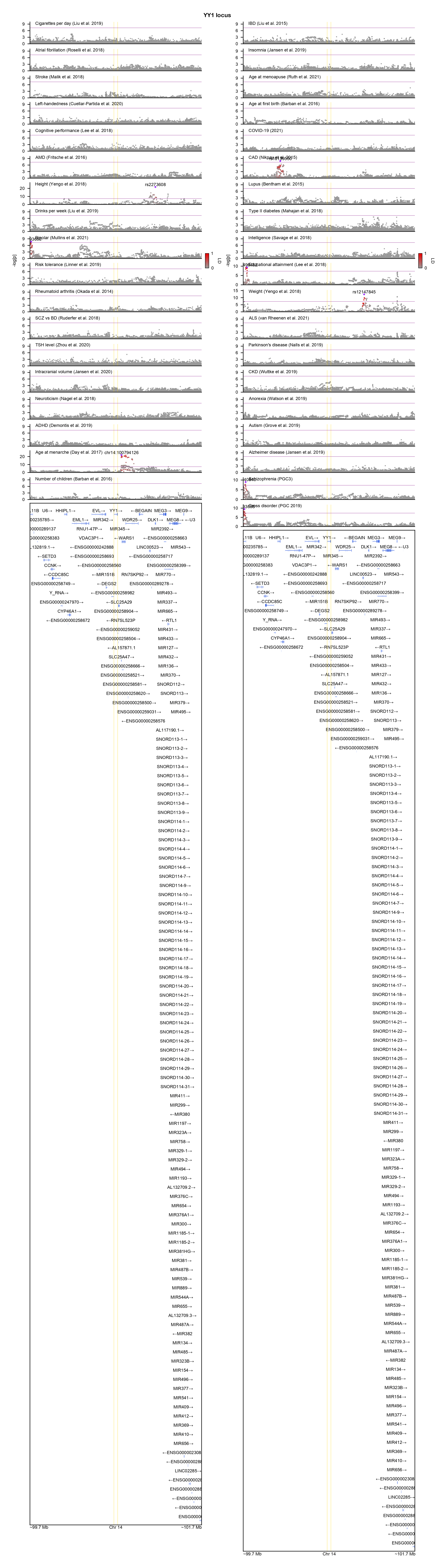

### ZNF804A-locuszoom.png

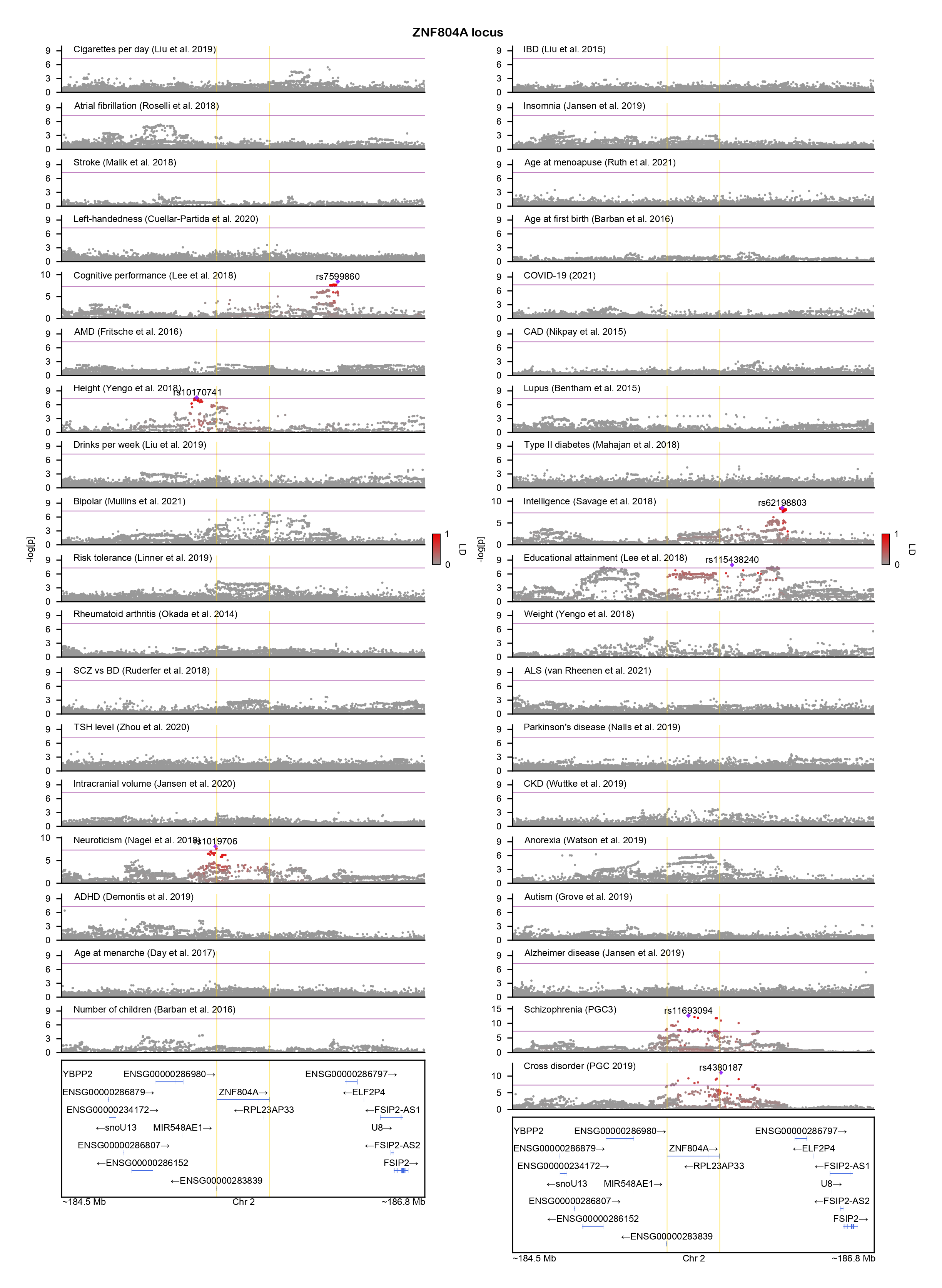

### ZNF835-locuszoom.png

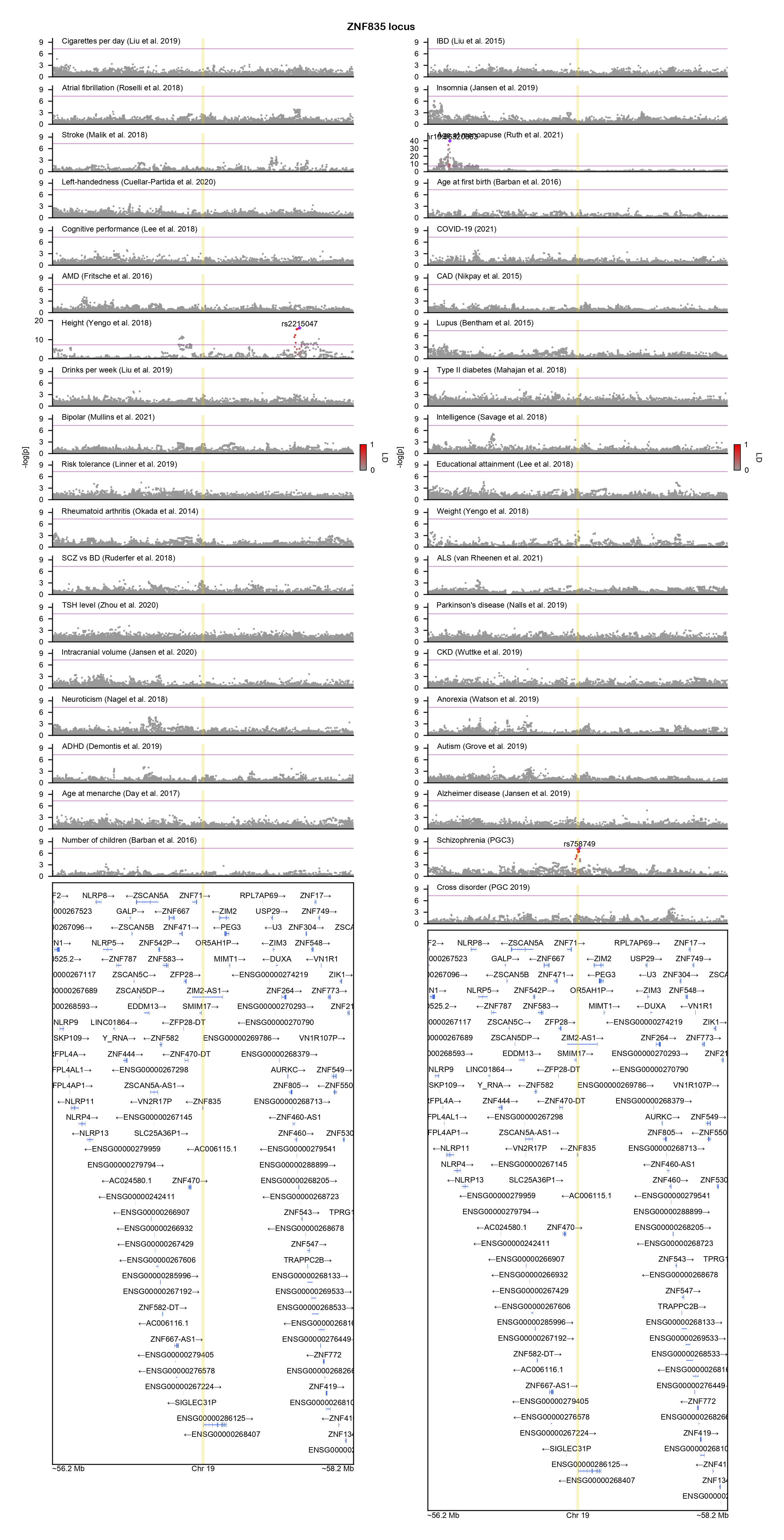
