## Supplementary figures and images for "GeneticsMakie.jl: A versatile and scalable toolkit for visualizing locus-level genetic and genomic data"

### AC106017.1-locuszoom.png

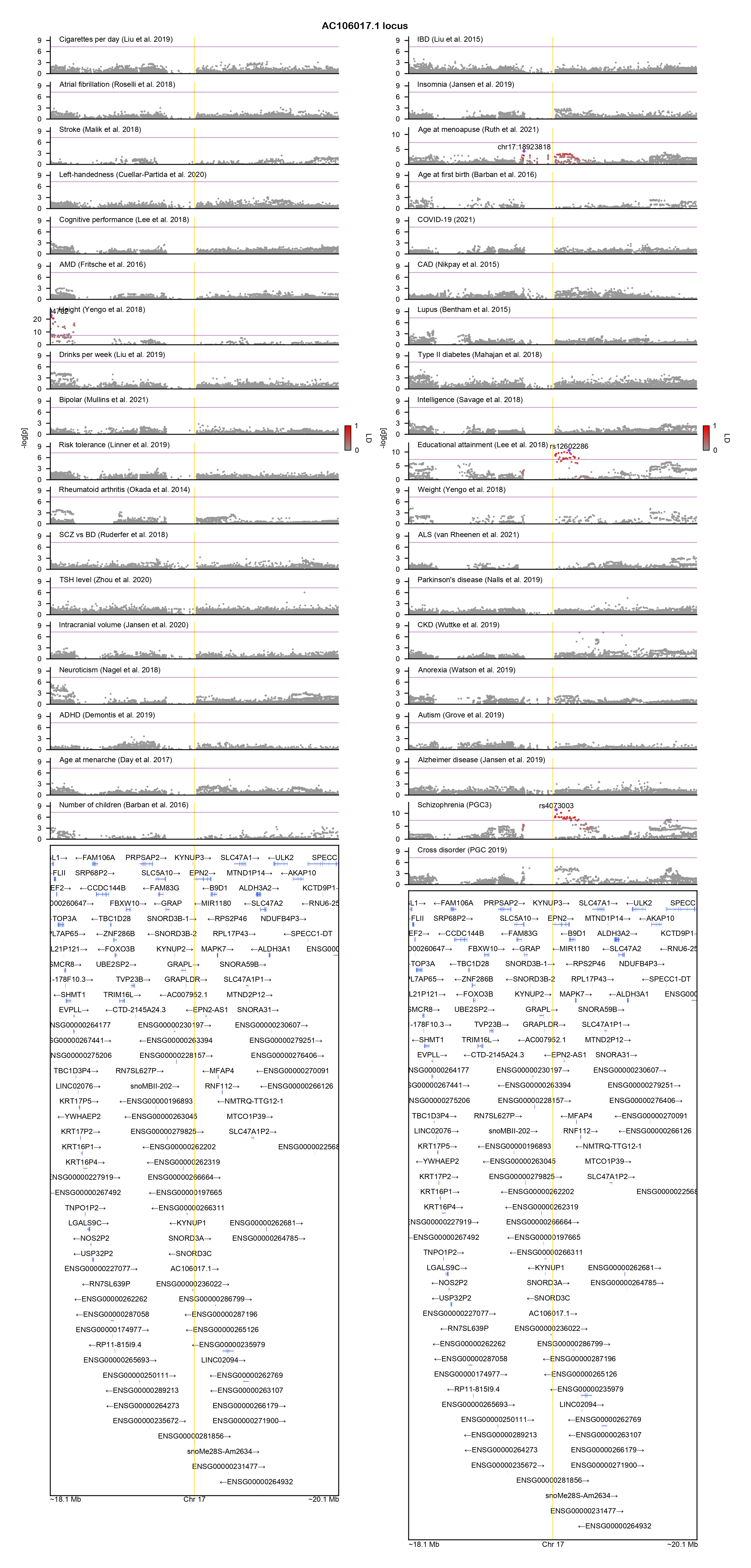

### ADCY2-locuszoom.png

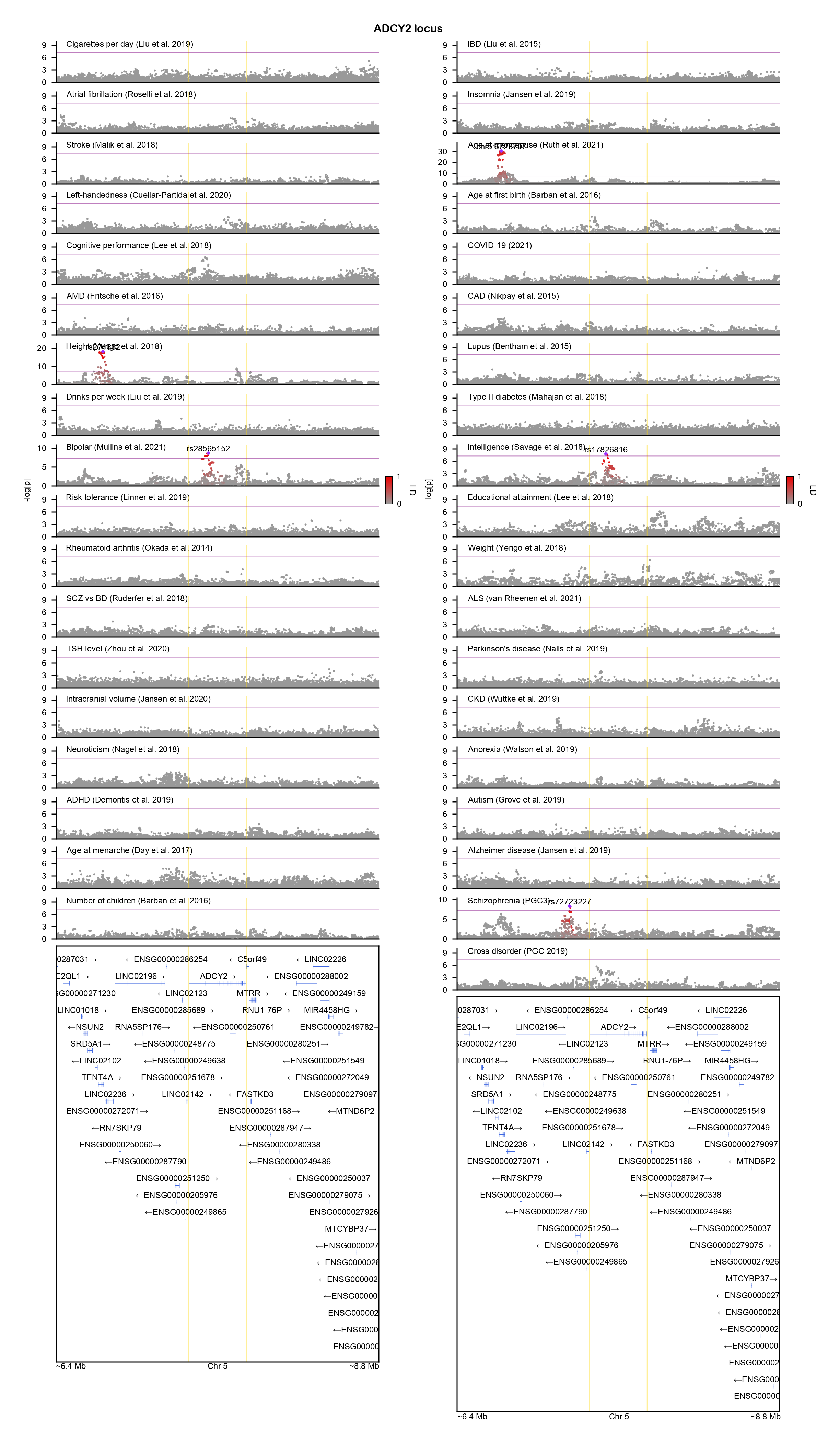

### ADO-locuszoom.png

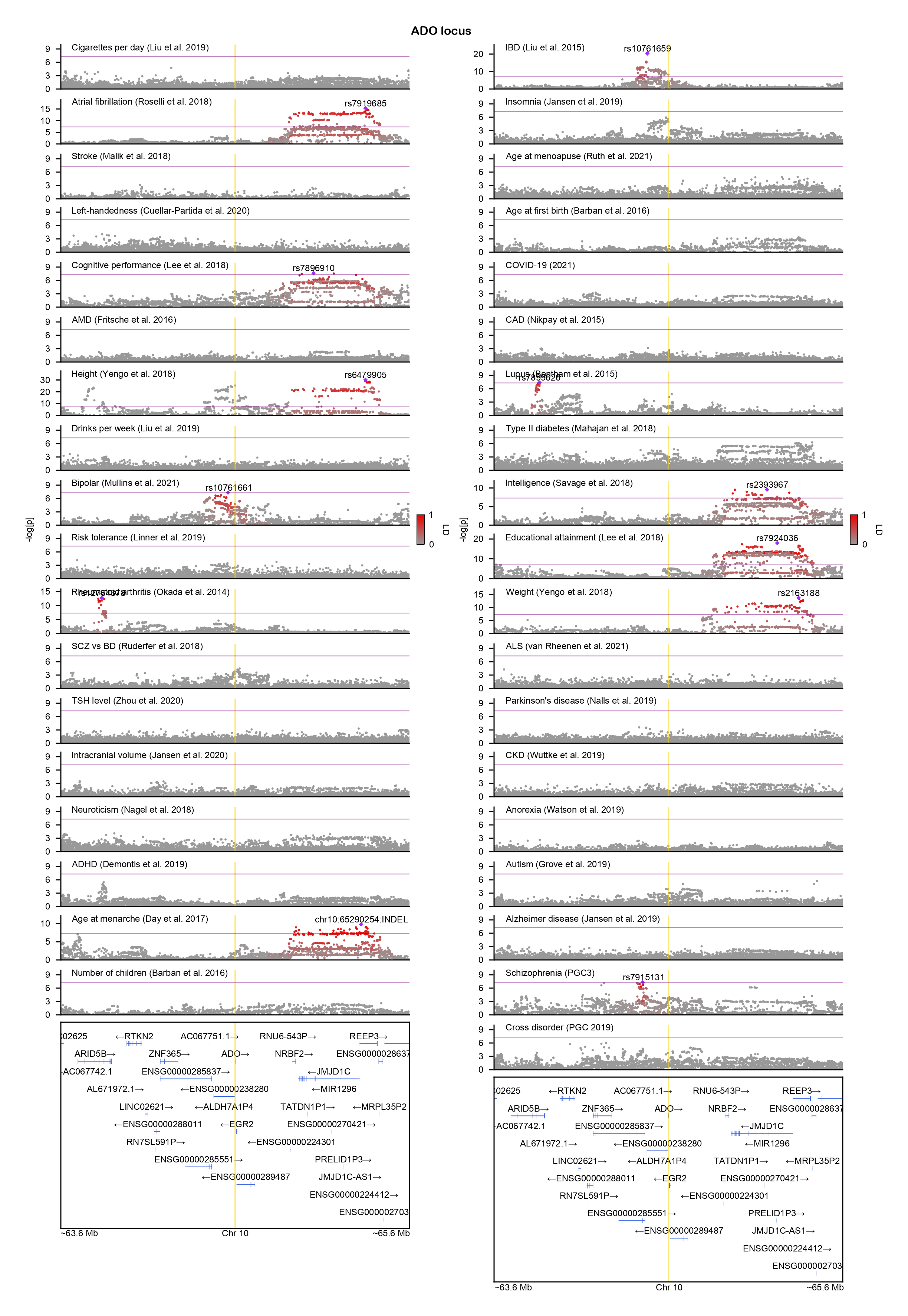

### AKT3-locuszoom.png

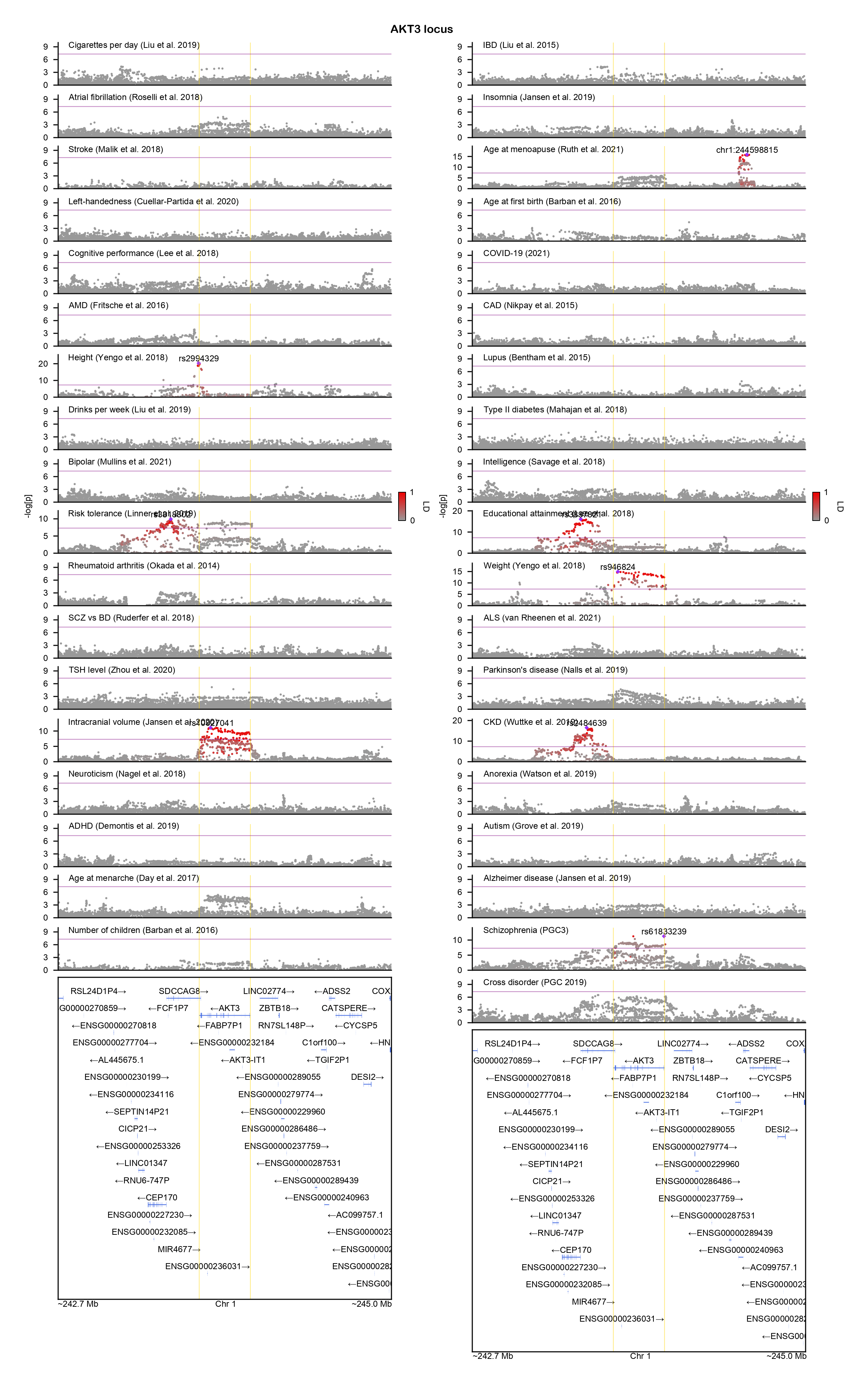

### AP3B2-locuszoom.png

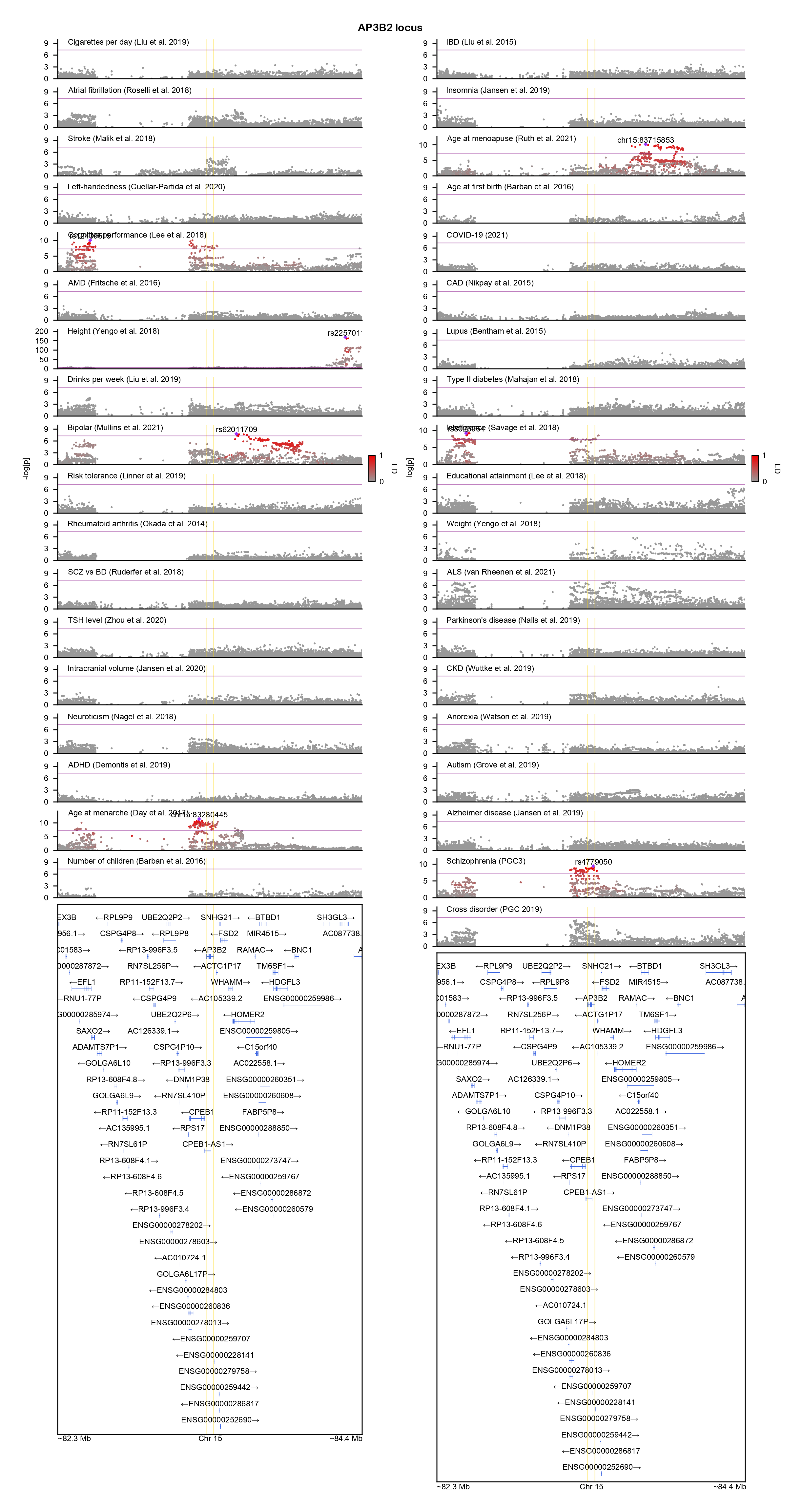

### ARRDC3-locuszoom.png

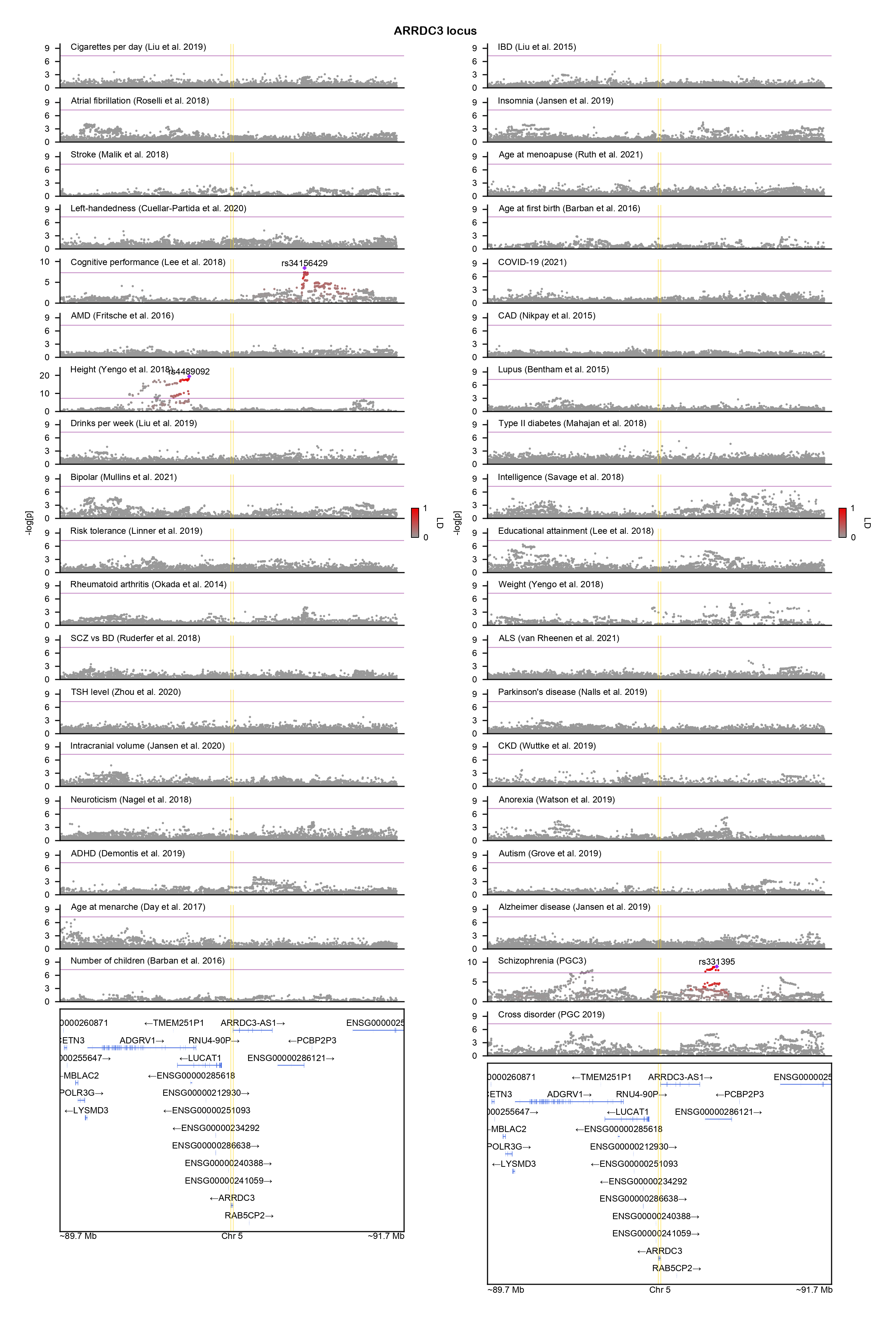

### ASIC1-locuszoom.png

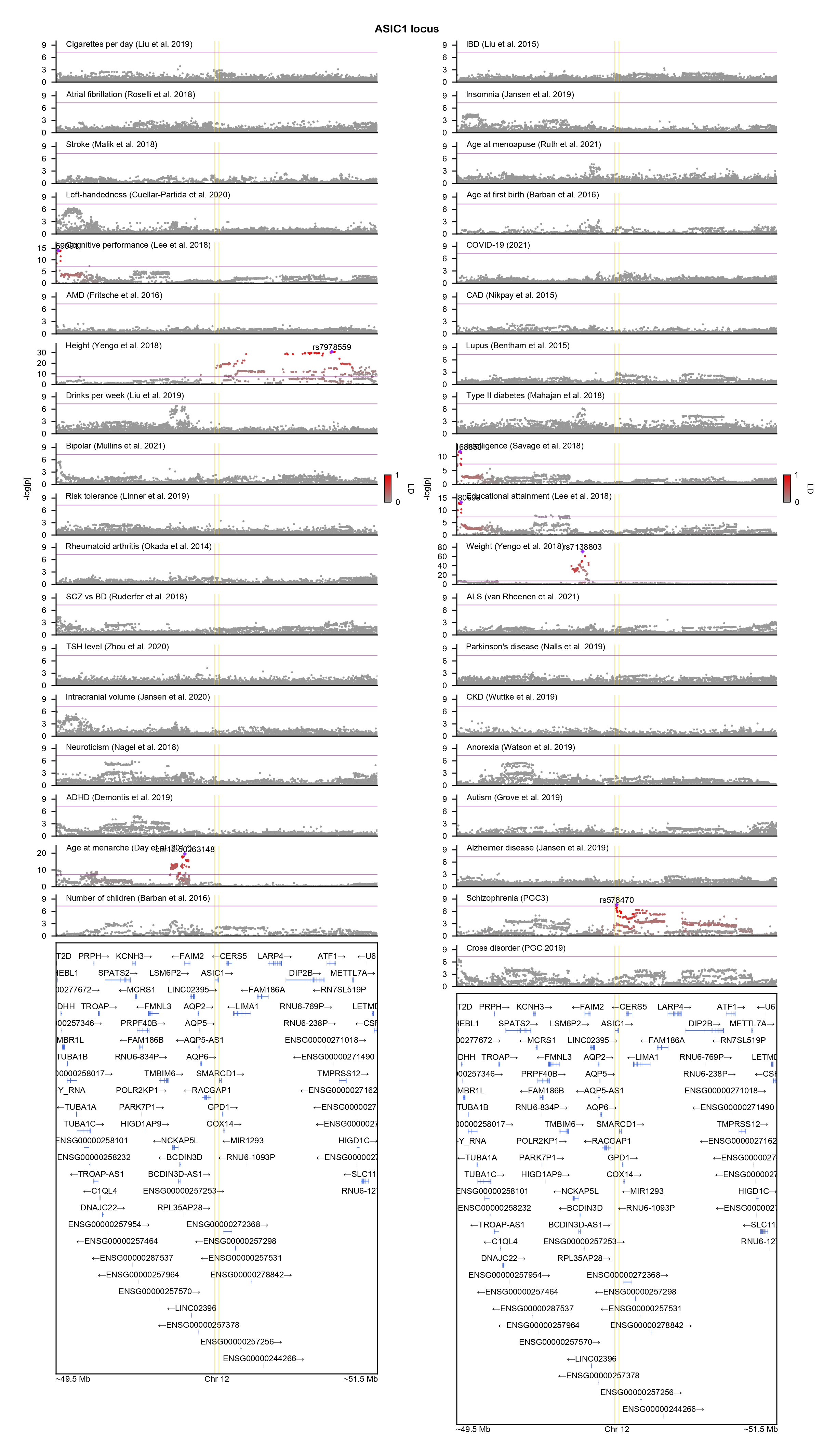

### ATF7IP-locuszoom.png

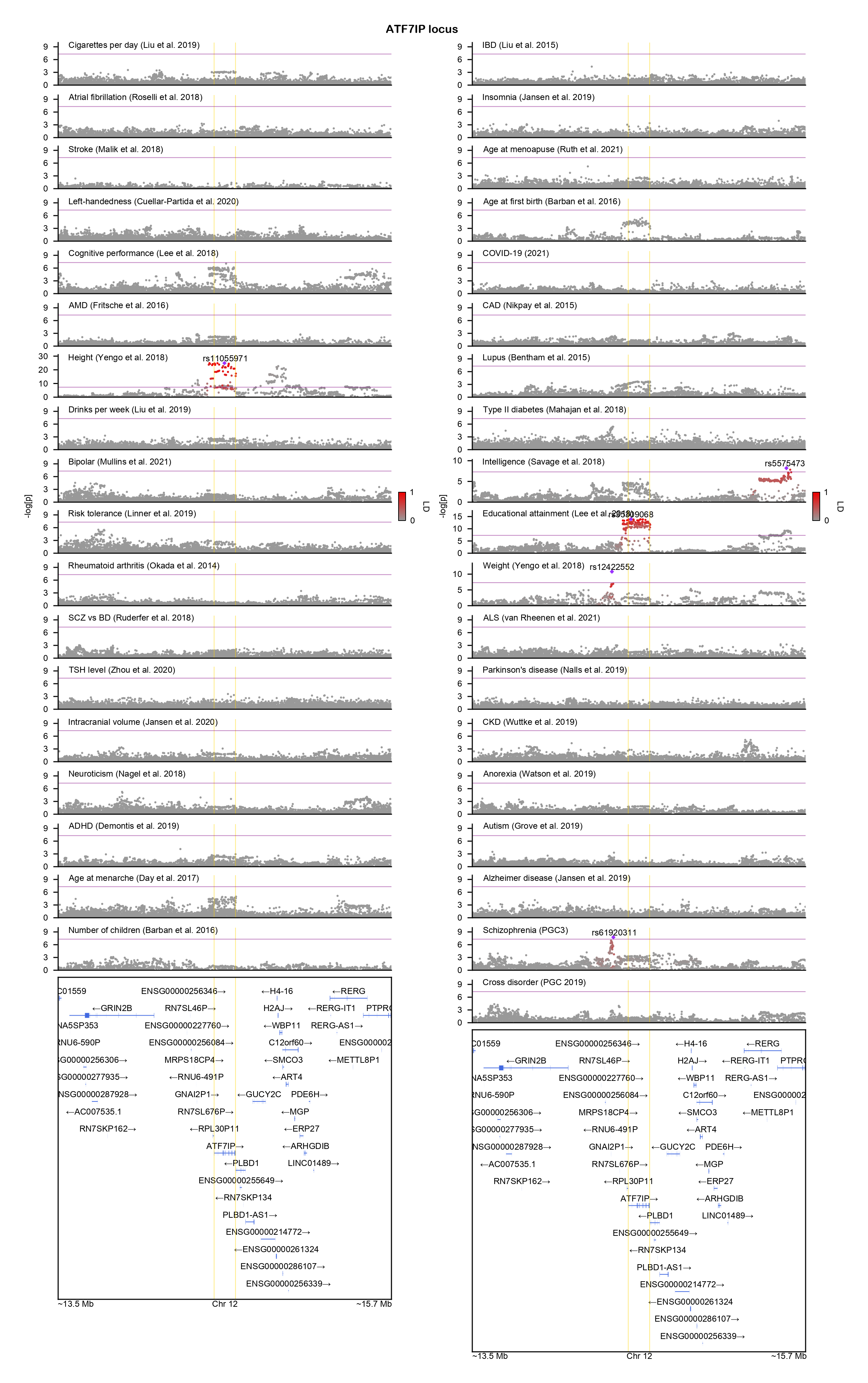

### ATP2A2-locuszoom.png

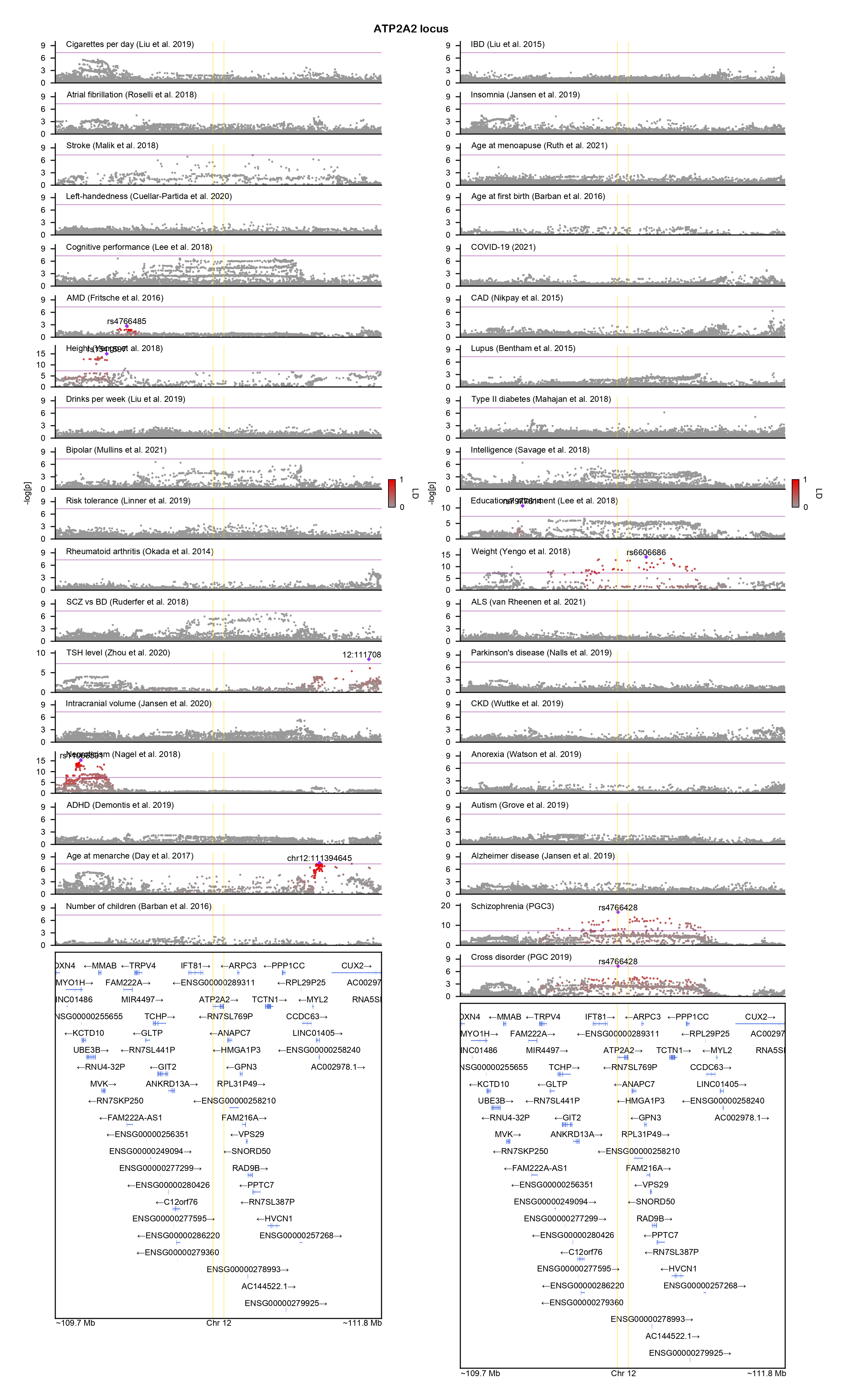

### BRINP2-locuszoom.png

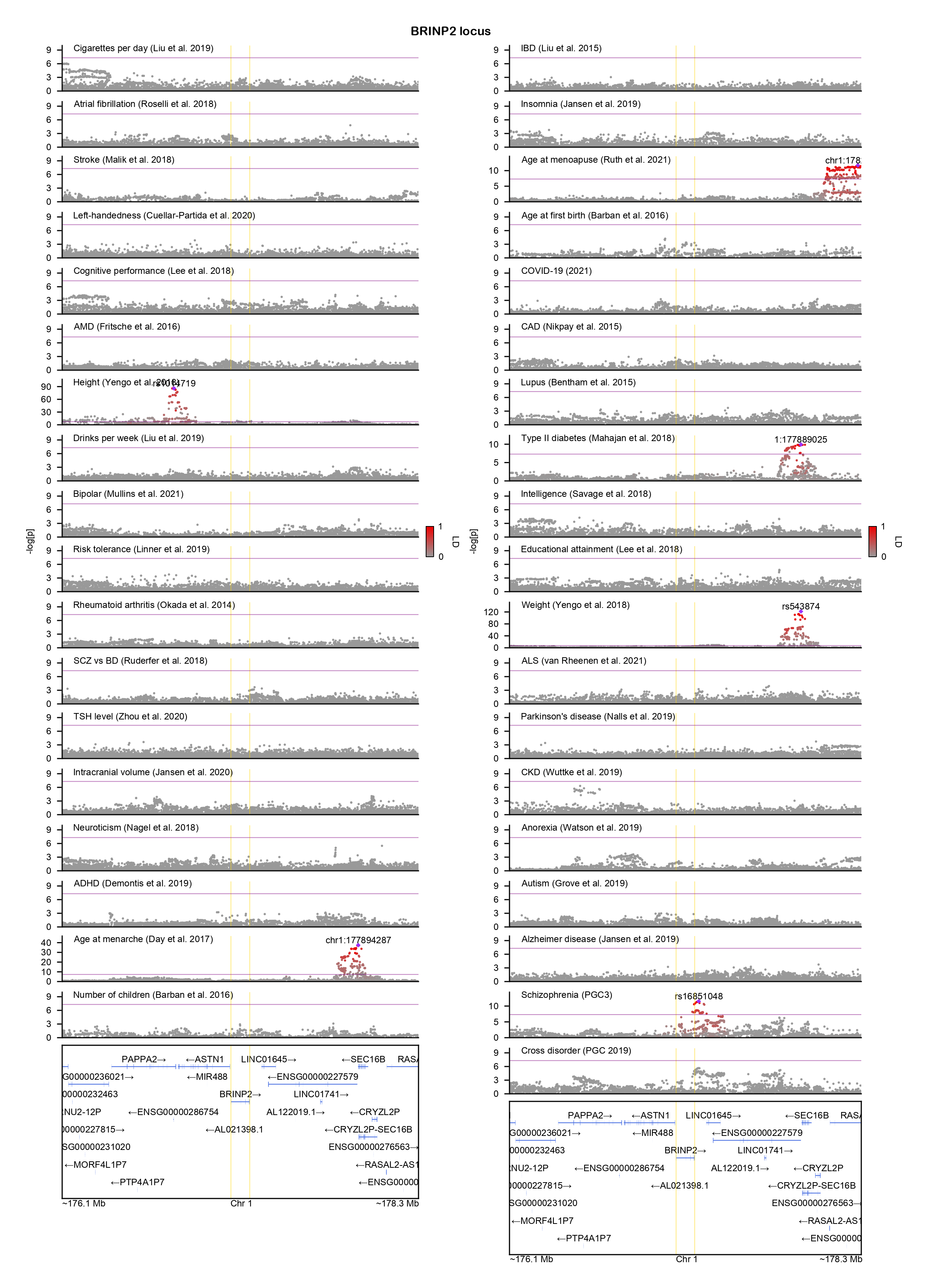

### BTG1-locuszoom.png

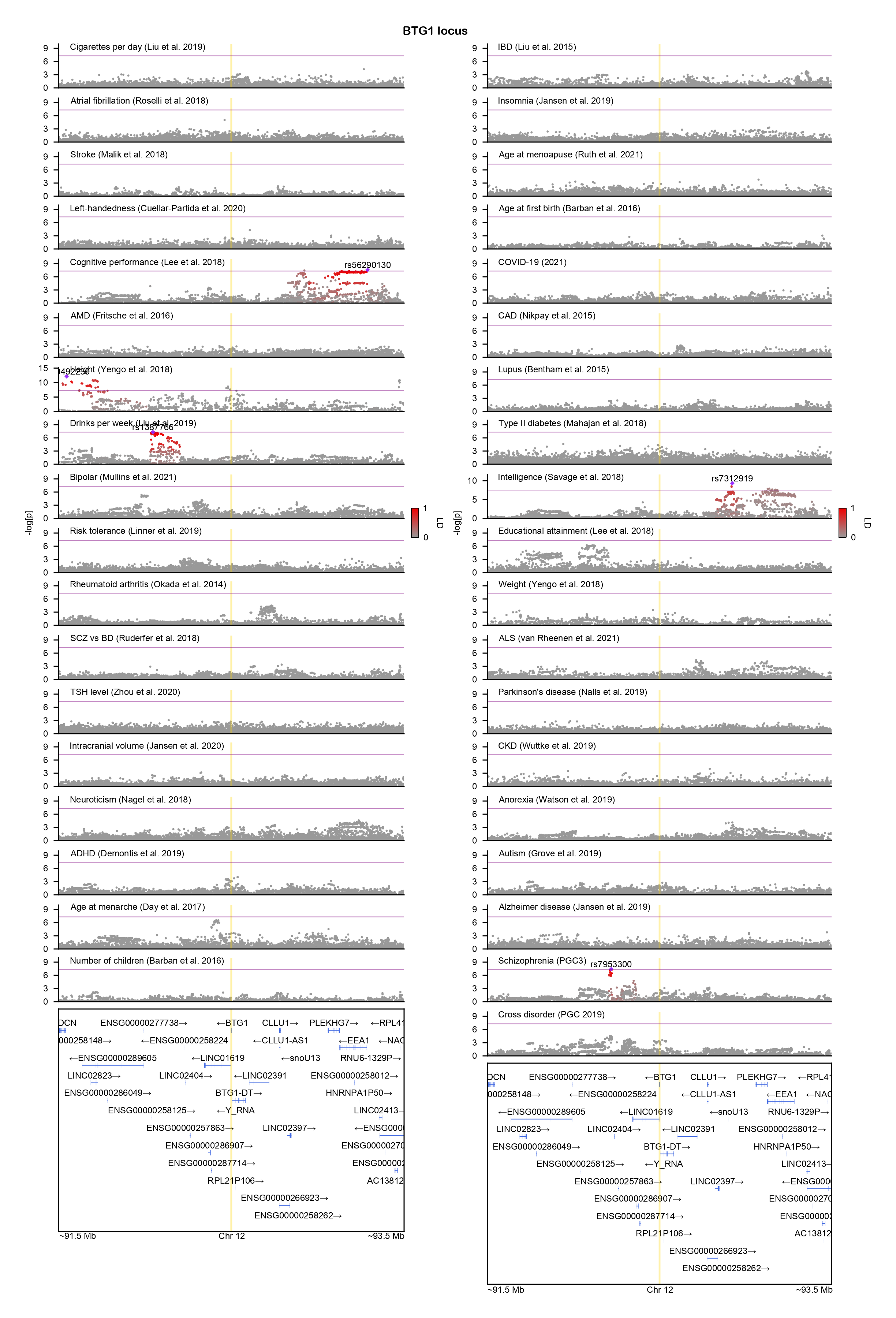

### C6orf118-locuszoom.png

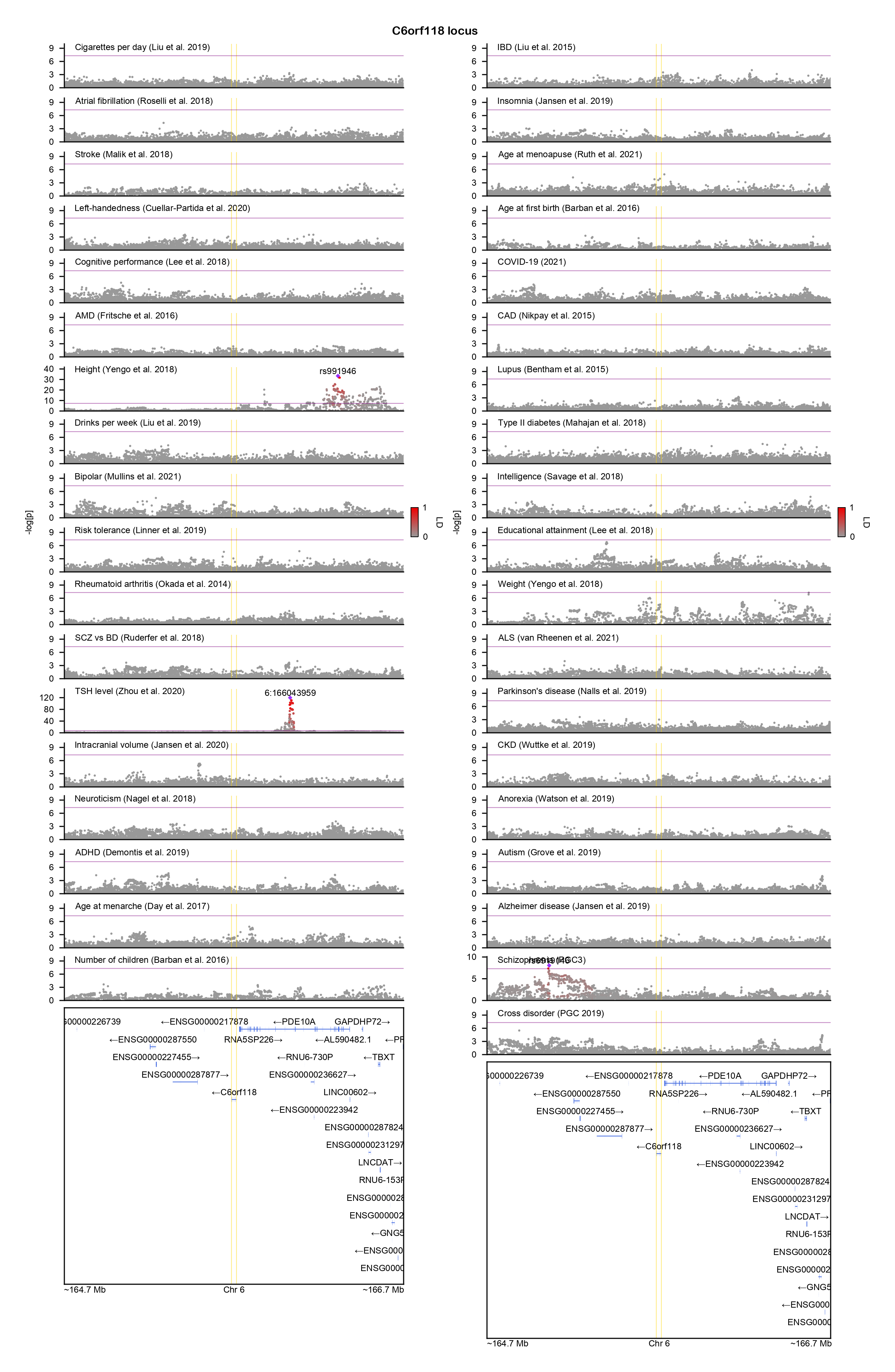

### CA8-locuszoom.png

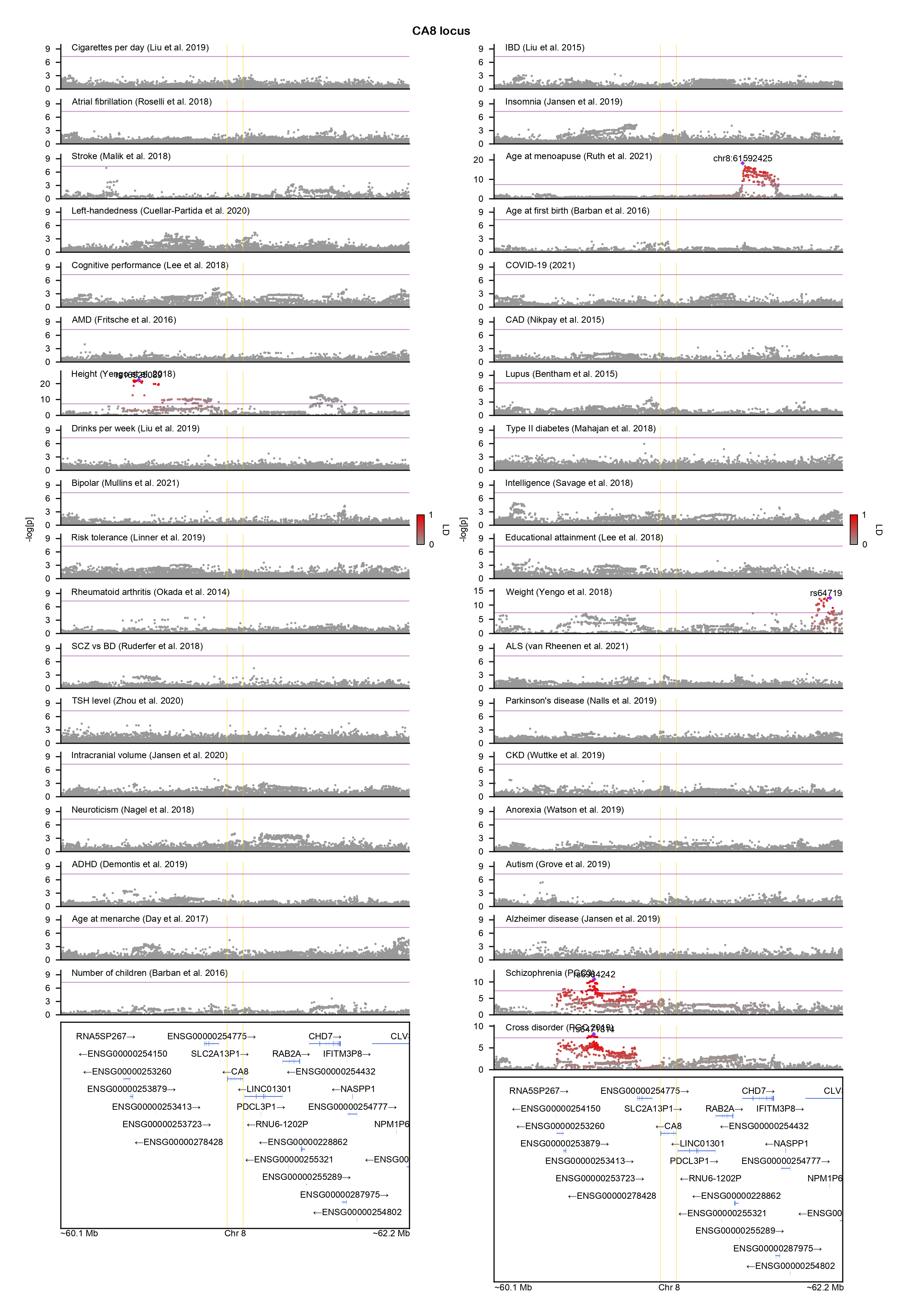

### CARMIL1-locuszoom.png

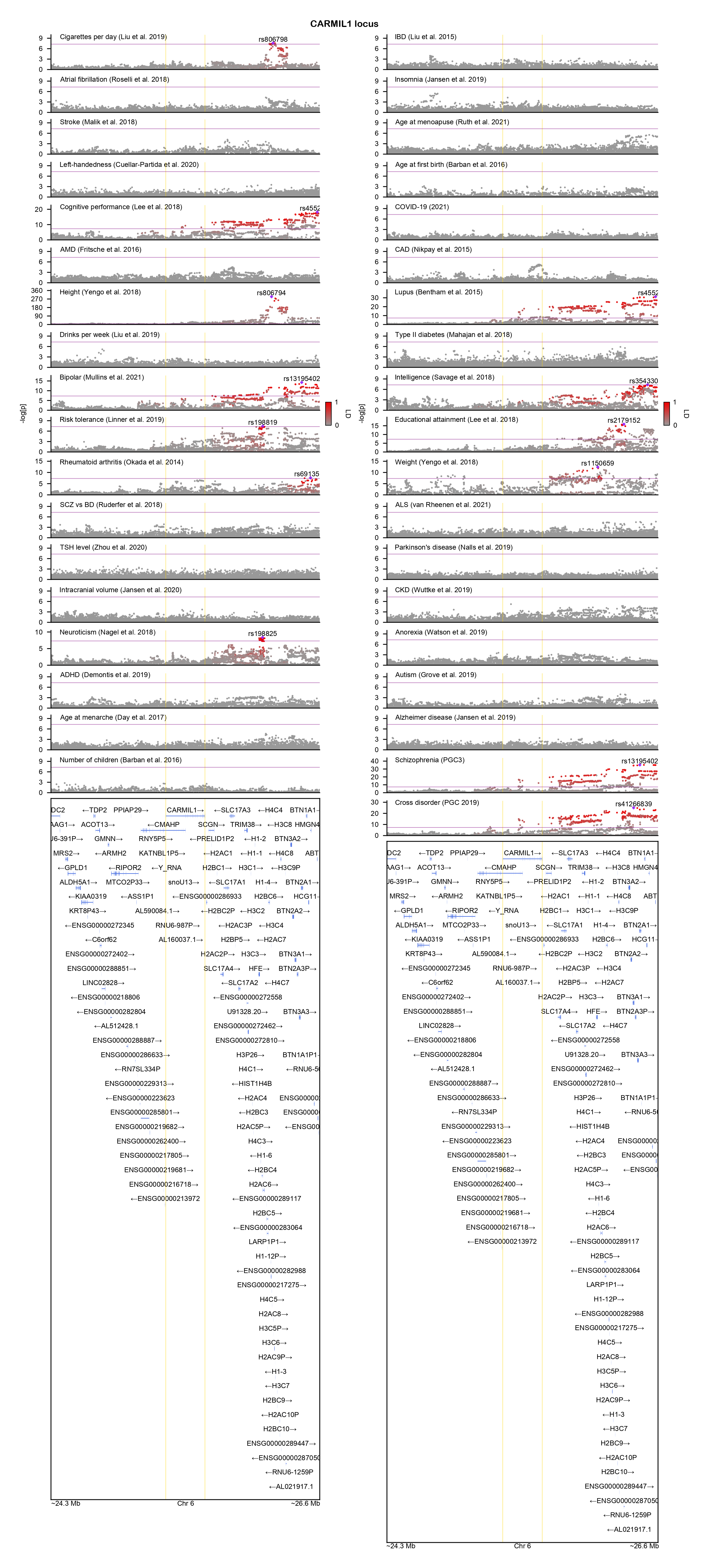

### CDH13-locuszoom.png

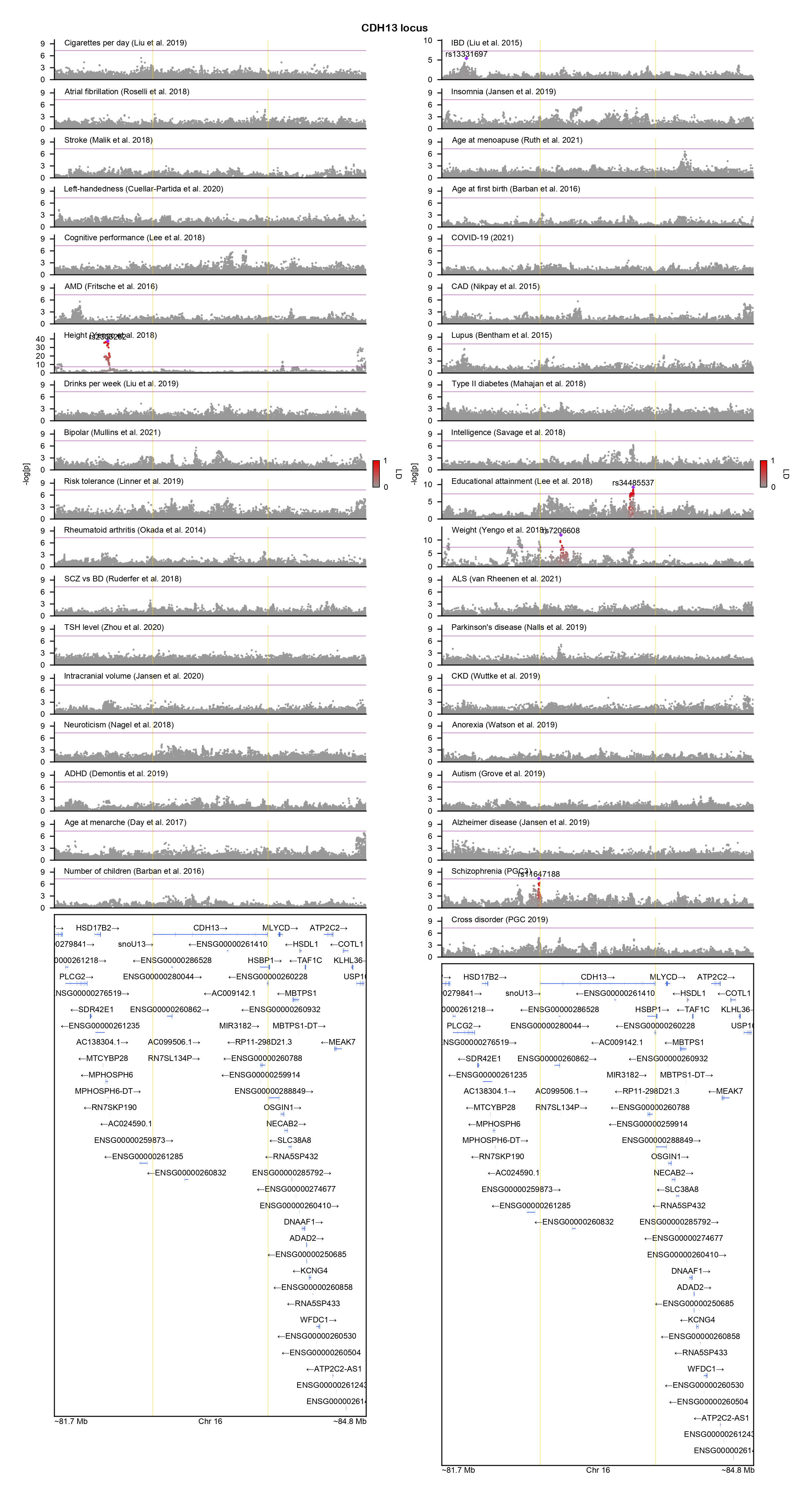

### CDH16-locuszoom.png

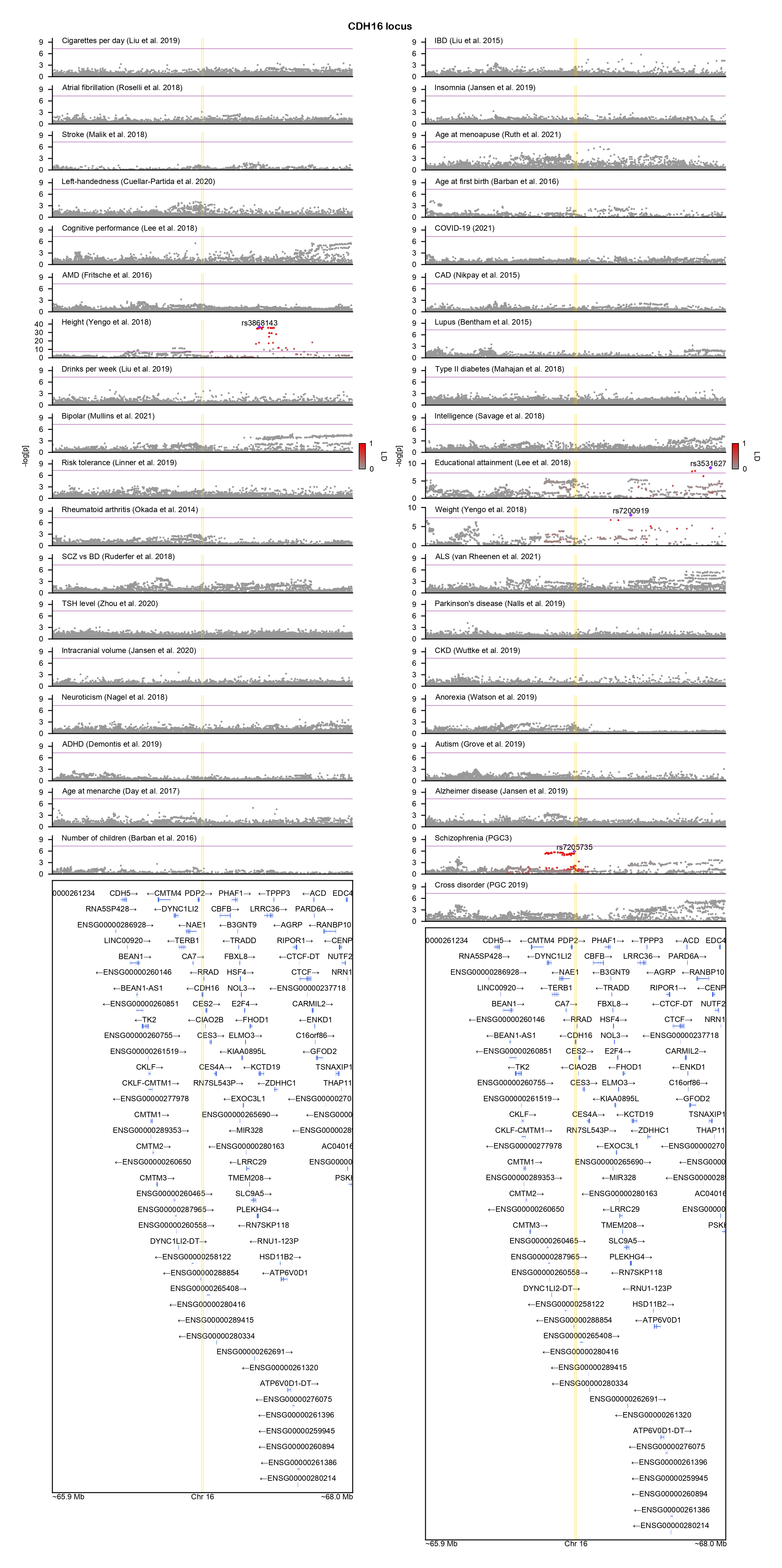

### CHRM3-locuszoom.png

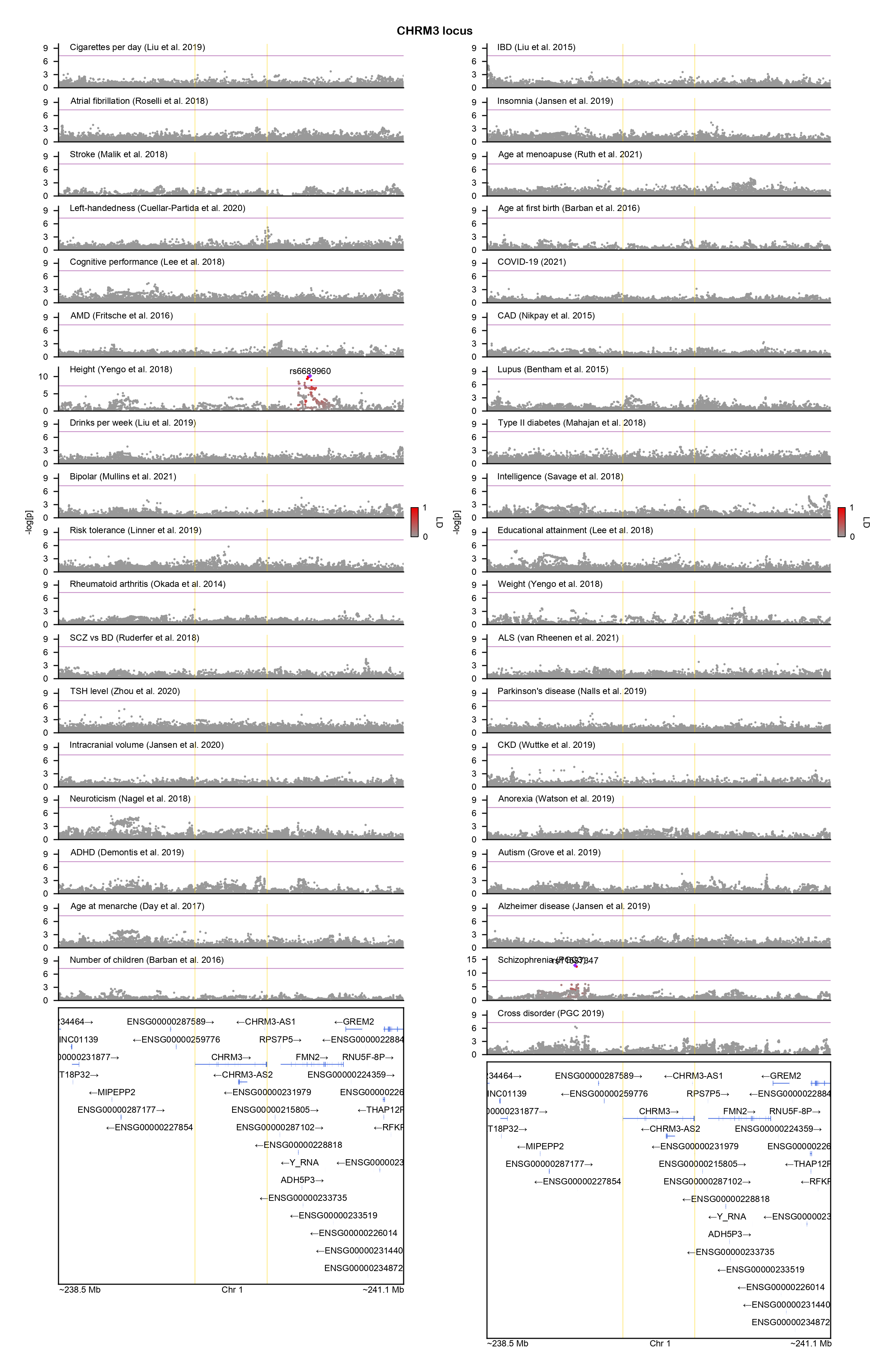

### CNNM2-locuszoom.png

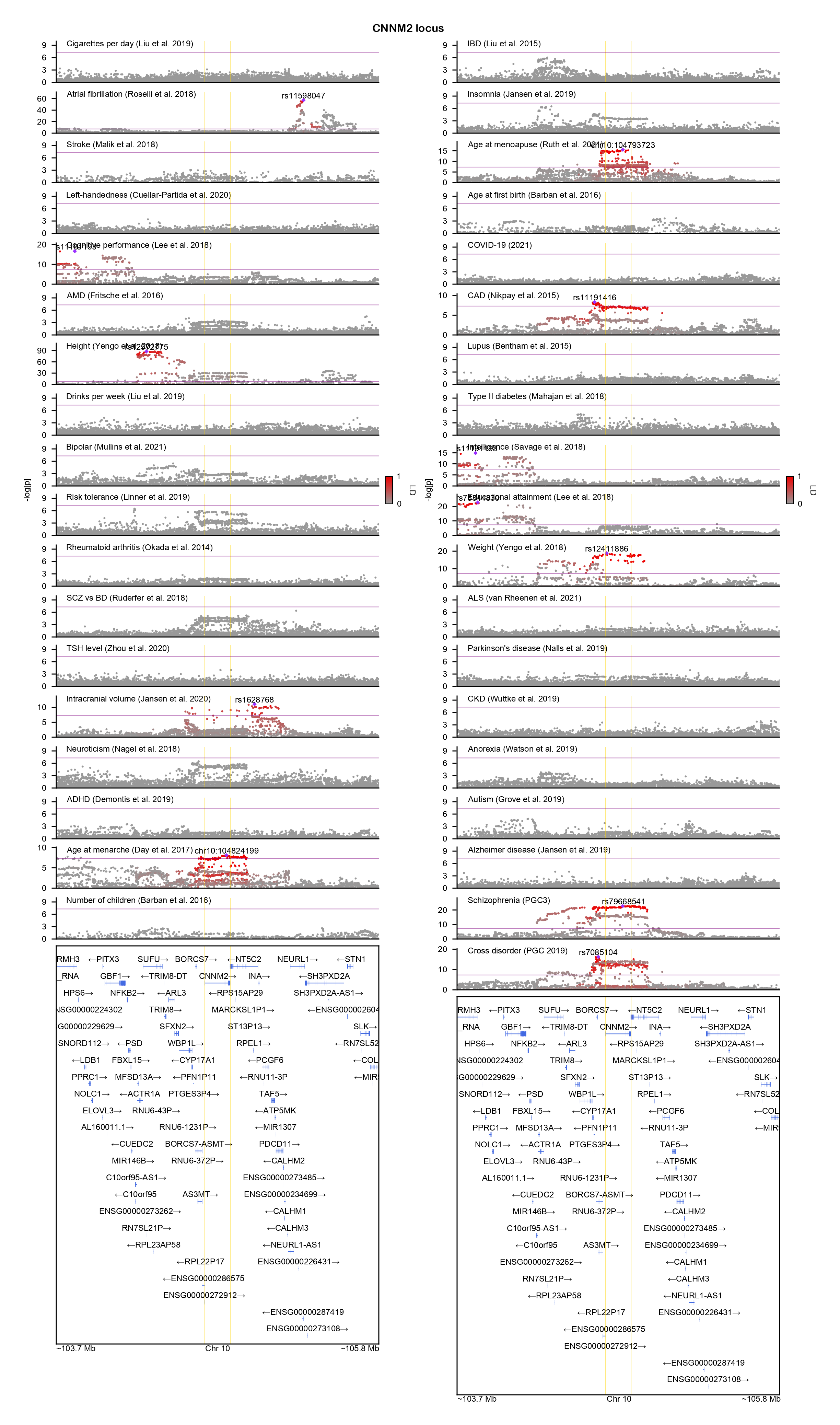

### CNOT1-locuszoom.png

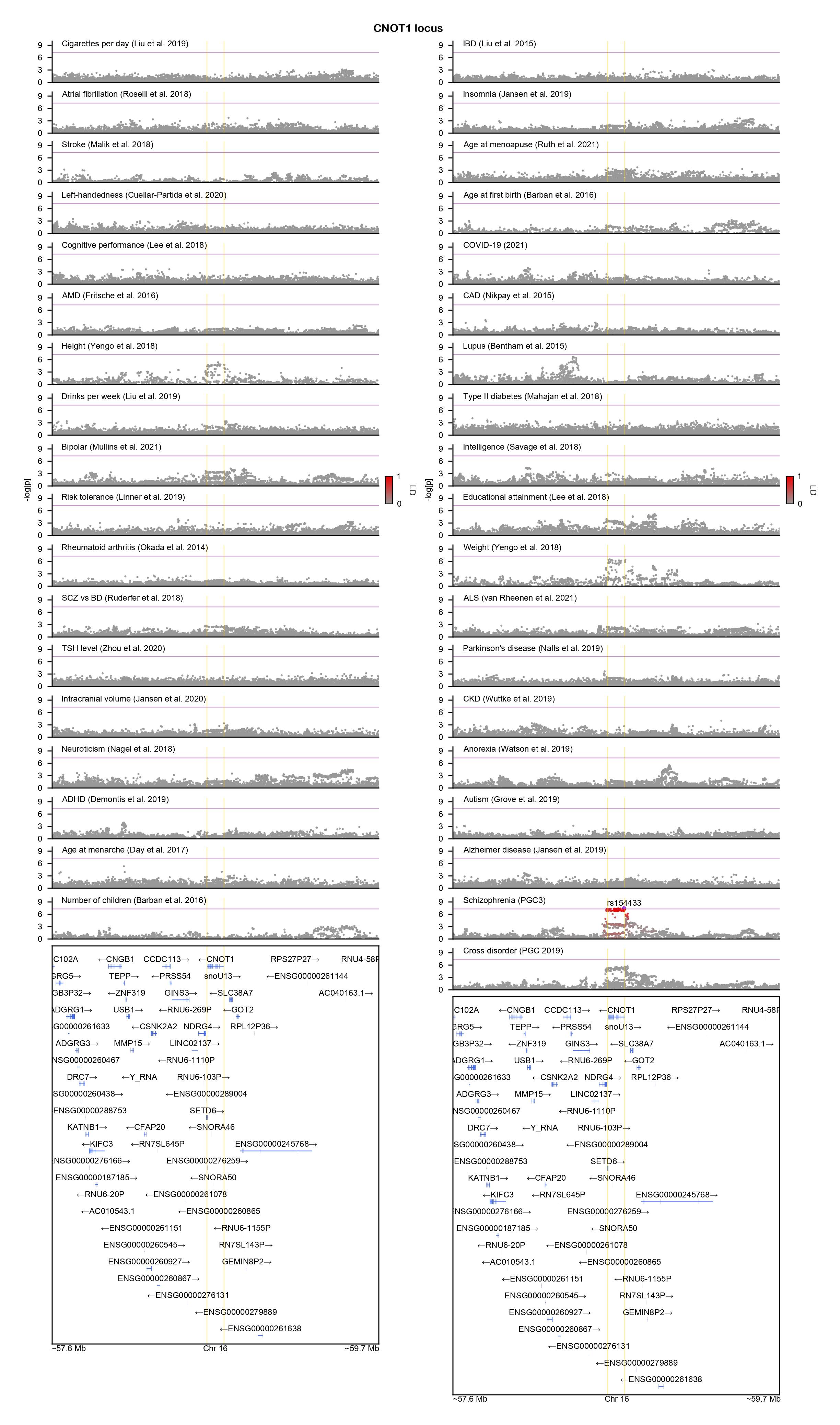

### COQ10B-locuszoom.png

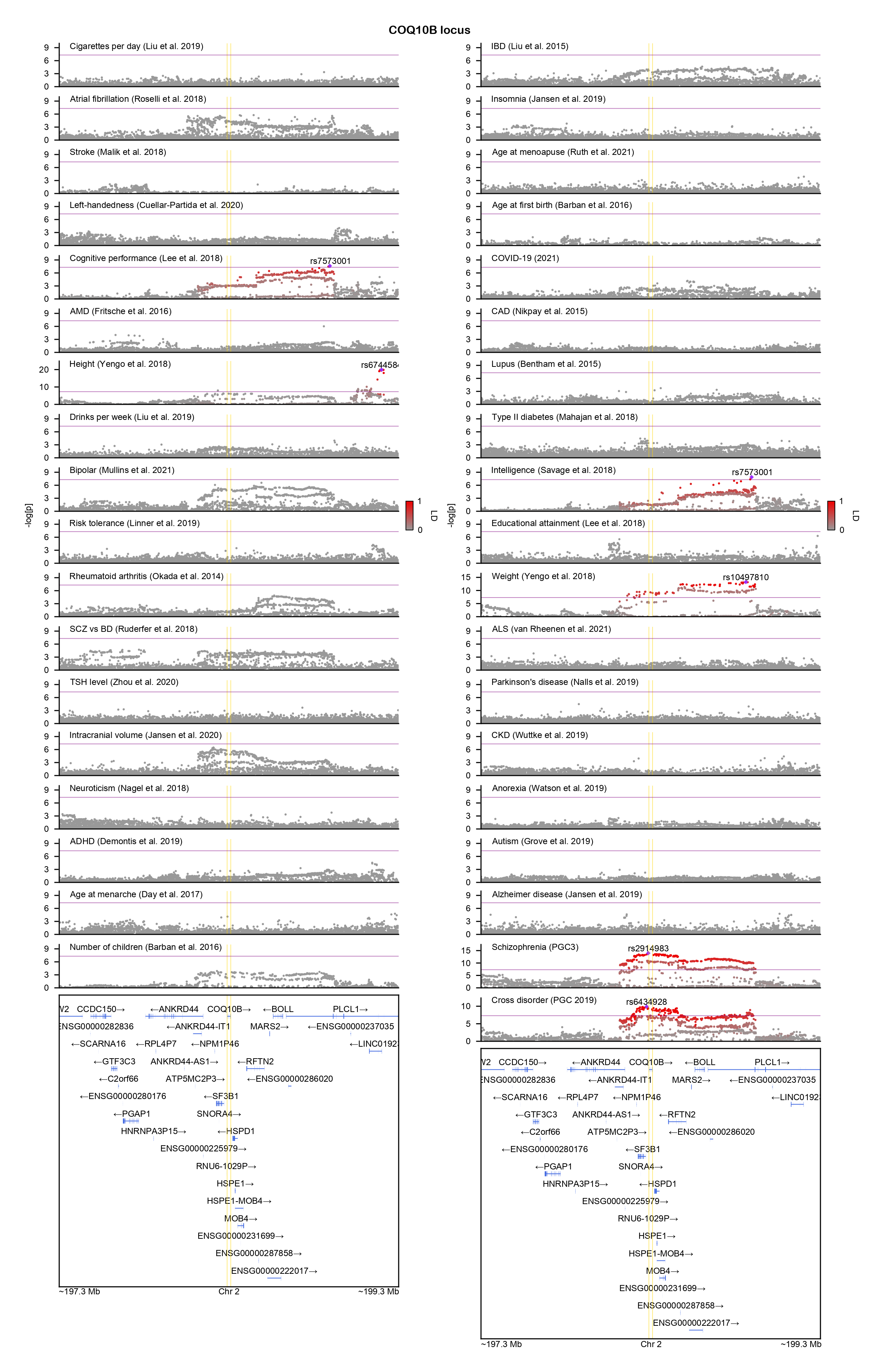

### CUL9-locuszoom.png

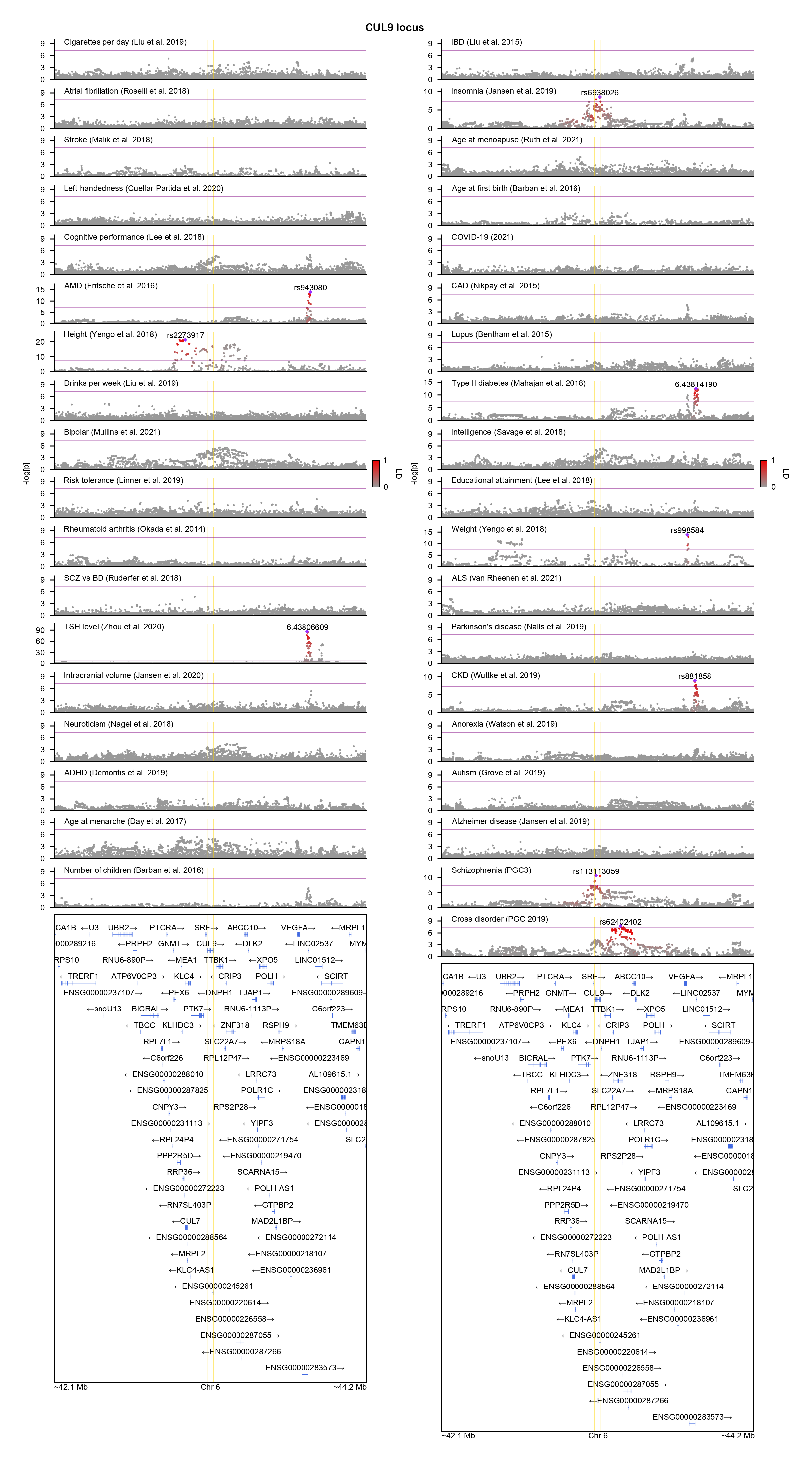

### CXXC5-locuszoom.png

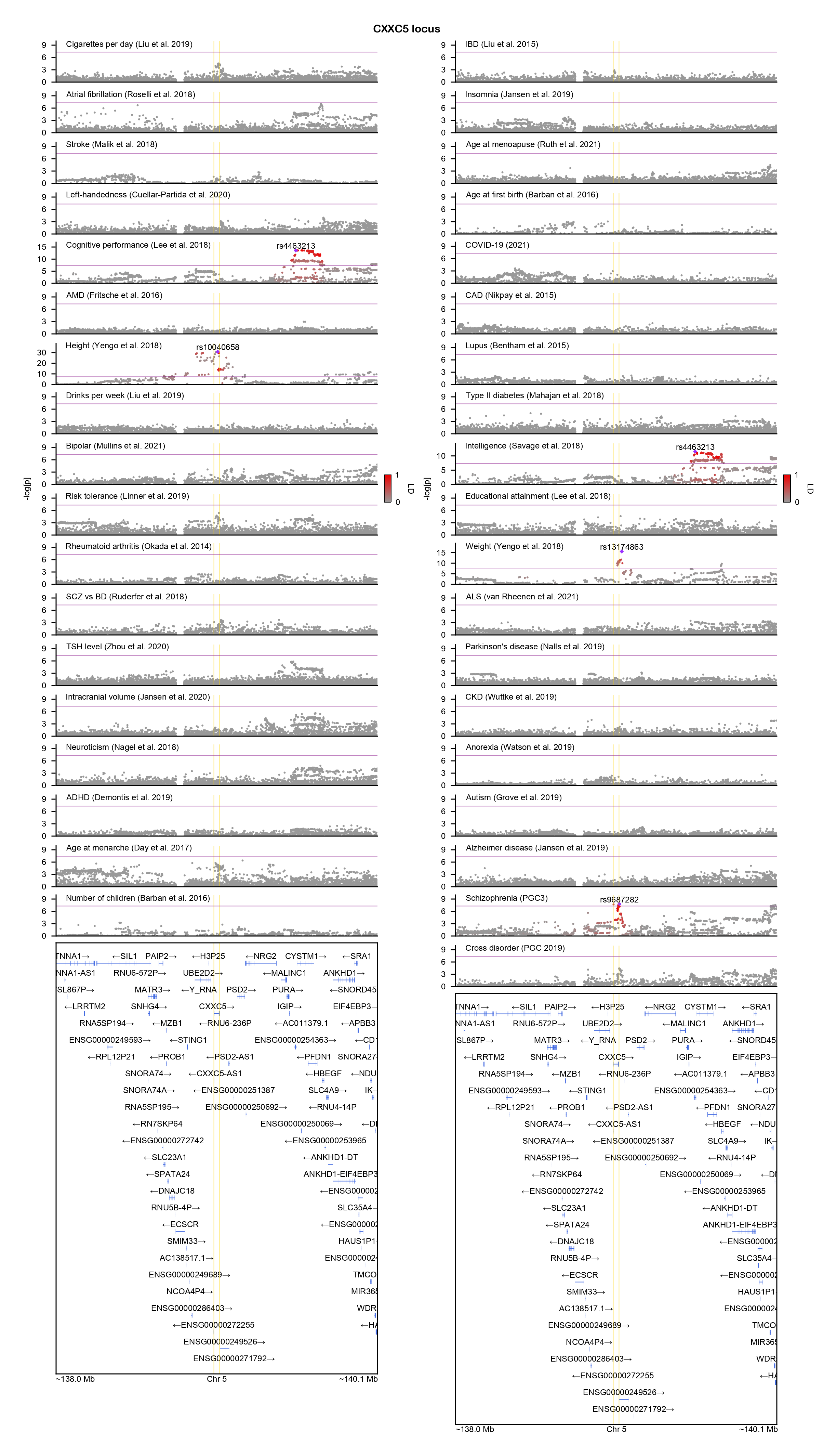

### DCLK3-locuszoom.png

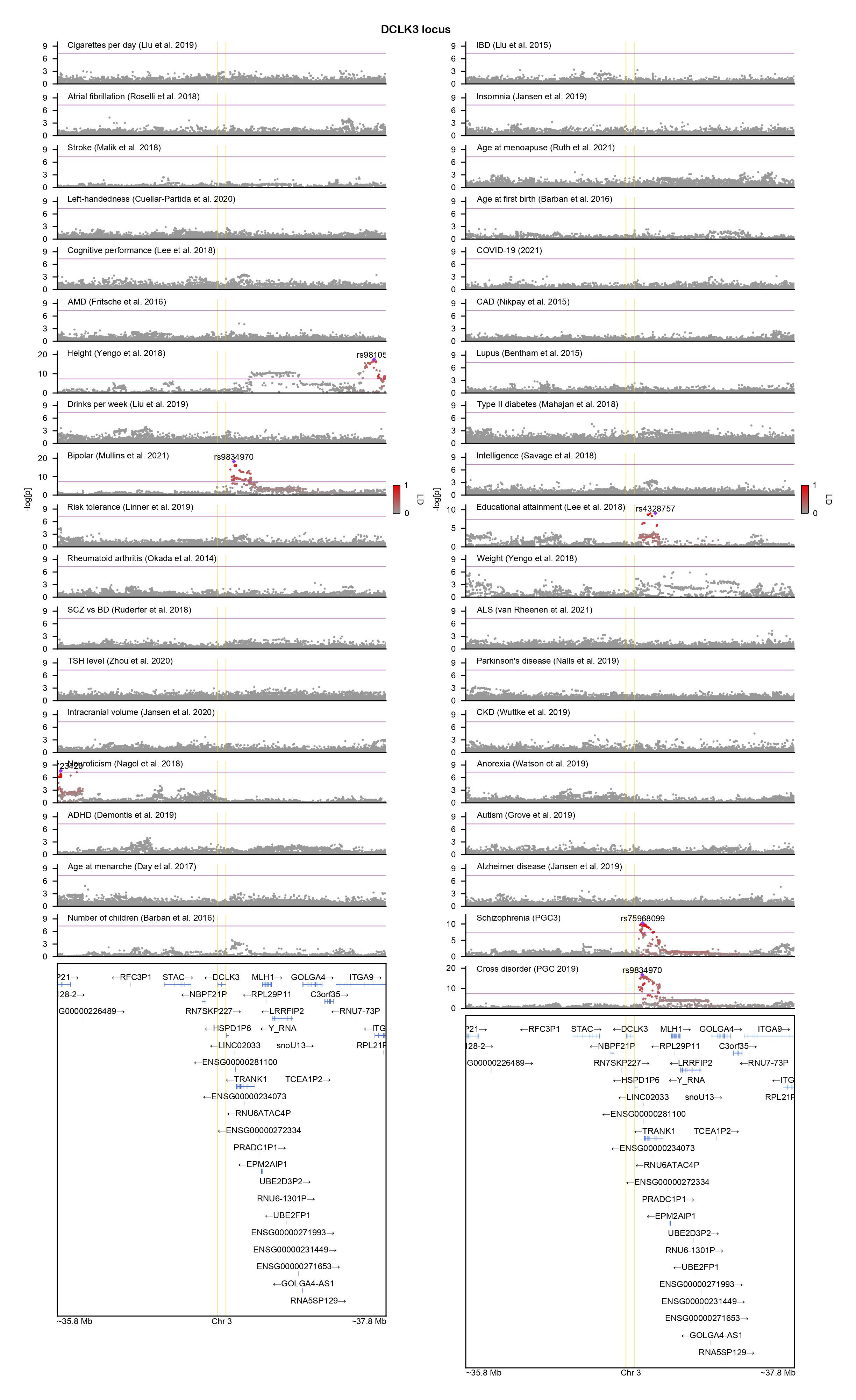

### DENND4B-locuszoom.png

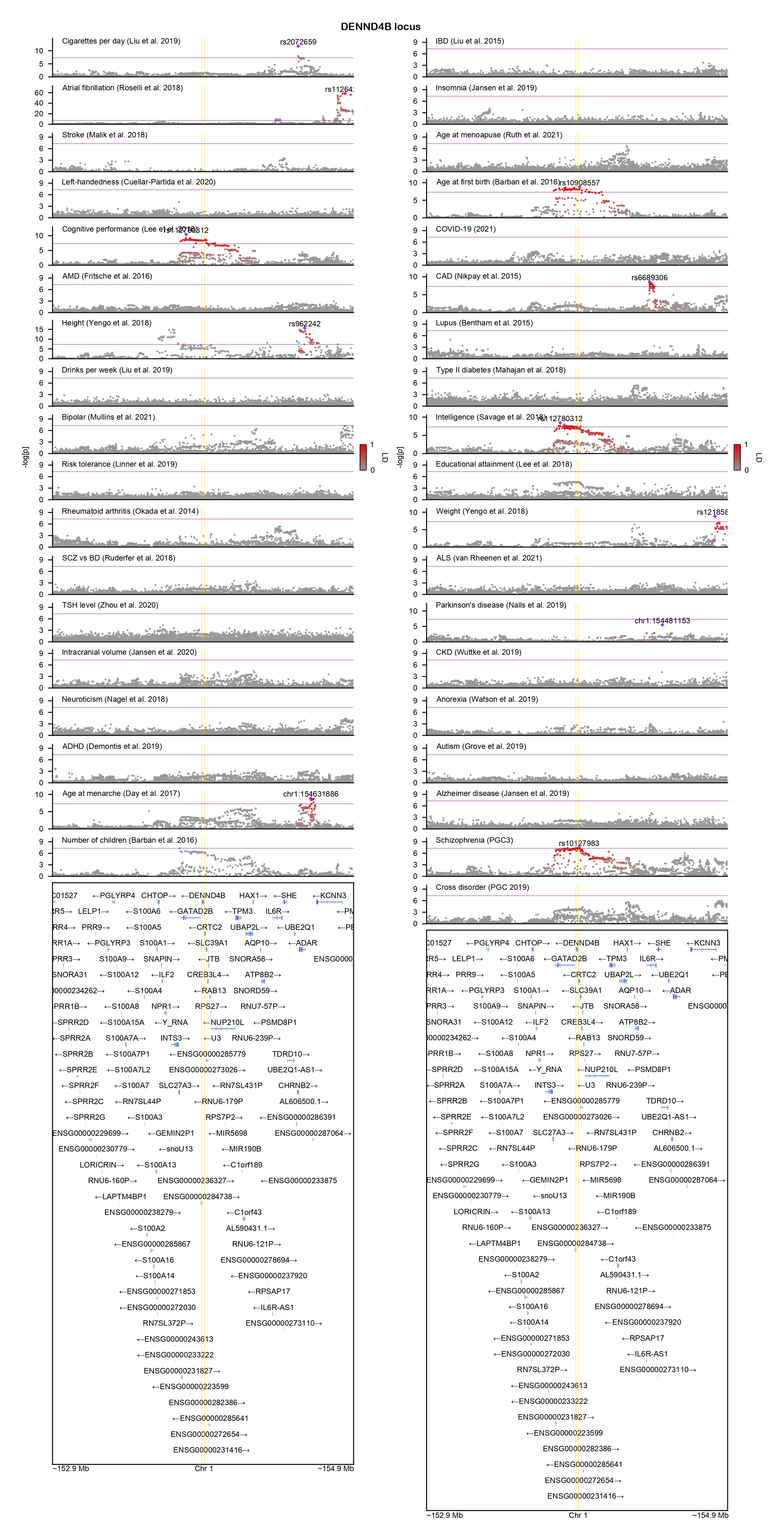

### DLX2-locuszoom.png

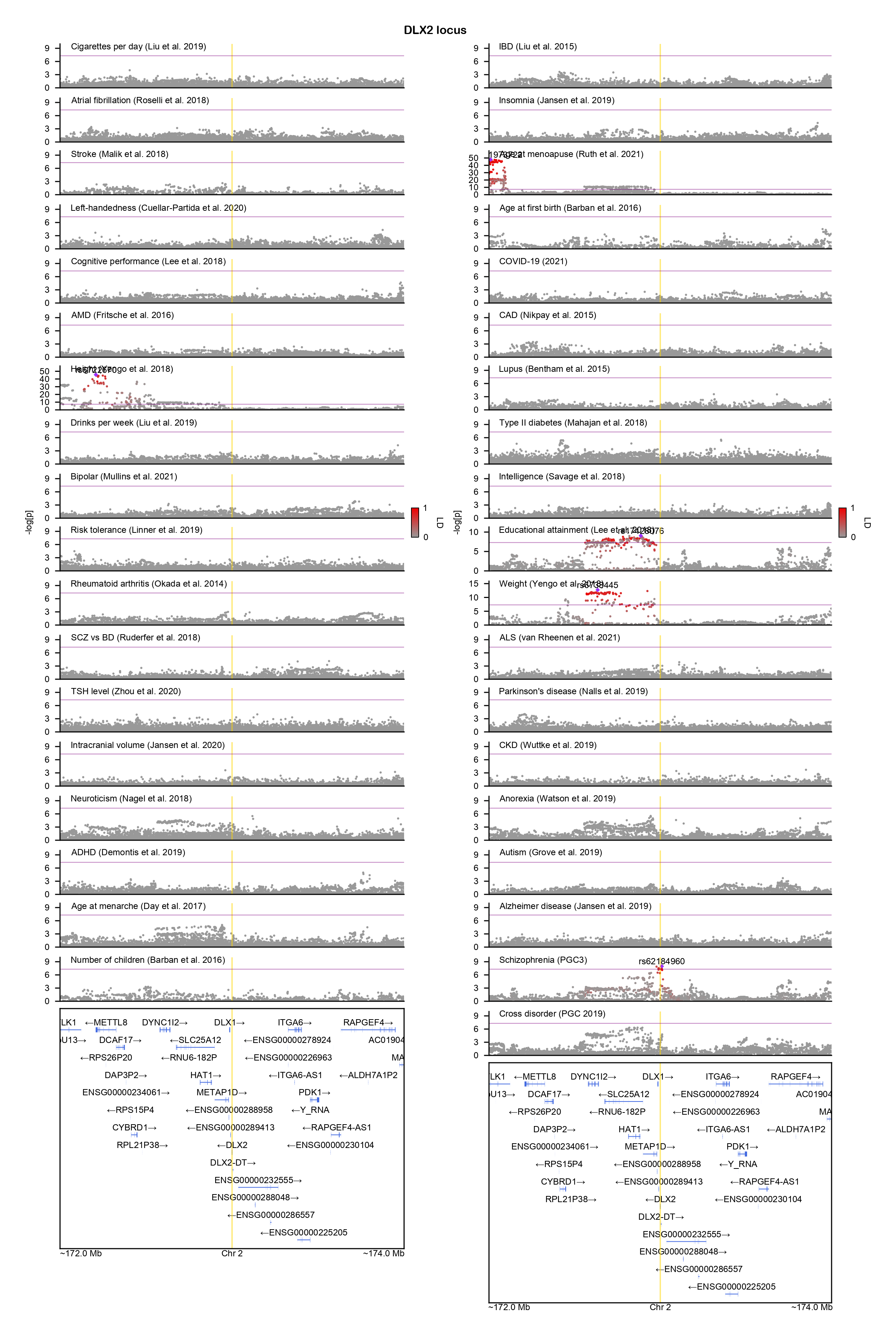

### DMRTA1-locuszoom.png

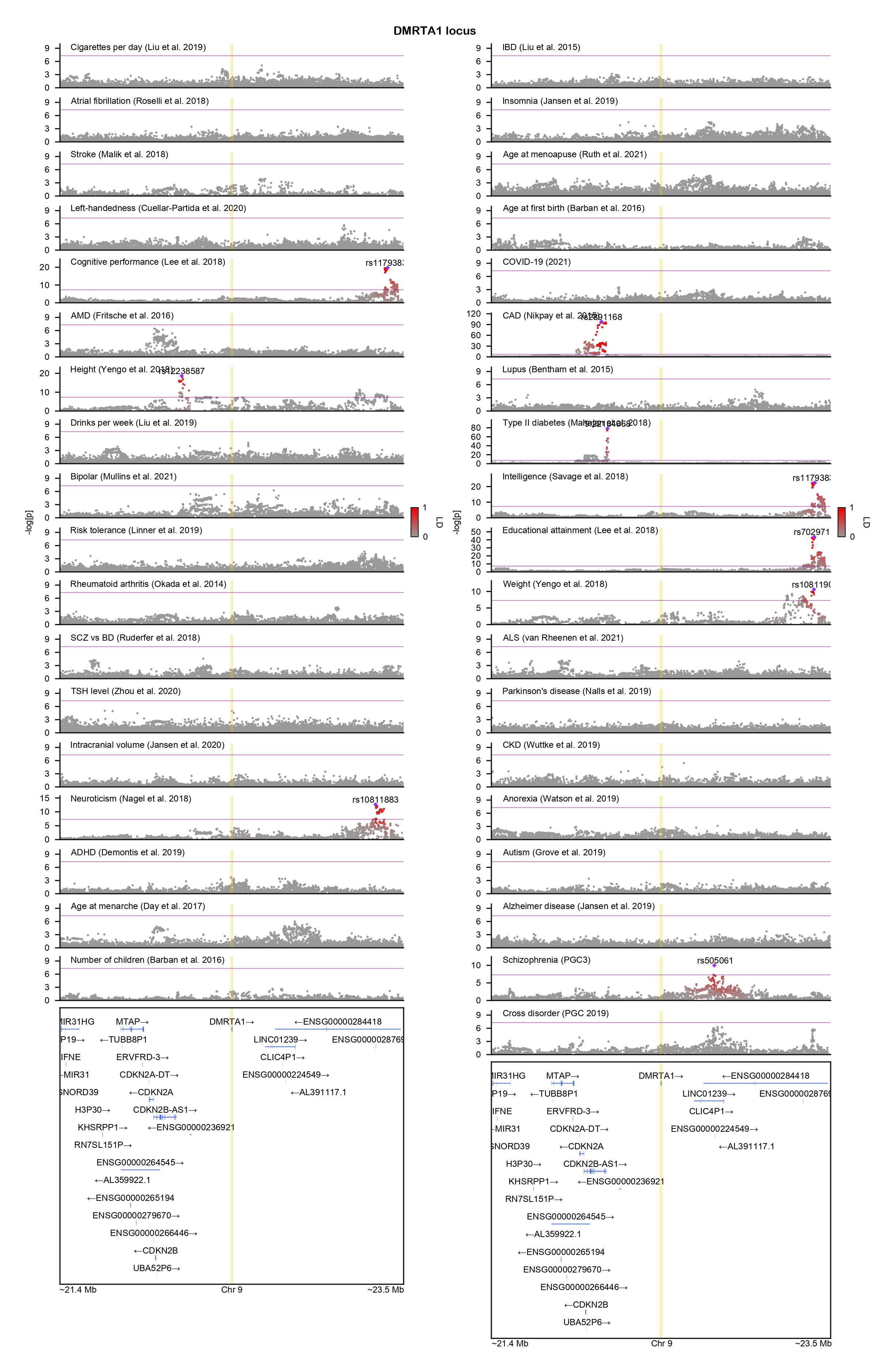

### DNAJA3-locuszoom.png

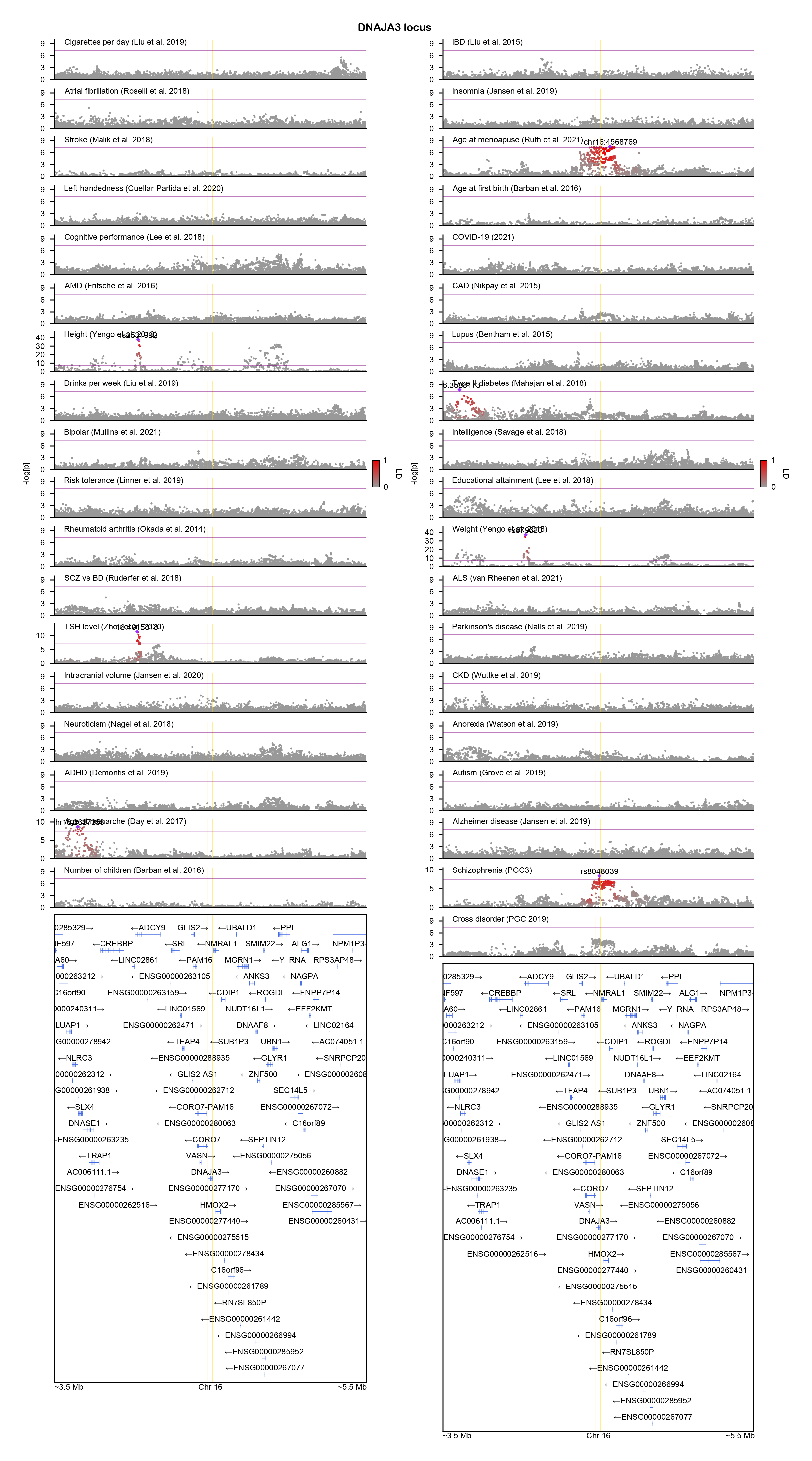

### DNAJC11-locuszoom.png

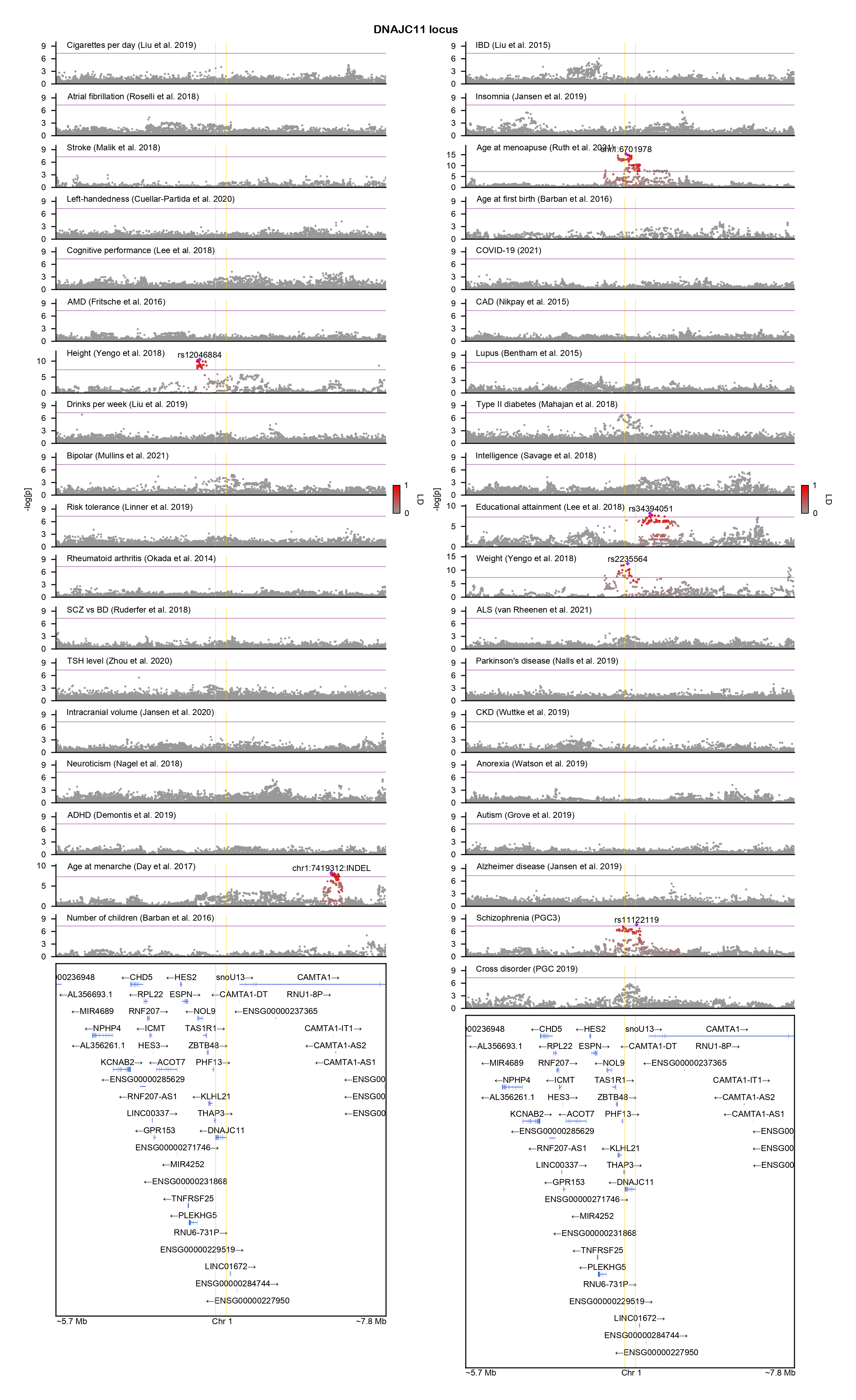

### DOT1L-locuszoom.png

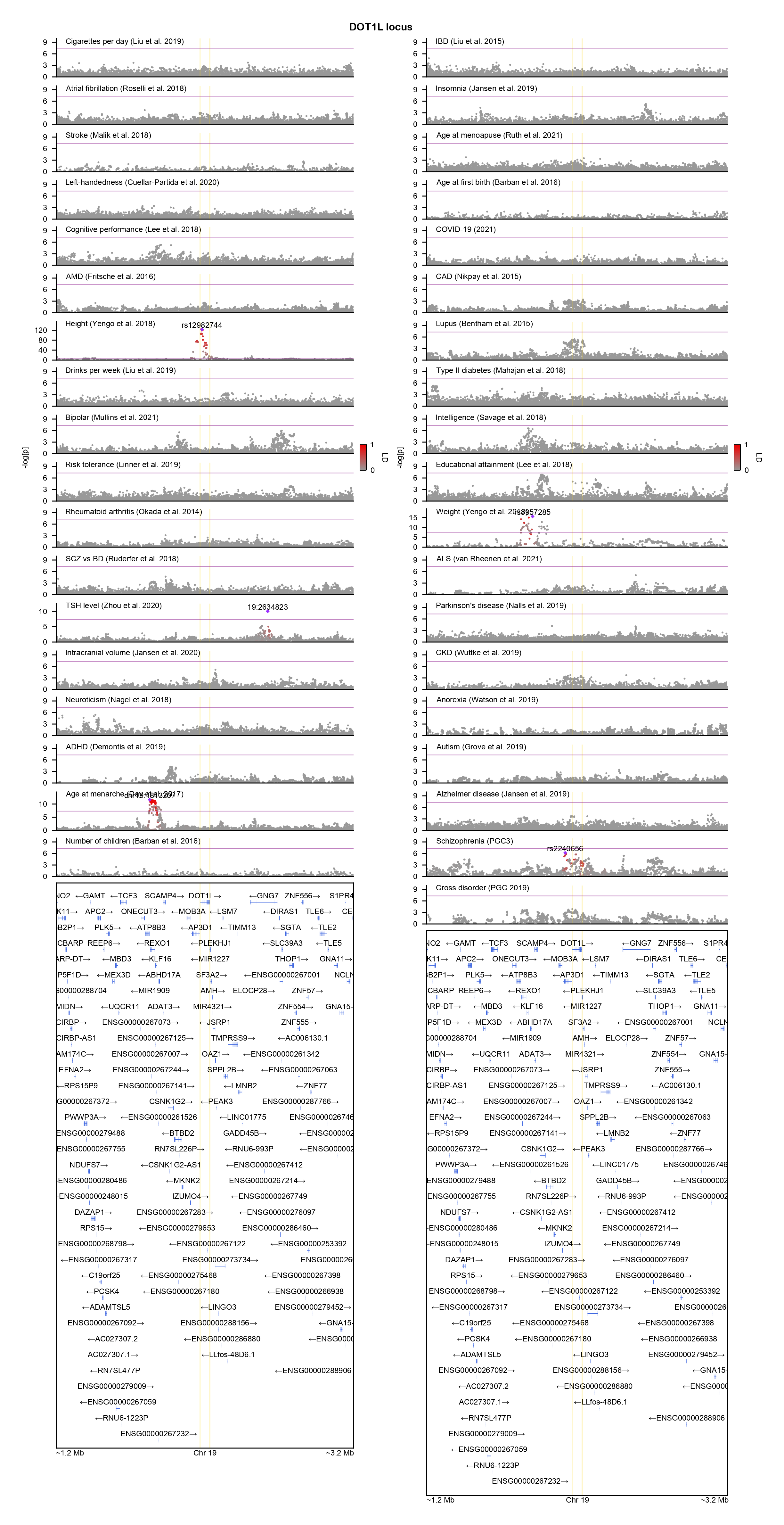

### DPYD-locuszoom.png

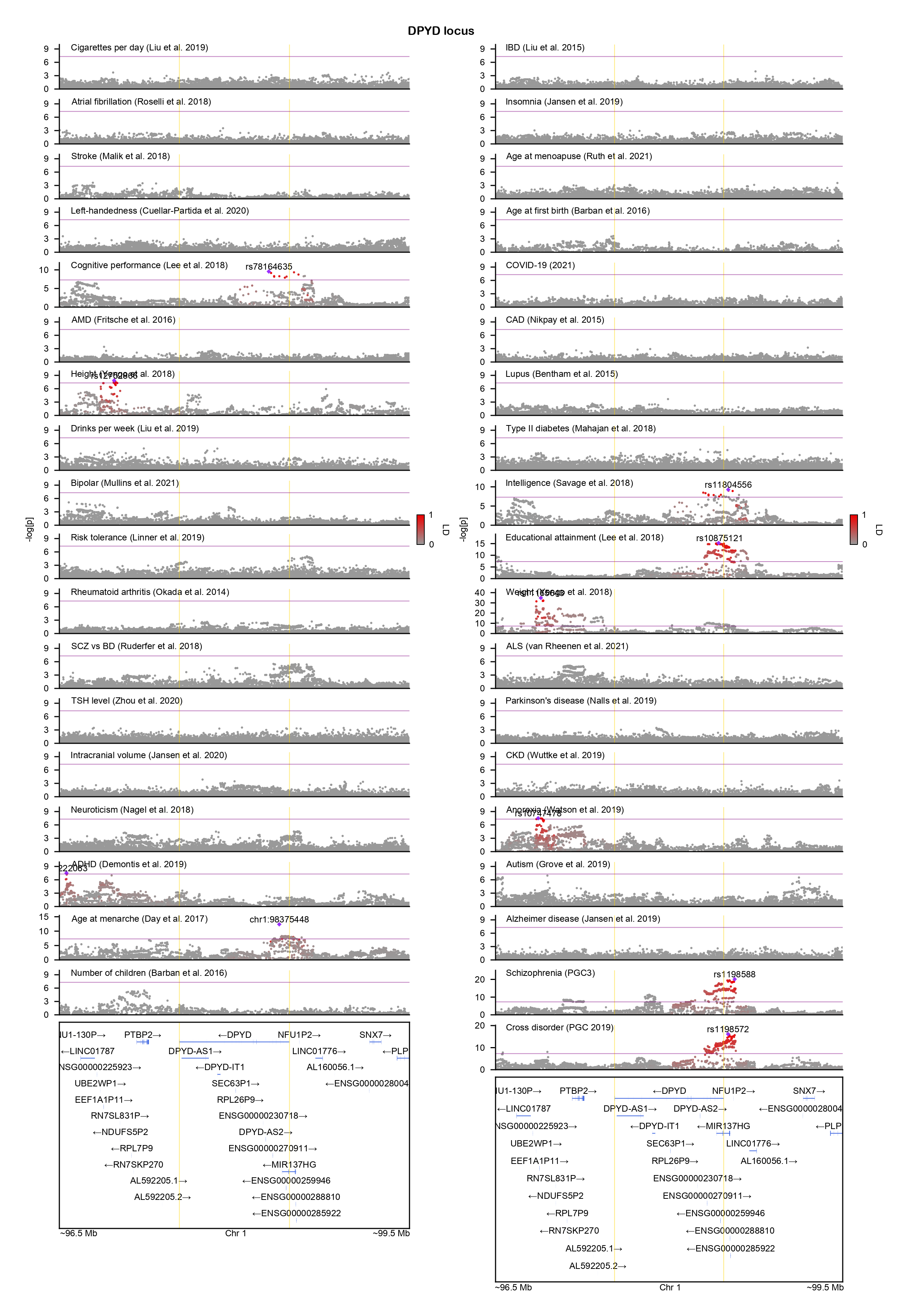

### DSC3-locuszoom.png

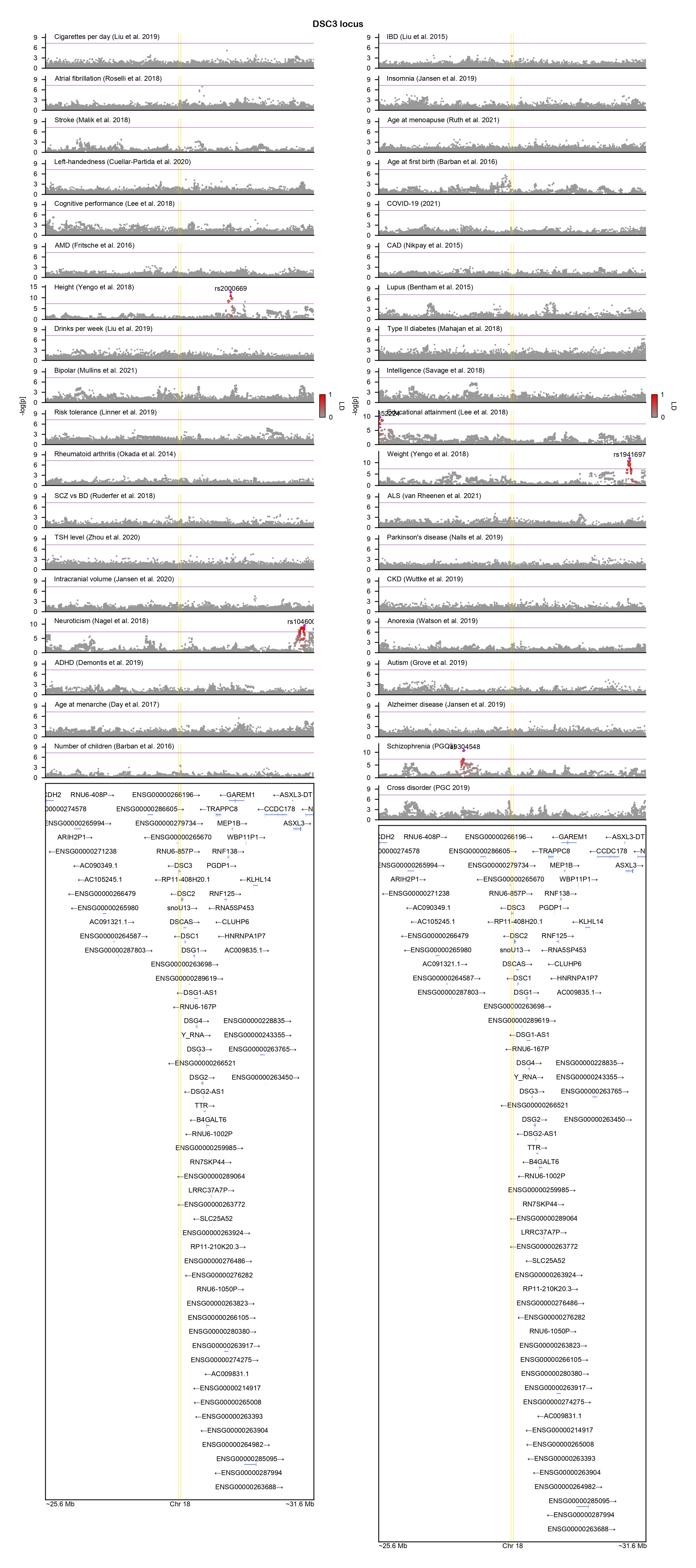

### EFNA5-locuszoom.png

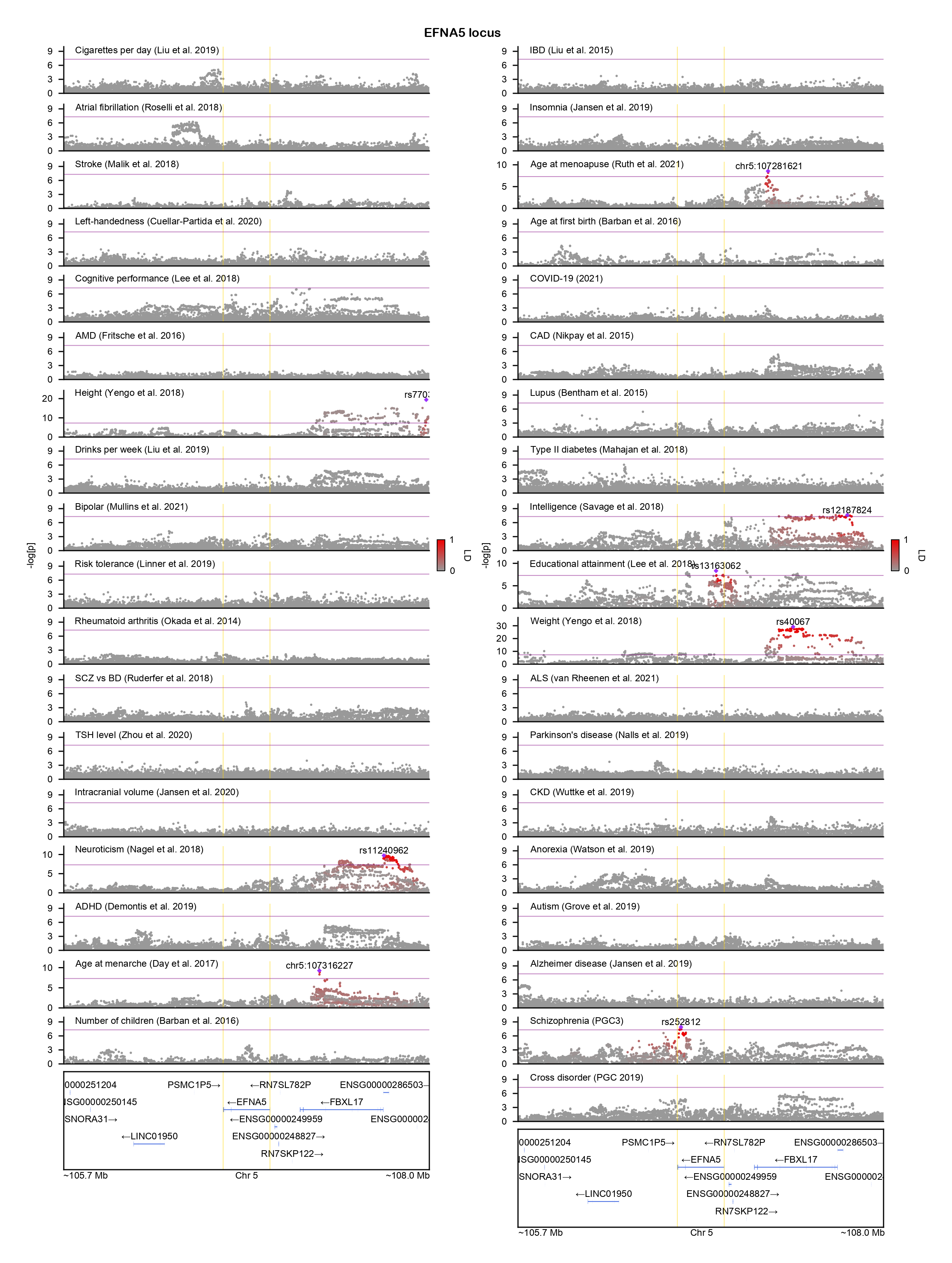

### ELAC2-locuszoom.png

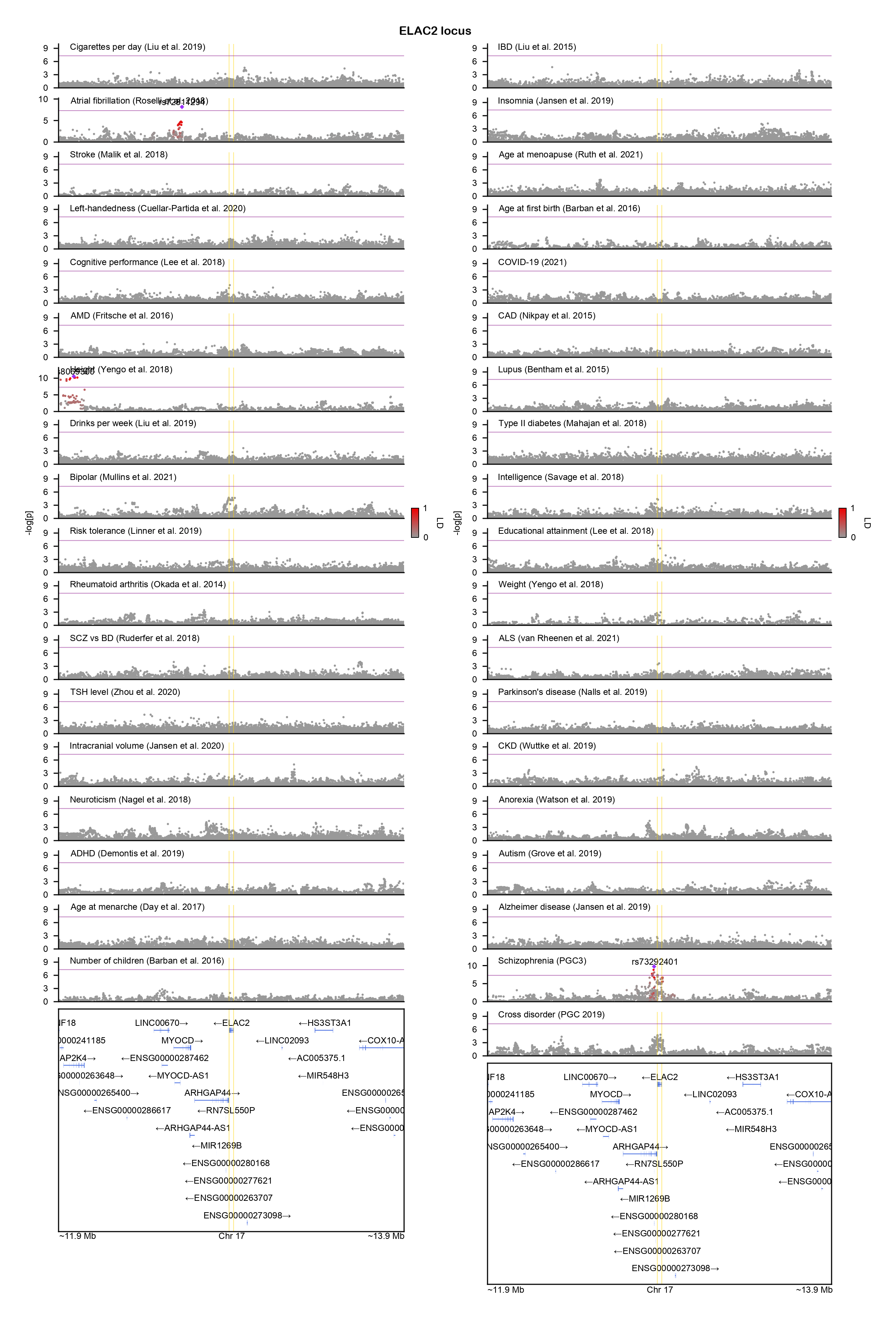

### EMB-locuszoom.png

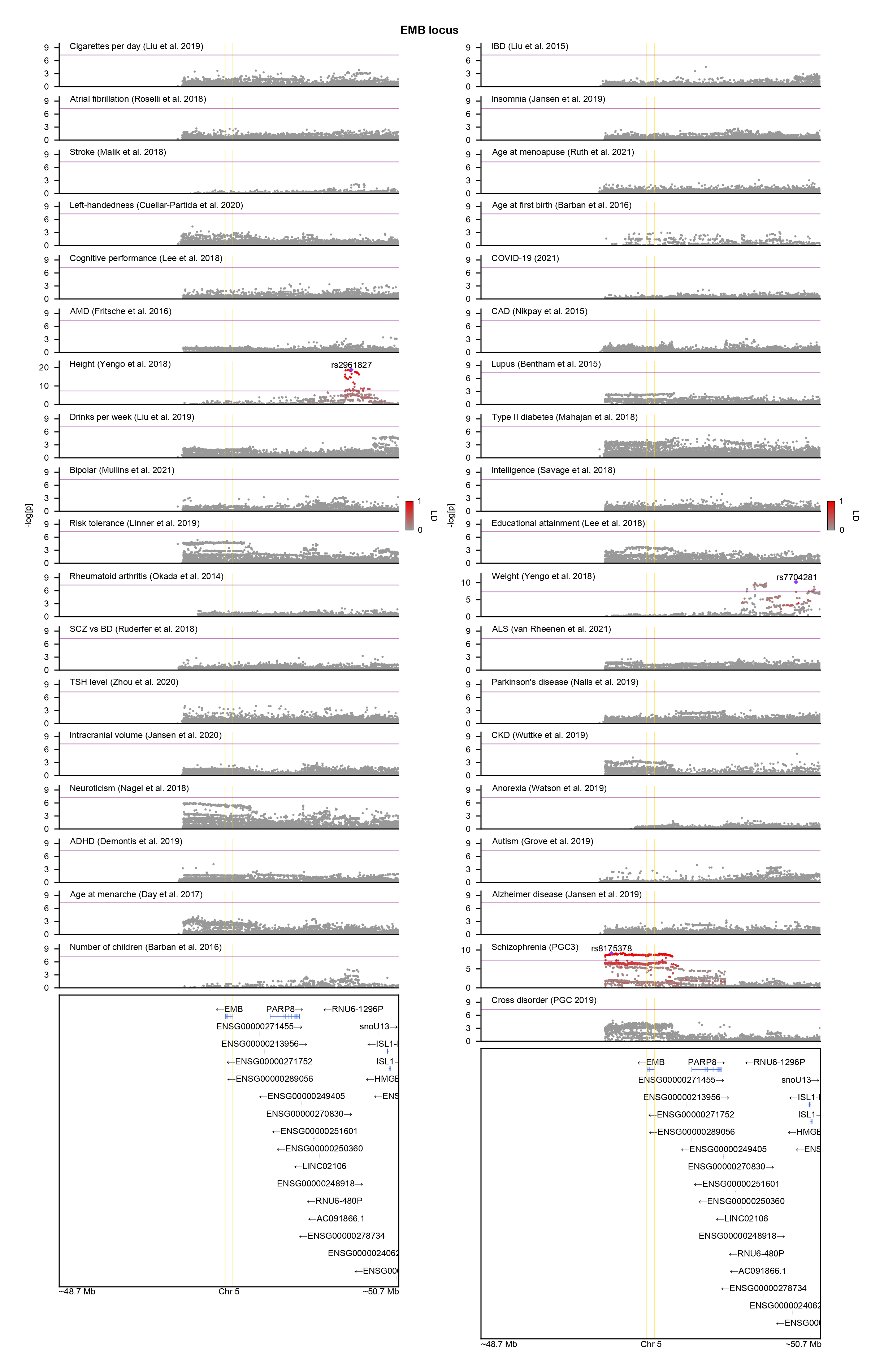

### ENOX1-locuszoom.png

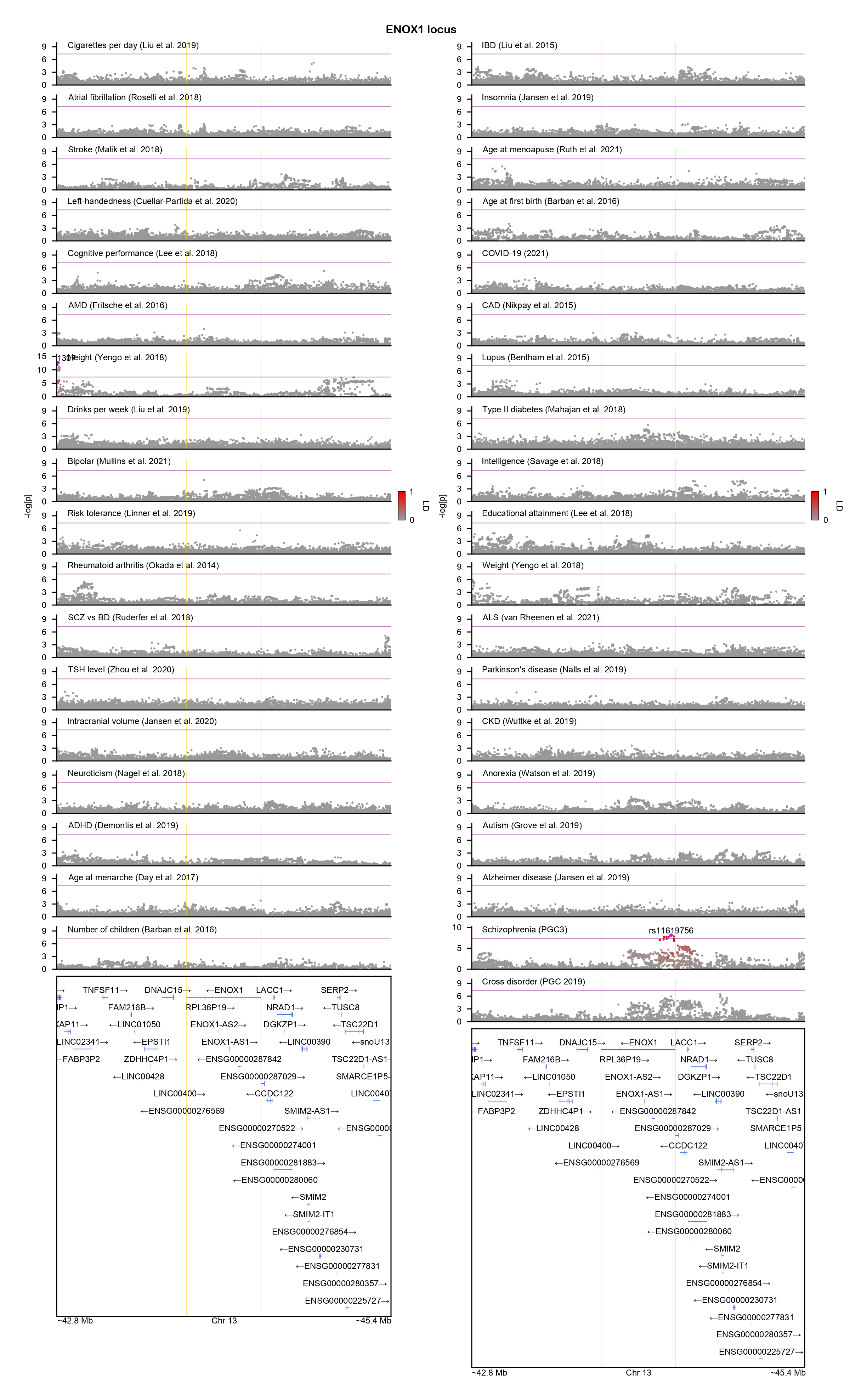

### ENSG00000258830-locuszoom.png

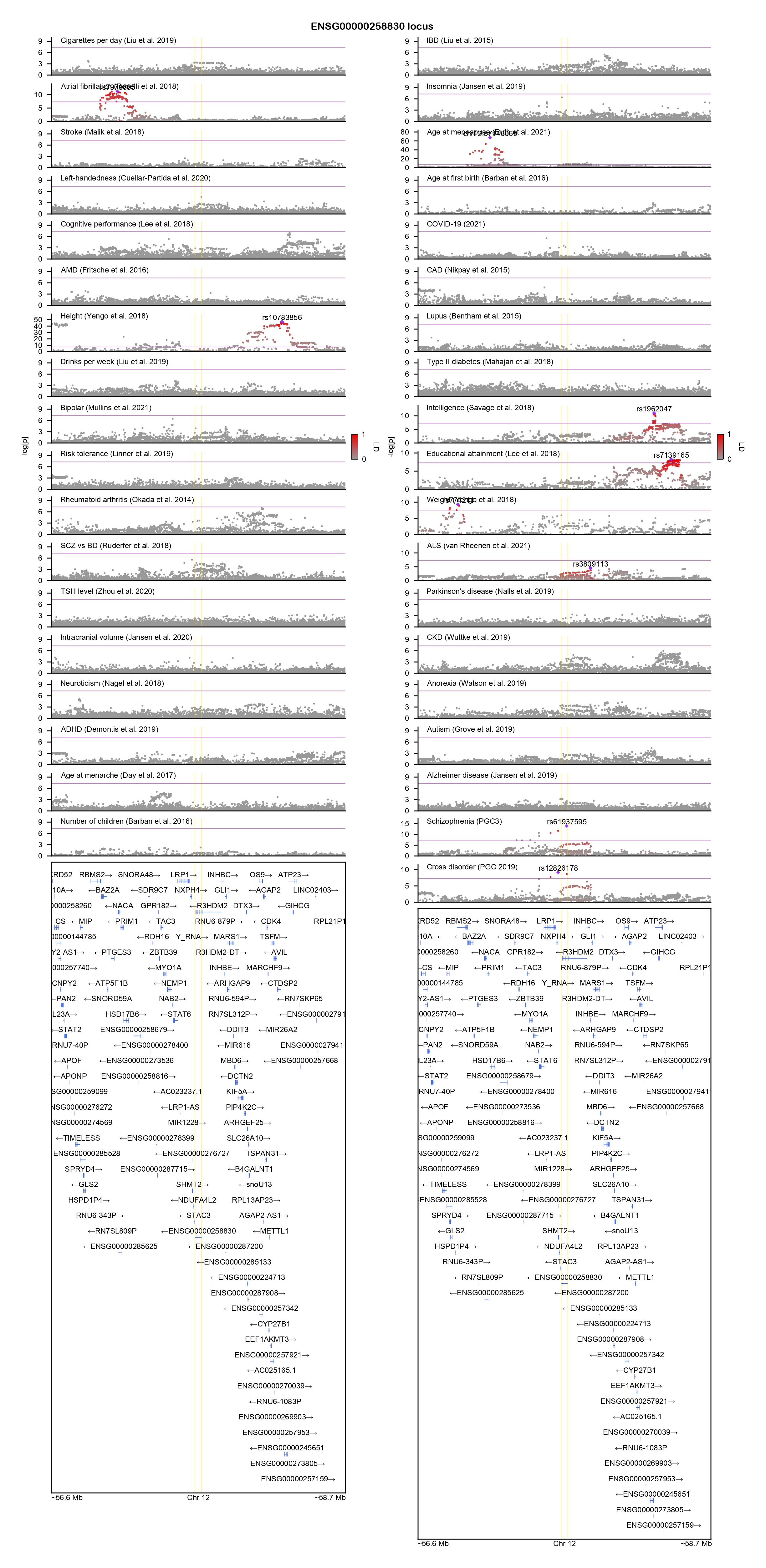

### EPHA7-locuszoom.png

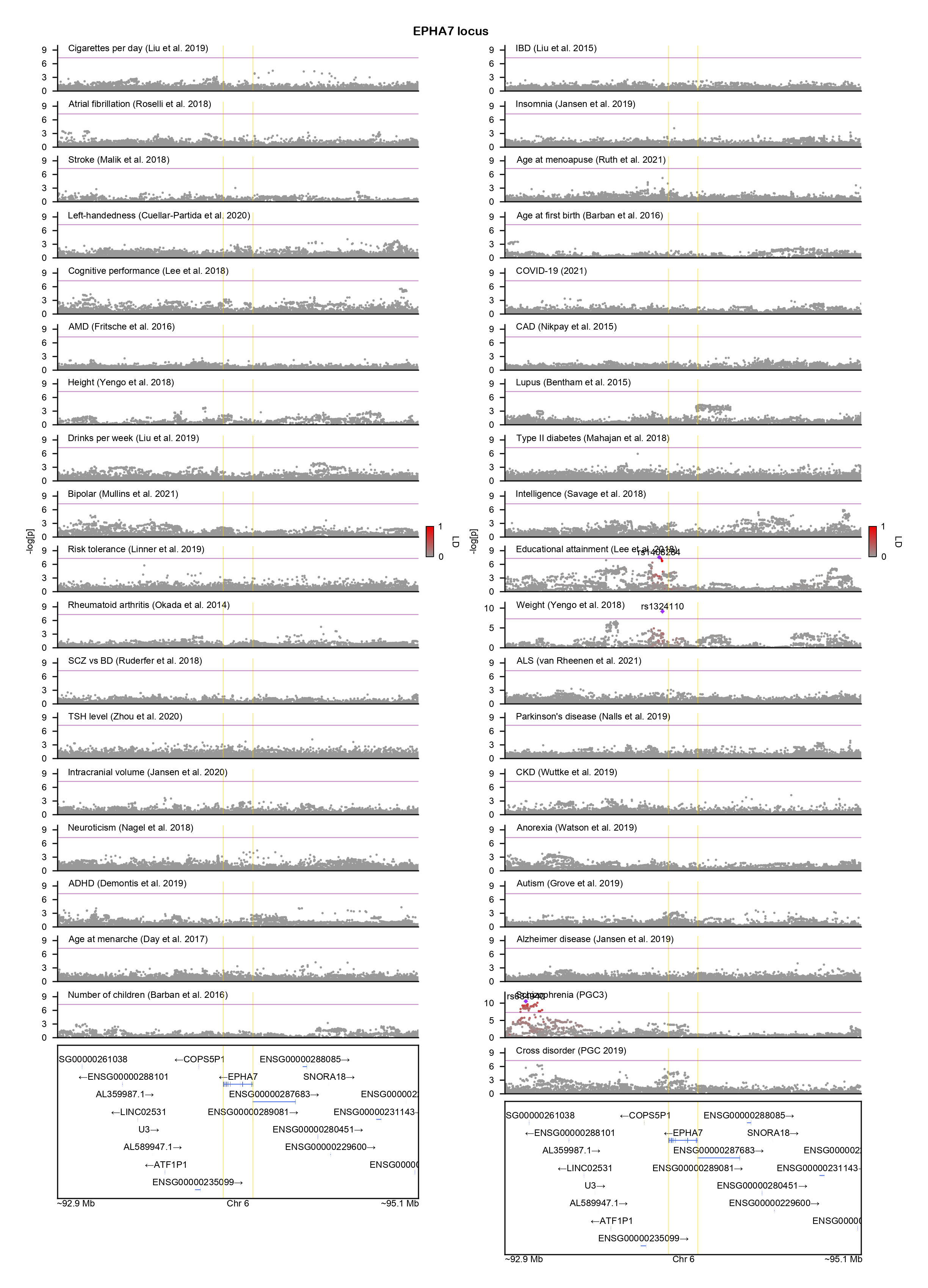

### ESAM-locuszoom.png

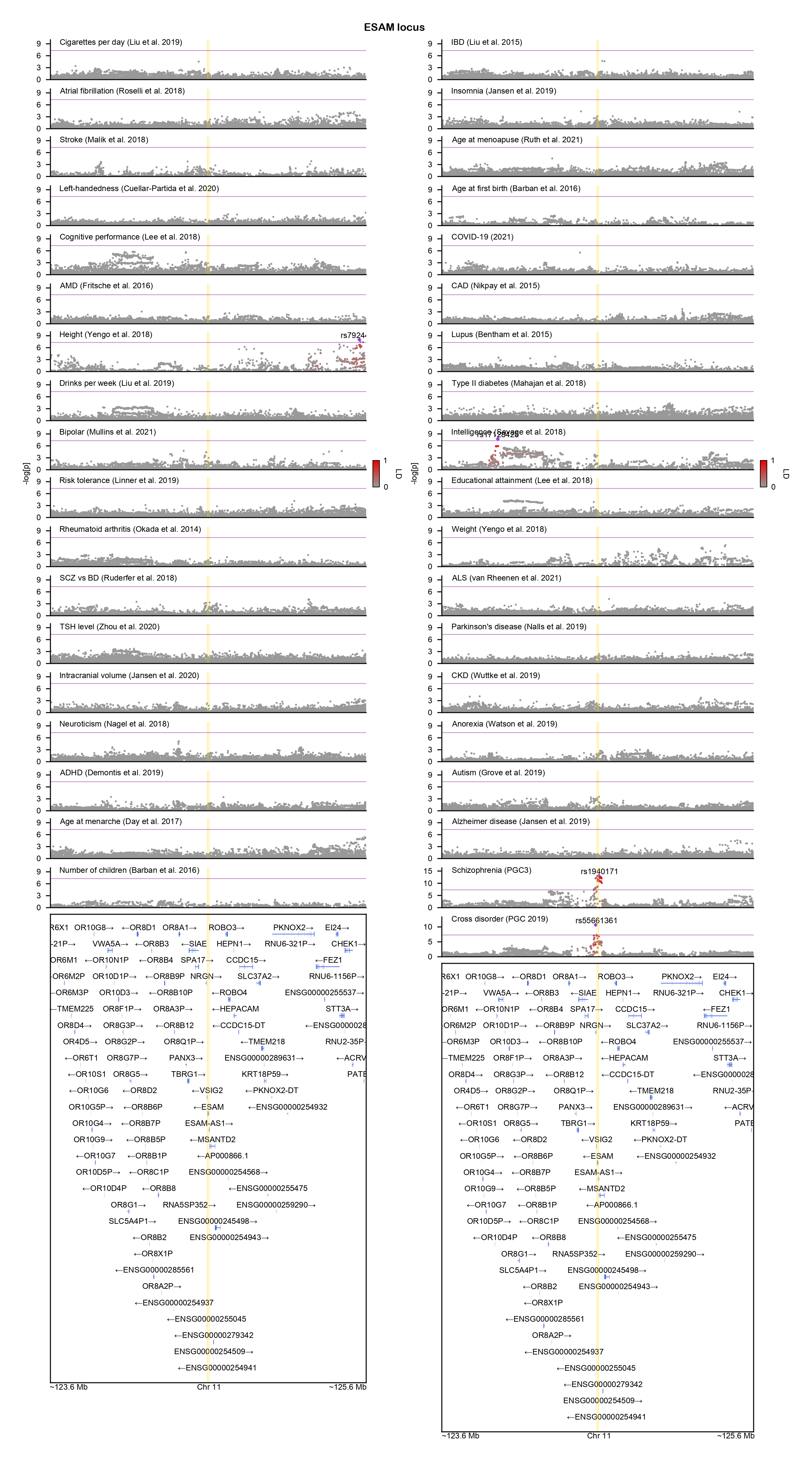

### ETF1-locuszoom.png

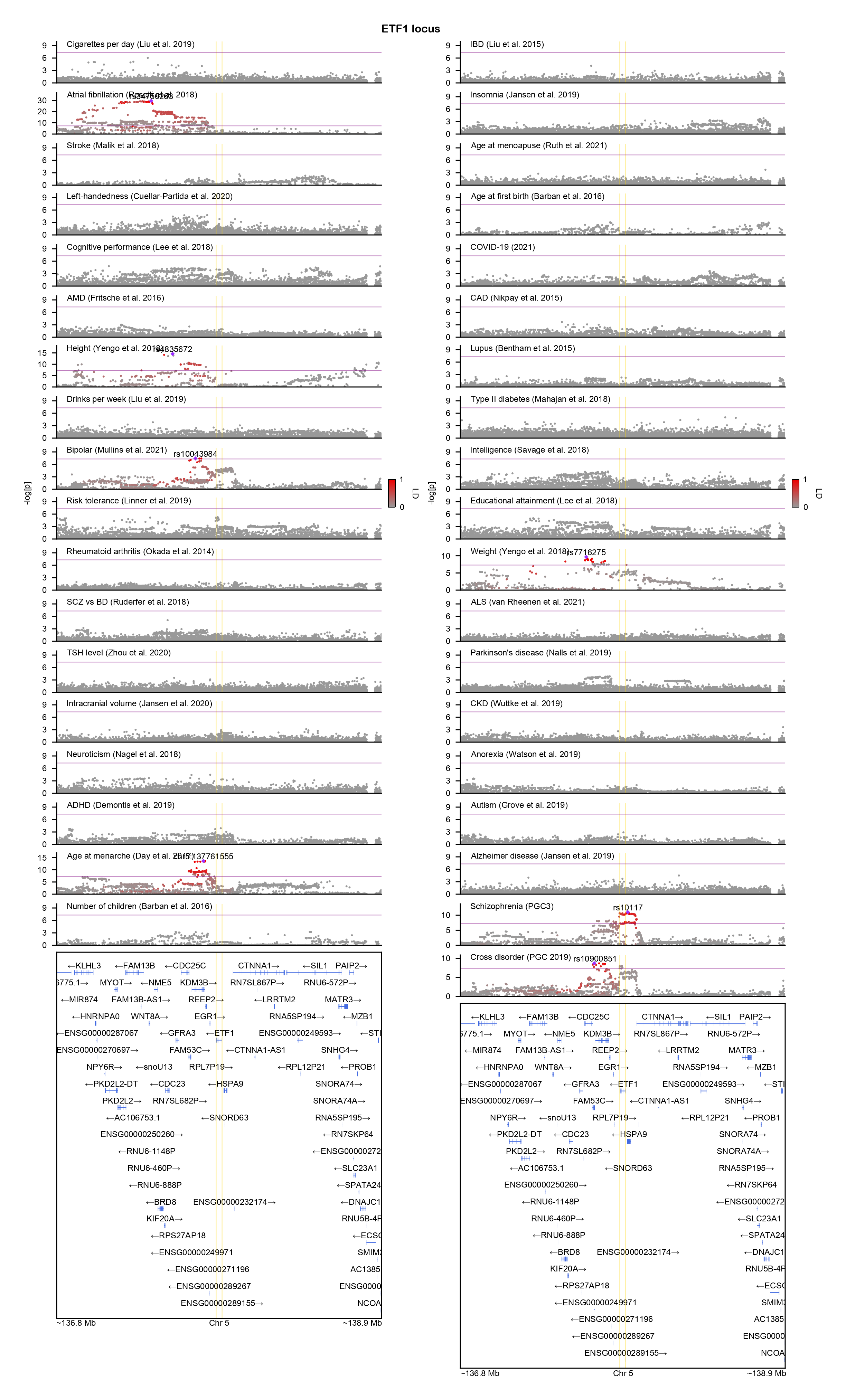

### FAHD2B-locuszoom.png

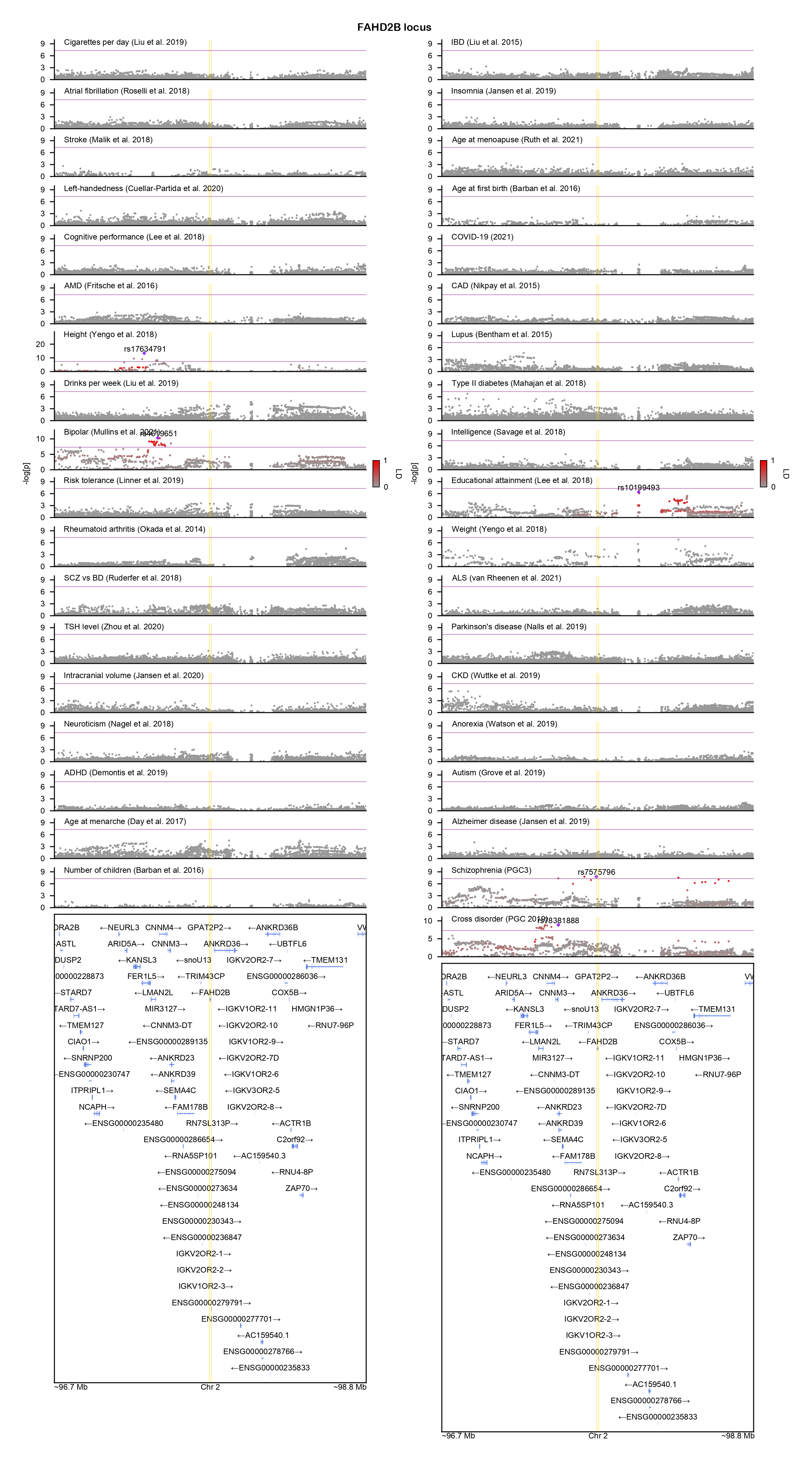

### FHIT-locuszoom.png

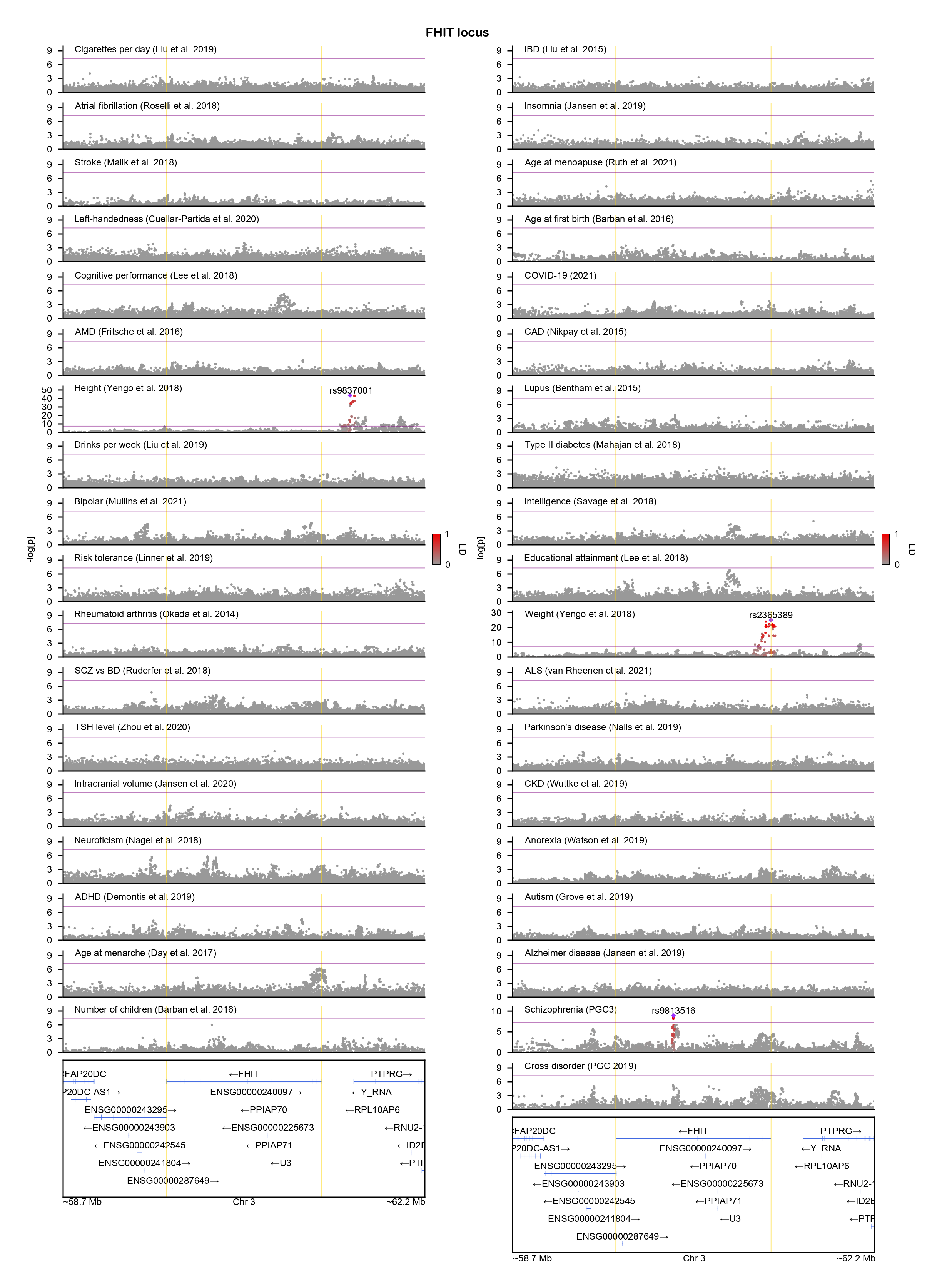

### FOXP1-locuszoom.png

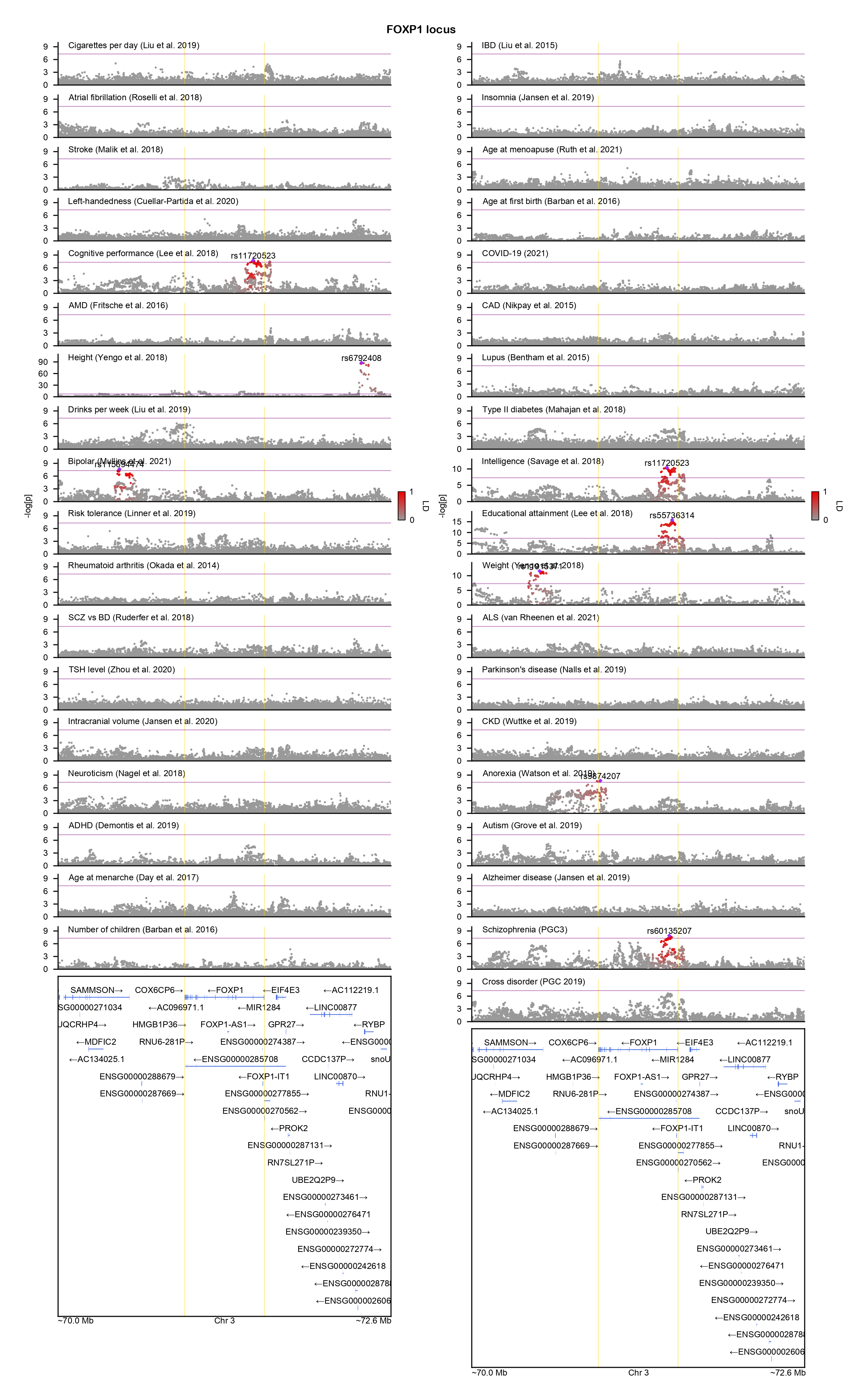

### FUT9-locuszoom.png

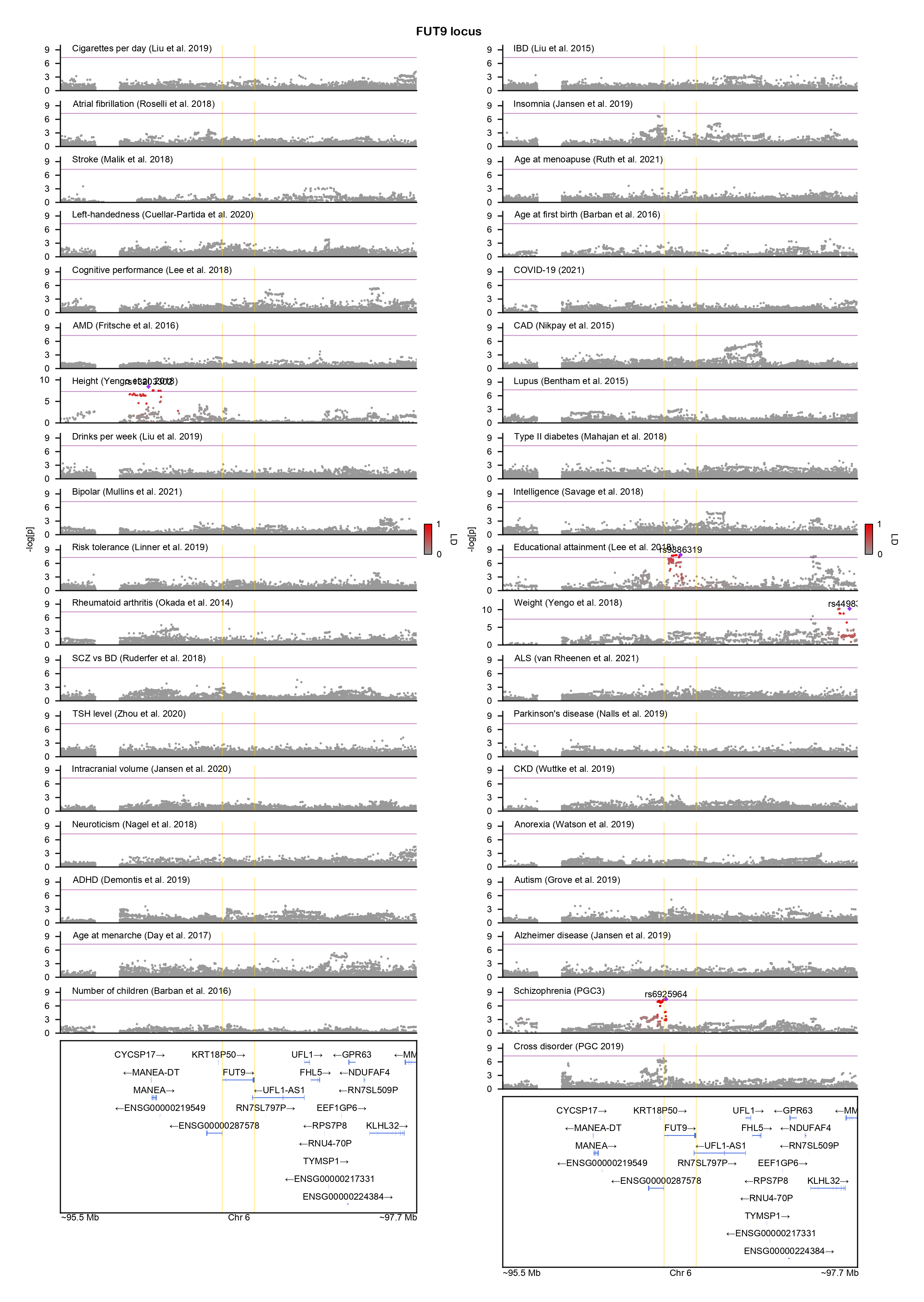

### GALNT10-locuszoom.png

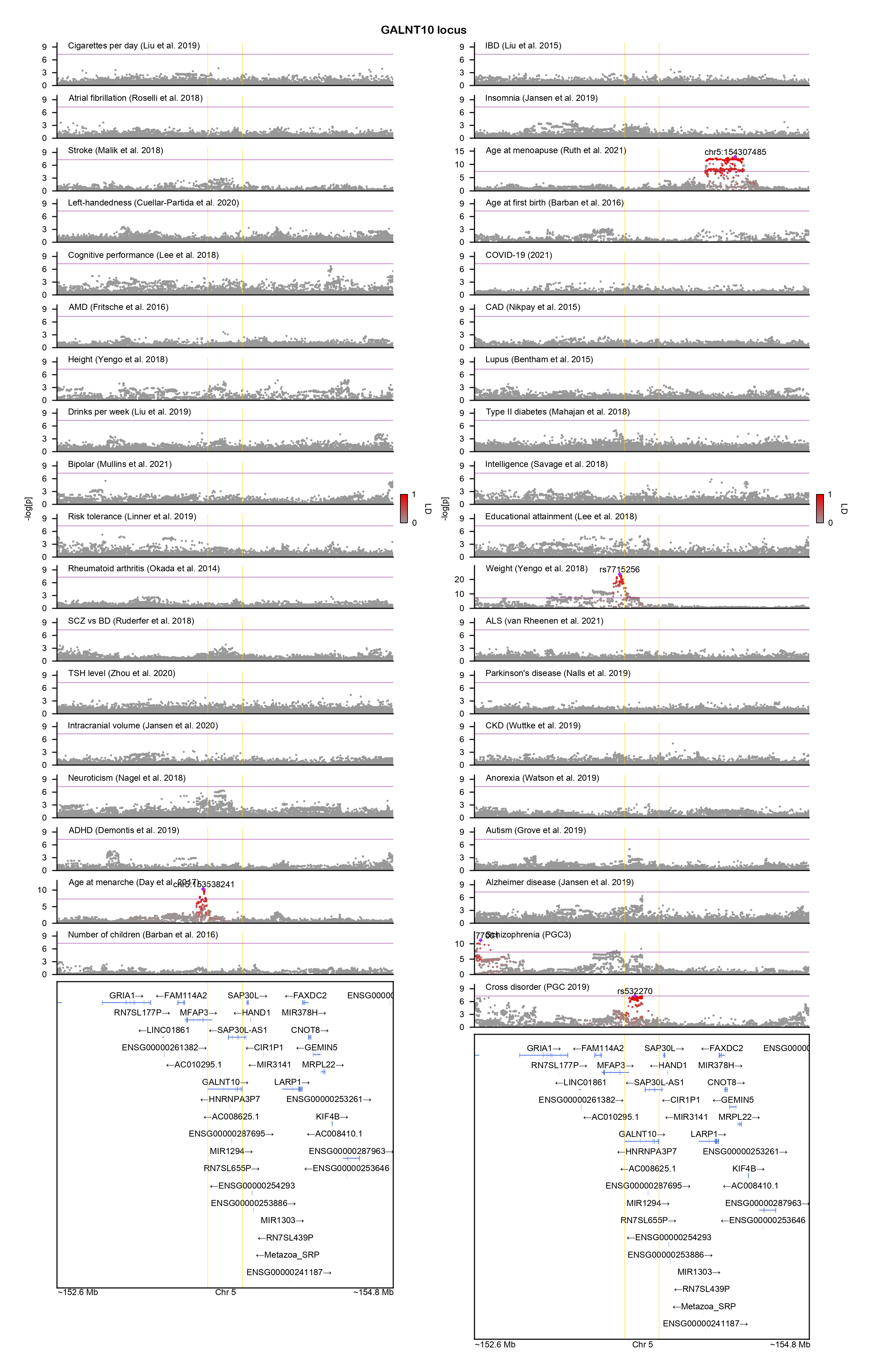

### GALNT17-locuszoom.png

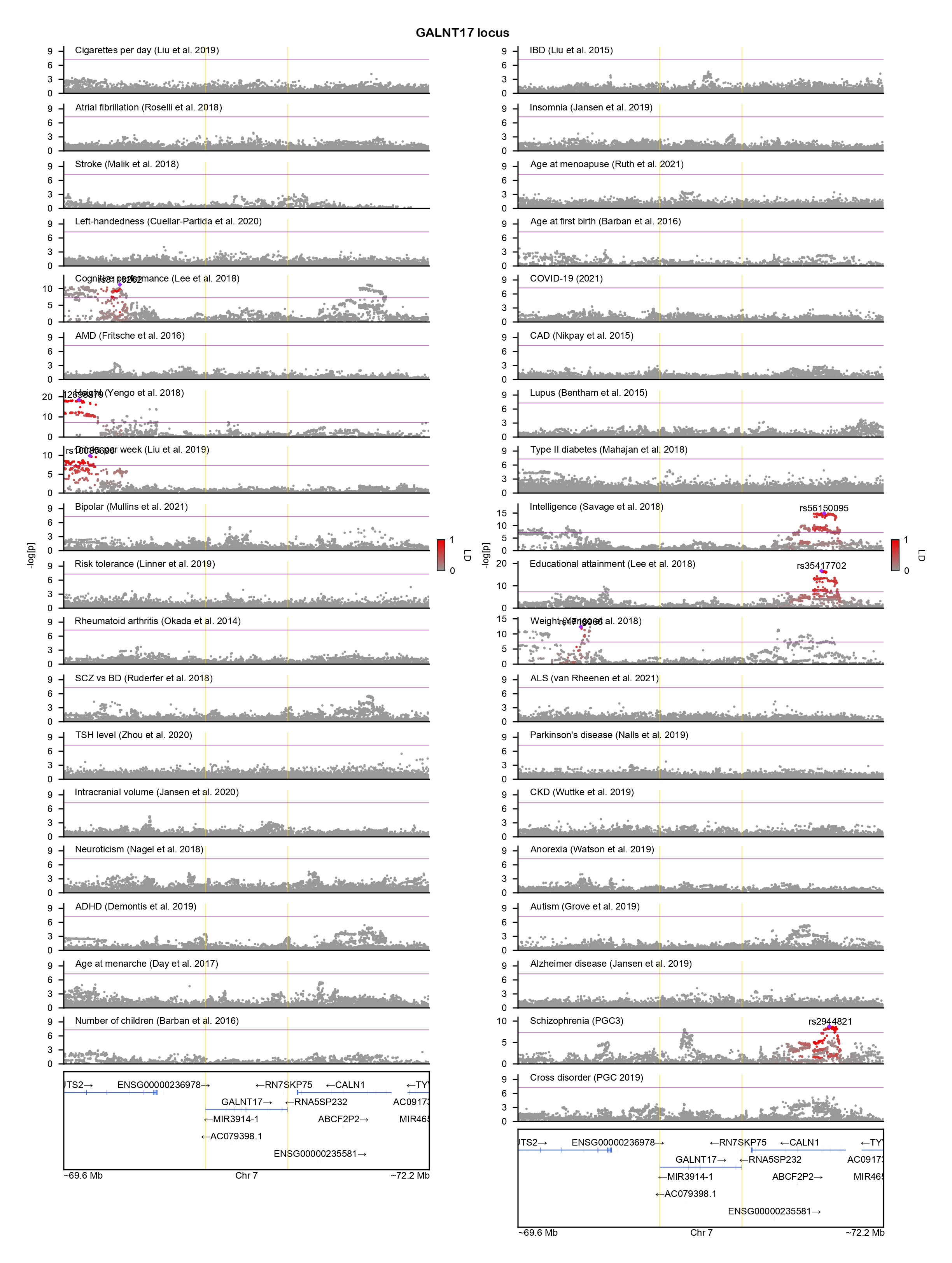

### GSDME-locuszoom.png

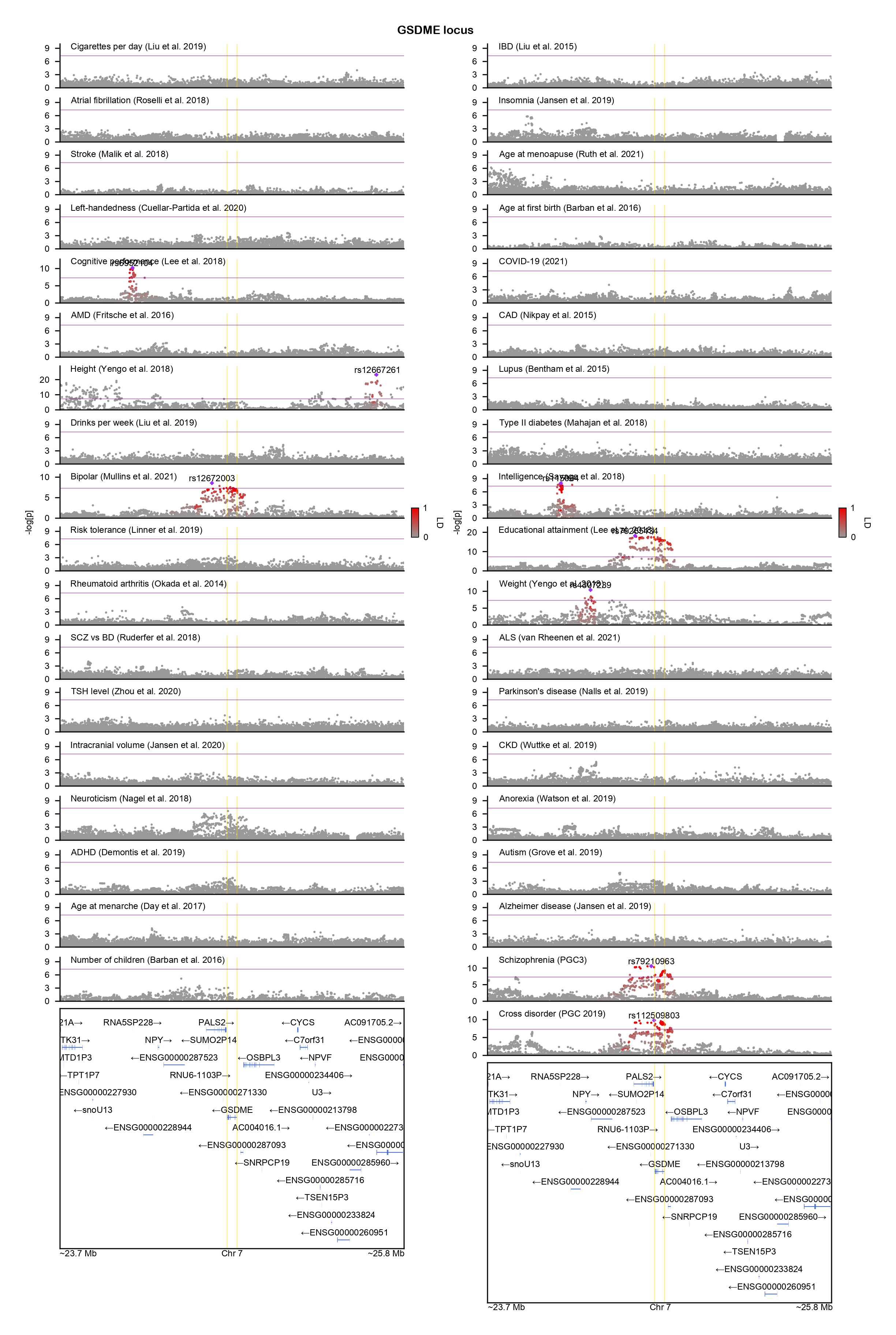

### GTDC1-locuszoom.png

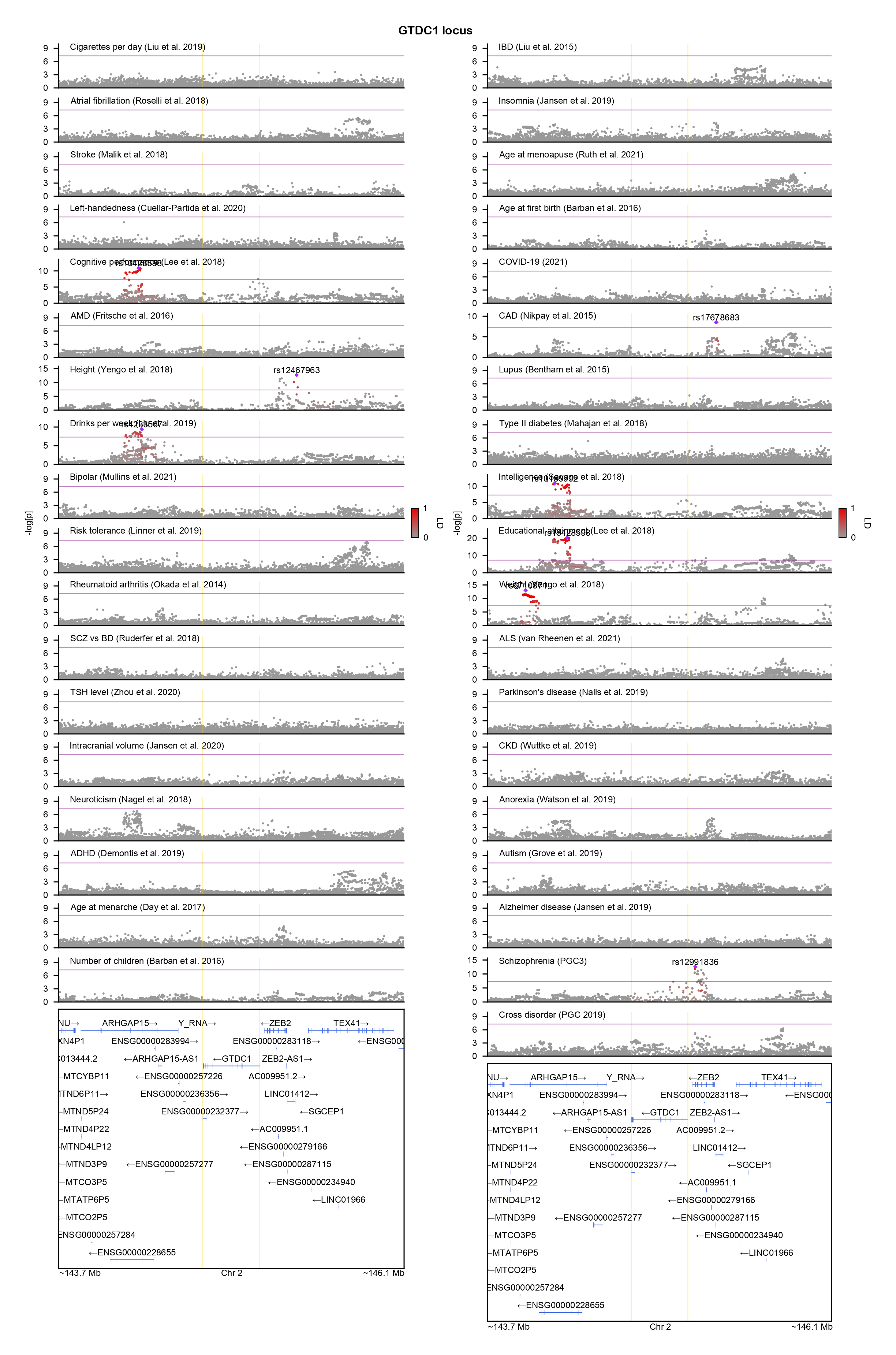

### HNF1A-locuszoom.png

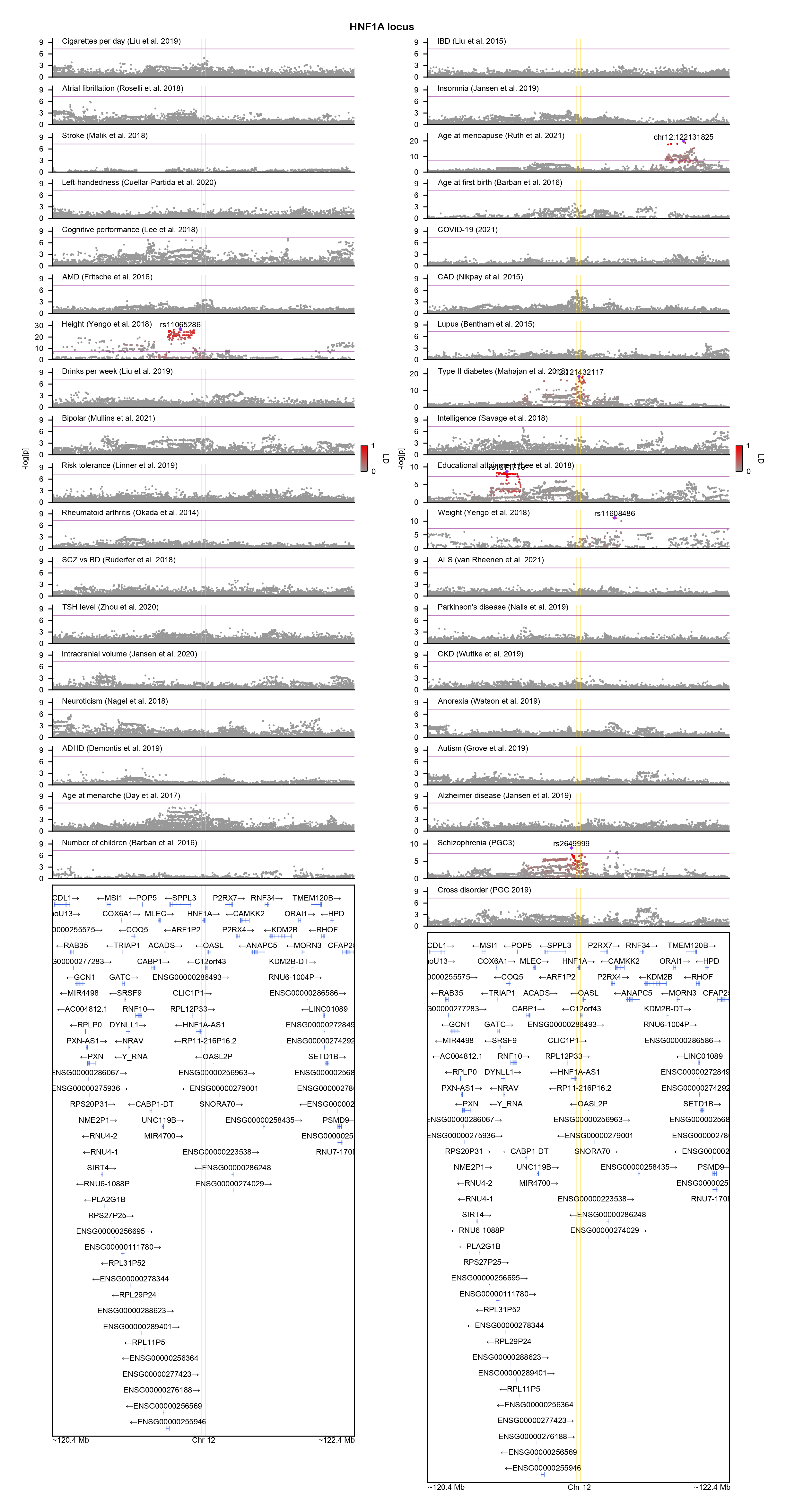

### IGSF9B-locuszoom.png

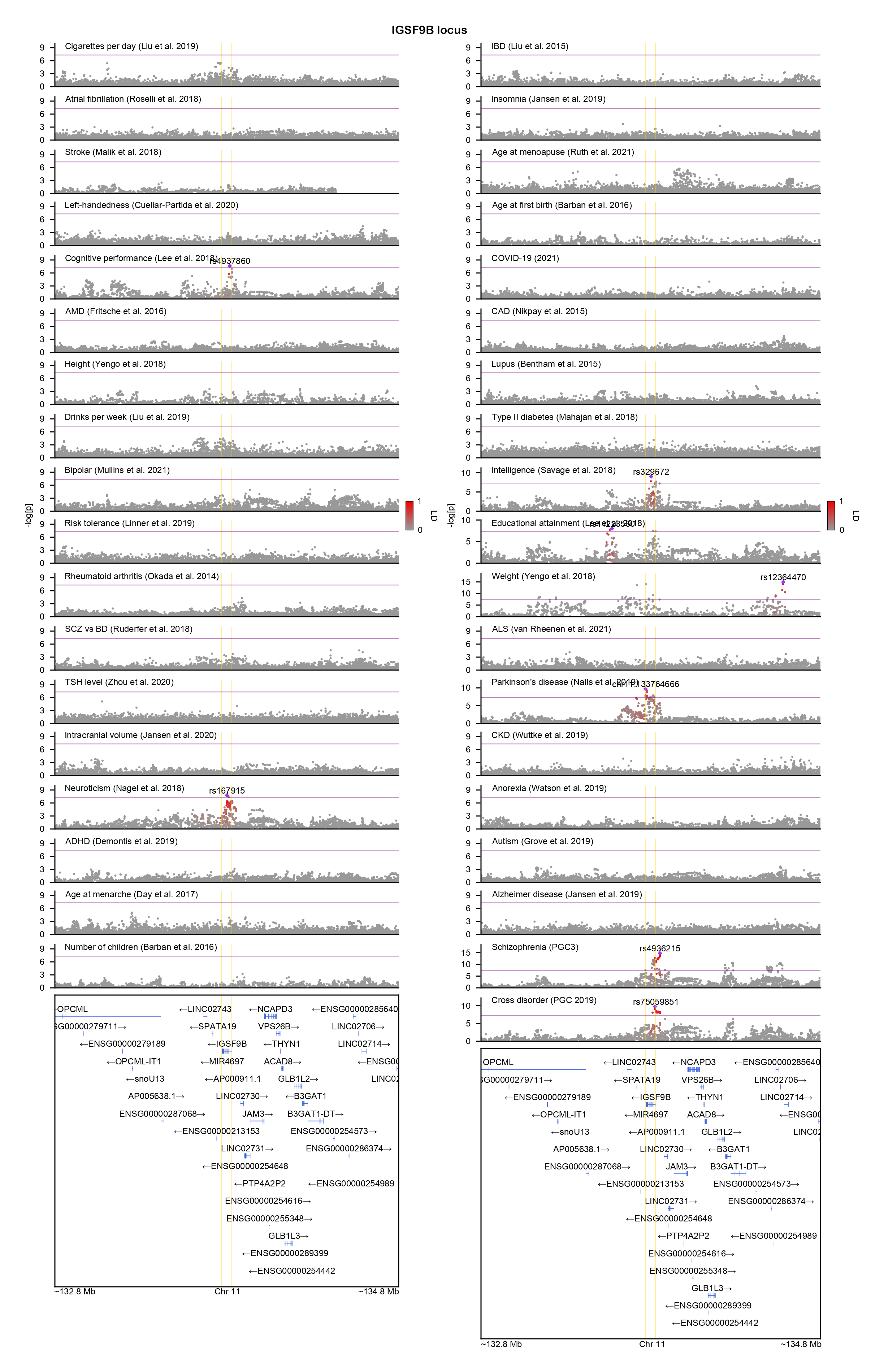

### INPP4B-locuszoom.png

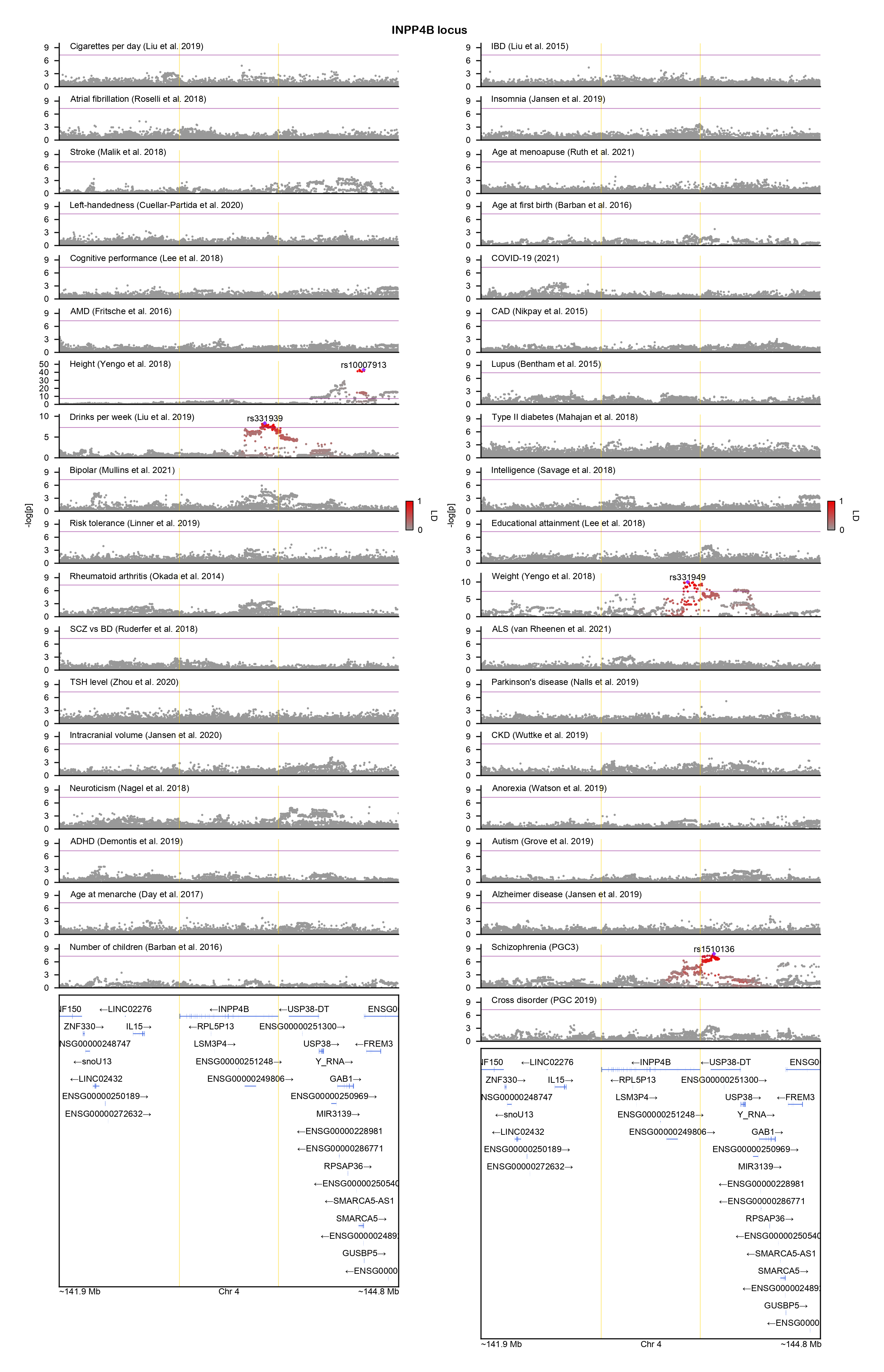

### KCNG2-locuszoom.png

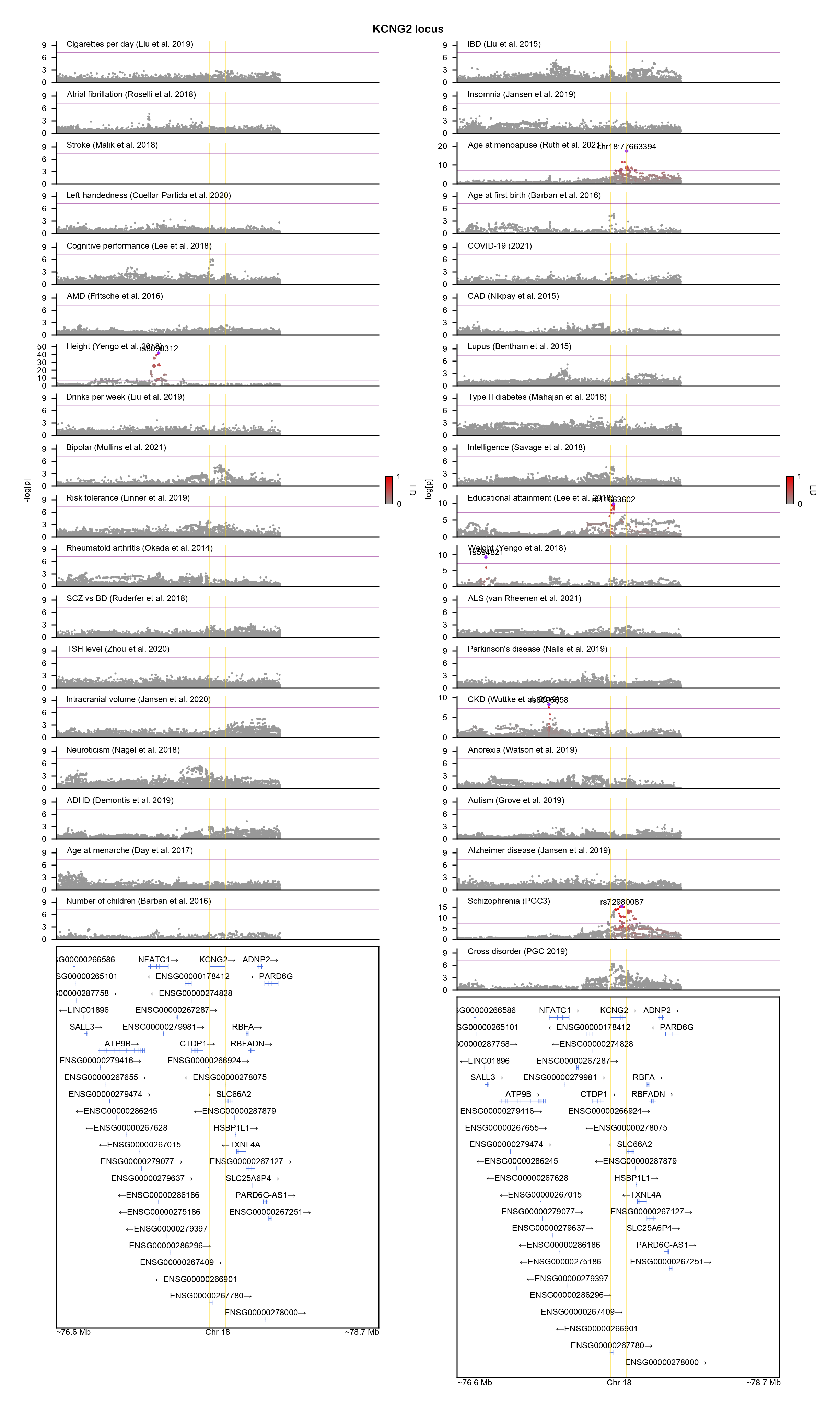

### KDM4A-locuszoom.png

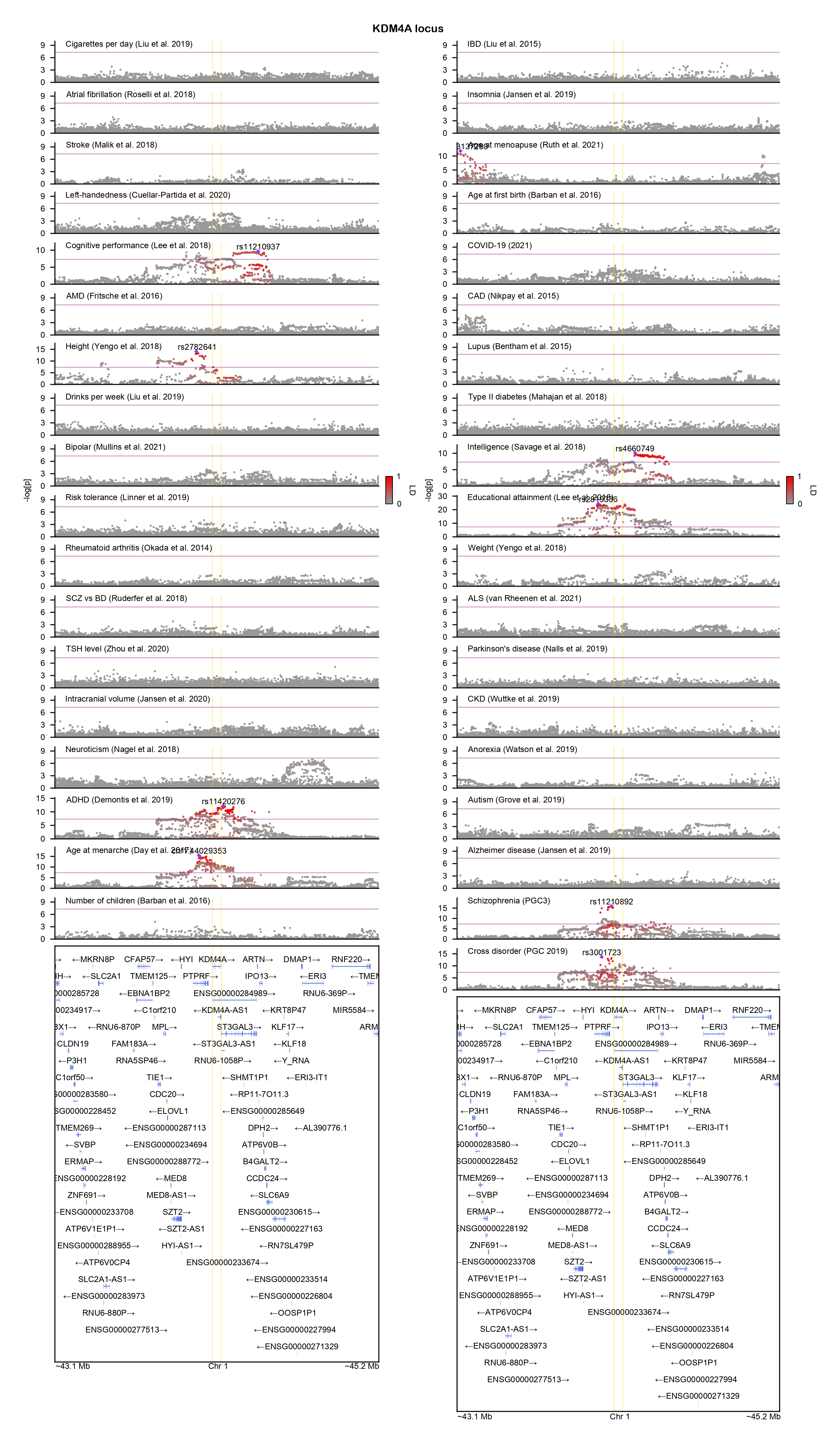

### KLHL29-locuszoom.png

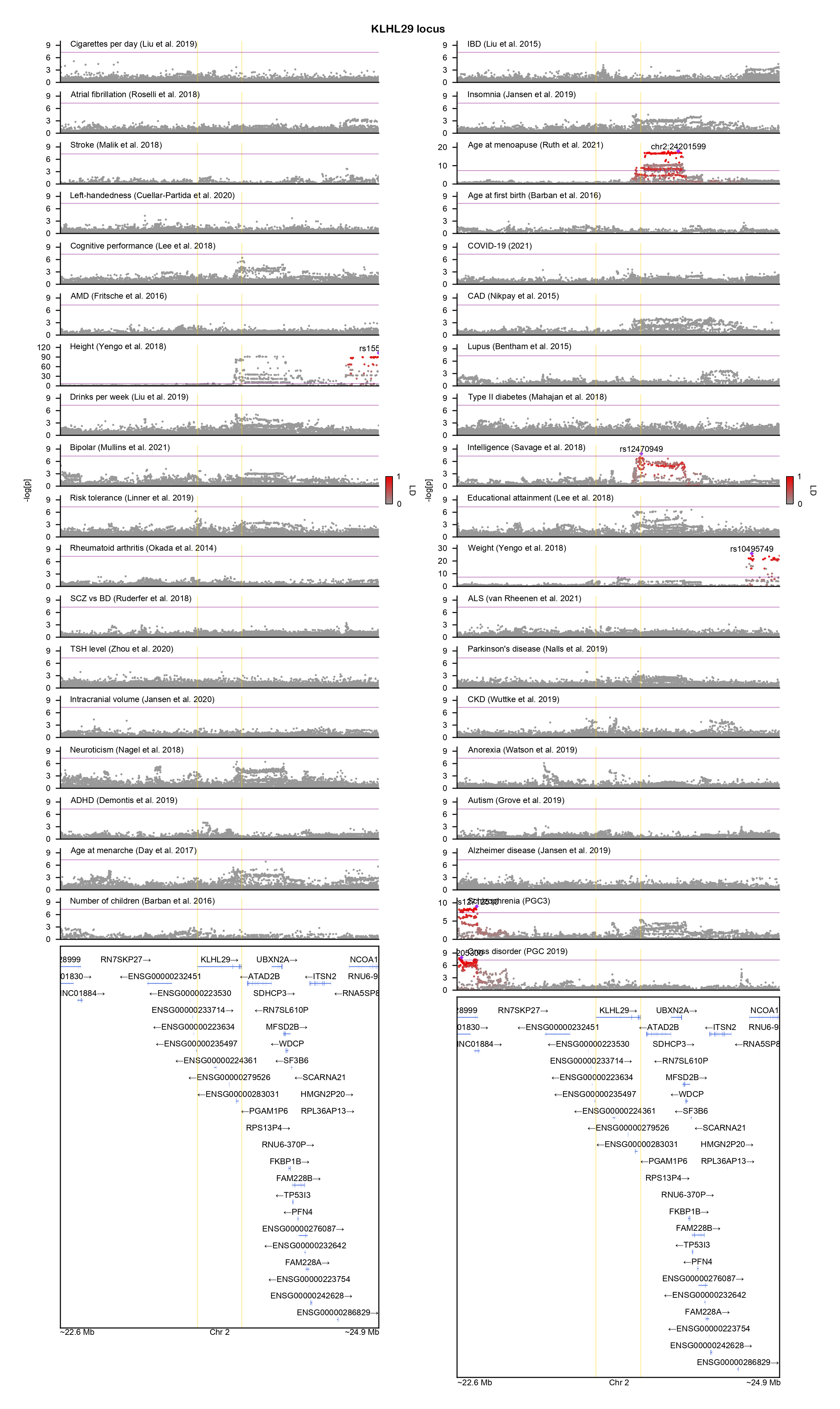

### L3MBTL2-locuszoom.png

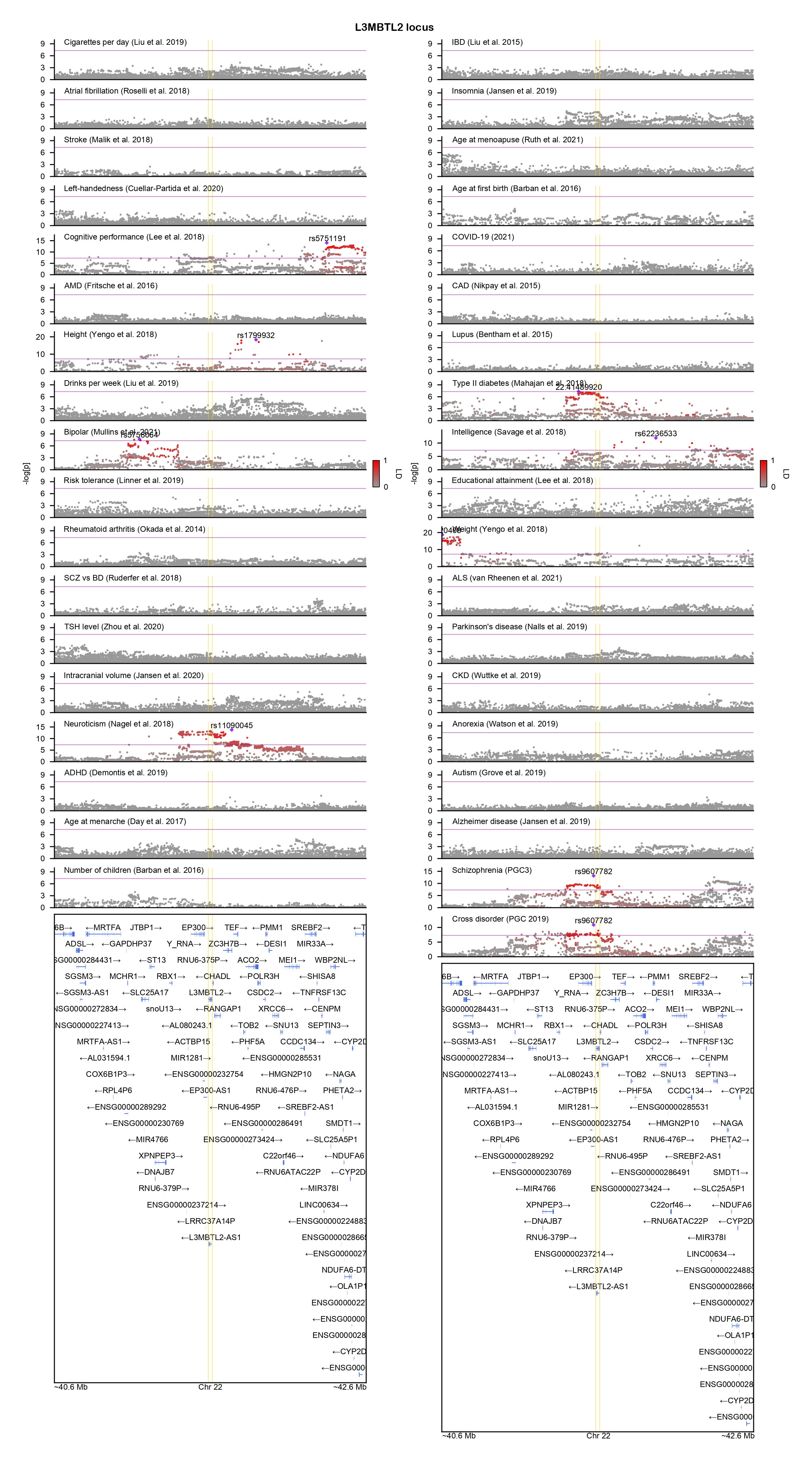

### LRRC4-locuszoom.png

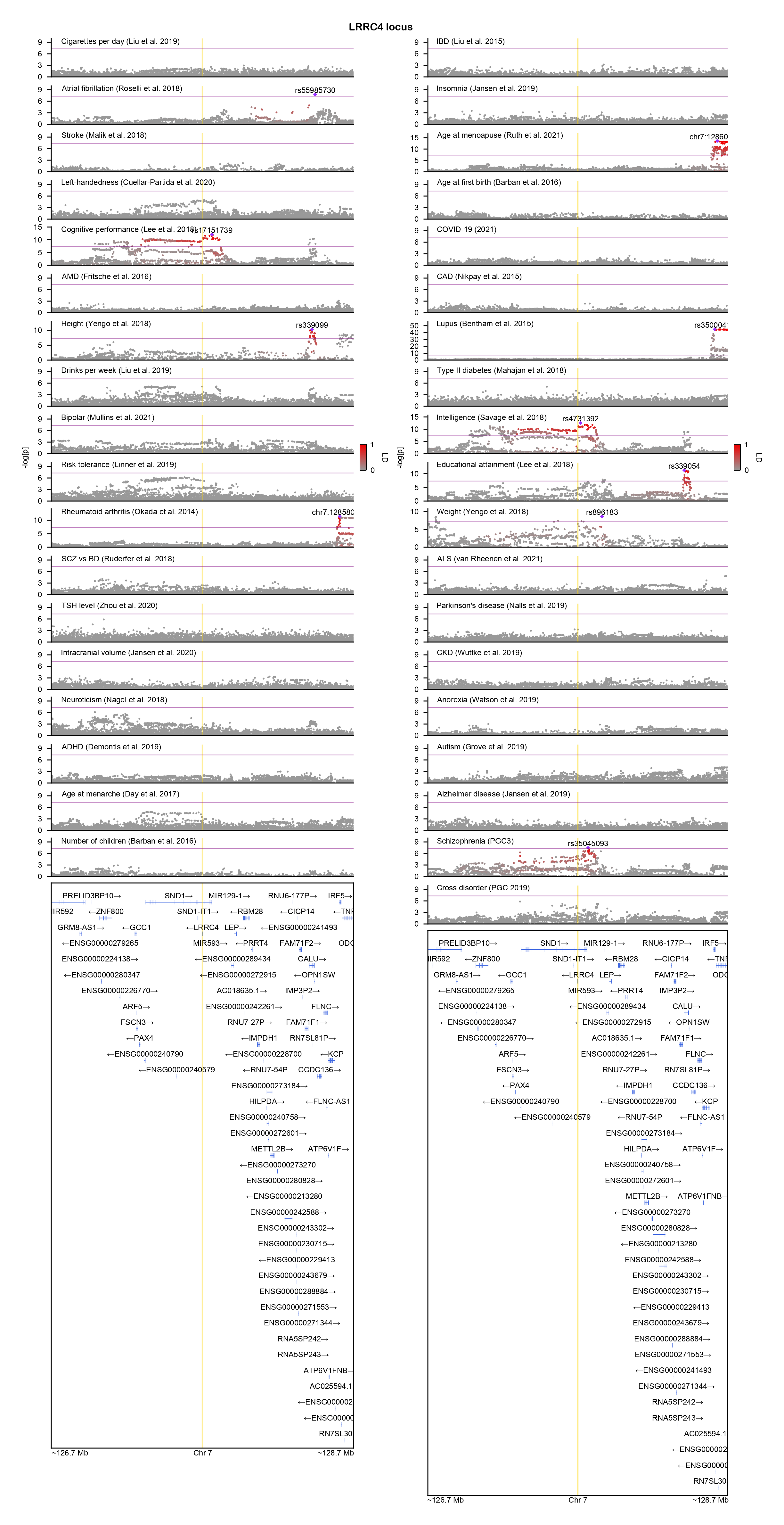

### LRRIQ3-locuszoom.png

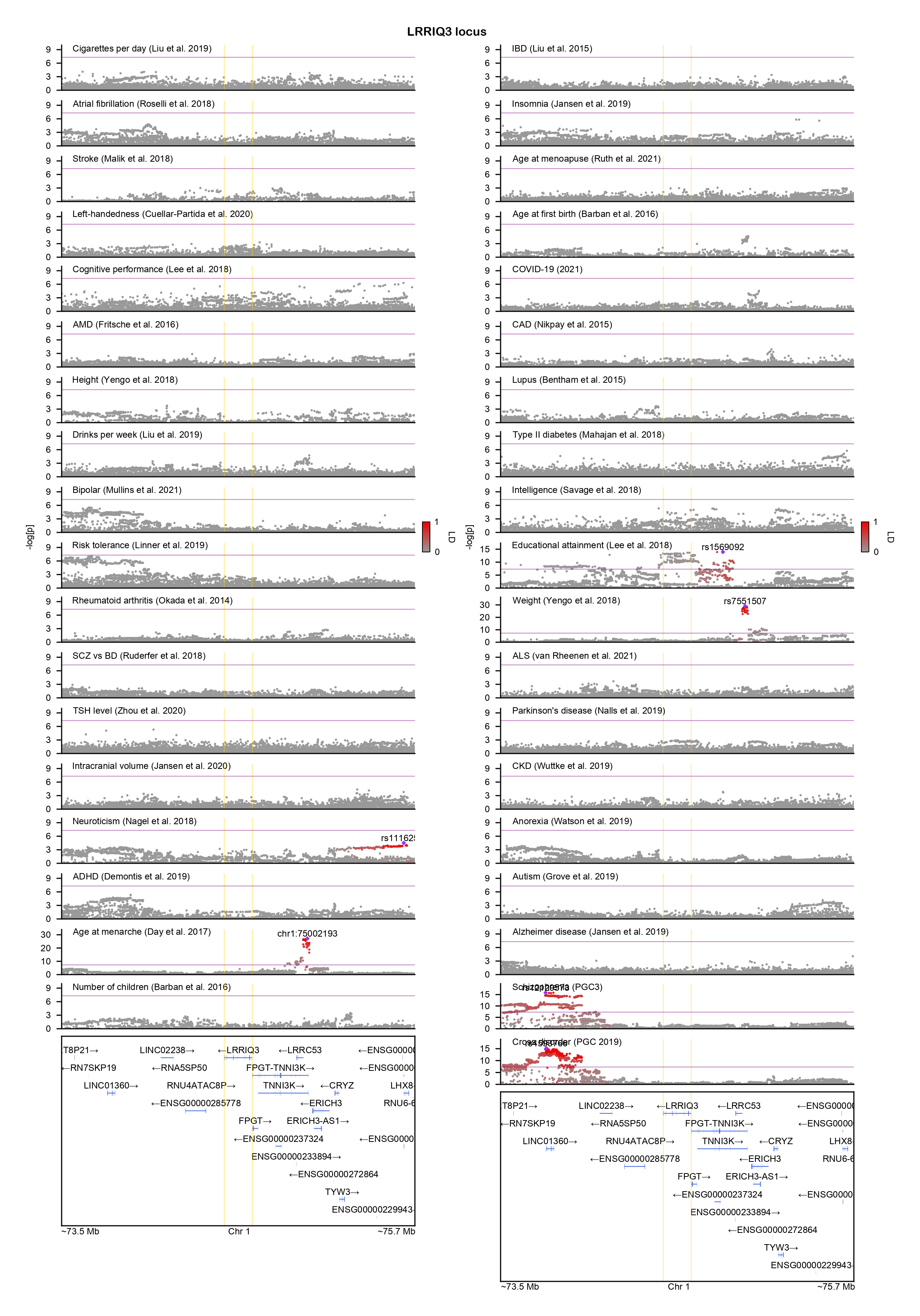

### LUZP2-locuszoom.png

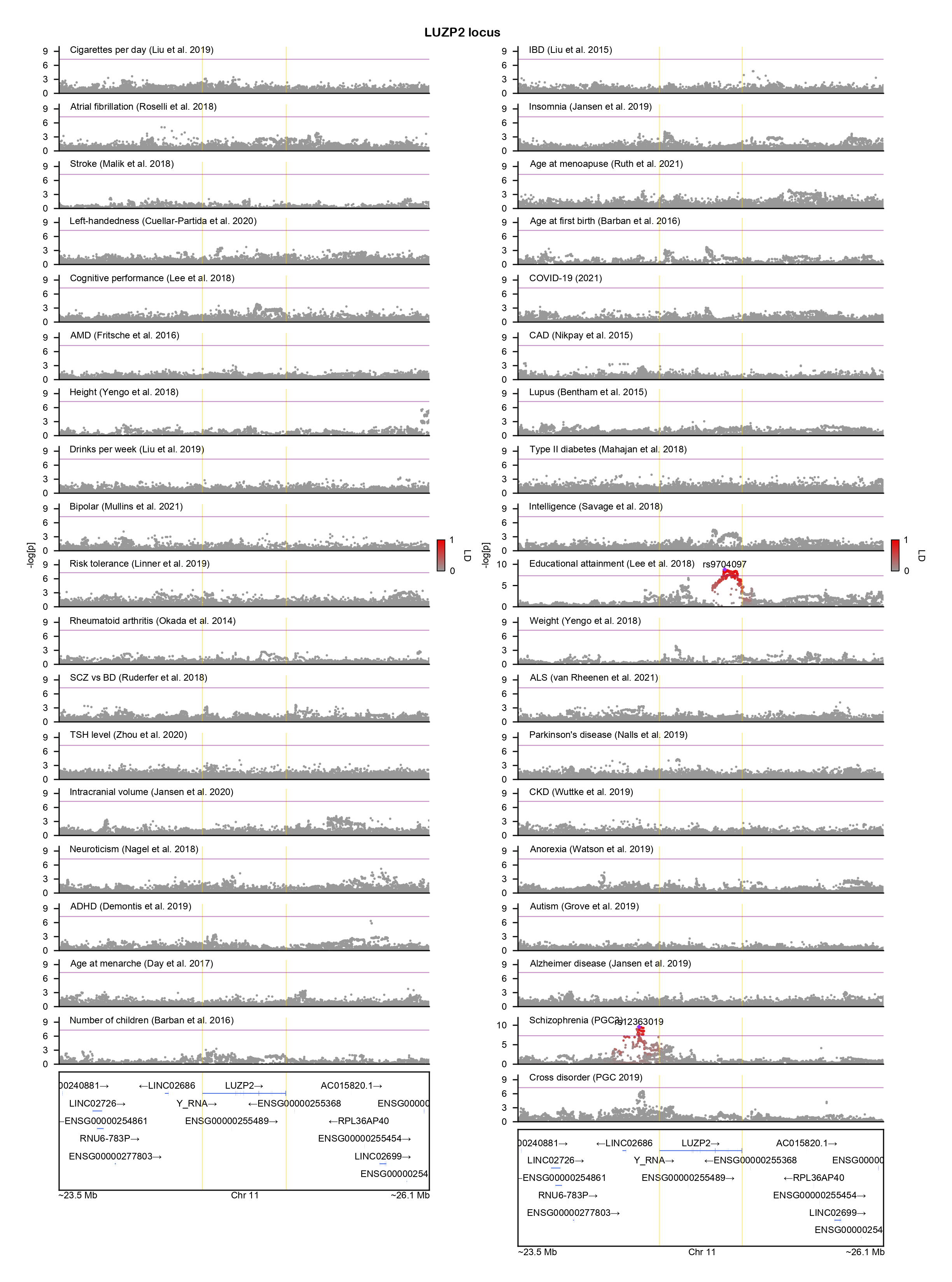

### MAN2A1-locuszoom.png

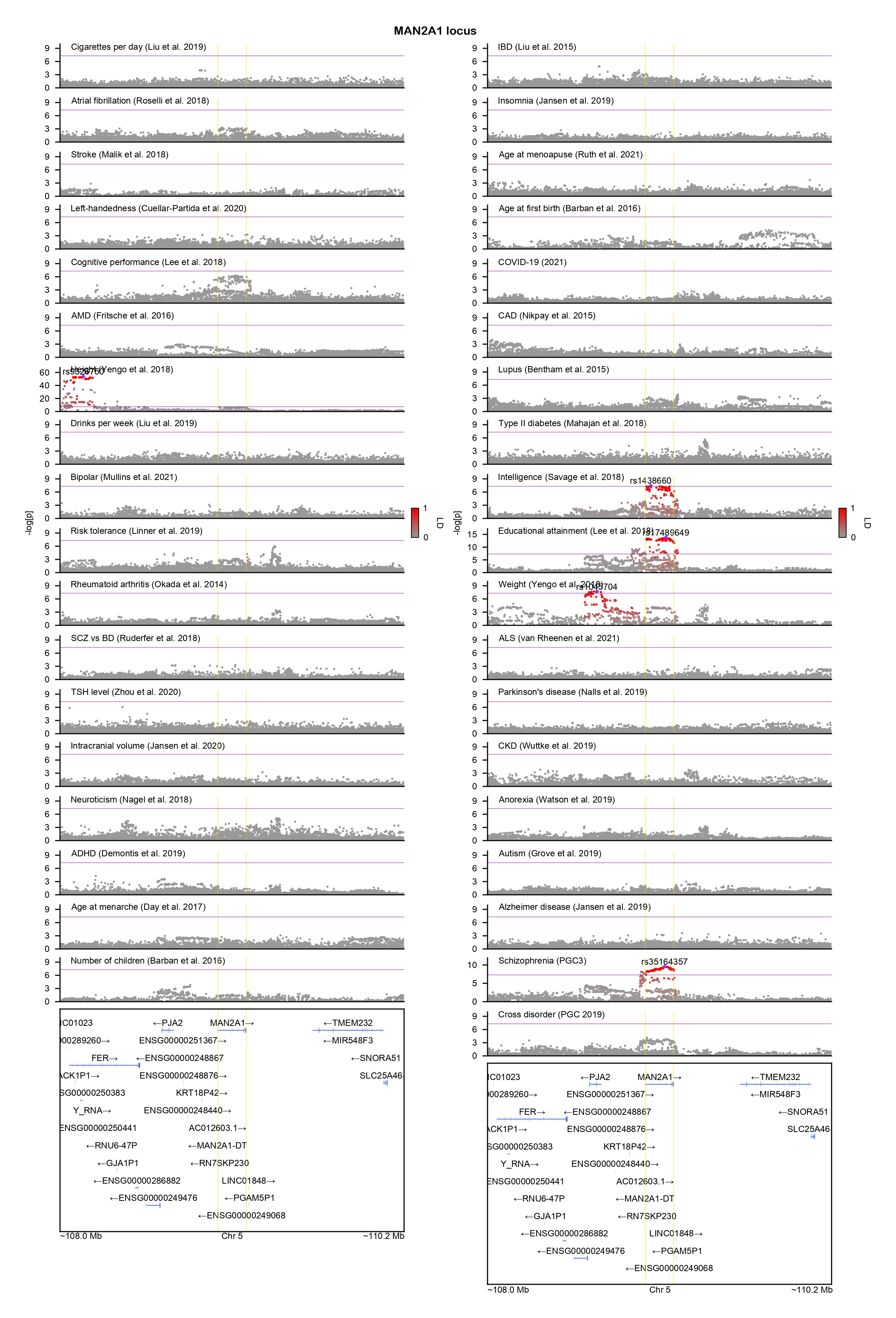

### MATN1-locuszoom.png

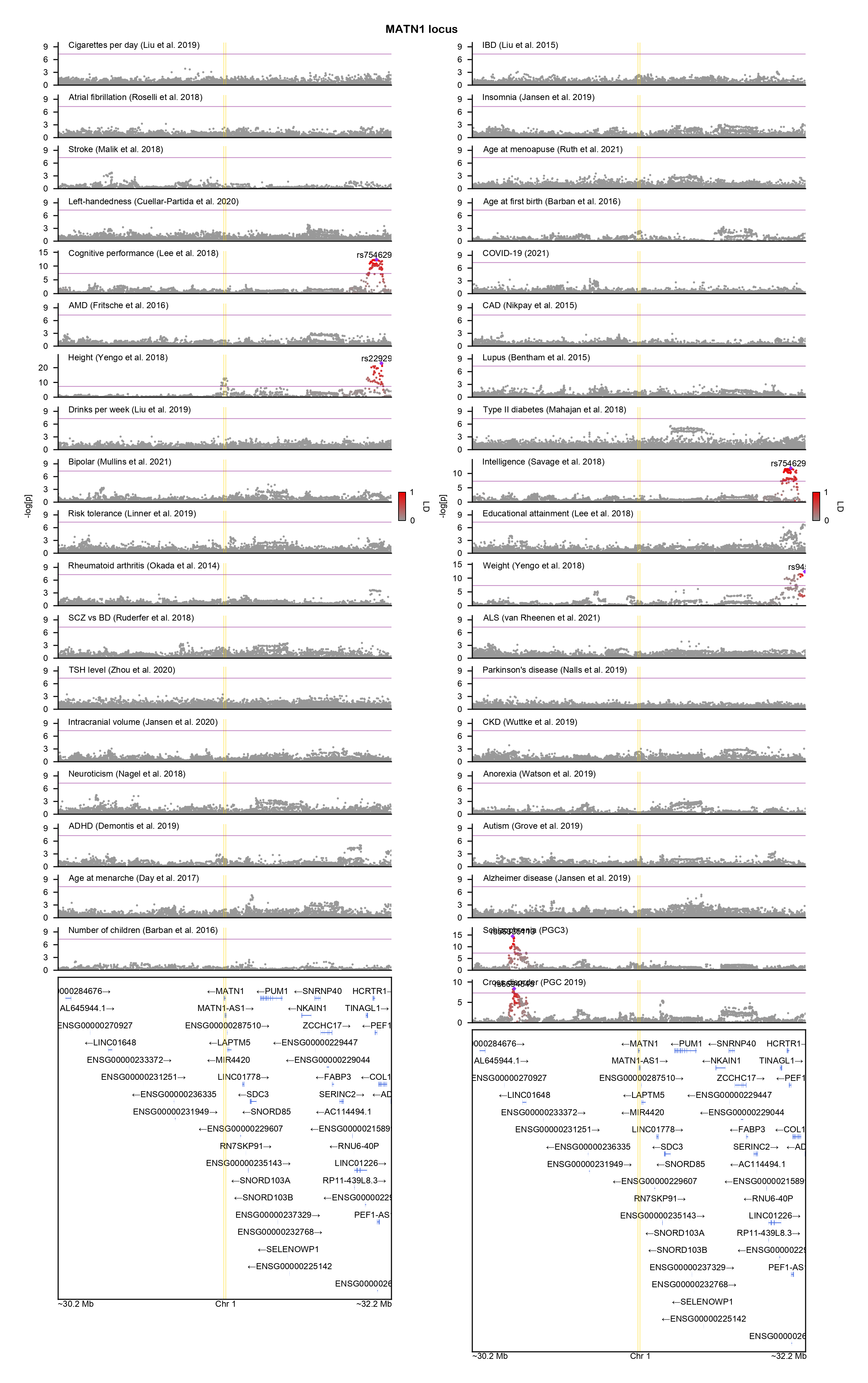

### MED27-locuszoom.png

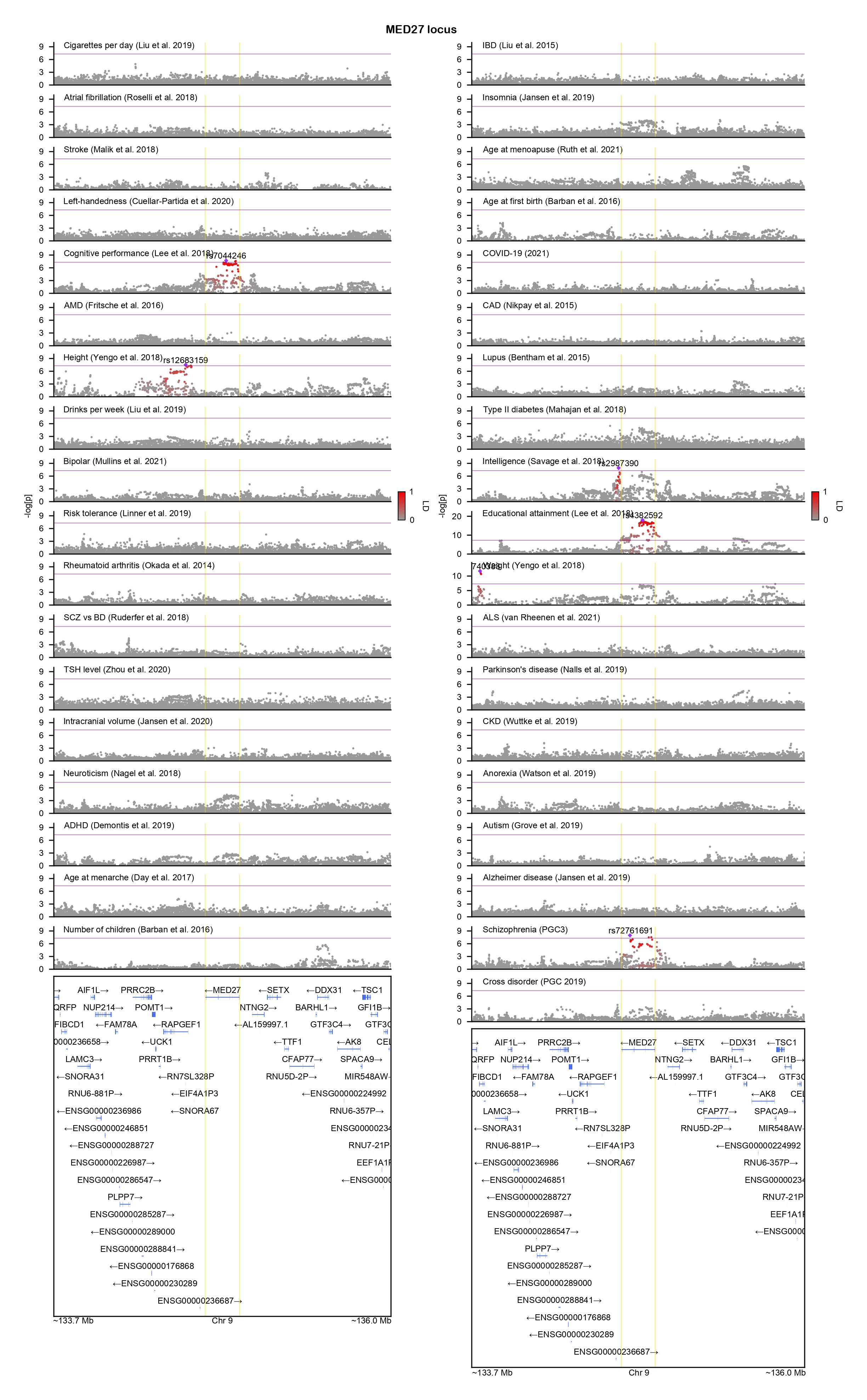

### METTL15-locuszoom.png

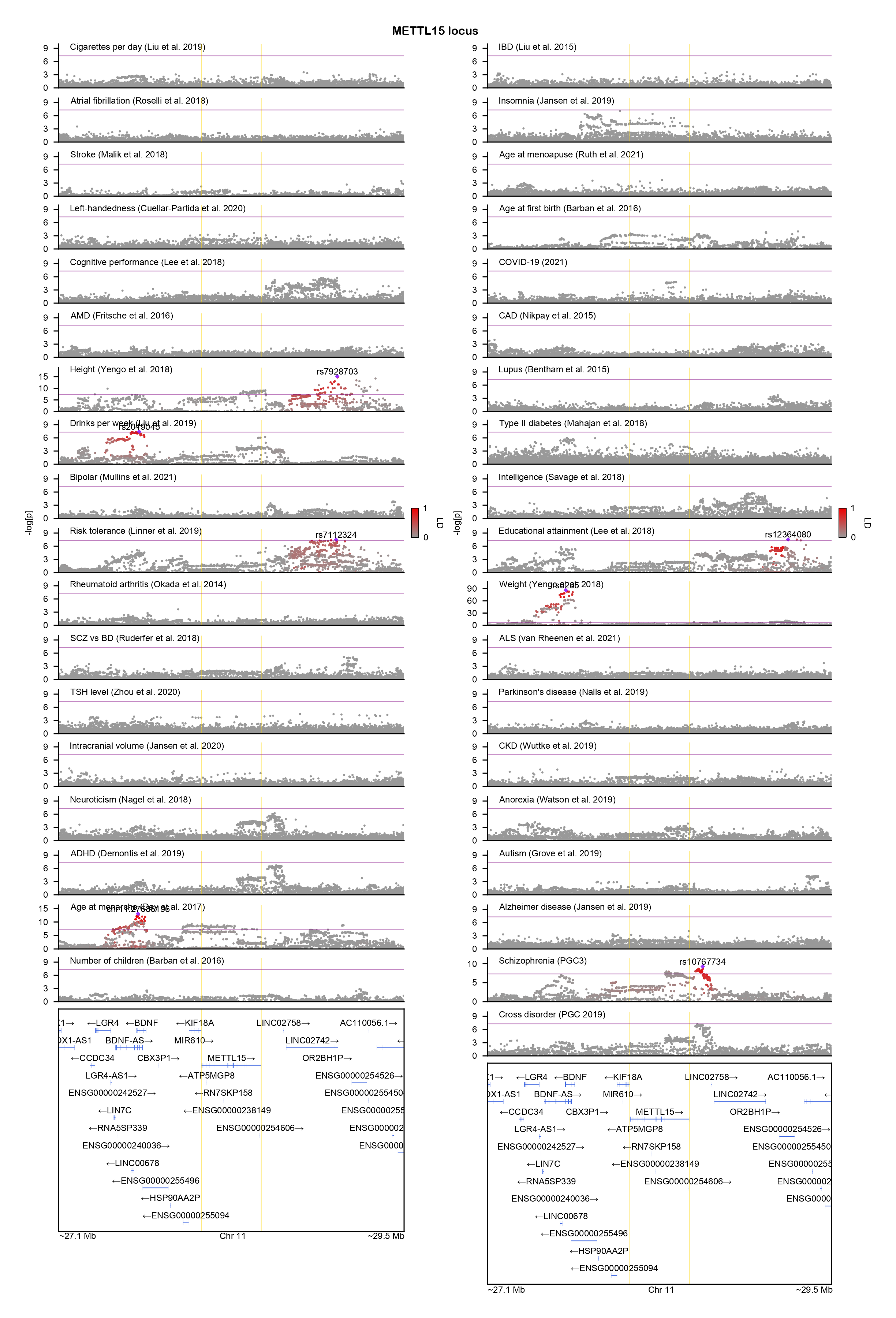

### MMP16-locuszoom.png

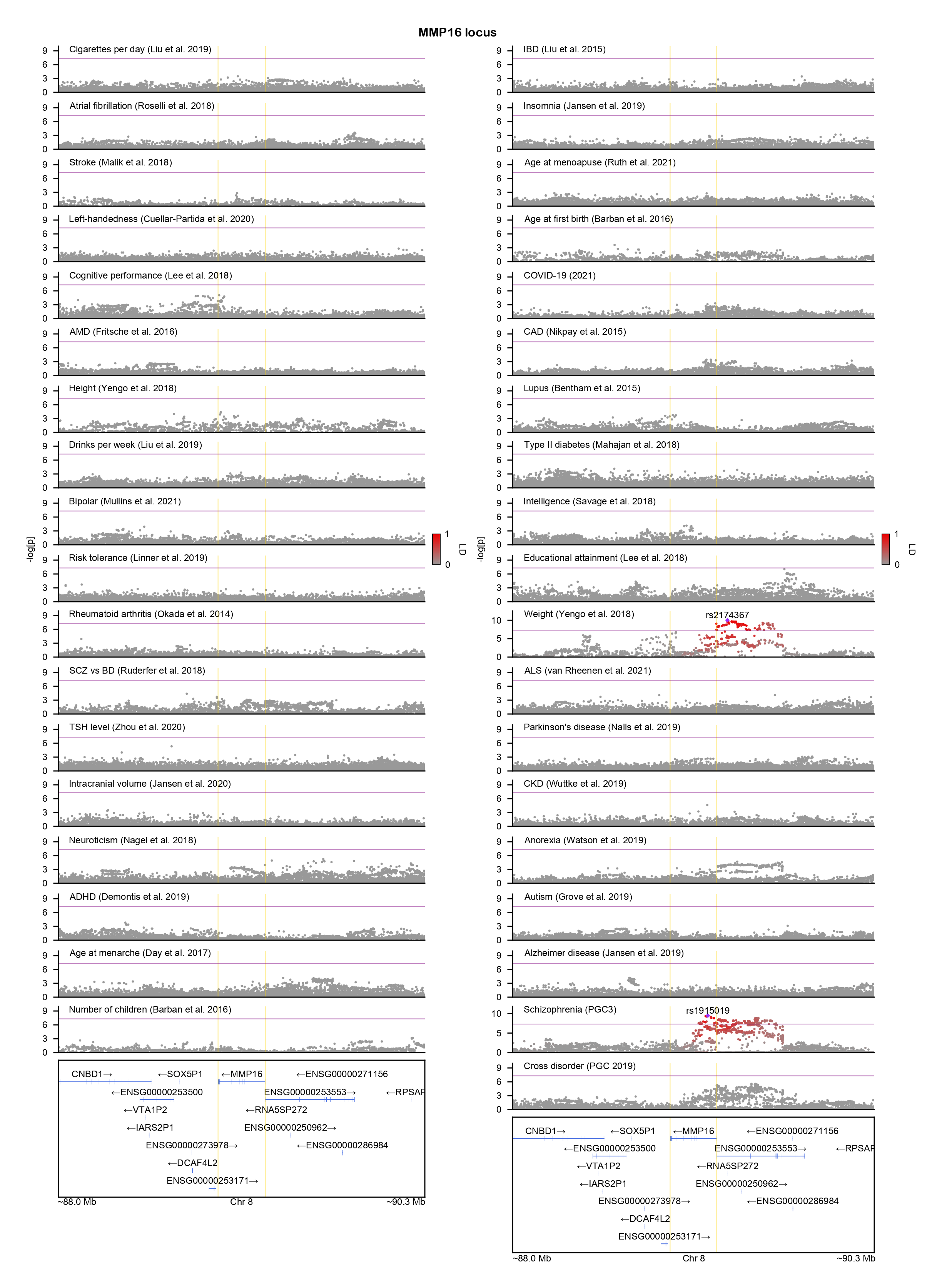

### MRPL33-locuszoom.png

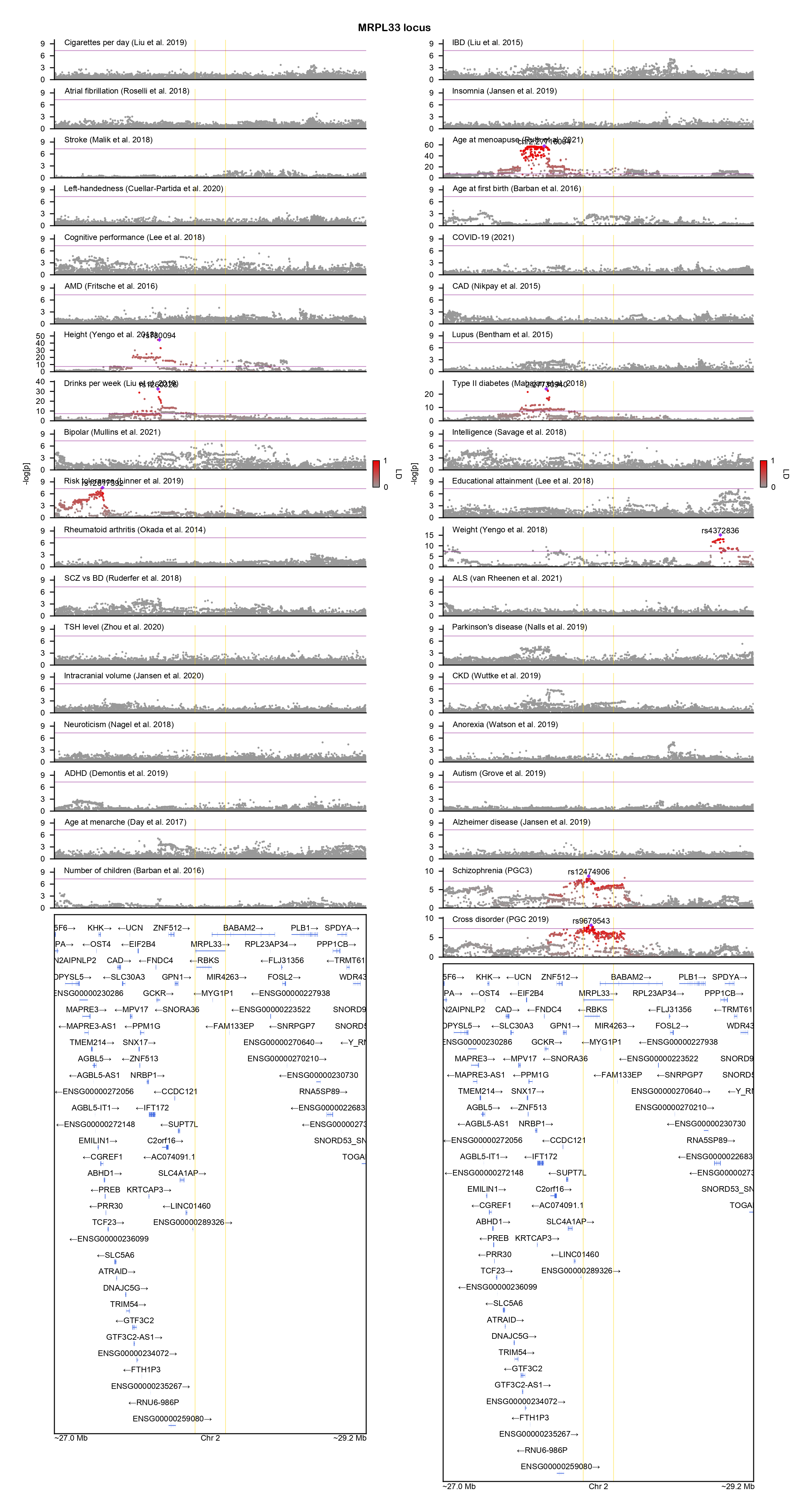

### MRPS30-locuszoom.png

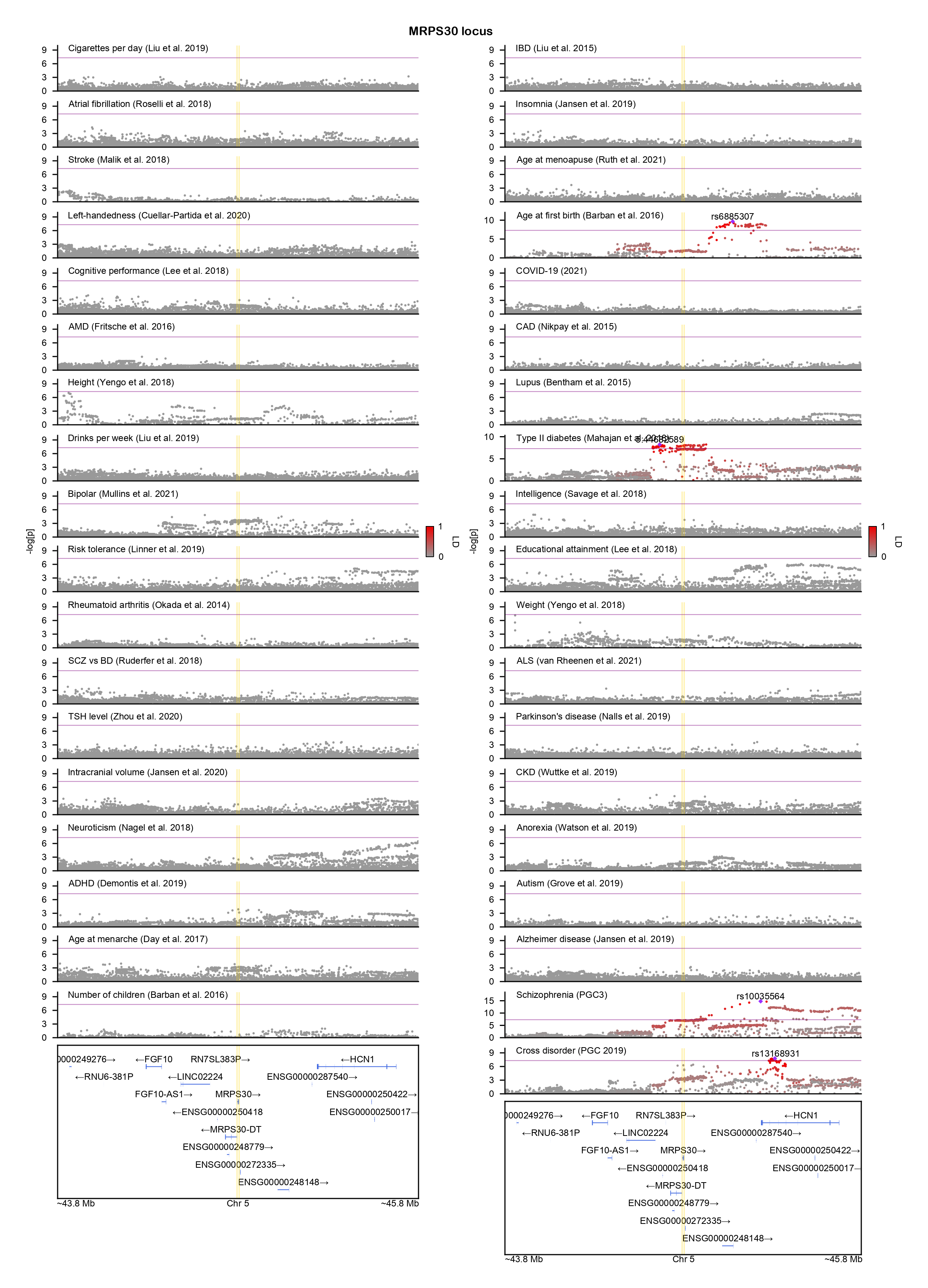

### MYBPC3-locuszoom.png

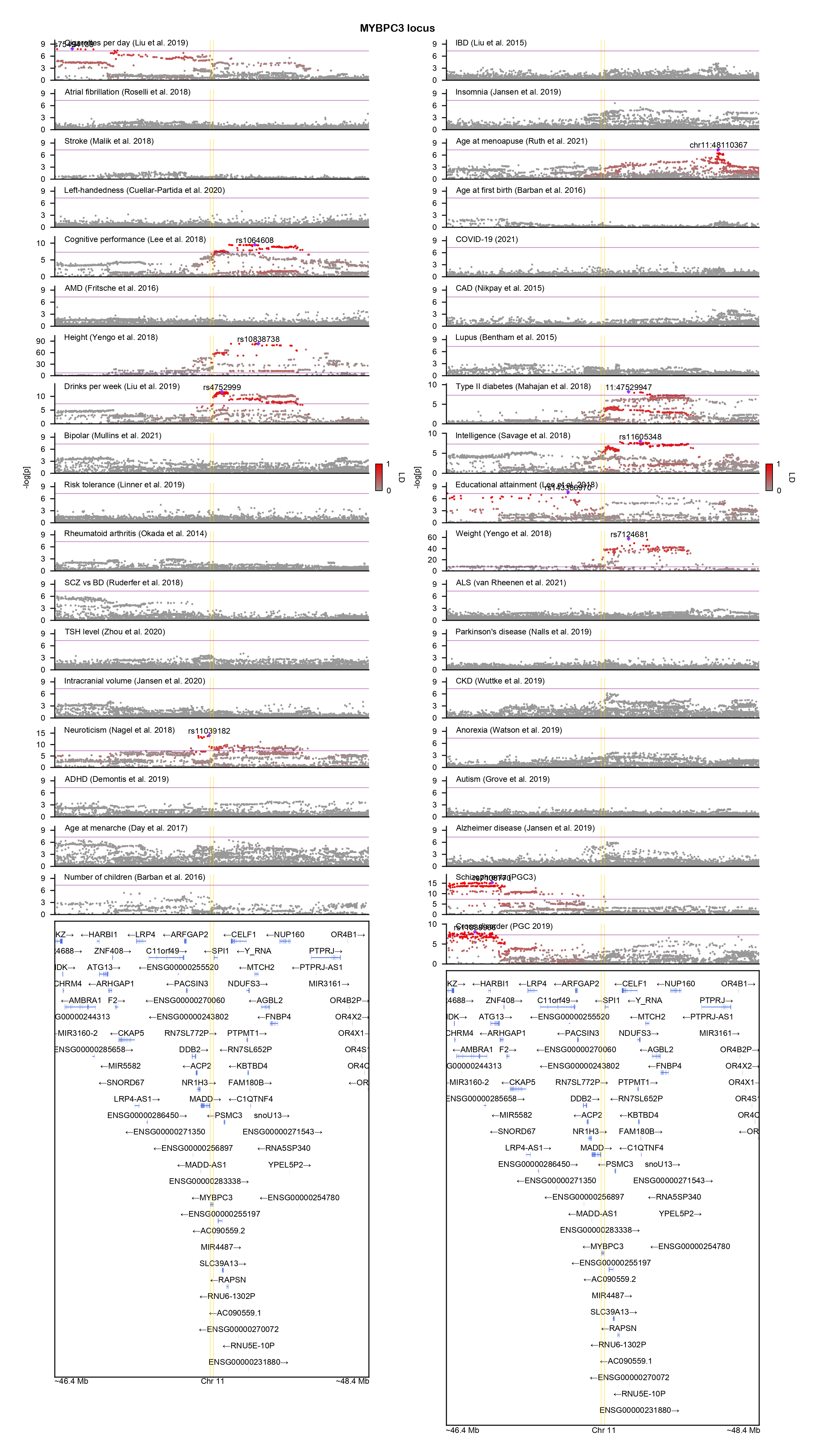

### NAA11-locuszoom.png

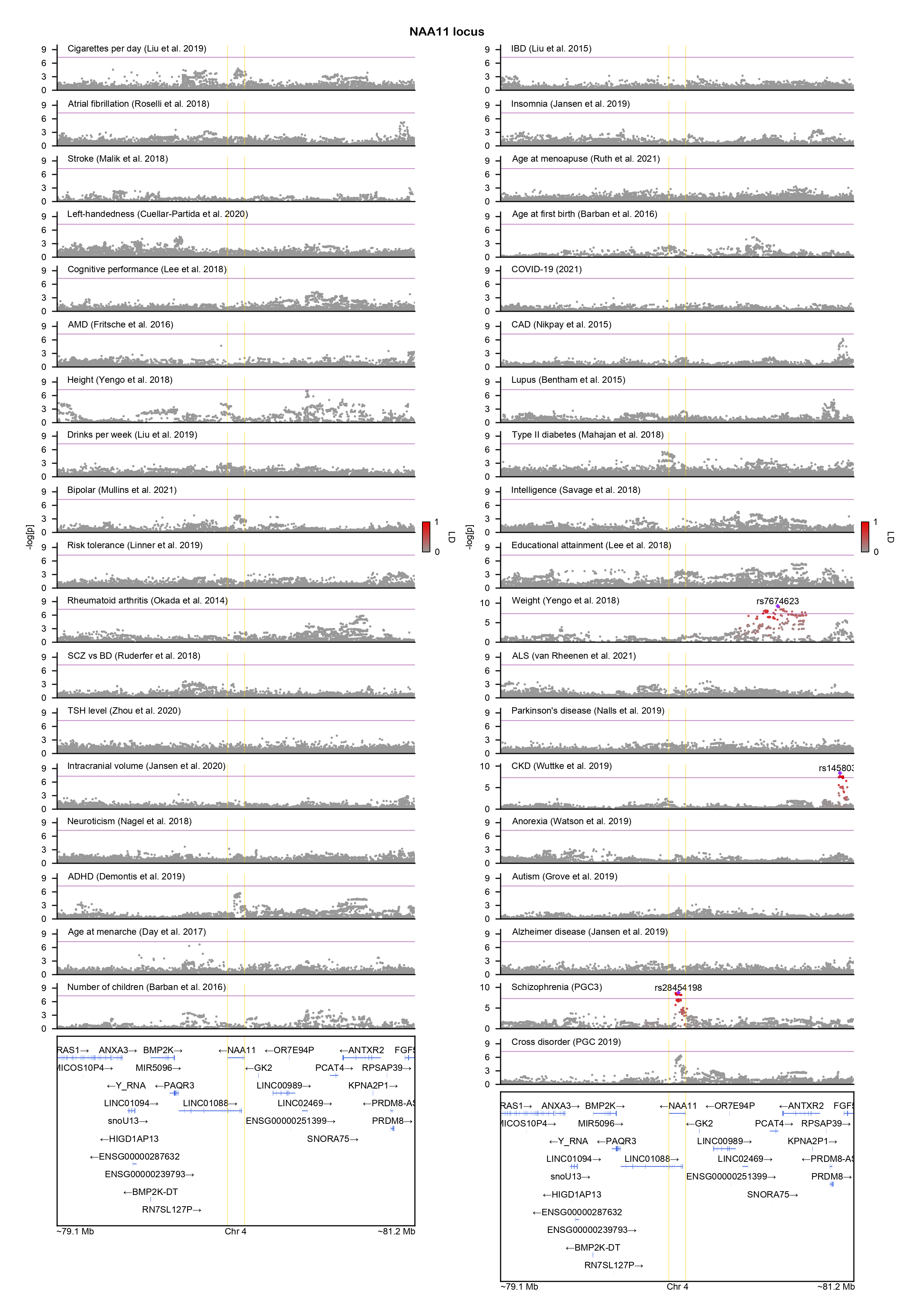

### NDST3-locuszoom.png

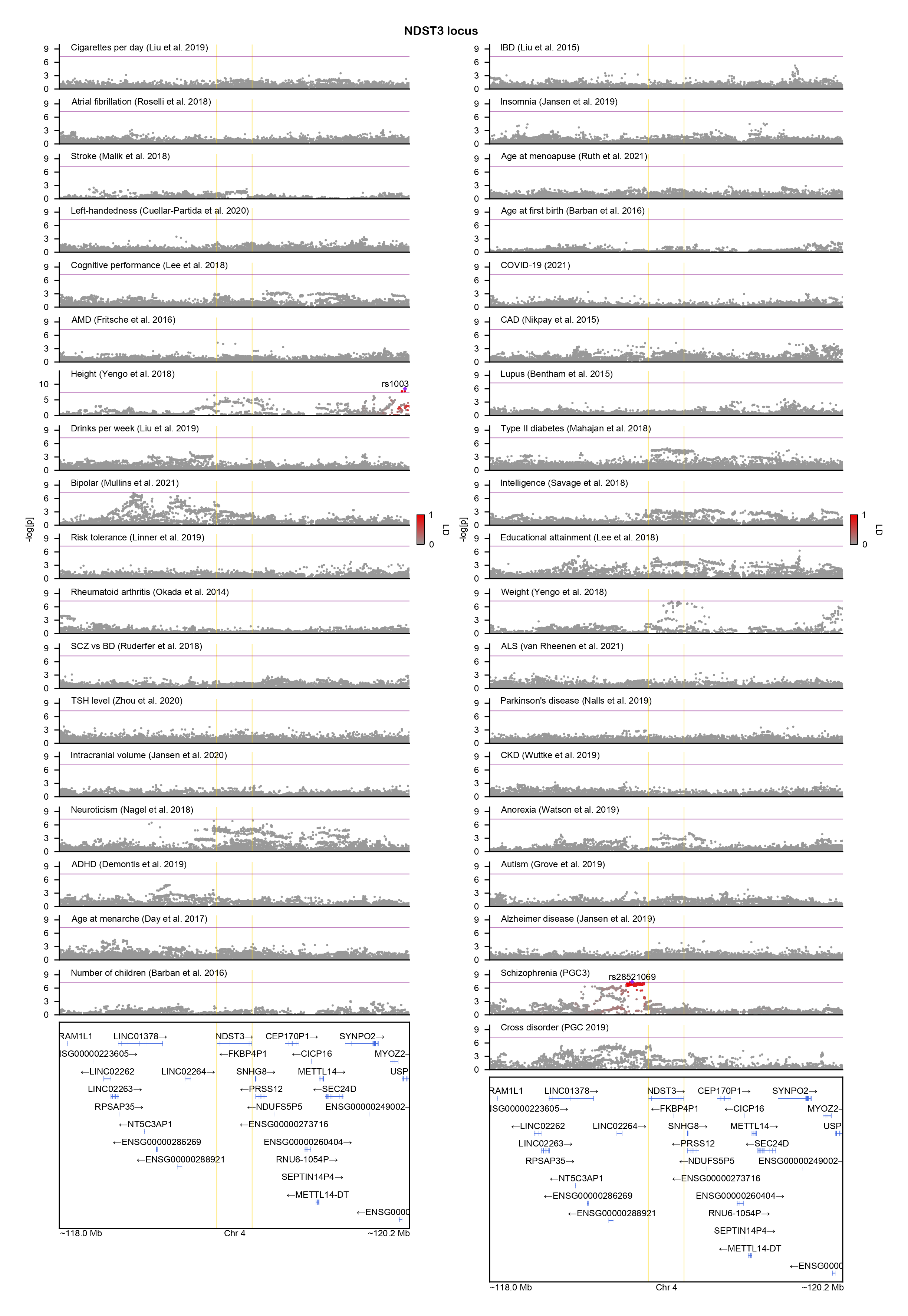

### NEBL-locuszoom.png

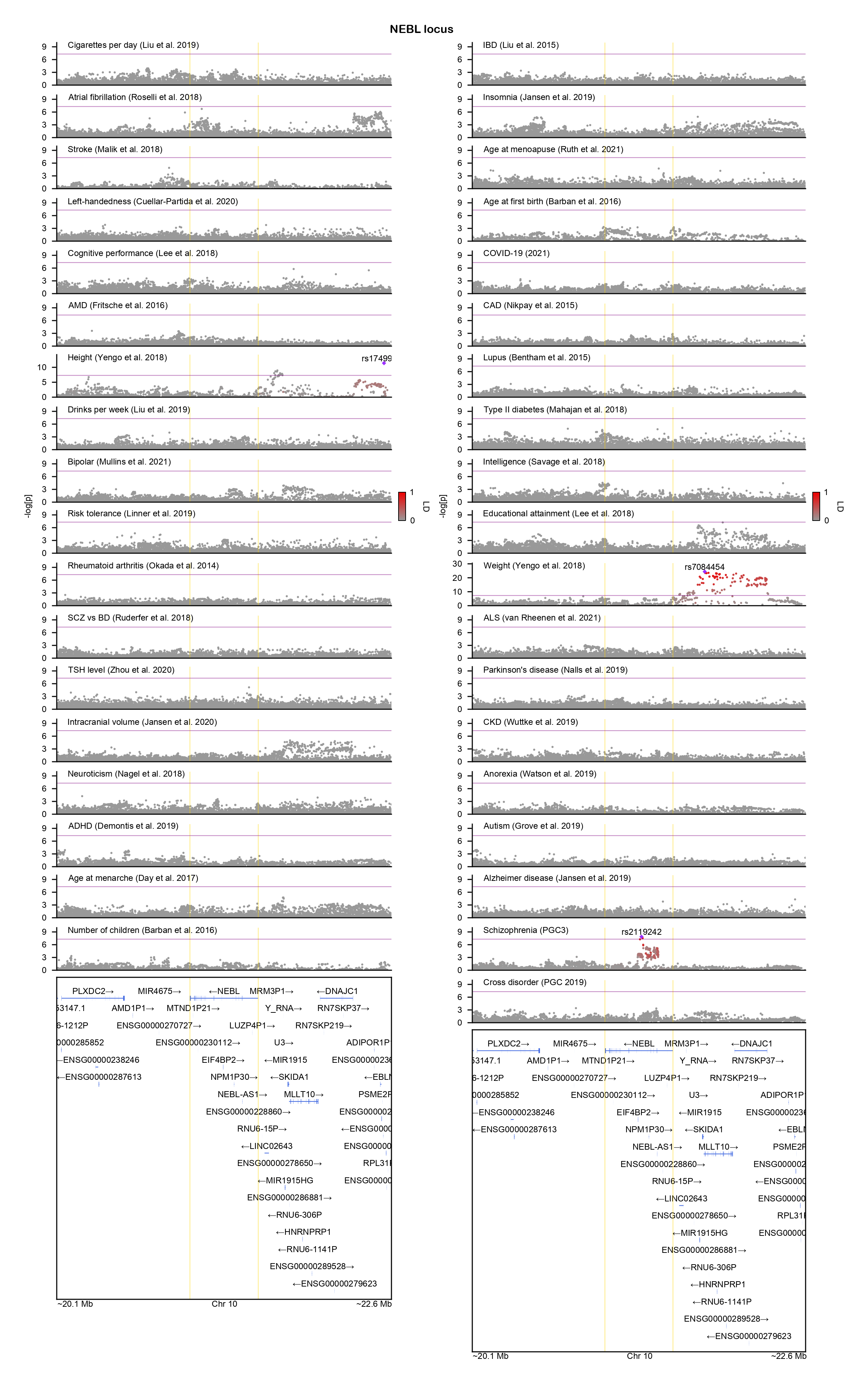

### NEGR1-locuszoom.png

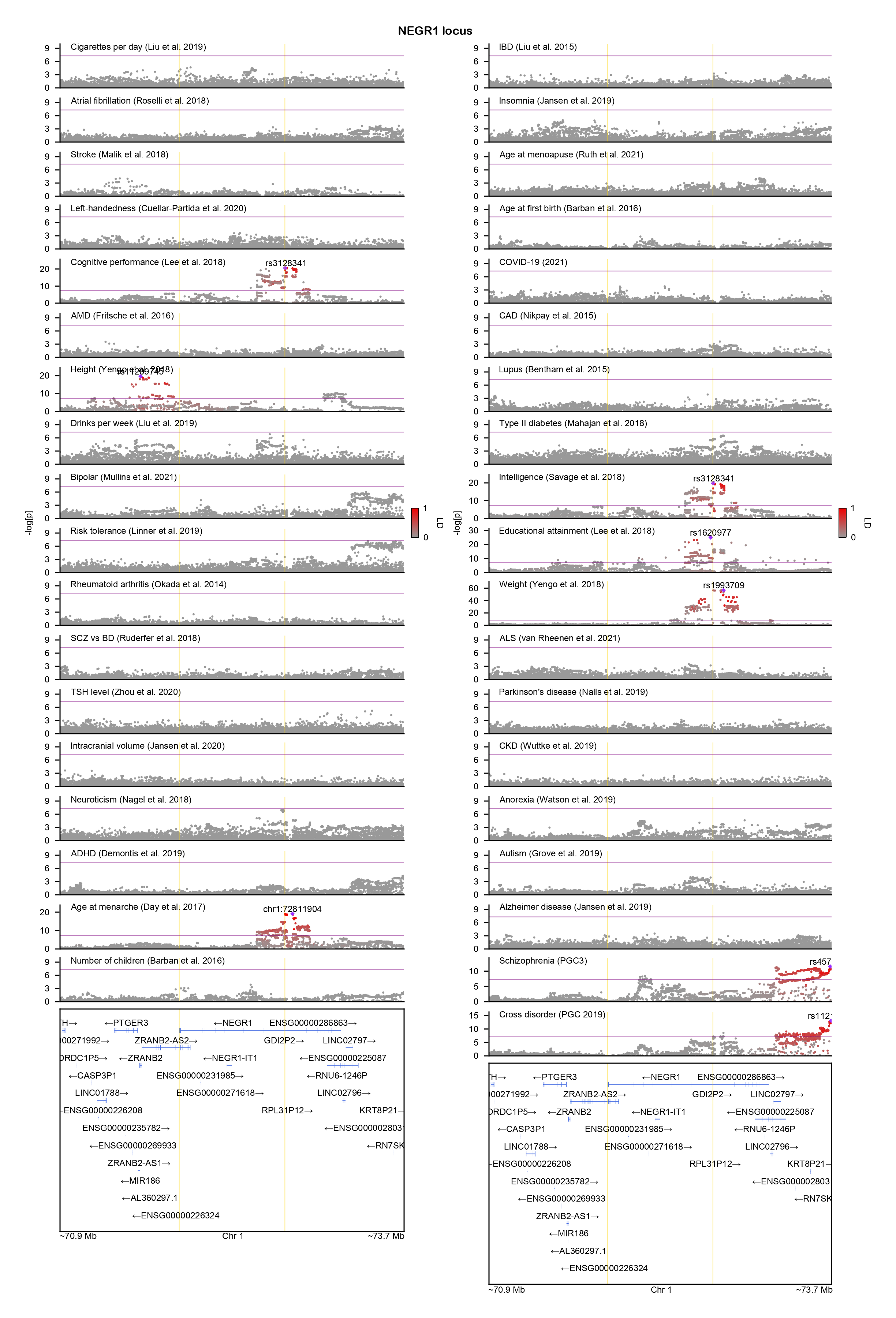

### NMUR2-locuszoom.png

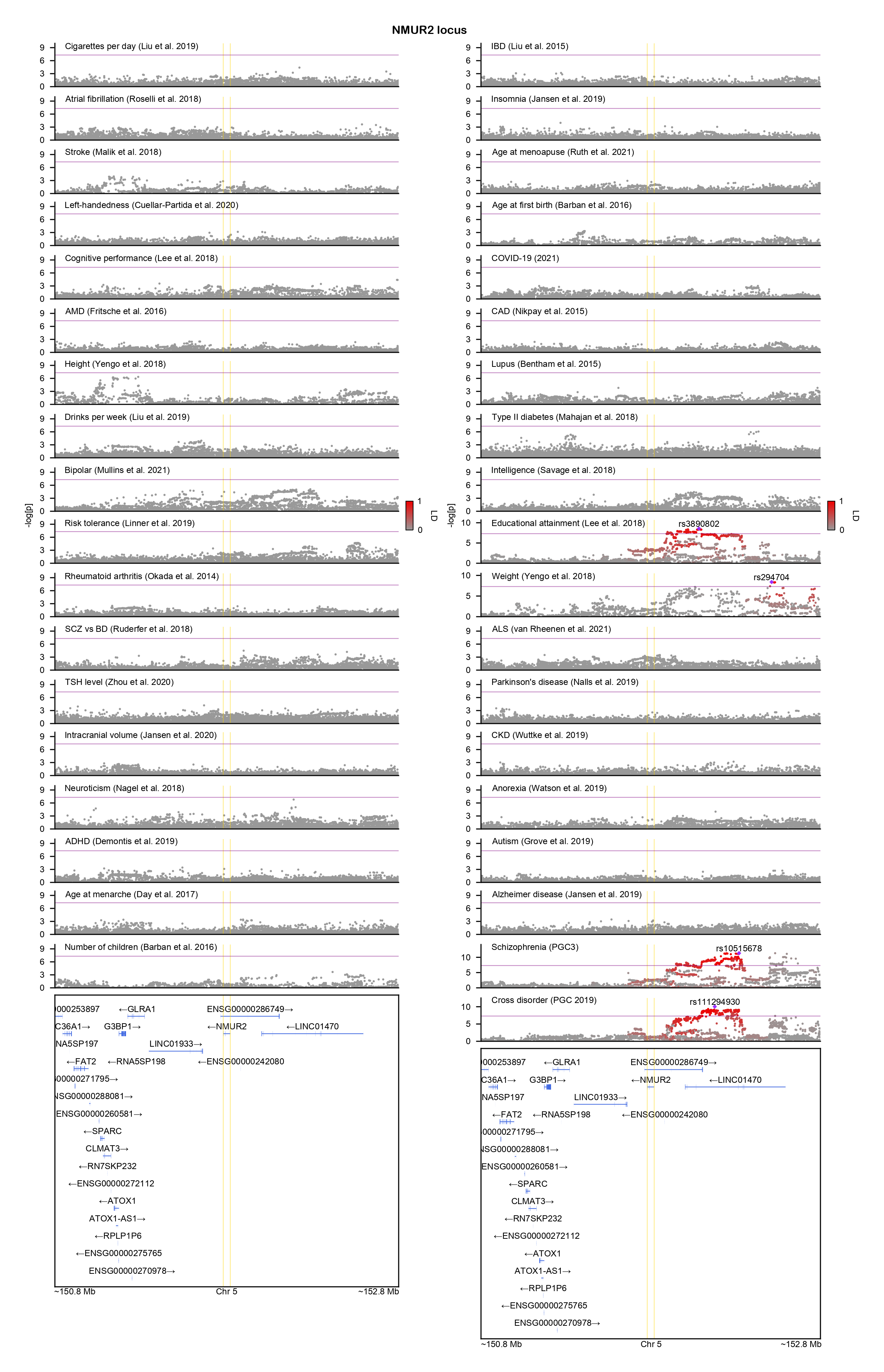

### NPAS3-locuszoom.png

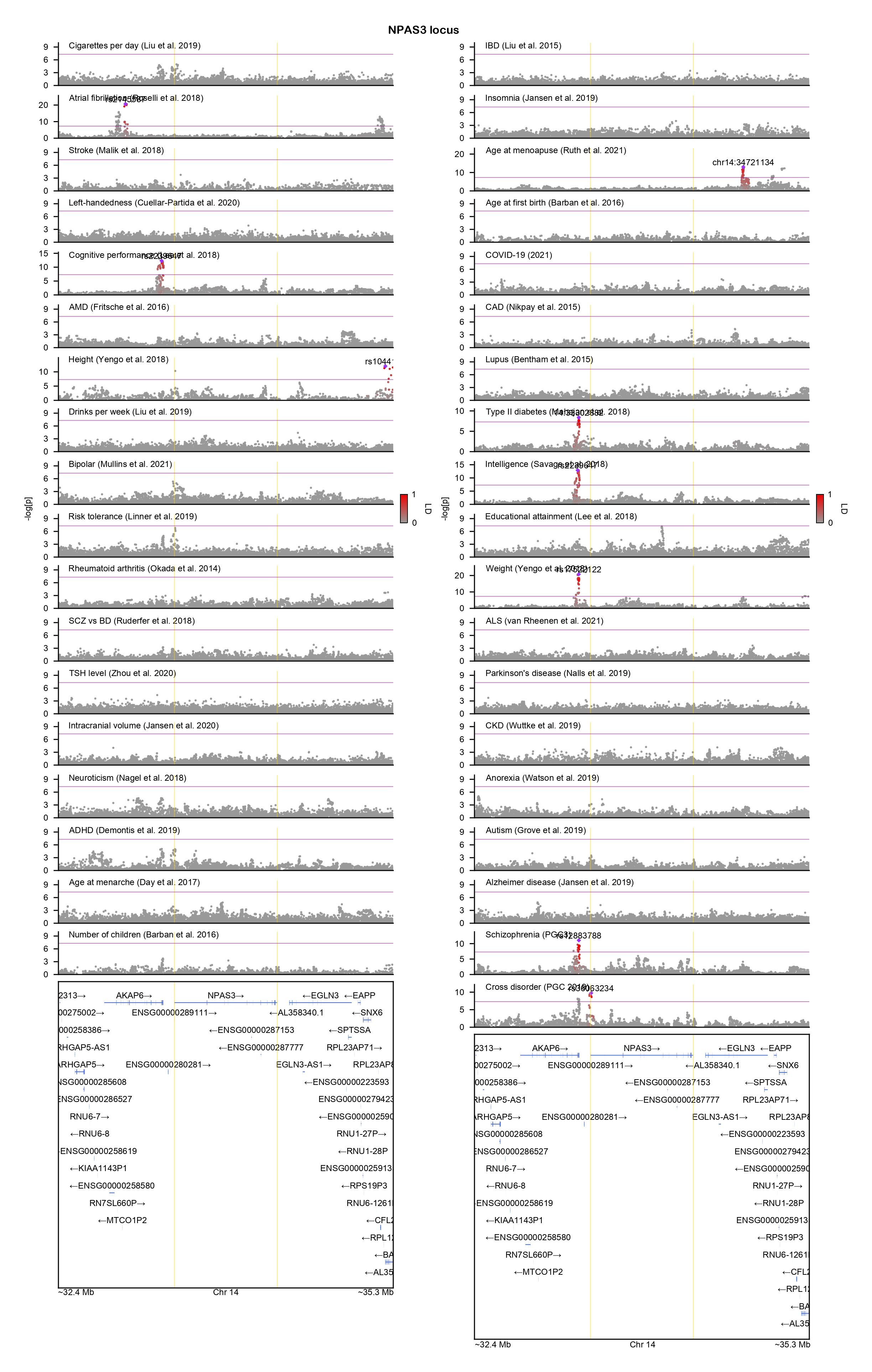

### NRIP1-locuszoom.png

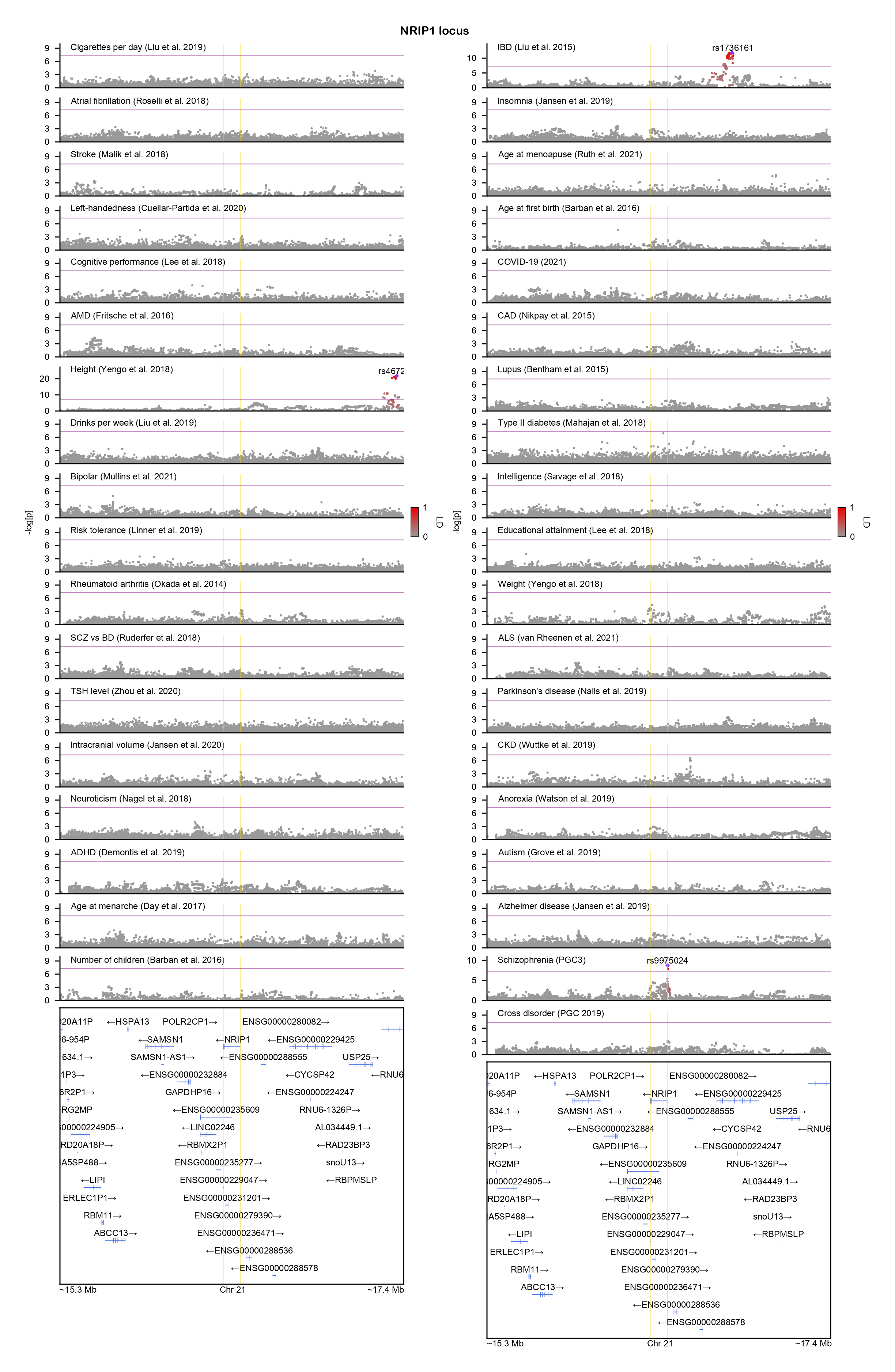

### OTOL1-locuszoom.png

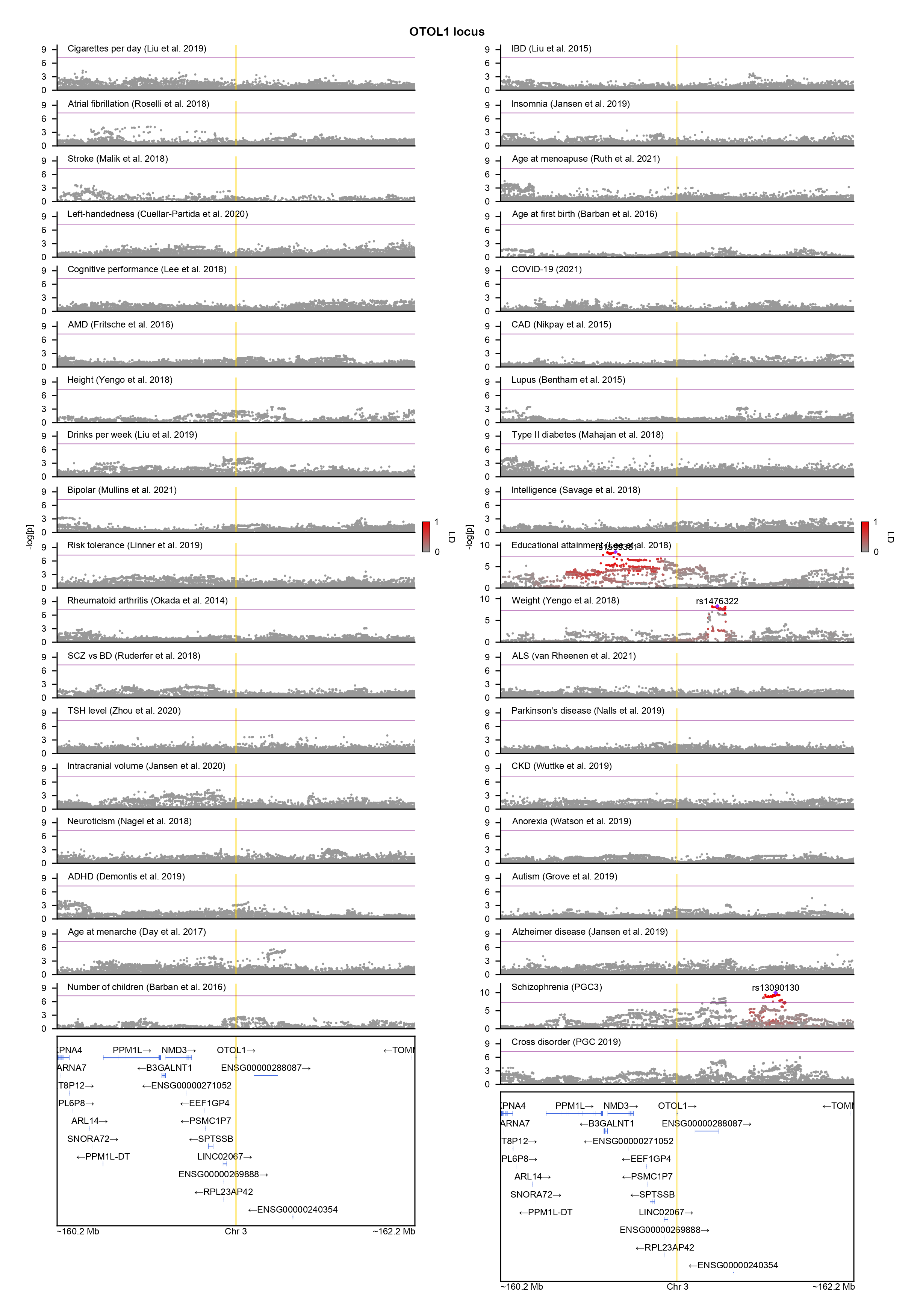

### PAH-locuszoom.png

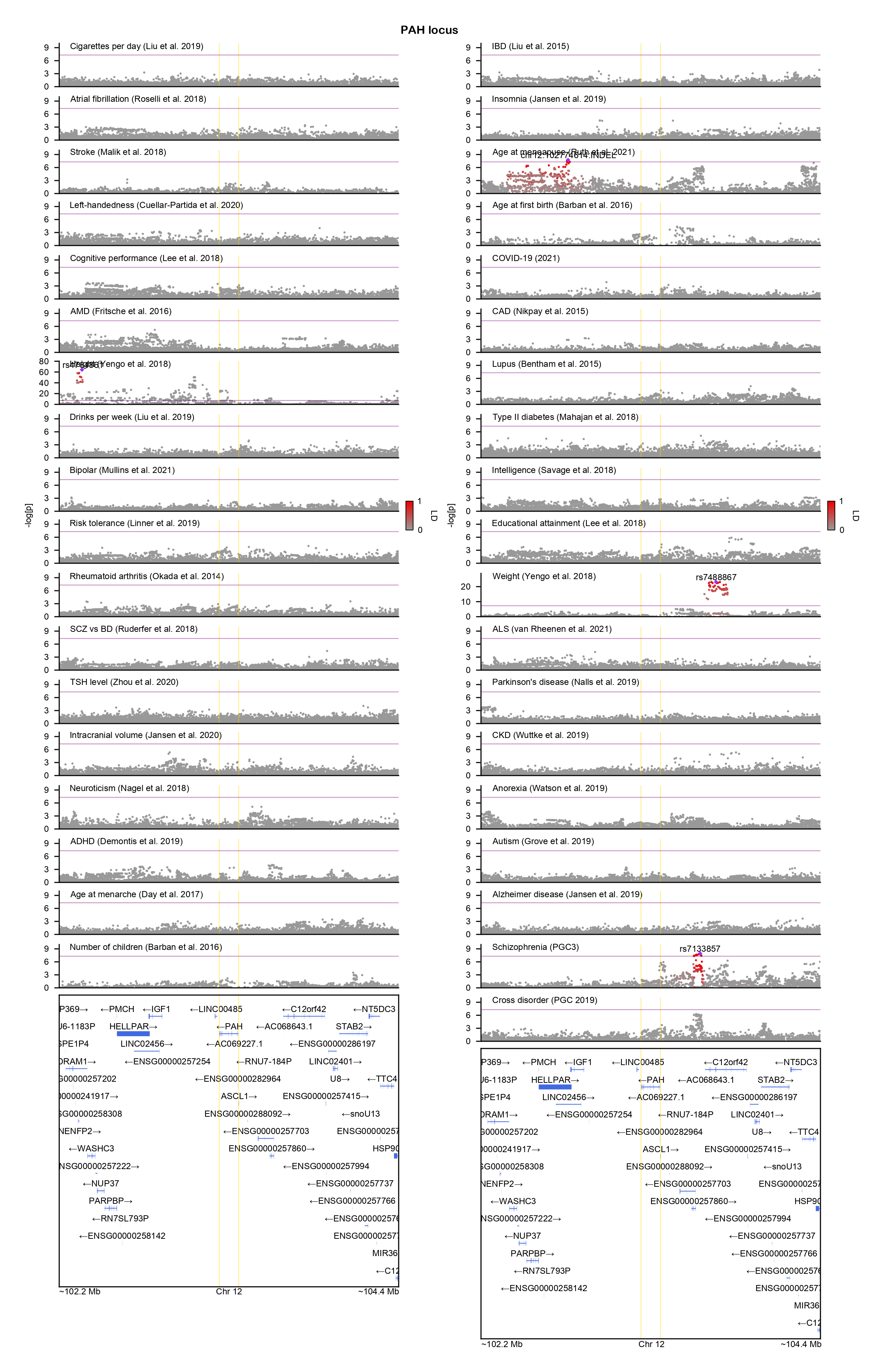

### PCDH17-locuszoom.png

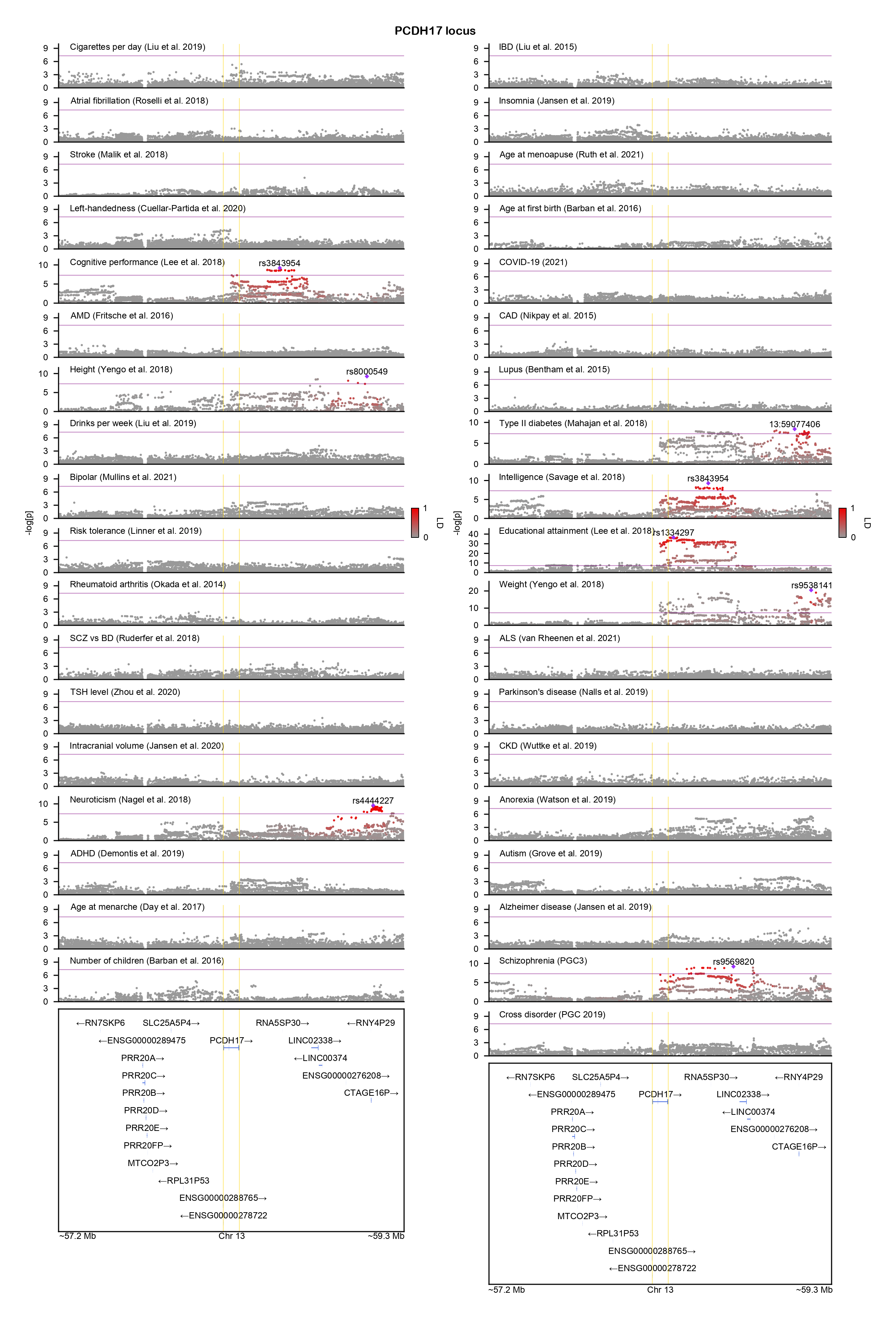

### PDE4B-locuszoom.png

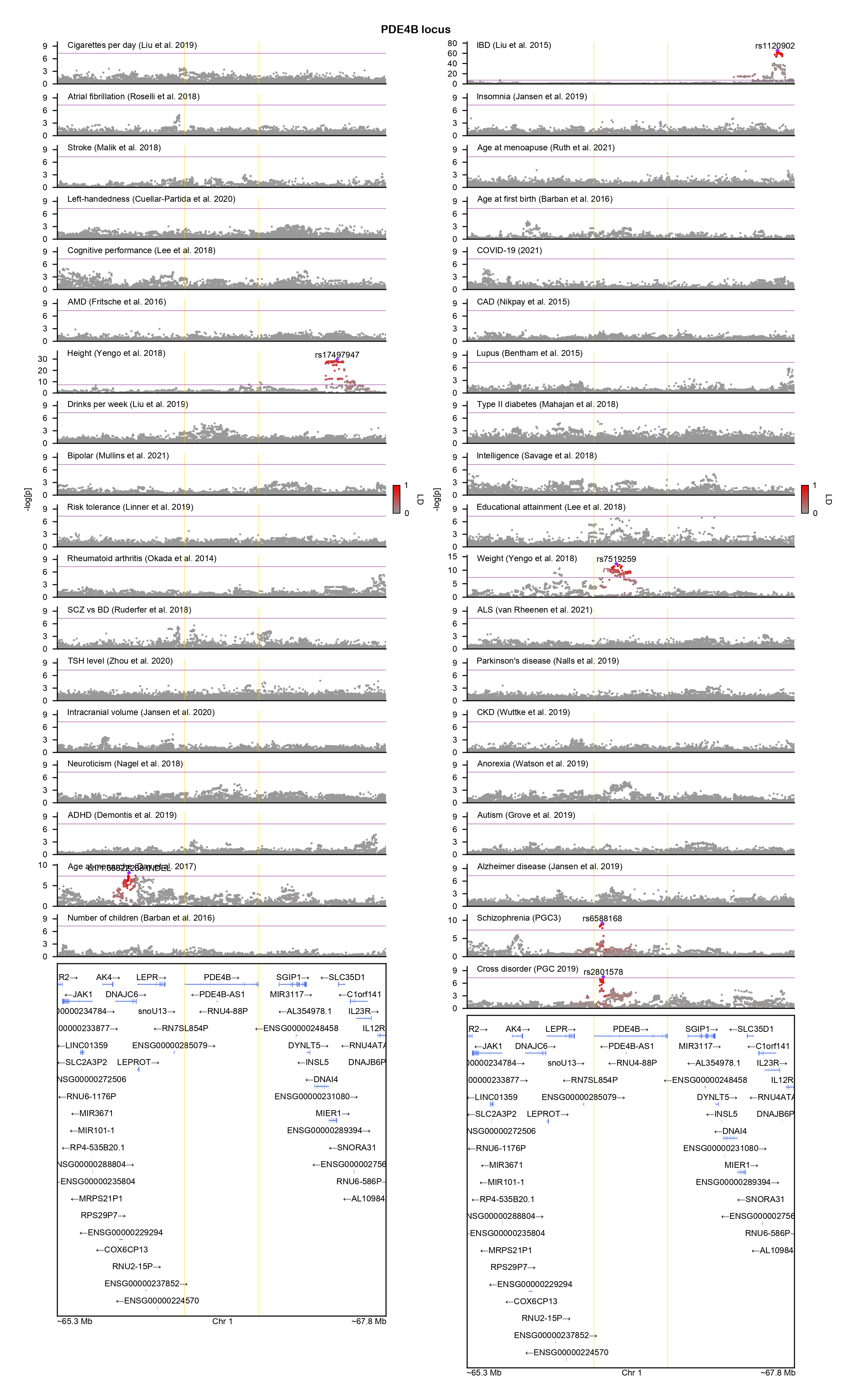

### PHF3-locuszoom.png

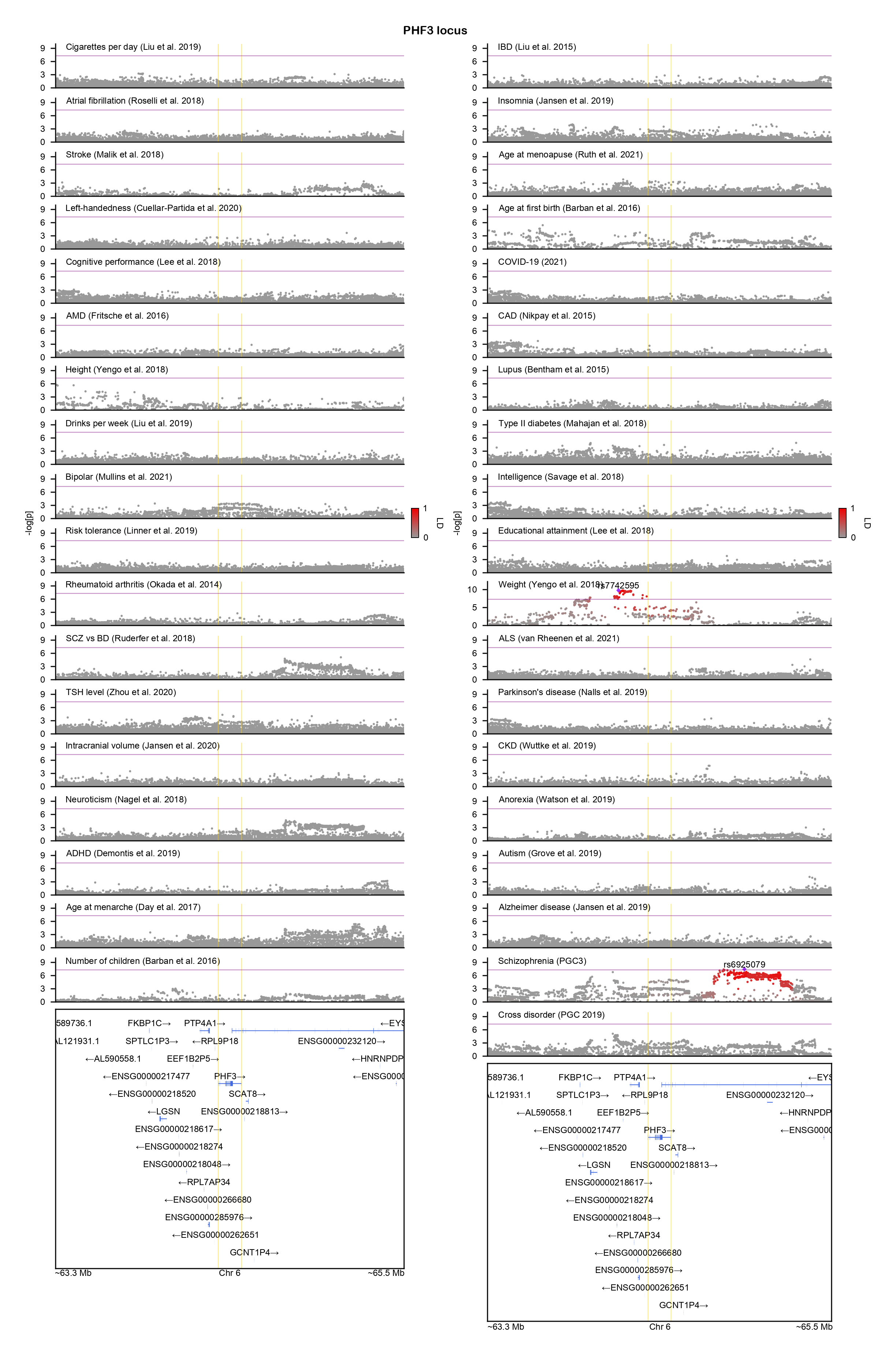

### PLCH2-locuszoom.png

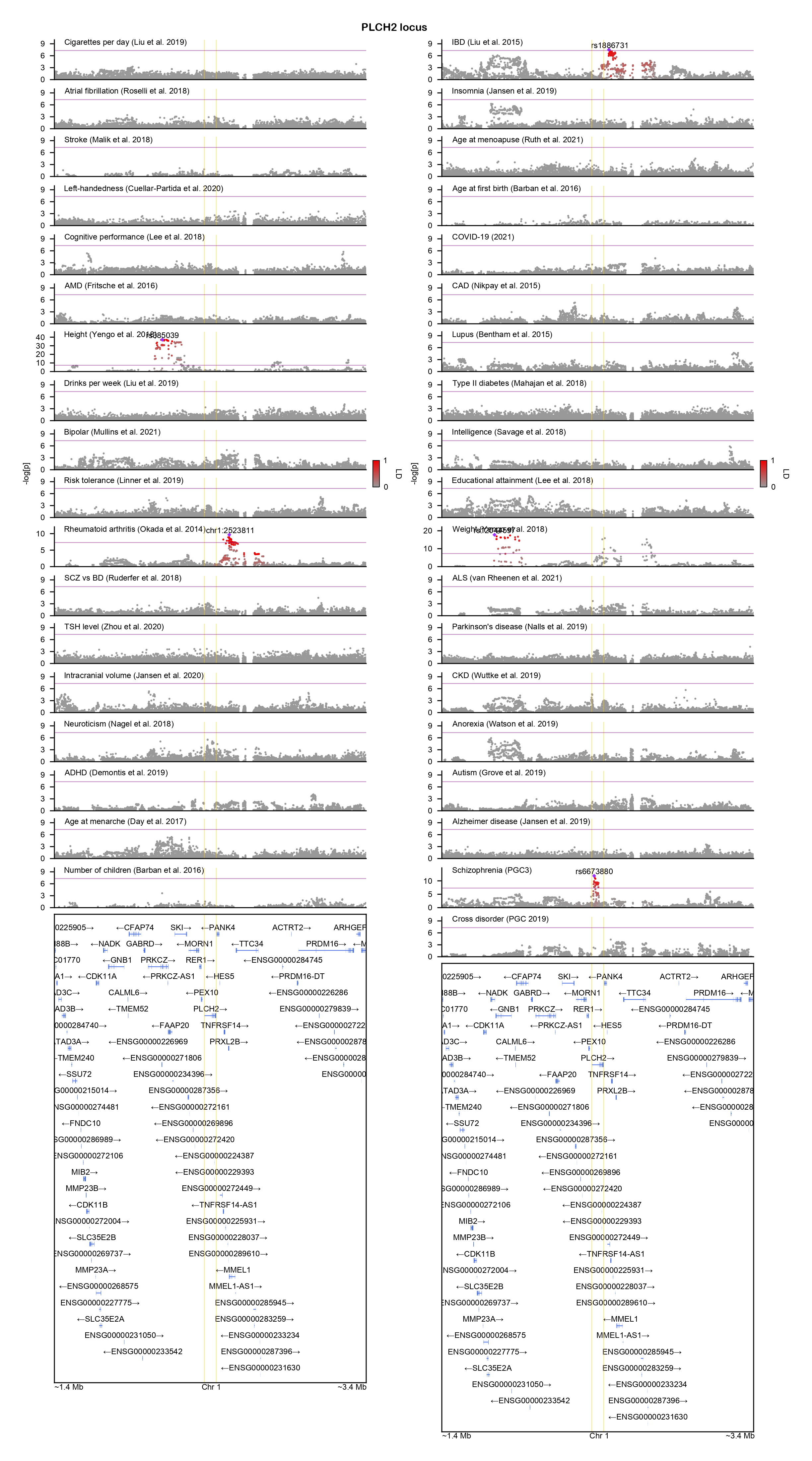

### PLK2-locuszoom.png

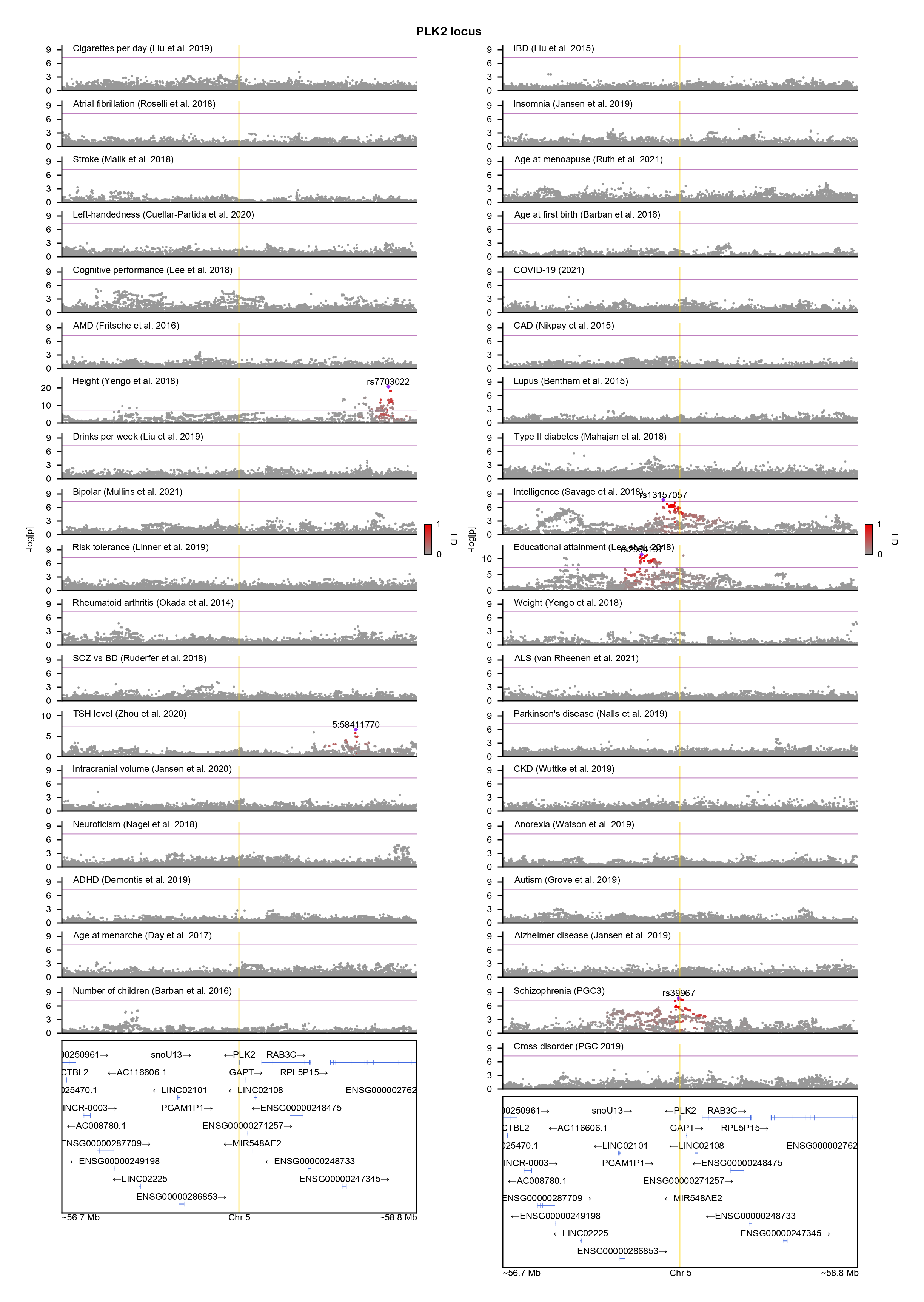

### POLG-locuszoom.png

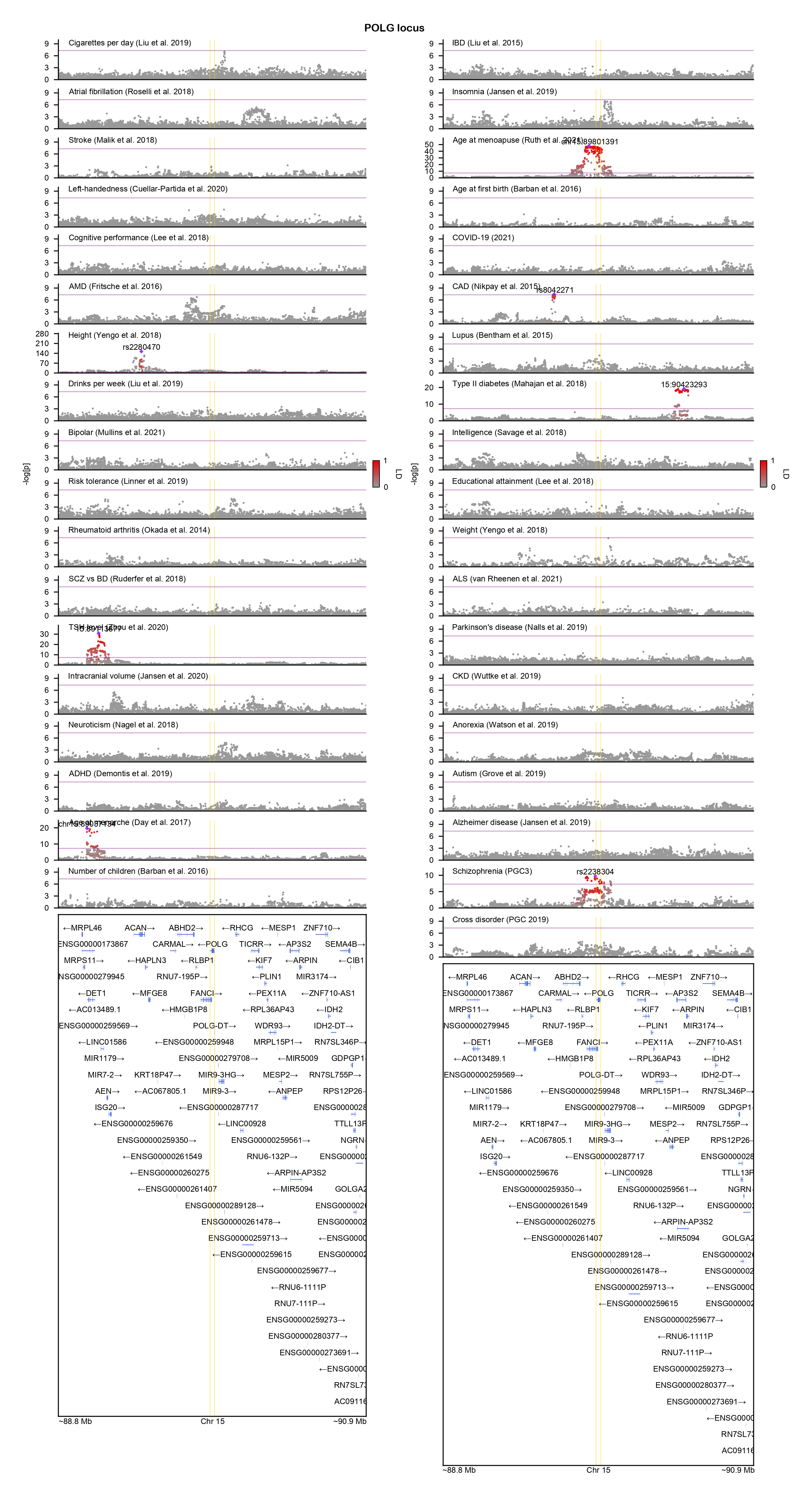

### PRKD1-locuszoom.png

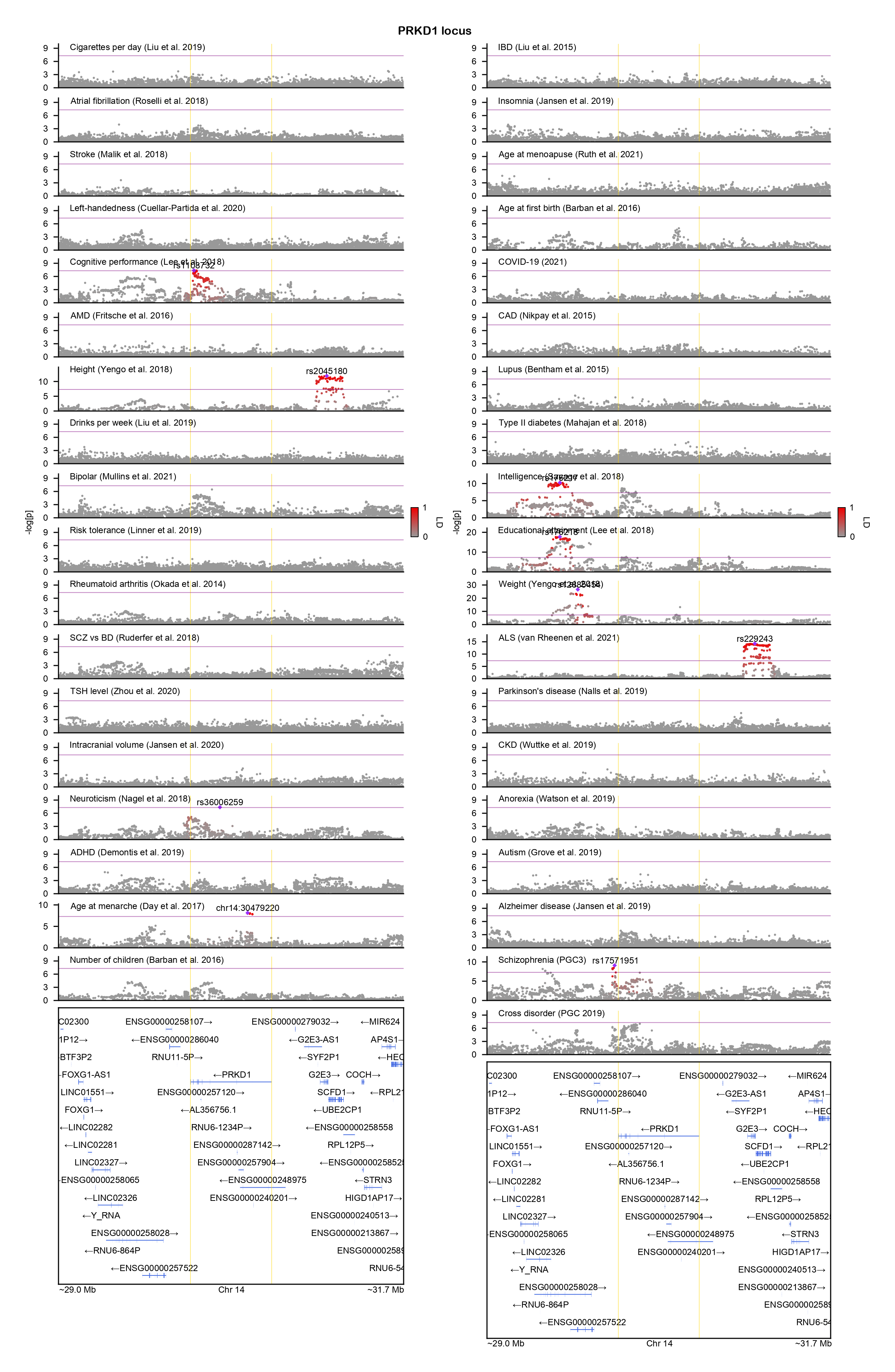

### PTBP2-locuszoom.png

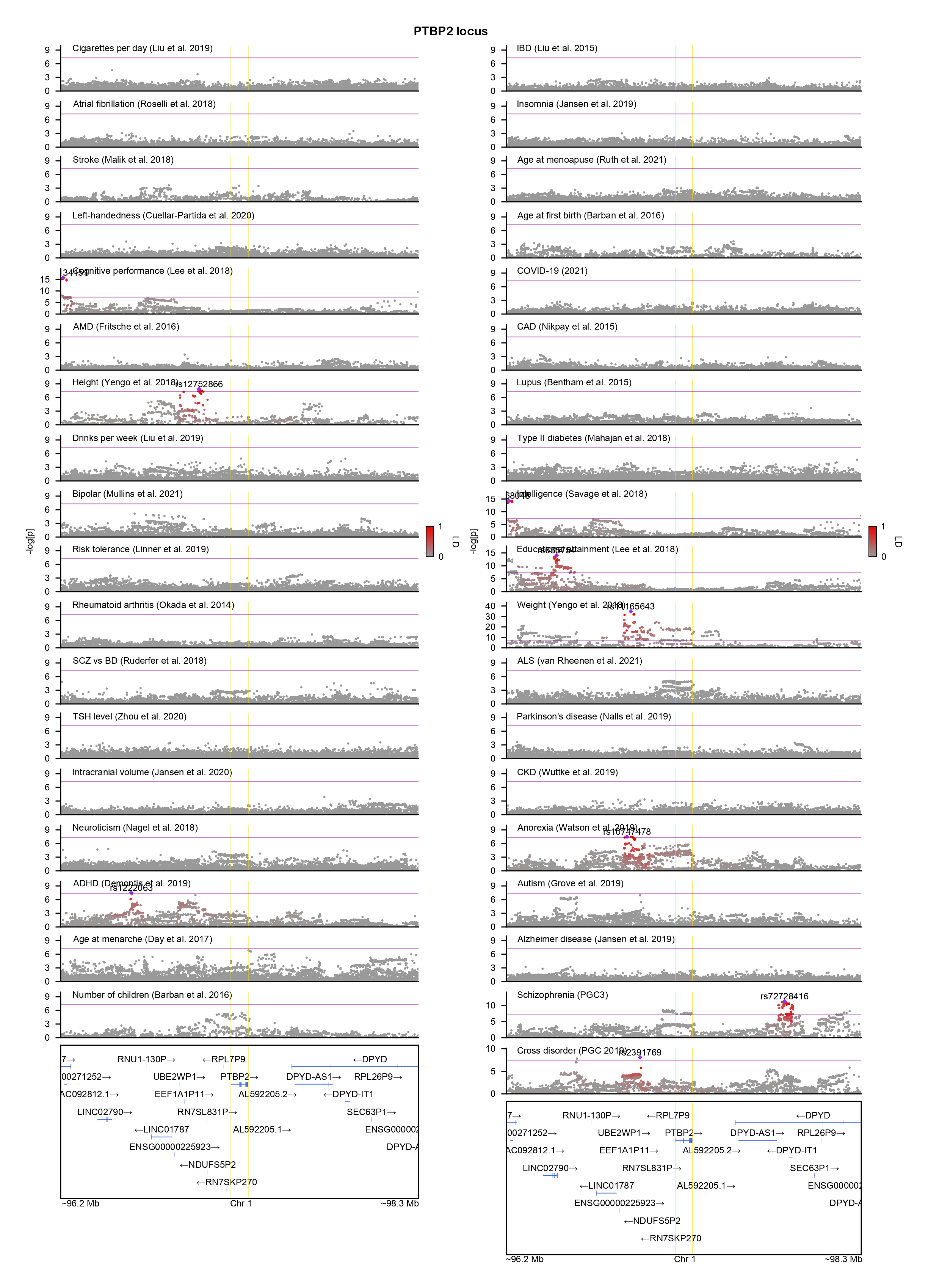

### PTPRD-locuszoom.png

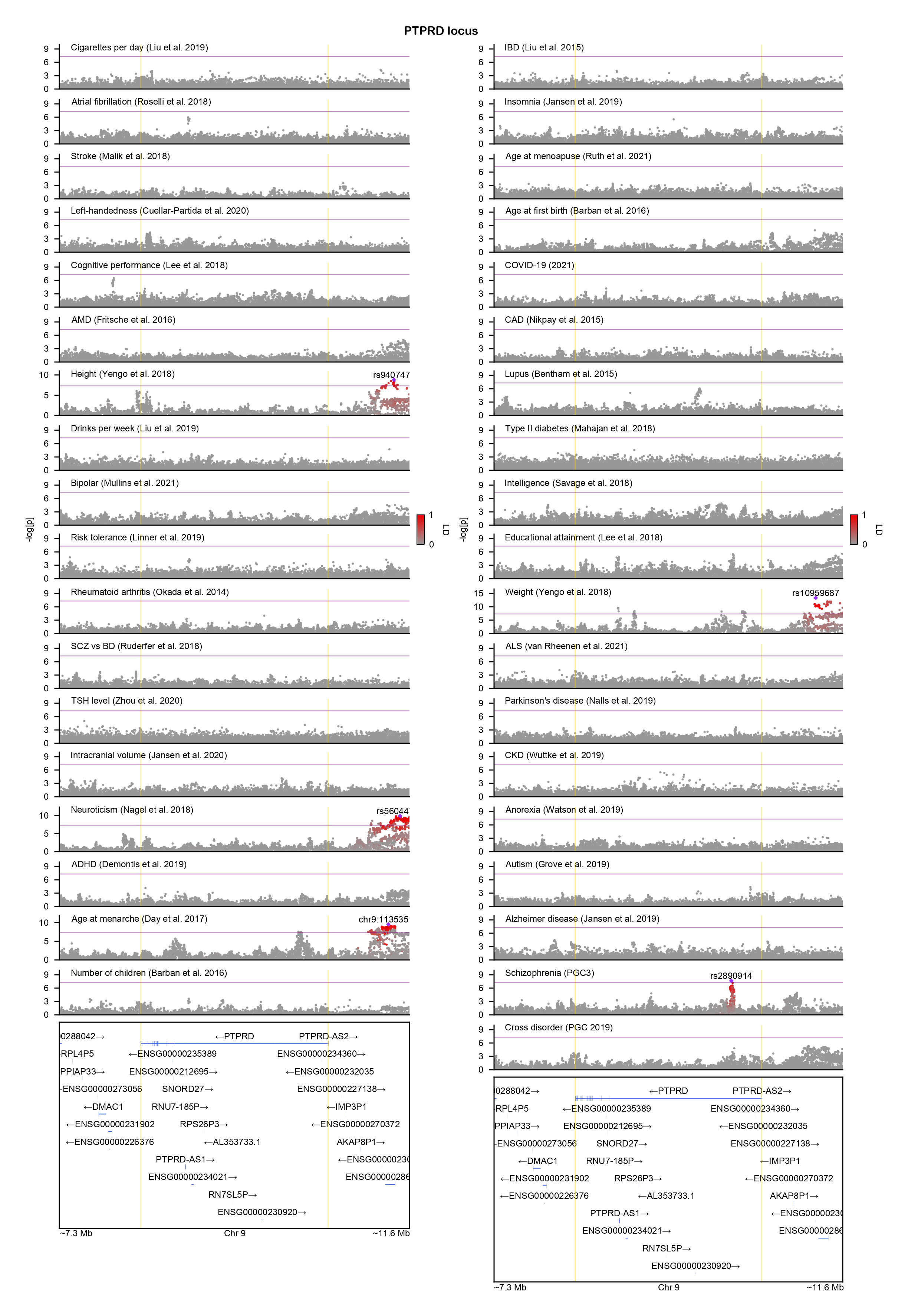

### RORA-locuszoom.png

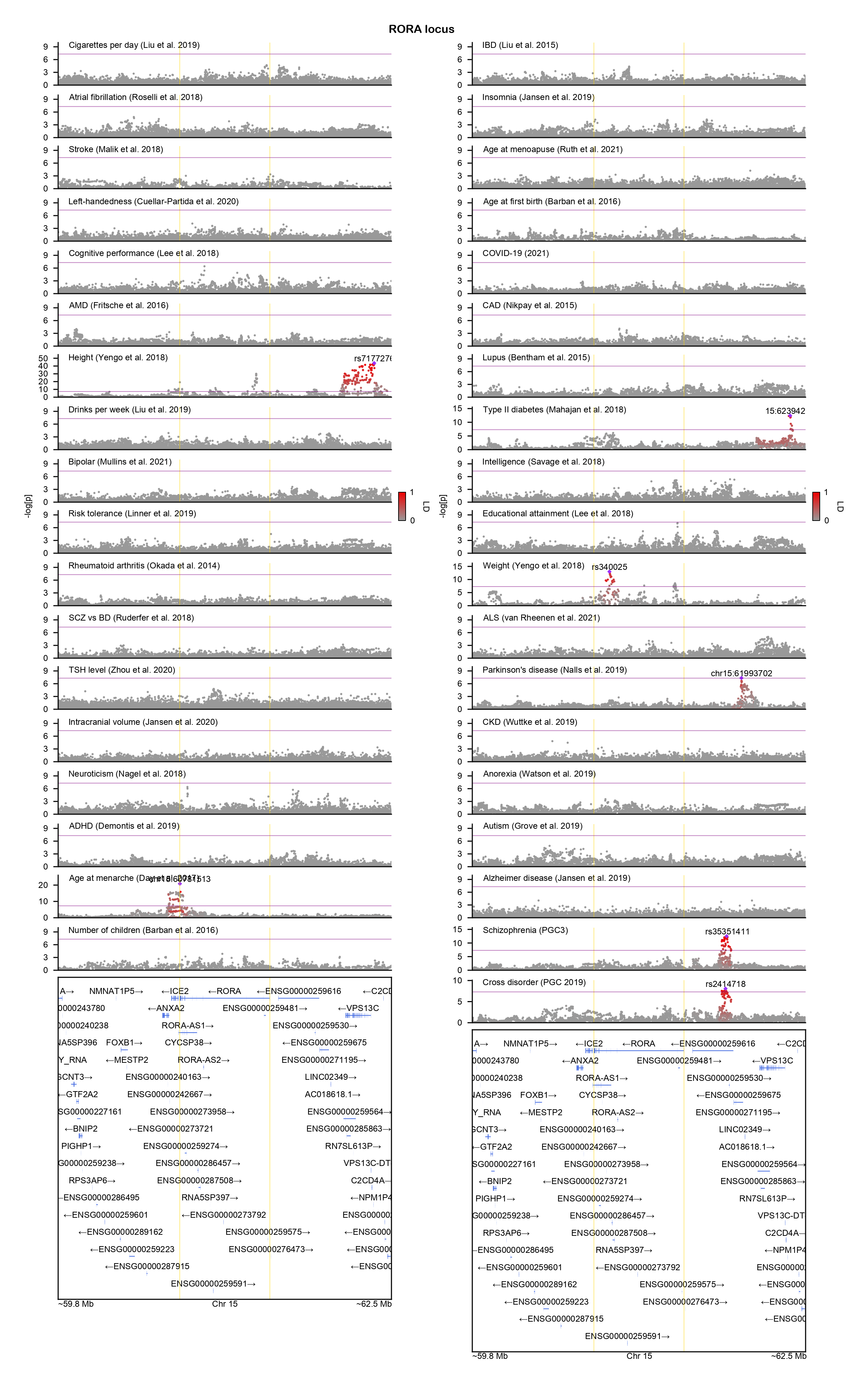

### SLC9C2-locuszoom.png

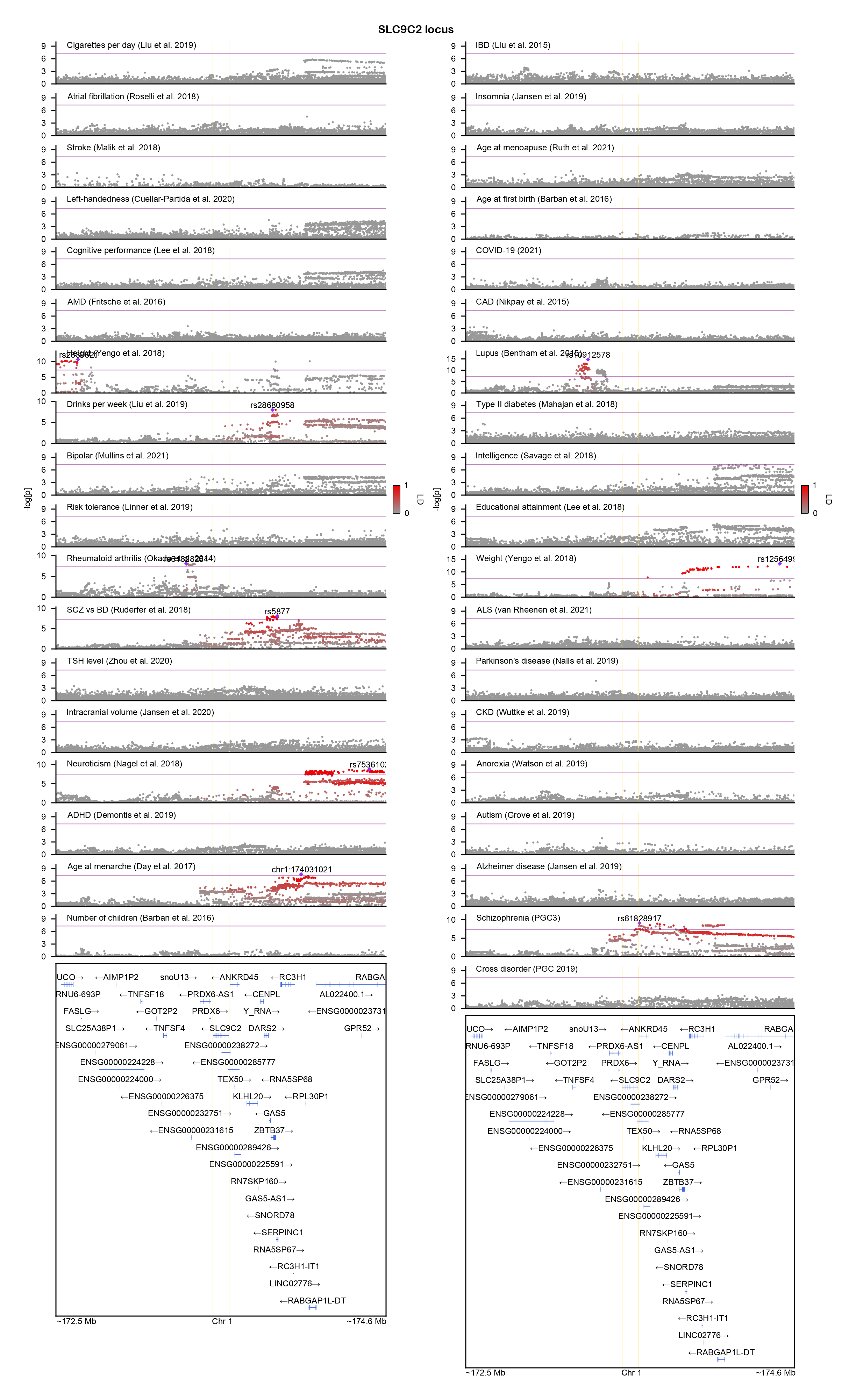

### SP3-locuszoom.png

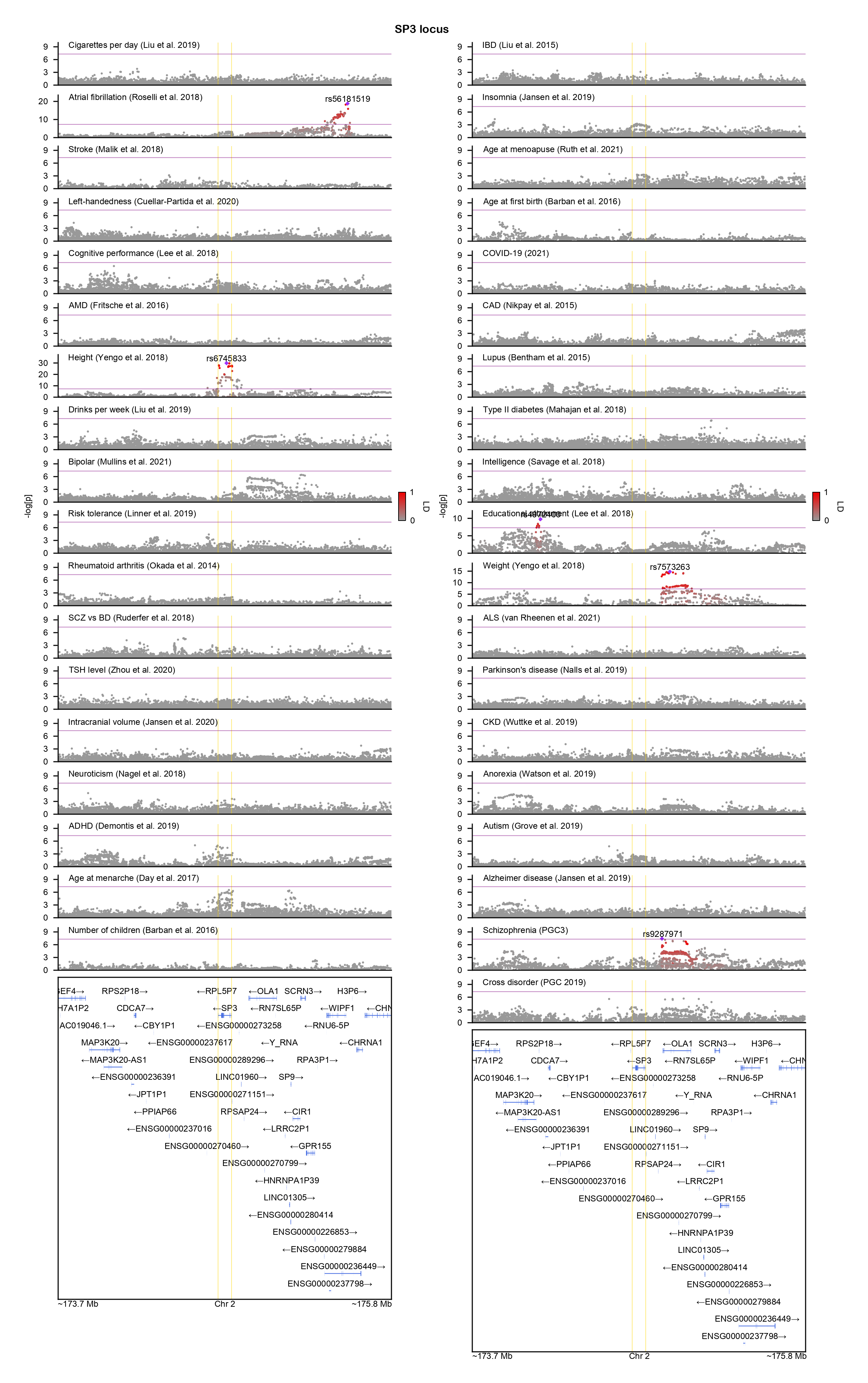

### SRR-locuszoom.png

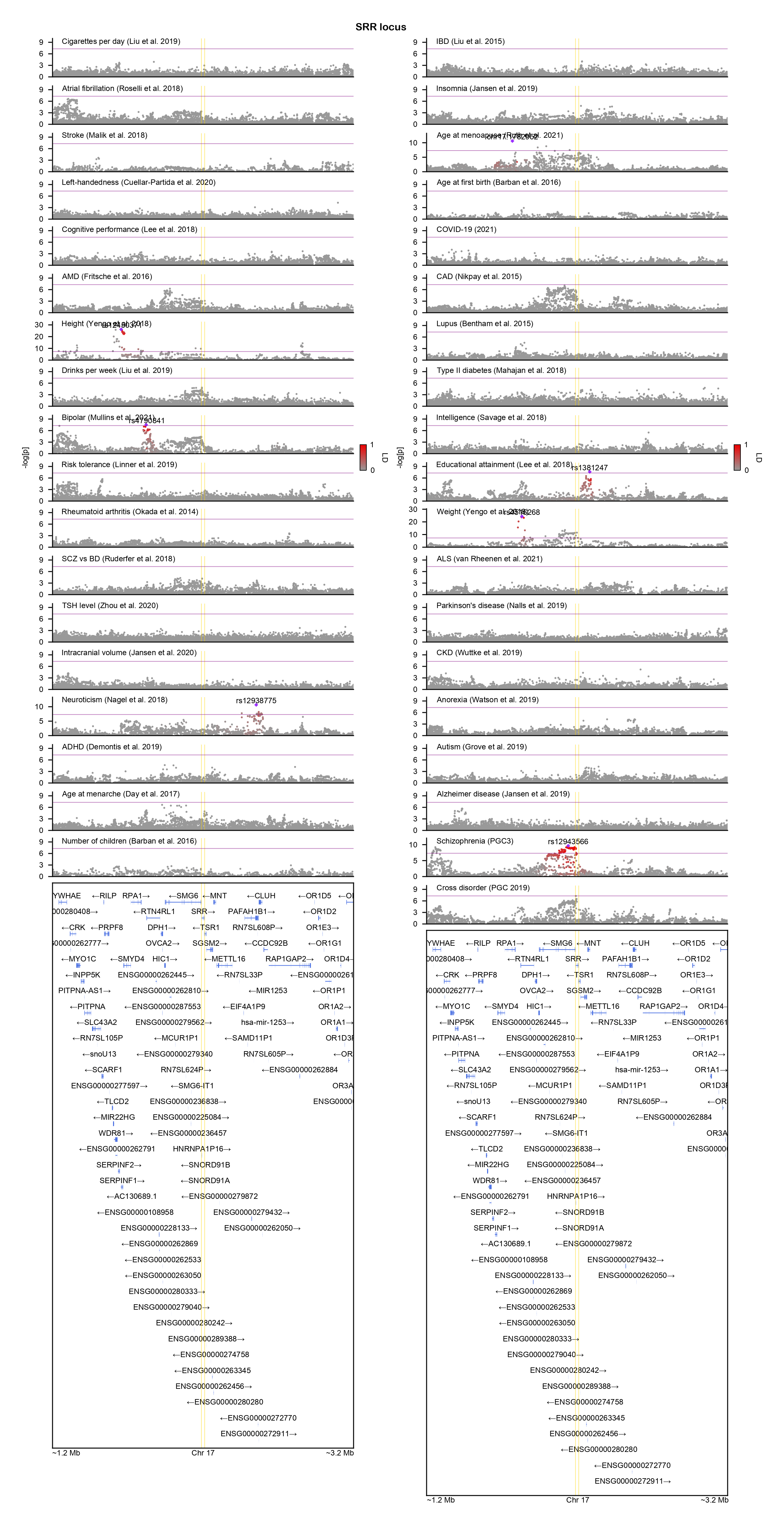

### SYNGAP1-locuszoom.png

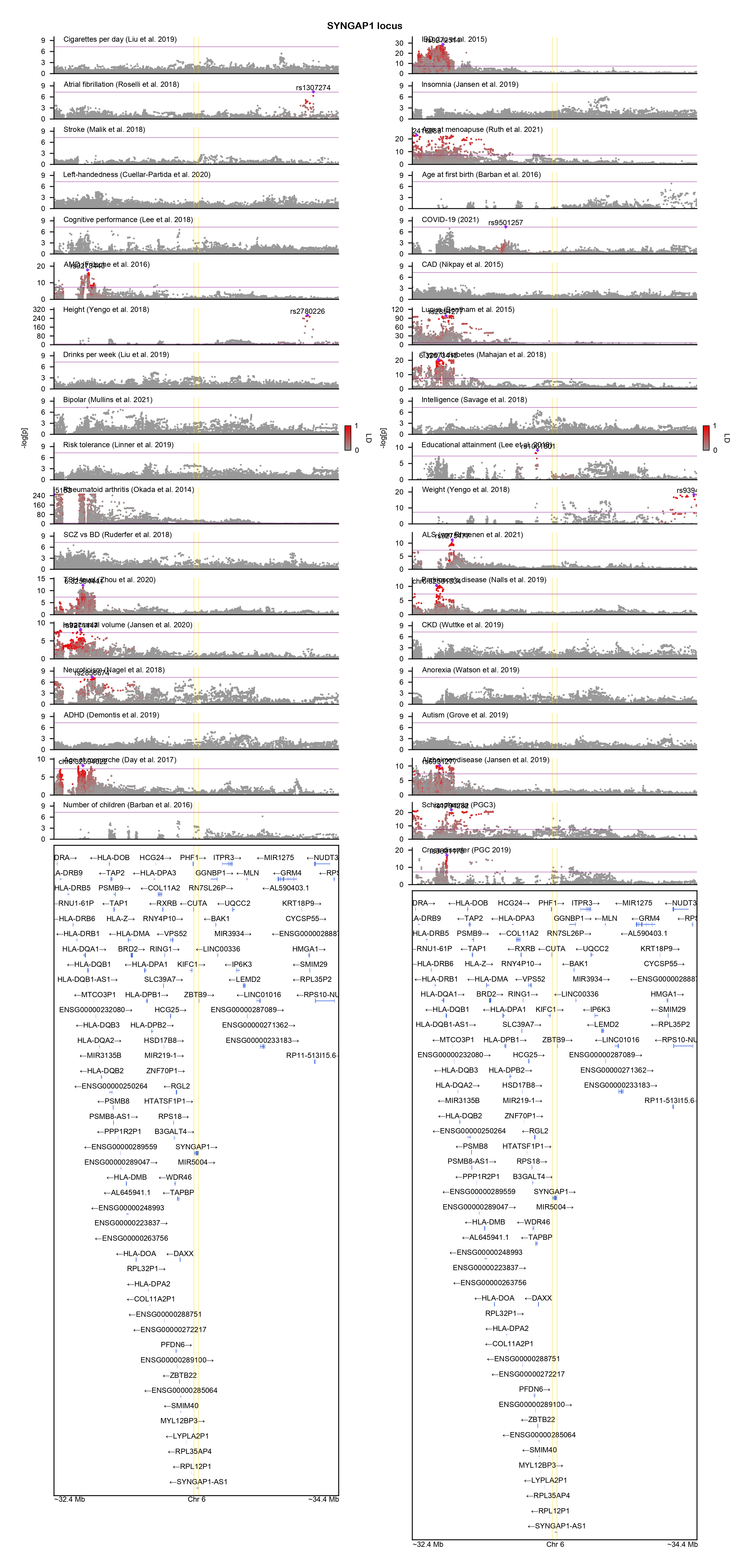

### TBC1D2-locuszoom.png

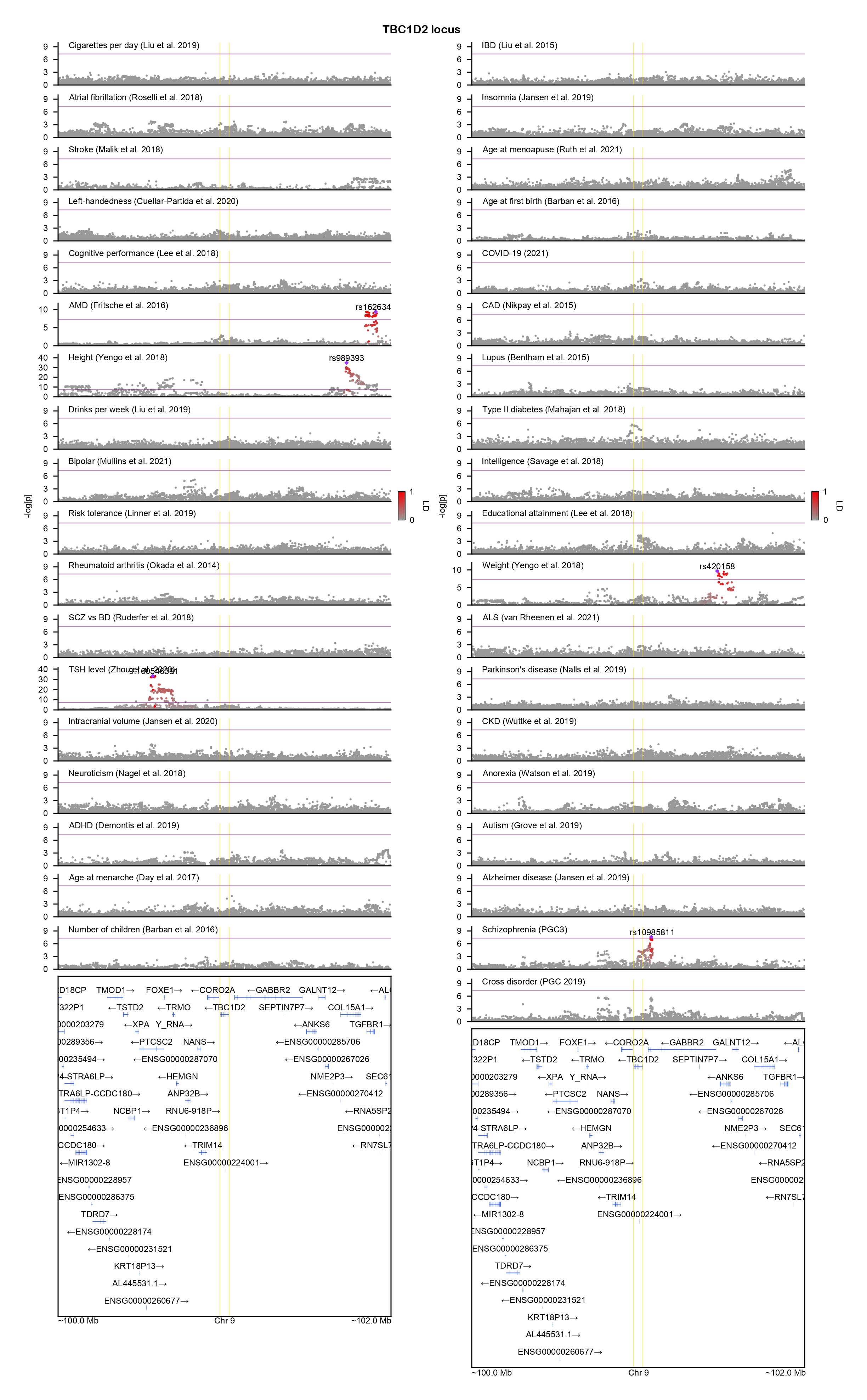

### TBC1D5-locuszoom.png

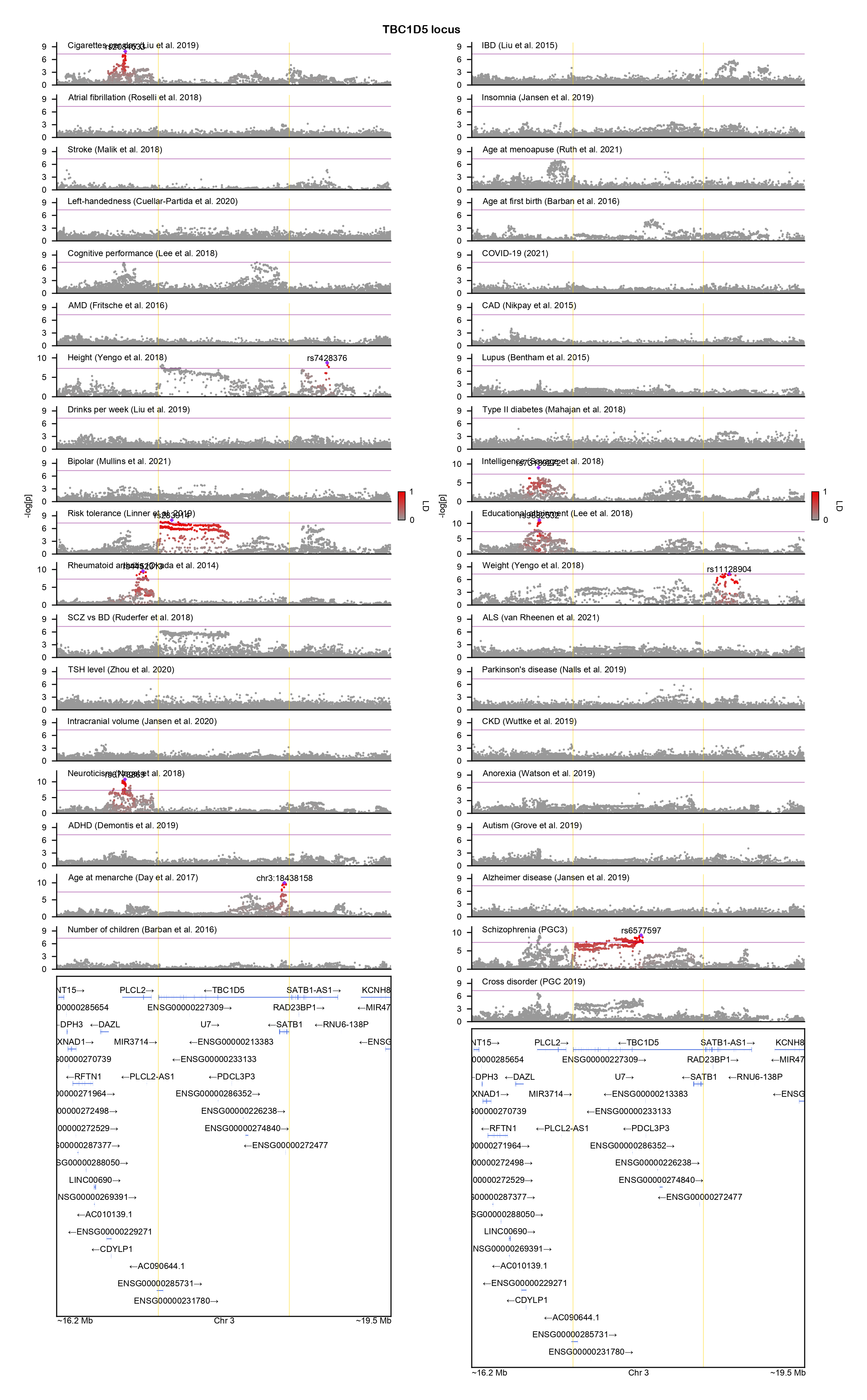

### TBL1XR1-locuszoom.png

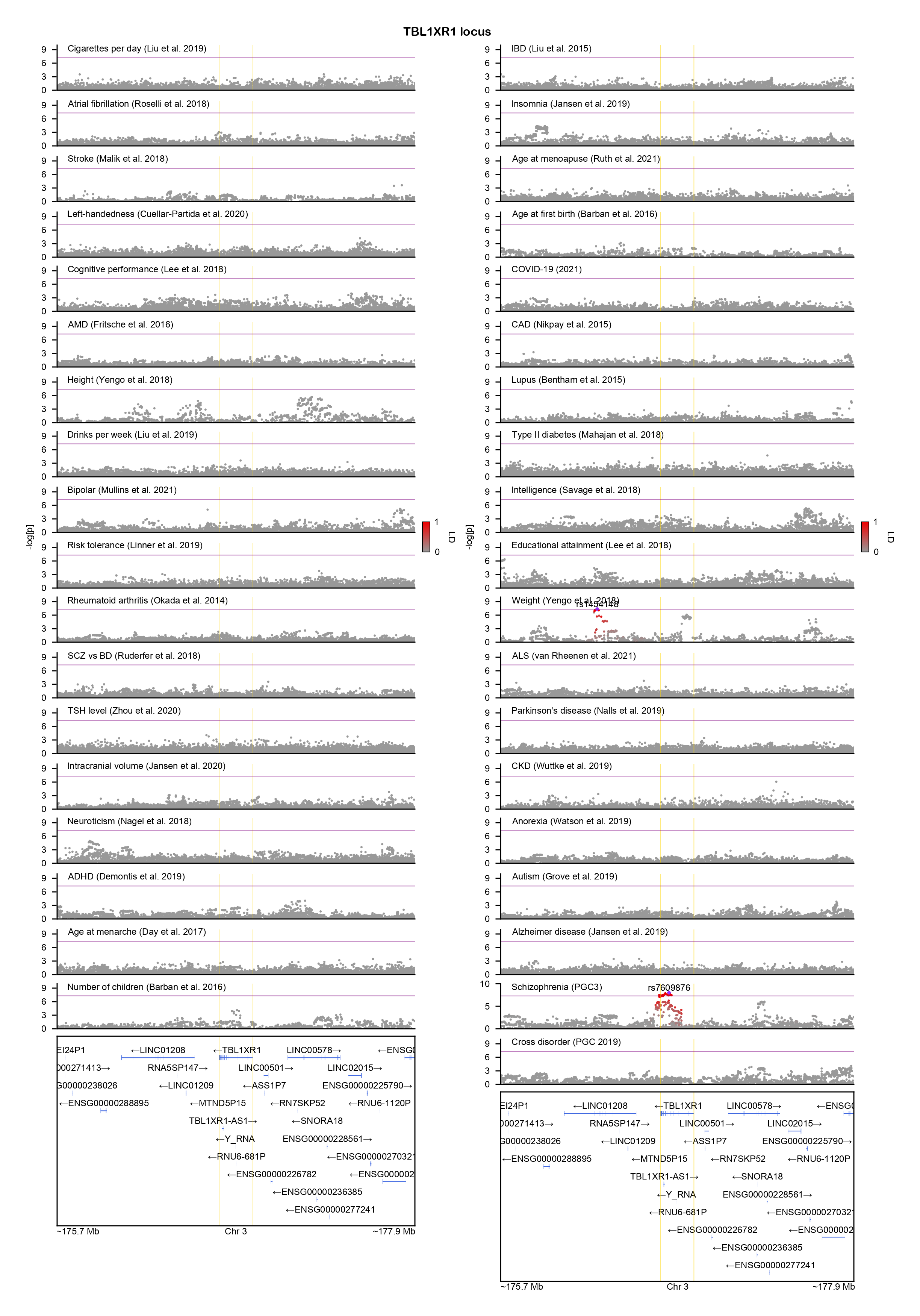

### THOC7-locuszoom.png

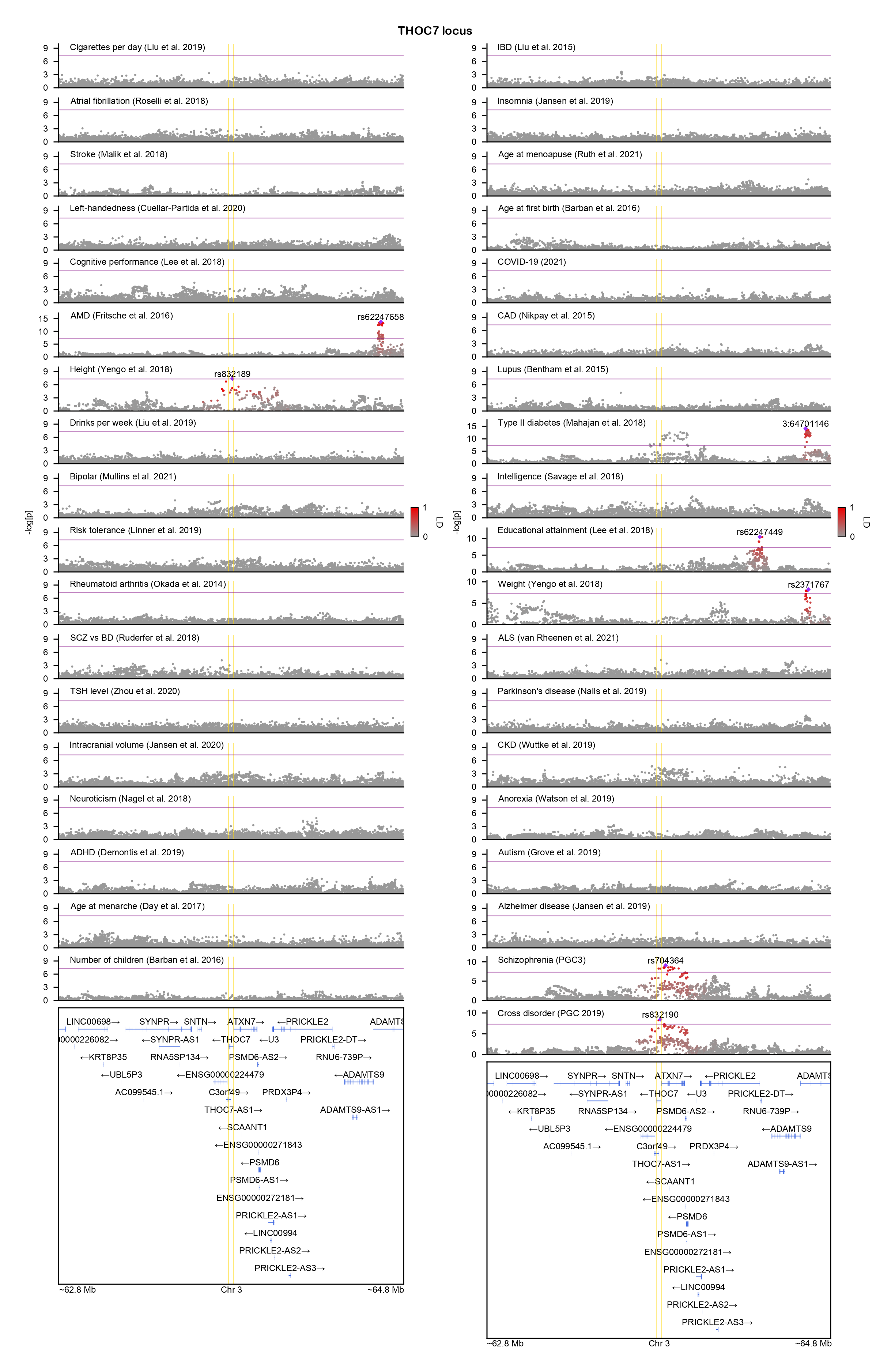

### TMCC3-locuszoom.png

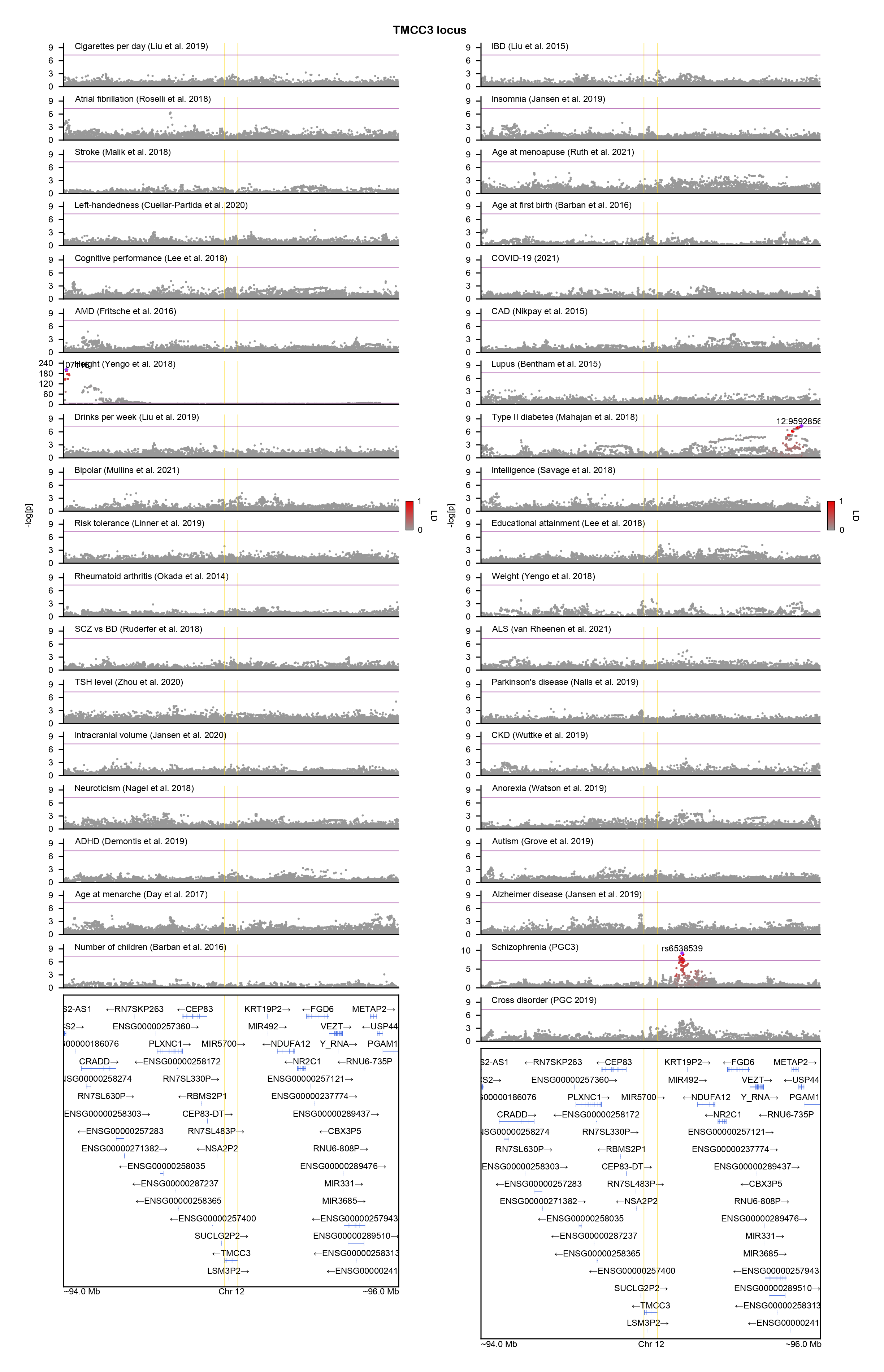

### TXNRD1-locuszoom.png

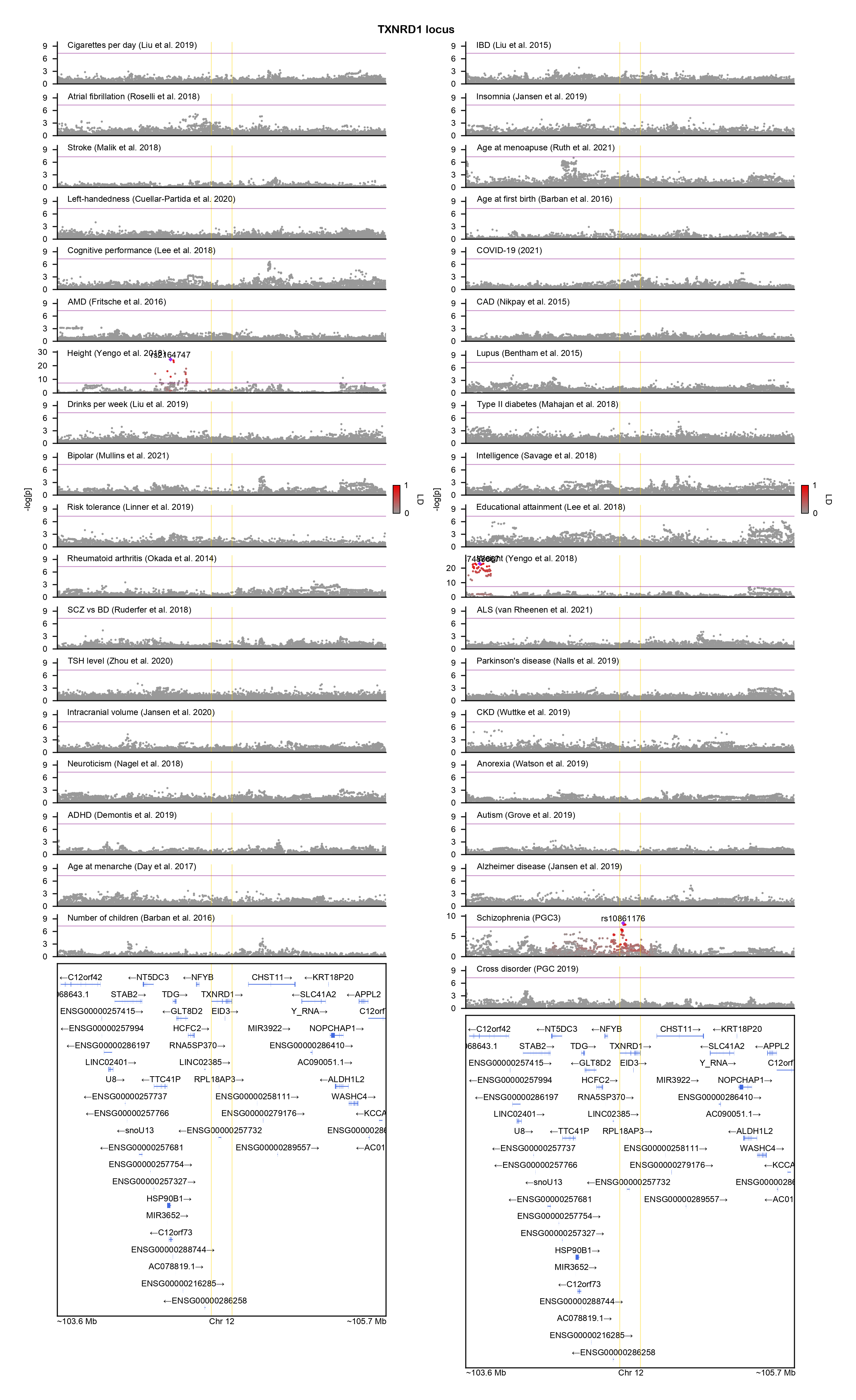

### VRK2-locuszoom.png

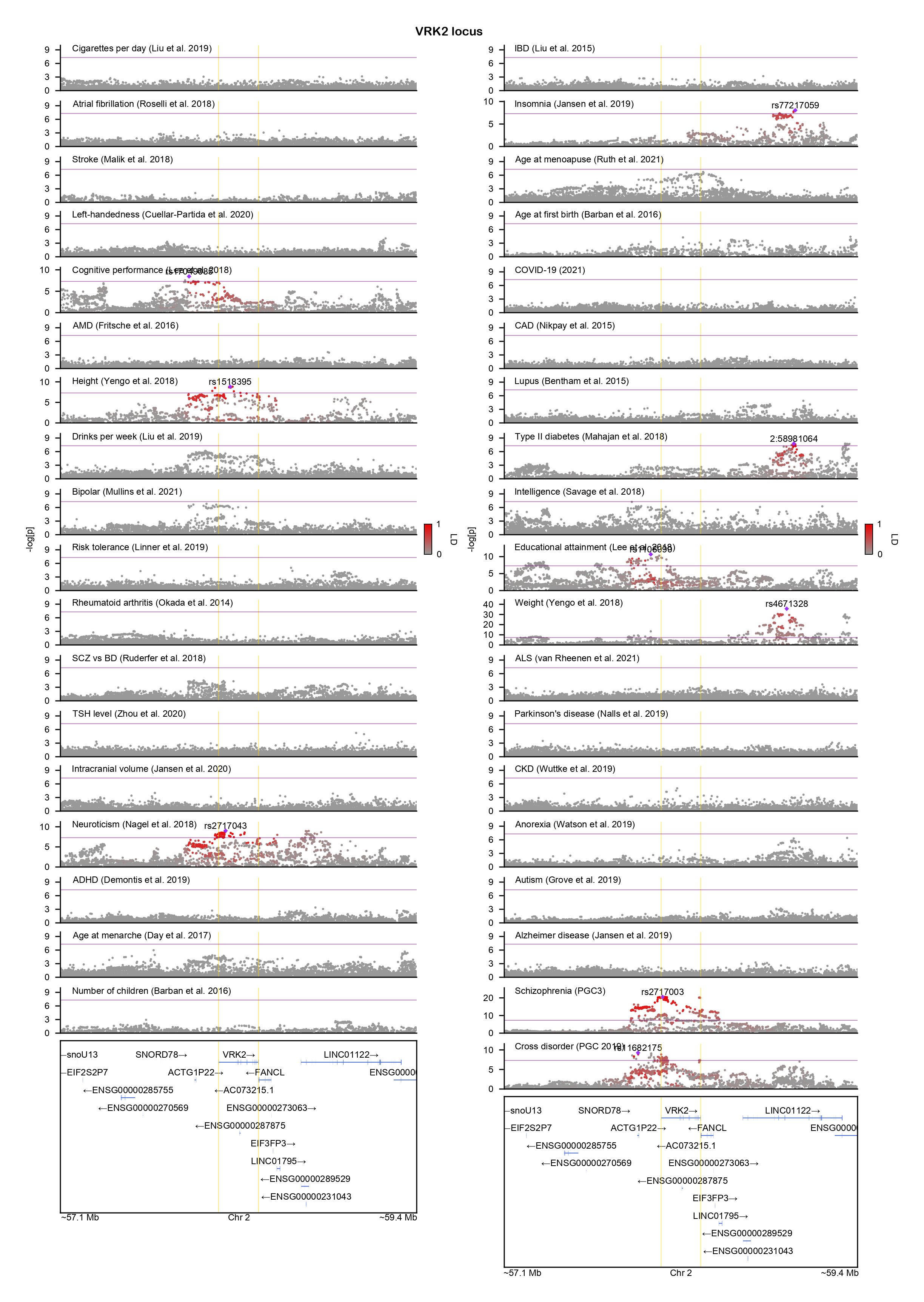

### WSCD2-locuszoom.png

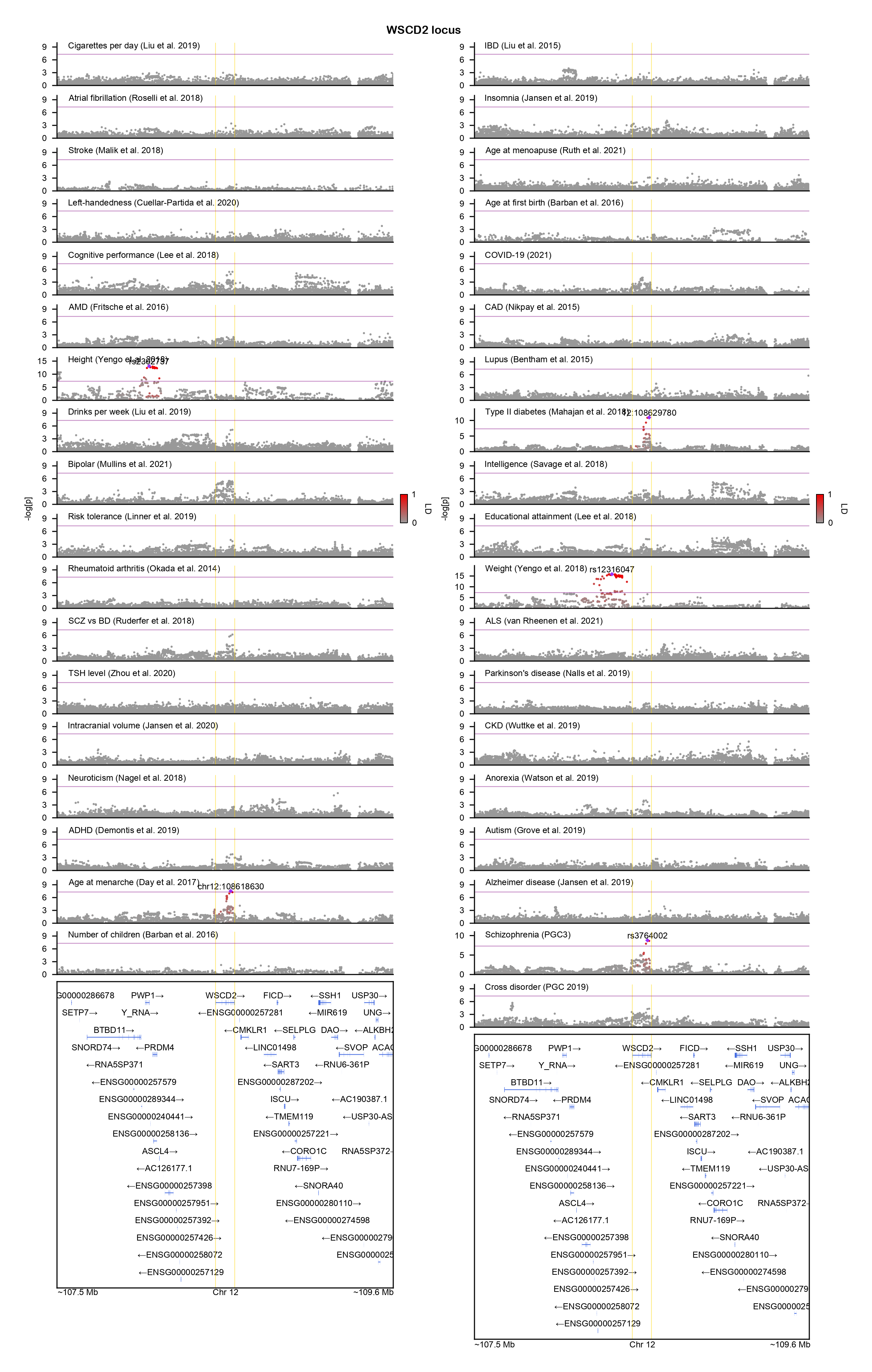

### ZEB2-locuszoom.png

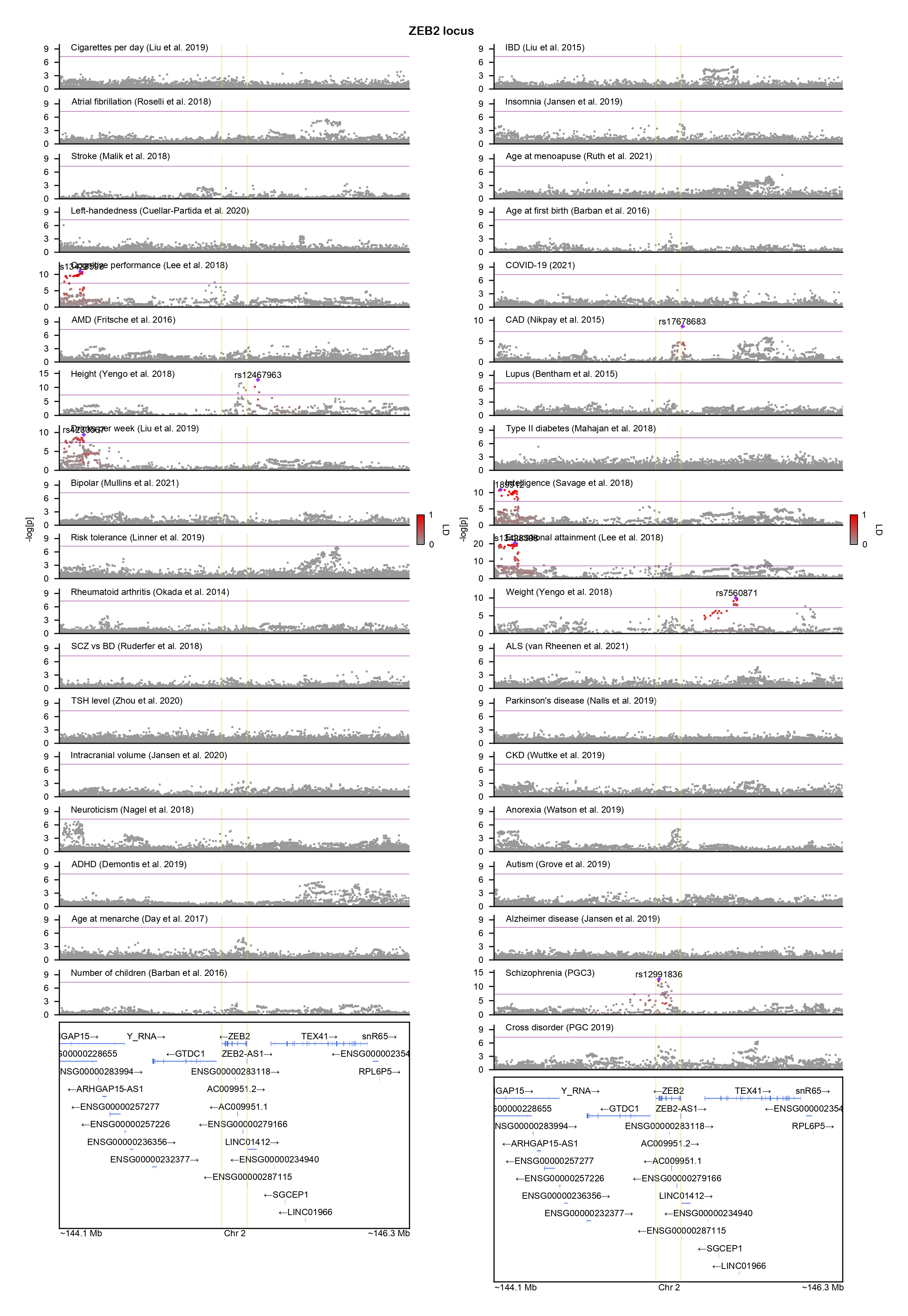

### ZNF281-locuszoom.png

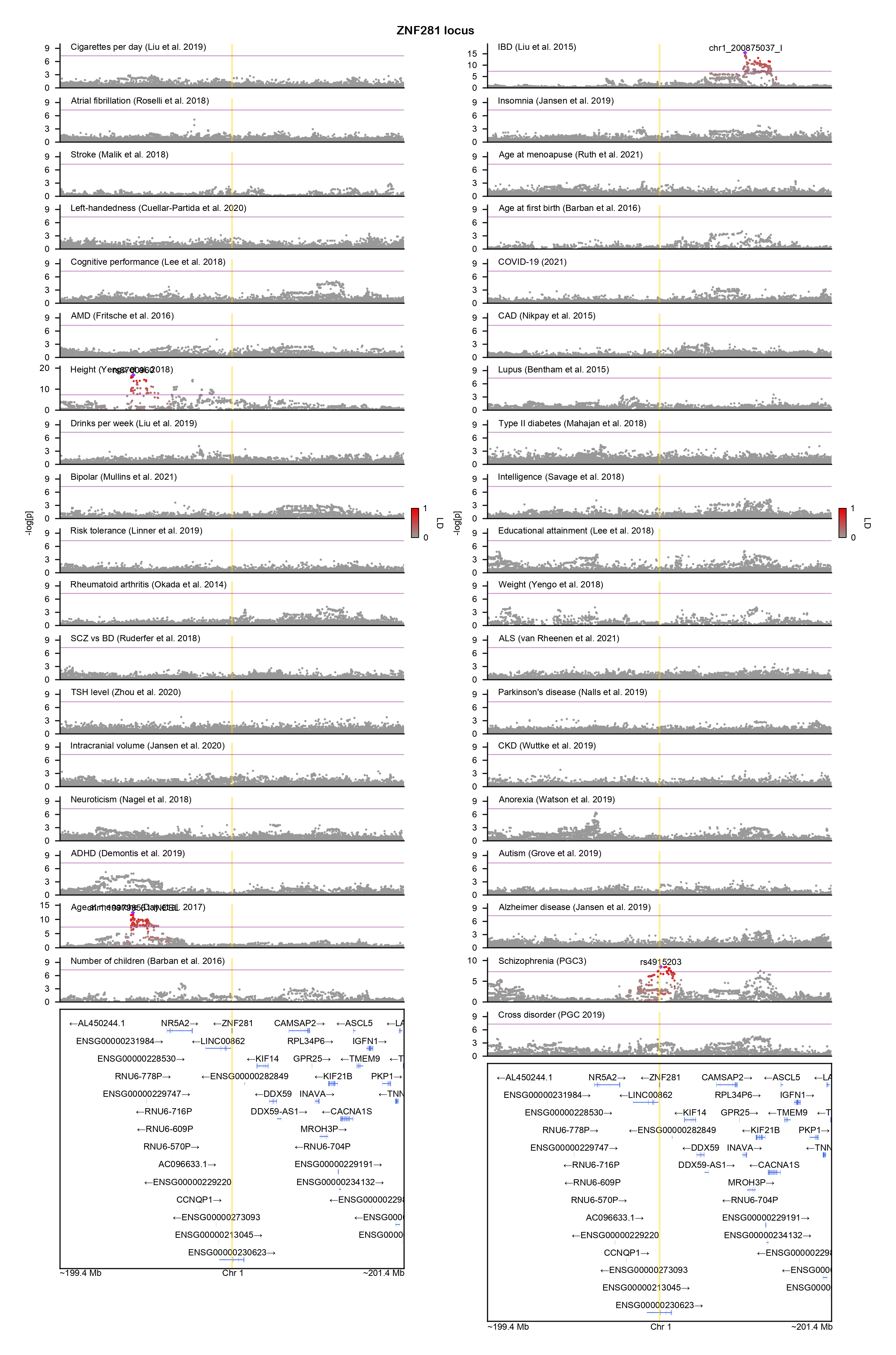

### ZSCAN2-locuszoom.png

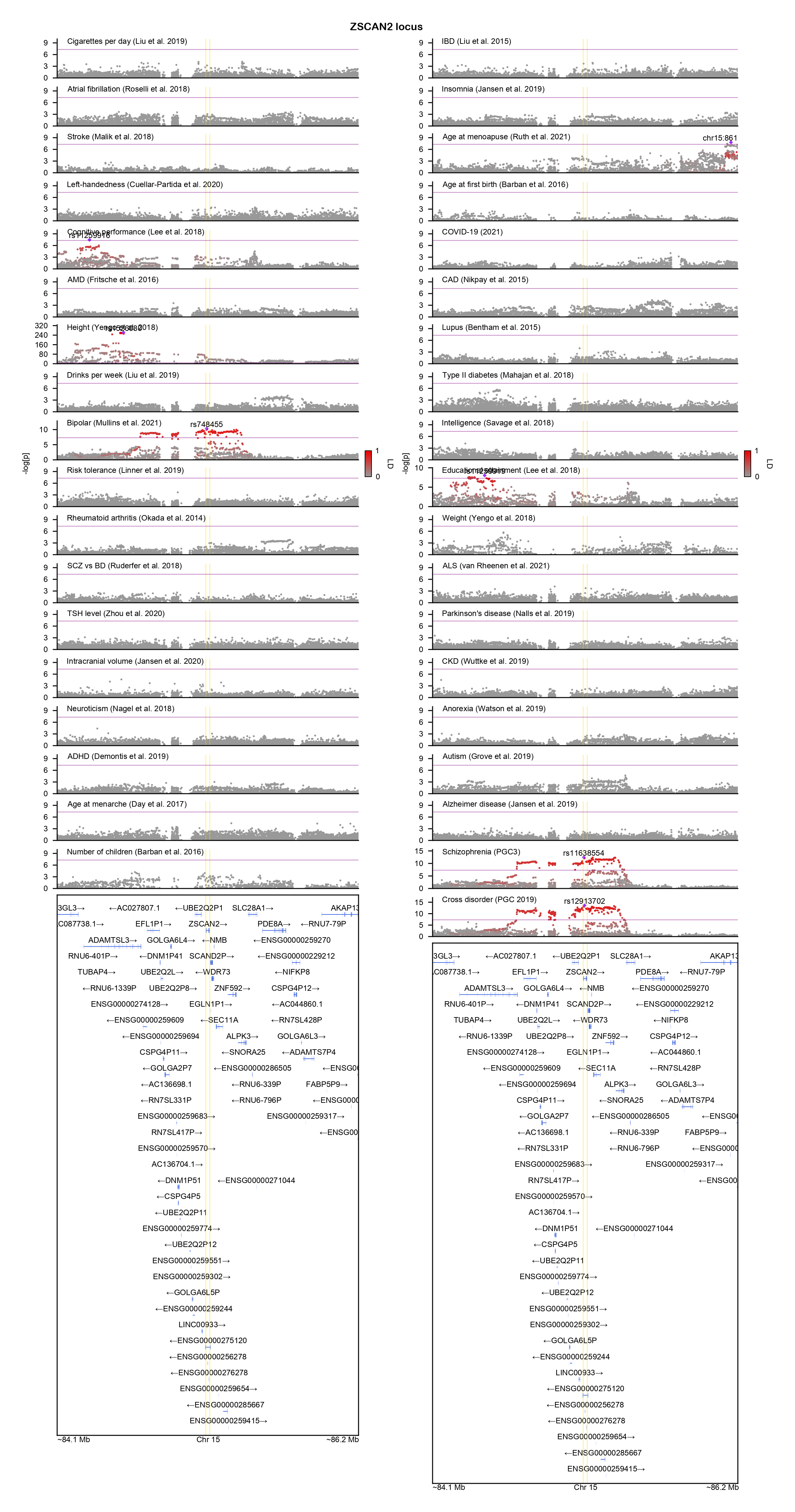

### ZSWIM6-locuszoom.png

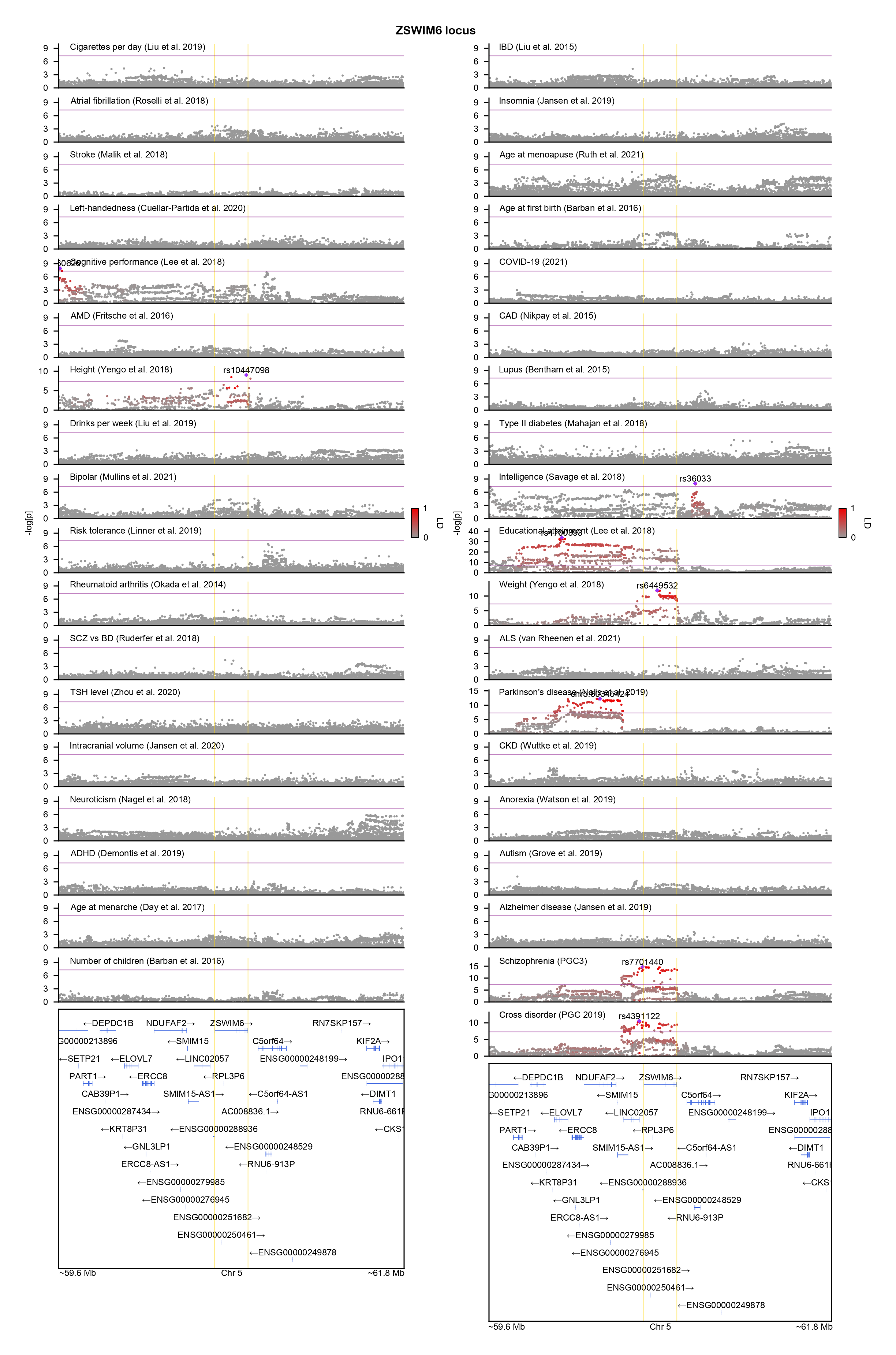
