## Supplementary figures and images for "GeneticsMakie.jl: A versatile and scalable toolkit for visualizing locus-level genetic and genomic data"

### AUTS2-locuszoom.png

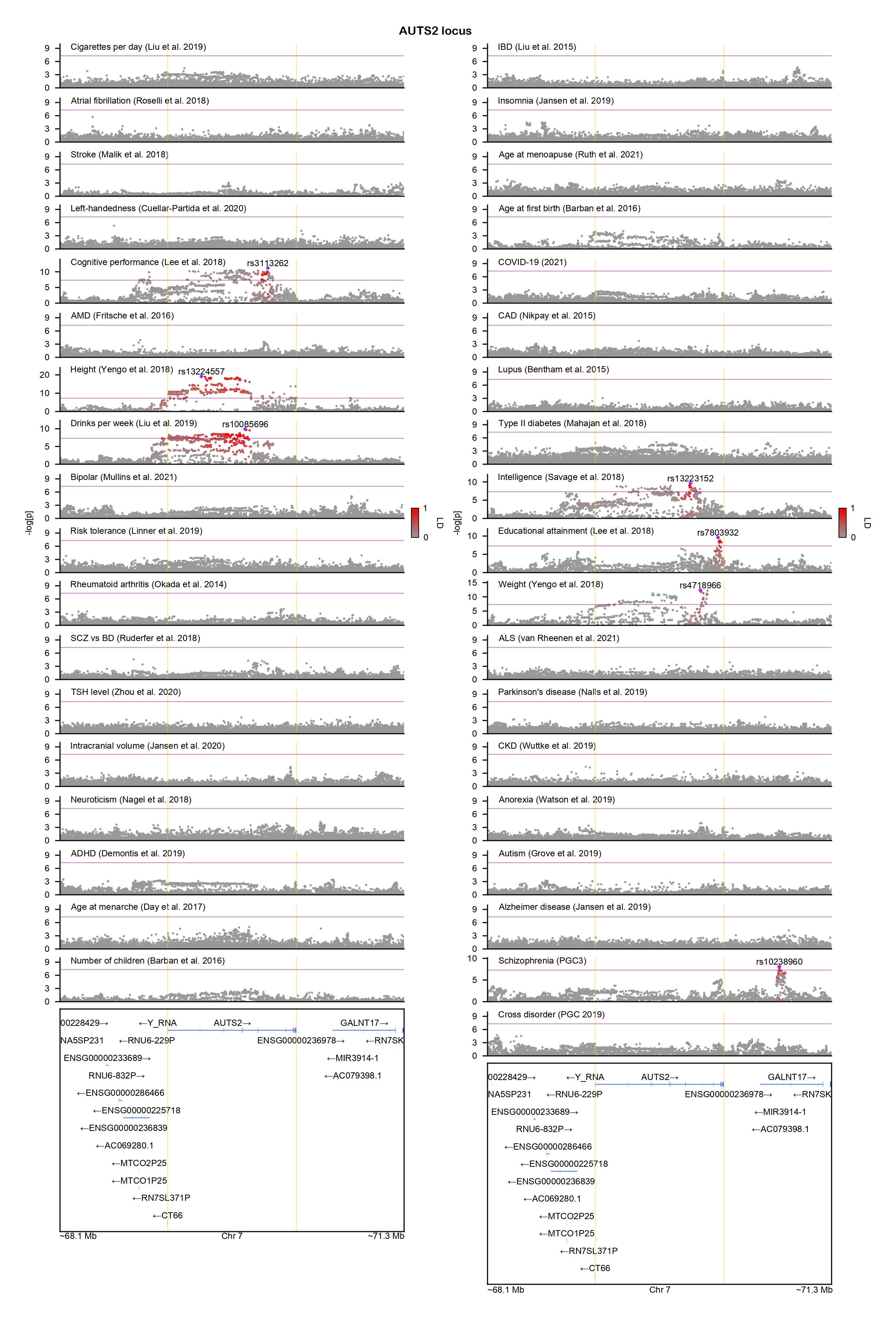

### CERT1-locuszoom.png

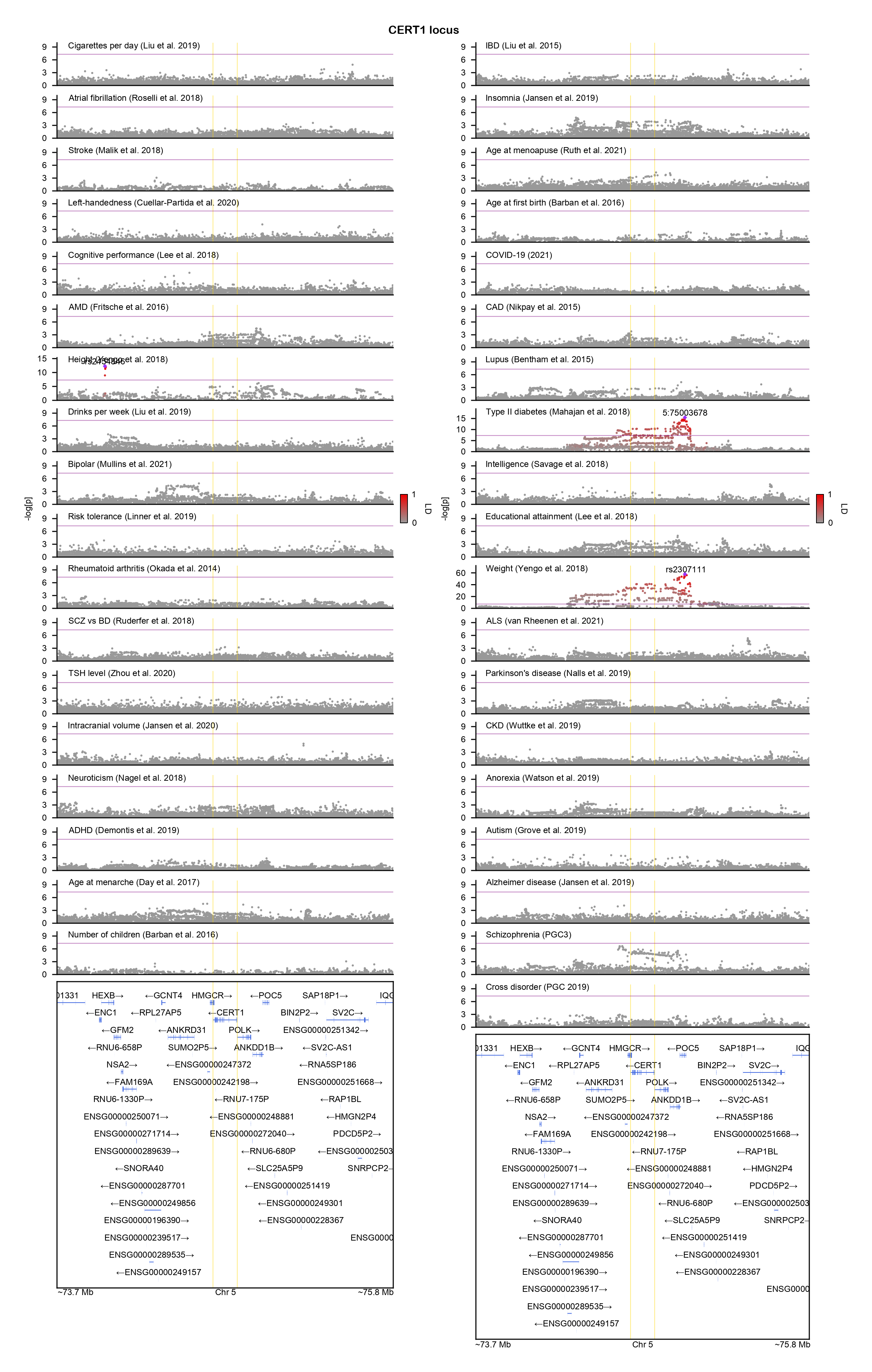

### EFTUD2-locuszoom.png

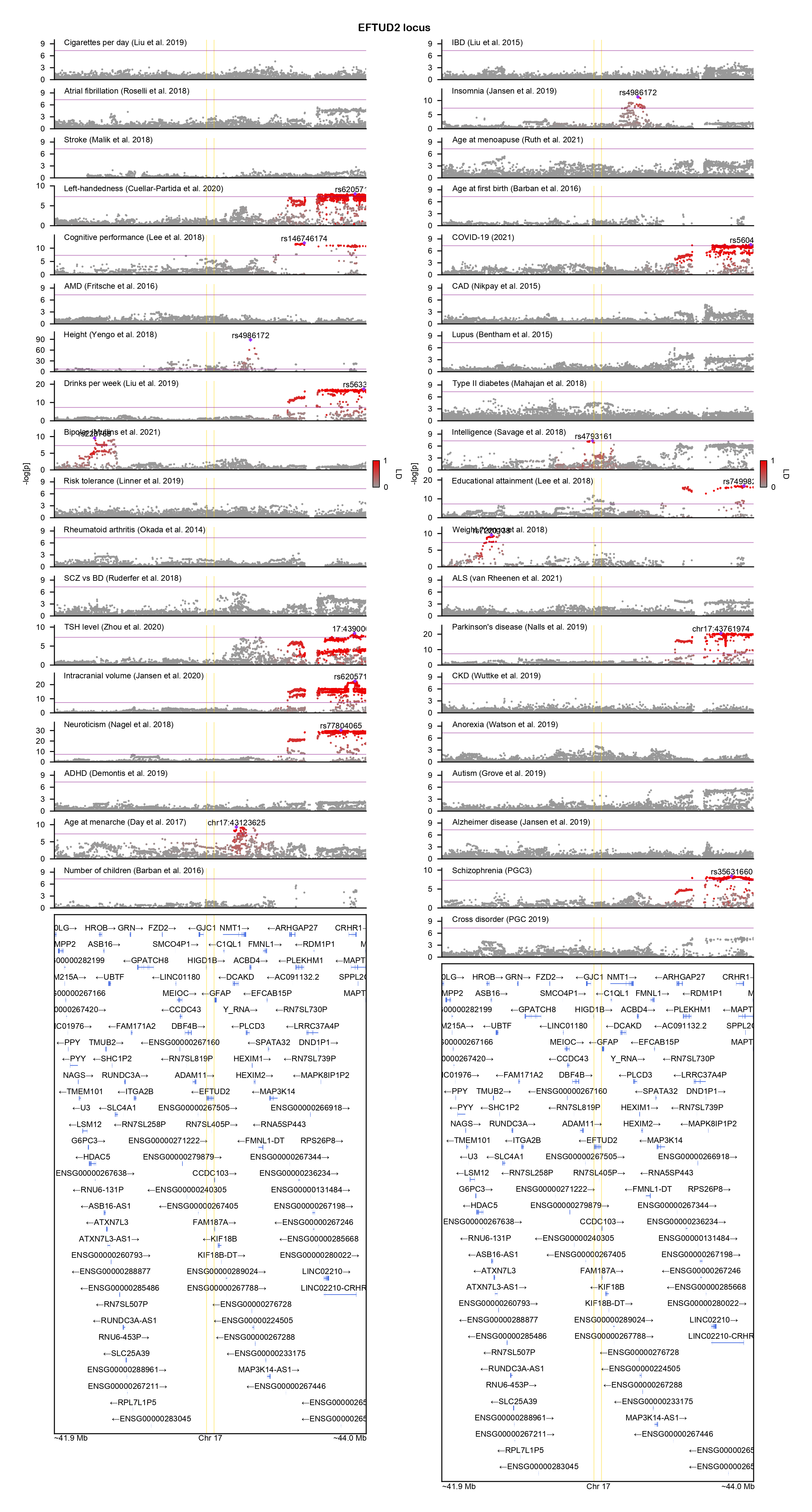

### GABRB2-locuszoom.png

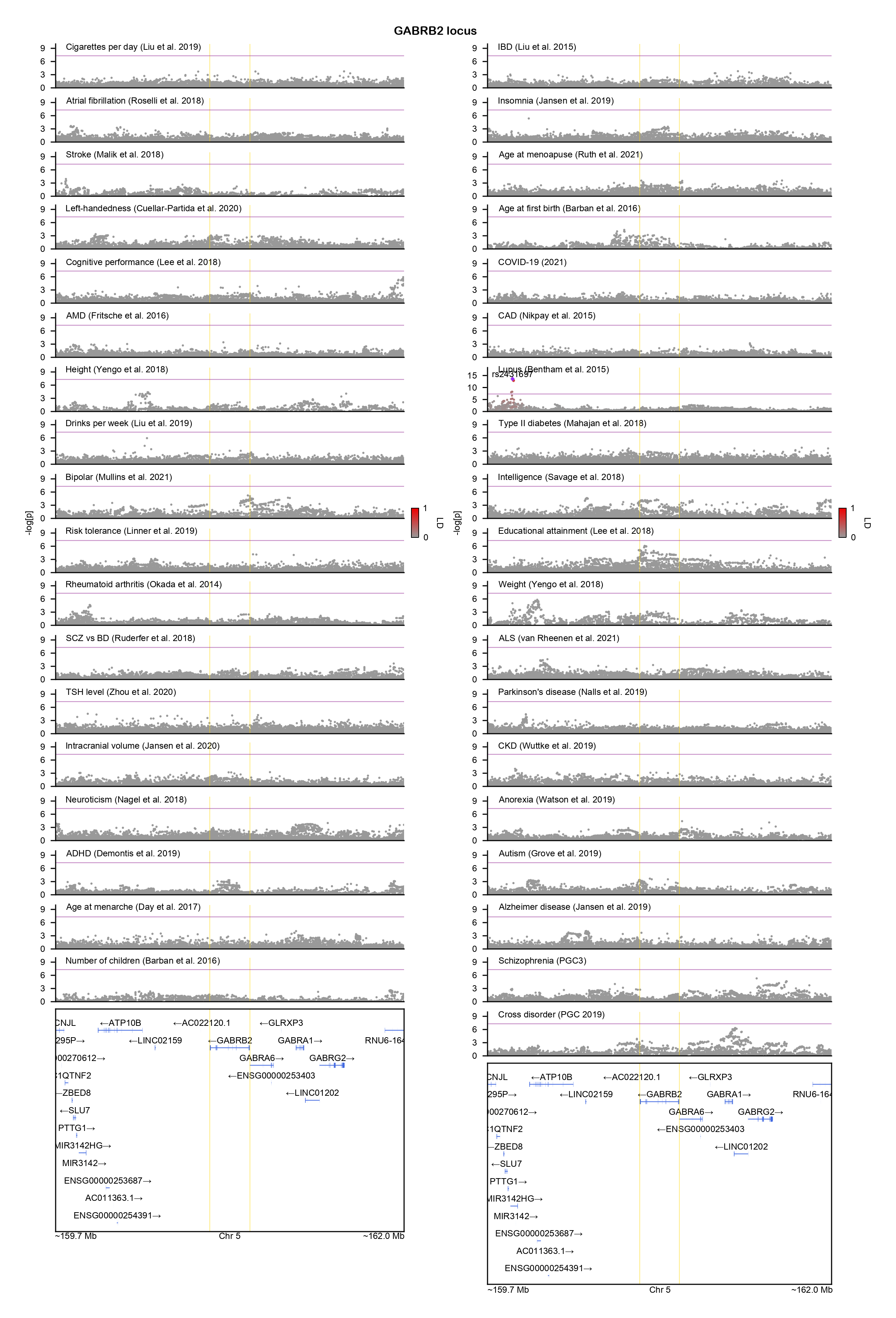

### ITPR1-locuszoom.png

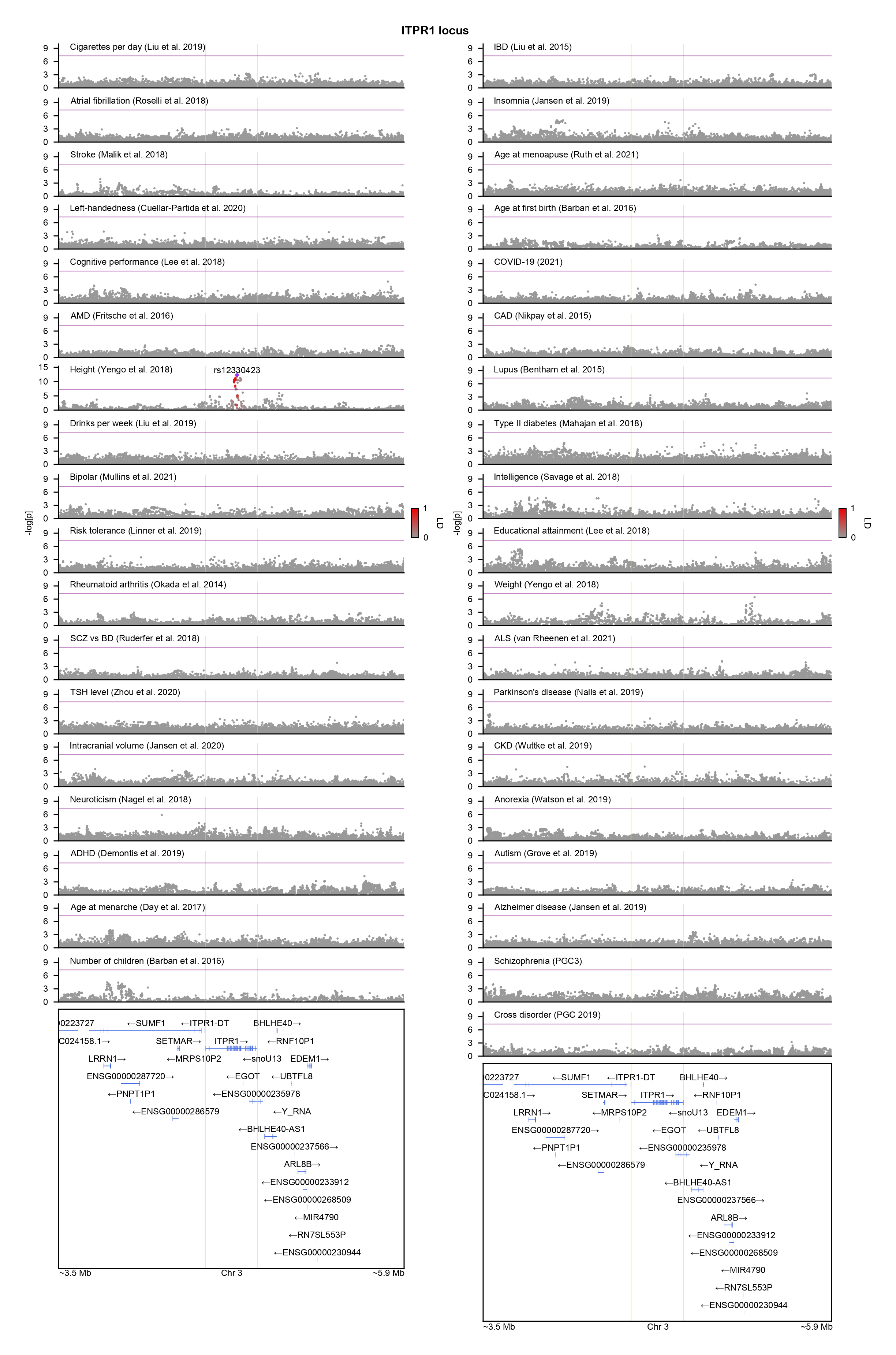

### KCNT1-locuszoom.png

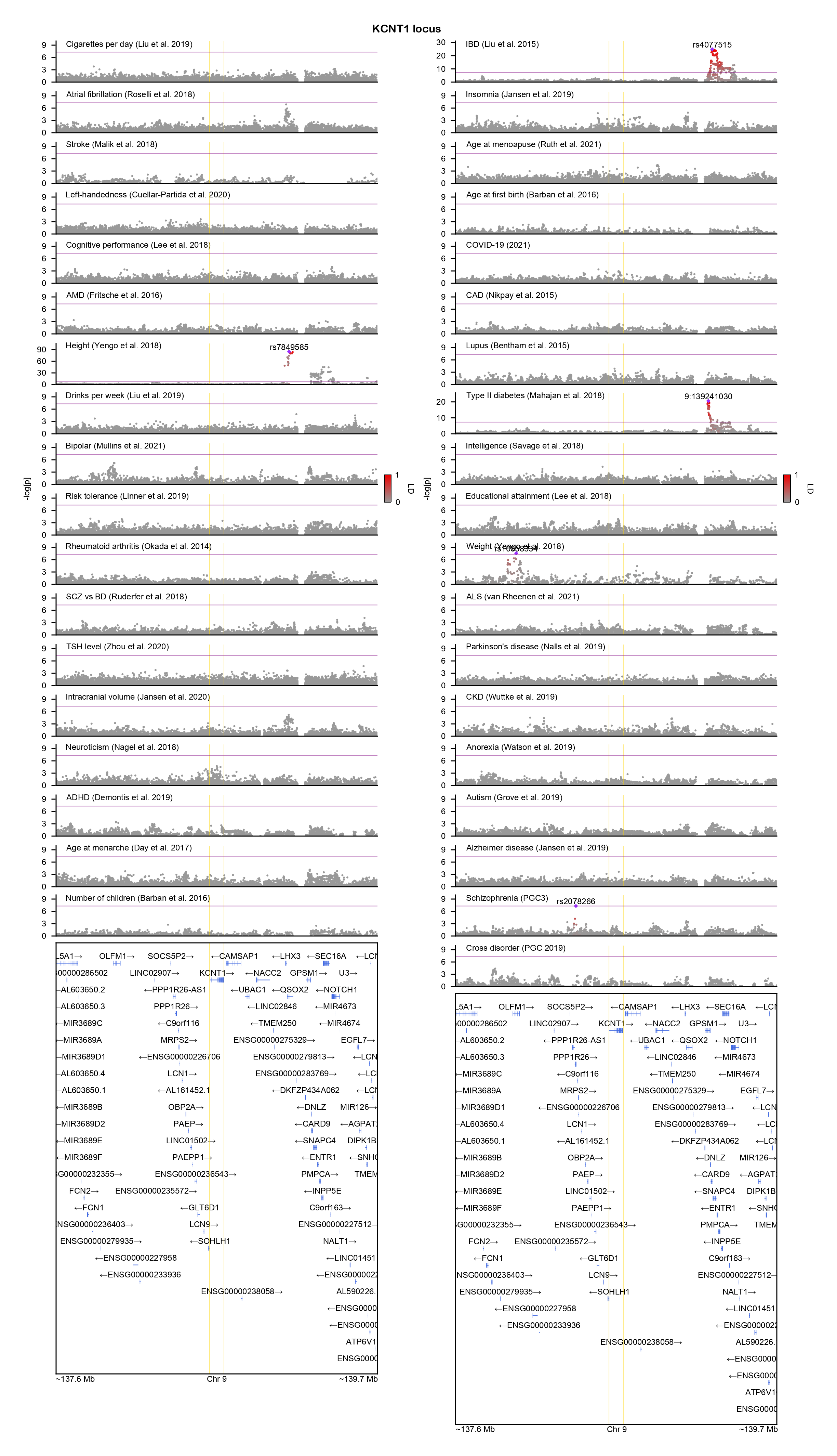

### KMT2C-locuszoom.png

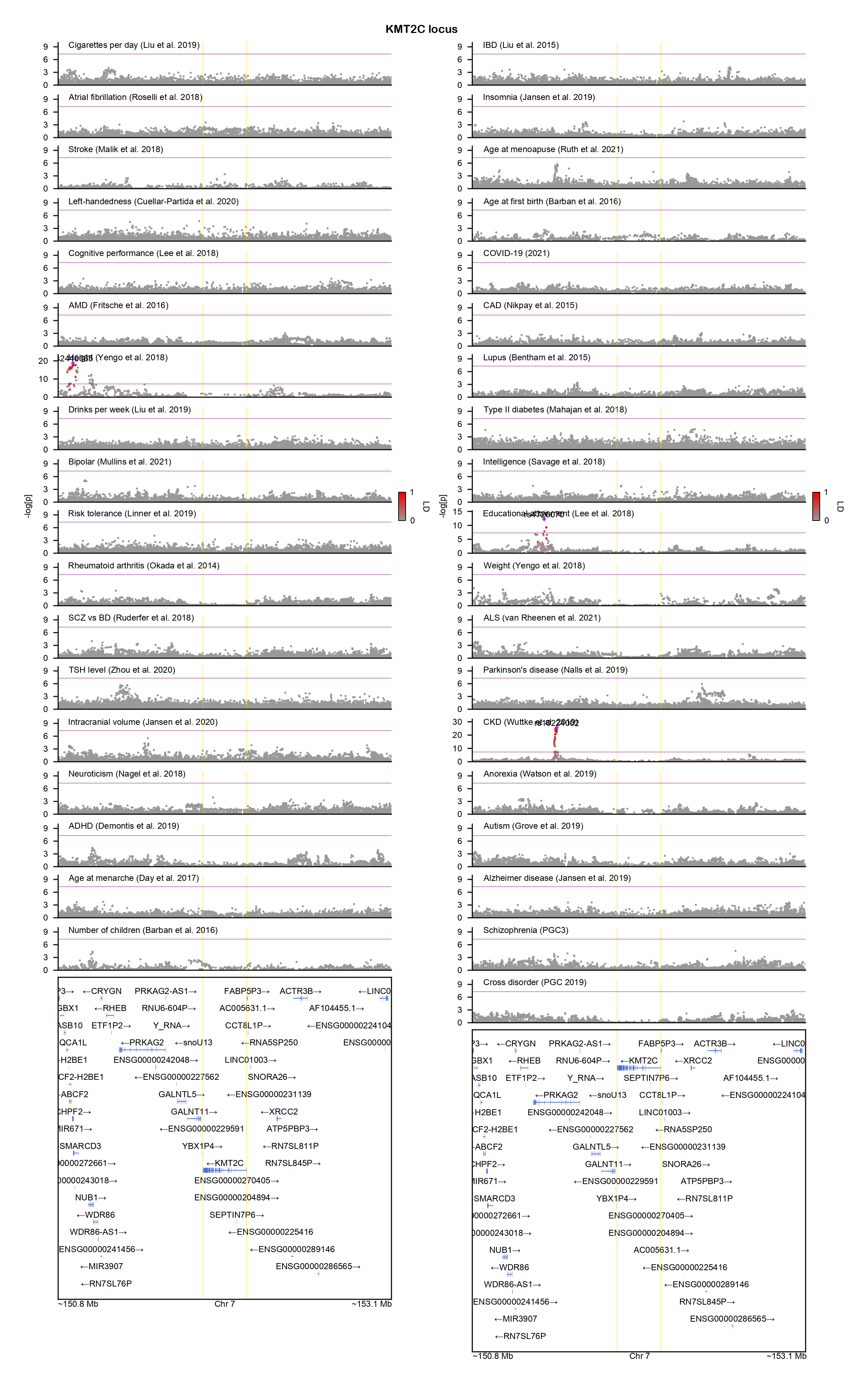

### LDB1-locuszoom.png

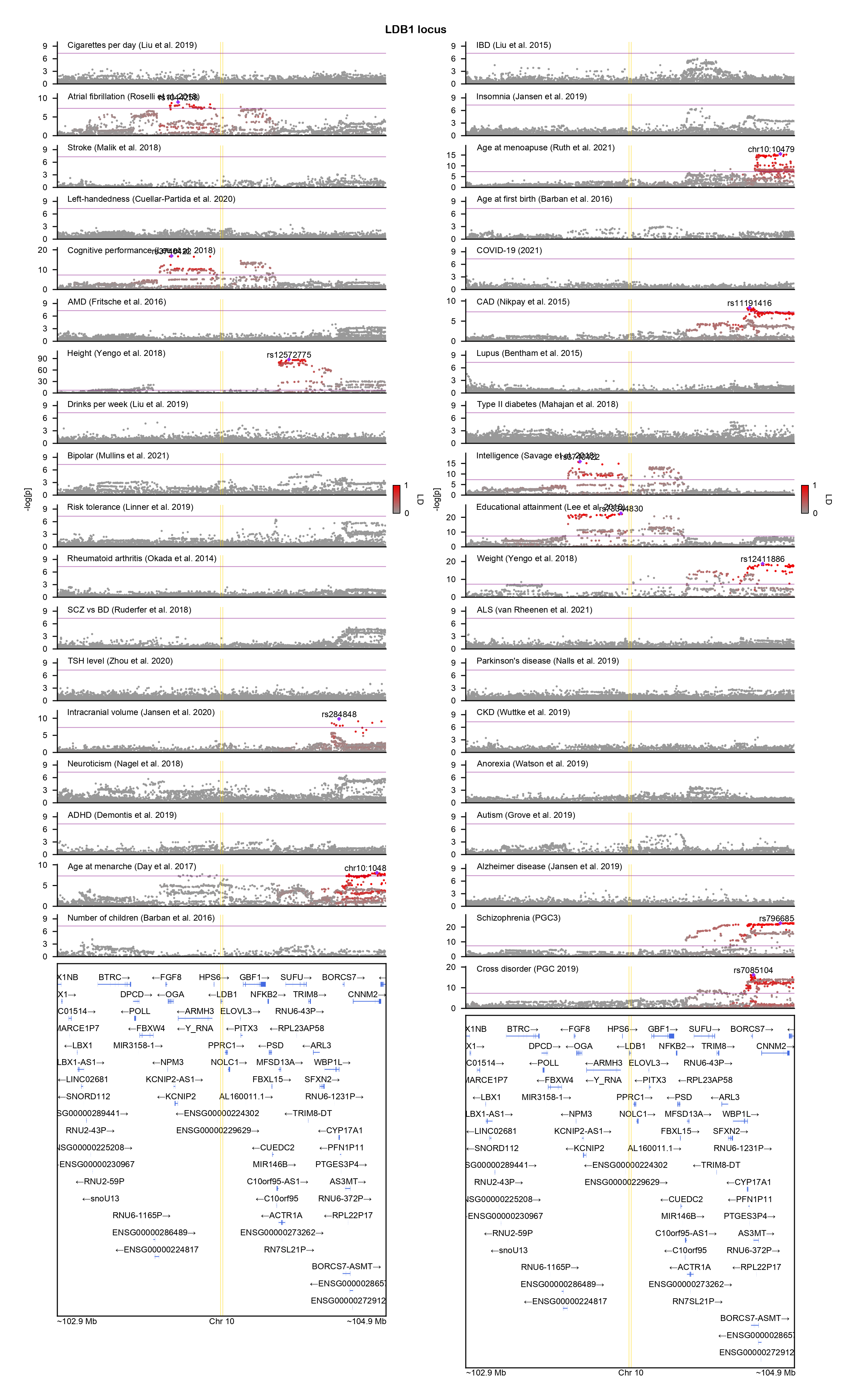

### MN1-locuszoom.png

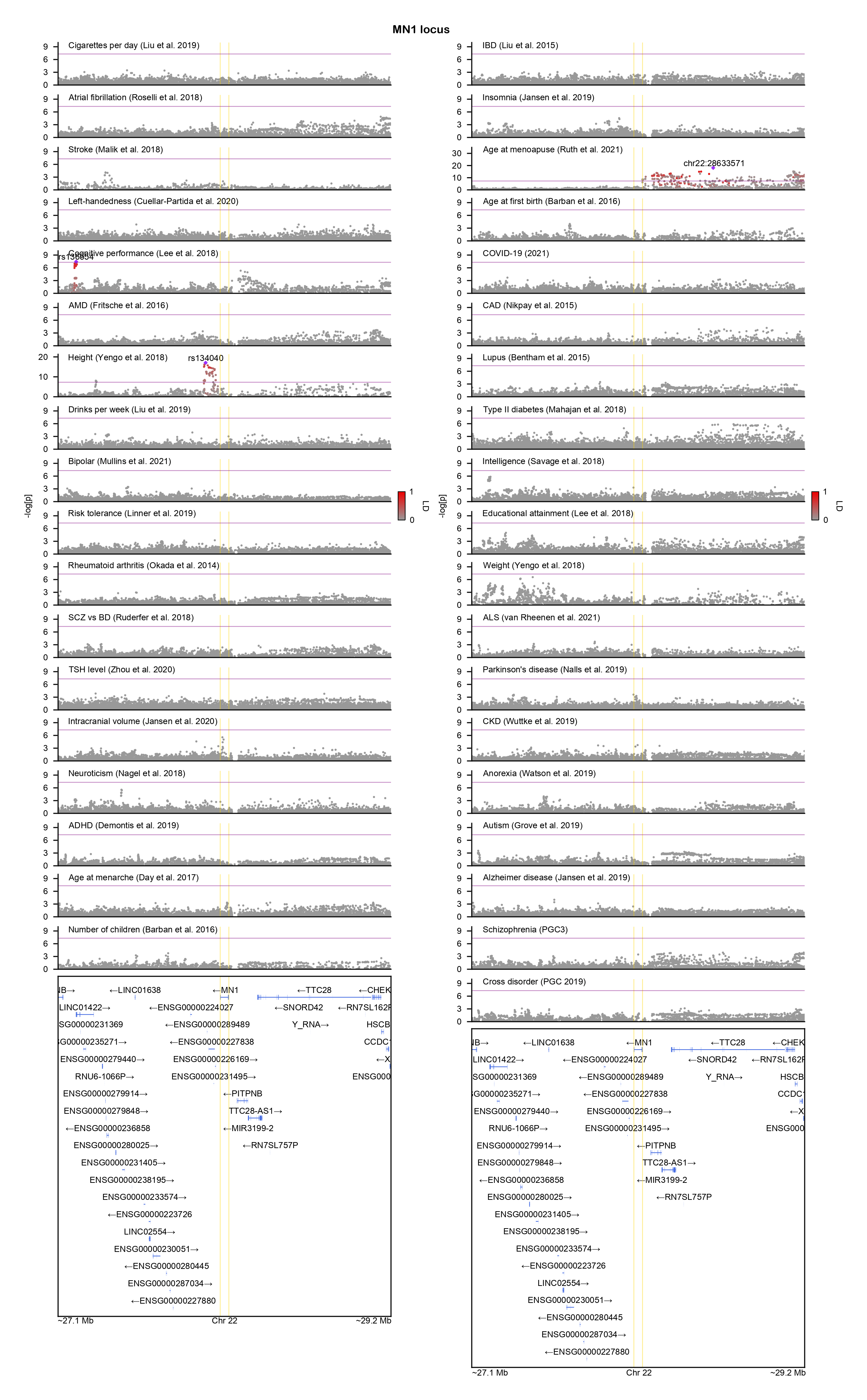

### PSMC5-locuszoom.png

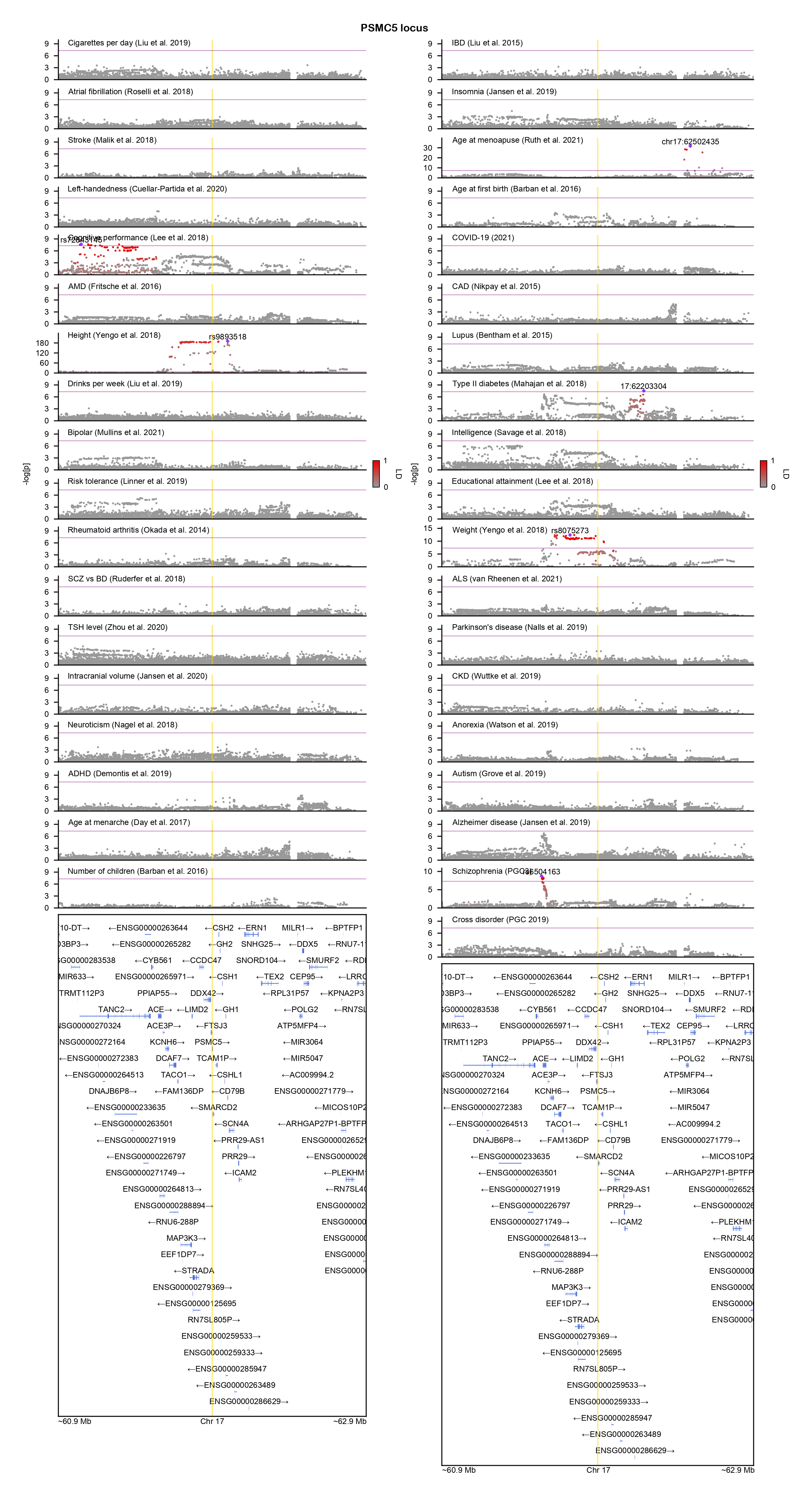

### SPAST-locuszoom.png

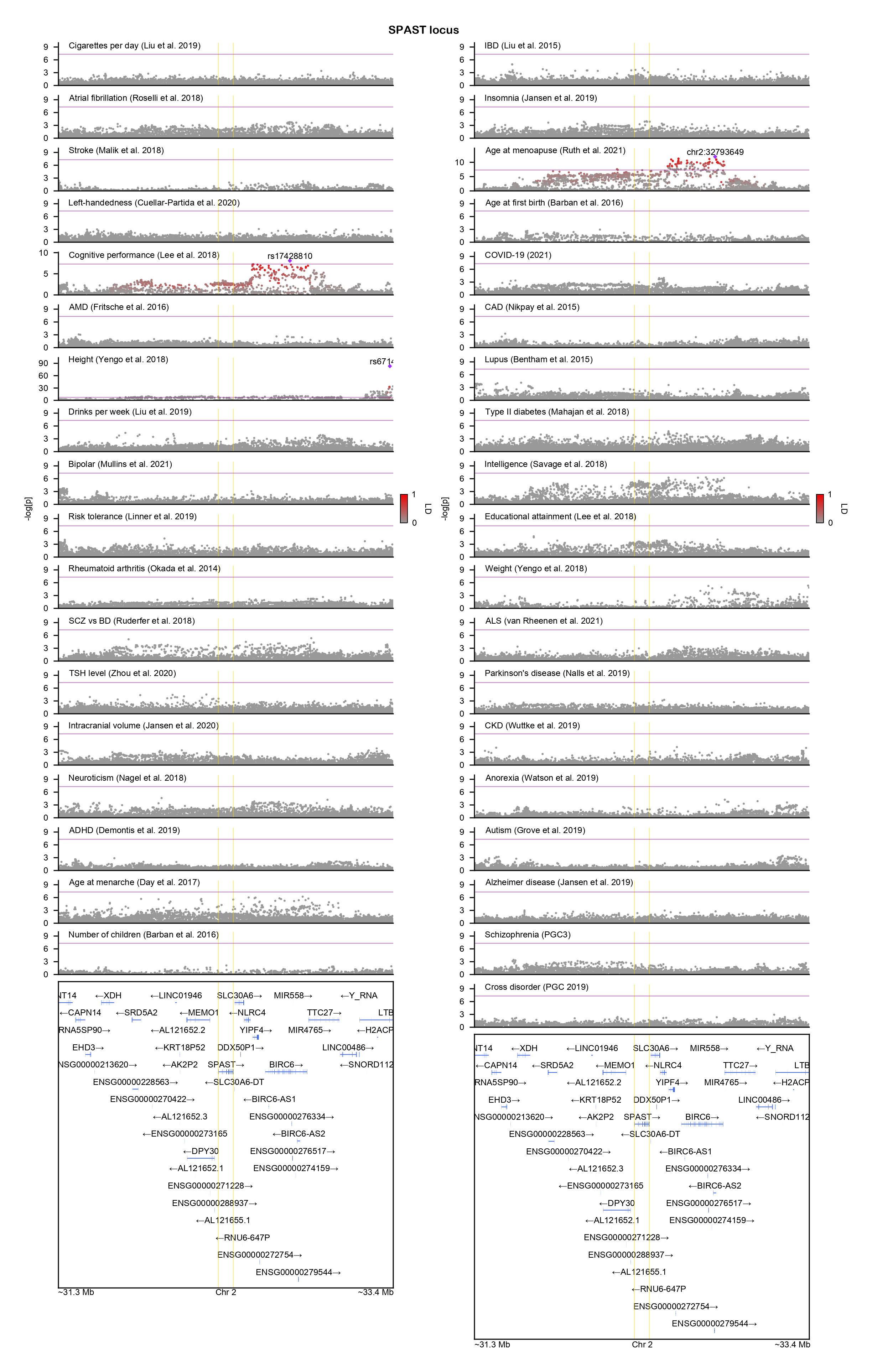

### TBR1-locuszoom.png

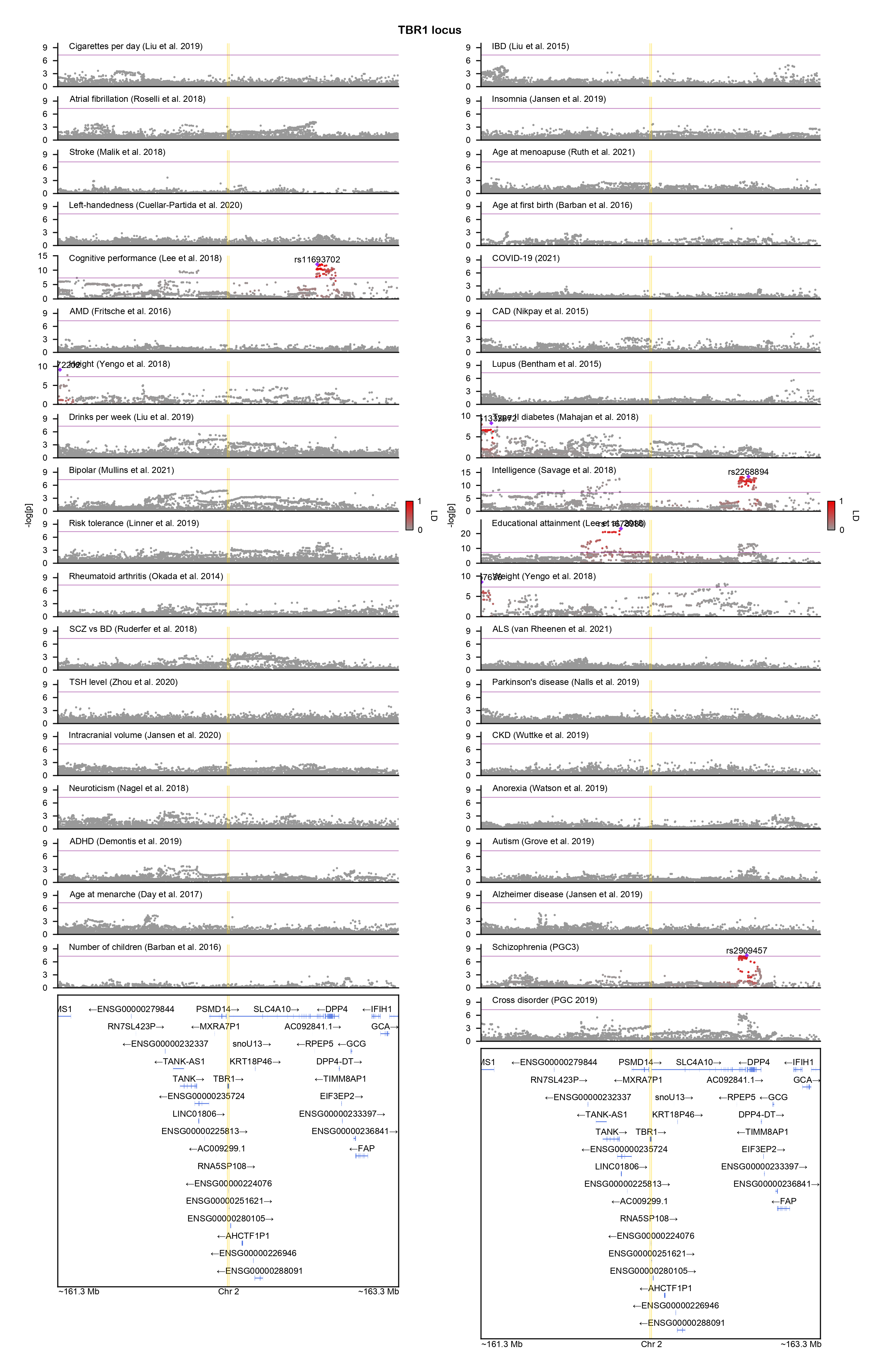

### TCF7L2-locuszoom.png

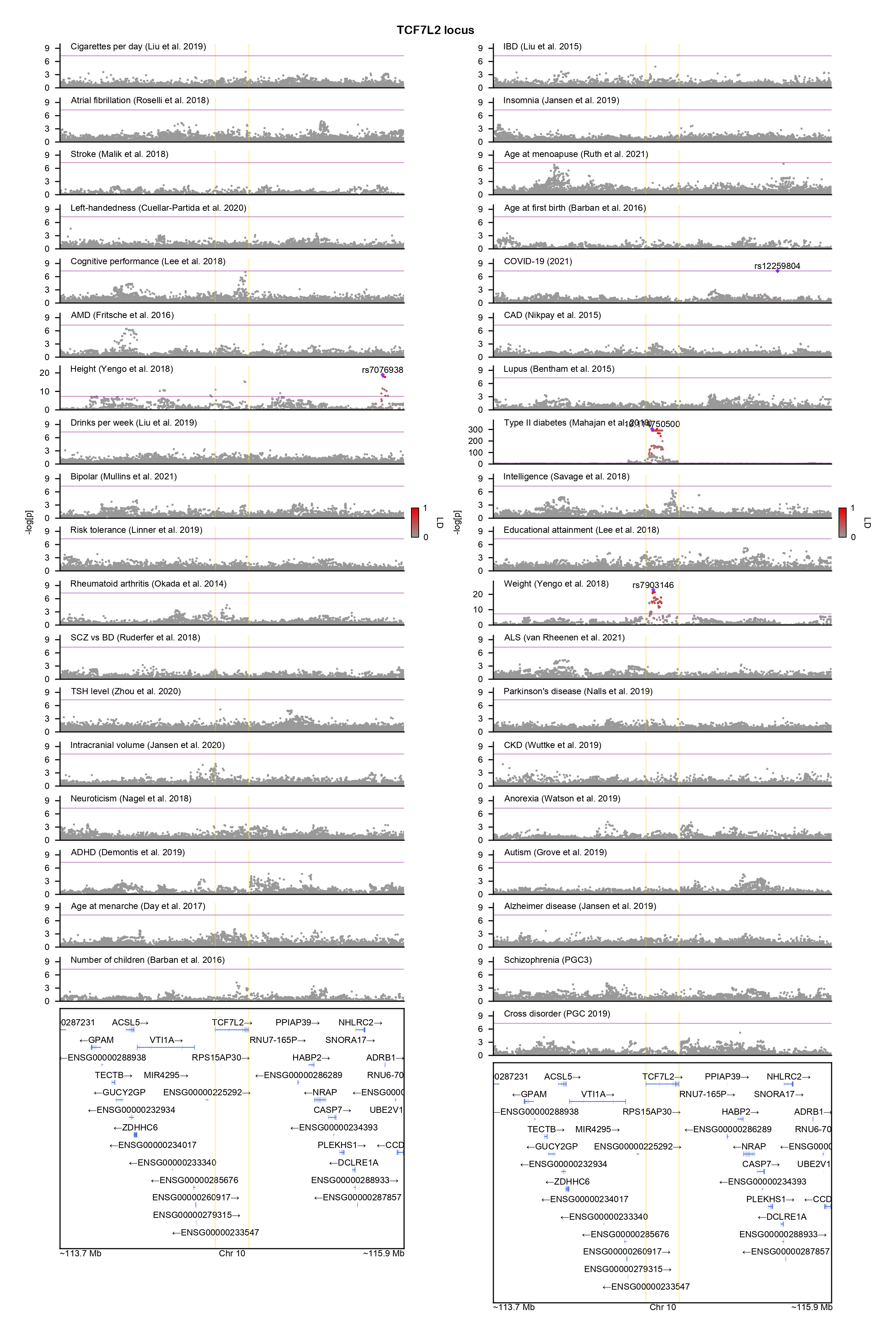
