## Supplementary figures and images for "GeneticsMakie.jl: A versatile and scalable toolkit for visualizing locus-level genetic and genomic data"

### ABL1-locuszoom.png

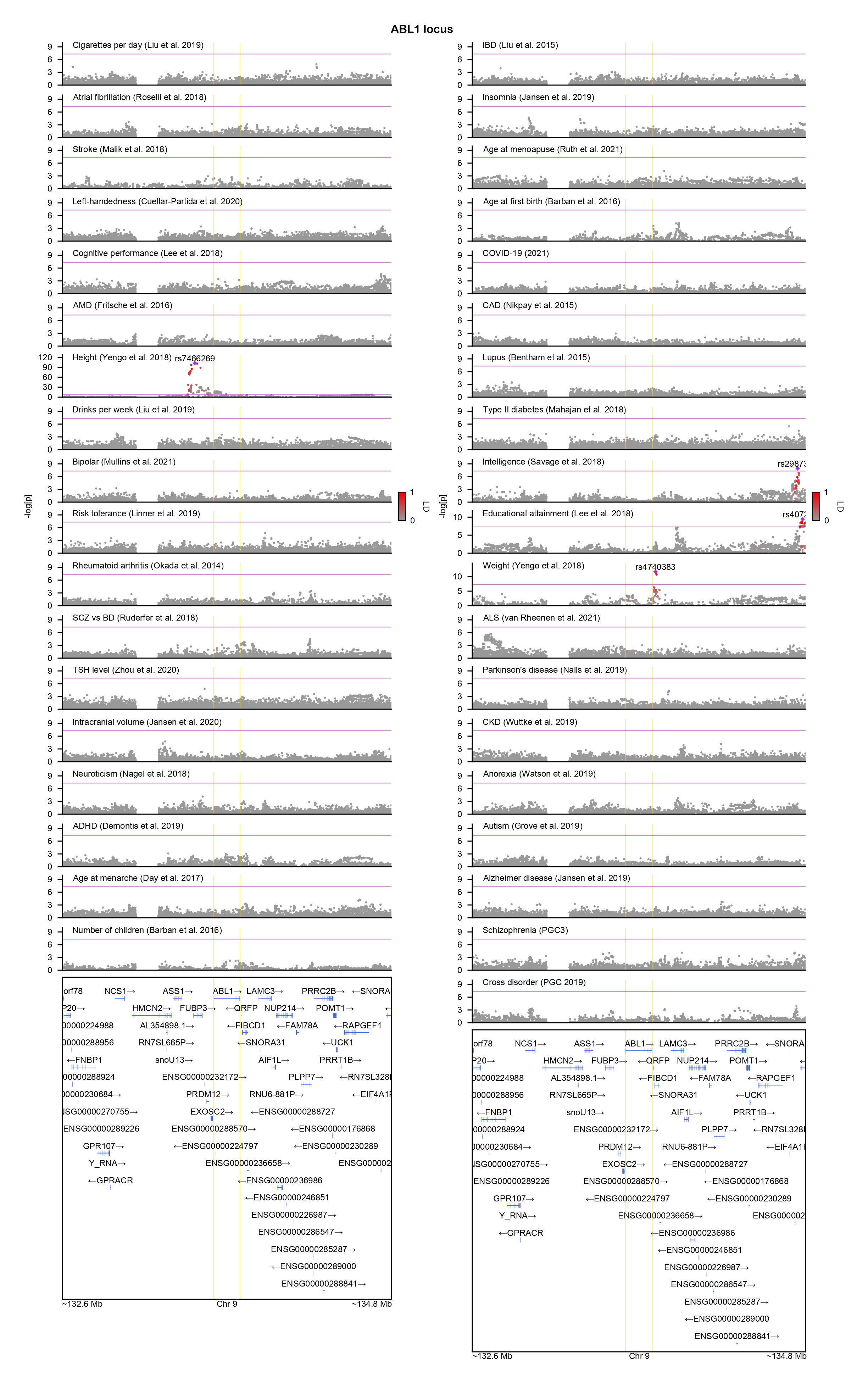

### ADCY5-locuszoom.png

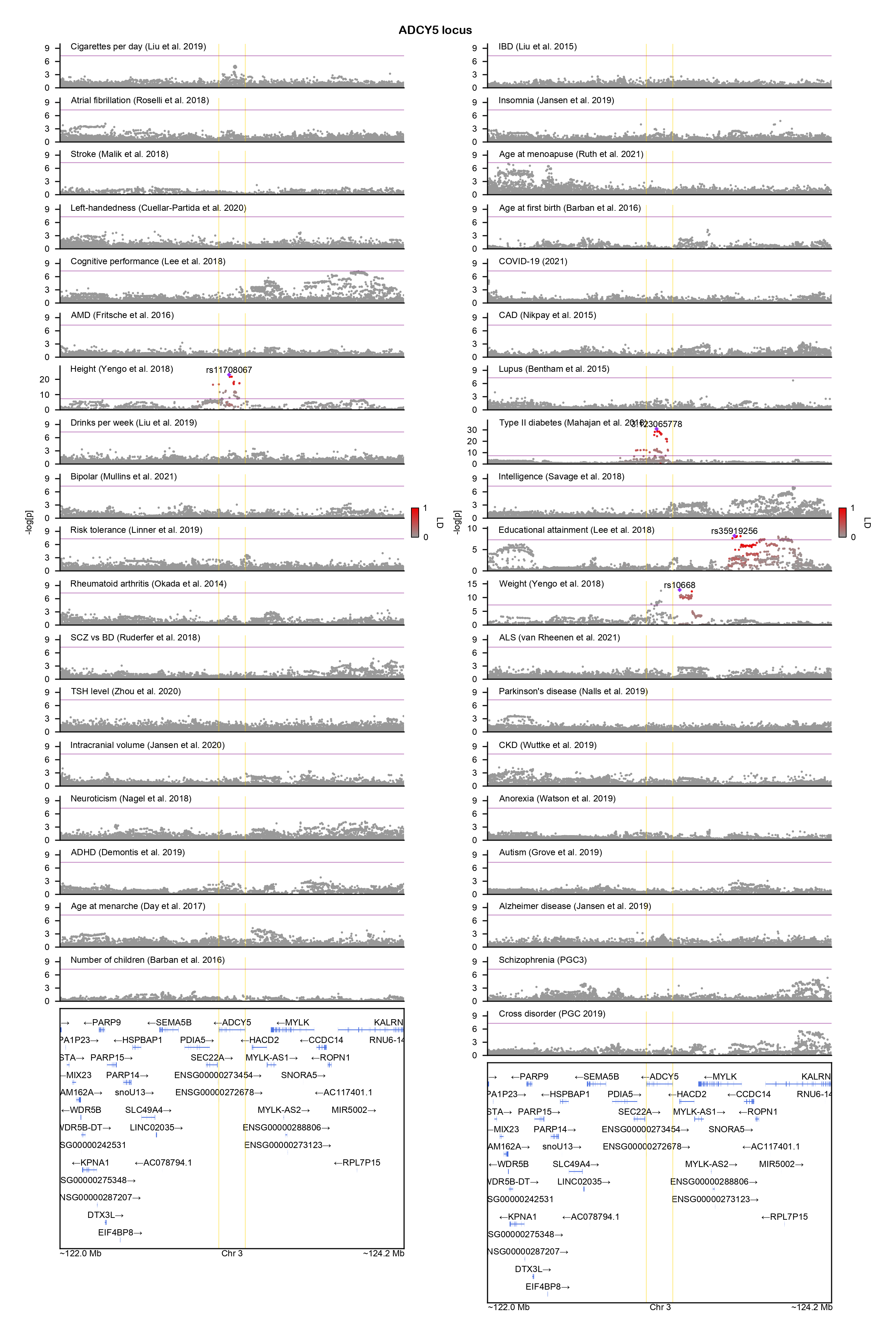

### AFF3-locuszoom.png

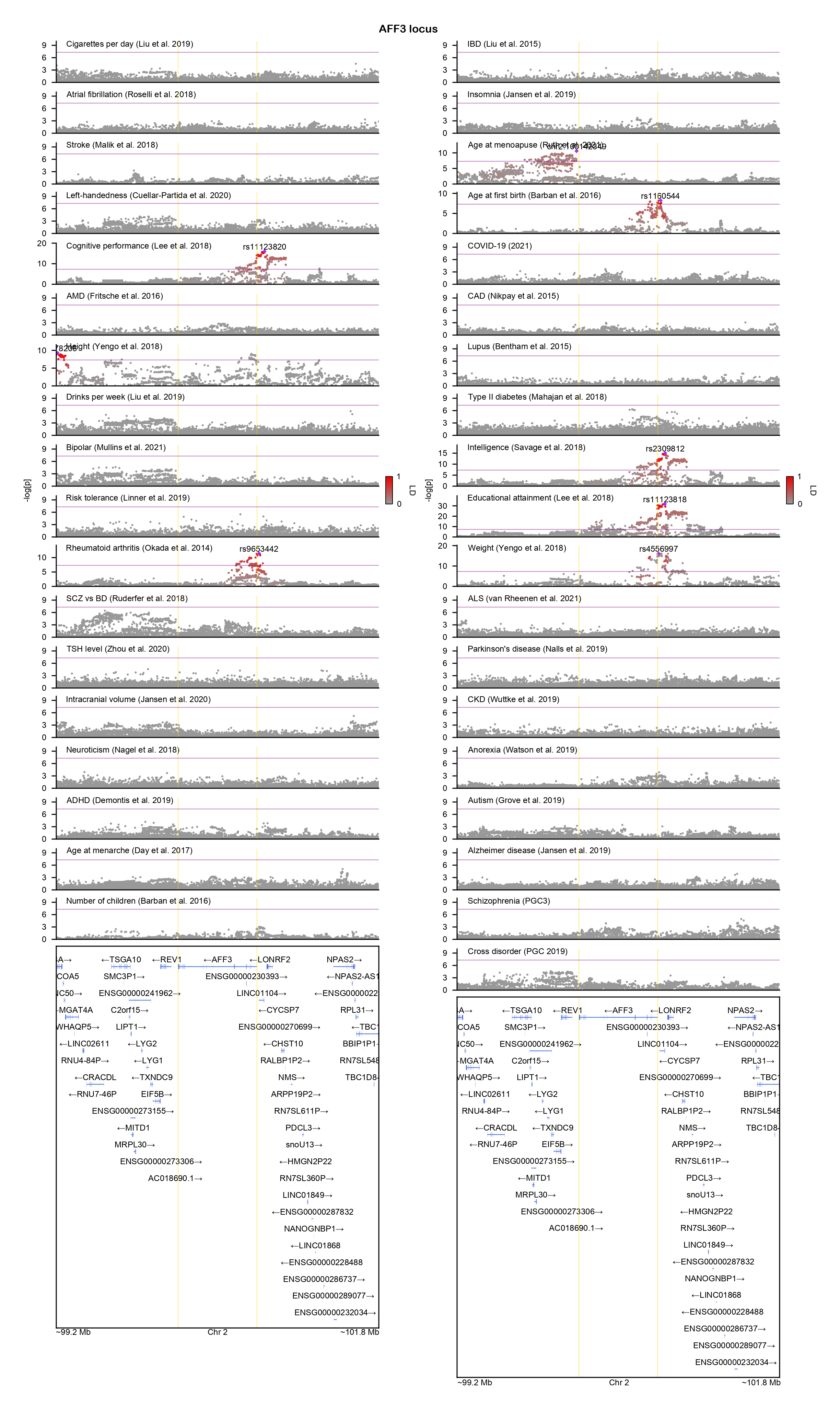

### AHDC1-locuszoom.png

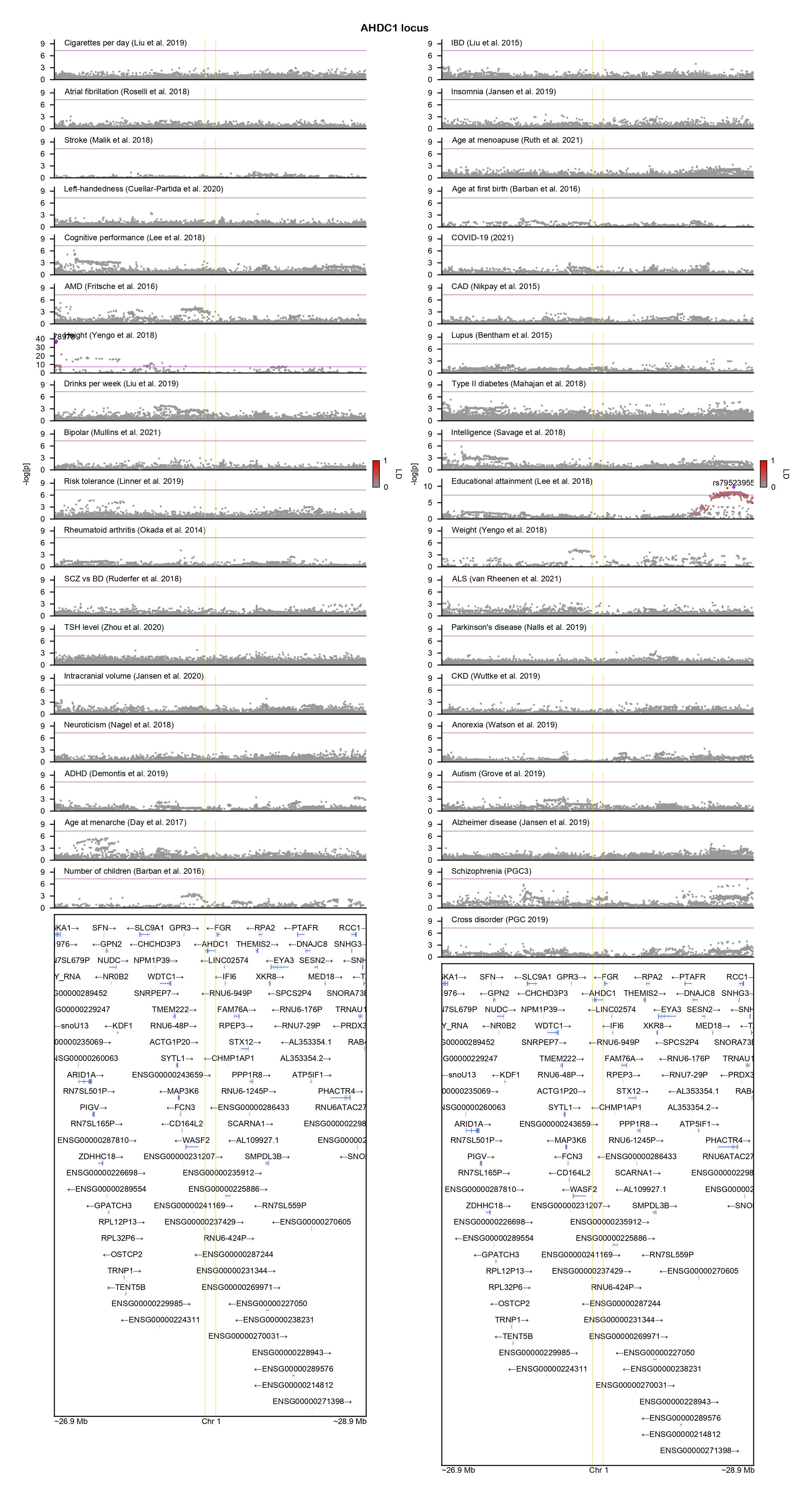

### ARHGAP35-locuszoom.png

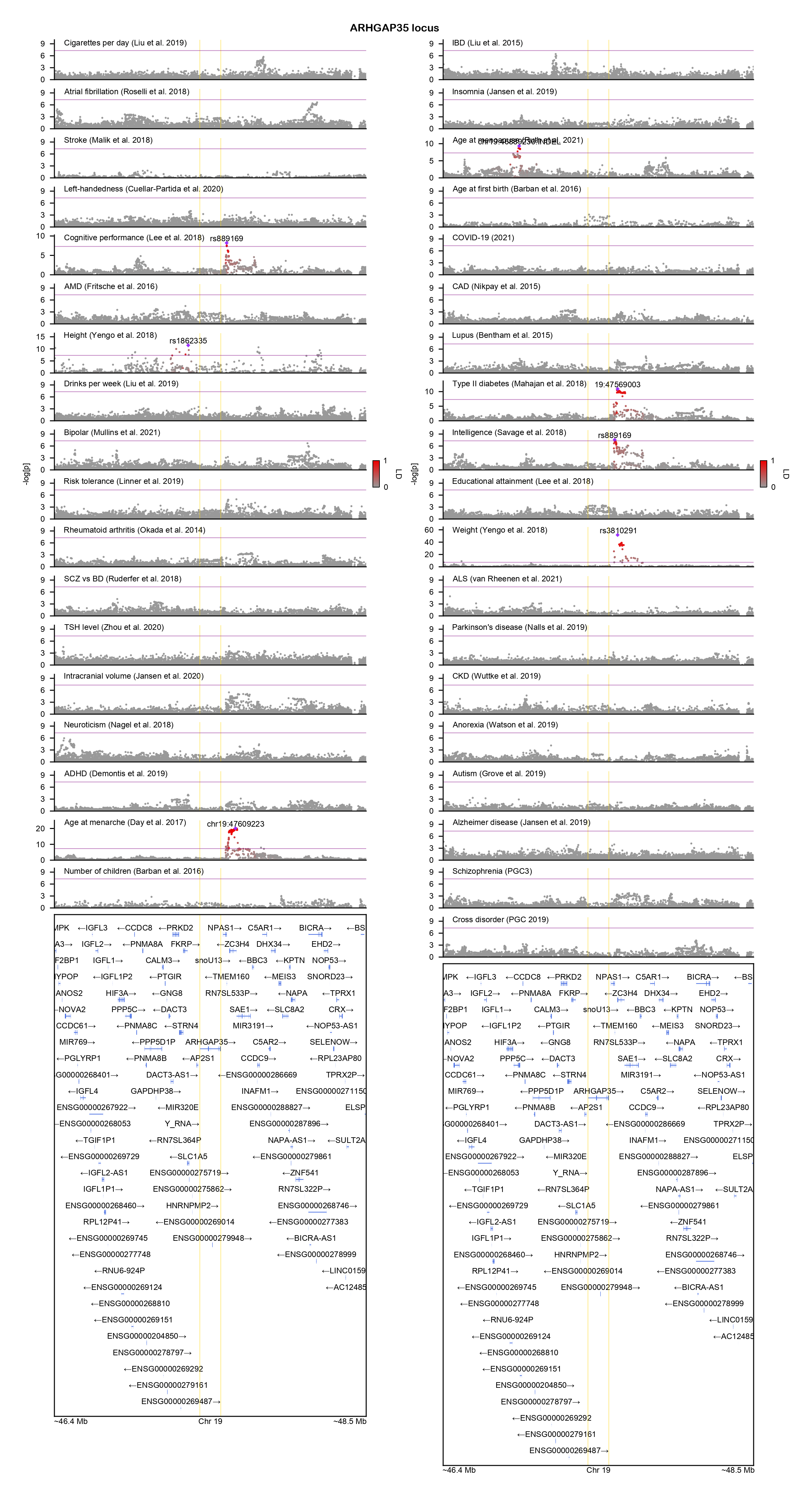

### ATP6V0A1-locuszoom.png

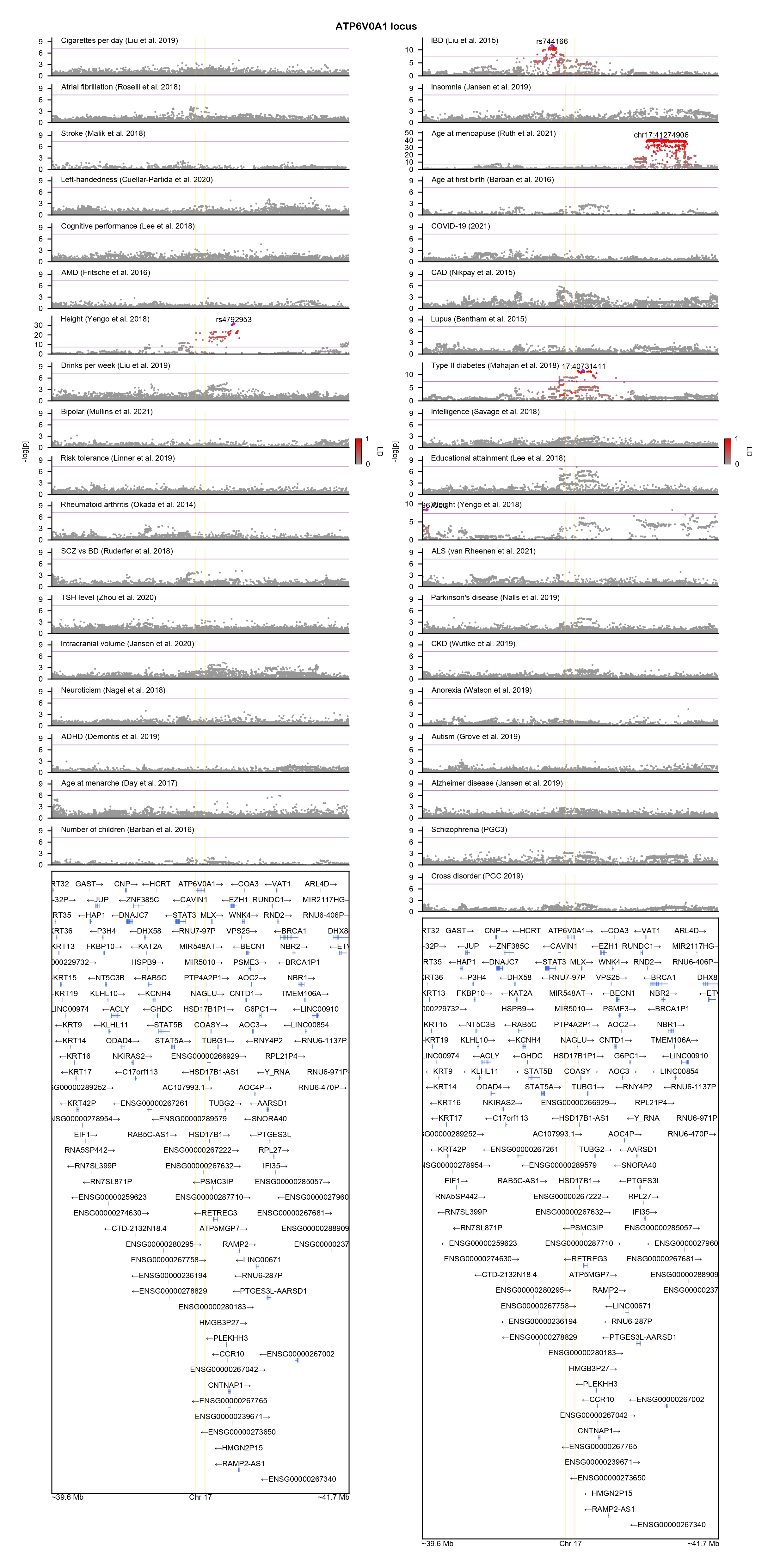

### ATRX-locuszoom.png

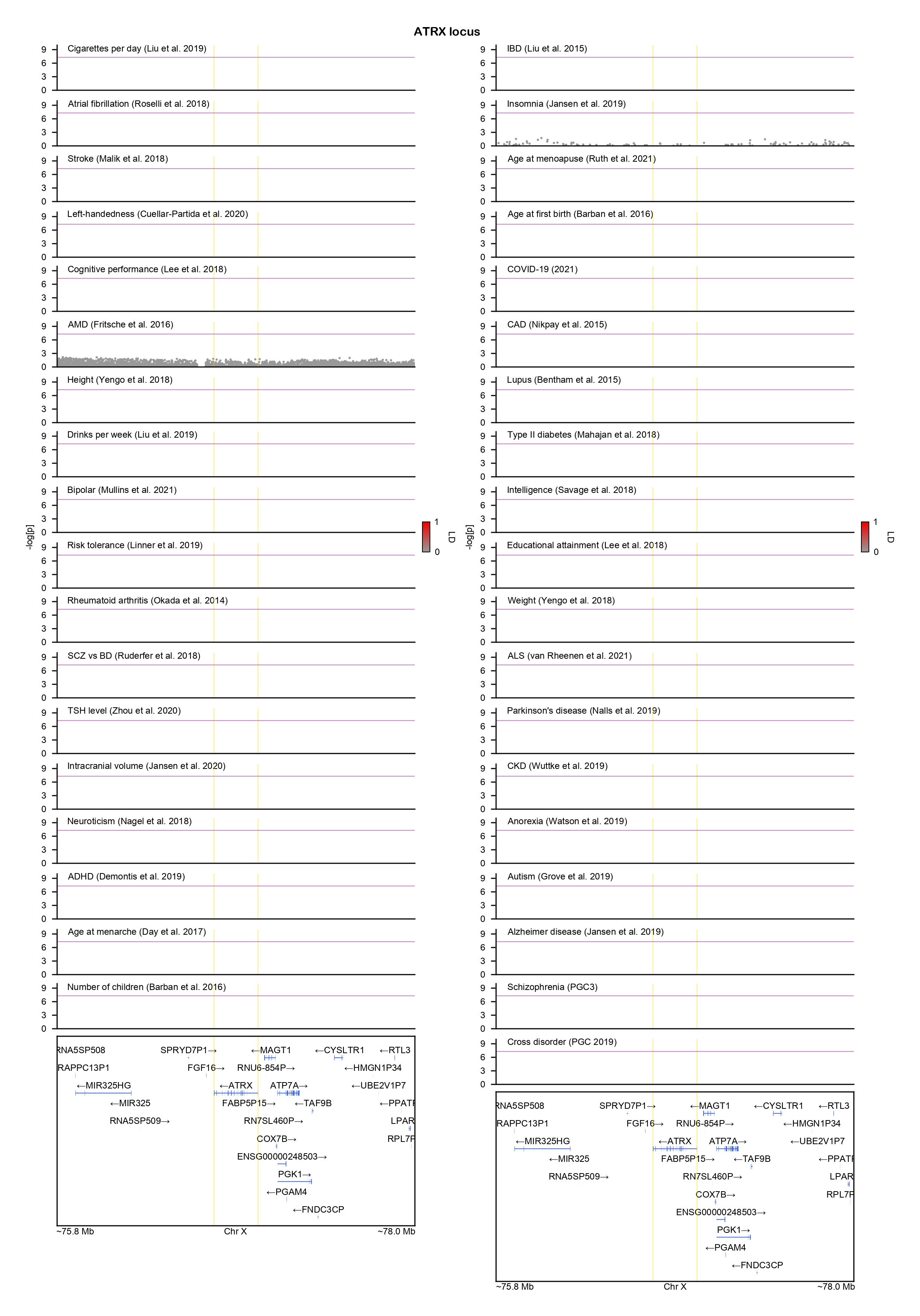

### BRAF-locuszoom.png

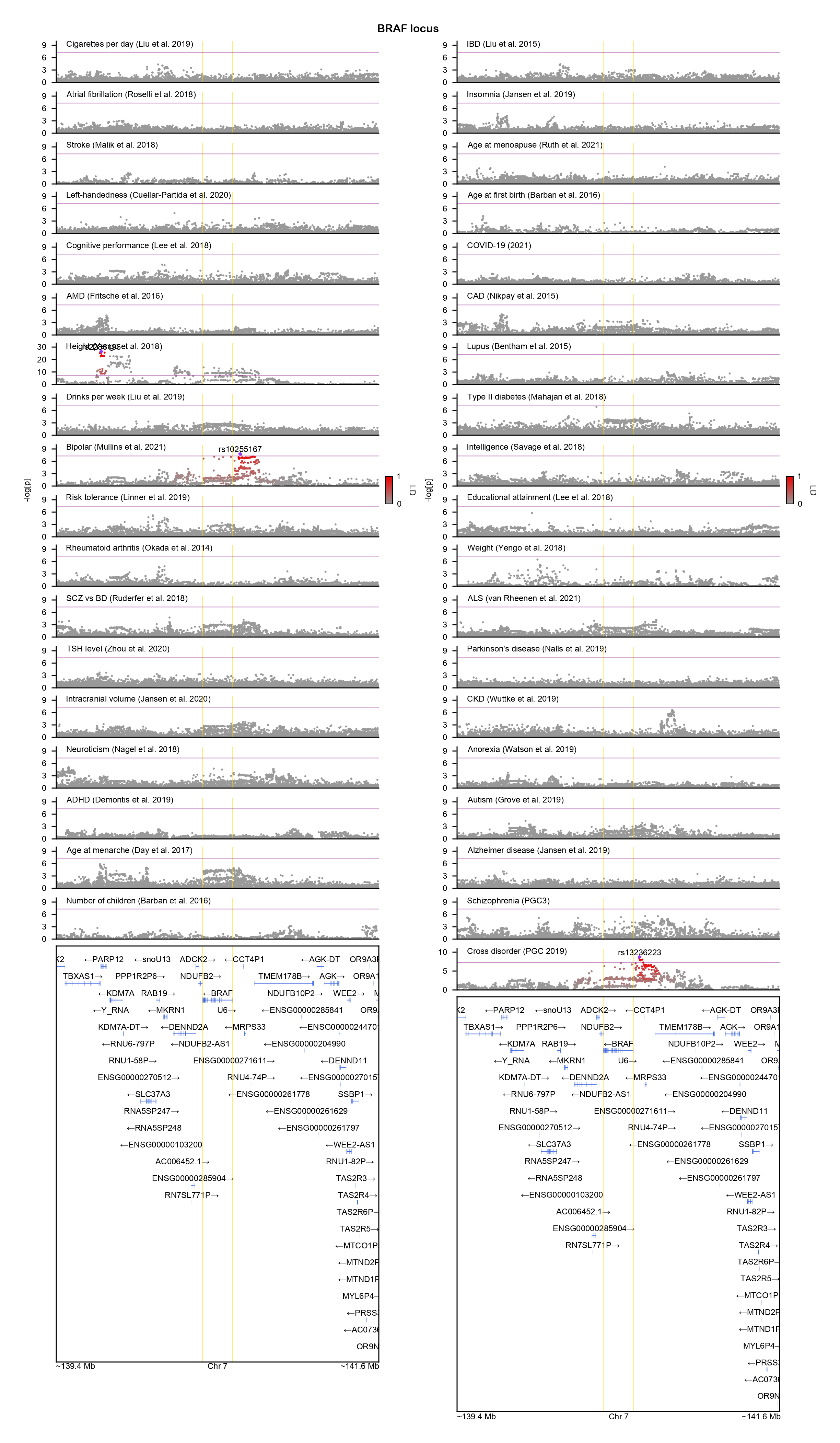

### BRPF1-locuszoom.png

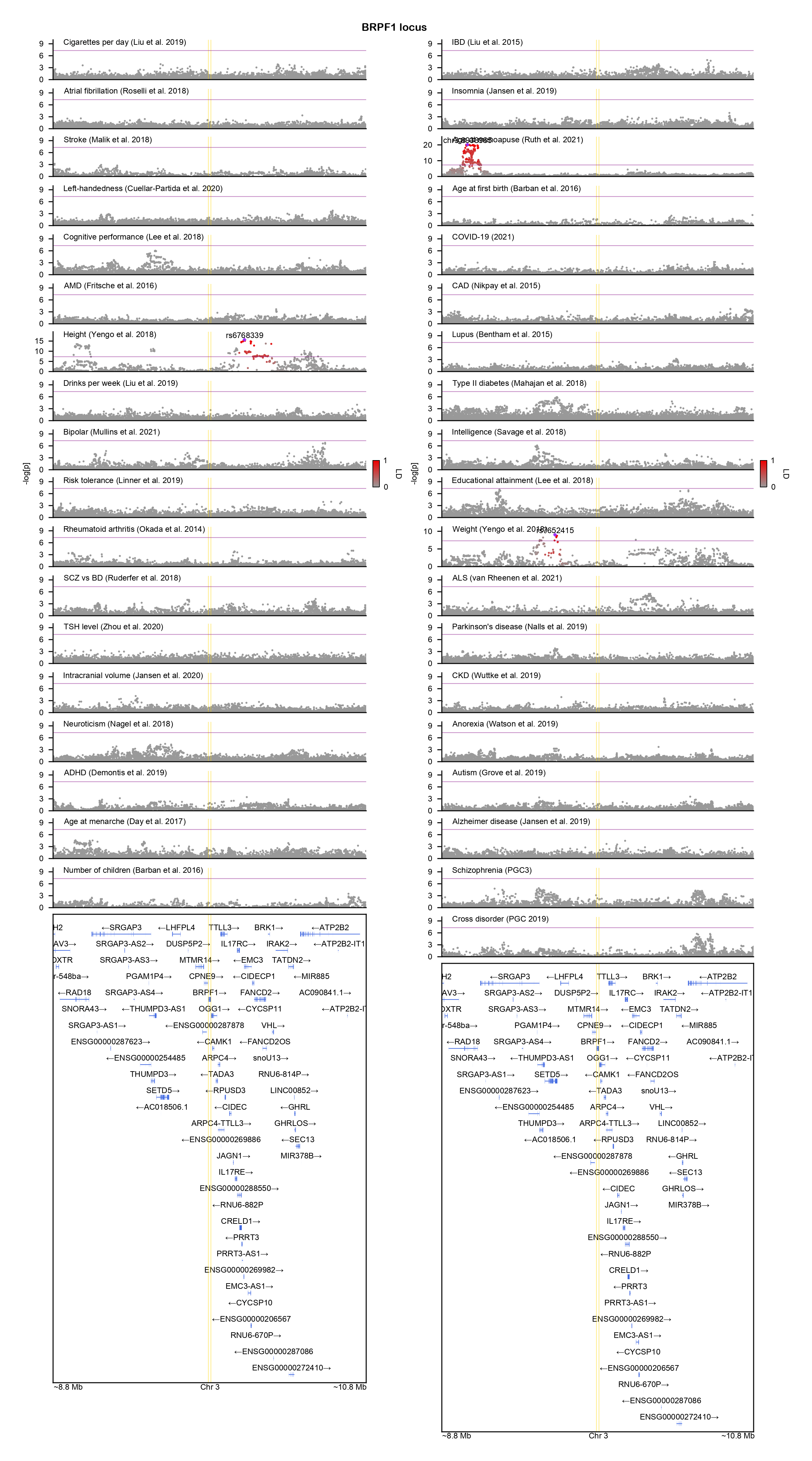

### CACNA1A-locuszoom.png

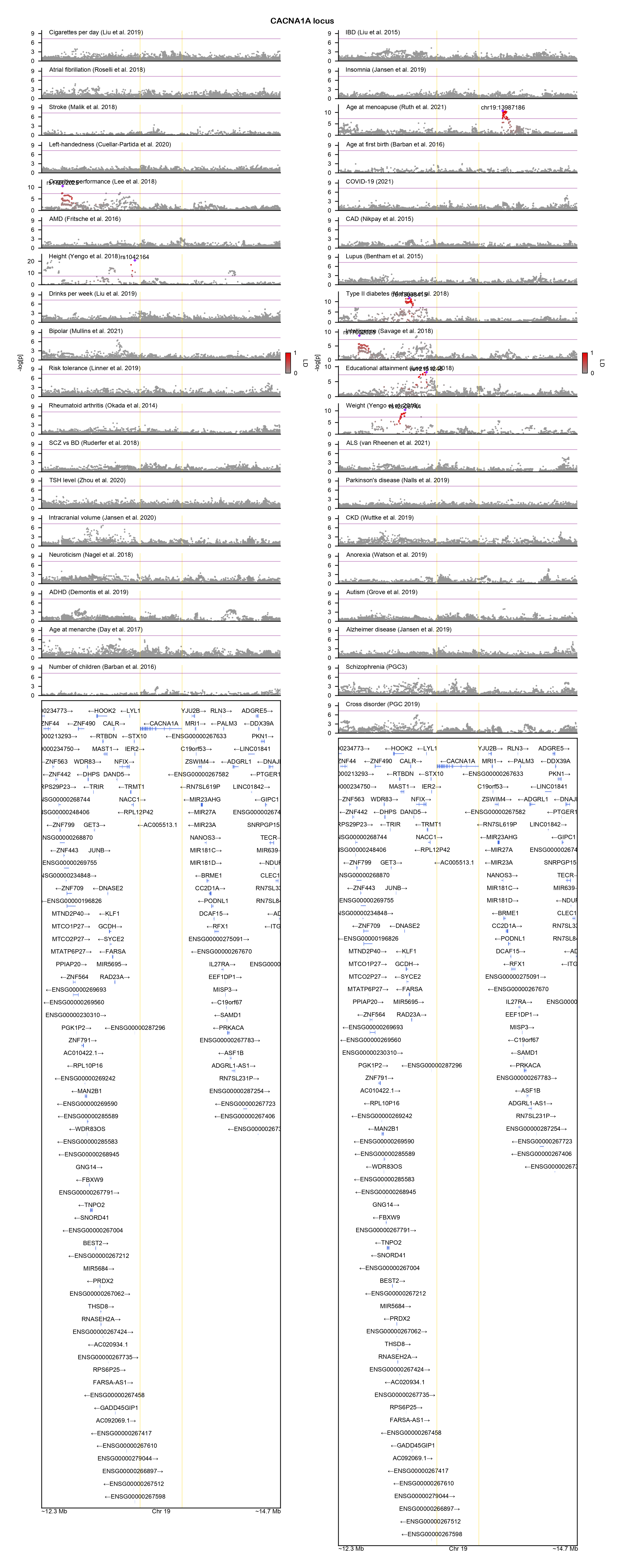

### CACNA1C-locuszoom.png

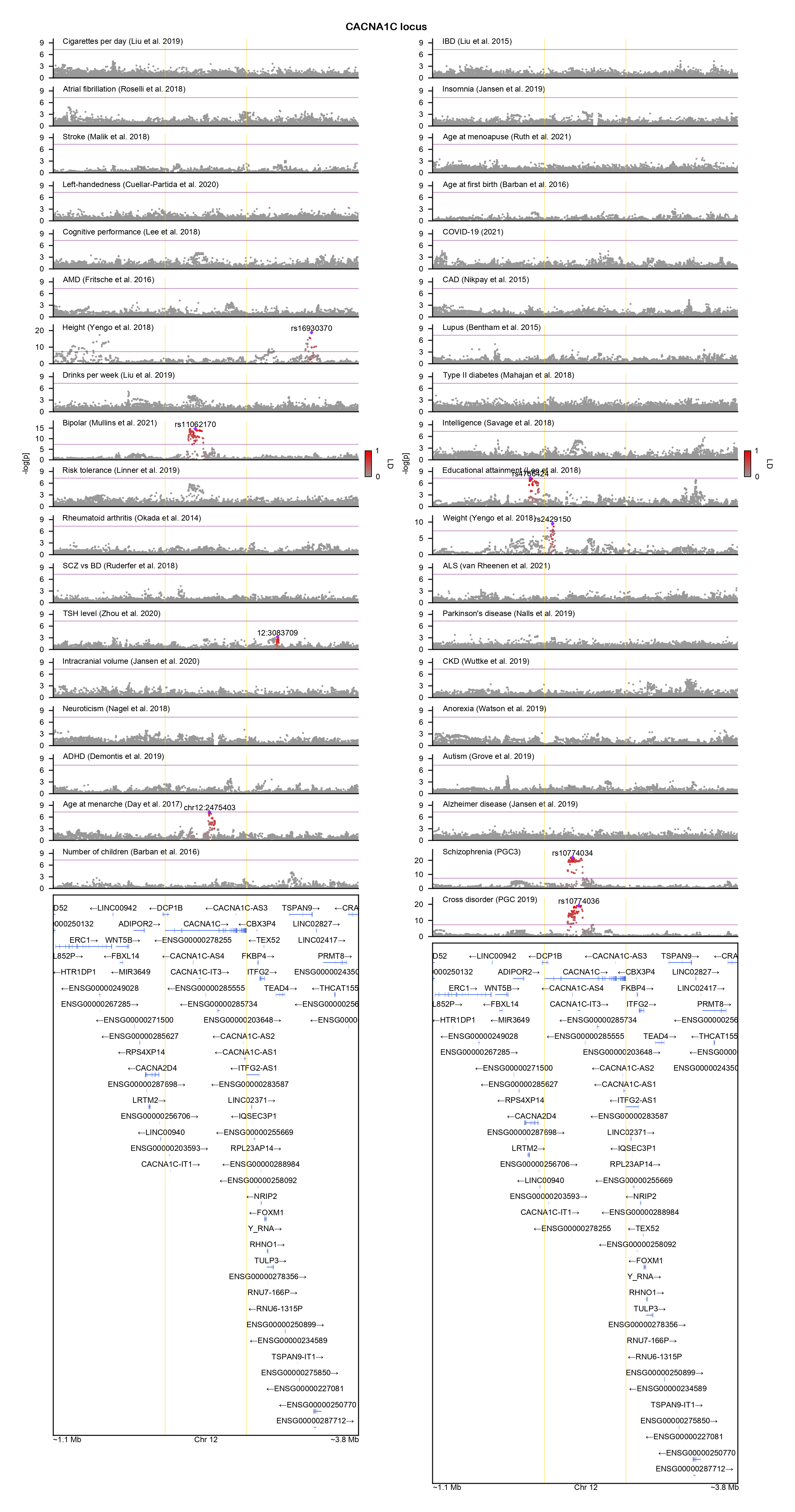

### CACNA1E-locuszoom.png

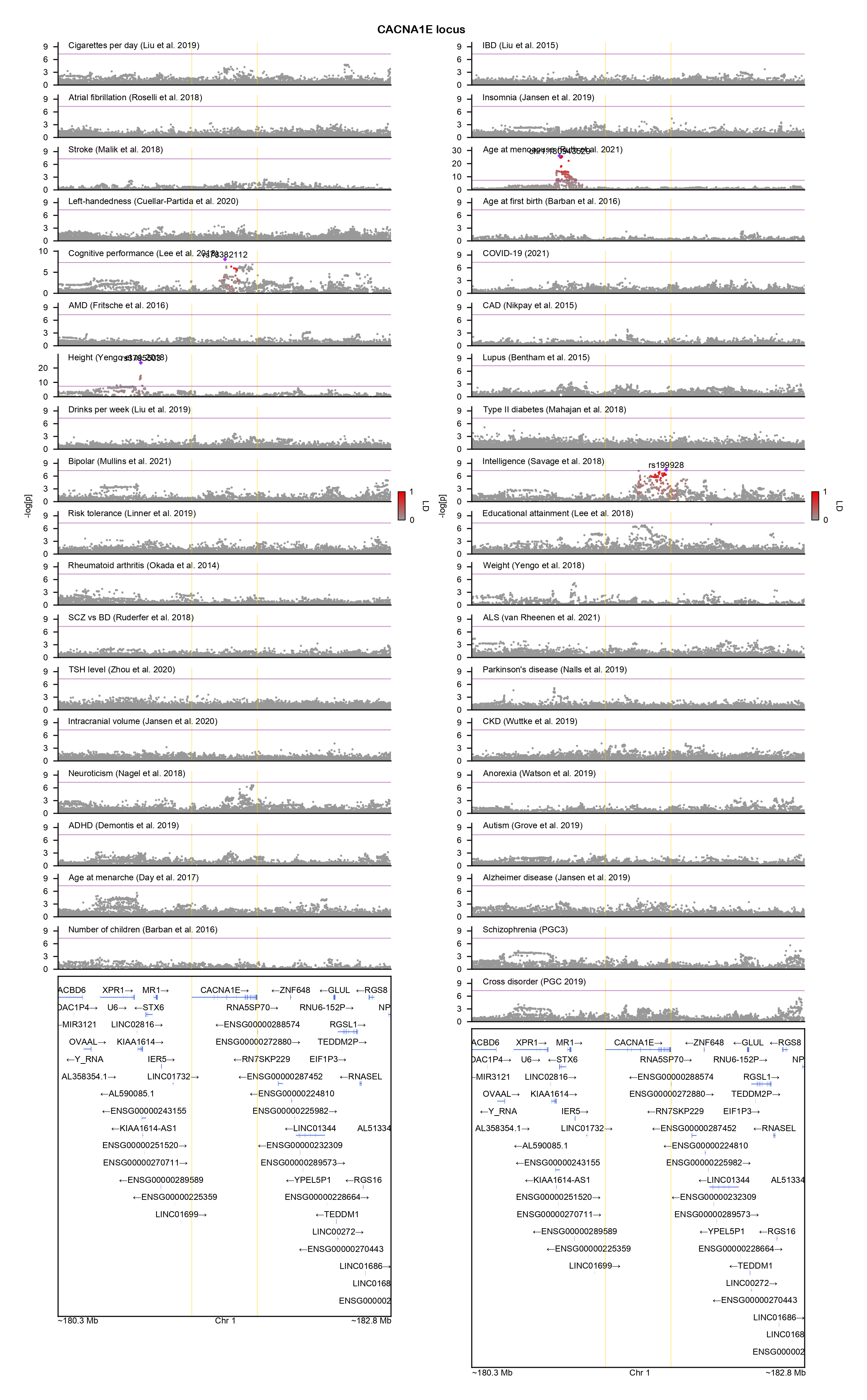

### CACNA1G-locuszoom.png

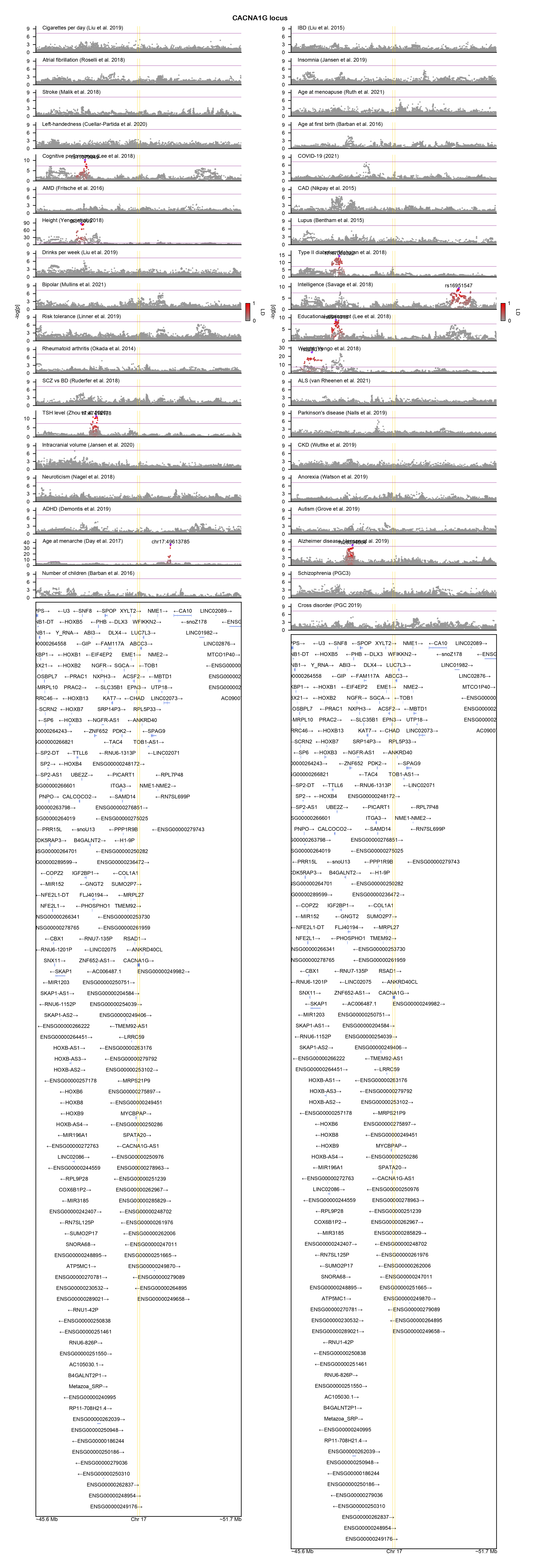

### CAMK2G-locuszoom.png

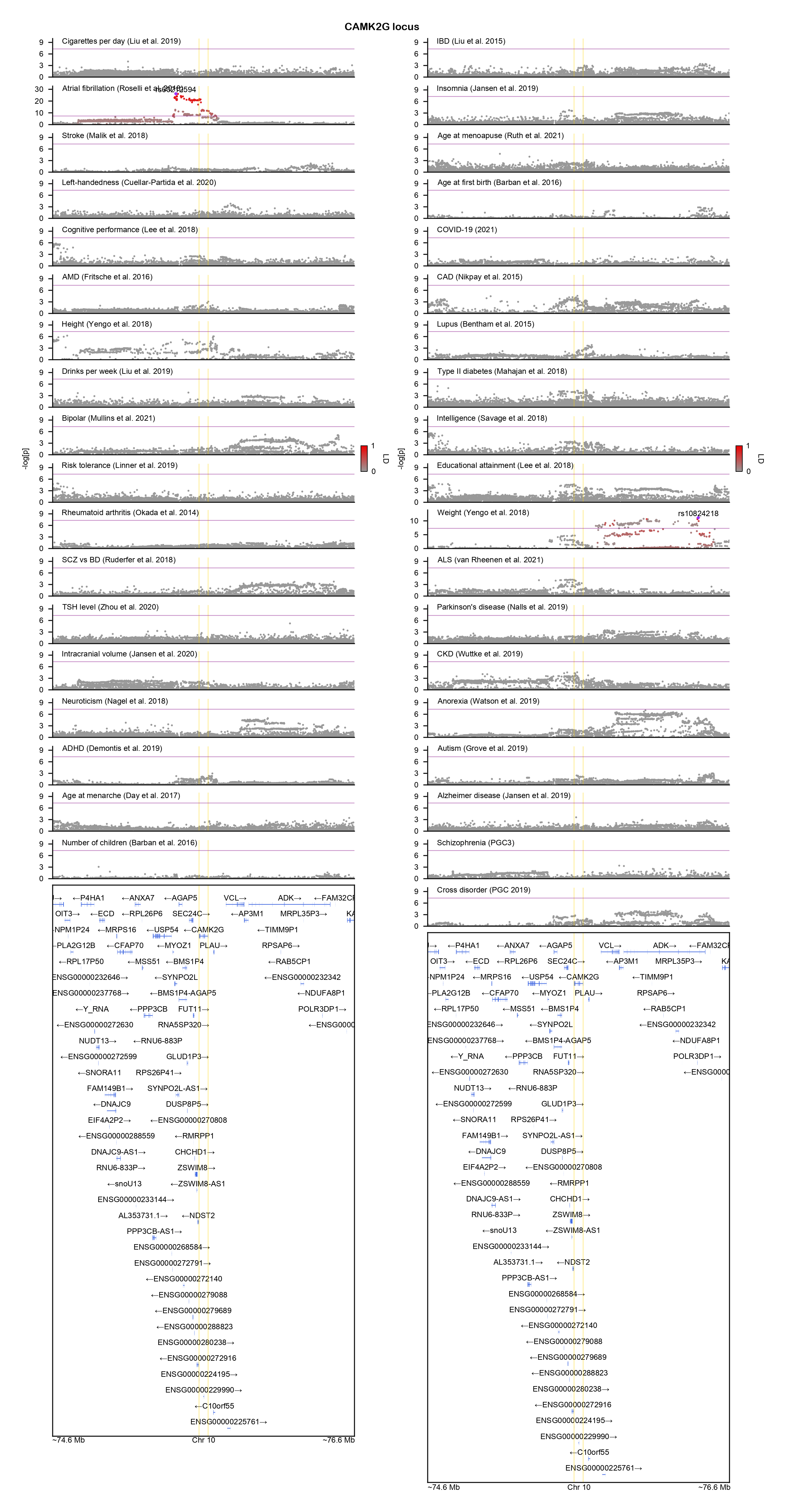

### CBL-locuszoom.png

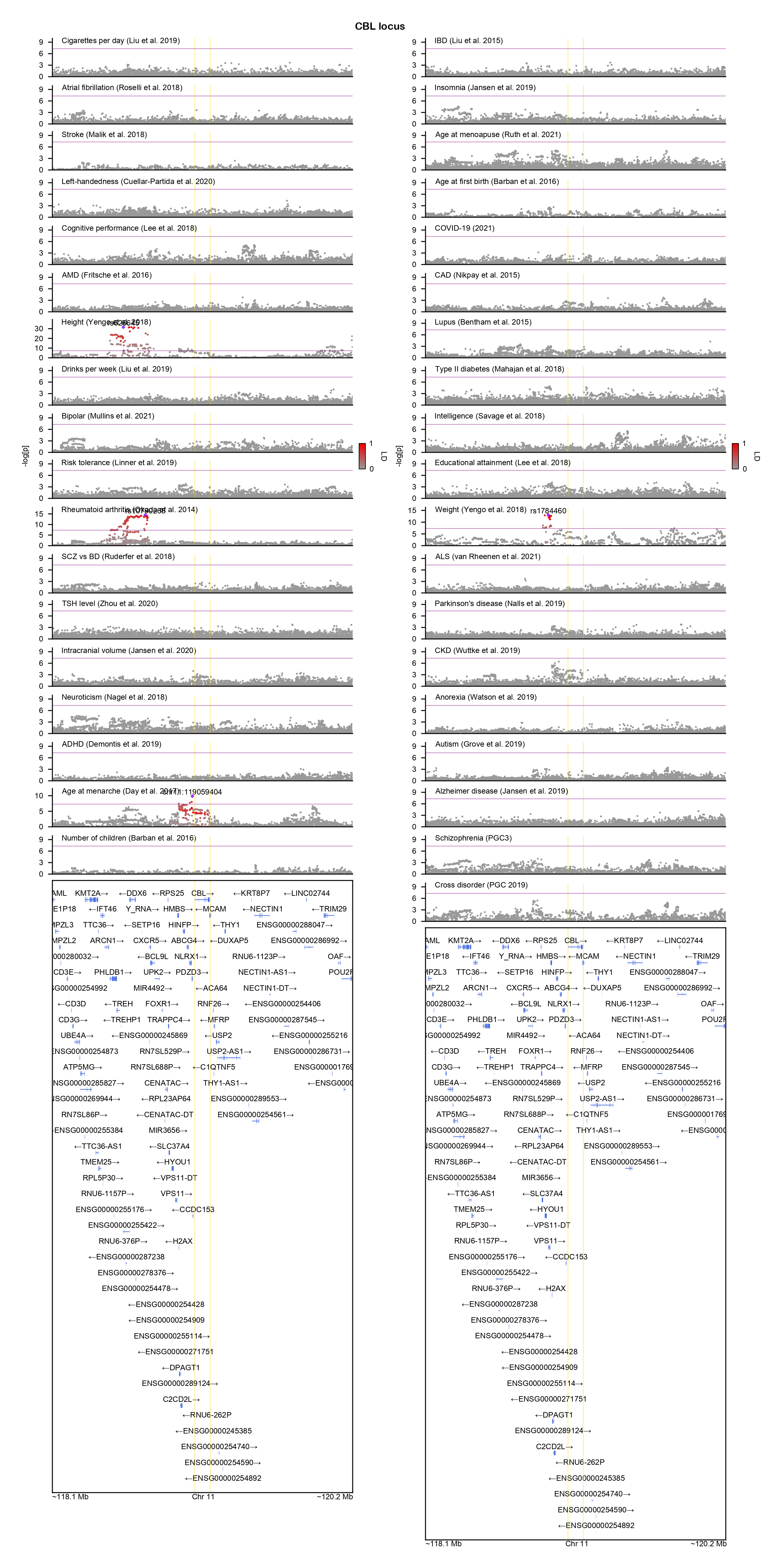

### CHD8-locuszoom.png

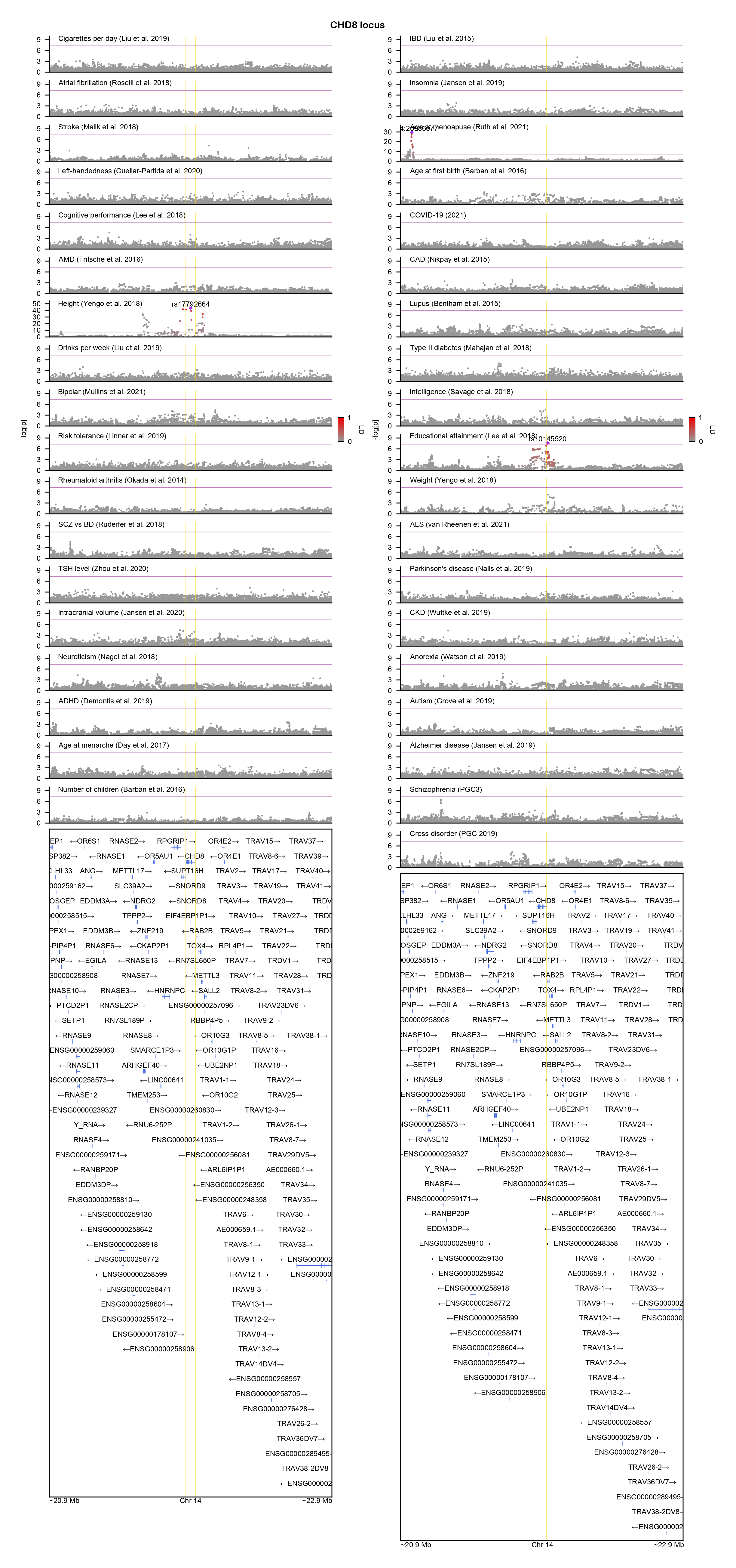

### CLTC-locuszoom.png

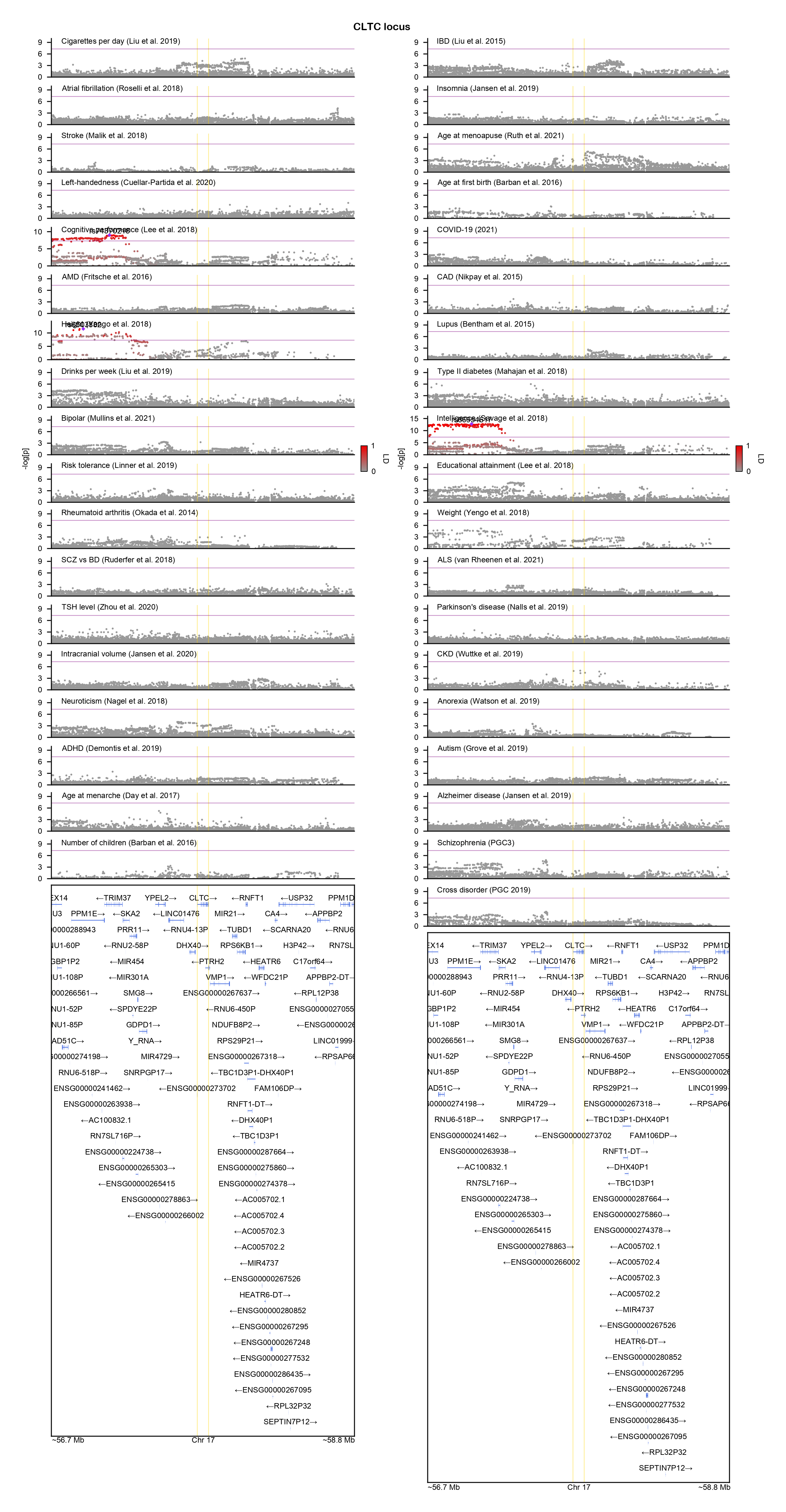

### CNOT3-locuszoom.png

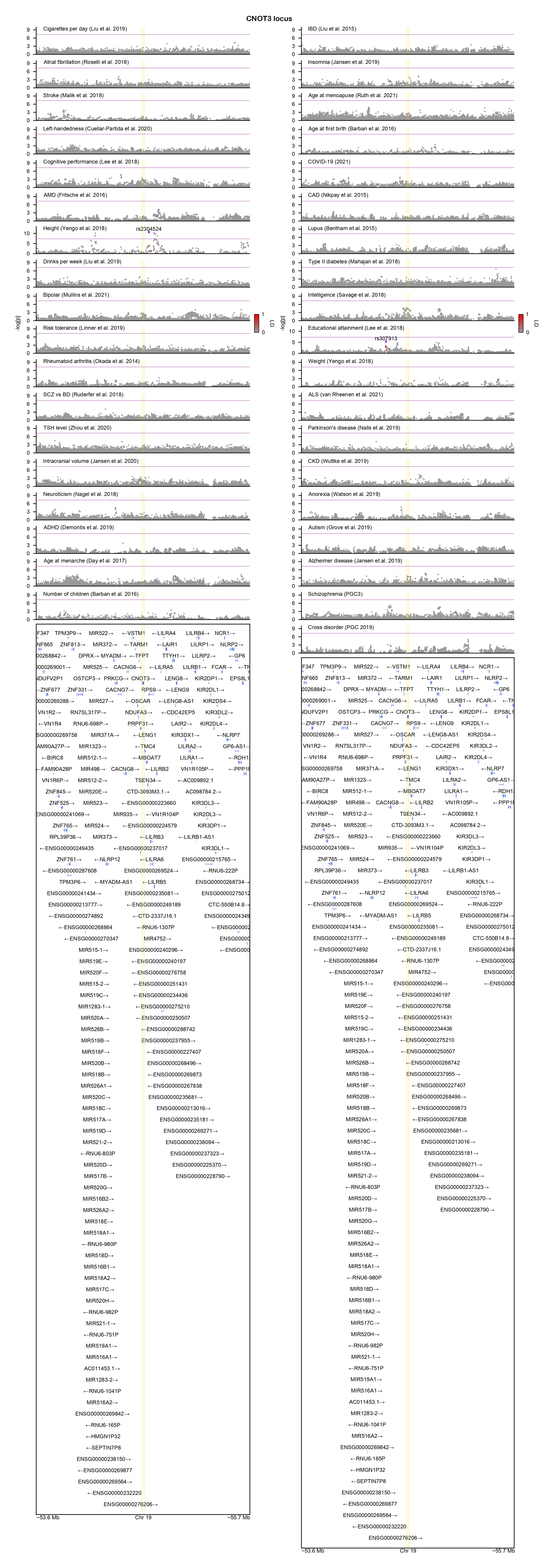

### CREBBP-locuszoom.png

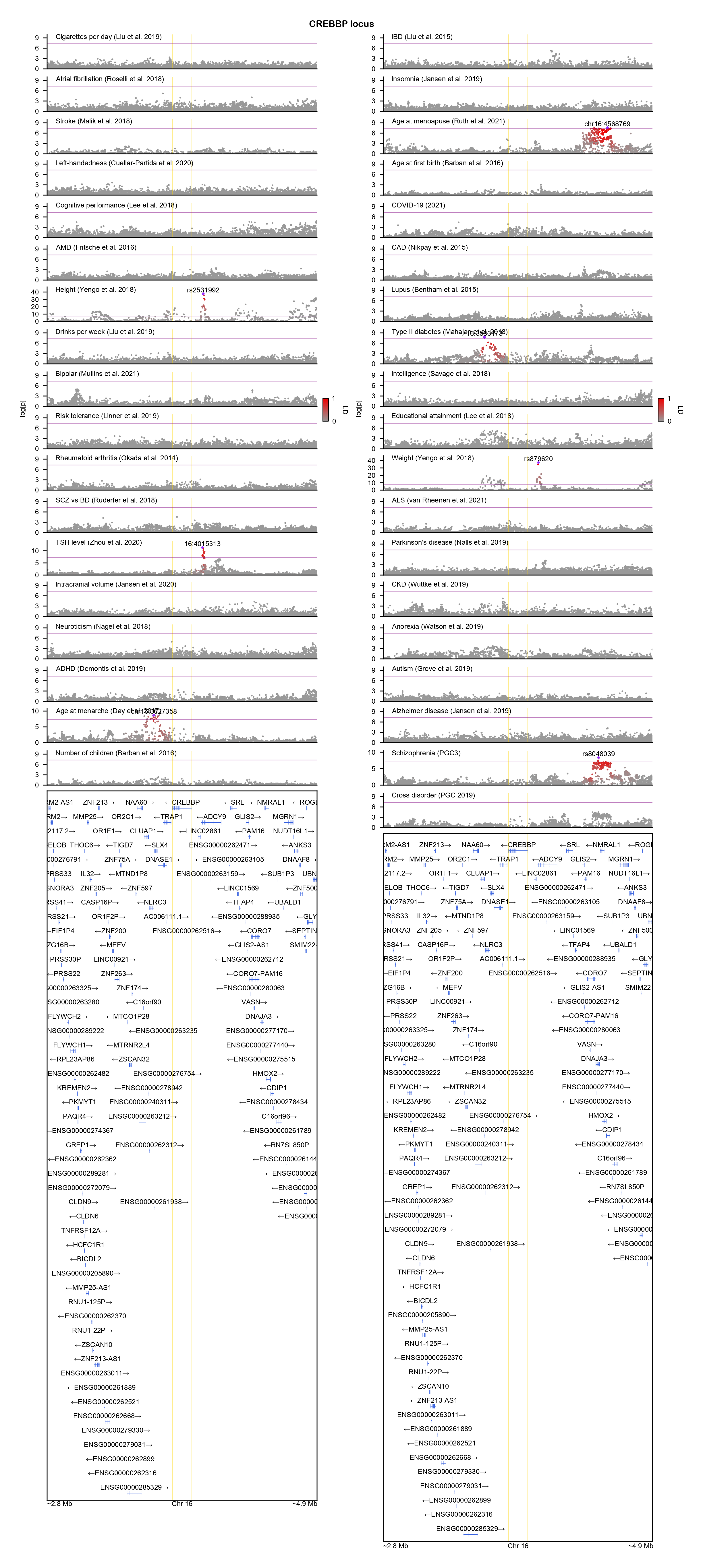

### CSNK2B-locuszoom.png

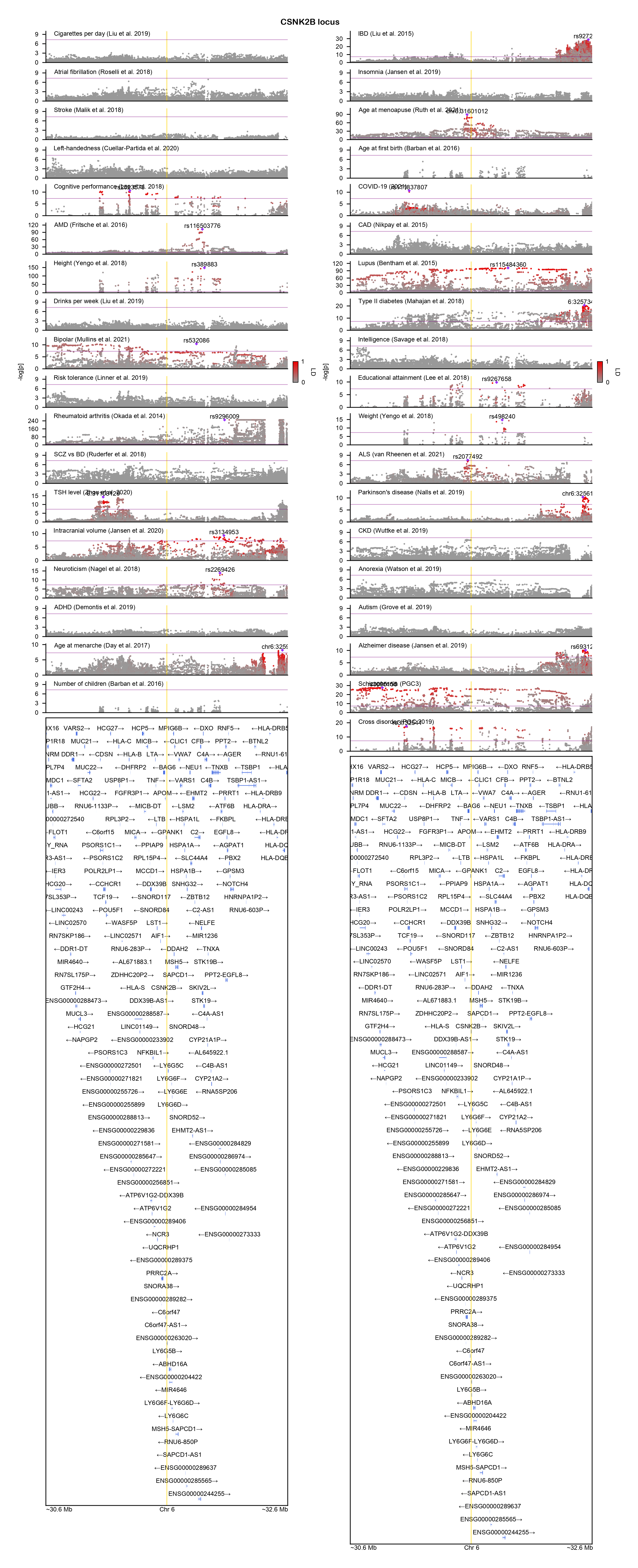

### CTBP1-locuszoom.png

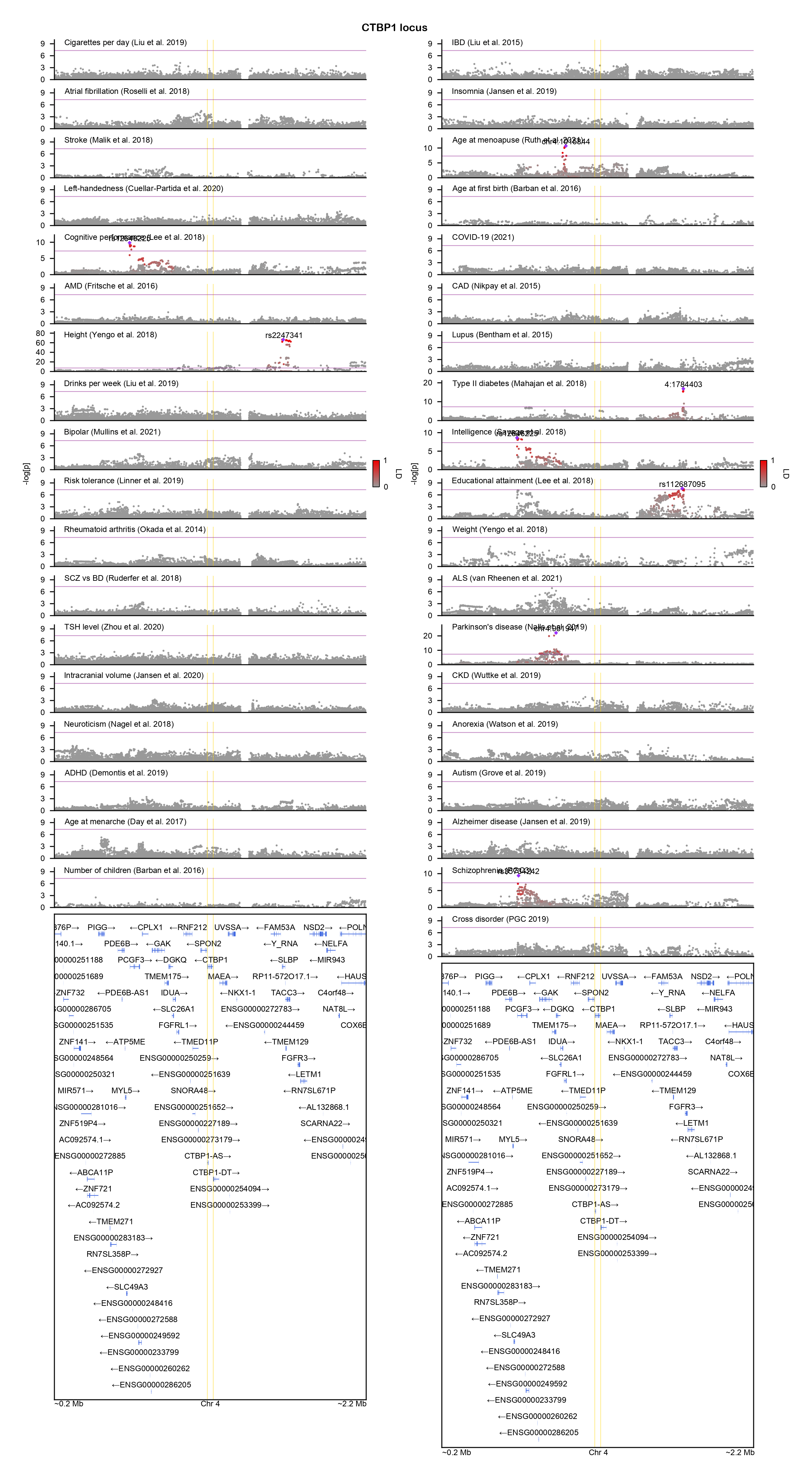

### DDX3X-locuszoom.png

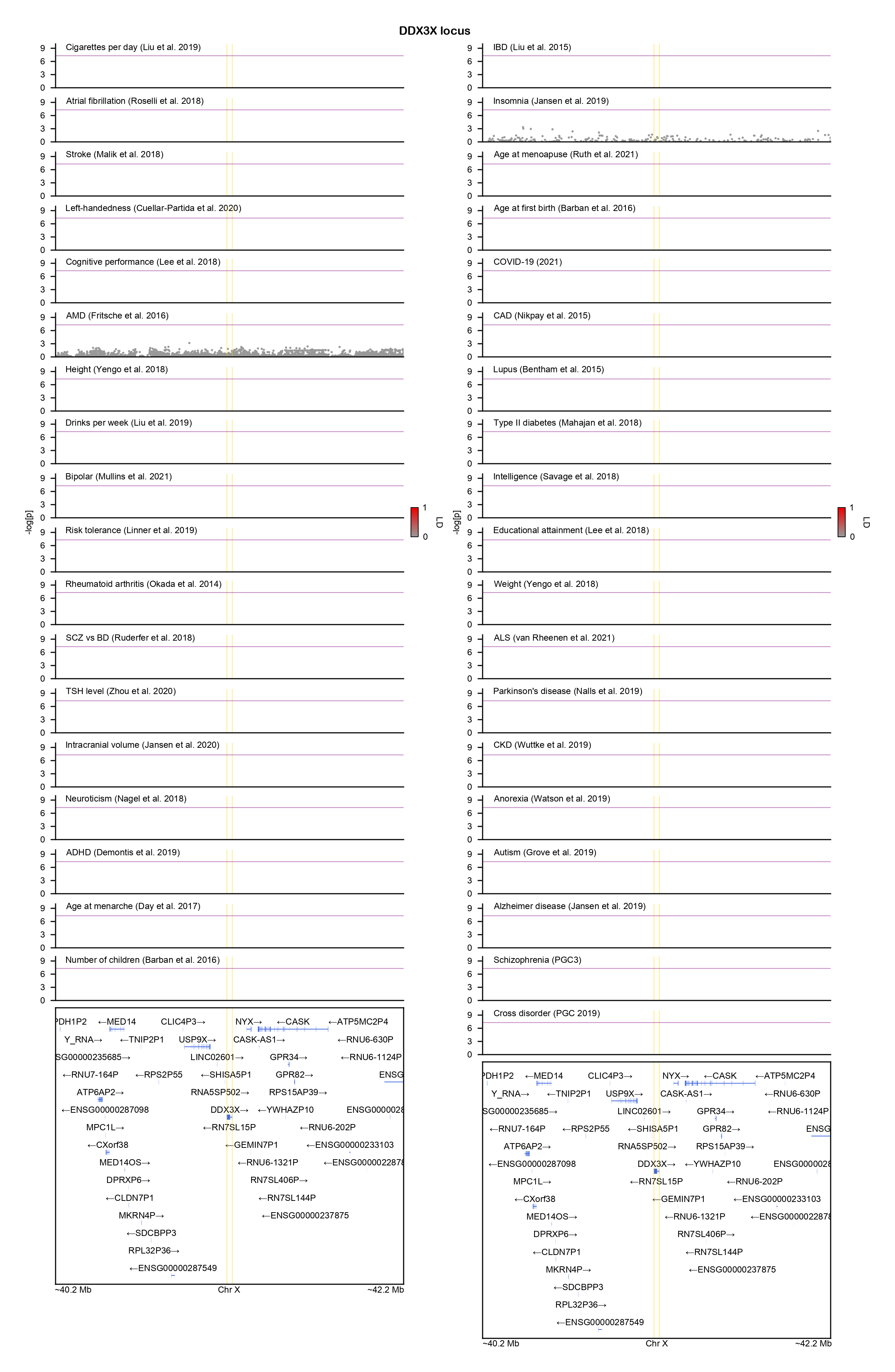

### DDX23-locuszoom.png

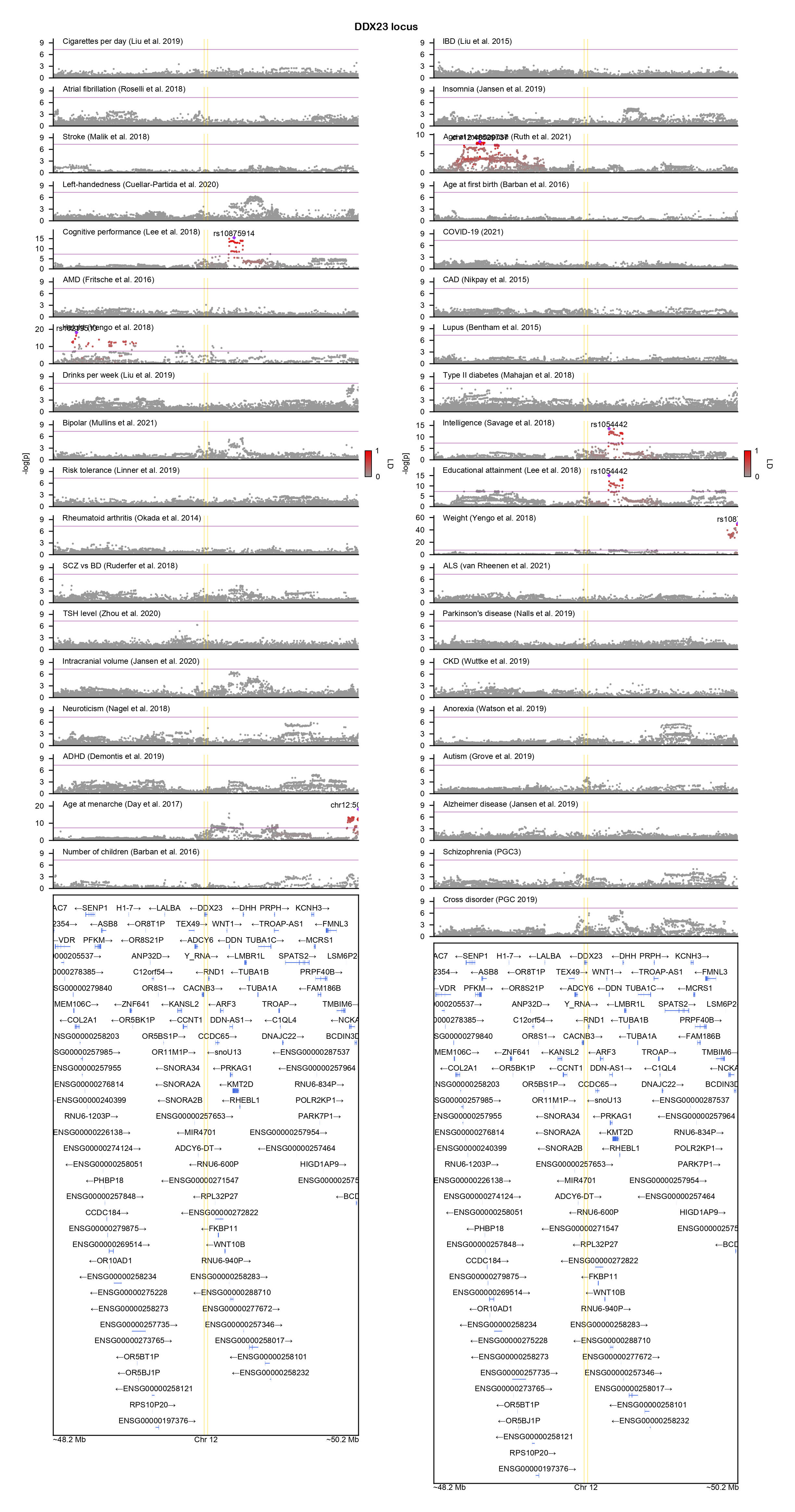

### DIP2A-locuszoom.png

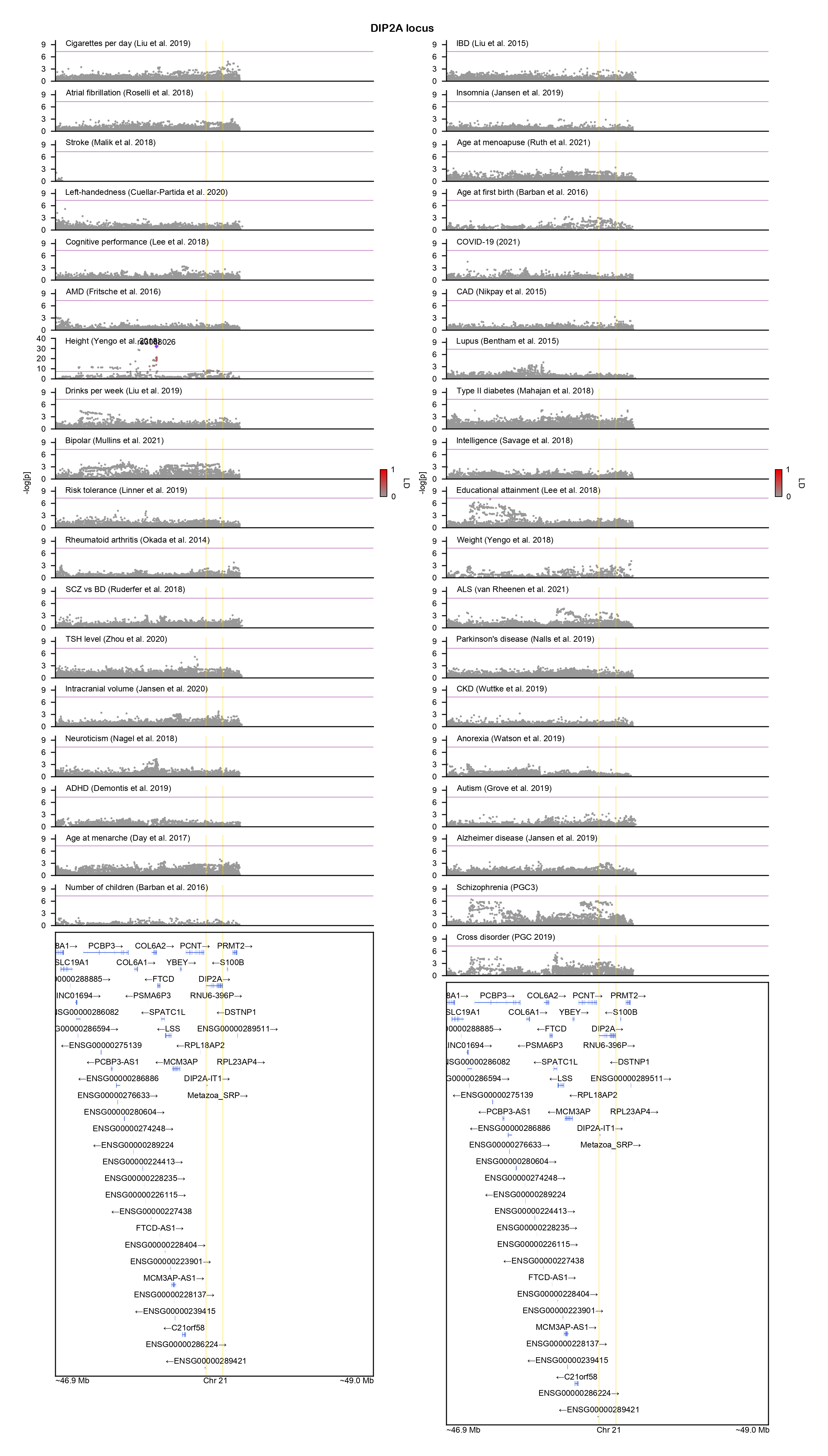

### DPF2-locuszoom.png

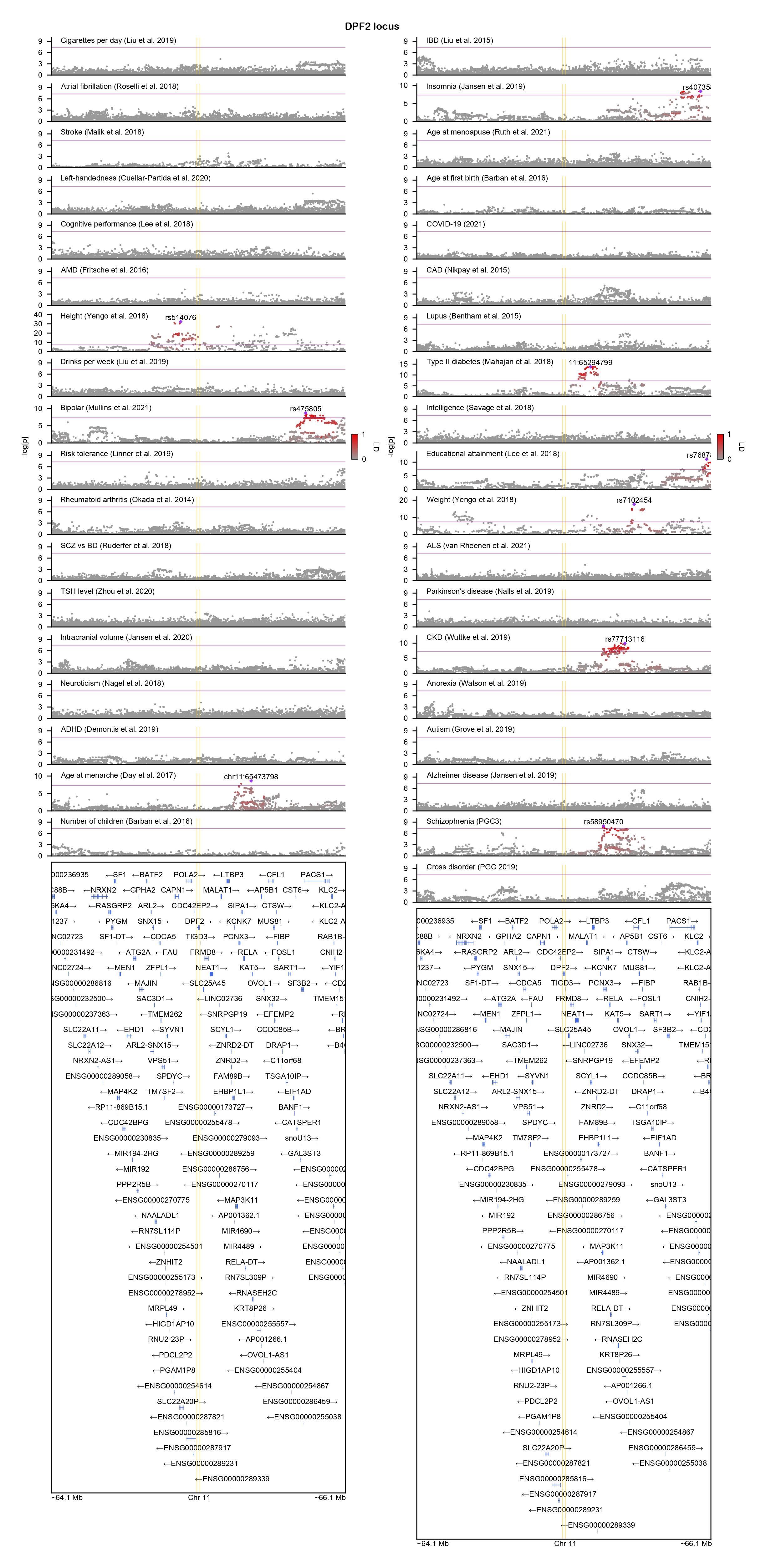

### EEF1A2-locuszoom.png

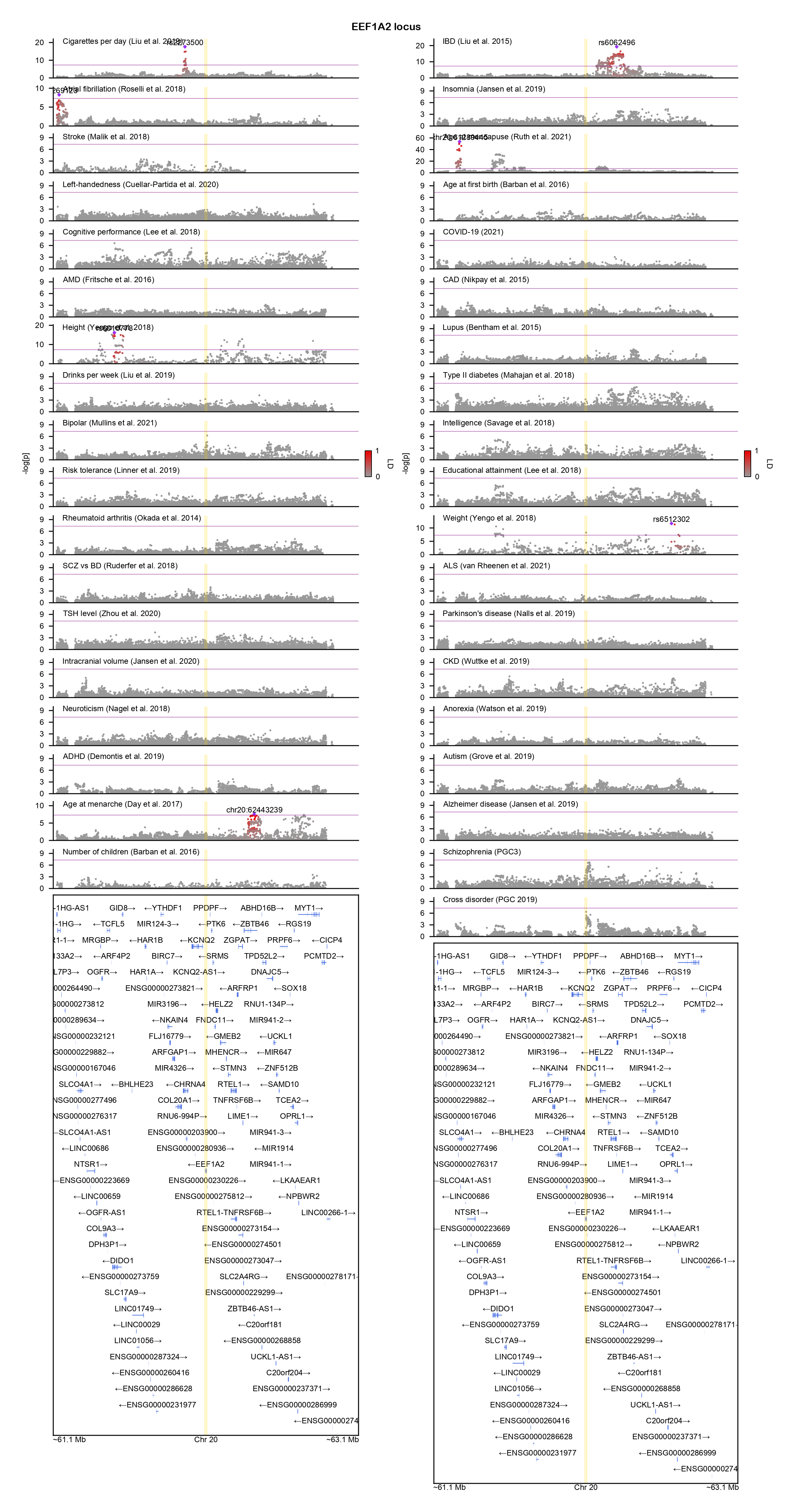

### EP300-locuszoom.png

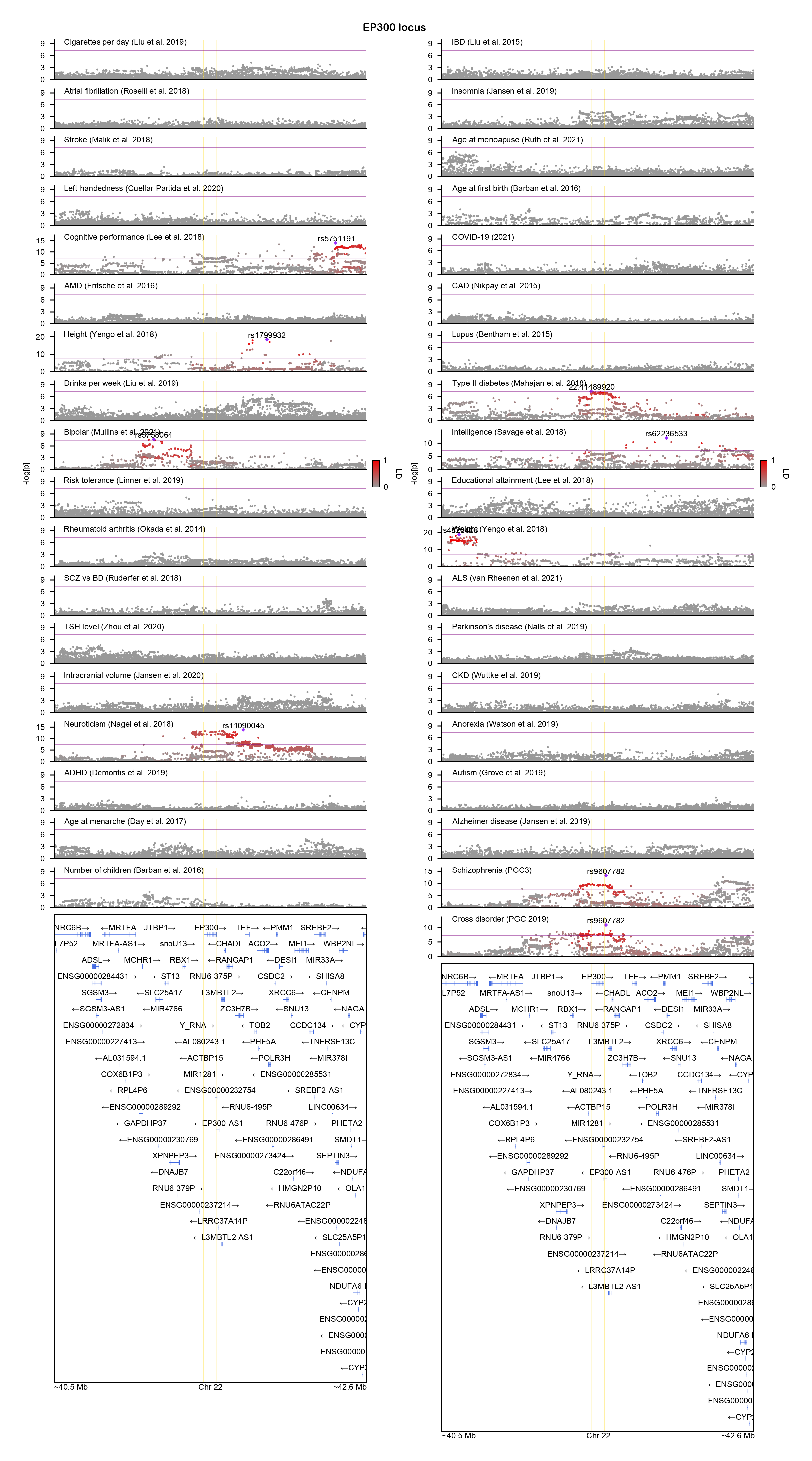

### FBXO11-locuszoom.png

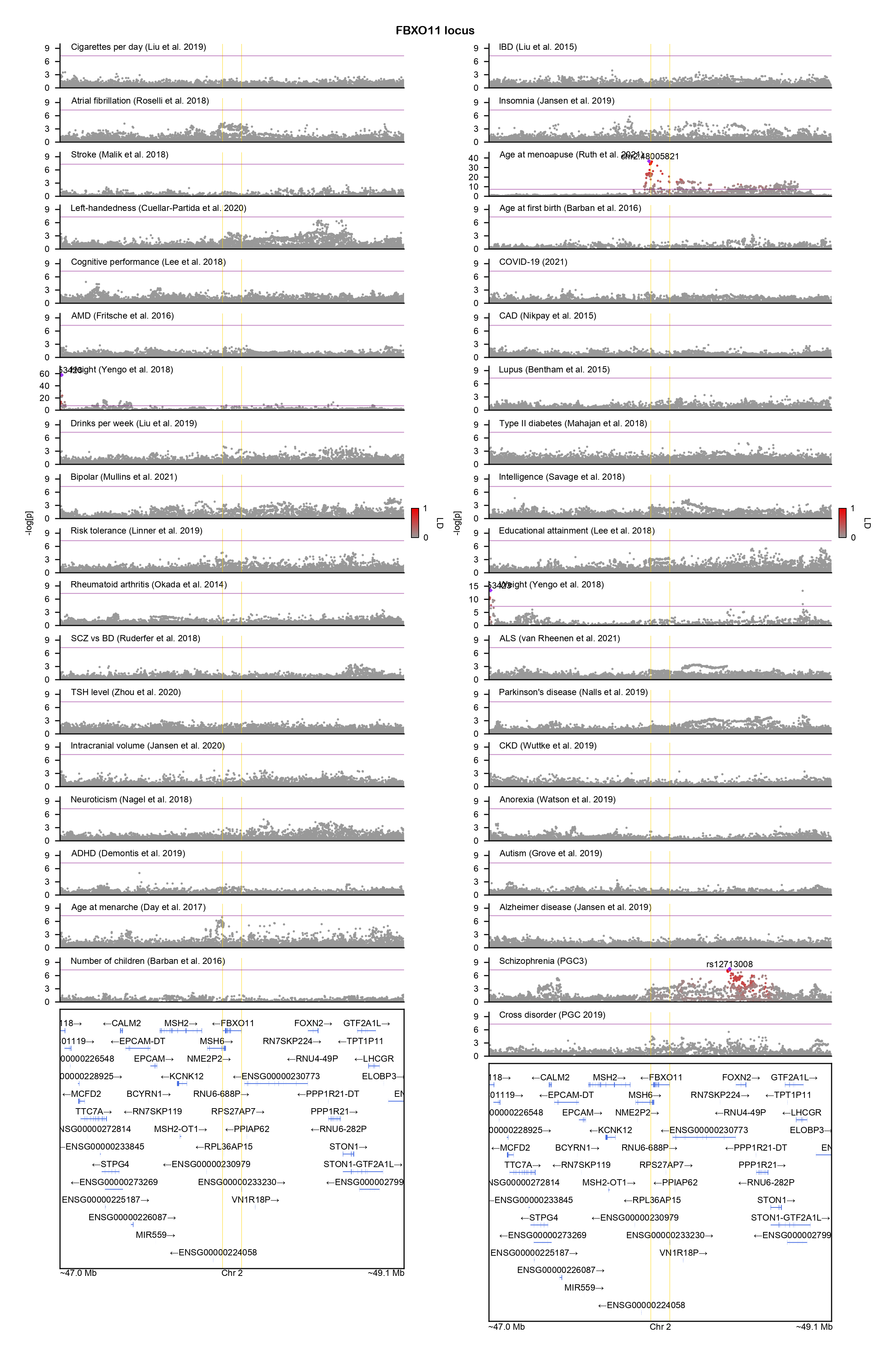

### FGFR2-locuszoom.png

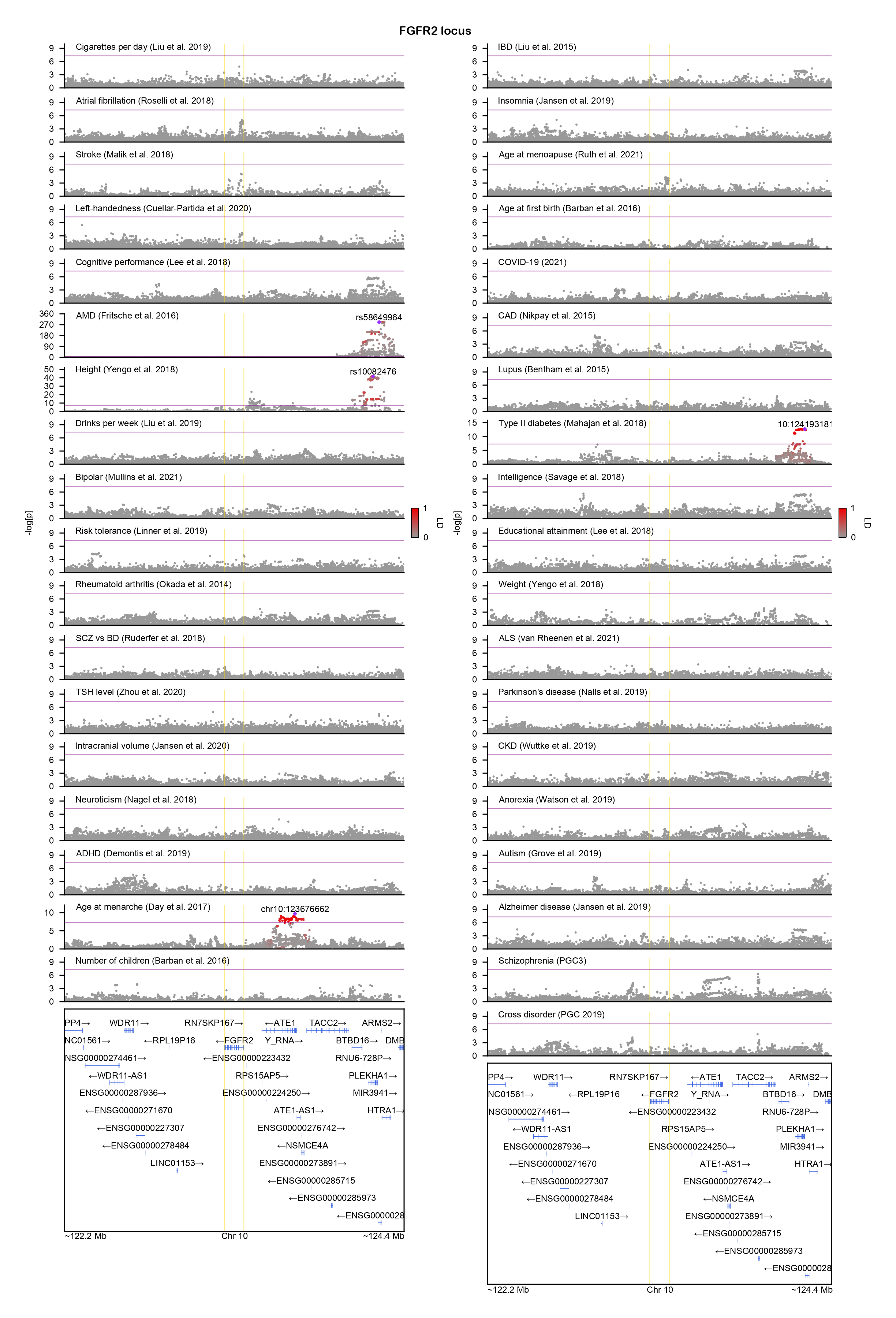

### FGFR3-locuszoom.png

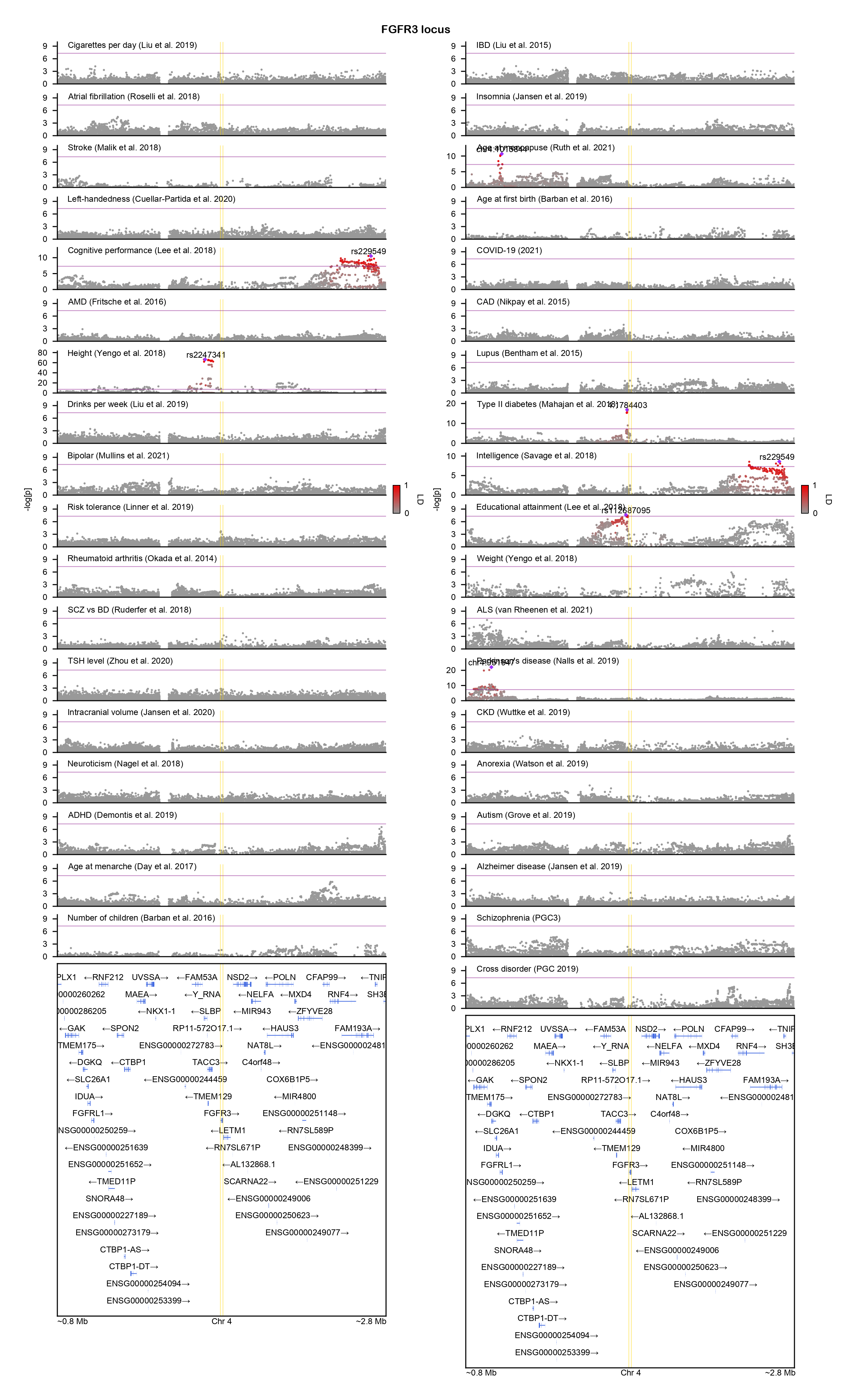

### FOXP2-locuszoom.png

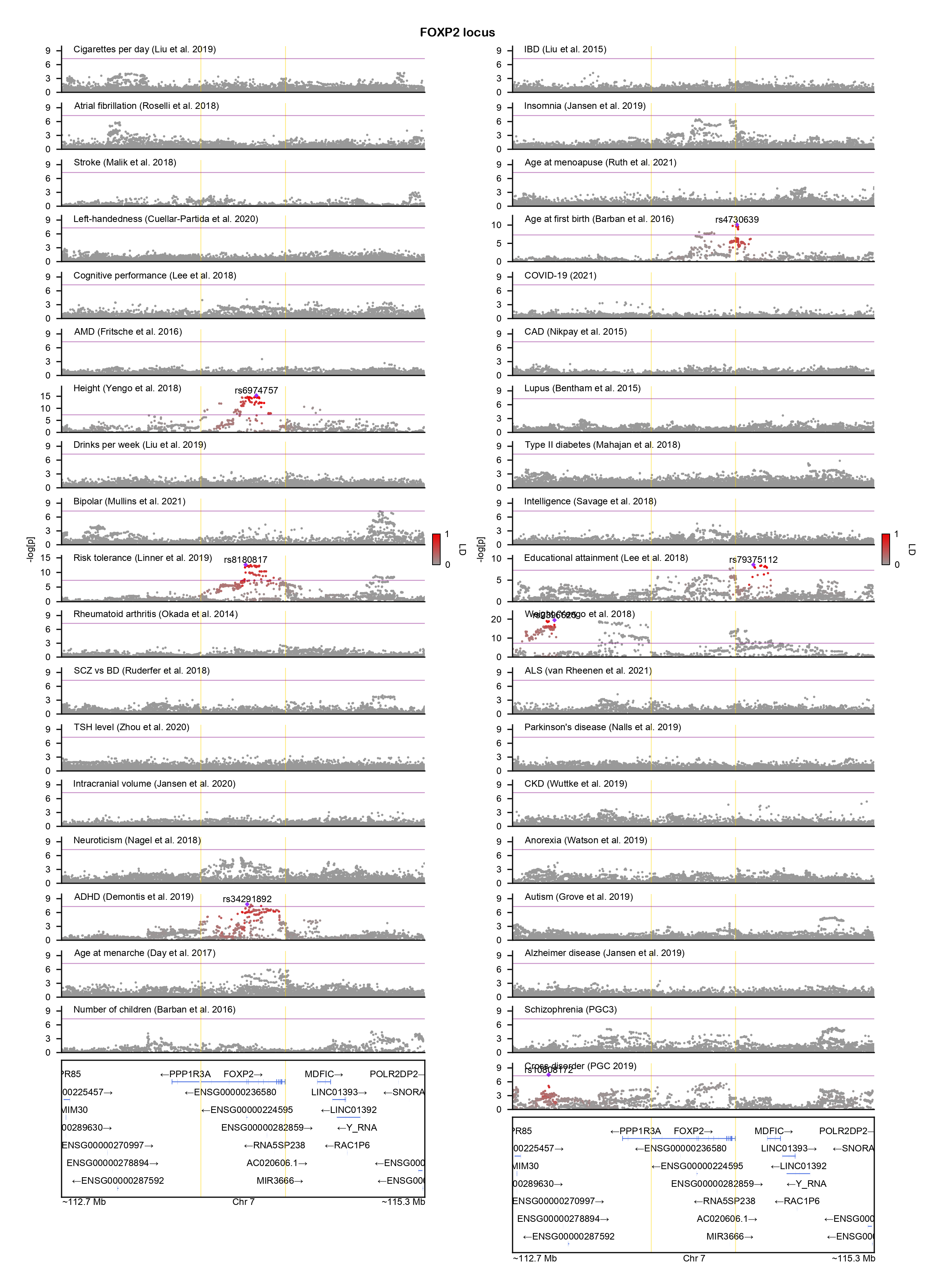

### GABRB3-locuszoom.png

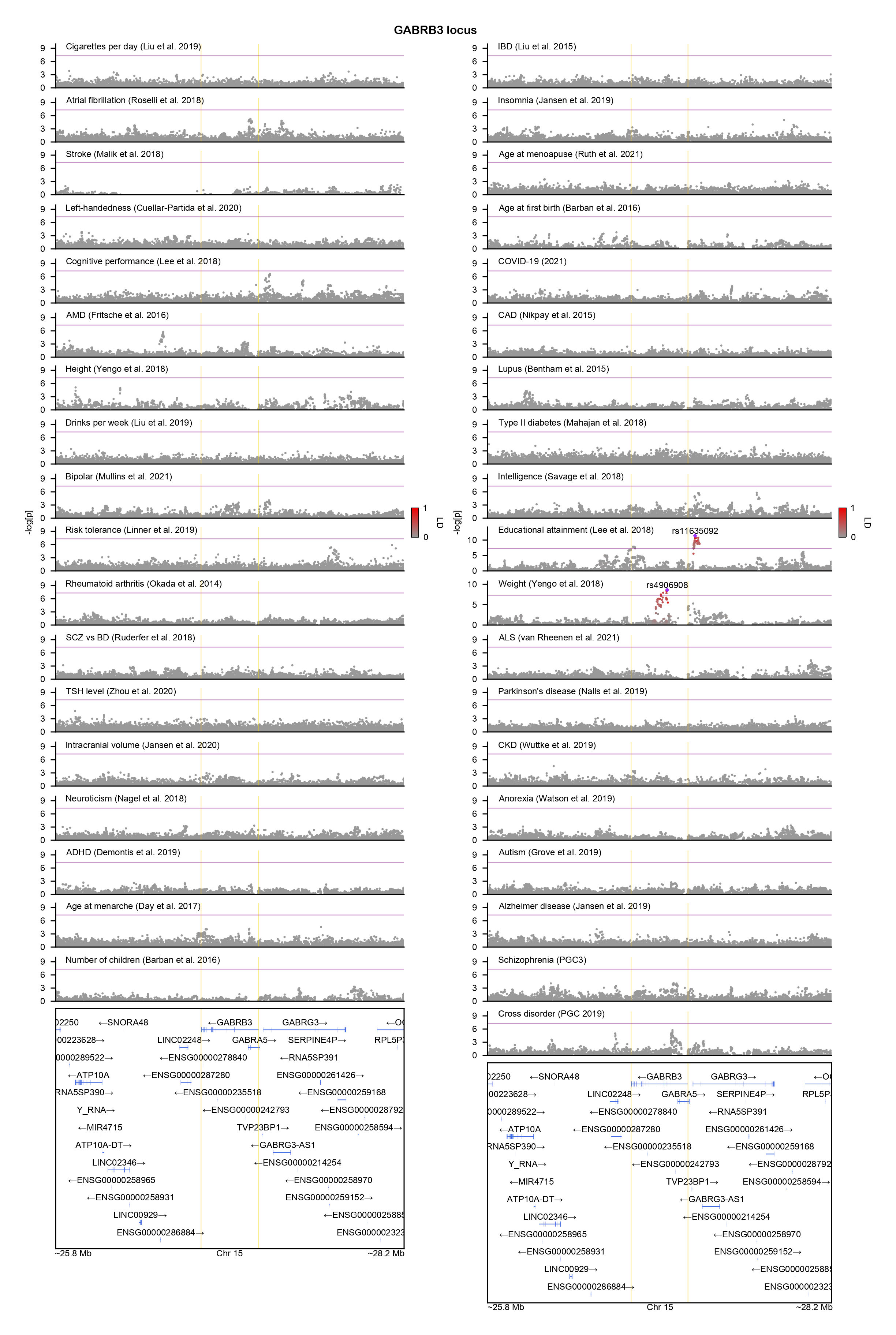

### GNB1-locuszoom.png

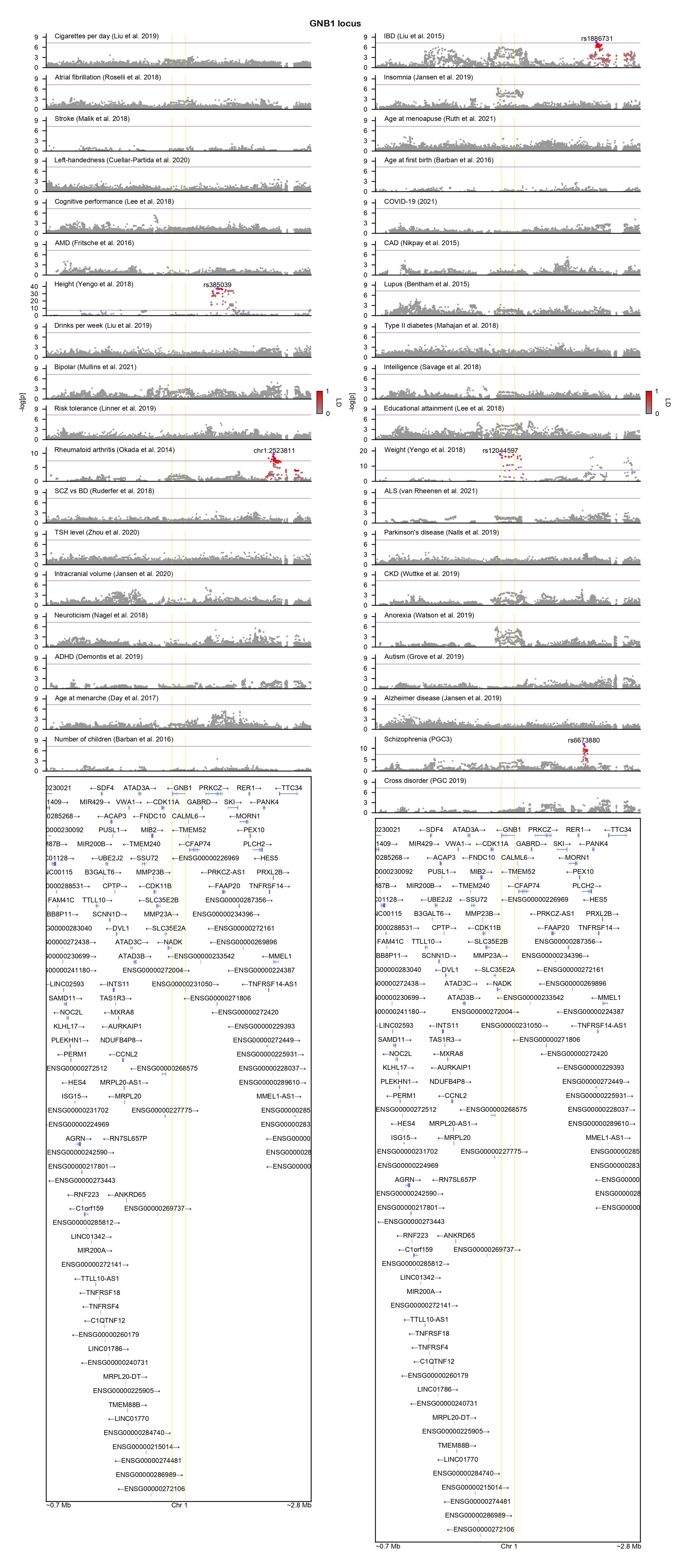

### GNB2-locuszoom.png

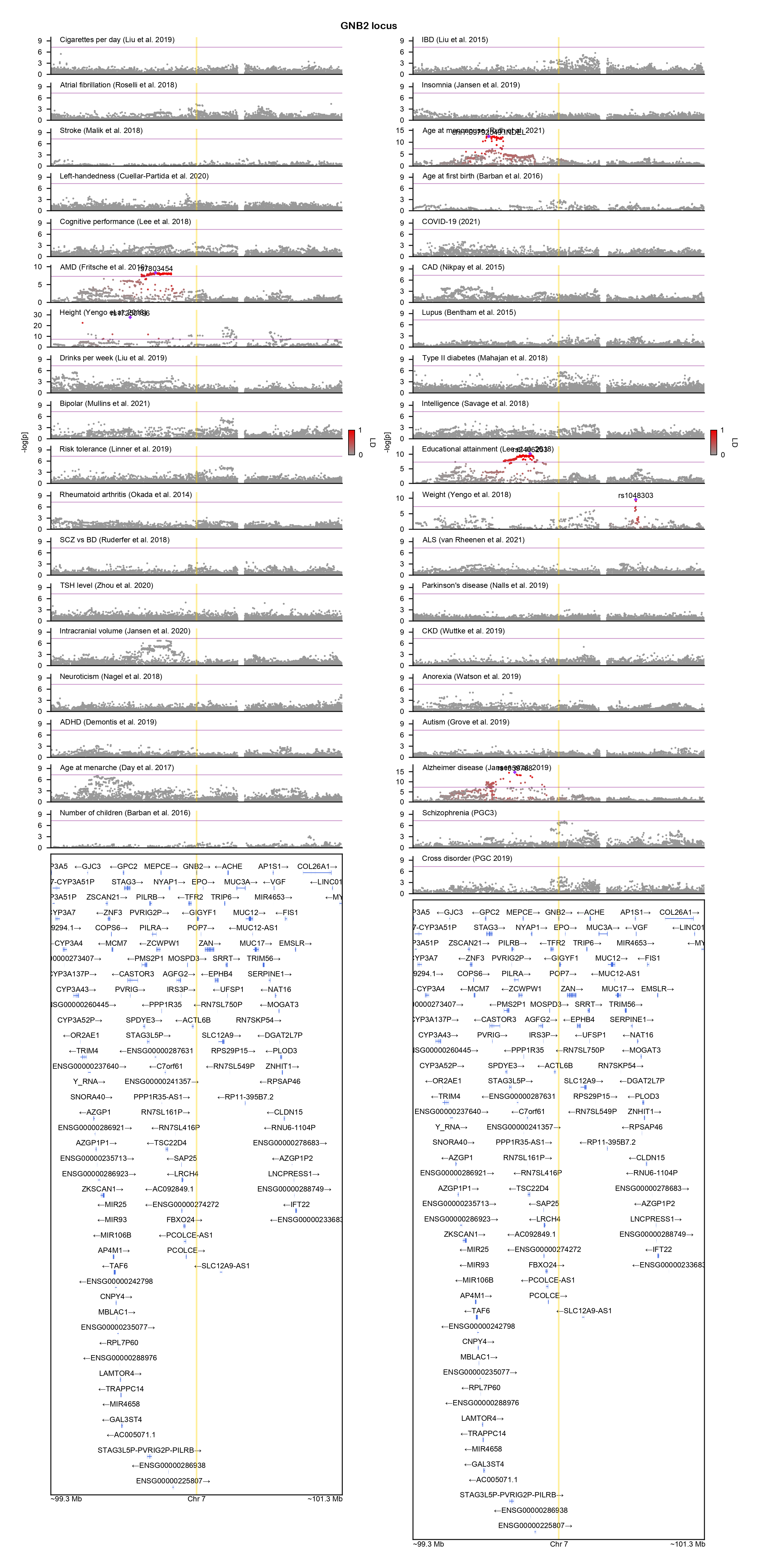

### H1-4-locuszoom.png

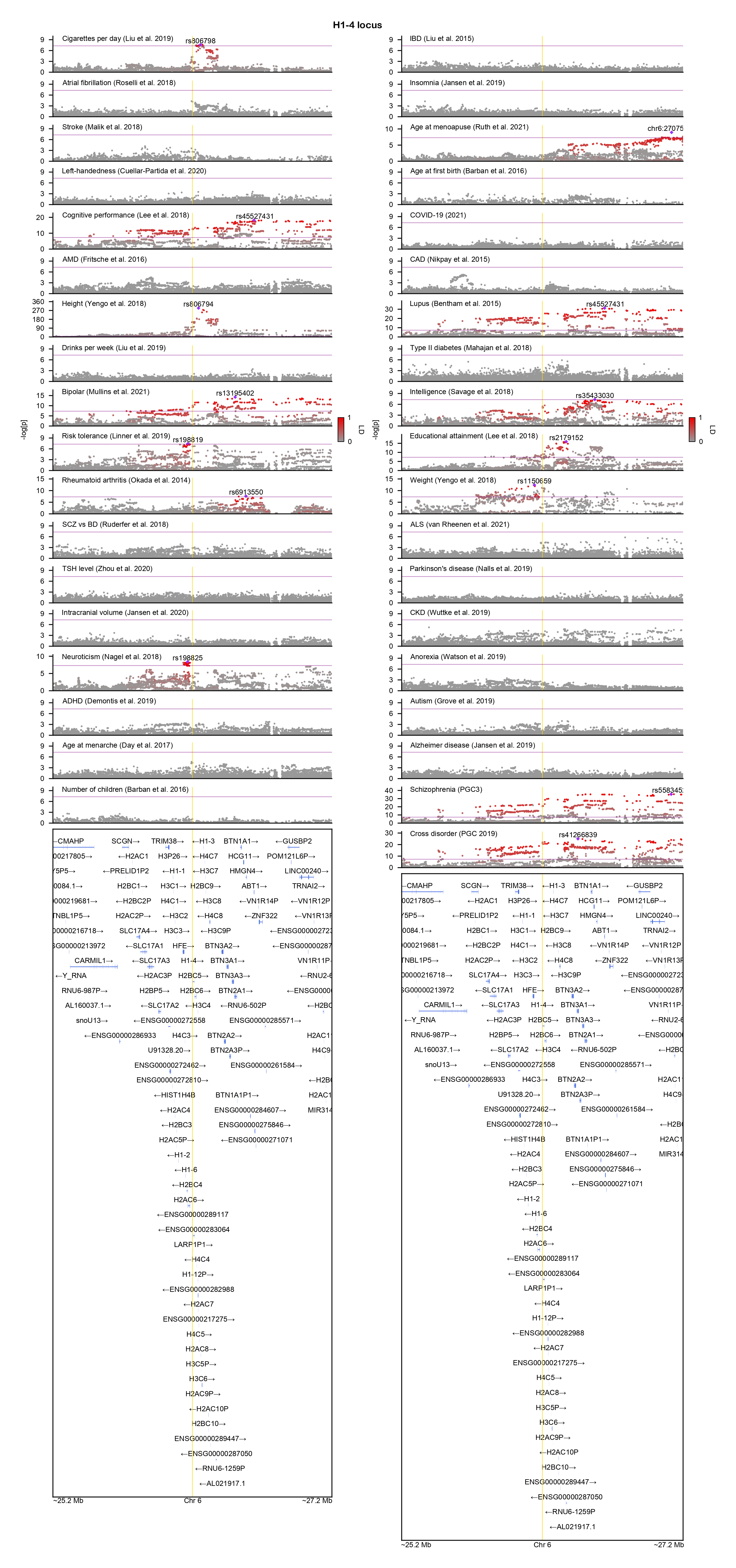

### H3-3A-locuszoom.png

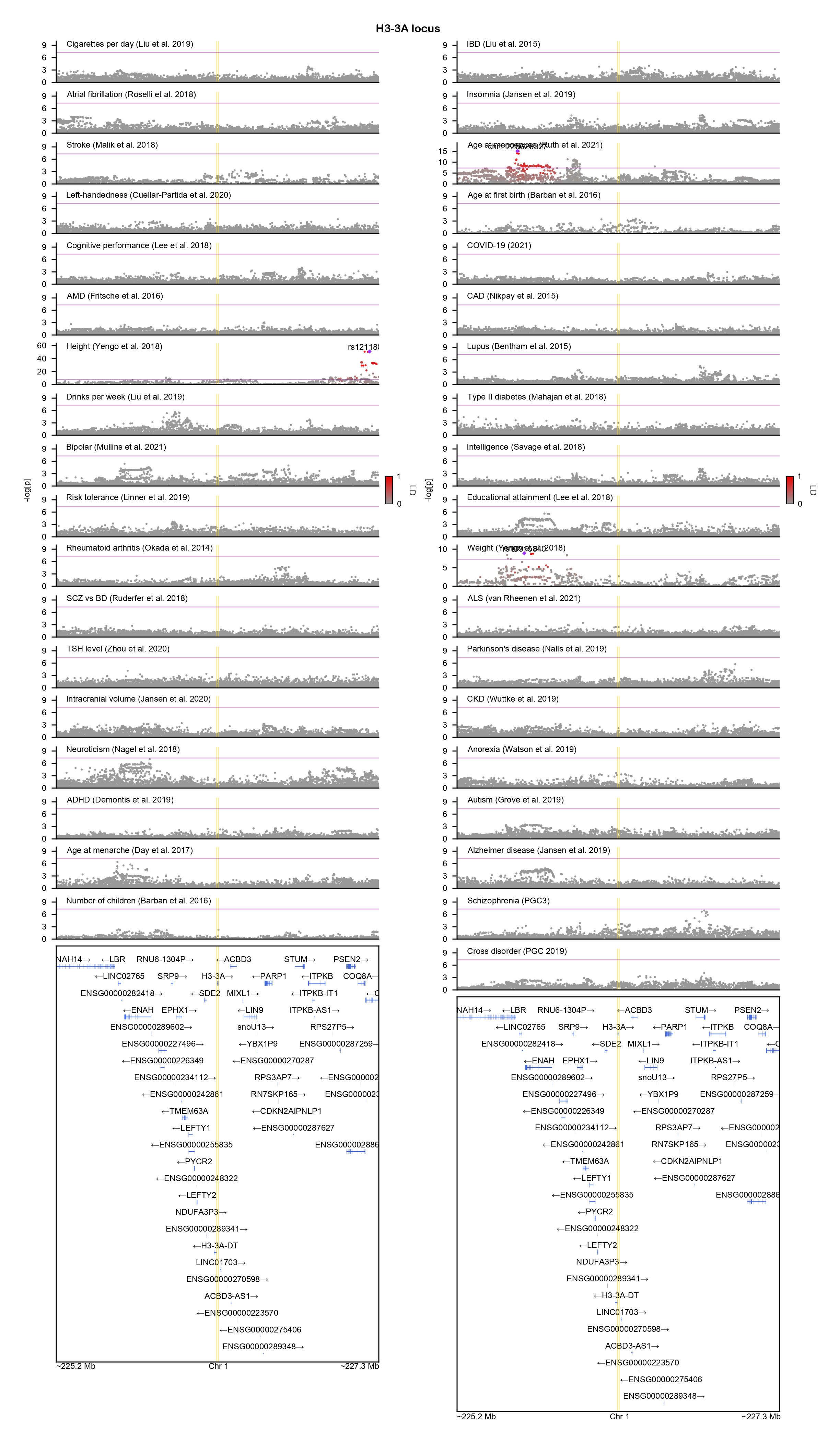

### HDAC4-locuszoom.png

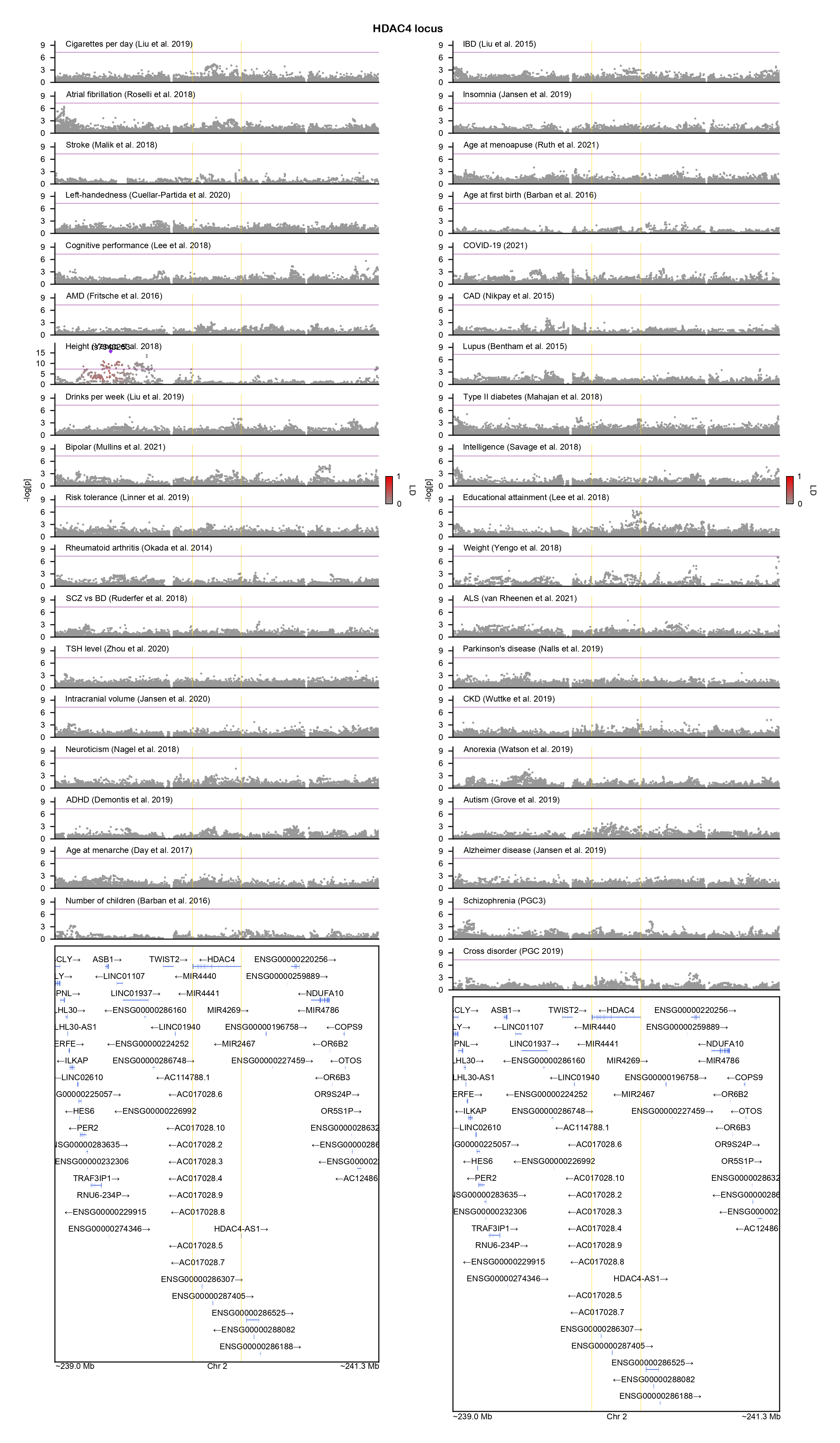

### HECTD4-locuszoom.png

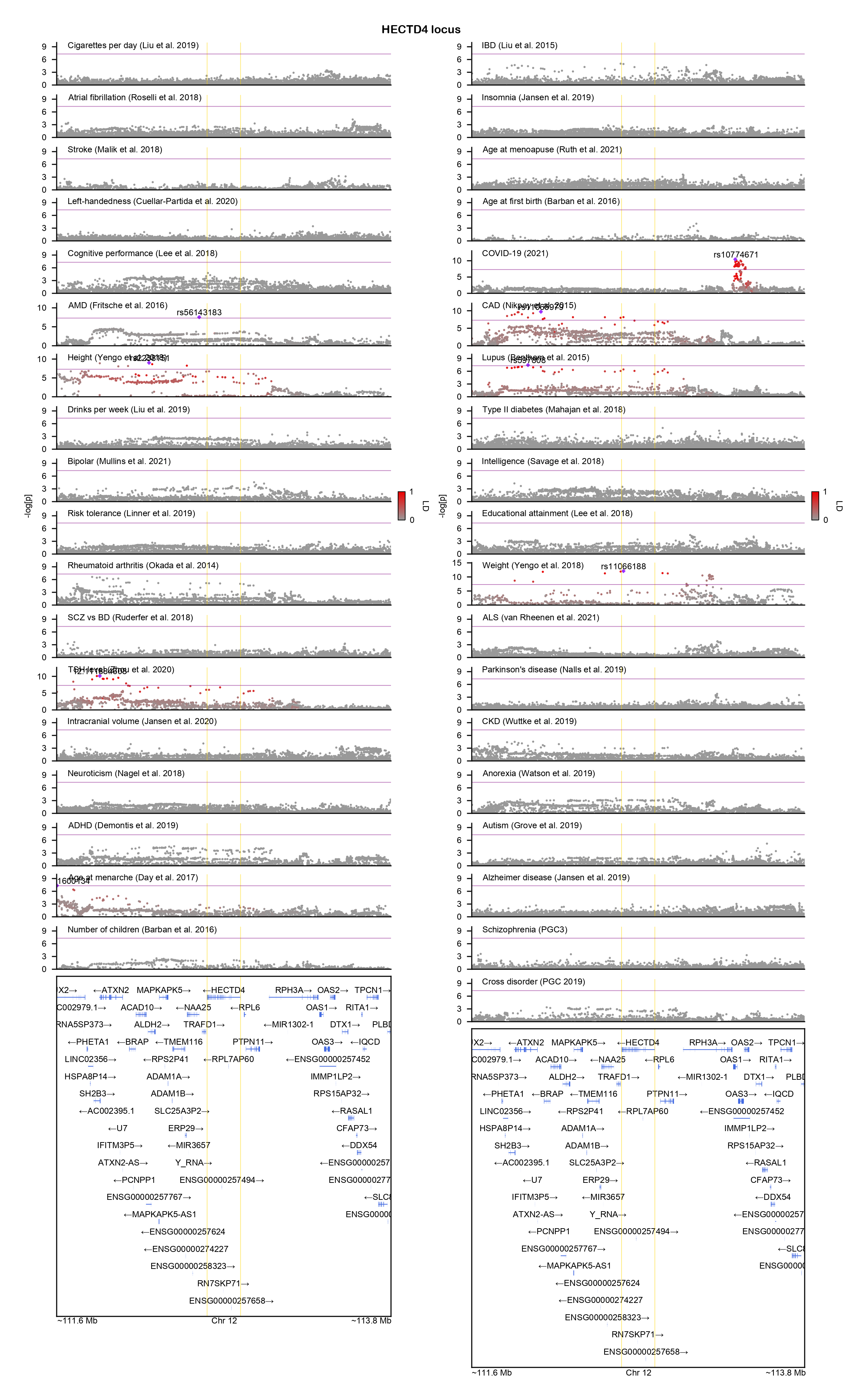

### HERC1-locuszoom.png

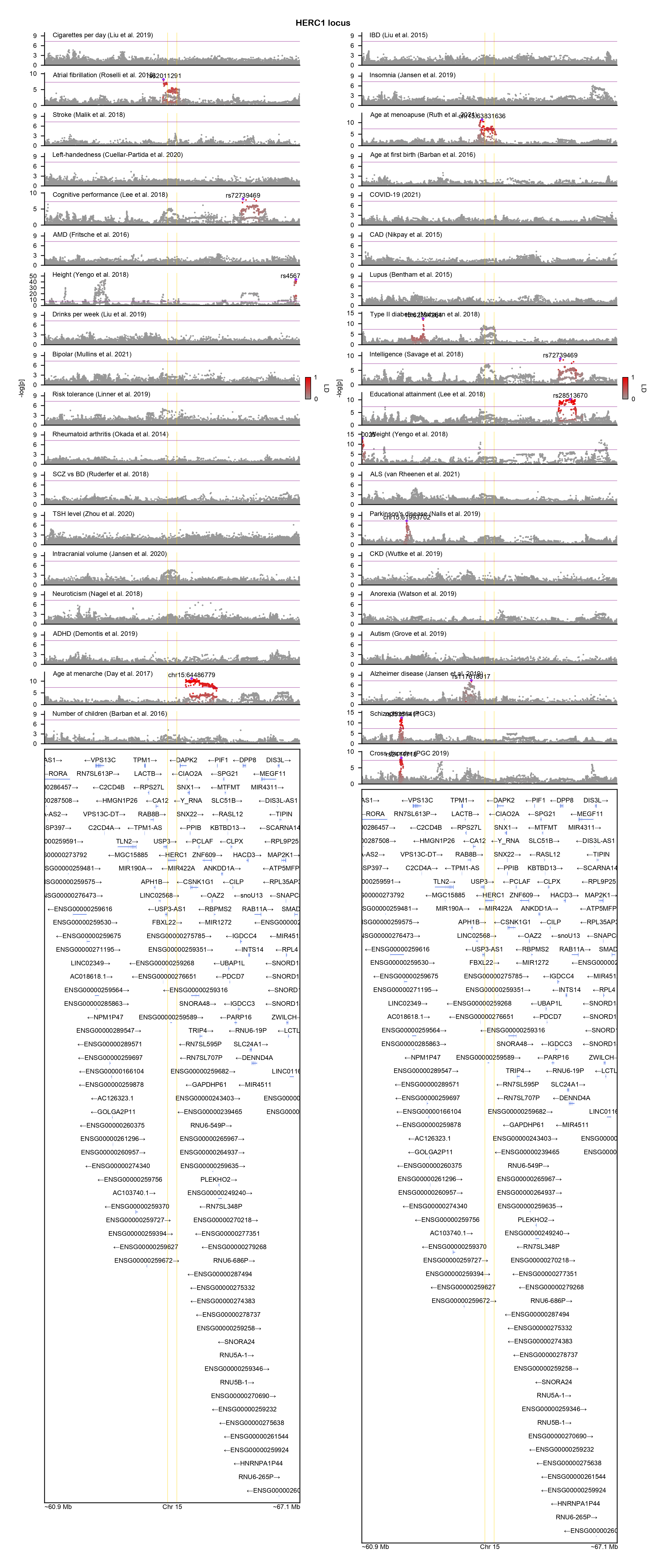

### HIVEP2-locuszoom.png

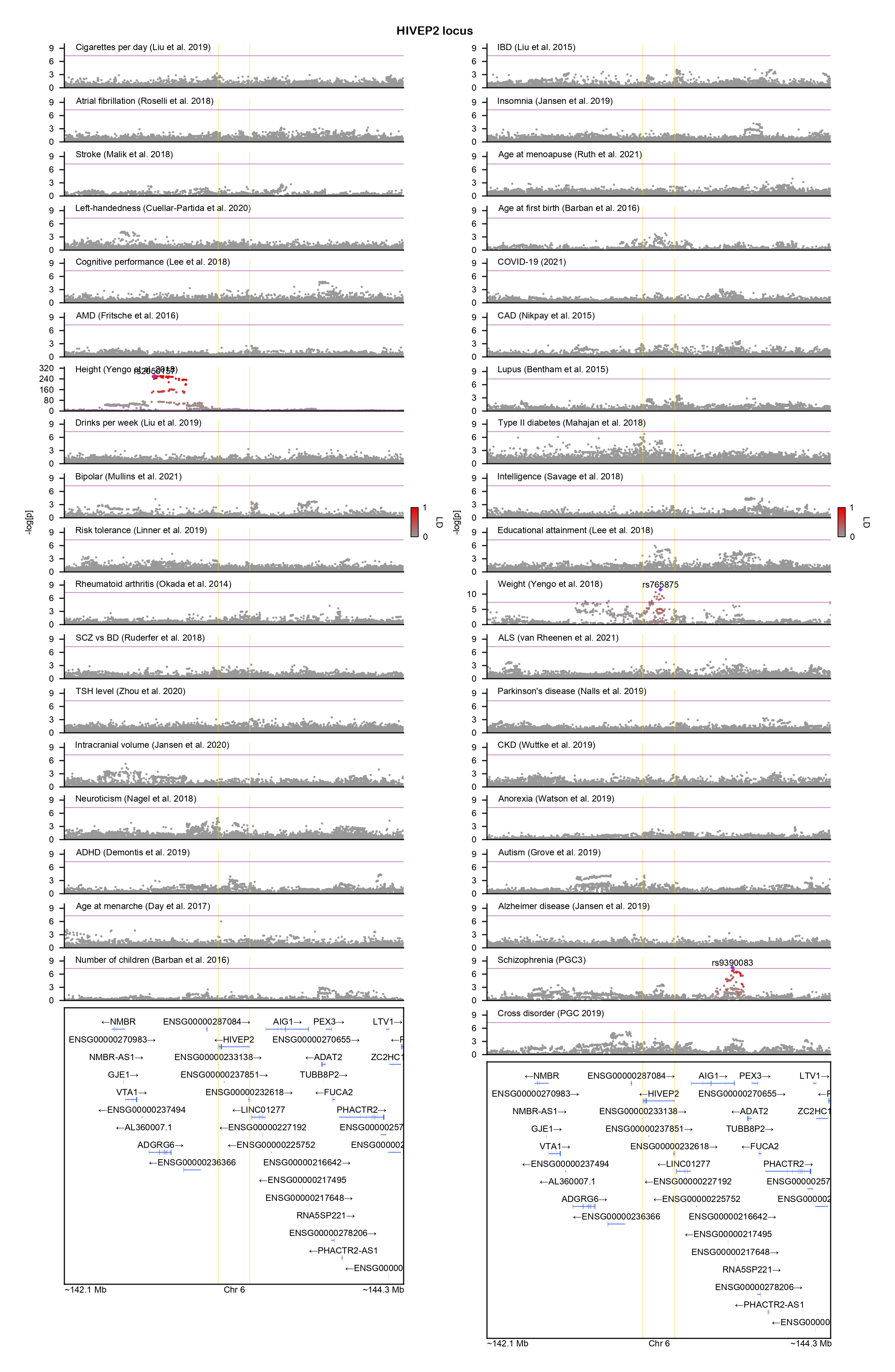

### HNRNPK-locuszoom.png

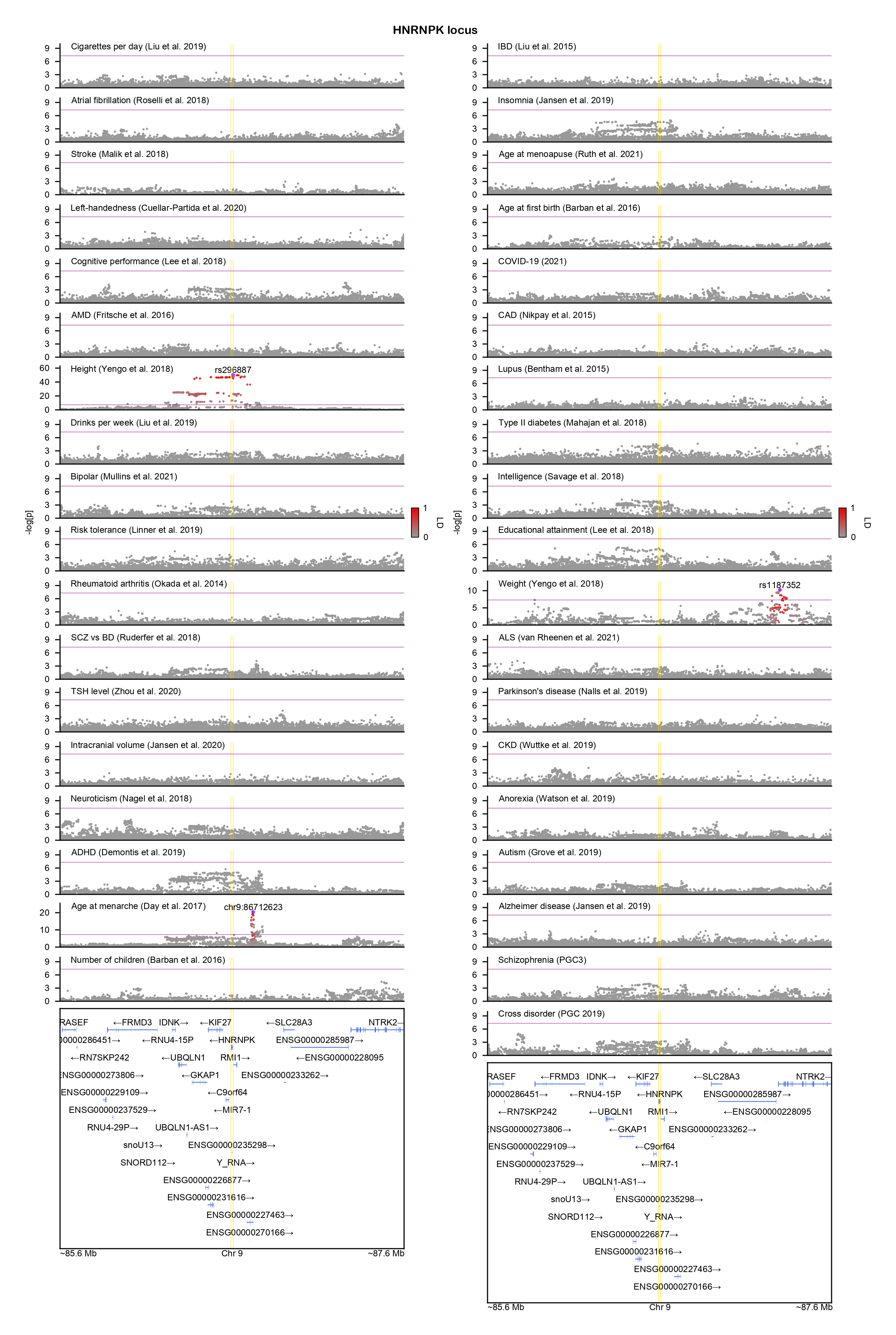

### HUWE1-locuszoom.png

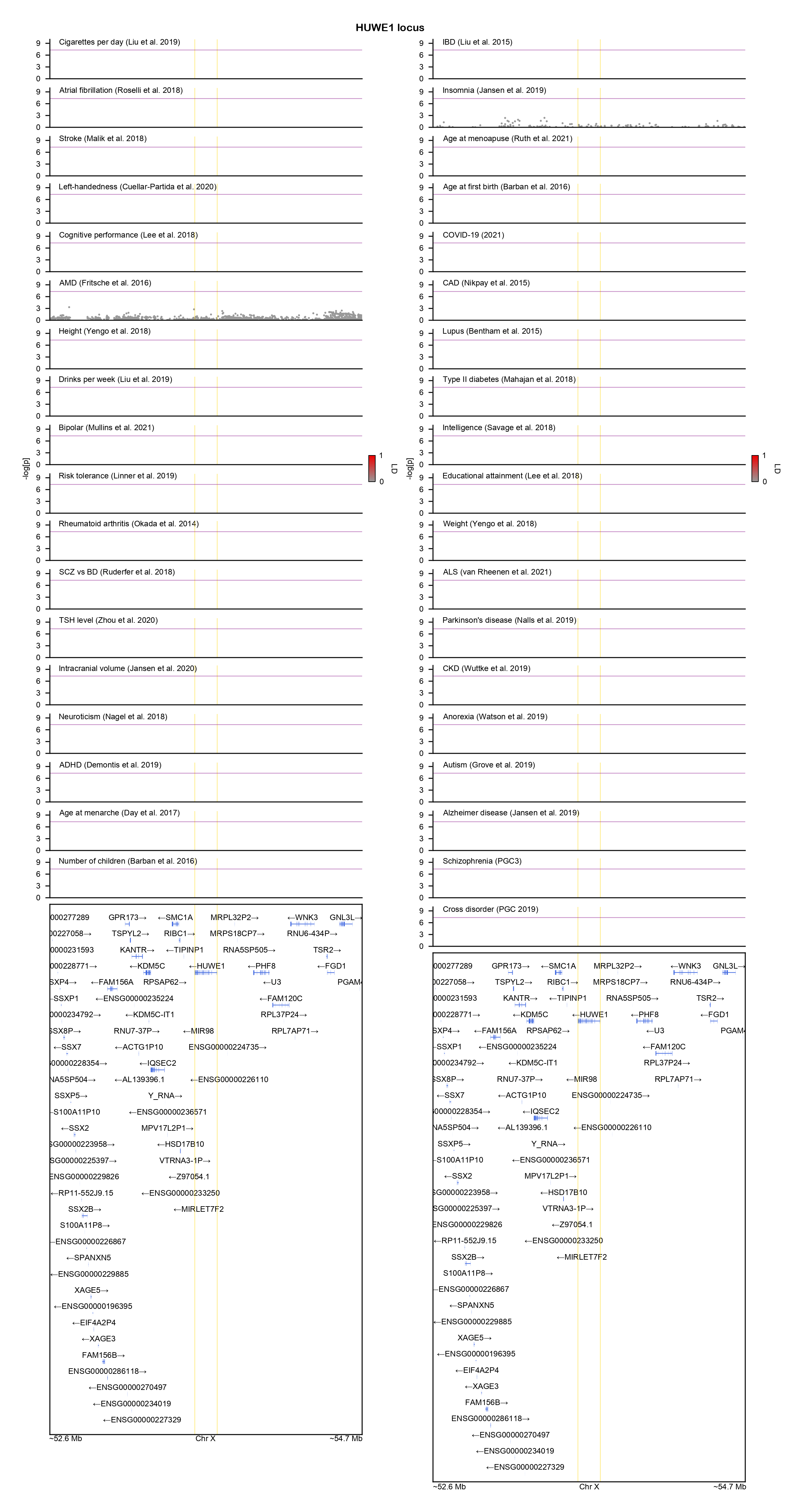

### IQSEC2-locuszoom.png

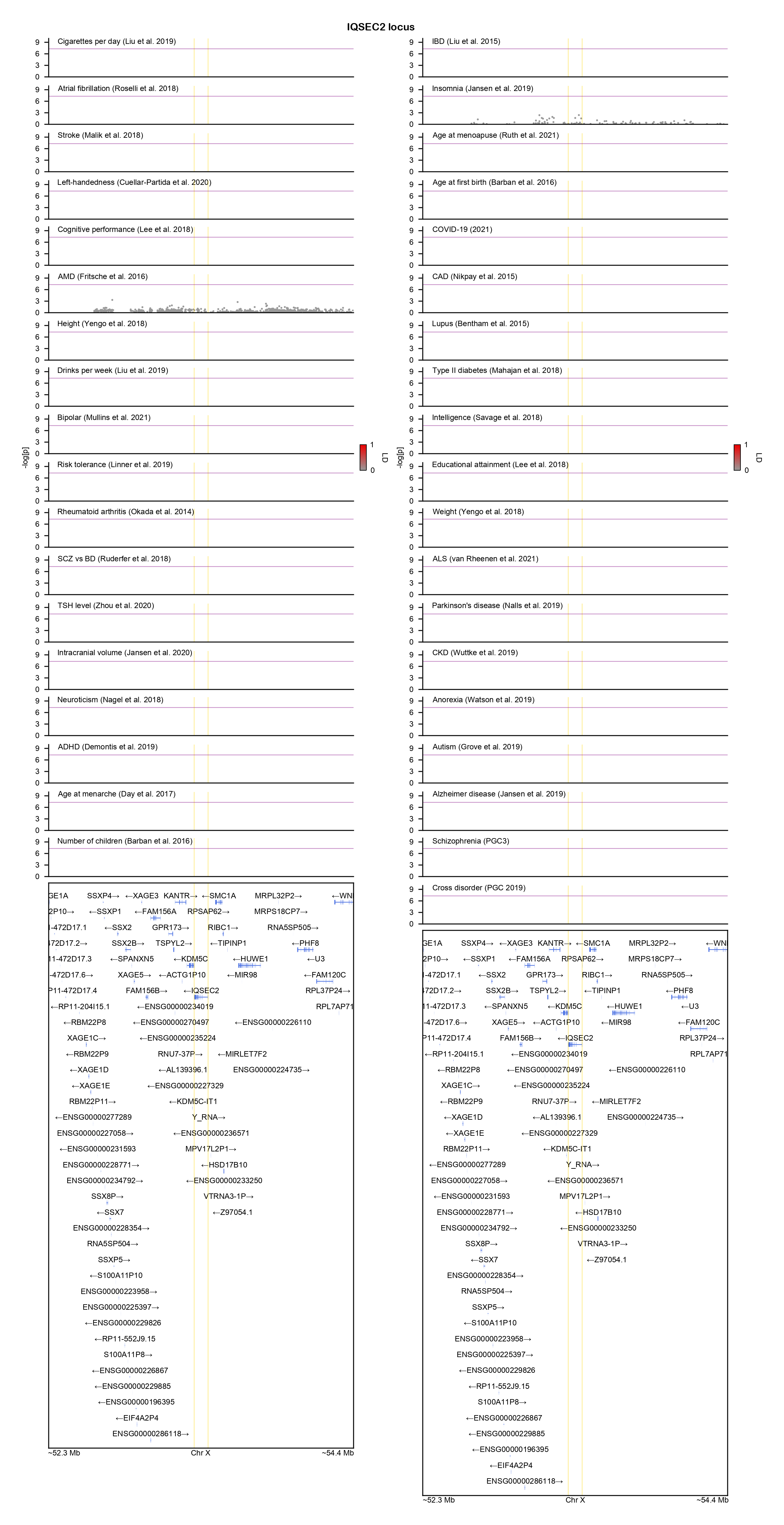

### KCNA2-locuszoom.png

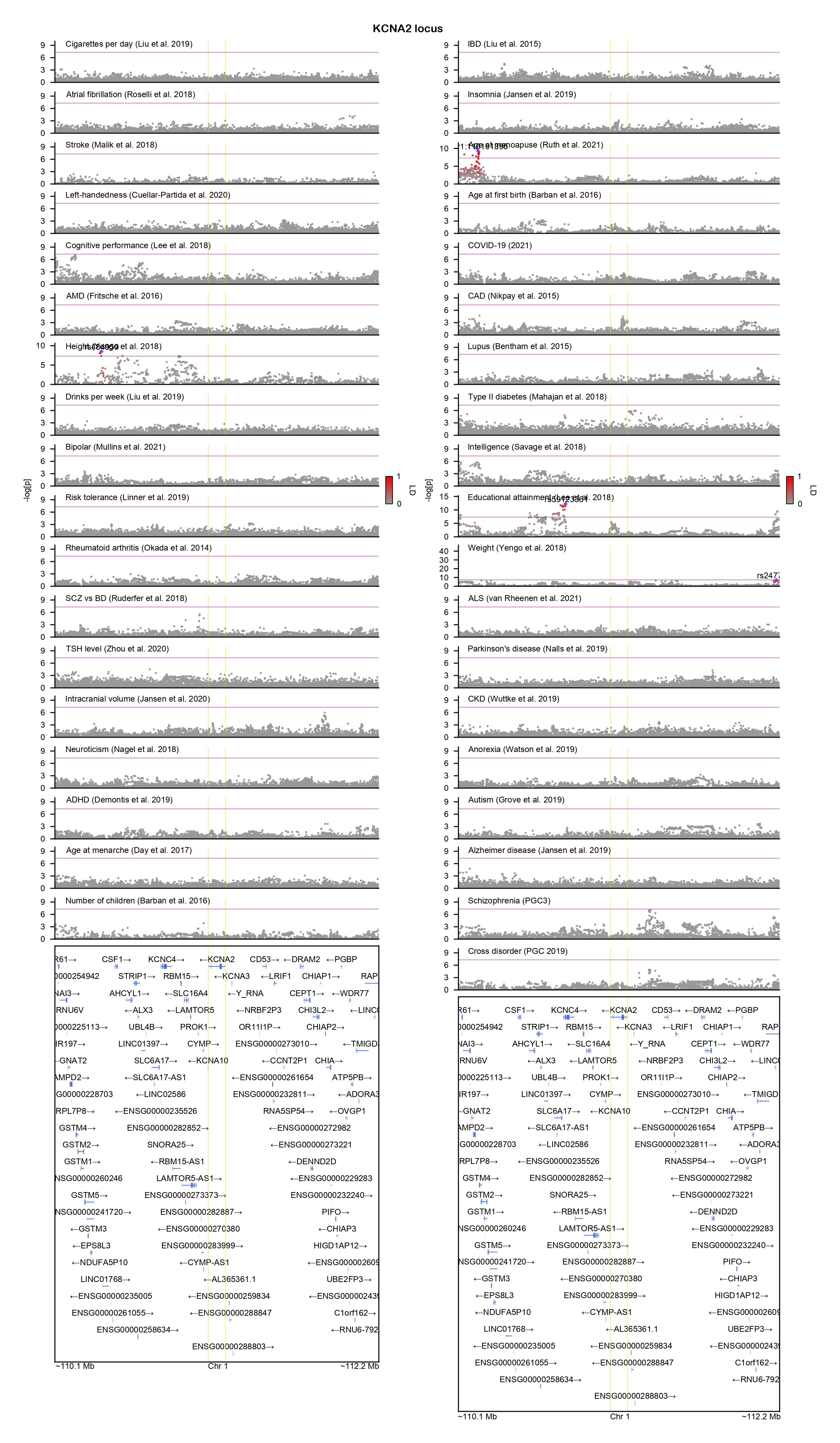

### KCNH1-locuszoom.png

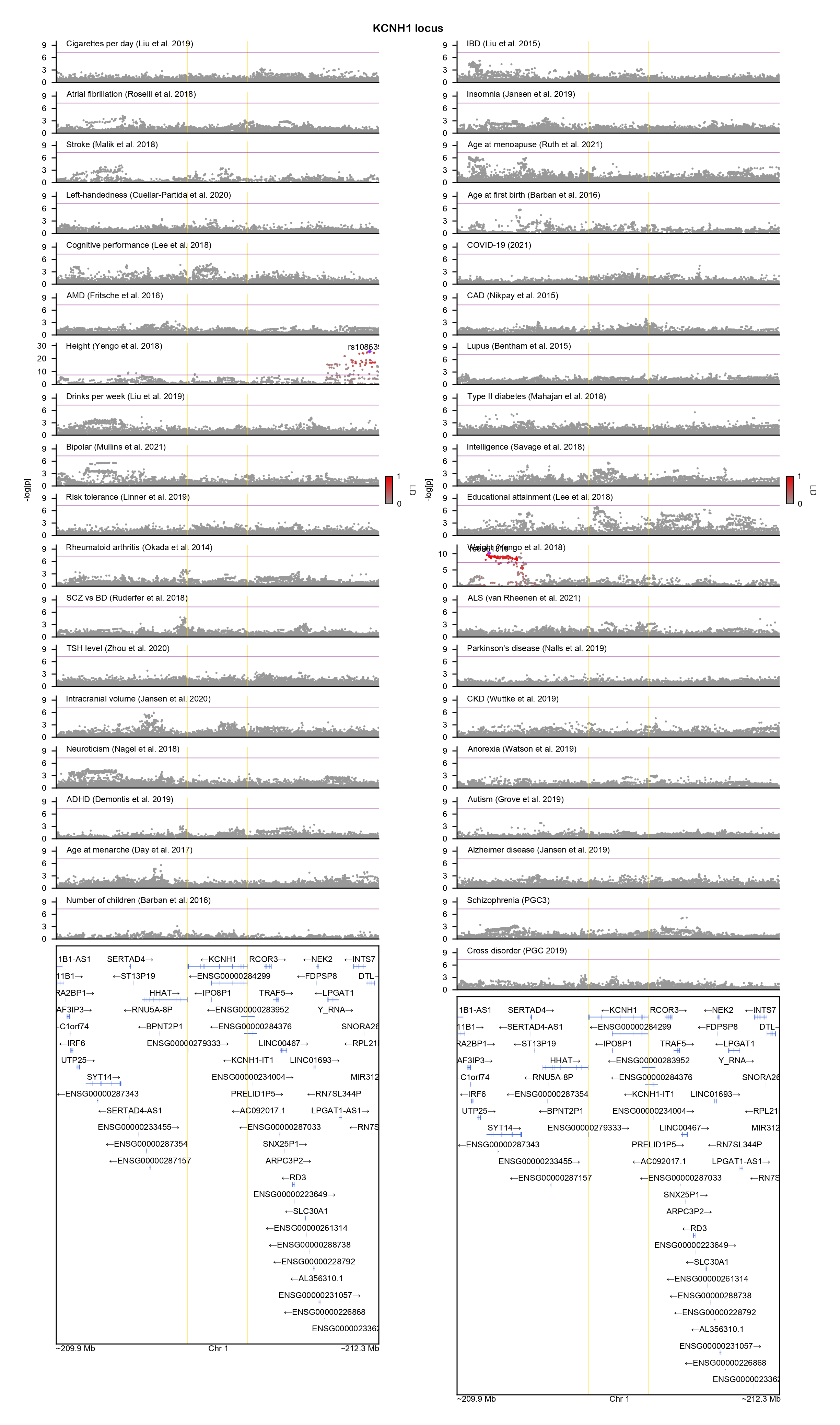

### KCNMA1-locuszoom.png

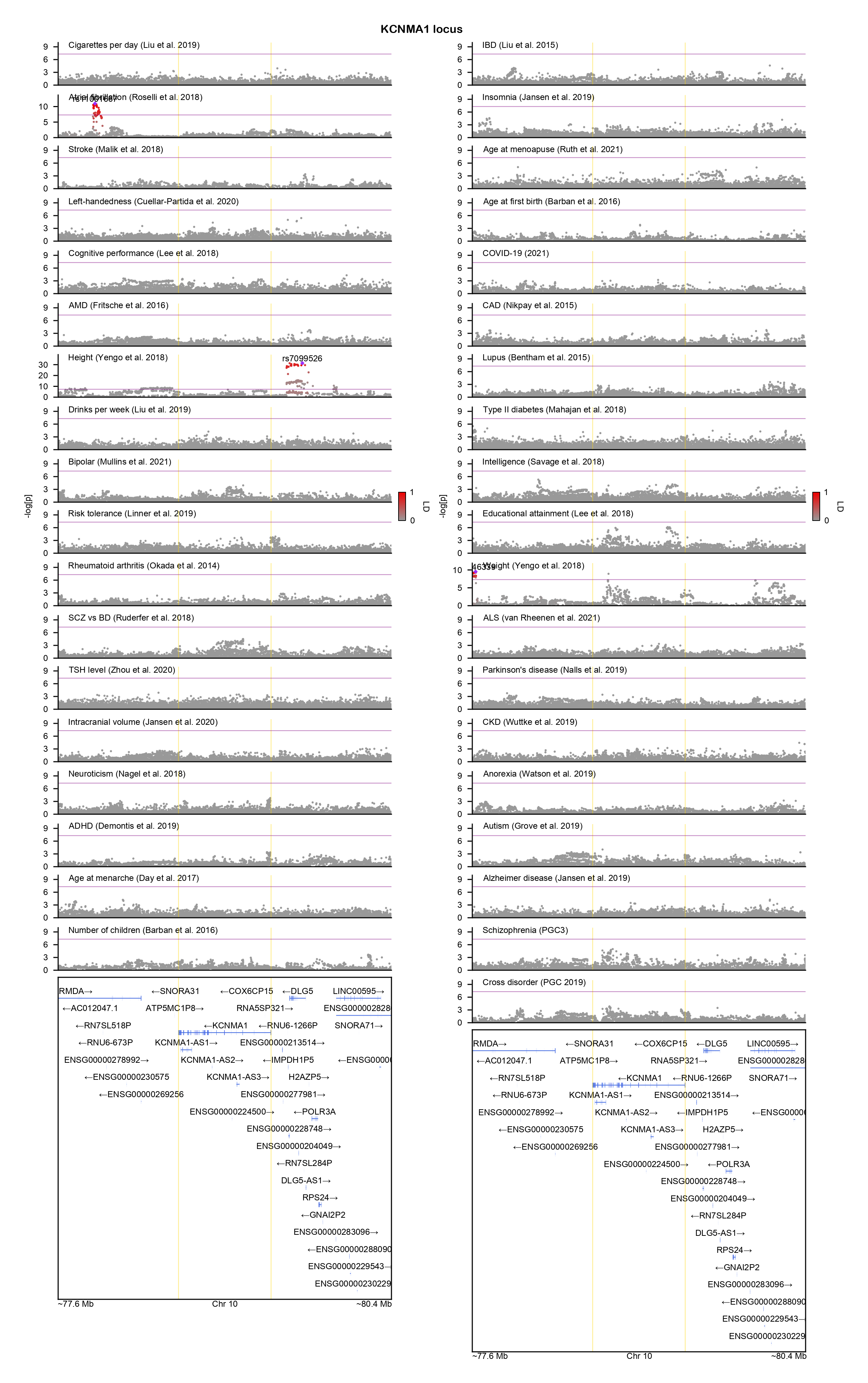

### KDM5B-locuszoom.png

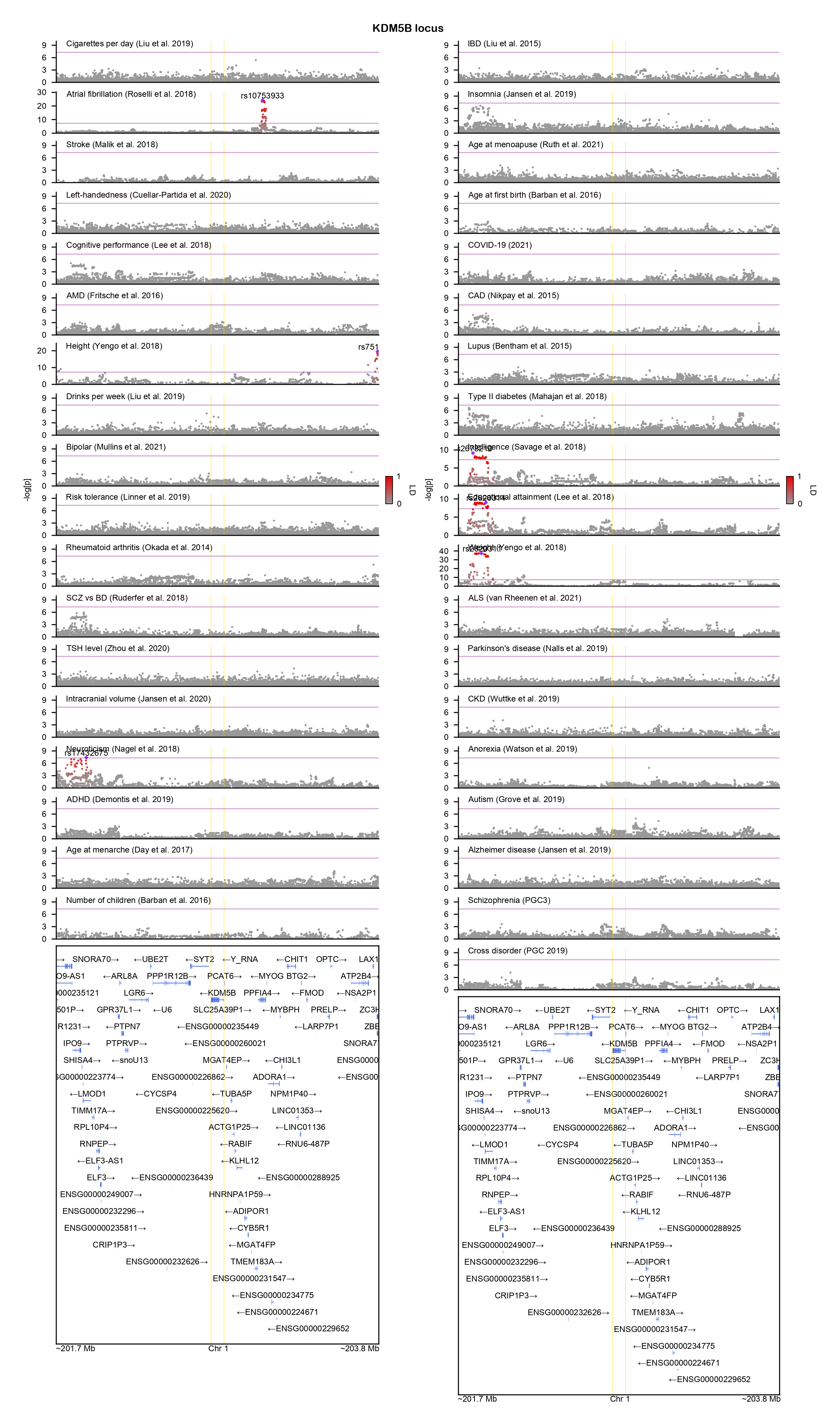

### KIF5C-locuszoom.png

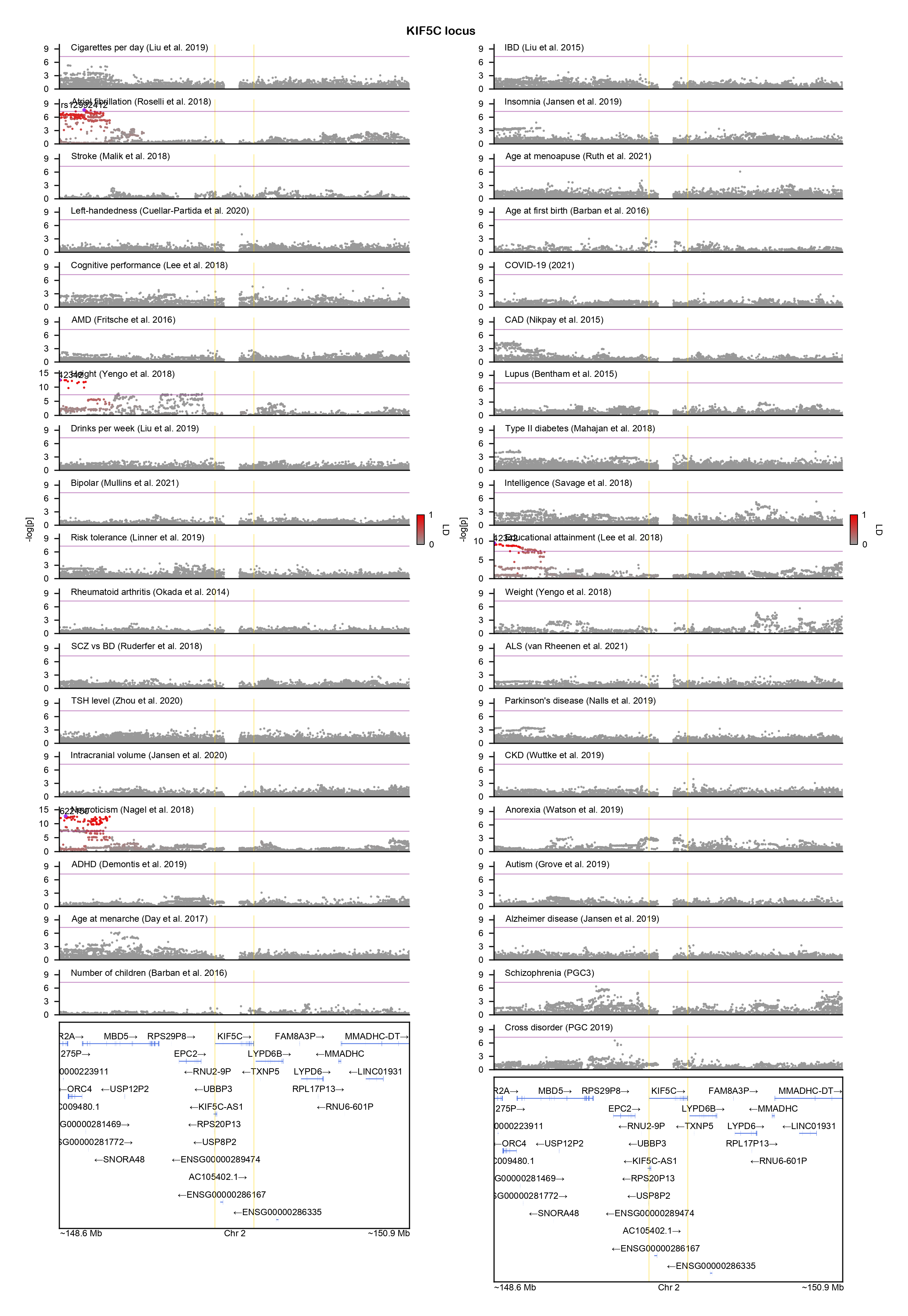

### KMT2A-locuszoom.png

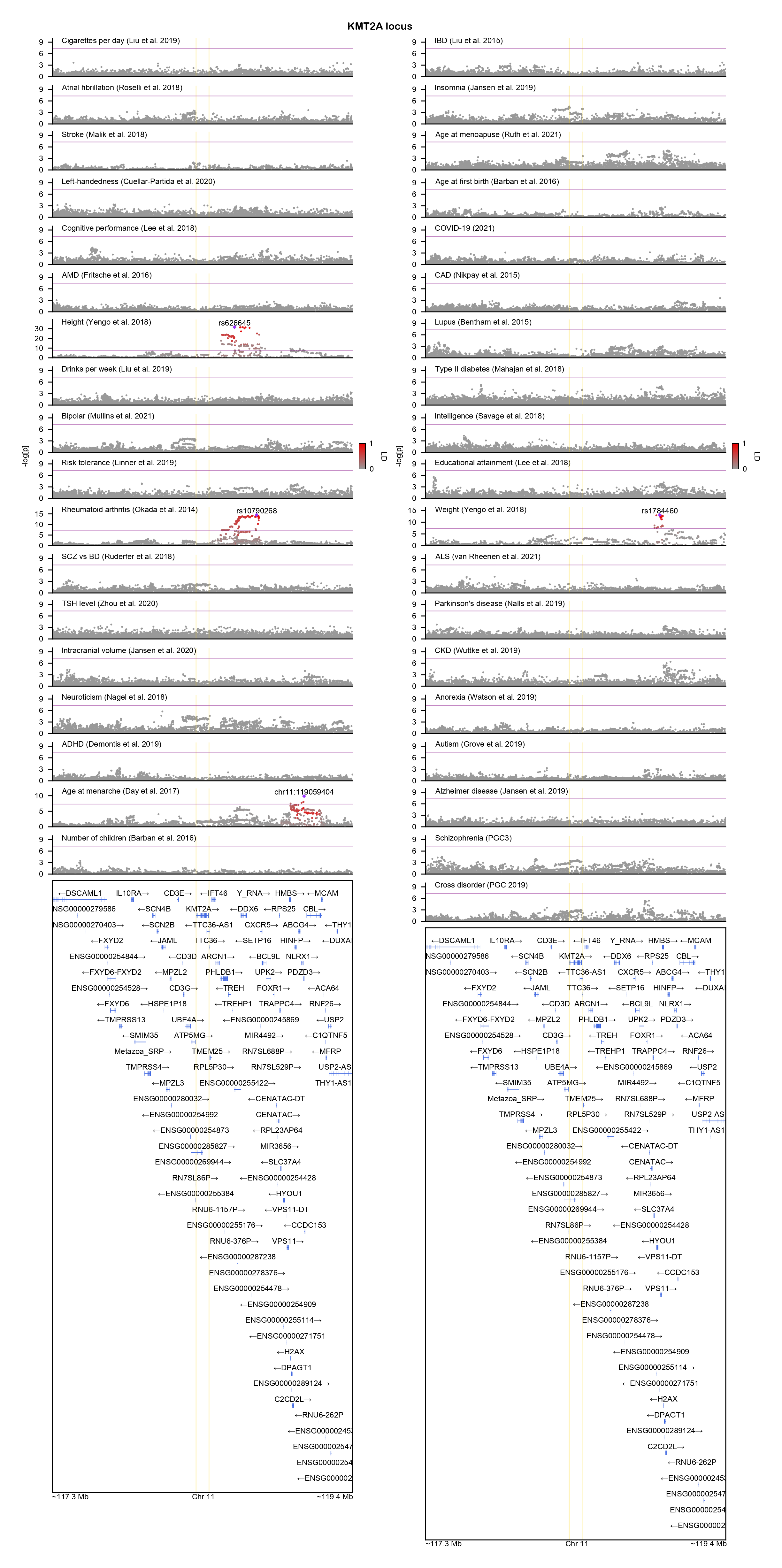

### KMT2B-locuszoom.png

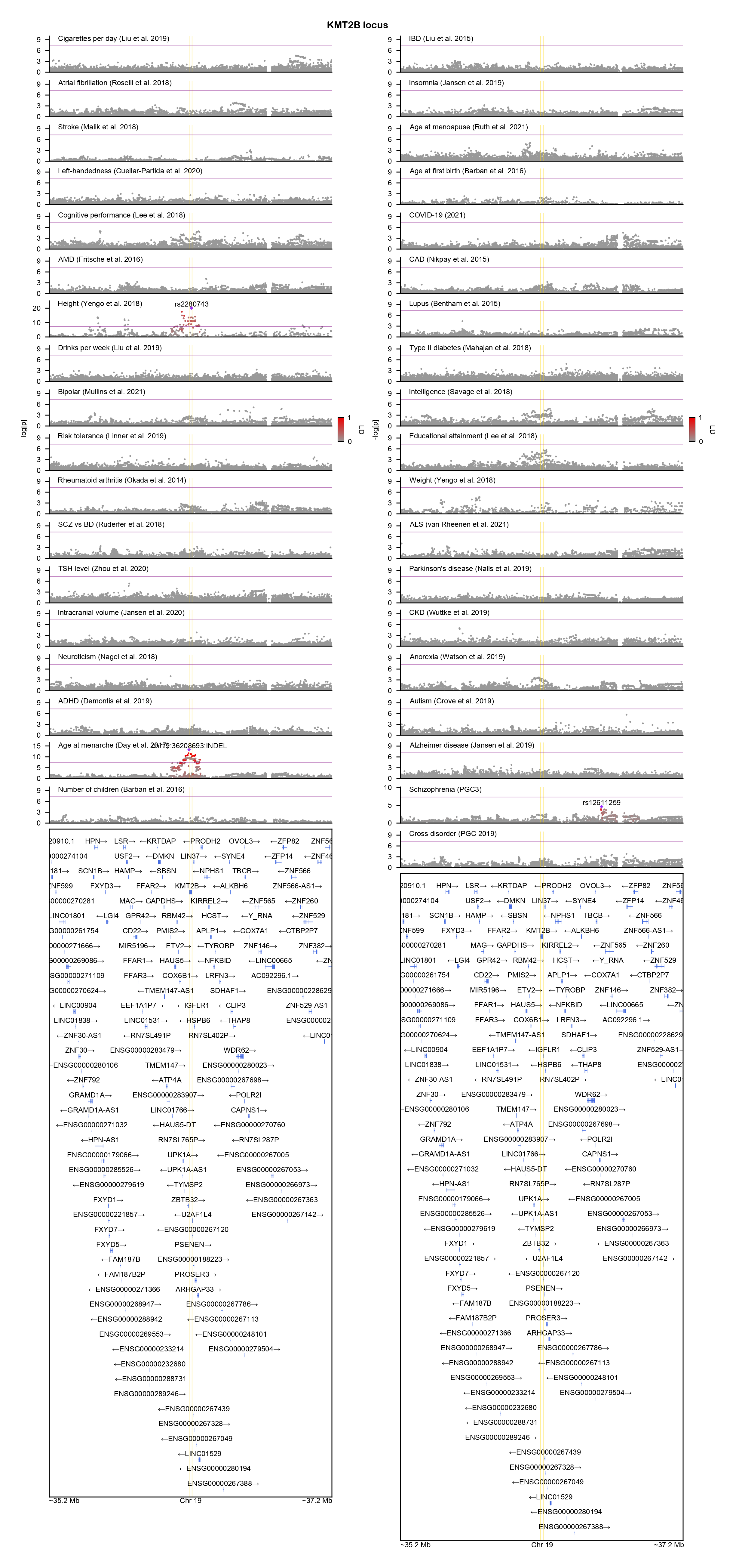

### KMT5B-locuszoom.png

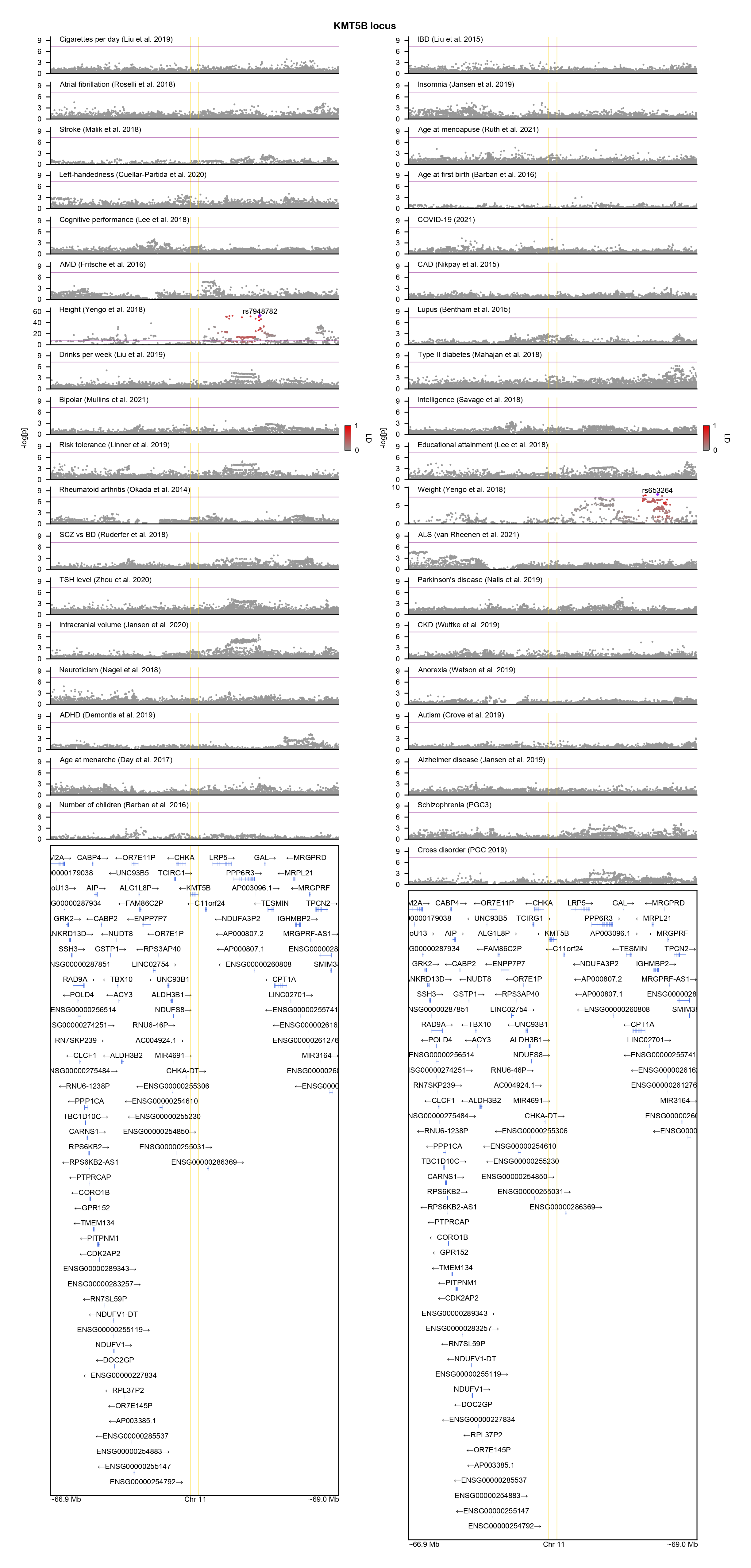

### LRRC4C-locuszoom.png

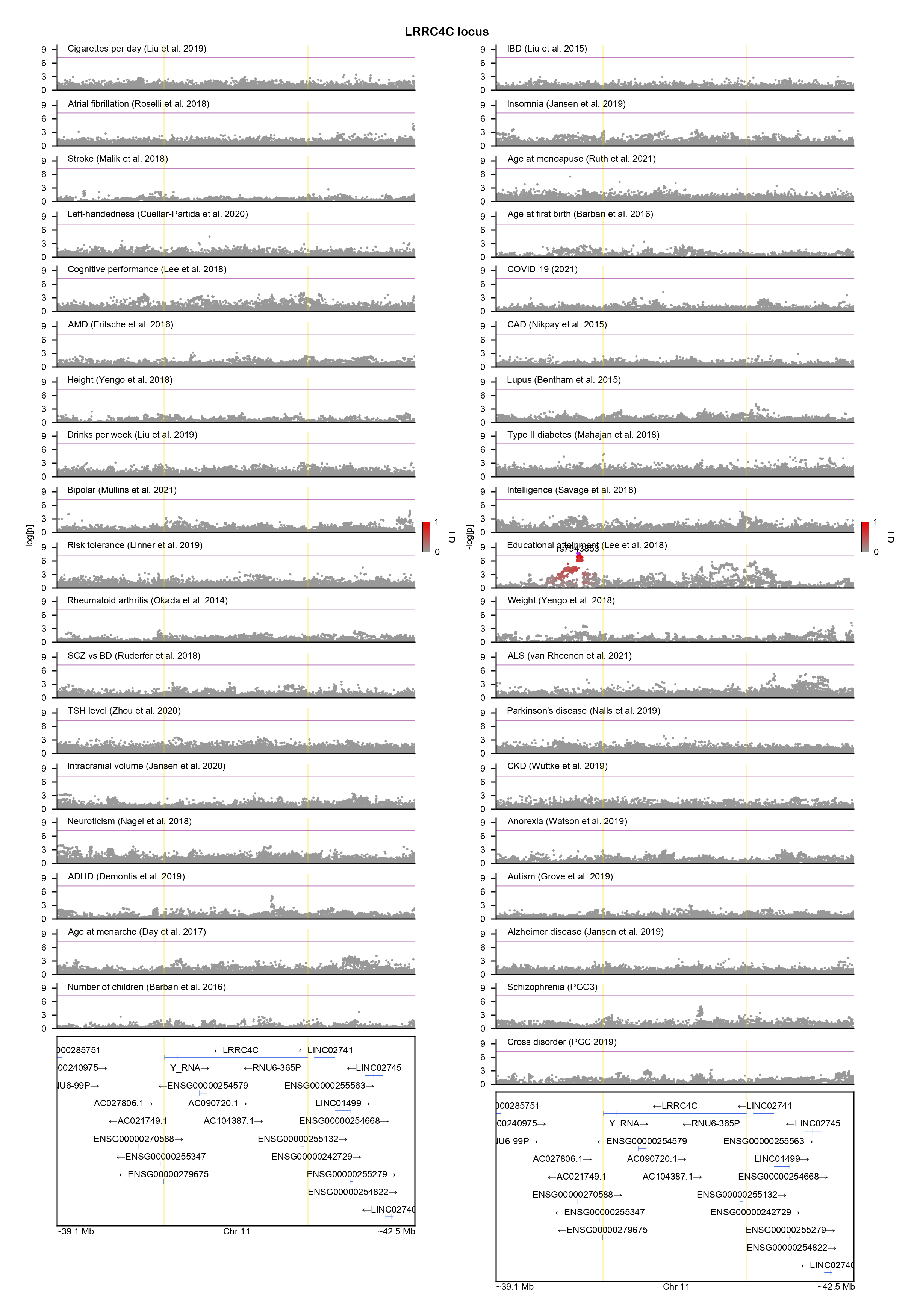

### LZTR1-locuszoom.png

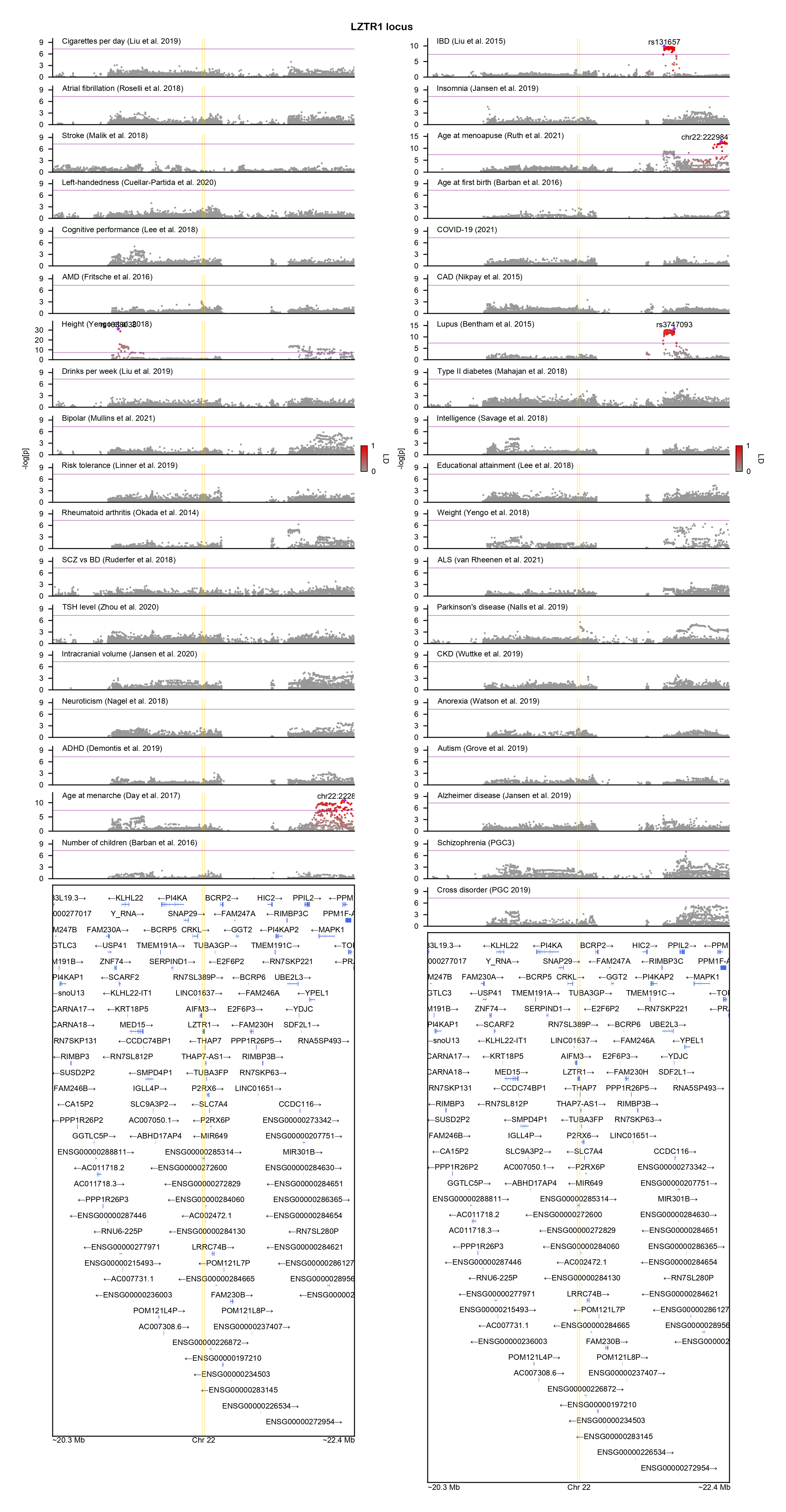

### MAGEL2-locuszoom.png

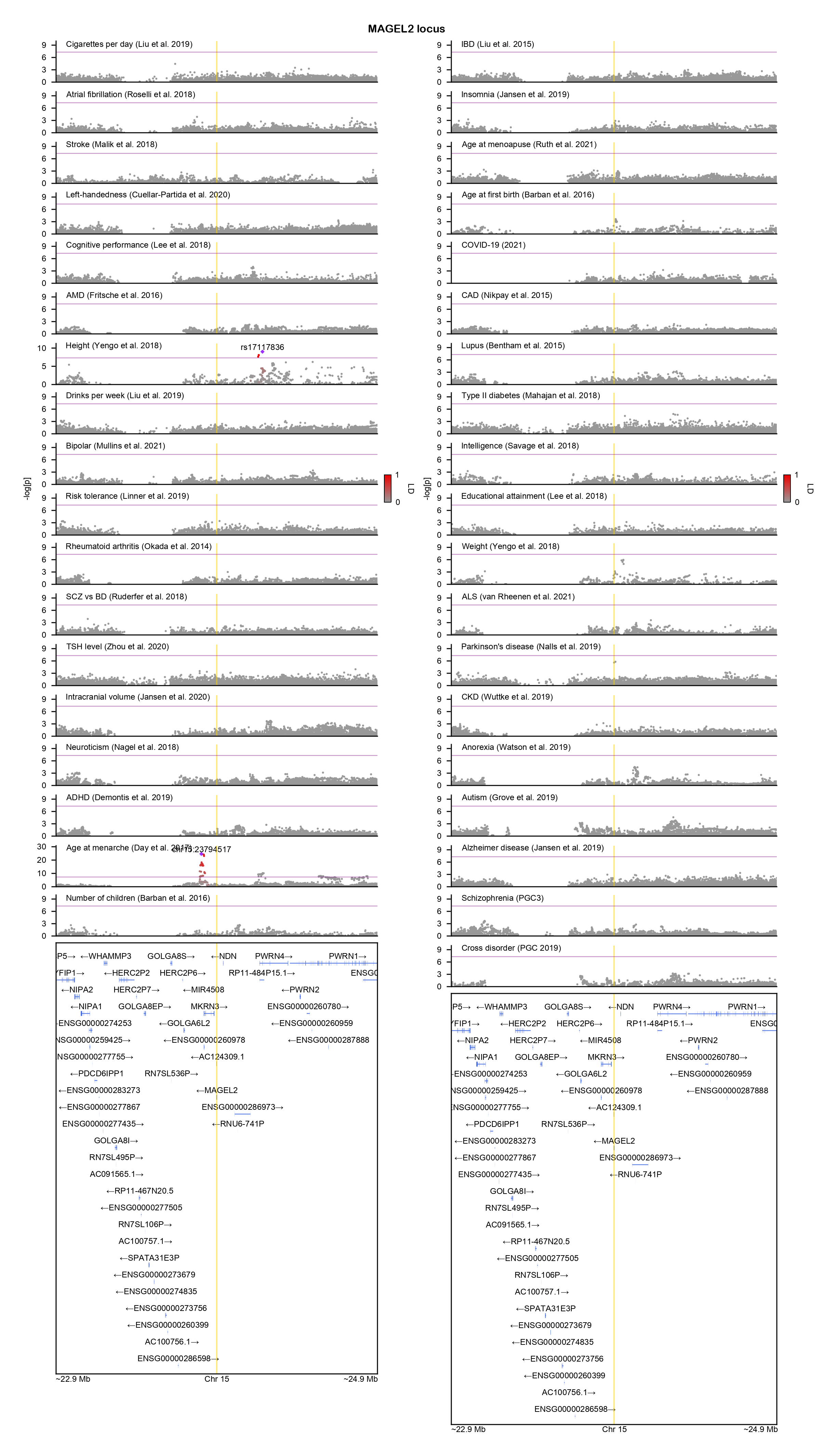

### MAP2K1-locuszoom.png

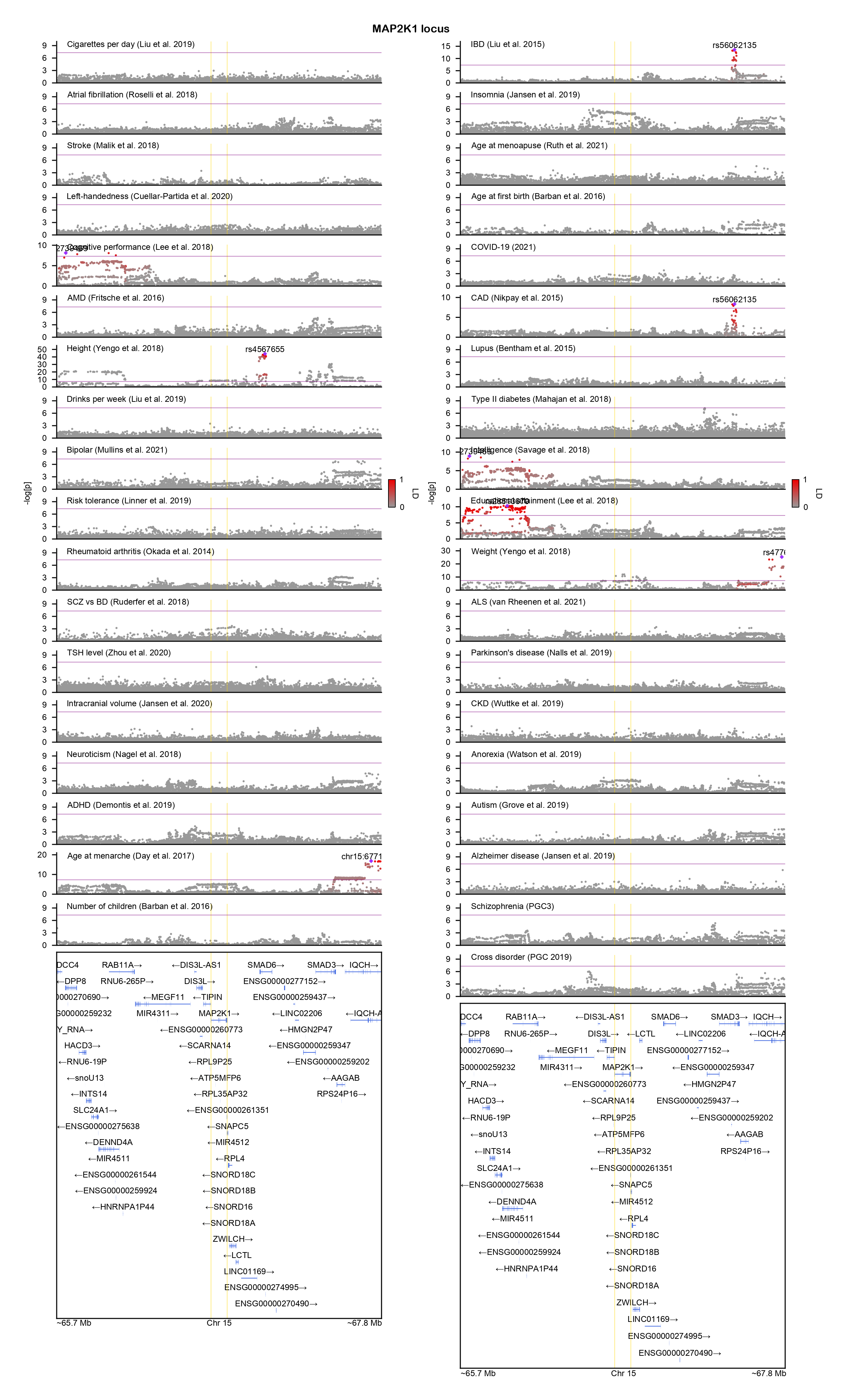

### MAST1-locuszoom.png

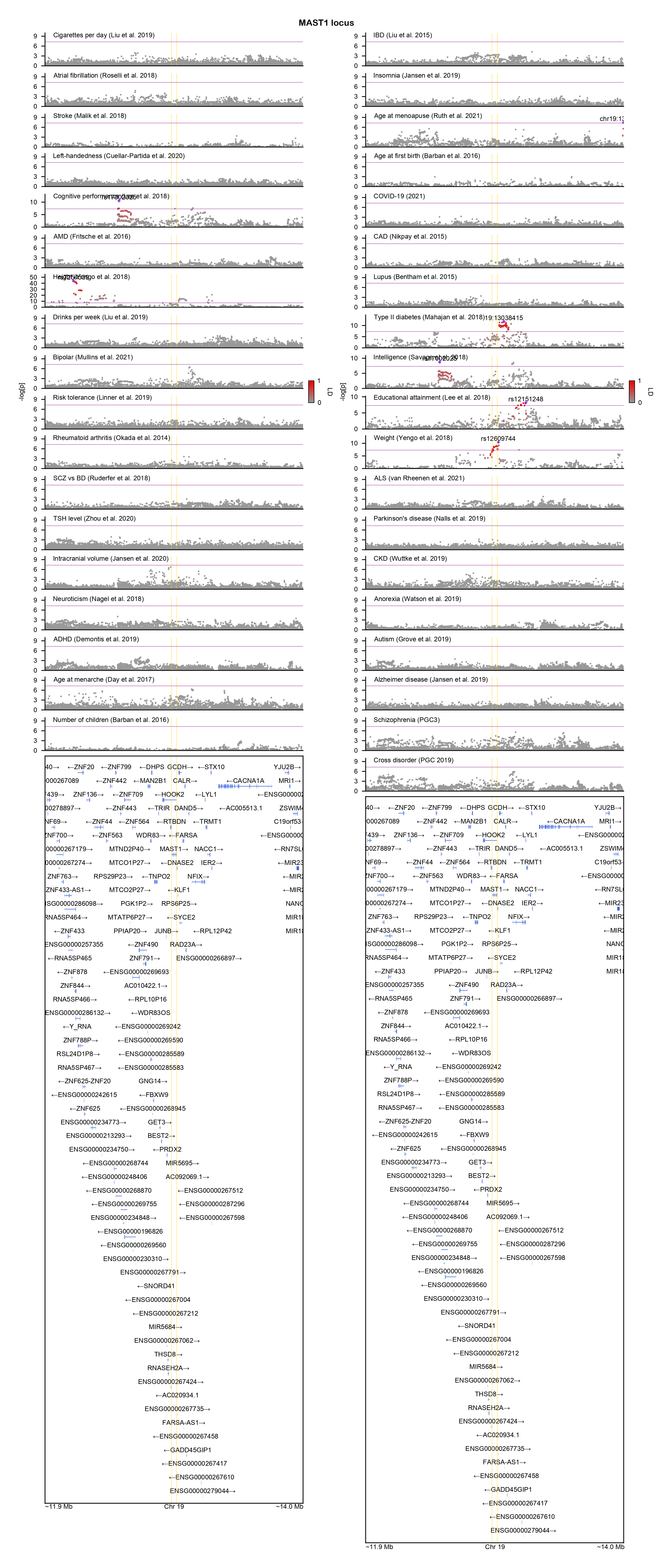

### MED13L-locuszoom.png

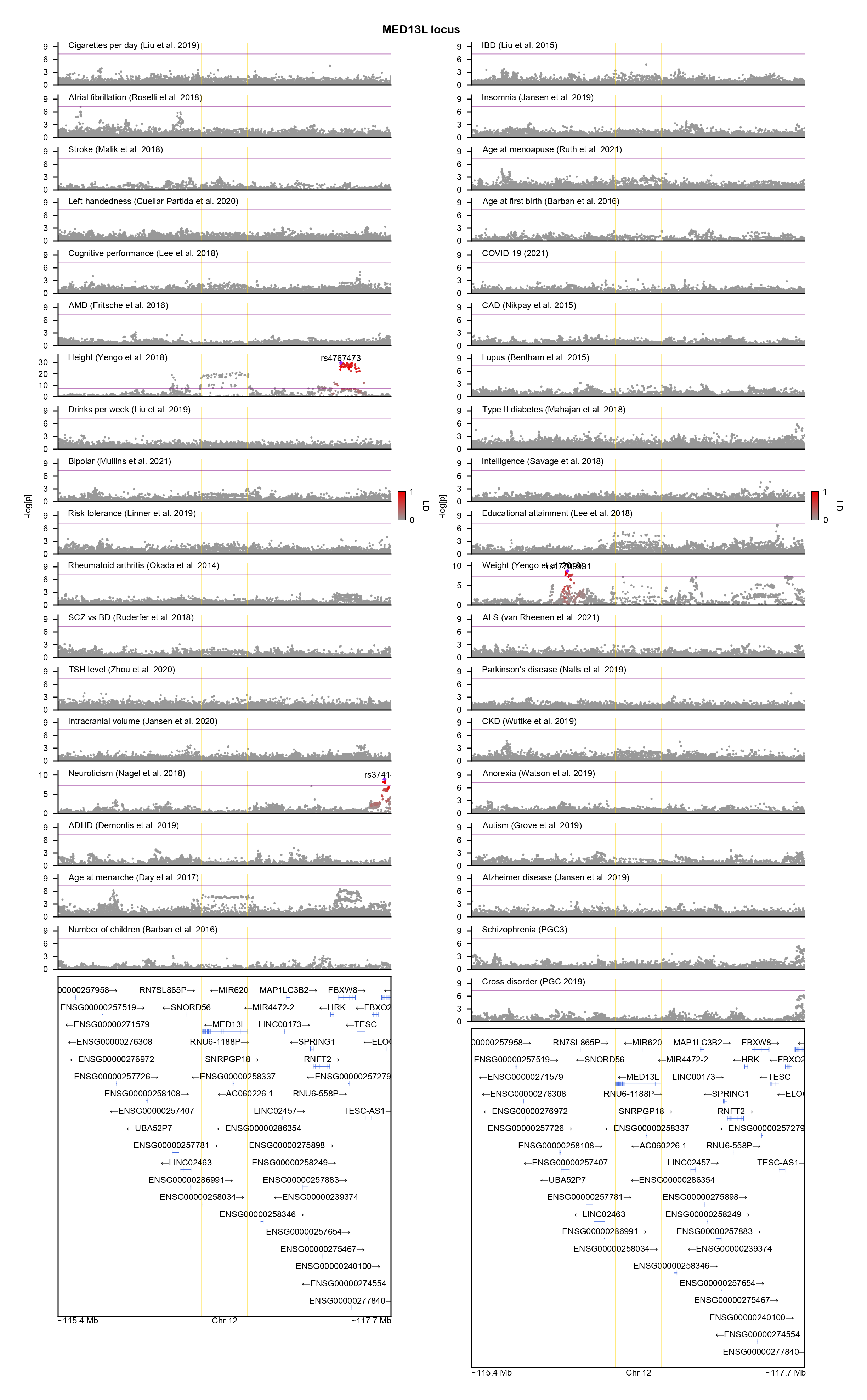

### MEIS2-locuszoom.png

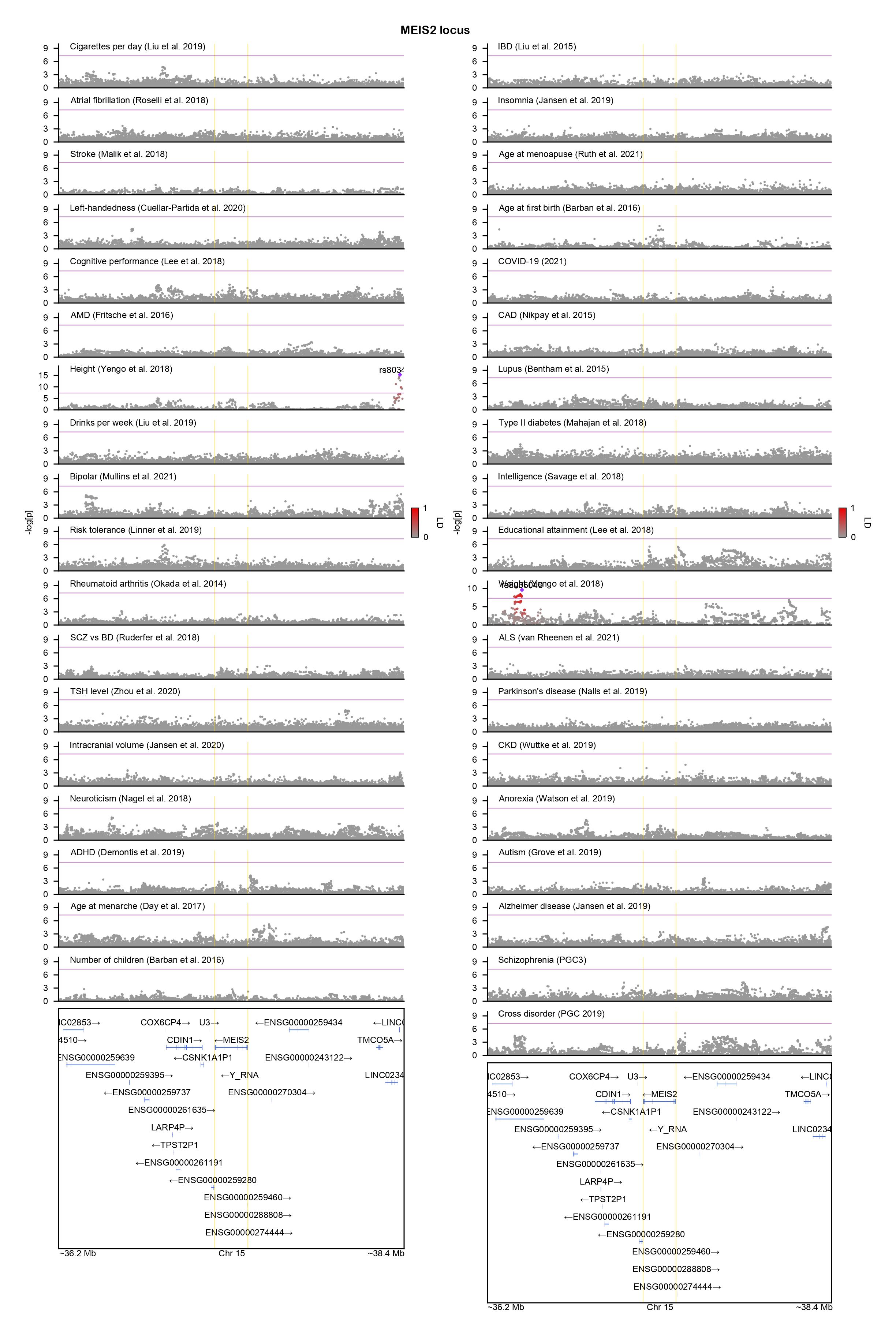

### MFN2-locuszoom.png

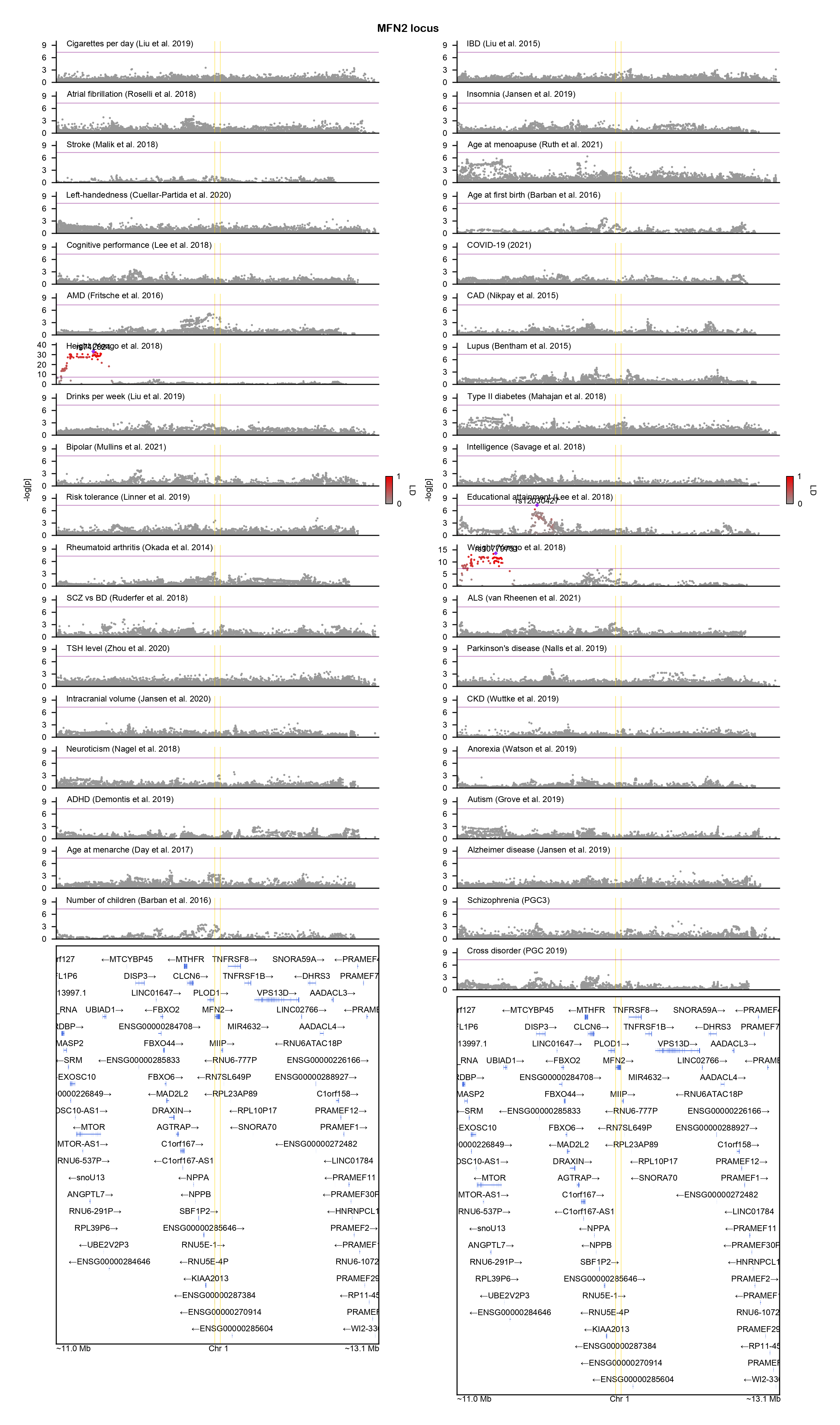

### NAA15-locuszoom.png

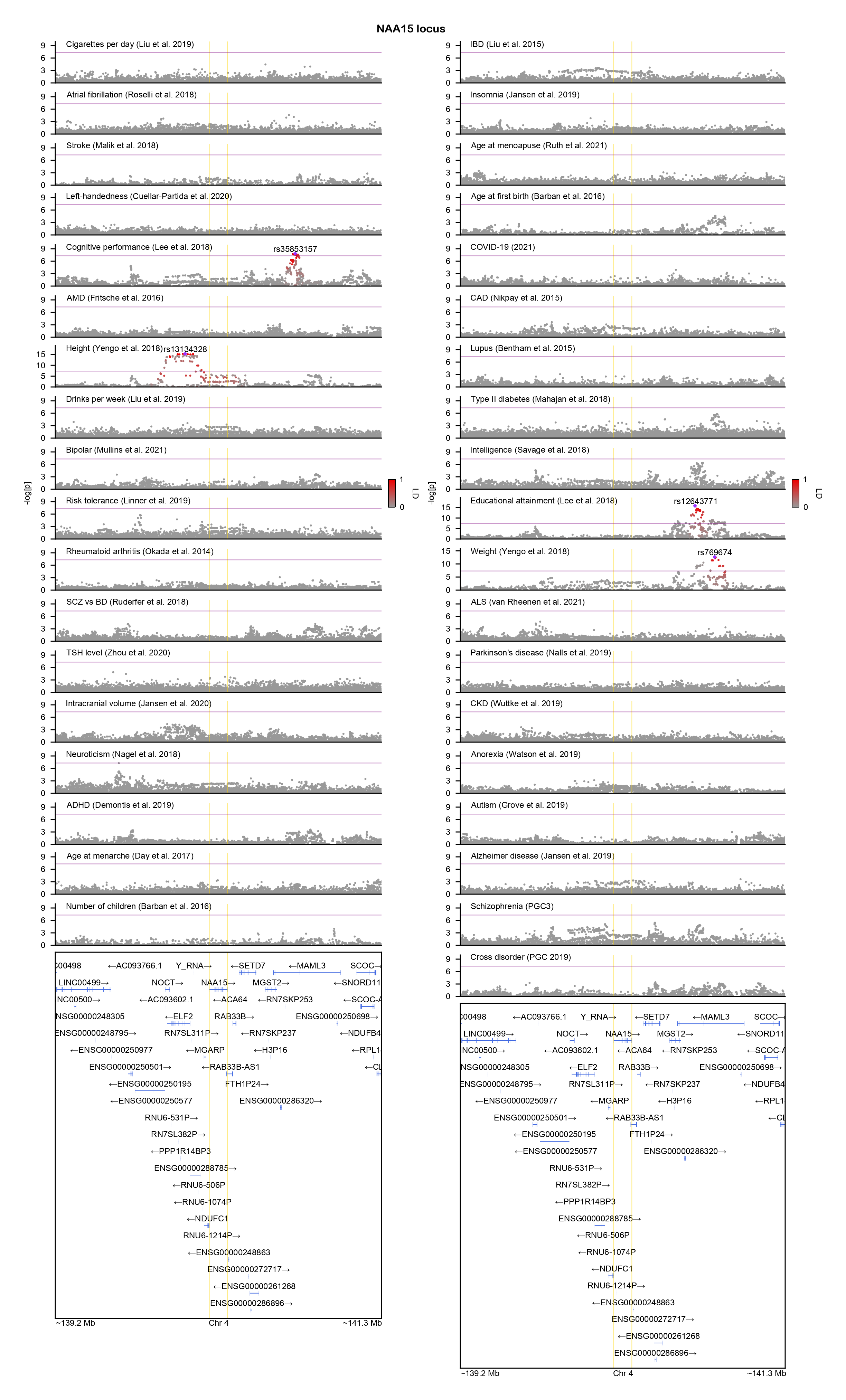

### NEXMIF-locuszoom.png

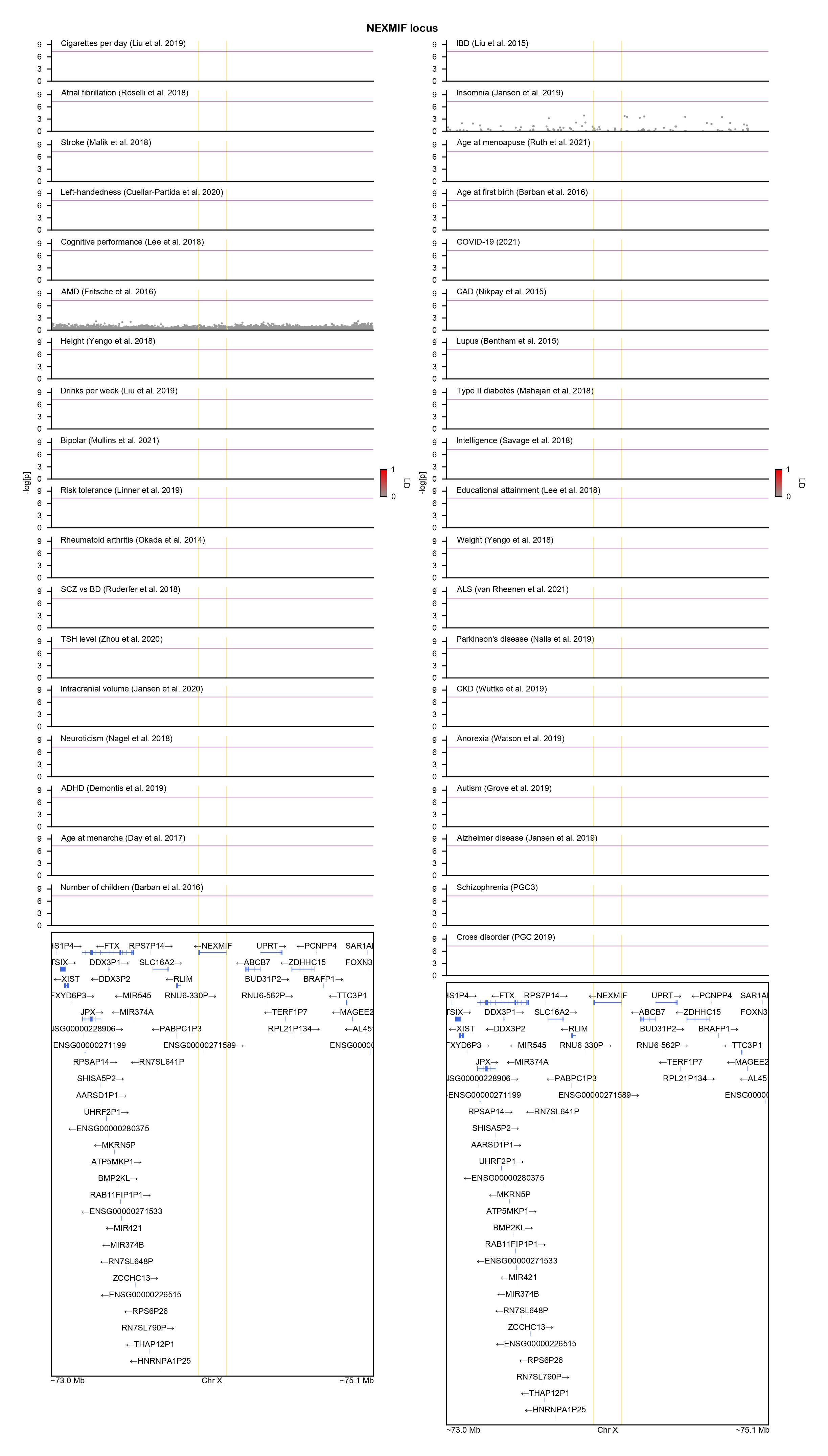

### NFIX-locuszoom.png

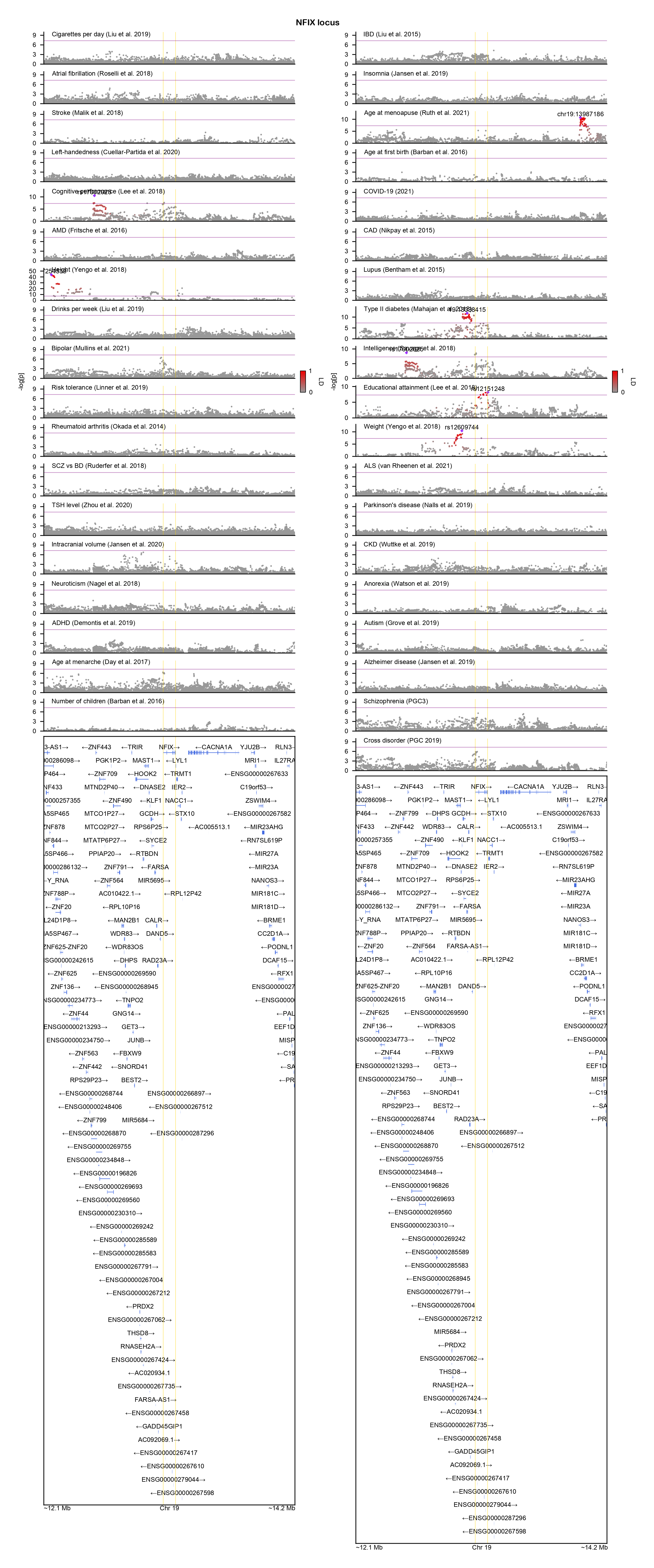

### NR2F1-locuszoom.png

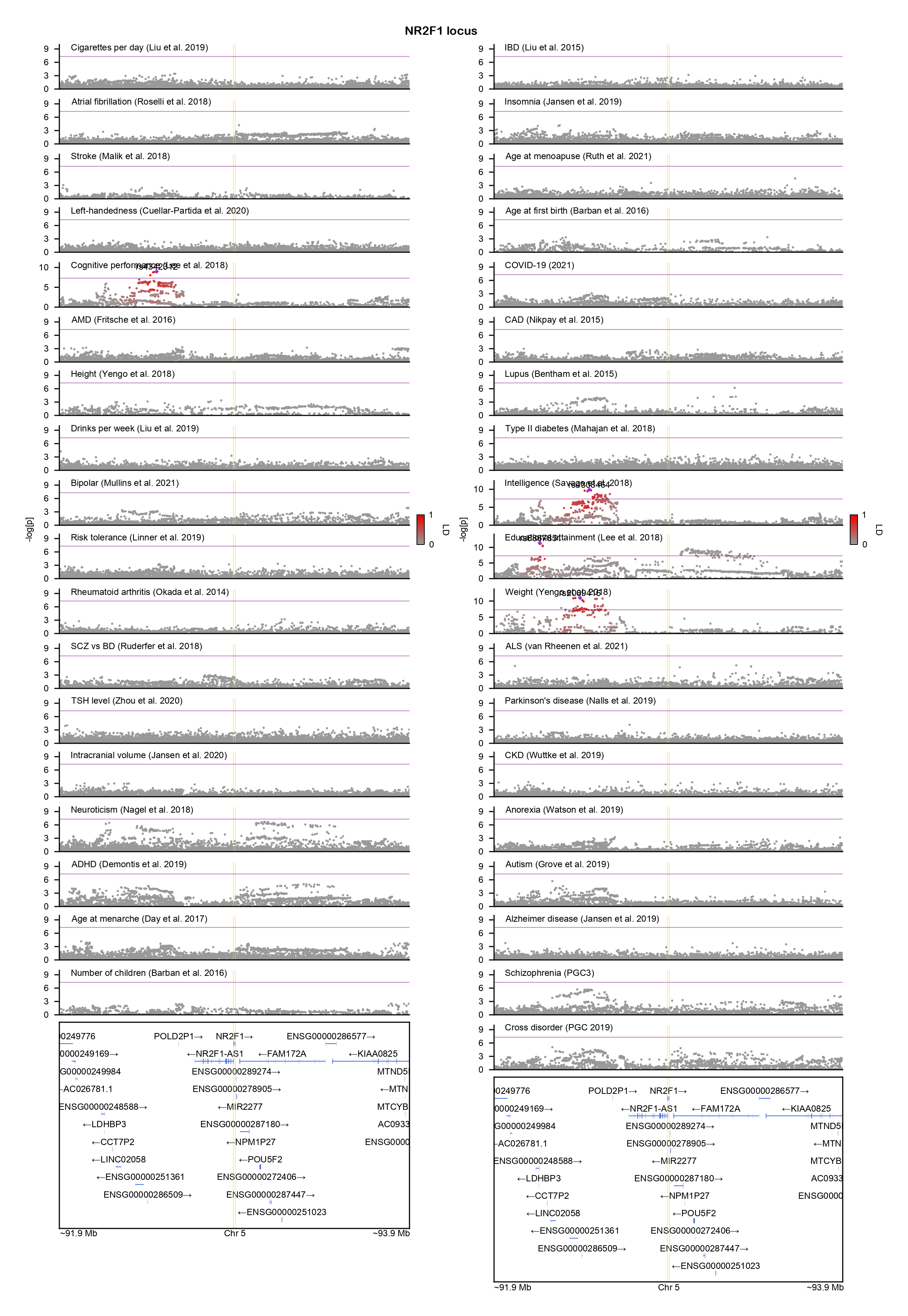

### NR3C2-locuszoom.png

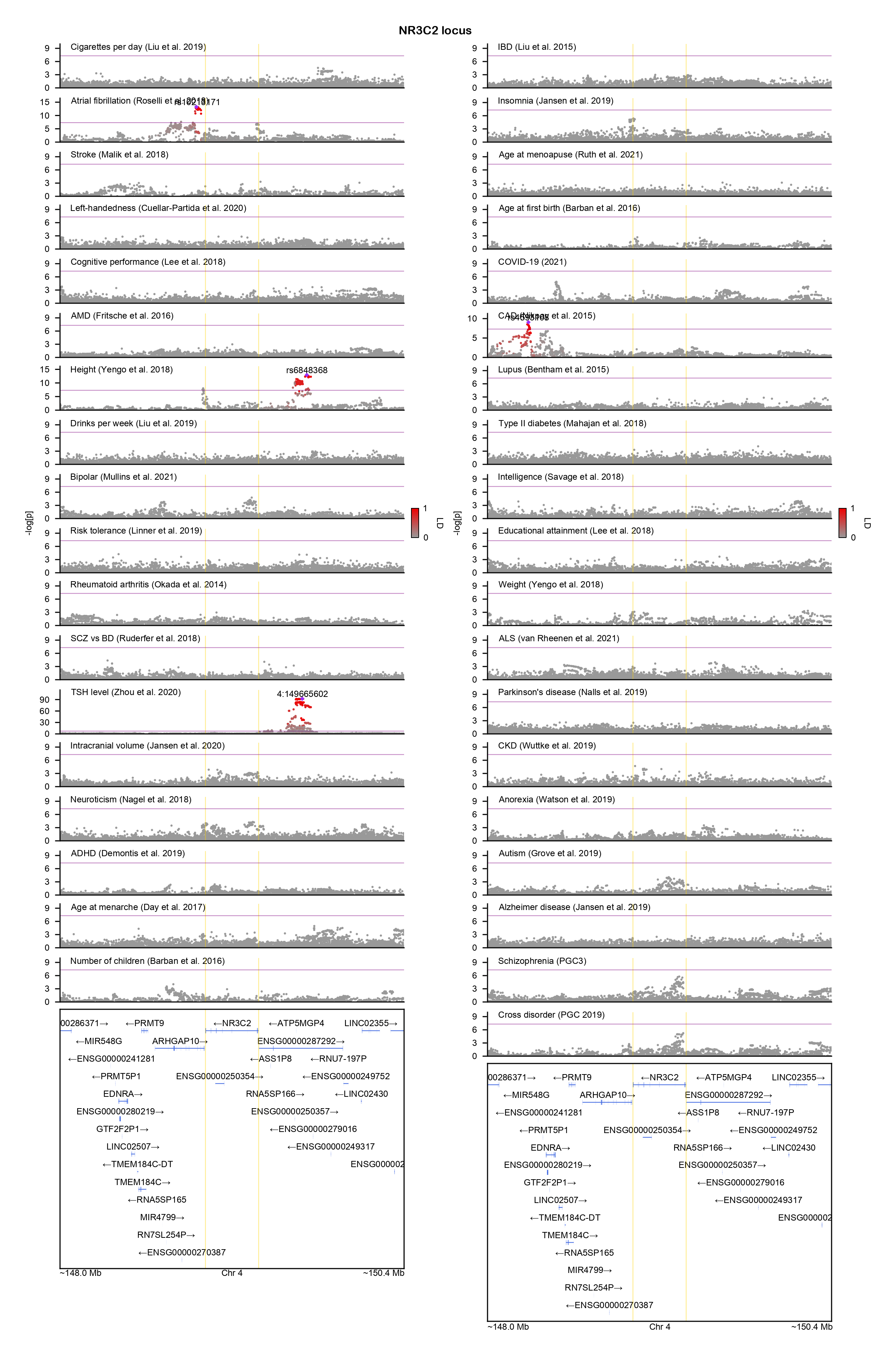

### ODC1-locuszoom.png

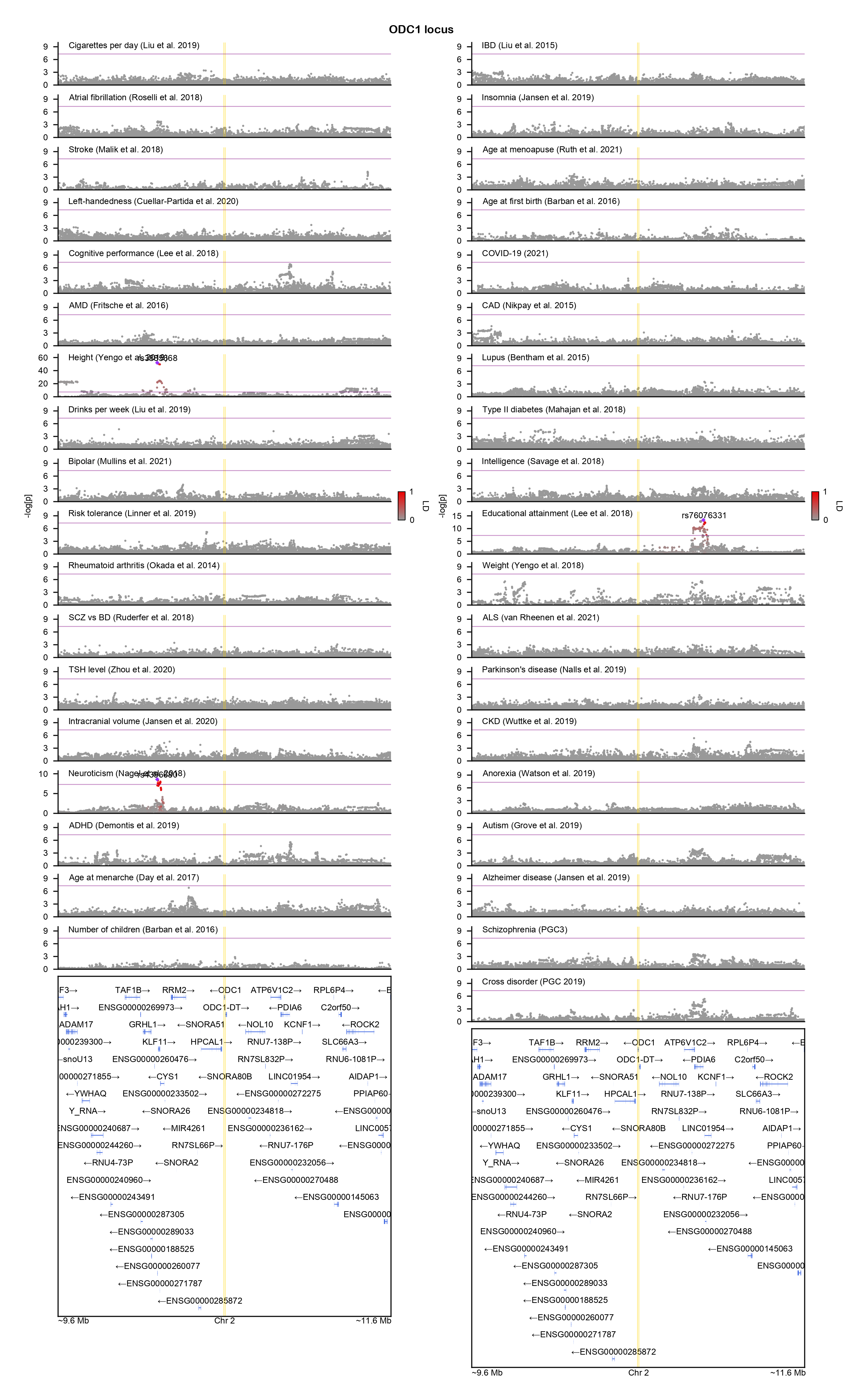

### PAX5-locuszoom.png

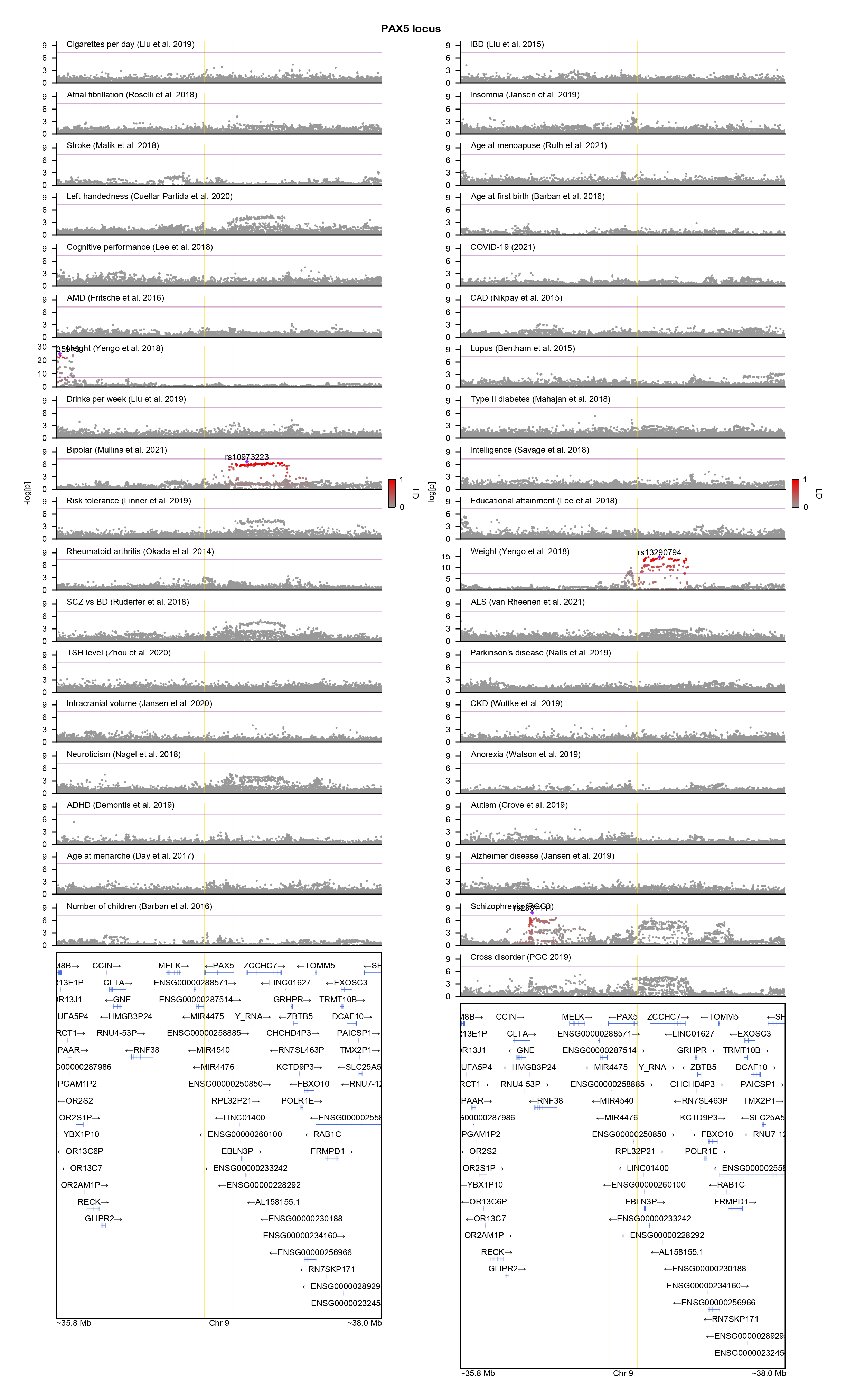

### PHF2-locuszoom.png

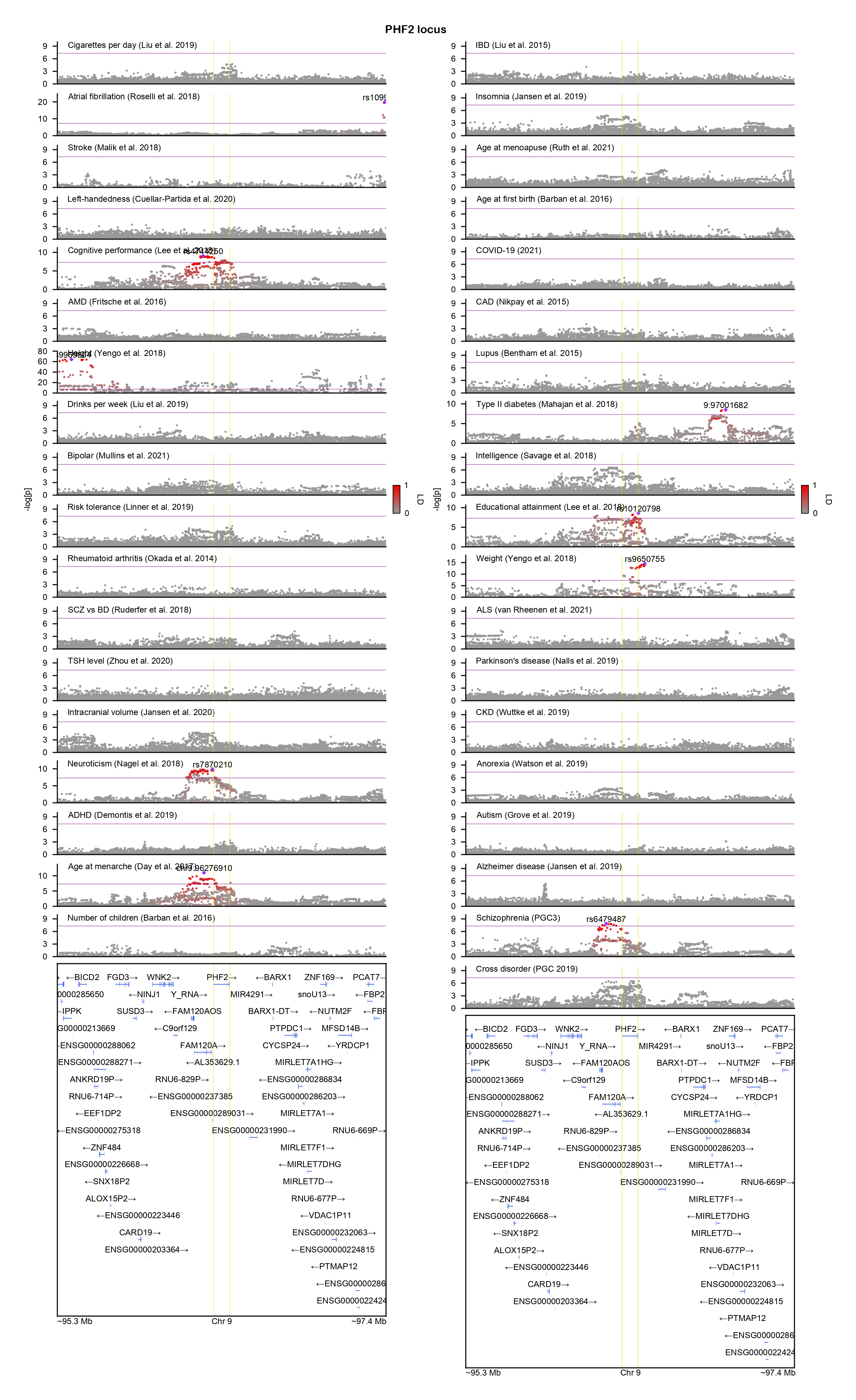

### PHF6-locuszoom.png

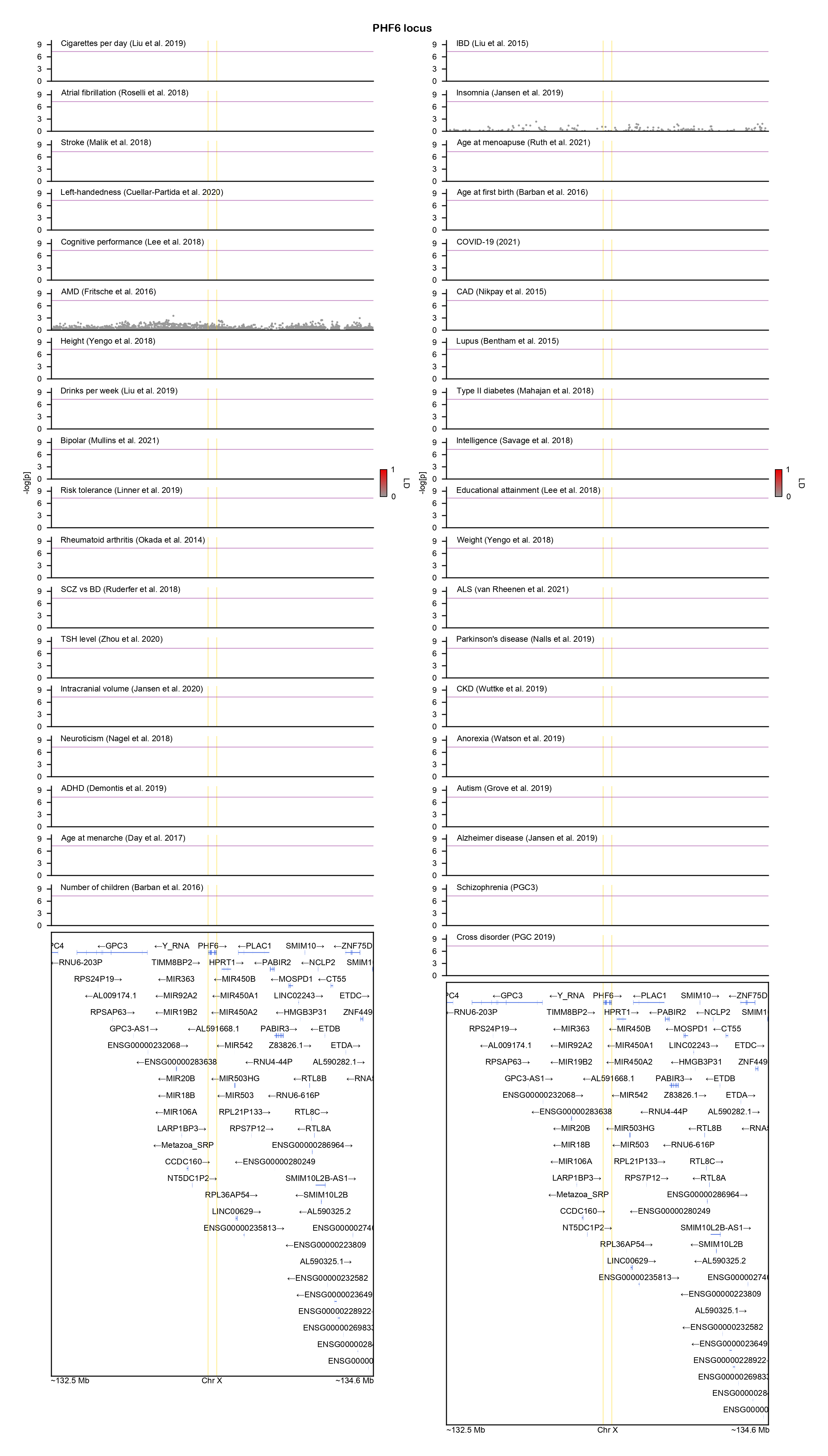

### POGZ-locuszoom.png

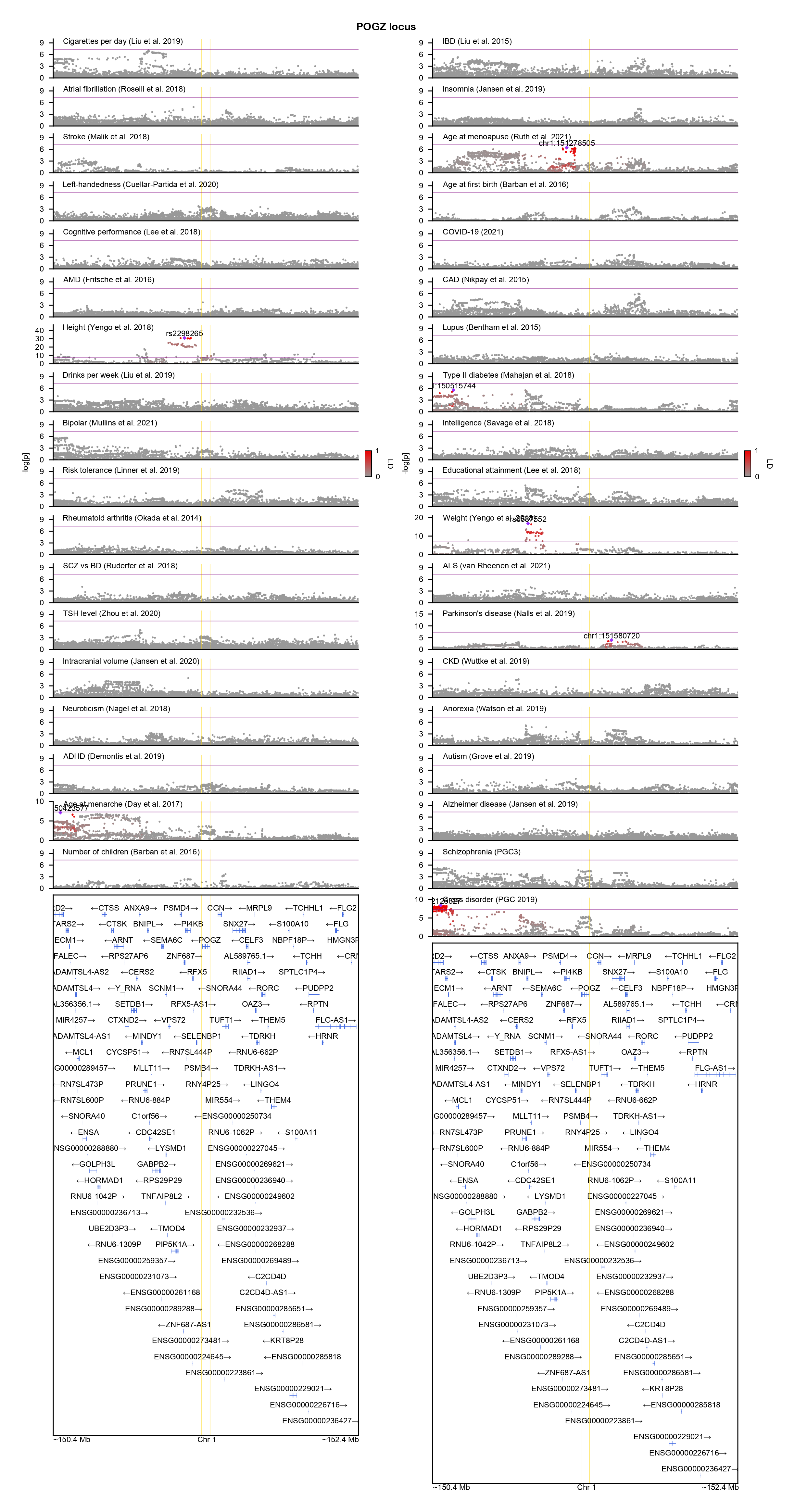

### PPP2R5D-locuszoom.png

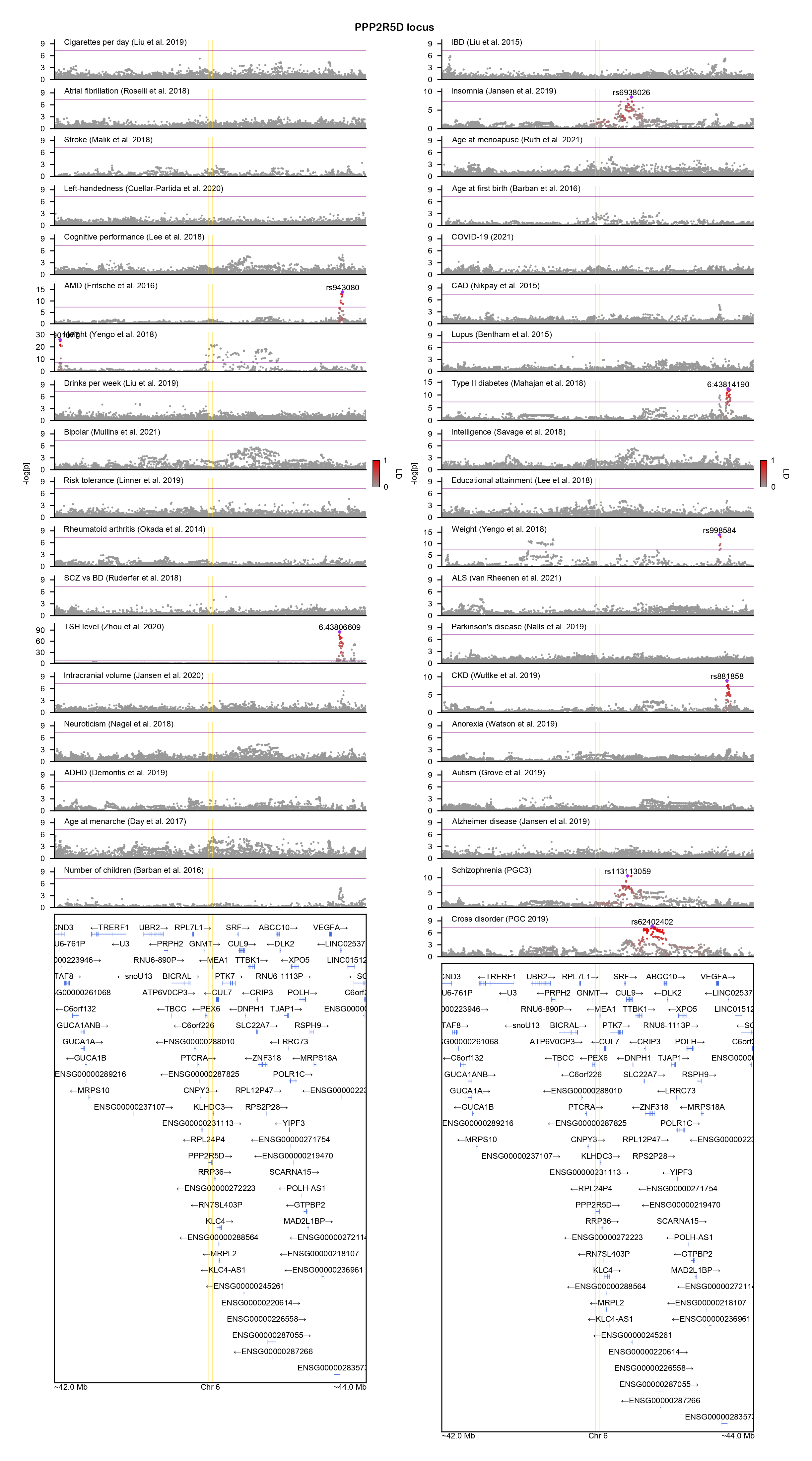

### PPP5C-locuszoom.png

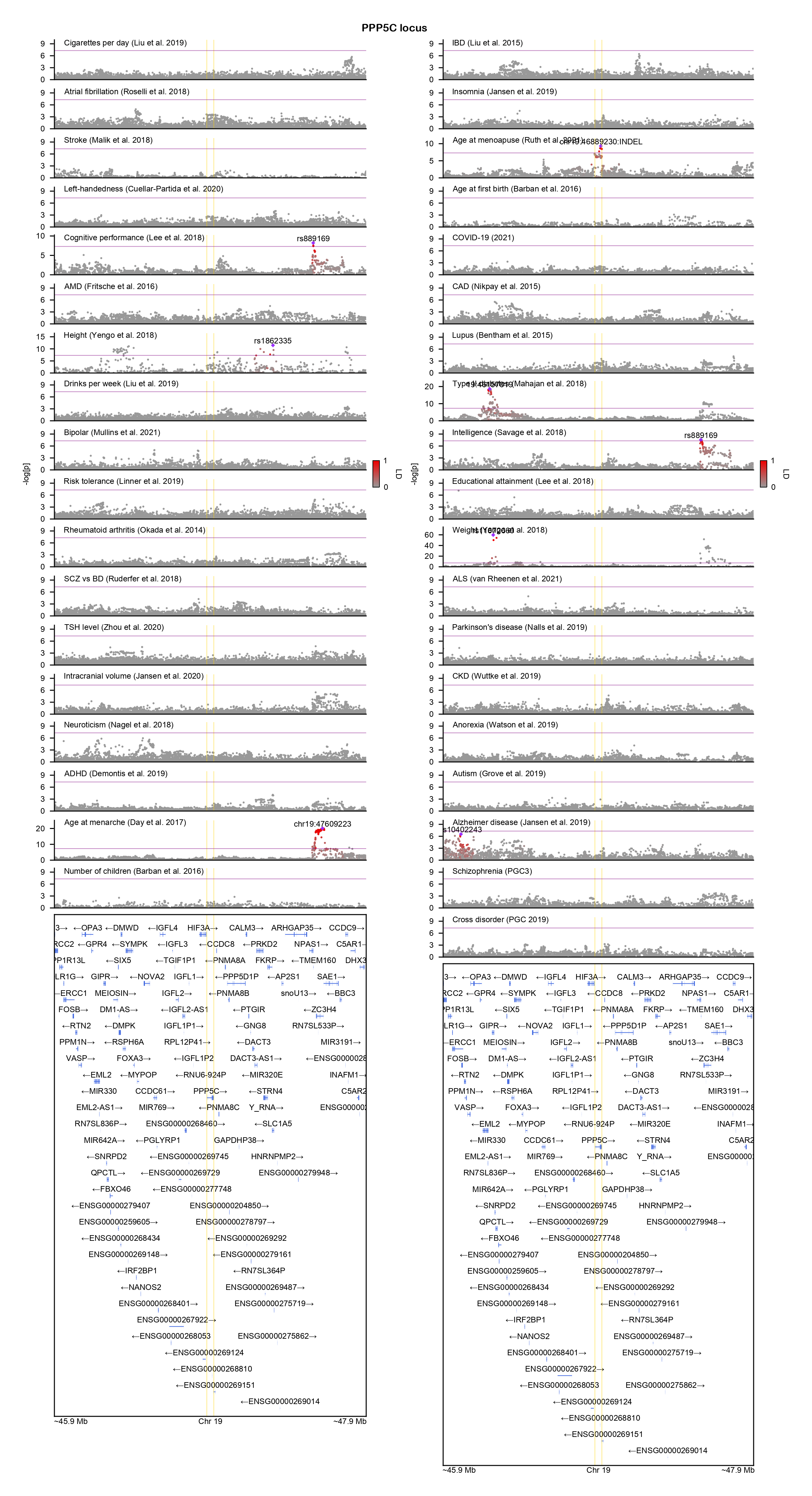

### PTEN-locuszoom.png

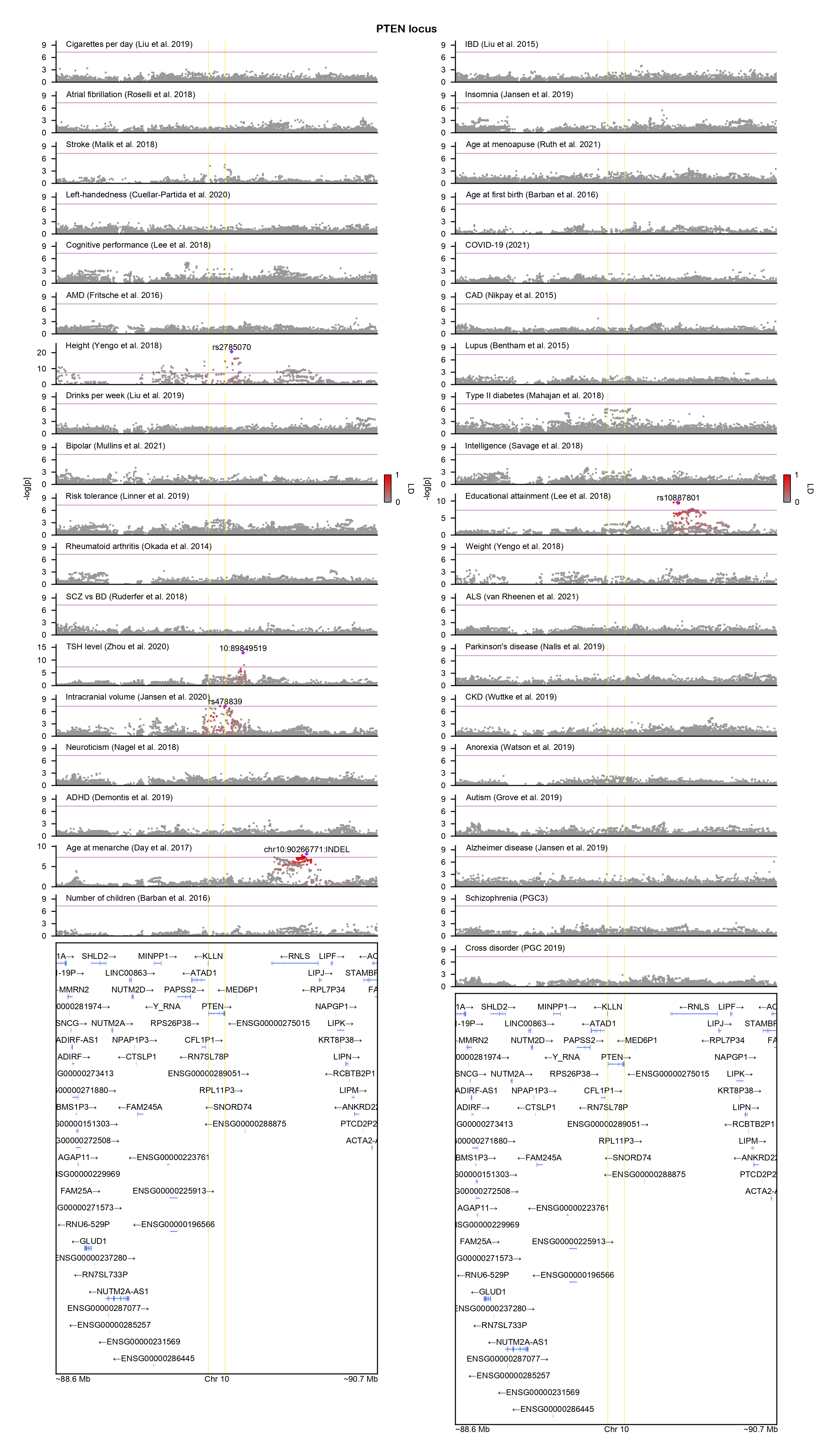

### PUF60-locuszoom.png

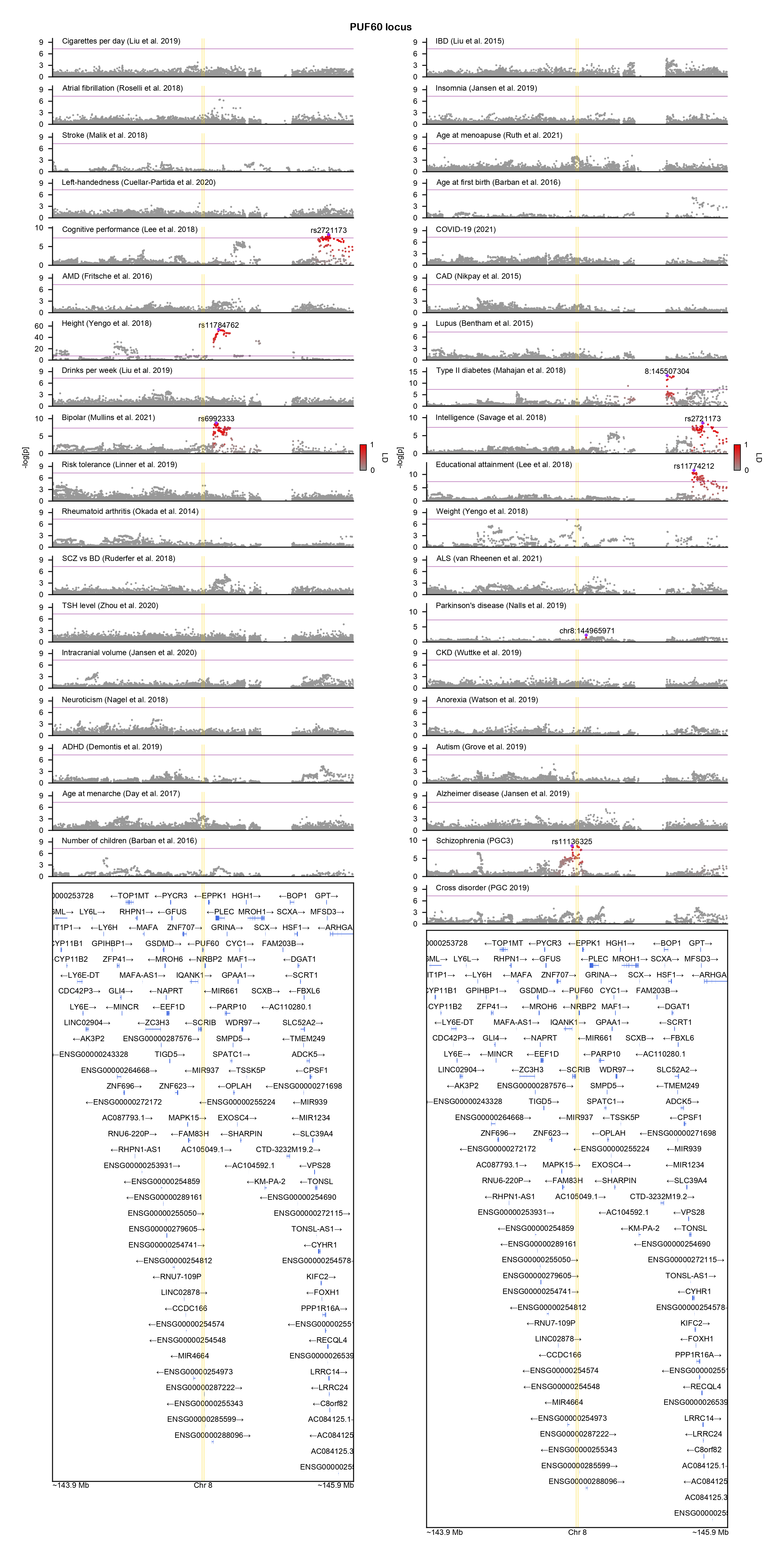

### QRICH1-locuszoom.png

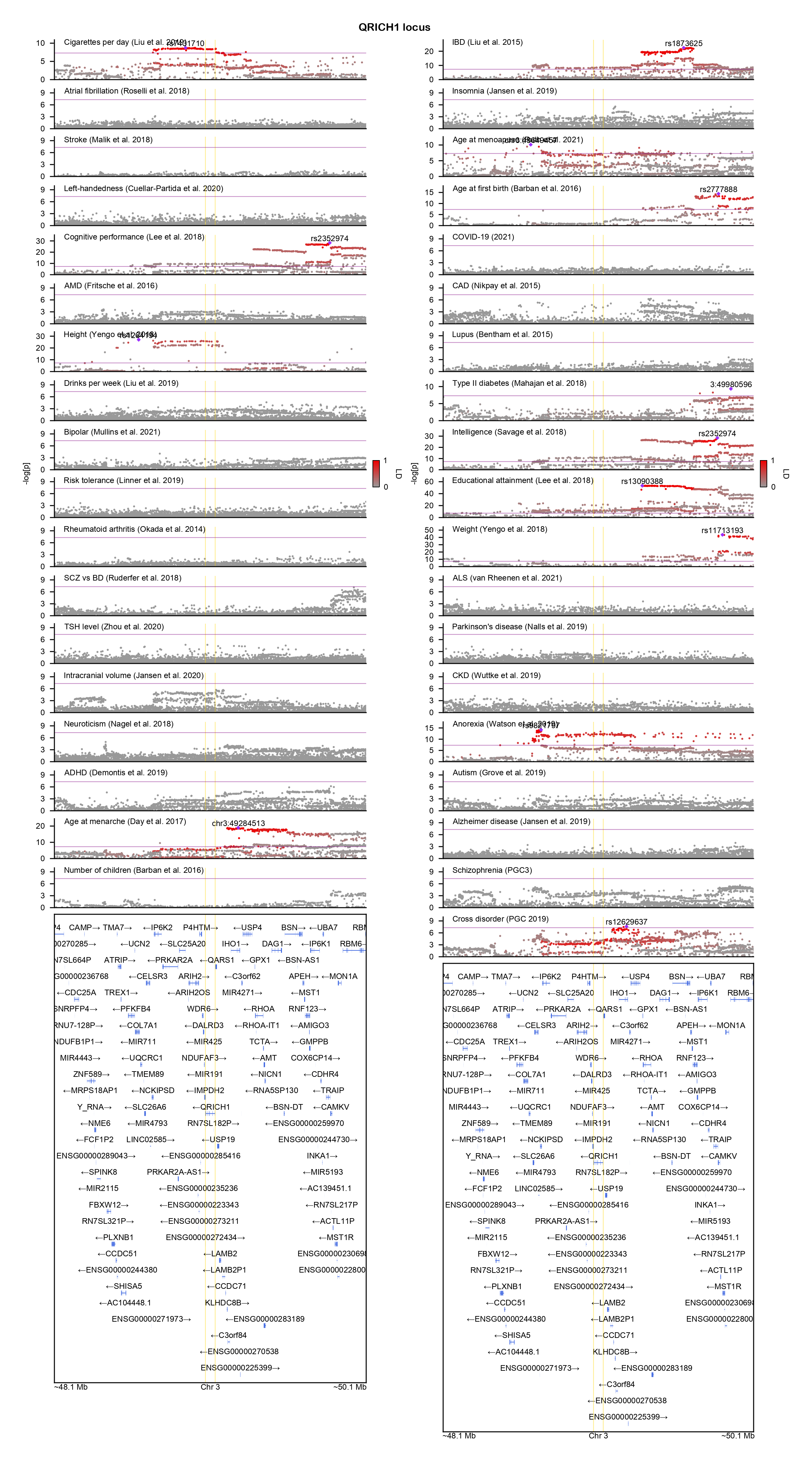

### RAB14-locuszoom.png

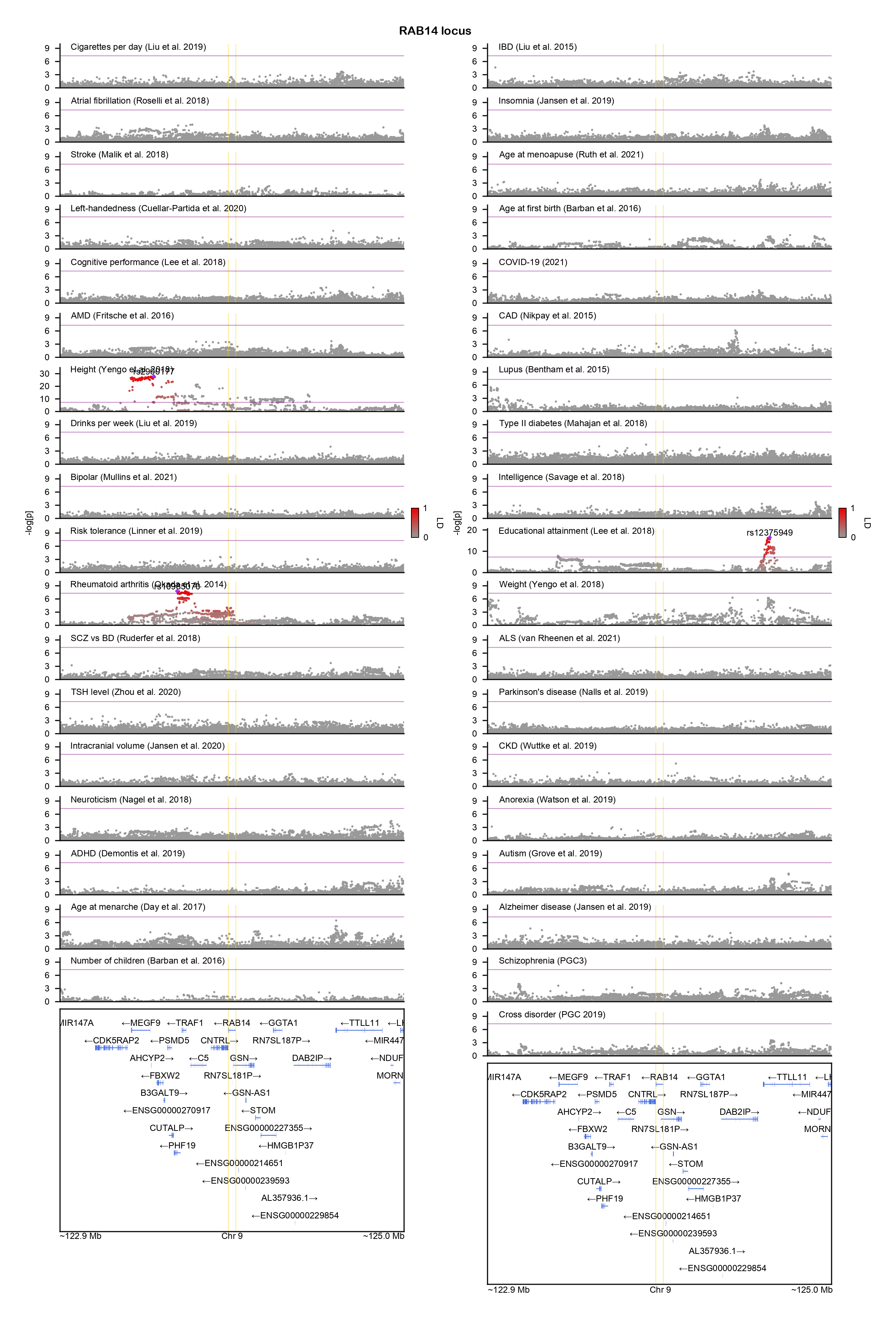

### RAI1-locuszoom.png

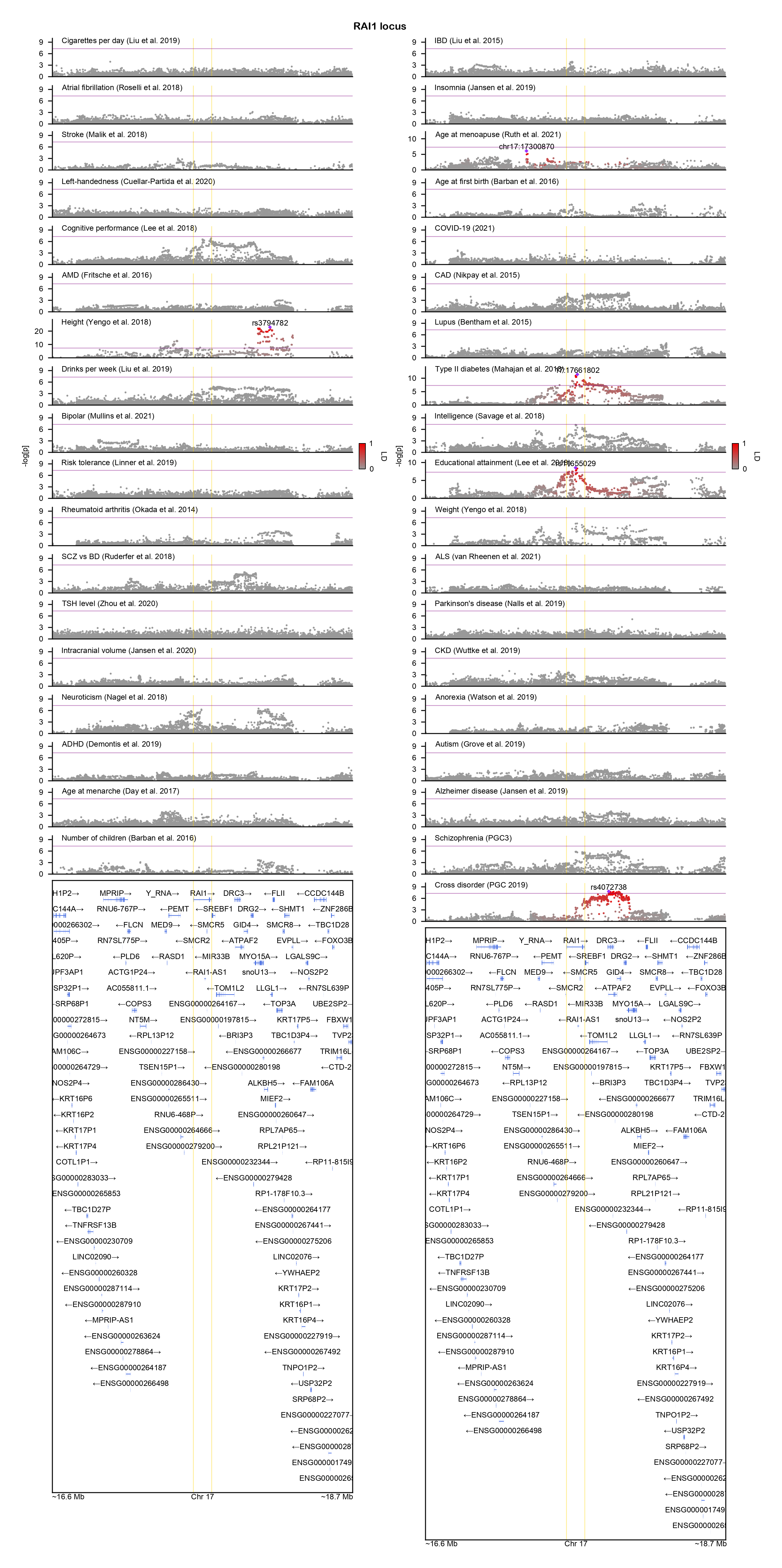

### RHOBTB2-locuszoom.png

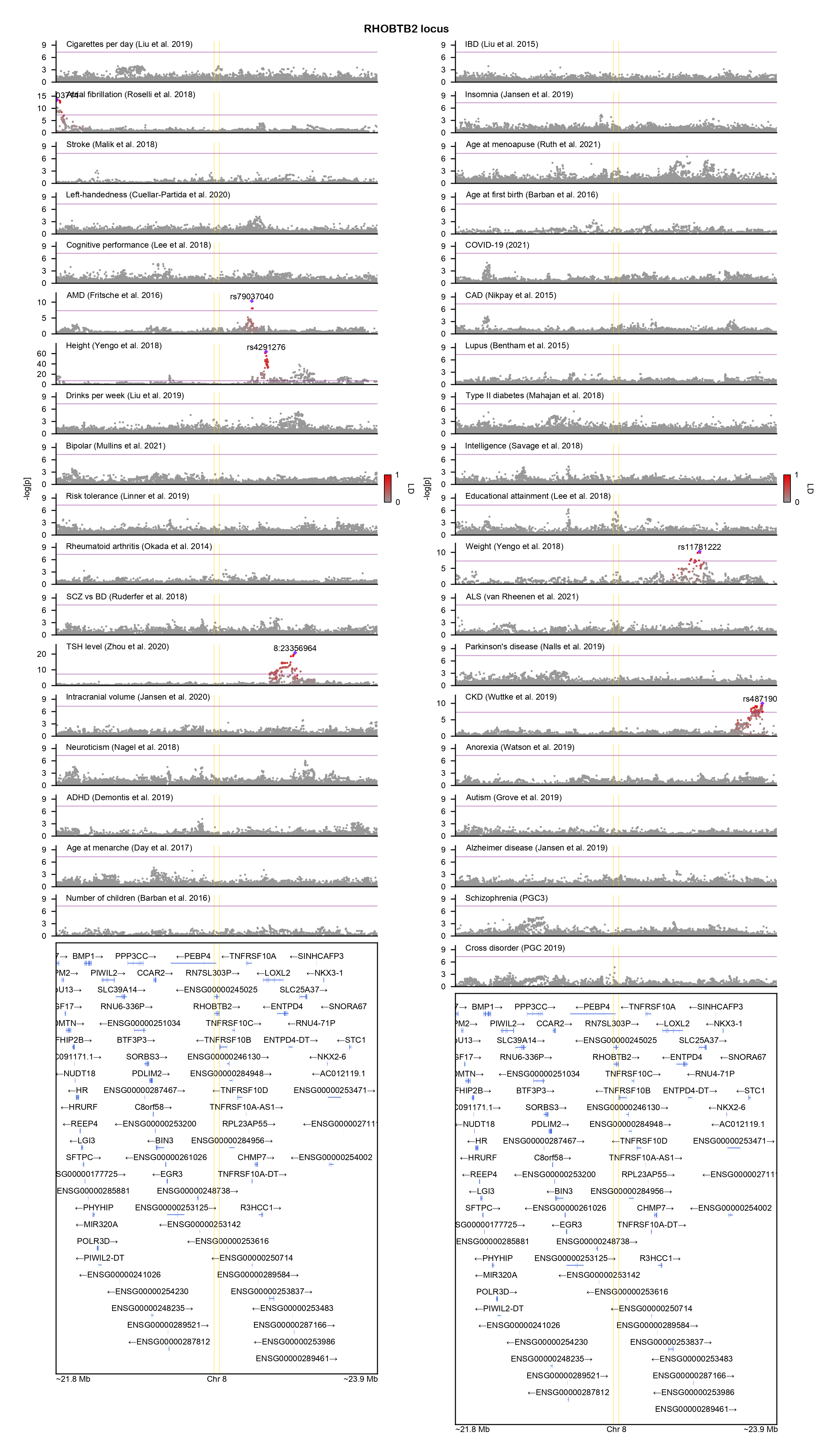

### RIT1-locuszoom.png

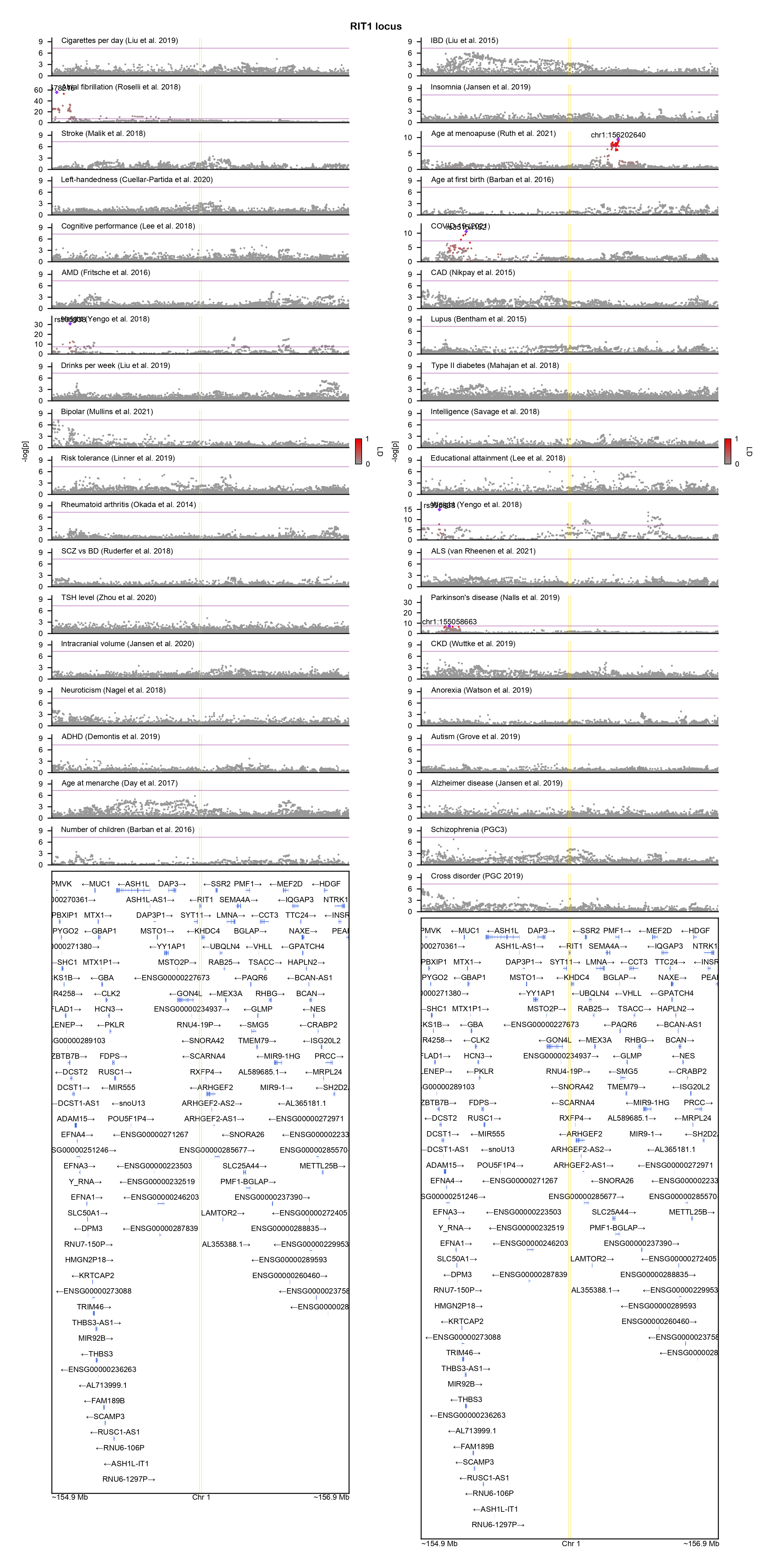

### RORB-locuszoom.png

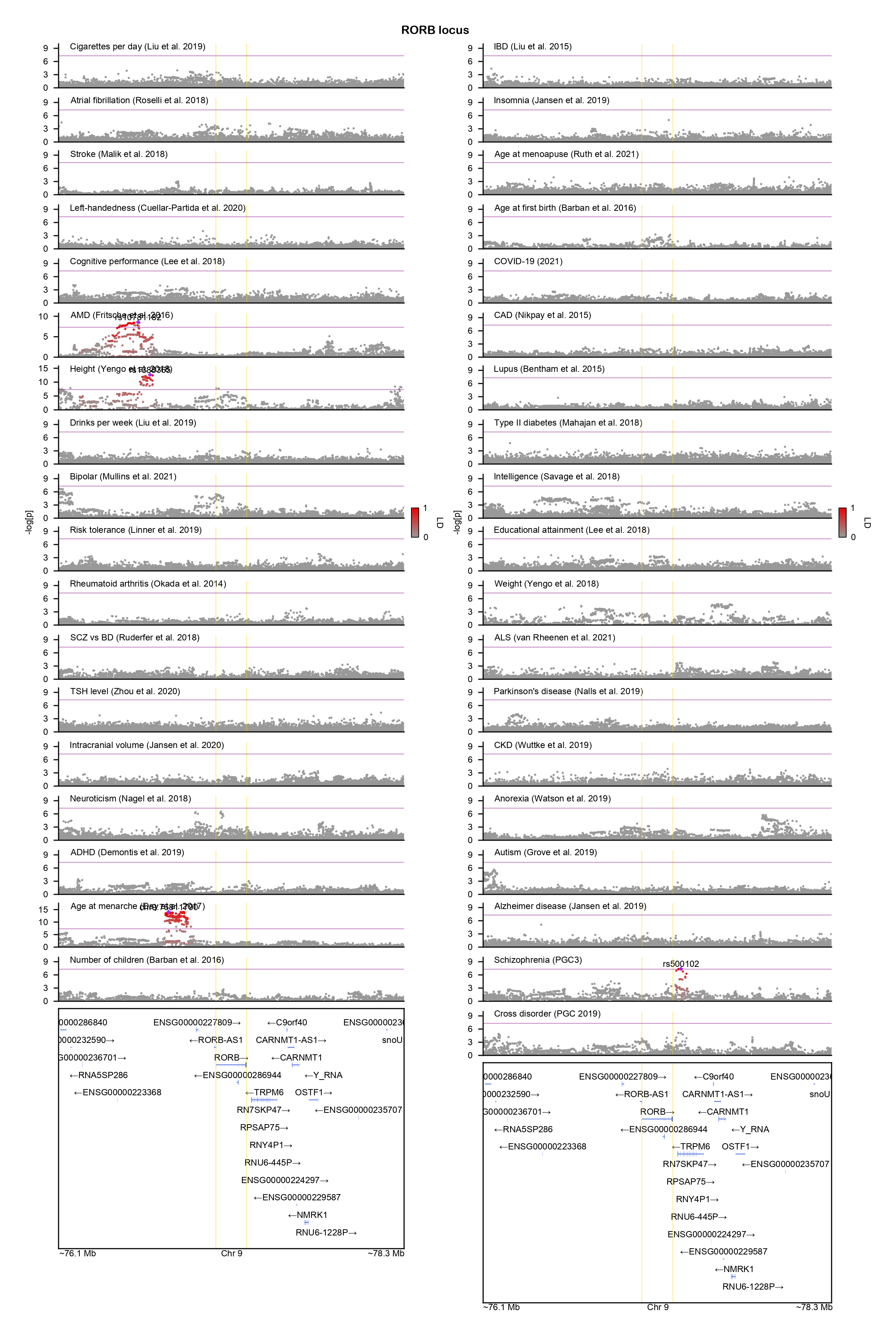

### SKI-locuszoom.png

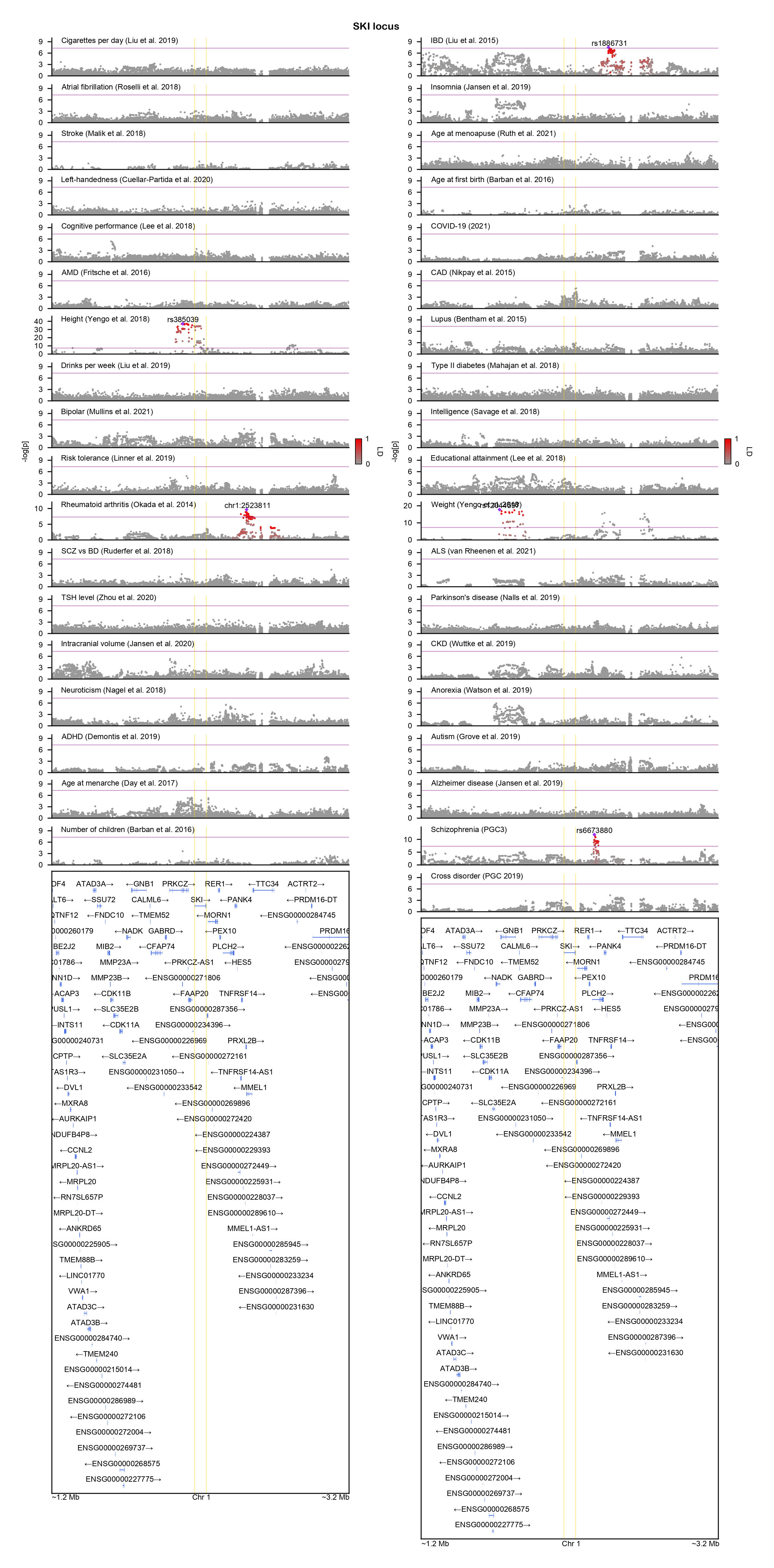

### SLC2A1-locuszoom.png

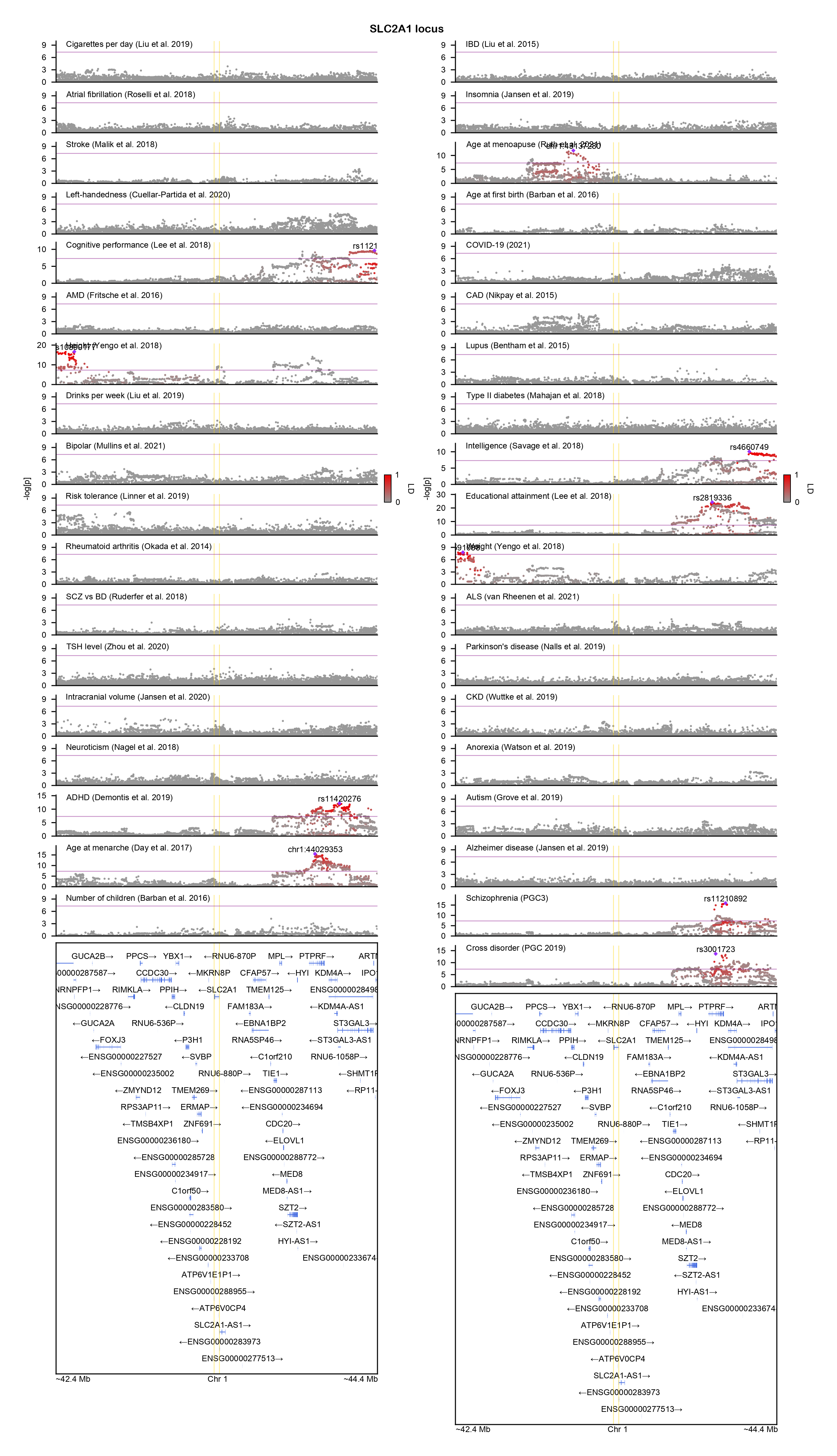

### SMARCB1-locuszoom.png

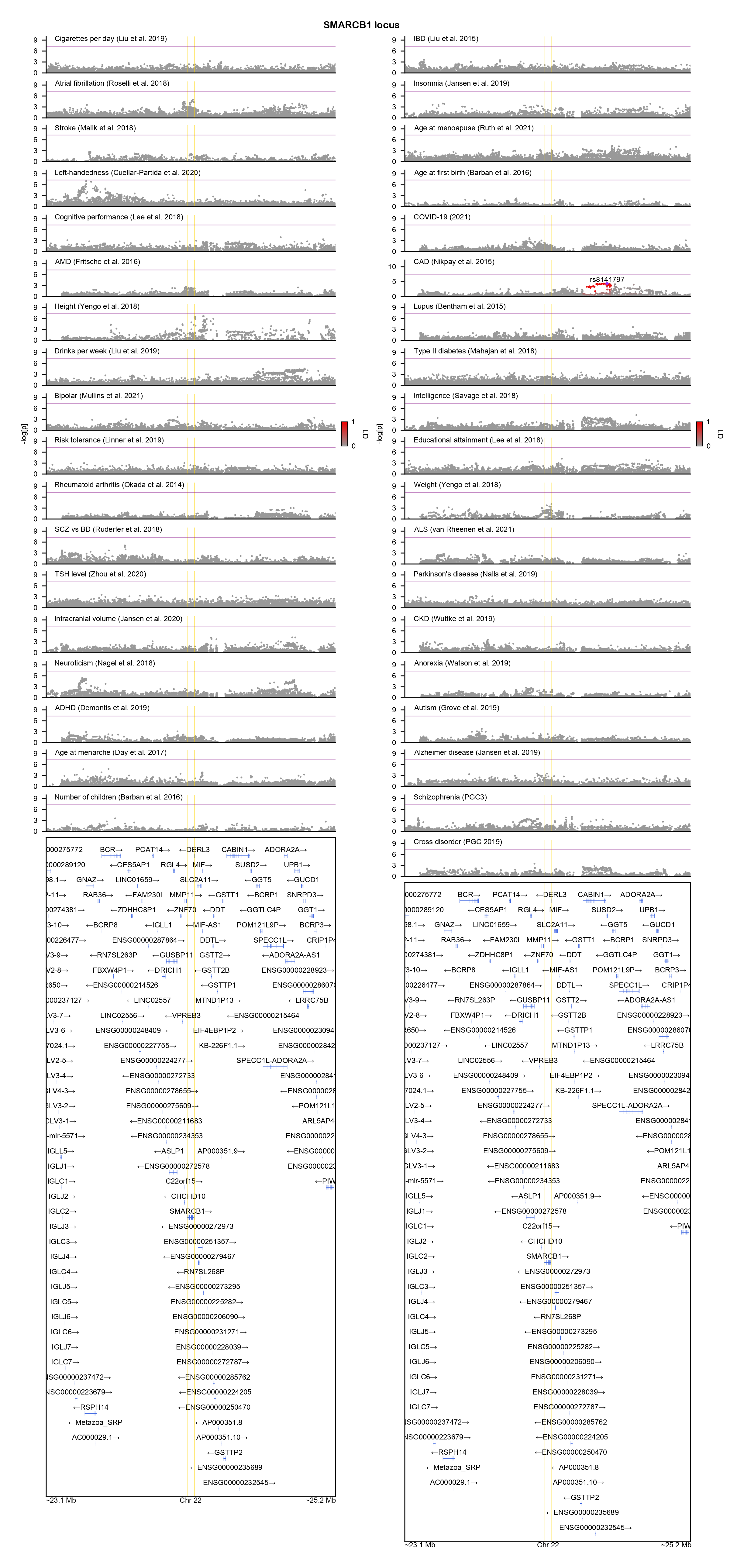

### SMARCC2-locuszoom.png

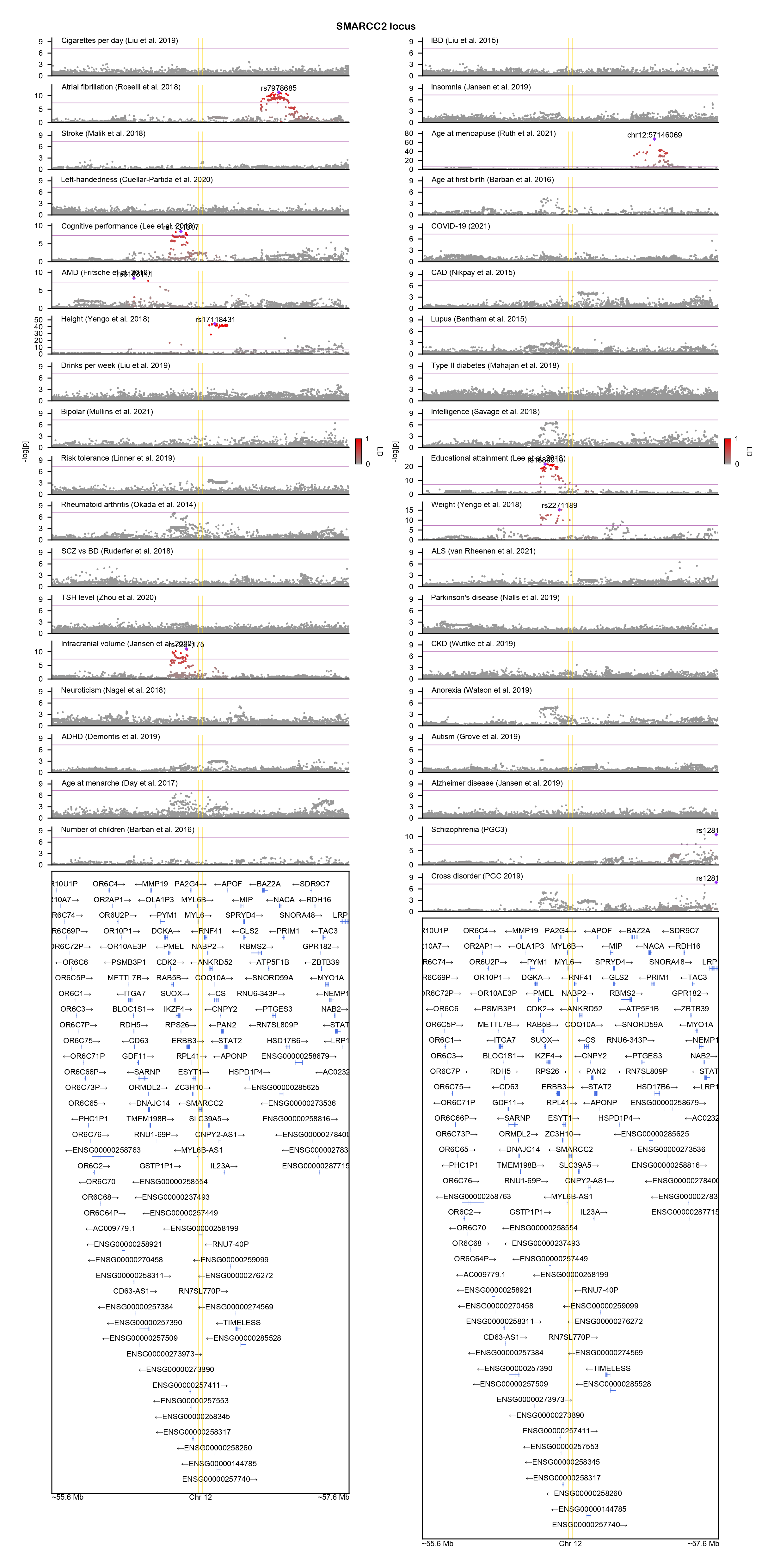

### SP4-locuszoom.png

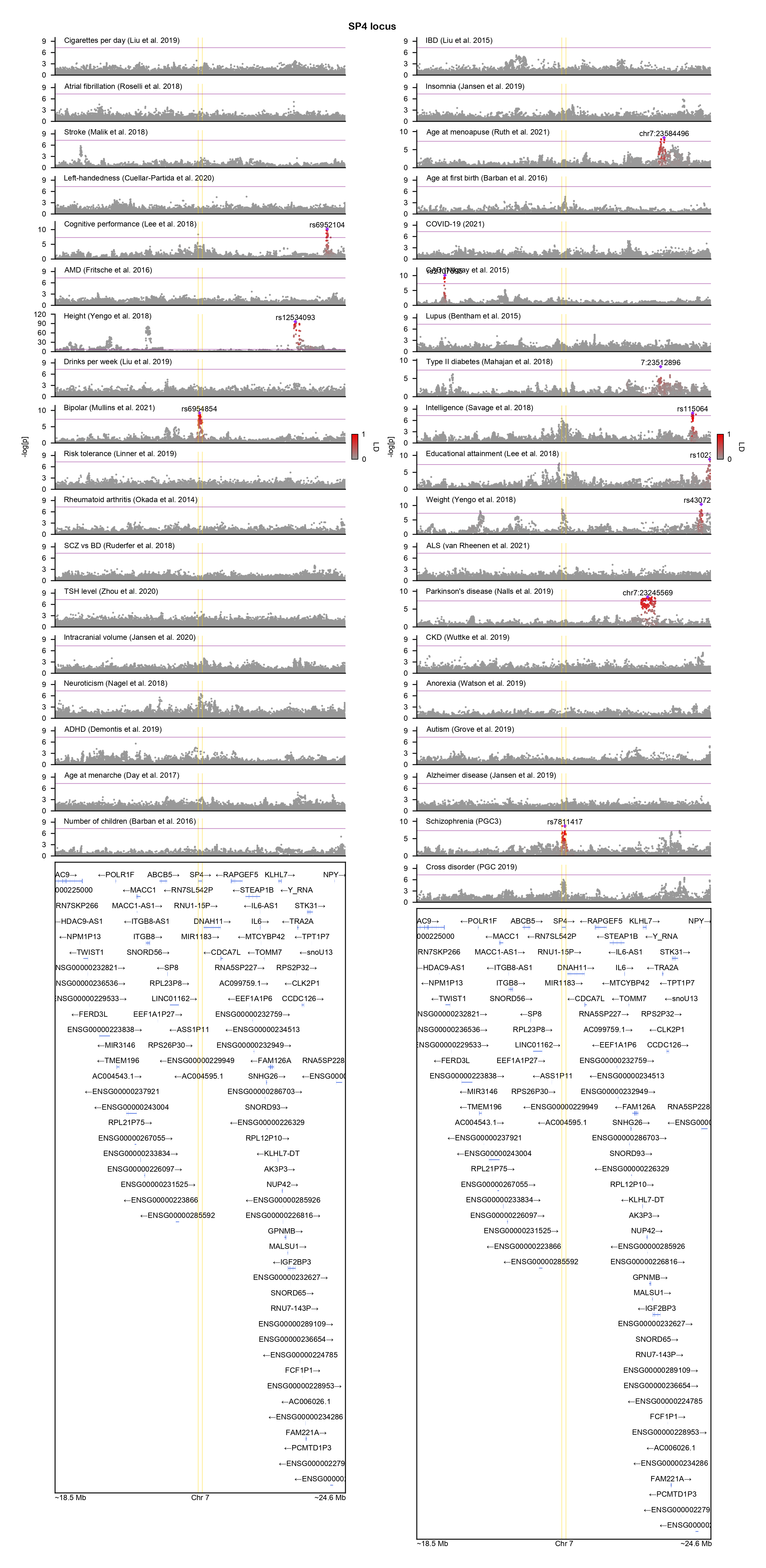

### SPTAN1-locuszoom.png

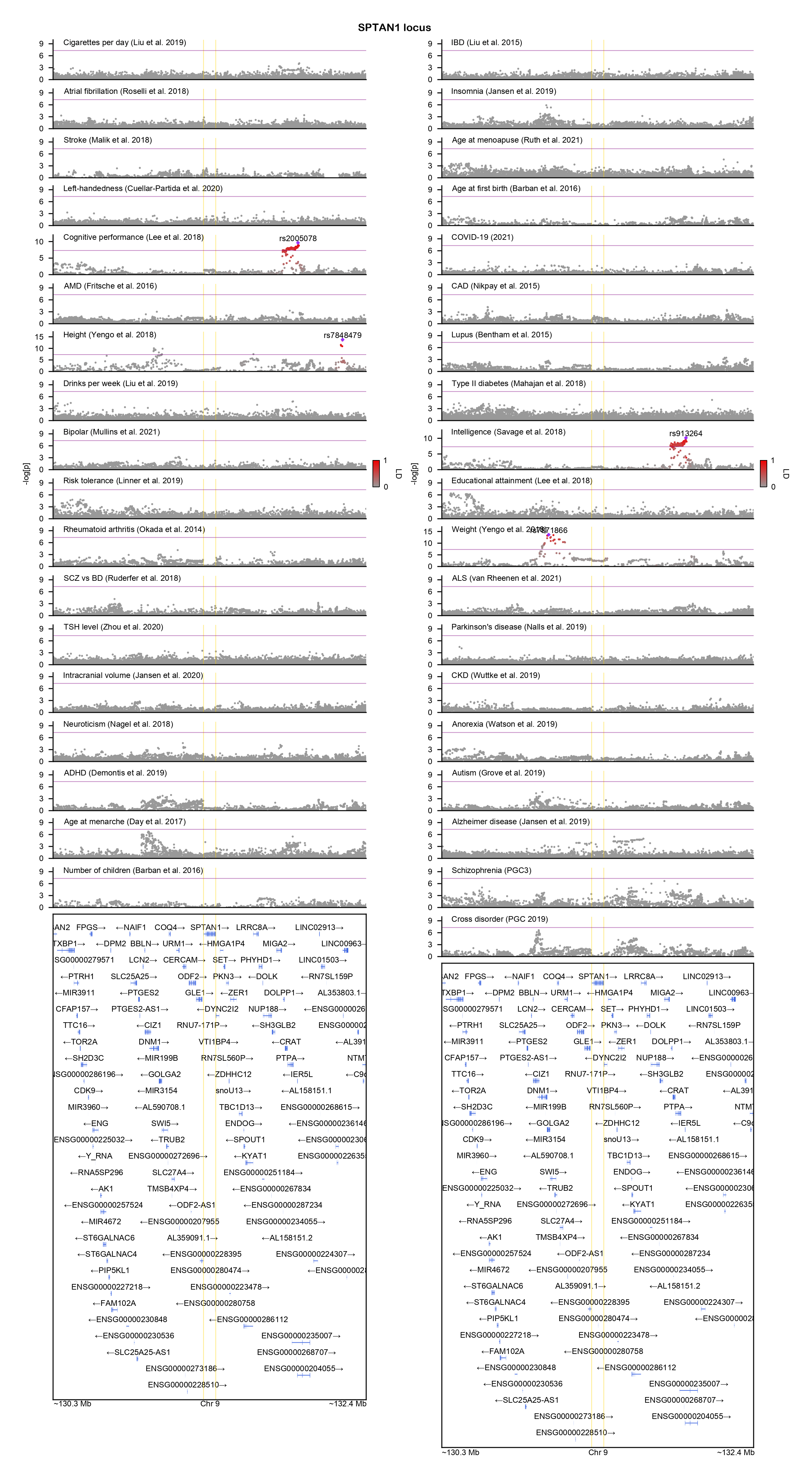

### STXBP1-locuszoom.png

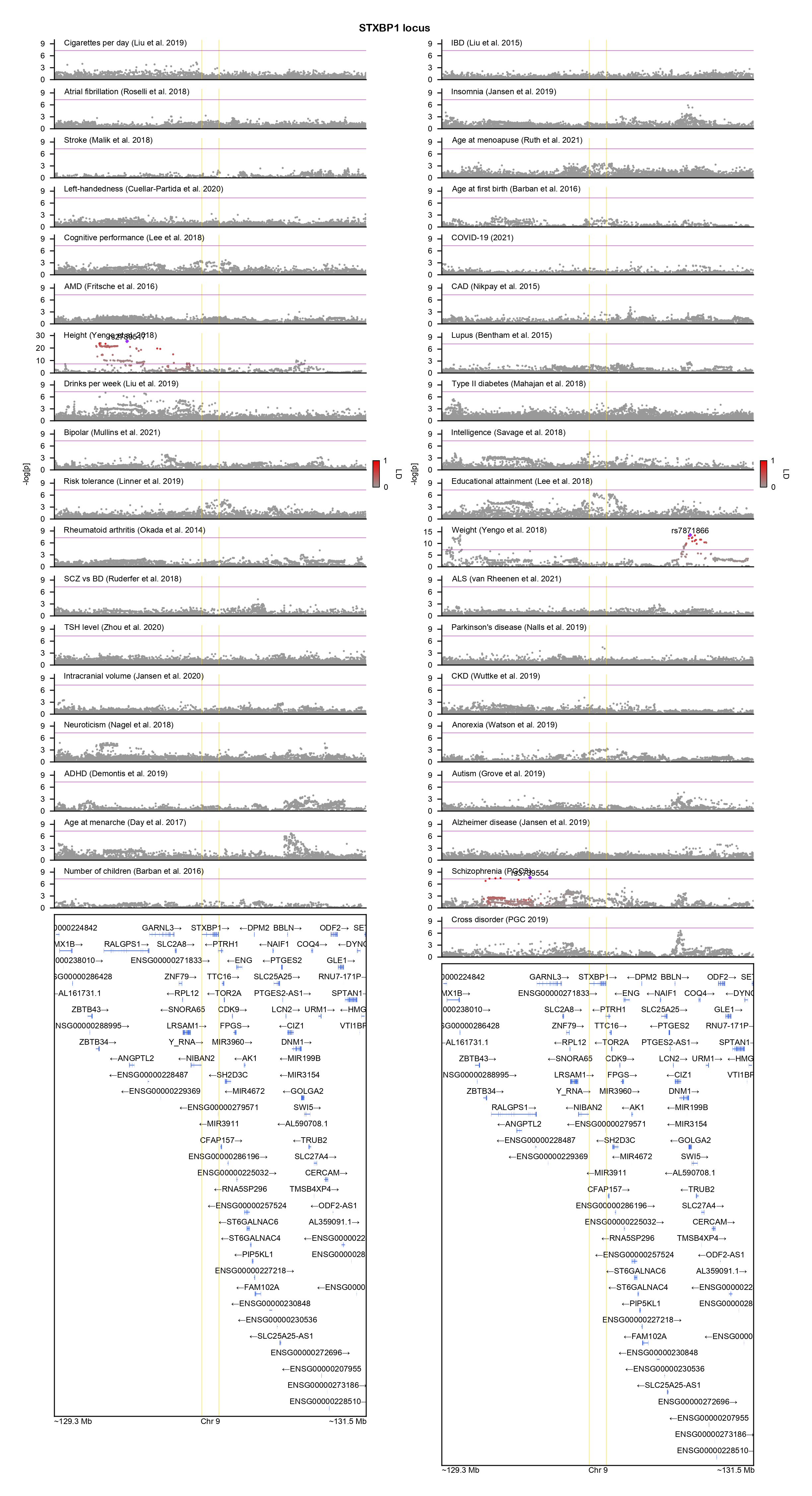

### TCF20-locuszoom.png

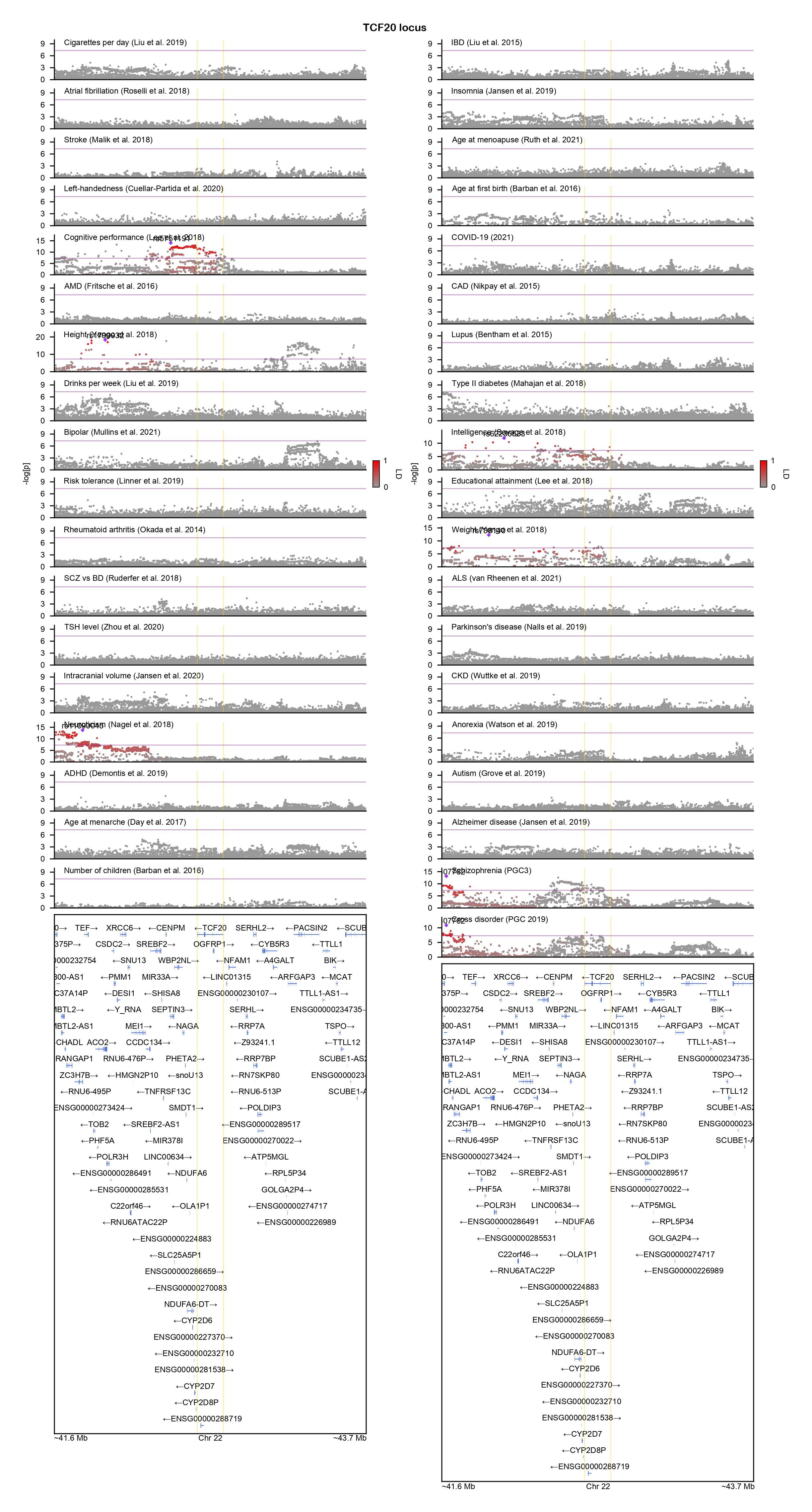

### TFE3-locuszoom.png

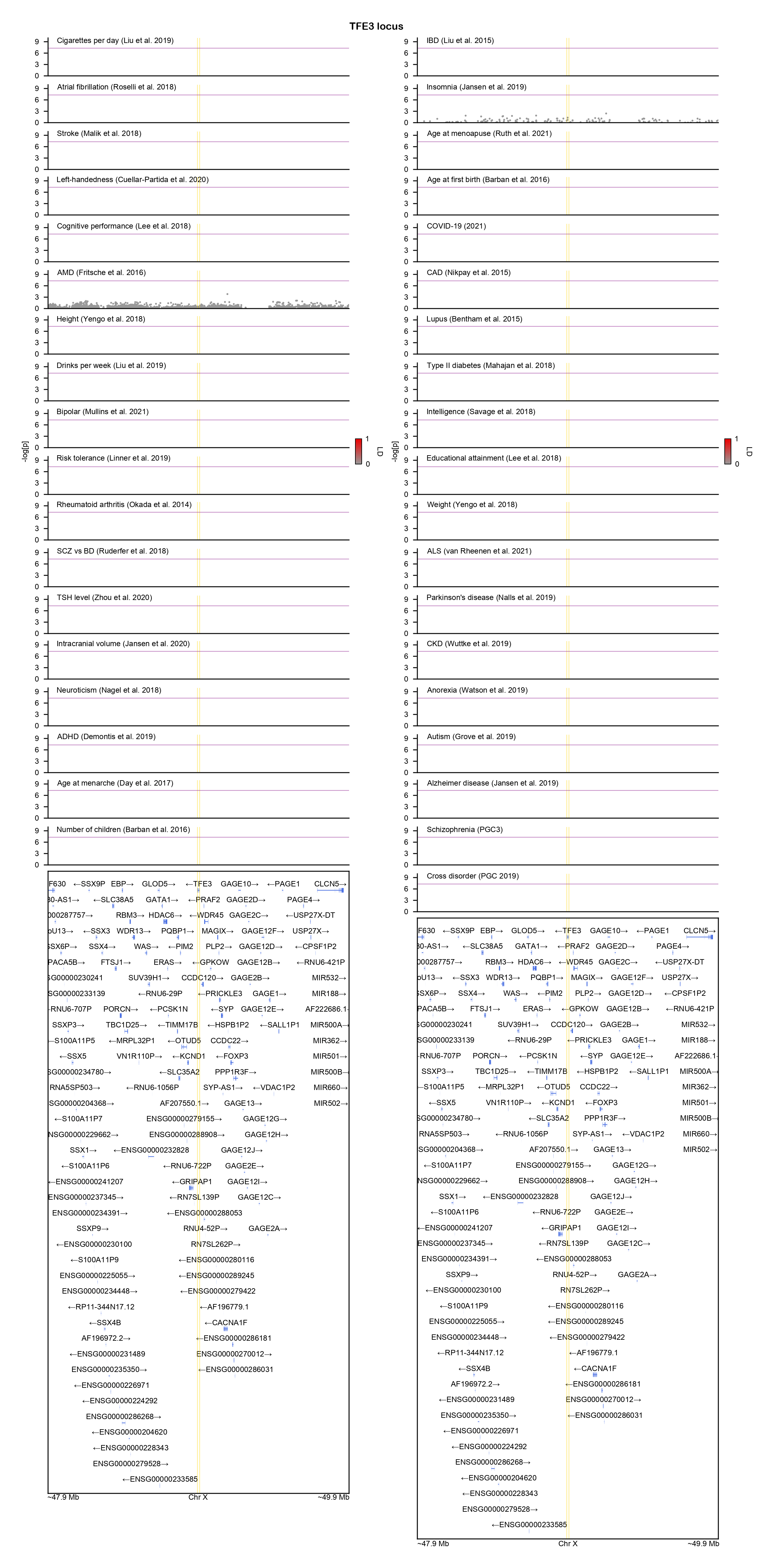

### TM9SF4-locuszoom.png

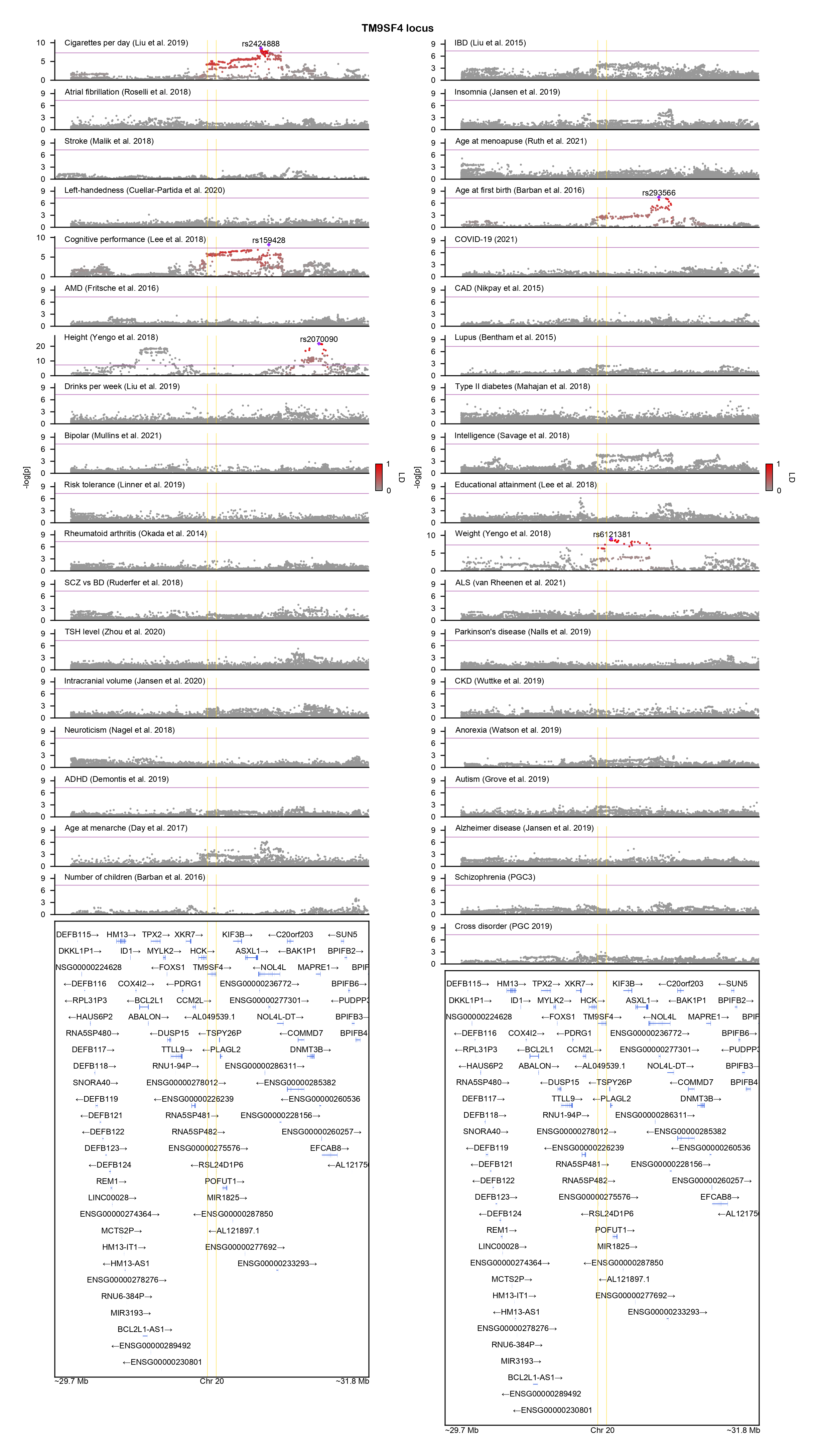

### TRIM23-locuszoom.png

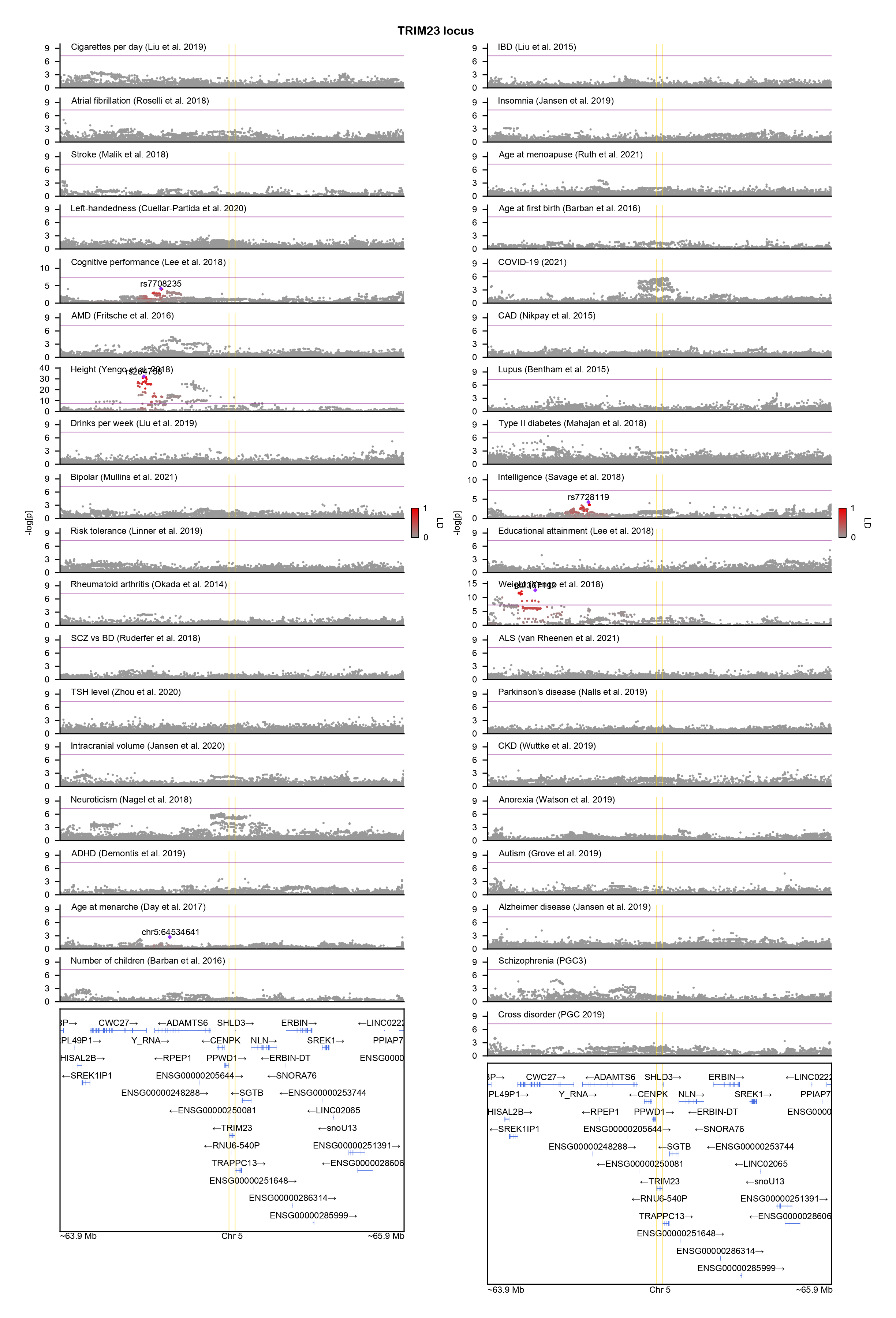

### TRIO-locuszoom.png

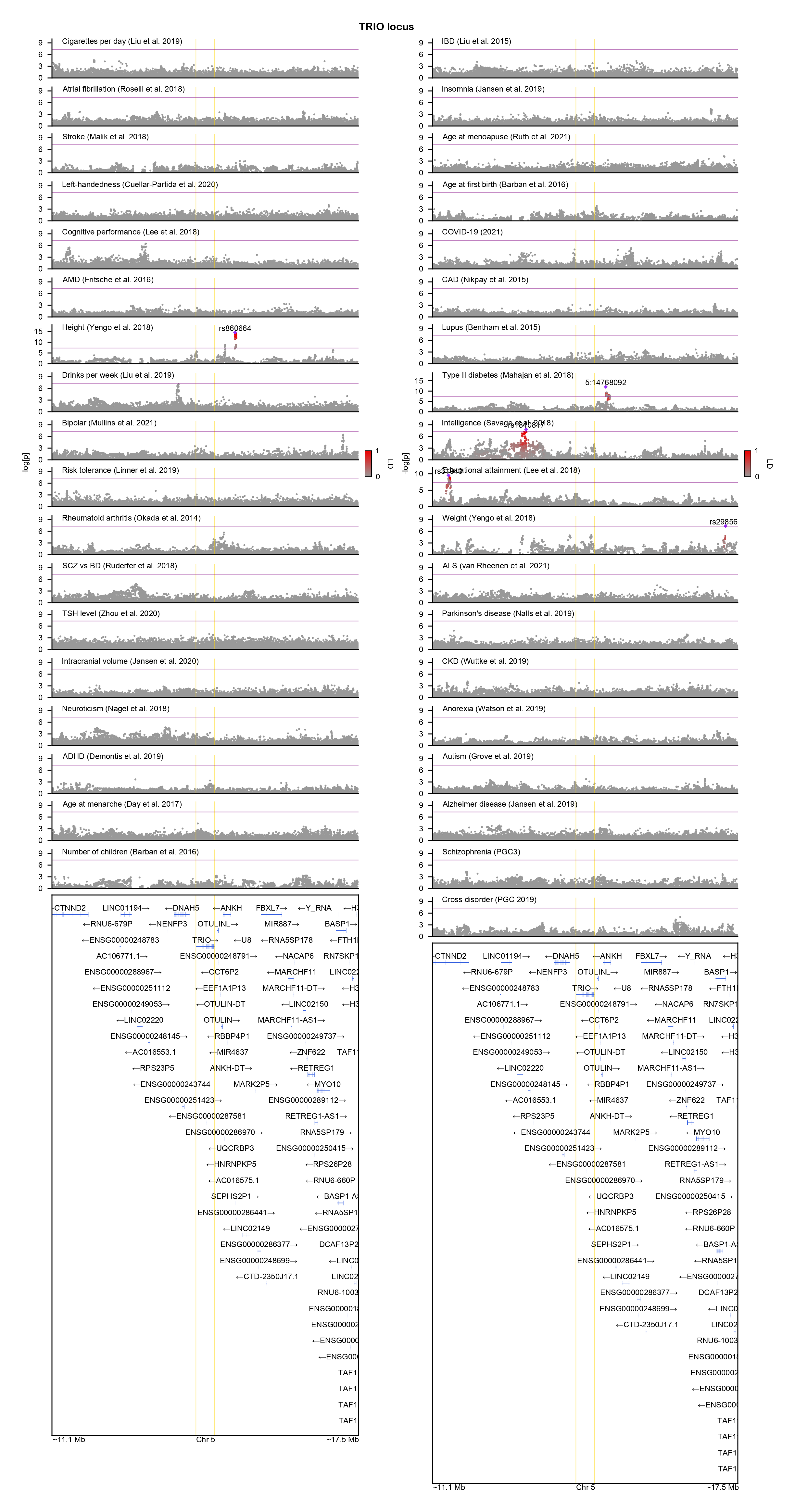

### UBTF-locuszoom.png

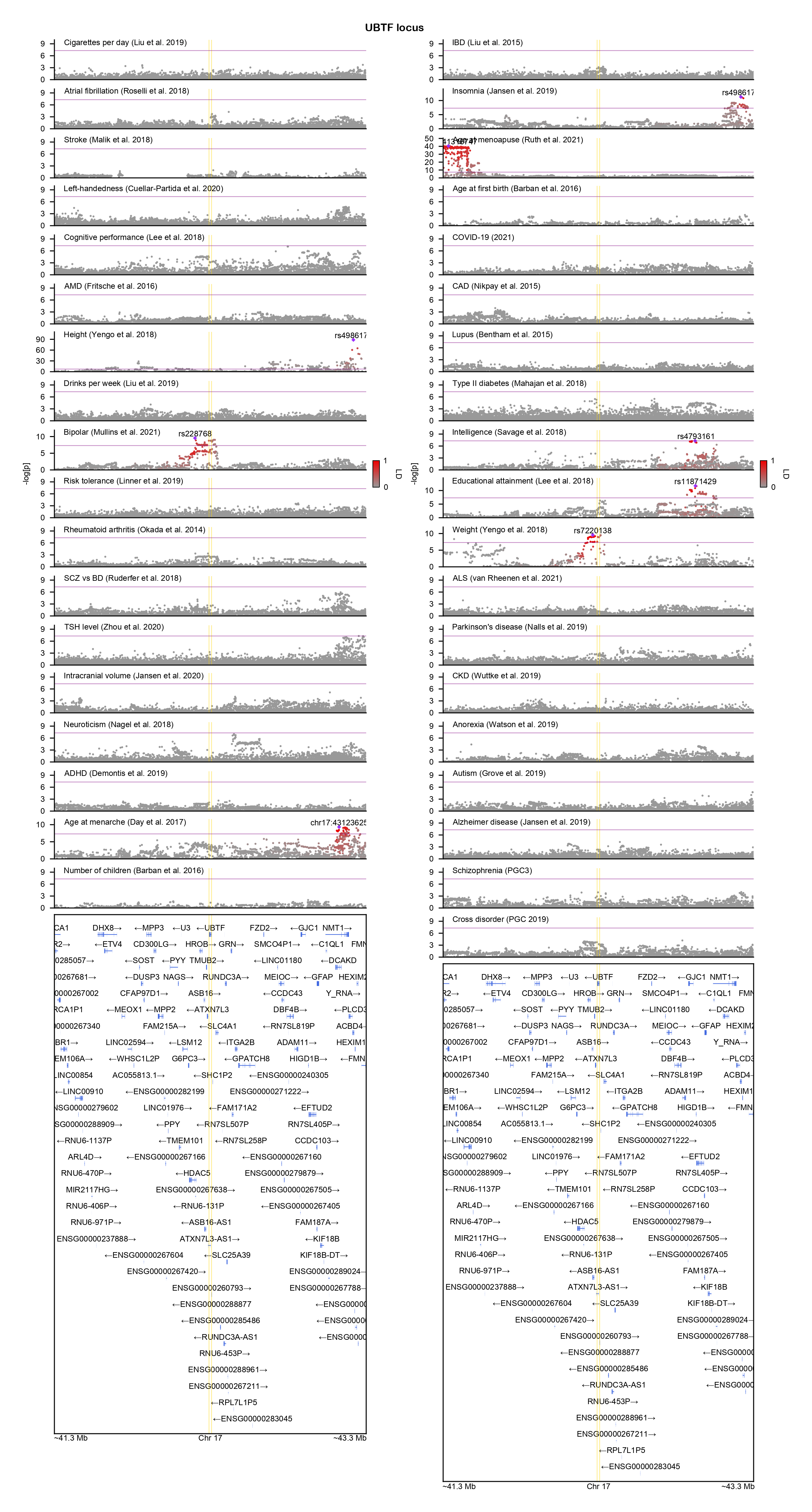

### USP9X-locuszoom.png

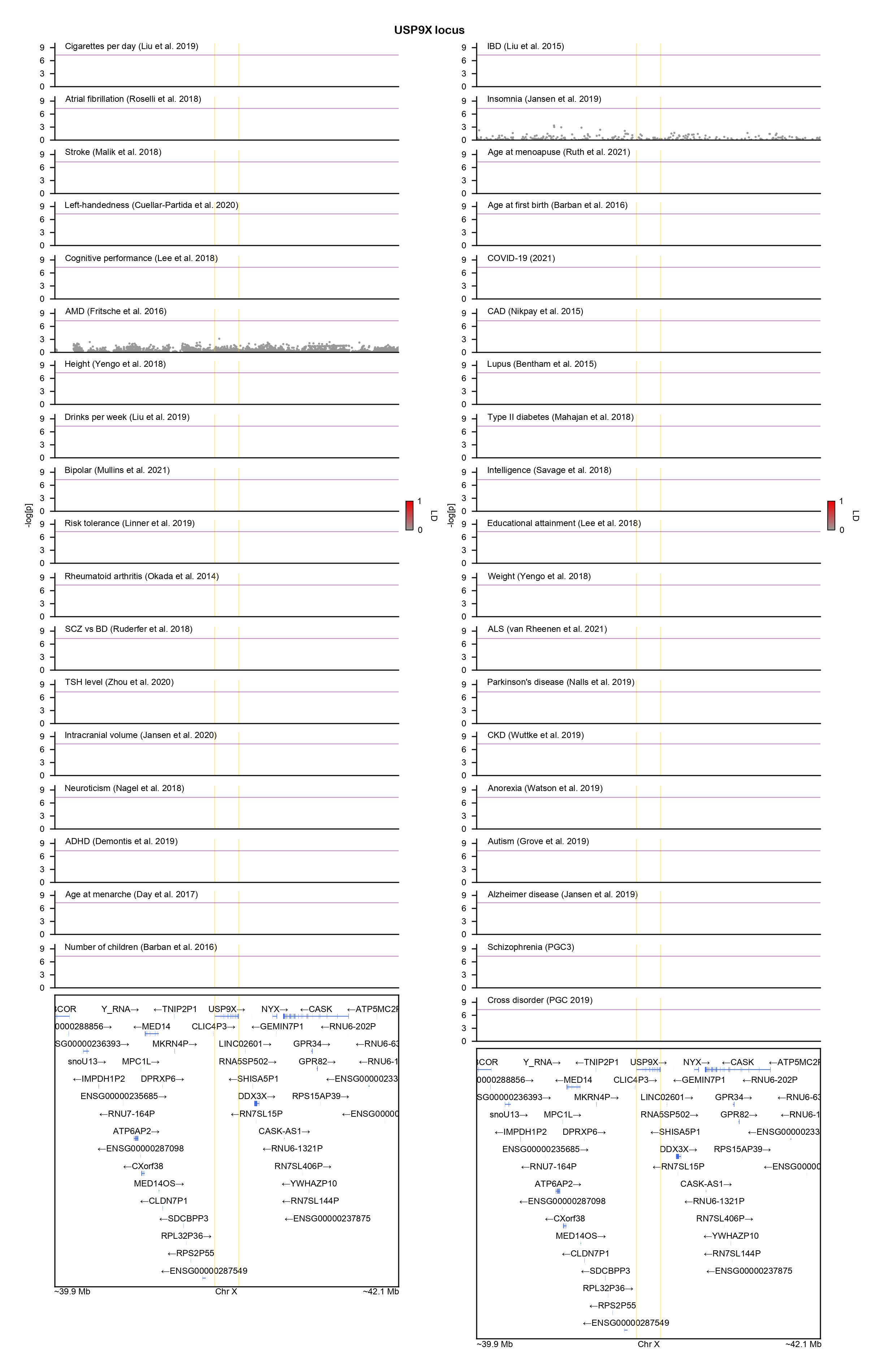

### VAMP2-locuszoom.png

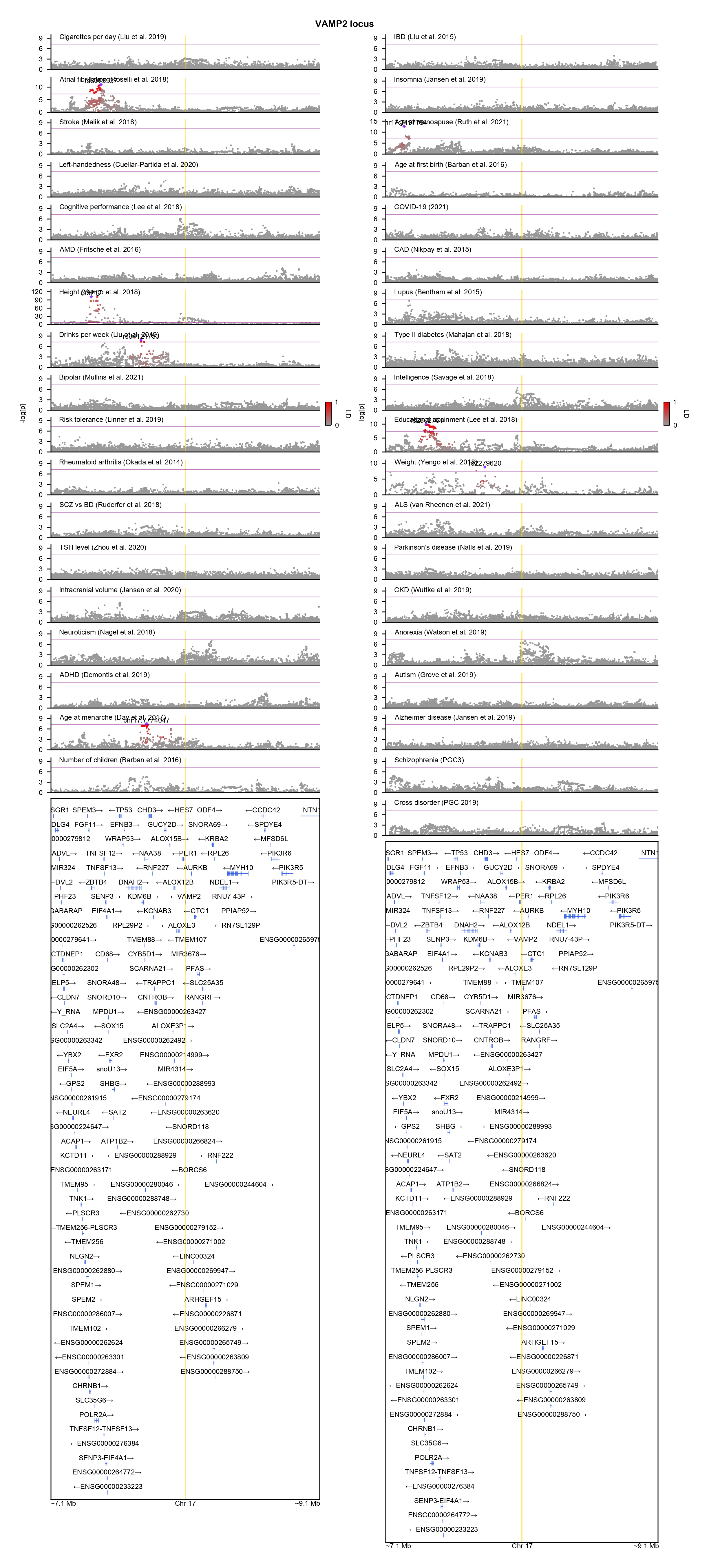

### WDR26-locuszoom.png

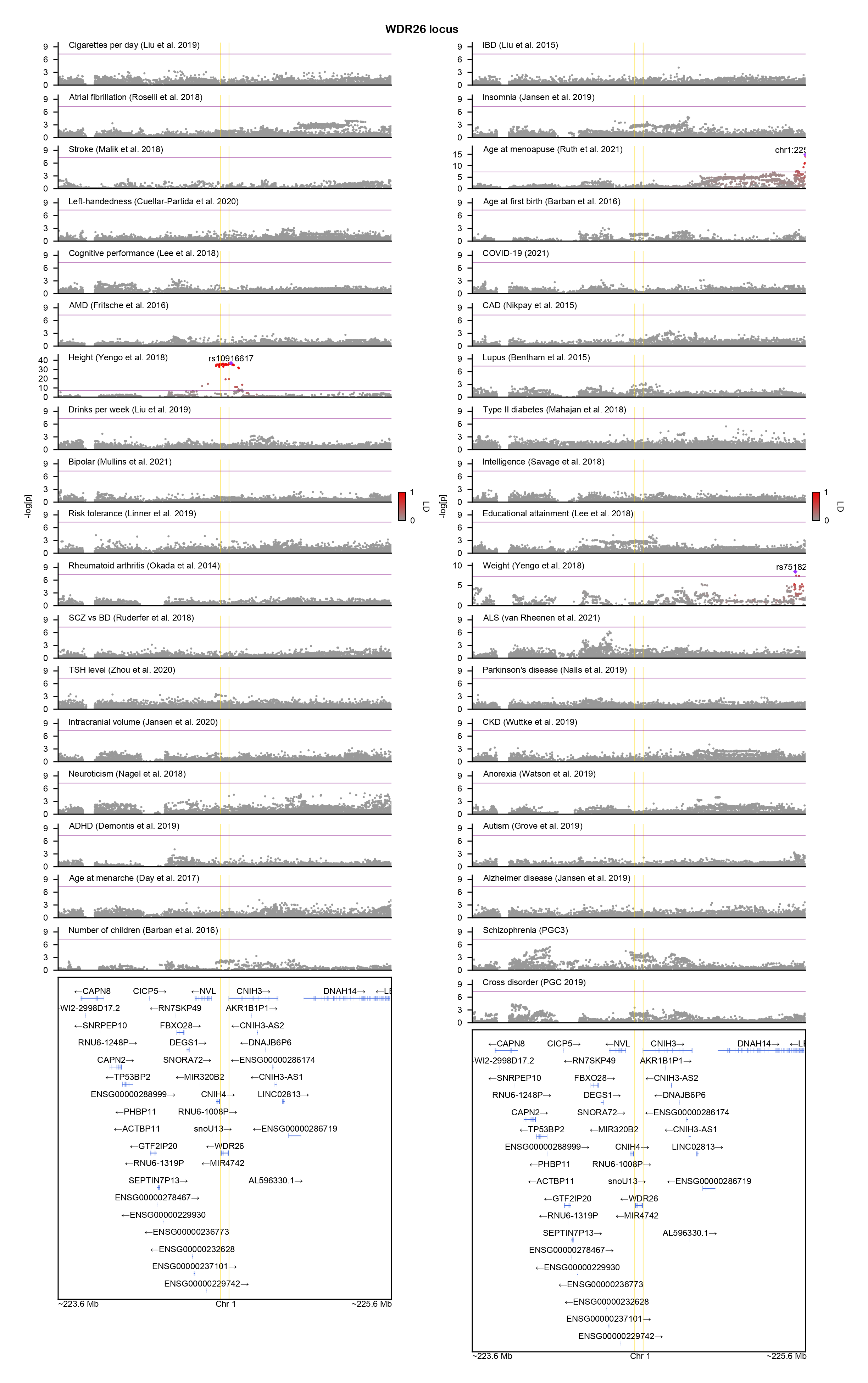

### WDR37-locuszoom.png

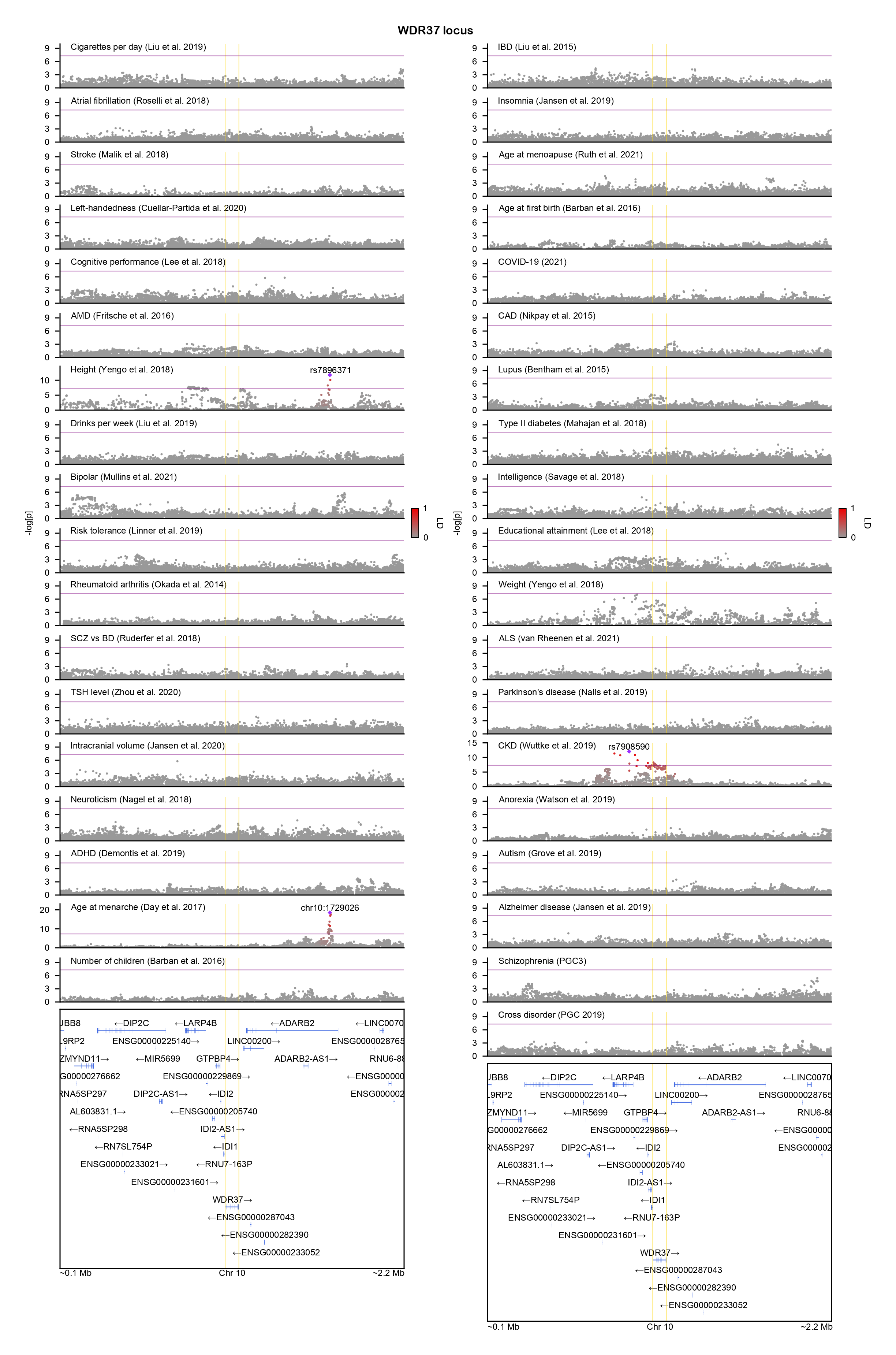

### ZBTB18-locuszoom.png

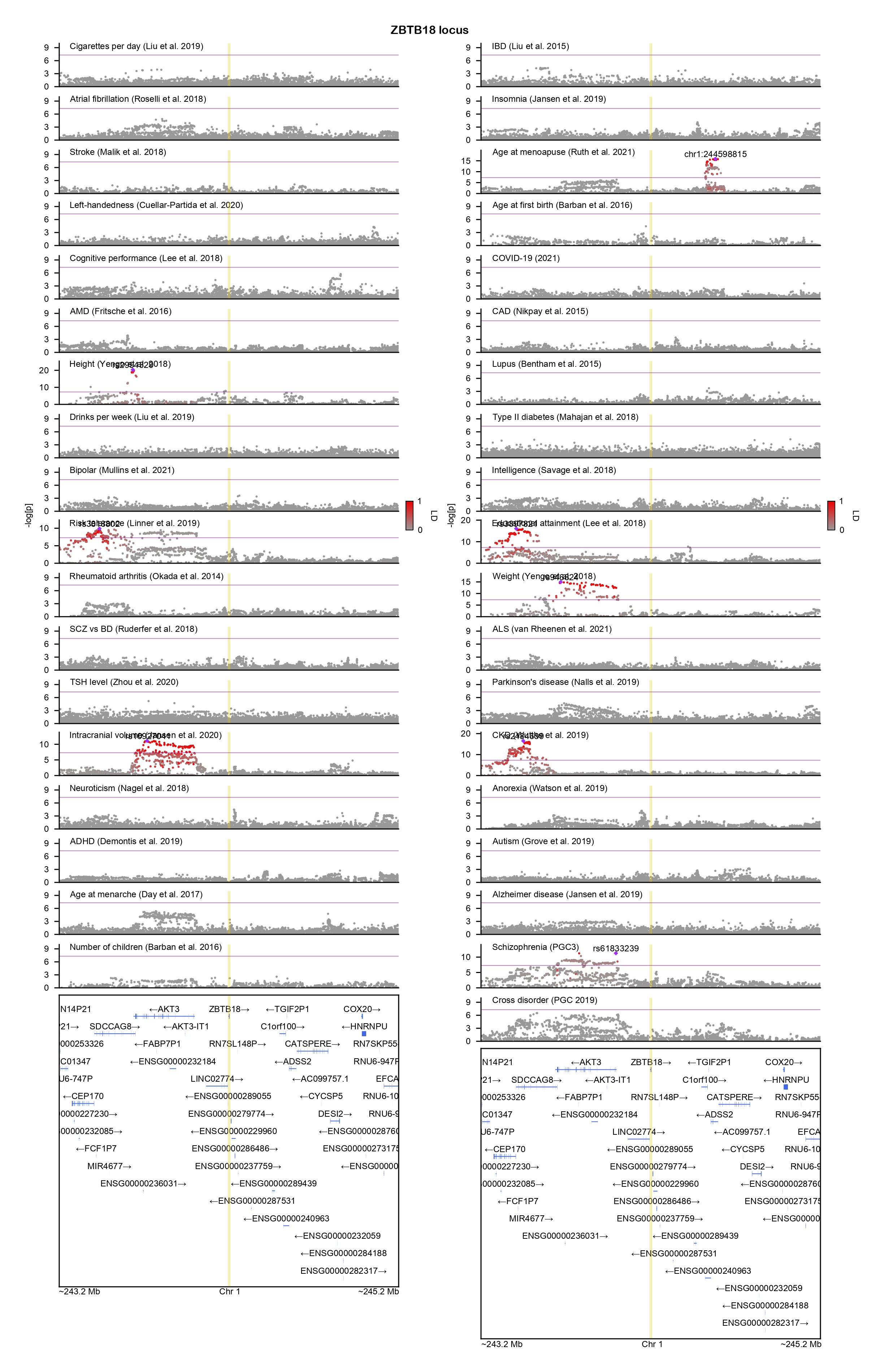

### ZBTB20-locuszoom.png

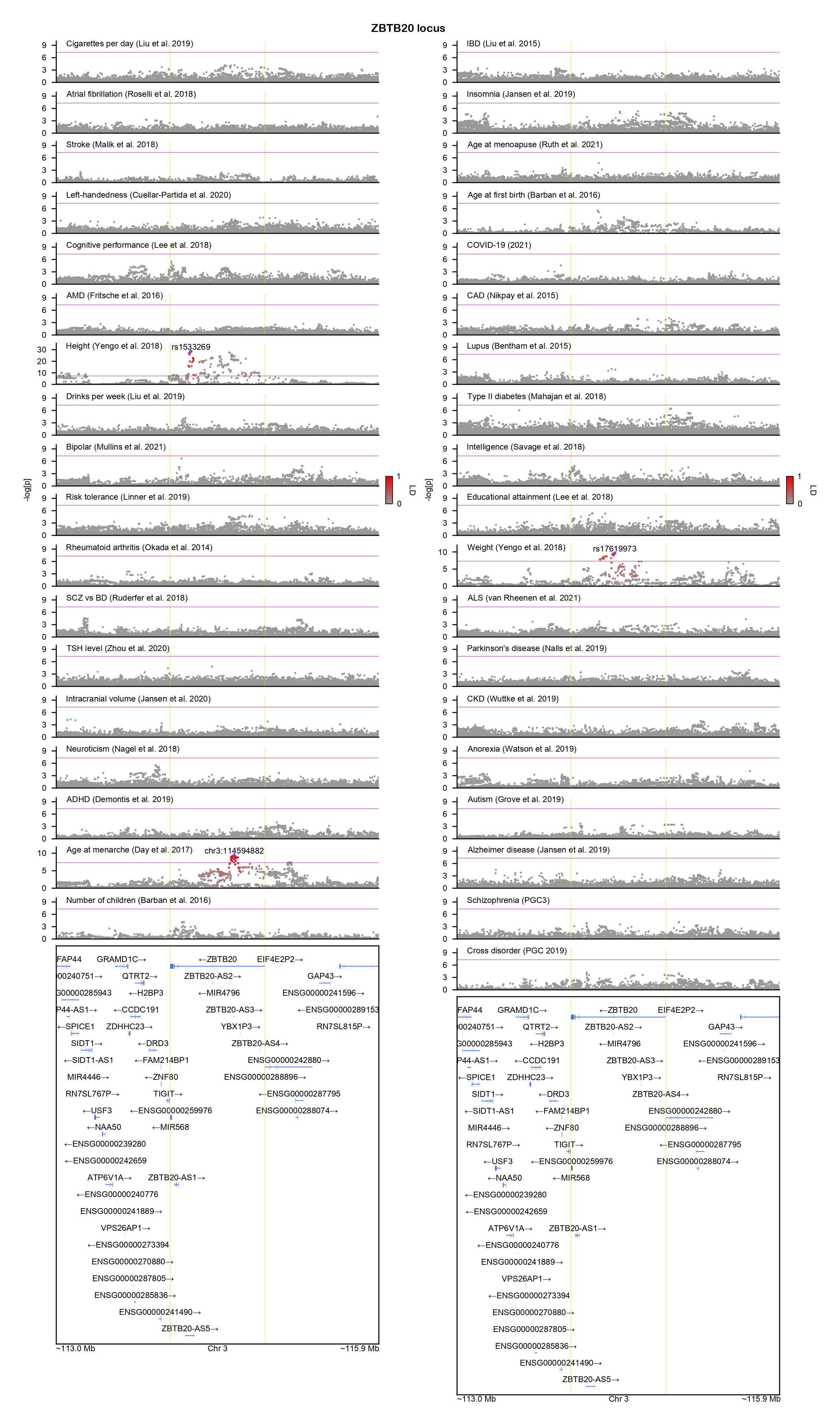

### ZMYND11-locuszoom.png

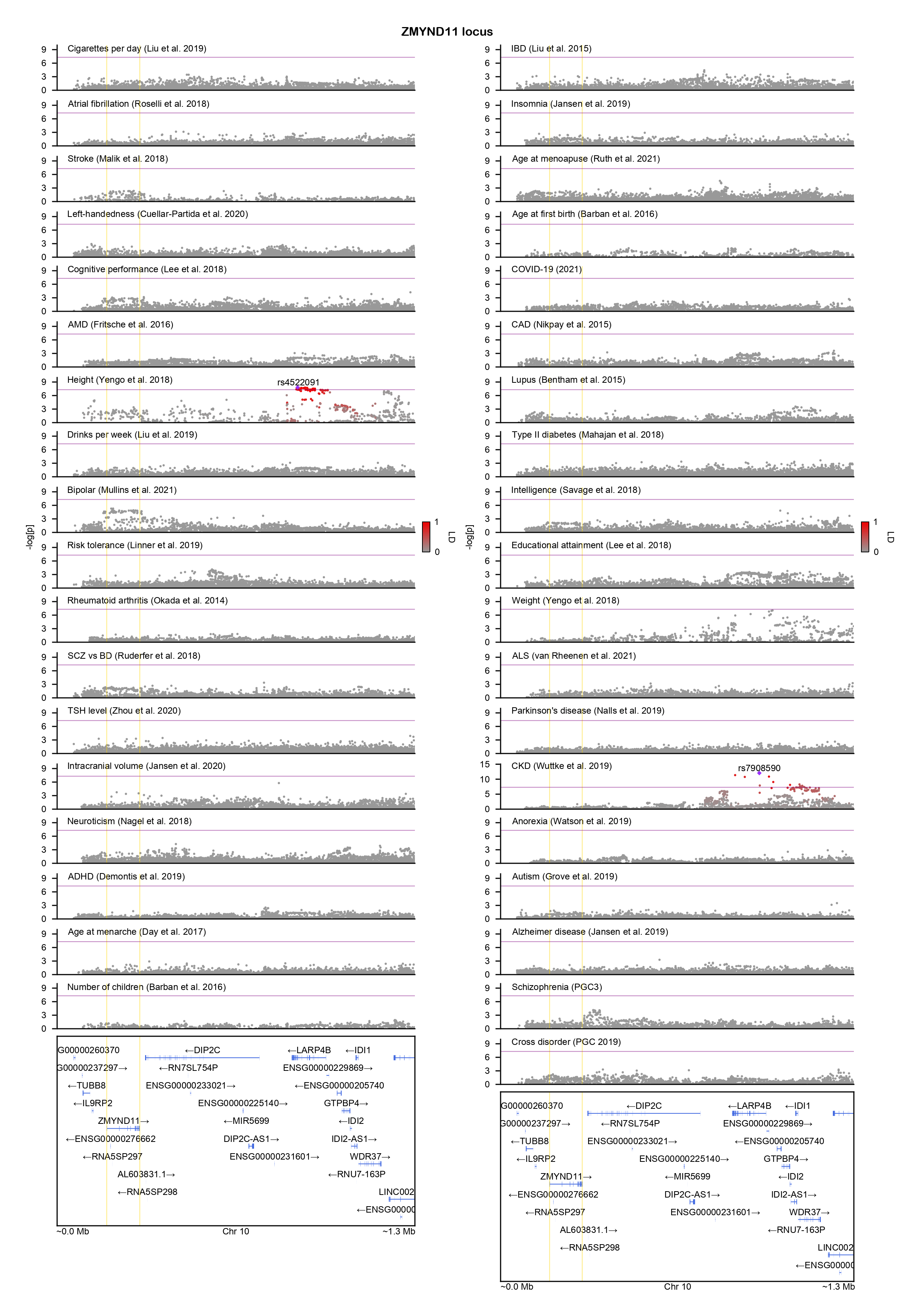

### ZNF292-locuszoom.png

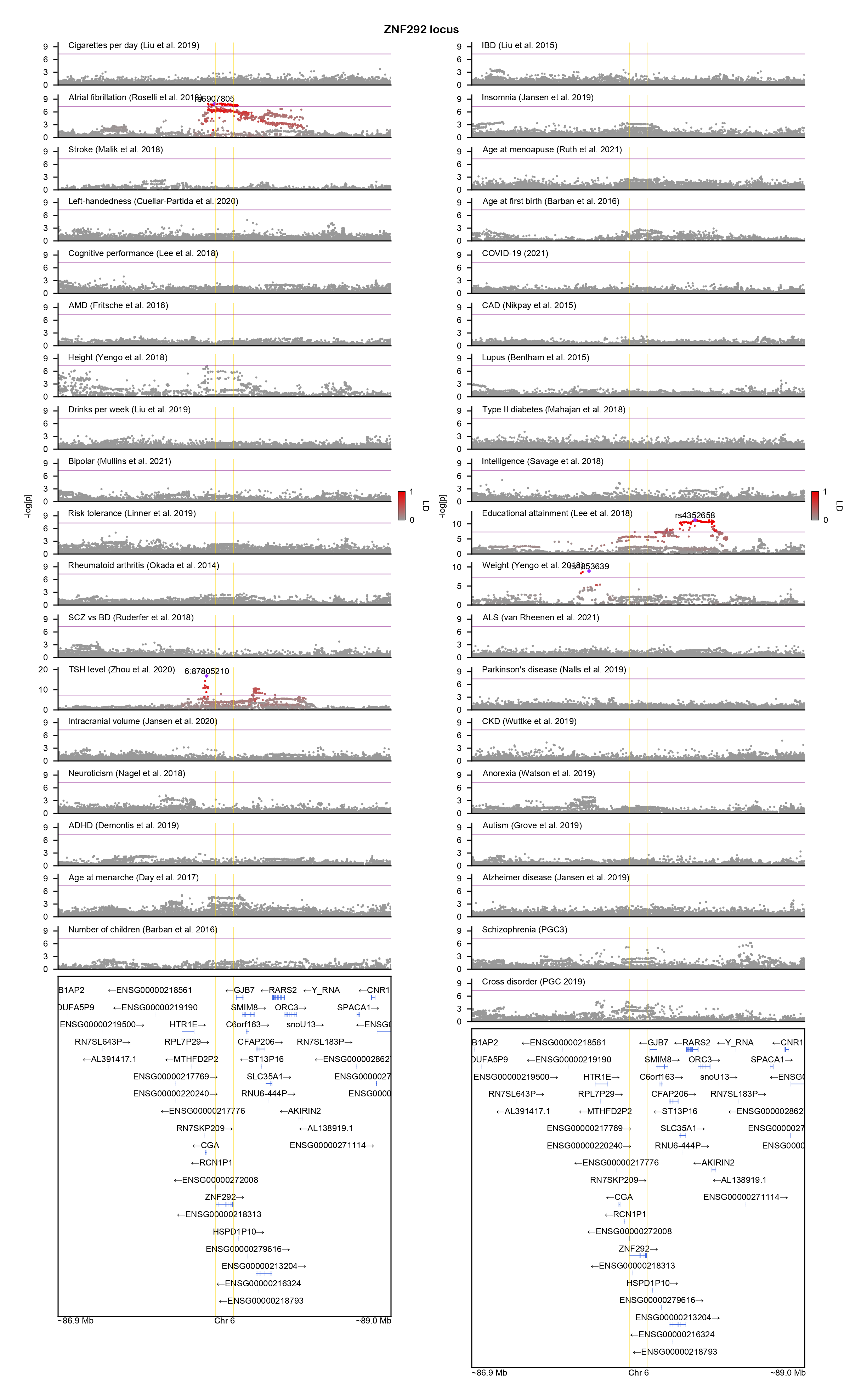
